# A multi-species co-occurrence index to avoid type II errors in null model testing

**DOI:** 10.1101/2021.11.03.467033

**Authors:** Vitalis K. Lagat, Guillaume Latombe, Cang Hui

## Abstract

Community structure is determined by the interplay among different processes, including biotic interactions, abiotic filtering and dispersal. Their effects can be detected by comparing observed patterns of co-occurrence between different species (e.g. C-score and the natural metric) to patterns generated by null models based on permutations of species-by-site matrices under constraints on row or column sums. These comparisons enable us to detect significant signals of species association or dissociation, from which the type of biotic interactions between species (e.g. facilitative or antagonistic) can be inferred. Commonly used patterns are based on the levels of co-occurrence between randomly paired species. The level of co-occurrence for three or more species is rarely considered, ignoring the potential existence of functional guilds or motifs composed of multiple species within the community. Null model tests that do not consider multi-species co-occurrence could therefore generate false negatives (Type II error) in detecting non-random forces at play that would only be apparent for such guilds. Here, we propose a multi-species co-occurrence index (hereafter, joint occupancy) that measures the number of sites jointly occupied by multiple species simultaneously, of which the pairwise metric of co-occurrence is a special case. Using this joint occupancy index along with standard permutation algorithms for null model testing, we illustrate nine archetypes of multi-species co-occurrence and explore how frequent they are in the seminal database of 289 species-by-site community matrices published by Atmar and Patterson in 1995. We show that null model testing using pairwise co-occurrence metrics could indeed lead to severe Type II errors in one specific archetype, accounting for 2.4% of the tested community matrices.

## 1. Introduction

A long-standing objective of ecology has been to infer the underlying community assembly processes that generate patterns of species coexistence in natural communities. These processes include environmental filtering, within/cross-guild biotic interactions, dispersal and disturbance (Latombe et al., 2021). One of the most commonly discussed patterns is species co-occurrence, which depicts the association and dissociation of species pairs in communities or in multiple sites/samples of a community (Camarota et al., 2016; Cordero and Jackson, 2019; D’Amen et al., 2018; Kohli et al., 2018). The level of non-random co-occurrence between species pairs has been interpreted as the result of two scale-dependent community assembly processes. First, a full spectrum of biotic interactions, from interference to facilitation, can strongly affect the level of co-occurrence at local- and mesoscales (e.g. one species can locally expel others with a similar niche requirement via limiting-similarity competition, or spatially synchronize its occurrence with other facilitative species; Hess et al., 2020; Meszéna et al., 2006; Schamp and Jensen, 2019). Second, environmental filtering can select a subset of species (typically within a functional guild) in heterogeneous environments and this process is thus considered as a major driver of species co-occurrence at meso-to broad-scales (Cadotte and Tucker, 2017). Two species can therefore frequently co-occur at broad spatial scales due to similar habitat requirements but locally exclude each other through competition. Drift and stochasticity can further interact with these scale-dependent processes to generate random patterns of co-occurrence (Hou et al., 2020; Måren et al., 2018), complicating our ability to discern community assembly processes and species coexistence mechanisms.

Testing for significant levels of species co-occurrence dates back to earlier debate on the role of interspecific competition in structuring ecological communities (Cordero and Jackson, 2019). In his analyses of the New Guinean avifauna, Diamond (1975) detected a checkerboard pattern of co-occurrence and subsequently concluded on the role of competition in driving species coexistence. Connor and Simberloff (1979, 1983) contested this conclusion by demonstrating similar co-occurrence patterns from null models with random reshuffling of species. This has led to a heated debate on the role of competition in community assembly and the importance of null model testing in statistical inference (especially in ecology). Despite such controversies, the use of null model testing to detect significant co-occurrence has become a central piece in many ecological theories and concepts (Sanderson, 2000; Veech, 2014). Resource differentiation, community assembly, ecological character displacement, the meta-community concept, equilibrium versus non-equilibrium organization of assemblages, and the local-regional species diversity relationship, are all notable examples of such theories (Davies *et al.,* 2007; Hunt *et al.,* 2008; Zelezniak *et al.,* 2015).

Co-occurrence has traditionally been measured and tested by considering only two species at a time, i.e. by assessing if the observed frequency of co-occurrence of all possible species pairs is higher or lower than expected by chance (e.g., Veech, 2013, 2014). Null model expectations are determined by shuffling an occupancy community matrix of species presence and absence over multiple sites/samples using a number of permutation algorithms. These null models therefore assume that levels of species co-occurrence only result from species relative occupancy (Gotelli, 2000; Gotelli and Sounding, 2001; Lehsten and Harmand, 2006). Null model tests normally rely on indices to summarize the co-occurrence patterns between species pairs in the community matrix [e.g., Stone and Roberts’ (1990) C-score or Sanderson’s (2004) natural co-occurrence metric] (Gotelli, 2000). Since a community matrix typically involves more than two species, values of pairwise metrics over all pairs of species are routinely averaged in the test. While these pairwise metrics have been instrumental in determining the drivers of ecological community structures, they are unable to discern any interactions for more than two species simultaneously (e.g. multiple species competing for the same resources or interacting with each other; Yang and Hui, 2021).

Null model testing using pairwise species co-occurrence metrics are constantly facing the challenge of conflating multiple contending community assembly processes (Cordero and Jackson, 2019). In particular, by averaging pairwise co-occurrence metrics, strong negative and positive associations between different pairs of species can cancel each other out, making it difficult to identify any significant non-random patterns, consequently often incurring type II statistical error. For example, the test for diffuse competition requires, intrinsically, the analysis of co-occurrence among multiple species and not just pairs of species within a community (Veech, 2014). Averaging pairwise co-occurrence indices does not capture information on the simultaneous co-occurrence.

Here, we present a multi-species co-occurrence index (‘joint occupancy’) that quantifies the occupancy of two or more species simultaneously in a community. The proposed metric takes the same presence-absence community matrix commonly used as inputs but captures co-occurrence patterns from two to n > 2 species (therefore generating n-1 values). We further derive the mean and variance of the proposed joint occupancy index to facilitate testing for statistical significance in observed co-occurrence patterns. Using this metric along with Gotelli’s (2000) recommended `sim2` permutation algorithm, we present nine possible archetypes of multi-species co-occurrence patterns and show how six of them can be observed in a dataset of 289 published species-by-site community matrices compiled by Atmar and Patterson (1995). We show that pairwise co-occurrence metrics could lead to severe type II (false negative) statistical errors during the null model test for matrices characterized by one of the six obtained archetypes. As such, indices of co-occurrence considering multiple species simultaneously, such as our multi-species joint occupancy index, should be preferred over pairwise metrics in future null model testing.

## 2. Materials and Methods

### 2.1. Null model testing

Null models have been developed to provide a reference point against which alternative hypotheses or inferences should be contrasted to detect non-random structures, and to infer causal relationships between ecological patterns and different community assembly processes. Null models have been especially instrumental in biogeography and ecology (Mora et al., 2019). Despite historical controversies about their utility (D’Amen et al., 2018), null models based on permutation algorithms have continued dominating the ecology literature and unearthing the solutions to many ecological problems that cannot be tested by conventional statistics.

At the mesoscale and higher, multiple candidate processes for structuring ecological communities can only be inferred from observed community patterns, including codistribution patterns, whereas processes at smaller scales can rely on experiments. Null models are crucial to discern non-random patterns from those random ones under certain constraints so that specific processes at play can be inferred. Using null models to detect non-random patterns in ecological communities requires a permutation algorithm that produces a sampling distribution for a particular metric, in the absence of a mechanism of interest, so that the observed patterns can be compared with those from the sampling distribution of the null model. Excluding a candidate mechanism in null model testing requires imposing specific constraints on the permutation algorithm, so that the ‘random’ patterns produced from the null model can be attributed entirely to the omission of the mechanism.

Nine basic permutation algorithms have been proposed (Gotelli, 2000). They range from no constraint at all (`sim1`) to those that are highly constrained on the row and column sums of the community matrix (e.g. `sim9`) in generating a sampling distribution. Two algorithms were recommended that can arguably avoid type II errors for co-occurrence patterns: `sim2` (which fixes the total occupancy of each species) and `sim9` (which fixes both the total occupancy of each species and the richness of each site). Algorithm `sim2` was found to produce congruent results when used with four co-occurrence indices (C-score, CHECKER, COMBO, and *V* ratio, respectively), while `sim9` worked best with C-score.

We corroborate previous and recent findings (Gotelli, 2000; Korňan et al., 2019; Maestre et al., 2008) on the use of `sim2` permutation algorithm on the premise that total number of species in an ecological community (which may or may not be influenced by their biotic interactions) determine the co-occurrence structure of ecological communities (Alexander et al., 2018). This means that if we were to fully attribute the co-occurrence structure of an ecological community to the effects of within-community biotic interactions, species must be allowed to move freely among sites but the total number of sites occupied by each species must be held constant during the permutation test. Rejecting the null hypothesis could signal either the role of biotic interactions or the effect of propagule fluxes in and out an open ecological community; consequently, a conclusion on the significant role of biotic interactions could face a type I error of false positive. In most cases, however, propagule fluxes serve to nullify the community structure imposed by biotic interactions, thus failing the detection on the role of biotic interactions in structuralizing co-occurrence, a type II error of false negative.

Because permutation algorithms that modify species occupancy while keep within-site richness constant (e.g., `sim3` and `sim9`) do not alter the expected number of sites occupied by different species, they are not suitable for testing mechanisms of co-occurrence. In an *i* by *n* species-by-site community matrix, the probability of finding any given number of species to co-occur in a particular site remains exactly the same for such vertical permutation algorithms that retain within-site richness. In other words, site-specific richness is not a reasonable constraint in null model testing for significant co-occurrence patterns. This, however, does not suggest that co-occurrence patterns are independent from the variation of alpha richness between sites. For instance, the level of resources, together with biotic interactions, can determine the number of species that a site can hold (i.e., alpha diversity), which may in turn influence species co-occurrence (see Appendix S2 for an examination of the relationship between co-occurrence and richness variation).

### 2.2. Pairwise co-occurrence metrics

One of the widely used species co-occurrence metrics is the *natural metric* proposed by Sanderson (2000, 2004). This metric simply counts the number of sites occupied by two species simultaneously and averages over all pairs of species. The observed value of the natural metric can then be compared to the sampling distribution obtained from randomizing the species-by-site matrix. Typically, if the observed value lies below the null model expectations (i.e., below the lower critical value of the confidence interval), the significance of negative species associations (i.e., segregations or dissociations) is confirmed, which normally implies the role of inter-specific competition (Cordero and Jackson, 2019) or other antagonistic interactions. The significance of positive species associations (or aggregation) can be inferred if the observed value lies above the null model expectations (i.e., above the upper critical value of the confidence interval), which has traditionally been used to imply the working of environmental filtering (Cadotte and Tucker, 2017) but can also signal facilitation or mutualism.

Many other pairwise co-occurrence metrics have been proposed; for instance, the number of unique species combinations (Pielou and Pielou, 1968), the number of species pairs forming perfect checkerboards (Diamond, 1975) and the checkerboard score (Stone and Roberts, 1990). However, these pairwise metrics can only detect patterns of co-occurrence between two species in community assemblages, but not patterns of three or more species occurring simultaneously in the same communities. As a result, they often suffer non-detection of non-stochastic processes (type II error).

### 2.3. Multi-species joint occupancy index

To overcome the limitations of pairwise metrics for detecting non-random higher-order cooccurrence patterns in structured community assemblages, we here propose the *multi-species joint occupancy index.* This index describes the average number of sites jointly occupied by a given number of species. To illustrate the limitations of pairwise metrics, let us consider the following two hypothetical cases. Three species (A, B and C) are distributed across seven sites [case (a) and (b) in Figure 1], with species A occurring in 4 sites, species B in 3 sites and species C in 5 sites. The joint occupancies (*J*) for all pairs of species: *J*^{A,B}^, *J*^{A,C}^ and *j*^{B,C}^ are 1, 3 and 2 respectively, in both cases. However, the joint occupancy of all three species is *J*^{A,B,C}^ = 1 in case (a) but 0 in case (b). This means by considering all possible orders of species combinations, the multi-species joint occupancy (defined below) can discern different patterns of species co-distributions, which may not be discernible by pairwise co-occurrence metrics. Consequently, all pairwise metrics either do not detect change in species distribution patterns [e.g. Pielou and Pielou’s (1968) metric] or they only consider two species at a time for its computation [for the case of Diamond’s (1975), Schluter’s (1984), and Stone and Roberts’ (1990) indices]. We therefore need to go beyond pairwise metrics to detect higher-order co-occurrence patterns and associated complex interaction structures (Dieckmann et al., 2000; Mayfield and Stouffer, 2017).

**Figure 1.**
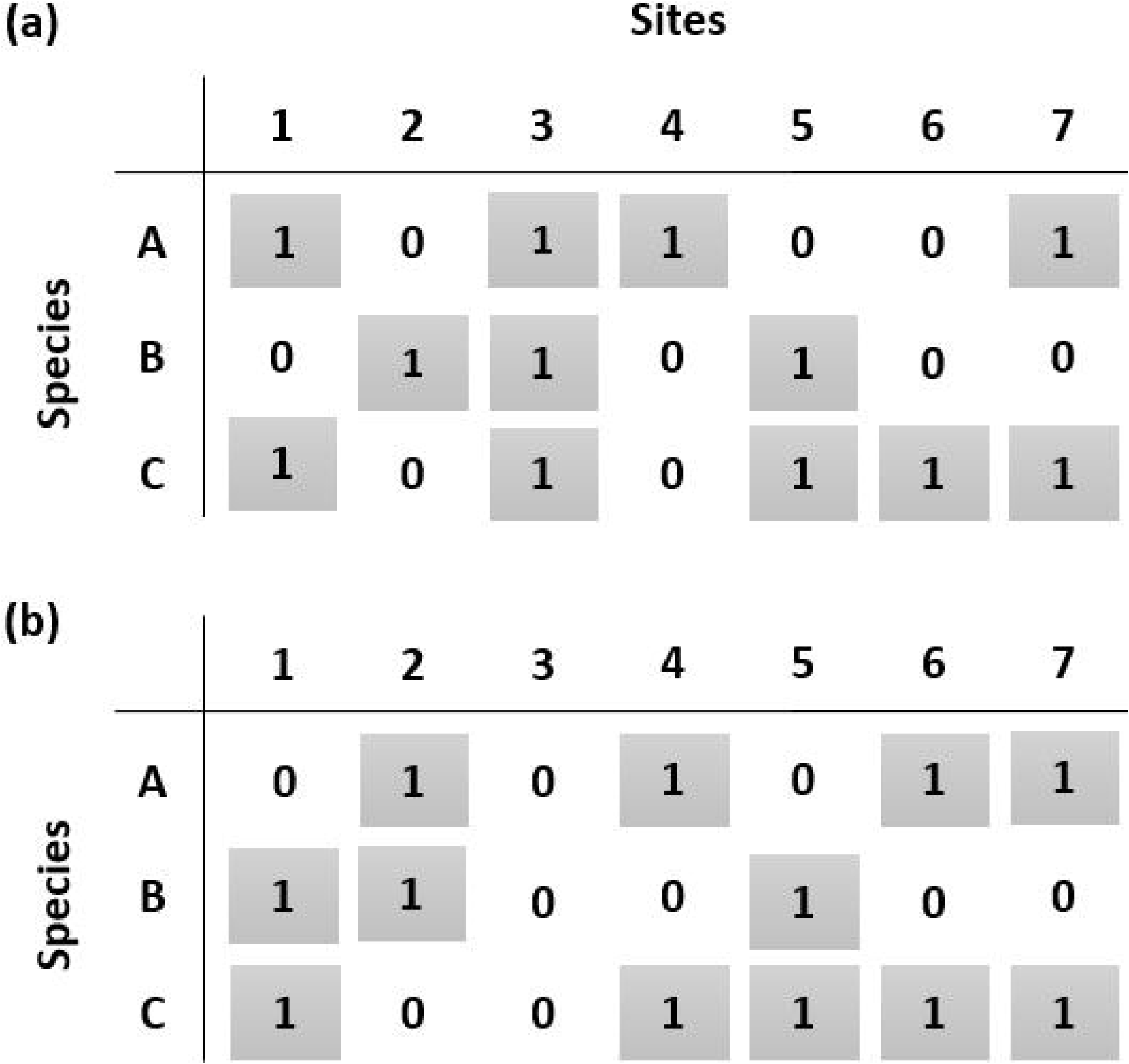
For the two communities (a) and (b), the values of the pairwise index (*i* = 2): *J*^{*A,B*}^, *J*^{*A,C*}^ and *J*^{*B,C*}^ are 1, 3 and 2, respectively. However, when the higher order of joint occupancy (*J*^{*A,B,C*}^; *i* = 3) is used, we get one for community (a) and zero for community (b). This means irrespective of the two different patterns in the two communities, the pairwise metric does not draw a distinction between them, unlike the higher order metric. It suffices then to say multi-species joint occupancy detects change in species distribution patterns across different sites while pairwise co-occurrence does not.

Given a species-by-site presence-absence matrix with *m* rows (species) and *n* columns (sites), we can count the number of sites occupied by two or more specific species simultaneously, which can be easily achieved using the concept of dot product in vector calculus. For instance, to compute the number of sites occupied by *i* out of a total of *m* species simultaneously, we can characterise every species (or row) by an occupancy vector, e.g. species A in Figure 1(a) has *V*_1_ = (1,0,1,1,0,0,1), and compute the dot products of all the *i* vectors. Summing the elements of the resulting vector gives the joint occupancy of *i* species. Specifically, let *V_i_* = (*v*_*i*,1_ *v*_*i*,2_, *v*_*i*,3_,..., *v_i,n_*) be the occupancy vector of species *i* among *n* sites. For a specific set of *i* species, denoted as {*i*}, we can assess whether or not this set of species occurs simultaneously in a specific site *j* by examining whether 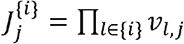 is equal to 1 or 0. Joint occupancy, *J*^{*i*}^ the number of sites occupied by this set of *i* species simultaneously, can be given by

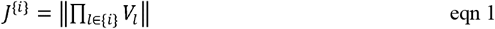

where ||·|| is the sum of all elements of the vector produced from the dot products of the *i* occupancy vectors.

To compute the joint occupancy of unspecified sets of species in the assemblage, we need to estimate the expected value of joint occupancy. Let *S_j_* be the number of species at site *j* and 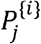 be the probability of *i* species simultaneously occurring in site *j*; we have

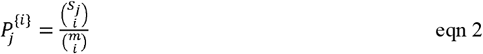

where 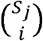 is the number of ways of selecting *i* species out of a total of *S_j_* species, and 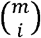 is the number of ways of selecting *i* species from the entire species assemblage (*m*). If 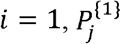 represents the probability of a particular species occurring in site *j*. If 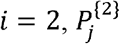 represents the probability of two species co-occurring in site *j*. If 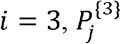, represents the probability of 3 species simultaneously occurring in site *j*, and so forth until *i* = *S_j_*, when 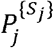 represents the probability of site *j* harboring *S_j_* species. The probability 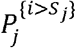 of site *j* harboring any number of species *i > S_j_* simultaneously is zero. Note that the probability 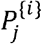 of *i* species occurring simultaneously in site *j* is also the expected value of joint occupancy of *i* species at site *j*.

Using the linearity property of expectation, and assuming the sites were sampled independently, we can conclude that the average number of simultaneous occurrences of *i* species (or the expected value of joint occupancy) in the entire assemblage is given by

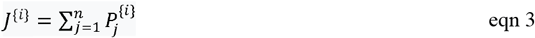

The number of species as superscript is referred to as the order of joint occupancy. Given the expected value of joint occupancy, we could calculate the variance of joint occupancy by summing up the covariance of occurrences between sites:

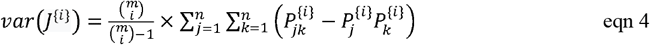

where 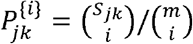 is the probability of *i* species jointly occurring in sites *j* and *k*, and *S_jk_* is the number of species occurring in both sites *j* and *k*.

Joint occupancy is not a single measure, but a set of measures that together provide insights on multi-species co-occurrence of different orders. *J*^{1}^ is the average number of sites occupied by a single species (i.e., average occupancy per species). *J*^{2}^ is the average number of sites occupied by two species simultaneously (i.e. the natural metric of co-occurrence). *J*^{3}^ is the average number of sites occupied by three species simultaneously, and so on. By considering the number of sites occupied by multiple species simultaneously, this index can assess the contribution of the spectrum from species-rich to species-poor sites, to multispecies joint occupancy, and overcomes limitations of pairwise co-occurrence indices to capture more precisely the non-stochastic drivers of community assembly.

### 2.4. Nine archetypes of multi-species co-occurrence

Statistical significance of pairwise co-occurrence metrics can be established by comparing its observed value to the sampling distribution of null expectations from the permutation of a focal species-by-site matrix. With the multi-species joint occupancy index, we can calculate the observed values and sampling distributions for a range of orders. Assuming incremental changes in the positions of observed joint occupancy values with respect to the sampling distributions of null expectations over the full range of orders, nine archetypes of cooccurrence can be defined (Figure 2). These archetypes range from those where all observed joint occupancy values lie above, within or below the null expectations for all orders (i.e., above the upper critical value of the confidence interval, within the upper and lower critical values, or below the lower critical value; archetypes A1, A5 and A9), to those where the positions of observed joint occupancy values differ from the null expectations only for a specific range of orders (Figure 2). Each archetype is representative of unique community assembly processes driving species co-occurrence (see Table 1 for interpretations and possible inferences for processes of all archetypes).

**Figure 2.**
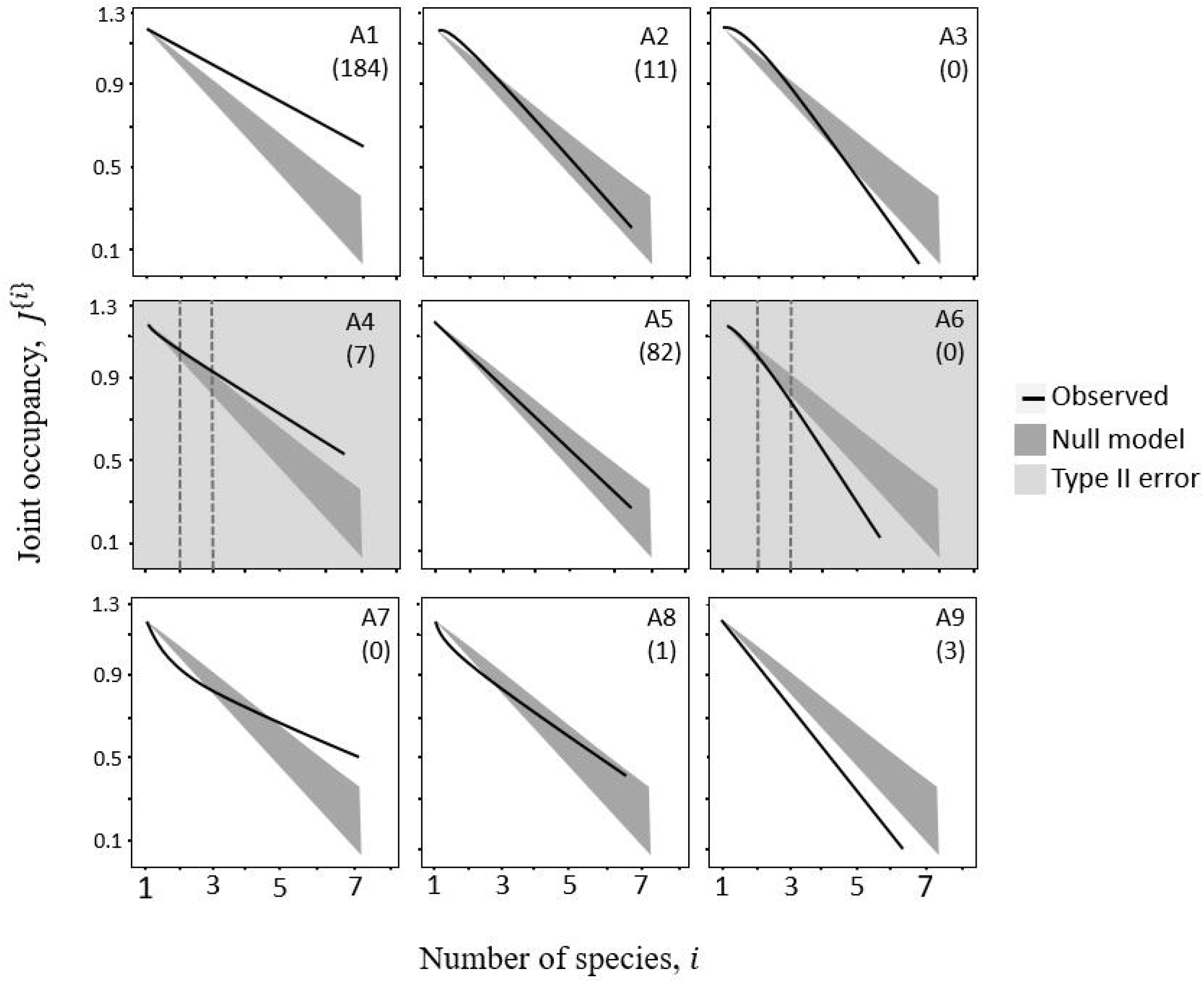
Nine possible archetypes of species co-occurrence patterns from the null model test of the same patterns. The number of communities under each archetype are given in brackets. The dark-gray ribbons represent the null model with its upper and lower boundaries being the critical values of the 95% closed confidence interval of the joint occupancy values (computed using eqn 3) of simulated data. Depending on the position of the black curves and lines (representing the joint occupancy values [from eqn 3] of observed data) relative to the null model, the test can be deemed statistically significant. As the number of species increase, the results and consequently, the inferences from the null model test, change in some archetypes. This is illustrated, for instance, by the gray dotted lines where two species considered simultaneously lead to different ecological interpretations as to what structures ecological communities compared to when three or more species are considered simultaneously.

**Table 1.**
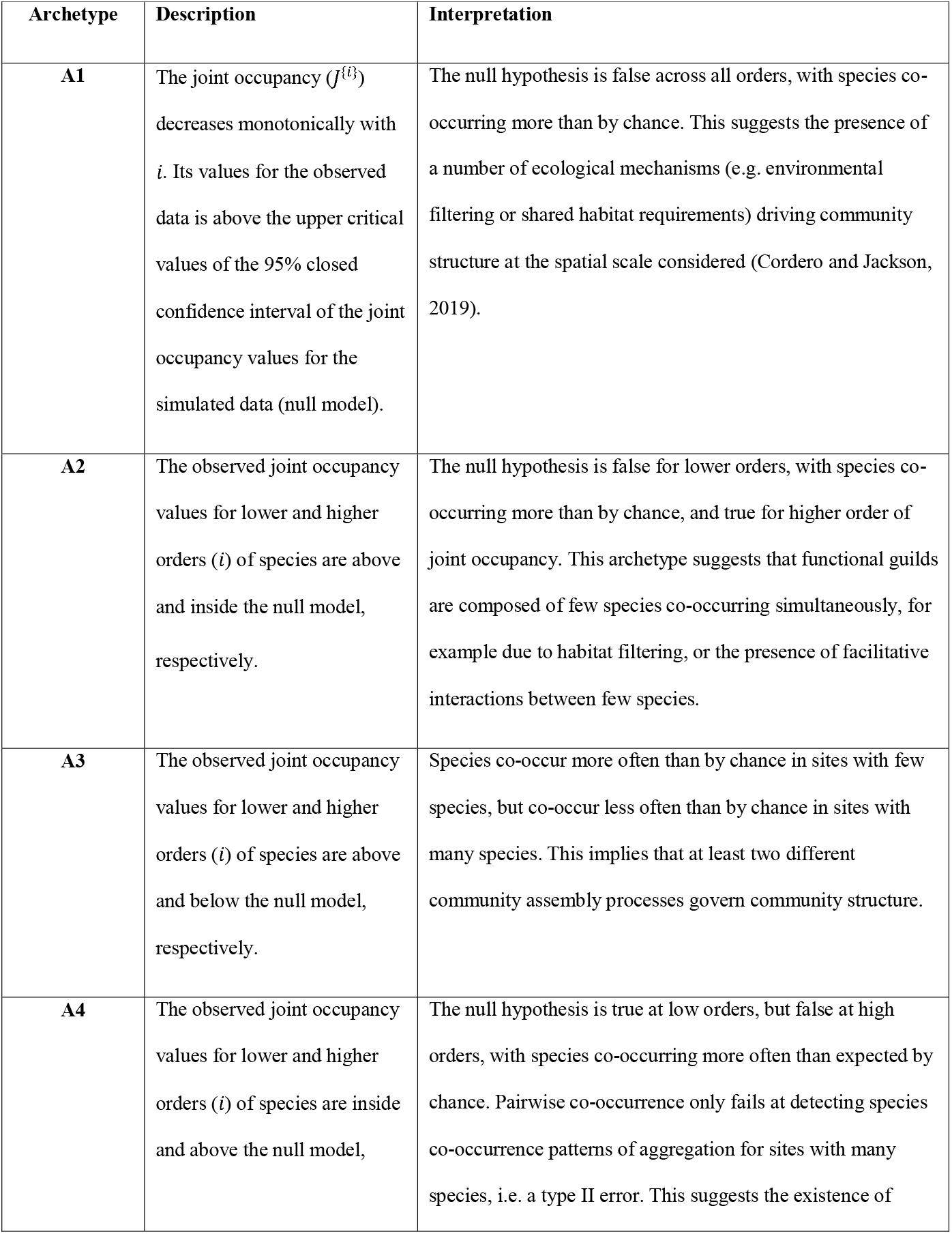
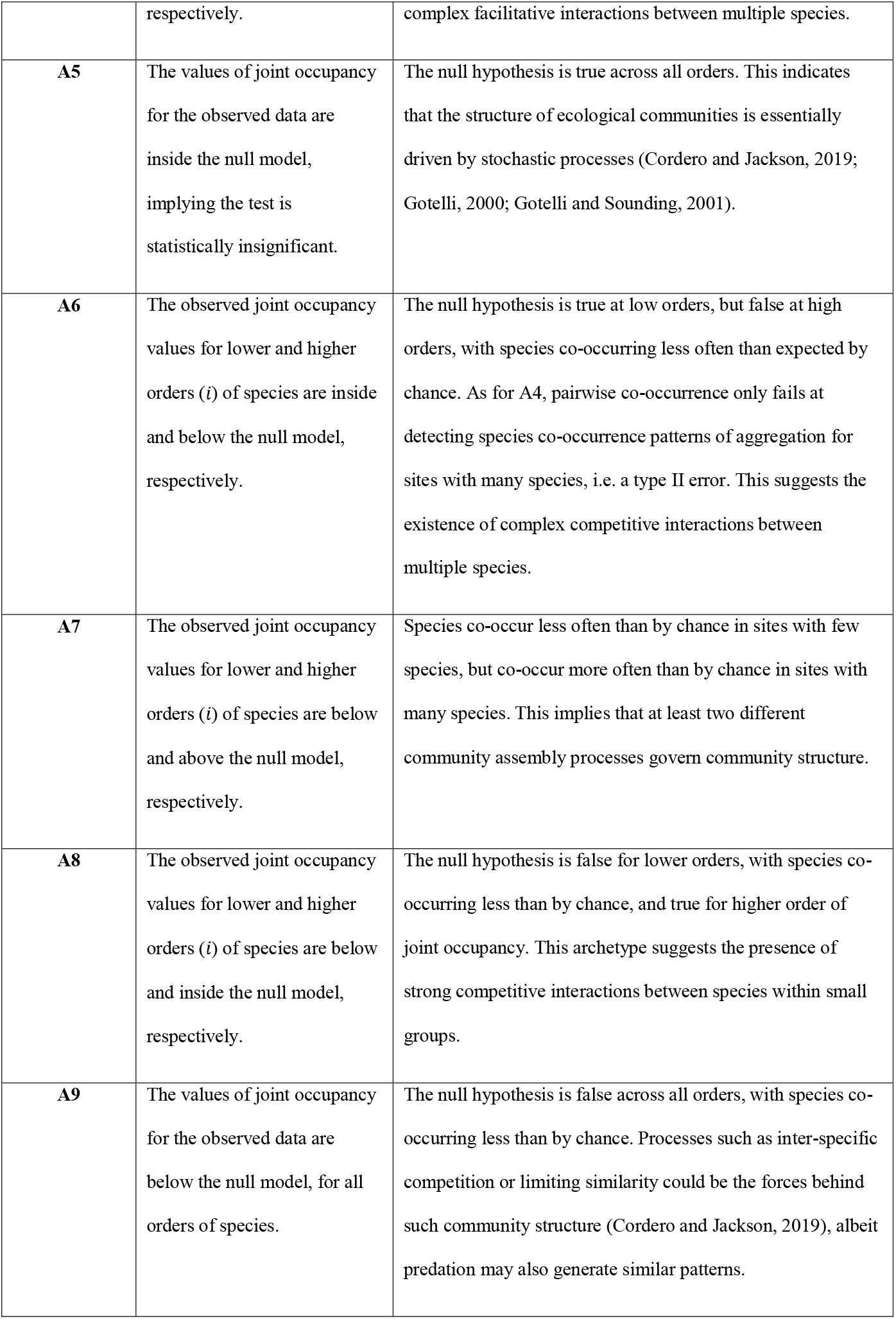
Archetypes of species co-occurrence patterns in ecological communities as brought out by a null model test using joint occupancy index and various permutation algorithms.

| Archetype | Description | Interpretation |
| --- | --- | --- |
| **A1** | The joint occupancy ($J^{(i)}$) decreases monotonically with $i$. Its values for the observed data is above the upper critical values of the 95% closed confidence interval of the joint occupancy values for the simulated data (null model). | The null hypothesis is false across all orders, with species co-occurring more than by chance. This suggests the presence of a number of ecological mechanisms (e.g. environmental filtering or shared habitat requirements) driving community structure at the spatial scale considered (Cordero and Jackson, 2019). |
| **A2** | The observed joint occupancy values for lower and higher orders ($i$) of species are above and inside the null model, respectively. | The null hypothesis is false for lower orders, with species co-occurring more than by chance, and true for higher order of joint occupancy. This archetype suggests that functional guilds are composed of few species co-occurring simultaneously, for example due to habitat filtering, or the presence of facilitative interactions between few species. |
| **A3** | The observed joint occupancy values for lower and higher orders ($i$) of species are above and below the null model, respectively. | Species co-occur more often than by chance in sites with few species, but co-occur less often than by chance in sites with many species. This implies that at least two different community assembly processes govern community structure. |
| **A4** | The observed joint occupancy values for lower and higher orders ($i$) of species are inside and above the null model, | The null hypothesis is true at low orders, but false at high orders, with species co-occurring more often than expected by chance. Pairwise co-occurrence only fails at detecting species co-occurrence patterns of aggregation for sites with many species, i.e. a type II error. This suggests the existence of |
|  | respectively. | complex facilitative interactions between multiple species. |
| **A5** | The values of joint occupancy for the observed data are inside the null model, implying the test is statistically insignificant. | The null hypothesis is true across all orders. This indicates that the structure of ecological communities is essentially driven by stochastic processes (Cordero and Jackson, 2019; Gotelli, 2000; Gotelli and Sounding, 2001). |
| **A6** | The observed joint occupancy values for lower and higher orders (*i*) of species are inside and below the null model, respectively. | The null hypothesis is true at low orders, but false at high orders, with species co-occurring less often than expected by chance. As for A4, pairwise co-occurrence only fails at detecting species co-occurrence patterns of aggregation for sites with many species, i.e. a type II error. This suggests the existence of complex competitive interactions between multiple species. |
| **A7** | The observed joint occupancy values for lower and higher orders (*i*) of species are below and above the null model, respectively. | Species co-occur less often than by chance in sites with few species, but co-occur more often than by chance in sites with many species. This implies that at least two different community assembly processes govern community structure. |
| **A8** | The observed joint occupancy values for lower and higher orders (*i*) of species are below and inside the null model, respectively. | The null hypothesis is false for lower orders, with species co-occurring less than by chance, and true for higher order of joint occupancy. This archetype suggests the presence of strong competitive interactions between species within small groups. |
| **A9** | The values of joint occupancy for the observed data are below the null model, for all orders of species. | The null hypothesis is false across all orders, with species co-occurring less than by chance. Processes such as inter-specific competition or limiting similarity could be the forces behind such community structure (Cordero and Jackson, 2019), albeit predation may also generate similar patterns. |

For instance, in Figure 2 A4, the observed value of multi-species joint occupancy (dark line) appears inside the null distribution for the pairwise metric (i.e., when order *i* = 2). But when a higher order of species (e.g., *i* = 3) is used, the observed value curve bends upward to reach values above the null model distribution. This would indicate the presence of non-random community assembly processes for groups of multiple species, that would have been unnoticed using only a pairwise metric. Similar conclusions apply to archetypes A3, A6, and A7, which all reveal different processes at low and high orders of joint occupancy (Figure 2). Note that in theory, the positions of observed joint occupancy values with respect to null expectations could vary in more complex ways as the joint occupancy order increases (for example above null expectations for *J*^{1}^ within null expectations for *J*^{3}^ and above null expectations again for *J*^{4}^. Nonetheless, we expect this kind of complex configurations to be rare.

### 2.5. Community matrix and testing

With the proposed joint occupancy index and `sim2` permutation algorithm, we performed null model testing of multi-species co-occurrence patterns for 289 species-by-site presenceabsence matrices compiled by Atmar and Patterson (1995). The numbers of species in these communities range from 3 to 309 and the numbers of sites from 3 to 202, representing about 40 taxonomic groups. Note, it is obvious that the number of sites harboring multiple species simultaneously (thus joint occupancy, *J*^{*i*}^ decreases with its order (i.e. the number of species, *i*). To understand the forms of this decline, we fitted exponential, power law and exponential-power law parametric forms to order-dependent joint occupancy. Fitting these parametric models to how joint occupancy declines with orders offer additional insights to forces behind species co-occurrence patterns and the role of biotic interactions. Similar to the fact that the parametric form of zeta diversity decline of multisite compositional similarity can unveil stochastic versus deterministic assembly processes at work in ecological communities (Hui and McGeoch, 2014; McGeoch et al., 2019), we could anticipate different orders of biotic interactions, other than just pairwise interactions, that are responsible to cooccurrence at different orders (e.g. neutral force of random encountering versus niche-based force of functional trait matching/complementarity). As higher order joint occupancy can only occur in species rich sites, we further anticipate the entangled concepts of species turnover and co-occurrence.

Specifically, using the joint occupancy index, we calculated *J*^{*i*}^ for an observed matrix. Using `sim2` permutation algorithm (which reshuffles the observed matrix while keeping species occupancy constant), we calculated *J*^{*i*}^ for 999 randomized matrices. We used these 1000 values of /W as its sampling distribution, from which the 95% [closed] confidence interval were used for hypothesis testing. All the results presented in this paper were computed using `msco` R Package in R version 4.1.1 (see Appendix S1).

## 3. Results

Joint occupancy, *J*^{*i*}^, declines monotonically with the number of species *i*. When two parametric forms [exponential: *J*^{*i*}^ = *a* · exp(*b · i*); and power law: *J*^{*i*}^ = *a·i*^*b*^] were fitted to how joint occupancy declines with increasing number of species (hereafter, ‘joint occupancy decline’), exponential form fitted with *R*^2^ > 0.95 in 257 (88.9%) of the 289 community matrices and power law with the same level of the goodness-of-fit in 243 (84.1%) cases. However, power law had lower AIC values than exponential in 55 (19%) communities, while exponential had lower AIC values than power law in 234 (81%) communities. When the two models were combined, the resulting model [exponential-power law: *J*^{*i*}^ = *a* · exp(*b · i*) · *i^c^*] provided the best fit (*R*^2^ > 0.95 in 285 communities). Among the three models, exponential-power law provided the lowest AIC values in 273 communities, while exponential and power law had the lowest AIC values in 11 and 5 communities, respectively (see summary in Table 2 and full results in Appendix S1). This means the exponential-power law was the best model (among the three) in fitting joint occupancy decline in 94% of communities.

**Table 2.** The frequency of the six realized archetypes of species co-occurrence patterns and the parametric forms of joint occupancy decline for all communities under each archetype. The three parametric forms fit extremely well (most *R*^2^ > 0.95) to joint occupancy decline. However, to further discern the best form, we based on AIC to select only the best form for each community. Exponential-power law provided the best fit in 94% of all communities, then exponential (4%) and power law (2%) respectively. When exponential and power law forms were compared exclusively, exponential was better than power law in 81% of communities, with 19% communities fitting better in power law than exponential [see Jo.res function in msco R Package]. The “abc” values in square brackets constitute the 95% closed confidence interval of the parameter estimates (of these regression models) for all communities under each archetype. Archetypes A3, A6, and A7 were not observed in our data.

| Archetype | n | Exponential<br>$J^{(i)} = a \cdot \exp(b \cdot i)$ | | | Power law<br>$J^{(i)} = a \cdot i^b$ | | | Exponential-power law<br>$J^{(i)} = a \cdot \exp(b \cdot i) \cdot i^c$ | | | |
| --- | --- | --- | --- | --- | --- | --- | --- | --- | --- | --- | --- |
|  |  | % | a | b | % | a | b | % | a | b | c |
| A1 | 184 | 73.4 | [1.63, 44.14] | [-1.65, -0.03] | 26.6 | [1.53, 18.67] | [-49.8, -0.43] | 97.8 | [1.9, 32.93] | [-1.19, 0.13] | [-1.53, -0.09] |
| A2 | 11 | 100 | [5.35, 32.45] | [-1.22, -0.67] | 0 | [1.78, 14.54] | [-2.05, -1.42] | 100 | [4.74, 31.24] | [-1.1, -0.6] | [-0.24, 0.03] |
| A4 | 7 | 71.4 | [4.88, 58.18] | [-1.69, -0.27] | 28.6 | [2.72, 18.62] | [-2.62, -0.59] | 85.7 | [3.33, 62.01] | [-1.28, -0.02] | [-1.22, 0.14] |
| A5 | 82 | 95.1 | [2.49, 59.06] | [-2.13, -0.11] | 4.9 | [1.26, 12.18] | [-3.2, -0.39] | 87.8 | [2.56, 75.45] | [-2.07, 0.02] | [-0.49, 1.14] |
| A8 | 1 | 100 | [24.17, 24.17] | [-2.29, -2.29] | 0 | [2.45, 2.45] | [-3.43, -3.43] | 100 | [30.6, 30.6] | [-2.53, -2.53] | [0.35, 0.35] |
| A9 | 3 | 100 | [8.46, 81.73] | [-2.57, -0.98] | 0 | [2.35, 5.96] | [-3.81, -1.79] | 66.7 | [13.72, 44.01] | [-2, -1.47] | [-0.83, 0.83] |
| NA | 1 | 100 | [43.54, 43.54] | [-1.63, -1.63] | 0 | [8.55, 8.55] | [-2.56, -2.56] | 100 | [37.19, 37.19] | [-1.47, -1.47] | [-0.24, -0.24] |

The null model tests identified six archetypes in the 289 communities (A1, A2, A4, A5, A8 and A9; see Figure 3 and Table 1 for interpretations of archetypes), ranging from those that indicate positive species associations (A1) to those of negative species association (A9) among species residing respective communities. Archetype A1, under which species cooccurring more than expected by chance over all orders, dominated 63.7% of these communities. Of these A1 communities, the exponential-power law parametric model had the lowest AIC values in 98% (180) of cases among the three parametric models. When only power law and exponential forms were compared, power law dominated 27% while exponential 73% of A1 cases.

**Figure 3.**
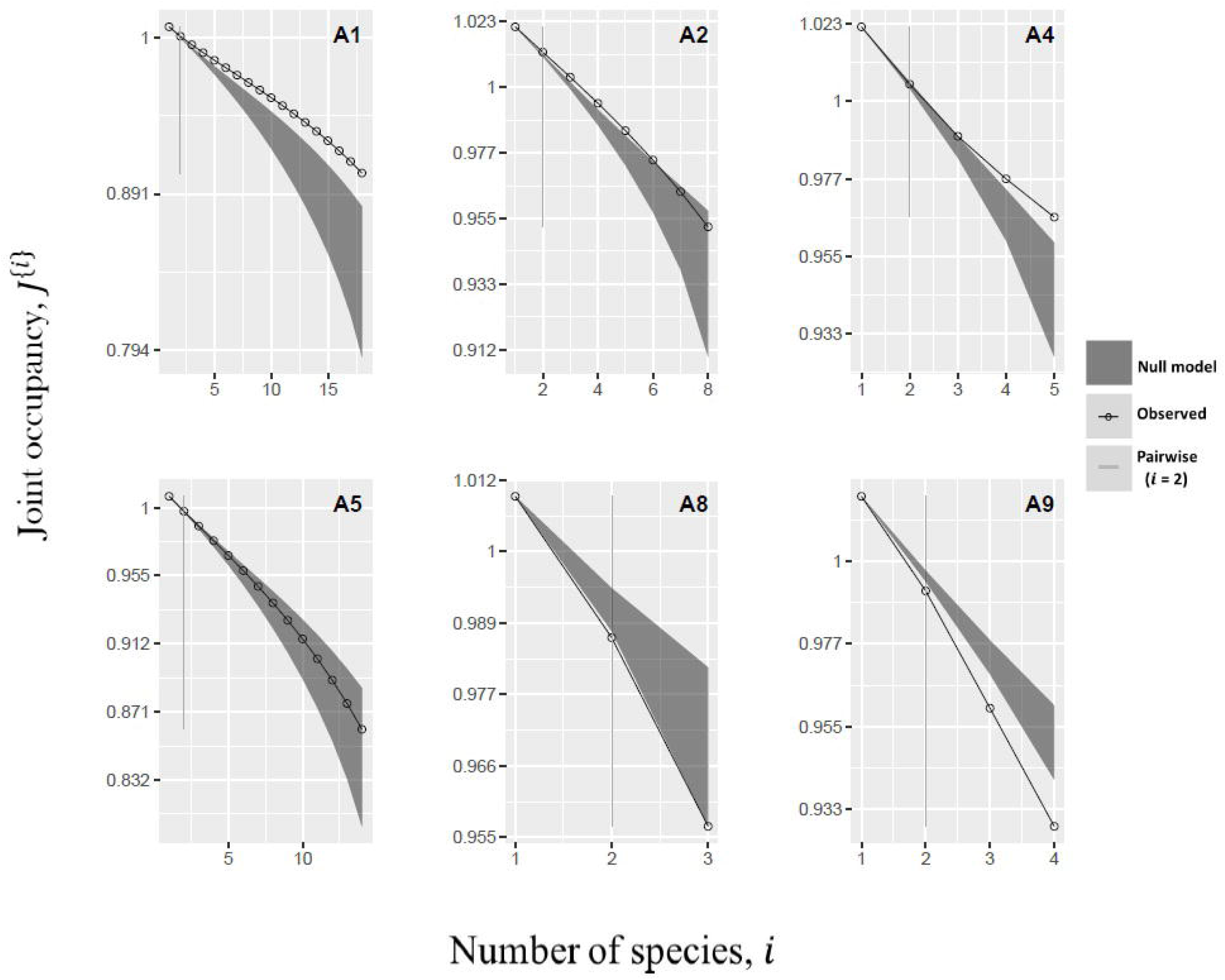
Archetypes of species co-occurrence patterns from the 289 empirical data matrices analyzed. The number of sites occupied by multiple species simultaneously (joint occupancy, *J*^{*i*}^; obtained from eqn 3) declines monotonically as the number of species (*i*) increase in all the archetypes. They range from communities whose species co-occurrences are positively associated more often than would be expected by chance (aggregation), to those that are negatively associated more often than by chance if species colonized sites randomly (segregation). A4 is typical of a community where a pairwise metric fails at detecting deterministic species co-occurrence patterns leading to type II statistical error. By using a multi-species joint occupancy index, the contributions made by the associations between multiple species towards the structure of ecological communities can be reckoned with.

A5 was the second most common archetype, characterizing 82 communities (28%). Under this archetype, observed co-occurrence does not differ significantly from null model expectations across all orders. Exponential form fitted better than power law in 95% of A5 communities (Table 2). There were only 22 communities following other four archetypes (A2=11, A4=7, A8=1, and A9=3; see Table 2). In particular, pairwise metrics (order 2) failed to detect non-stochastic forces at work for higher-order co-occurrence in the 7 A4 communities (i.e. type II error of false negative; Figure 3). There was one community that did not follow any of the nine archetypes (‘NA’ in Table 2), representing the combinations of several proposed archetypes and highlighting the possibility of rare but highly complex cooccurrence patterns in real communities (see community 80 in Appendix S3).

## 4. Concluding remarks

Inferring underlying processes from patterns requires to either examine multiple patterns capturing different pieces of information about the community structure (Grimm and Railsback, 2012; Latombe et al., 2011) or an informative pattern that can reflect and bridge many facets of biodiversity structure (Hui and McGeoch, 2014). The proposed multi-species joint occupancy index (*J*^{*i*}^) is such an informative and complete set of indices for describing species co-occurrences. Instead of only calculating the average over a set of pairwise (or any order *i* ≥ *m*) co-occurrences, we can compute the average over all sets of simultaneous species occurrences, and compare the joint occupancy decline to its null form, therefore capturing information on the identity of sites occupied by multiple species simultaneously. Since it reckons with all possible species simultaneous occurrences, this index can be used in testing for significant co-occurrence patterns among multiple species (e.g., diffuse competition) within a community, contrary to pairwise co-occurrence metrics.

The gain of information captured by our new joint occupancy metric is apparent in the nine basic archetypes of species co-occurrence we could identified, instead of three for pairwise metrics. Of the six archetypes detected in the 289 matrices, archetype A1 and A5 were the most common ones. Archetype A5 indicates that species co-occur as expected by chance when species occupancy is fixed, across all orders of joint occupancy. As archetype A5 depicts species co-occurrence patterns conforming to null model expectations across the full spectrum of orders, it represents the dominance of stochastic/neutral ecological processes in a community. Because the exponential form of joint occupancy decline fitted better than power law in 95% of A5 communities (Table 2), this particular form could signal stochastic co-occurrence among species. Hui and McGeoch (2014) found that stochastic community assembly can give rise to an exponential decline of compositional similarity, our results here point out to the possible connection between species turnover and co-occurrence in a community: an exponential decline of compositional similarity with zeta diversity order could correspond to an exponential decline of joint occupancy.

Archetype A1 indicates stronger species association than expected by chance across all orders of joint occupancy. This suggests that habitat filtering is an important driver of community assembly (Cordero and Jackson, 2019). A1 communities tend to have the largest variation in their within-site richness (Figure S1), whereas A5 communities, characterized by stochastic processes have some of the lowest variations in within-site richness. These results corroborate previous findings that environmental filtering can enhance habitat heterogeneity and consequently the gradient of species richness (Schuler et al., 2017). In contrast, archetype A9 indicates stronger species segregation than expected by chance across all orders of joint occupancy, signaling the work from intense inter-specific competition or other antagonistic interactions. Because A1 communities are much more common than A9 communities, environmental filters could be, in general, a more dominant driver of species co-occurrence patterns than competition, at least at the spatial scales represented in the dataset.

The fact that the vast majority of datasets belongs to categories A1, A5 or A9 indicates that the processes structuring a community tend to apply to all groups of species (e.g. functional guilds) composing the community, regardless of their size. The other archetypes observed in the dataset (A2, A7 and A8) are characteristic of communities with rarer assembly processes that change for different groups of species within the communities. Although they are quite rare, they represent special cases that can enable us to better understand when and why community assembly processes diverge from expectations, and may represent unique types of ecosystems. Archetypes A2 and A8 represent communities for which non-random co-occurrence of species is only observed for low orders of joint occupancy, and characterized 11 and one community, respectively. These archetypes indicate communities for which niche processes are at play, but only for part of the community, between a small number of species.

Two of the nine archetypes (A4 and A6) depict patterns of co-occurrence different from null expectations at only high orders, i.e. a type II error. Only one of these archetypes (A4) was observed in the dataset, characterizing 7 of the 289 matrices. It indicates the presence of complex interactions between large groups of species that are absent for small groups, an aspect that is not commonly explored in community ecology. Importantly, despite being relatively rare, the theoretical and empirical possibilities of encountering these archetypes illustrates the importance of using an appropriate metric for null model testing of species cooccurrence patterns to obtain a comprehensive perspective on community assembly enabling us to identify unique ecosystems.

The nine archetypes are certainly not exhaustive, and more complex patterns can exist (e.g. see community 80 in Appendix S3). Although there are still notable limitations to how different archetypes can be interpreted (even at orders >2, we can still face some of the same issues as pairwise indices, such as the fact that different groups of species may be driven by opposing positive and negative associations that can cancel each other out, making it difficult to identify any significant non-random patterns), multi-species joint occupancy nonetheless provides a comprehensive description of co-occurrence patterns and represents a firm step forward to better deciphering ecological processes at play in communities.

## Supporting information

Appendix S1: Results replication

Appendix S1: msco R package manual

Appendix S3: Results from 289 communities analyzed

## Acknowledgements

VKL is supported by the DSI-NRF Centre of Excellence in Mathematical and Statistical Sciences (CoE-MaSS) at Wits University and the Science Faculty Scholarship at Stellenbosch University. CH is supported by the National Research Foundation of South Africa (grant no. 89967). Opinions expressed and conclusions arrived at, are those of the authors and are not necessarily to be attributed to the CoE.

