## Appendix S1: Results replication for "A multi-species co-occurrence index to avoid type II errors in null model testing"

To access all the results presented in this paper, install the msco R Package (available on GitHub) using the following code:

```
devtools::install_github(repo = "vitaliskim/msco",
             dependencies = TRUE)
```

and execute the code below in R computational environment:

msco::Jo.res()

See the value of the Jo.res function in the msco R package manual attached with a code to replicate these results [which may also be accessed using: ?msco::Jo.res]. The manual can be recreated by running:

setwd("~/R/win-library/4.1/msco")

devtools::build_manual(pkg = ".")

The working directory in the above code can be set to where the package was installed, if it is not the default path specified above (for Windows OS, for example), assuming the R version used is 4.1
