## Appendix S1: msco R package manual for "A multi-species co-occurrence index to avoid type II errors in null model testing"

### Package ‘msco’

October 27, 2021

**Type** Package

**Title** Multi-Species Co-Occurrence Analyses

**Version** 0.1.0

**Author** Vitalis K. Lagat [aut, cre]
(<https://orcid.org/0000-0002-9063-9187>),
Guillaume Latombe [aut, ths]
(<https://orcid.org/0000-0002-8589-8387>),
Cang Hui [aut, ths]
(<https://orcid.org/0000-0002-3660-8160>)

**Maintainer** Vitalis K. Lagat <>

**Description** Co-occurrence patterns are typically explored between species pairs. However, doing so inevitably ignore the possible higher-order interference and interactions among three or more species in a community. To address this, the multi-species co-occurrence (msco) index, which calculates the joint occupancy, was recently proposed <DOI:to be added>. By reckoning with possible higher-order species interactions, the msco index can encapsulate nine possible archetypes of species co-occurrence patterns and avert the statistical type II error by examining patterns that would otherwise not be detected by any pairwise co-occurrence metrics. To dissect the effects of neutral encounter versus trait-based processes on multi-species co-occurrence and related higher-order interactions, generalised B-spline model has also been proposed <DOI: to be added>. The R package "msco" implements the multi-species co-occurrence index and includes a list of functions for calculating this index for different orders, as well as testing and visualizing species co-occurrence archetypes. This package also implements the generalized B-spline model and related analyses.

**Depends** R (>= 3.5.0)

**License** GPL-3

**Imports** scales, ggplot2, dplyr, minpack.lm, cowplot, gtools, tibble, tidyr, caret, pagenum, ape, import, taxize, glm2, splines2

**Encoding** UTF-8

**LazyData** true

**RoxygenNote** 7.1.2

### R topics documented:

**Index** **34**

---

`Arch_schem` *A schematic figure of the archetypes*

---

##### Description

A schematic diagram illustrating nine possible archetypes (from the null model test) of the patterns of species co-occurrences in ecological communities. The archetypes are denoted $\{Ai : i \in (1 : 9)\}$. See the details below.

##### Usage

```
Arch_schem()
```

##### Value

The `Arch_schem` function returns a **schematic diagram** of the archetypes of species co-occurrence patterns (denoted by $\{Ai : i \in (1 : 9)\}$), with the following components:

| **Archetype** | **Description/Interpretation** |
|---|---|
| `A1` | The joint occupancy value of the observed community matrix (observed; dark solid line) is above the null model. This means the null hypothesis (i.e. a statement that imply any change in the observed patterns do not reflect any community assembly process as underlying cause) should be rejected, confirming the presence of a mechanism of interest being tested (Lagat *et al.,* 2021a). It is typical of a community whose species are positively associated (or aggregated) more often than would be expected by chance. Such patterns of community structure may arise from a number of ecological mechanisms including environmental filtering or shared habitat requirements (Cordero and Jackson, 2019). |
| `A2` | The observed is greater than null expectation for `i = 2` but within null expectation for $i \geq 3$. This implies a pairwise metric detects a non-random pattern of the community structure, but when higher order species are considered, a random pattern is produced. This is typical of a community whose species are aggregated more often than by chance in sites with few species than in sites with many species (Lagat *et al.,* 2021a). |

| | |
|---|---|
| `A3` | The observed is greater than null expectation for lower orders, within null expectation for medium orders, and less than null expectation for higher orders. This means species co-occur more often than by chance in sites with few species, but are segregated more often than by chance in sites with many species, depicting a community structured by two different community assembly processes (Lagat *et al.,* 2021a). |
| `A4` | The observed is within null expectation for `i = 2` but greater than null expectation for $i \geq 3$. This means when use pairwise co-occurrence is used, the null hypothesis is not rejected, but when joint occupancy is used, the same null hypothesis is rejected. I.e., pairwise co-occurrence fails at detecting patterns of aggregation for sites with many species, i.e. a type II error or false negative (Lagat *et al.,* 2021a). |
| `A5` | The observed is within the null expectation for all orders $i \geq 2$, implying the test is not statistically significant. This has been ecologically inferred to mean ecological communities are random and that no community assembly processes or mechanisms influence their structure (Lagat *et al.,* 2021a; Cordero and Jackson, 2019; Gotelli and Sounding, 2001). |
| `A6` | The observed is within the null expectation for `i = 2` but less than the null expectation for $i \geq 3$. This means when use pairwise co-occurrence is used, the null hypothesis is not rejected, but when joint occupancy is used, the same null hypothesis is rejected. I.e., pairwise co-occurrence fails at detecting patterns of segregation for sites with many species, i.e. a type II error or false negative (Lagat *et al.,* 2021a). |
| `A7` | The observed is less than null expectation for lower orders, within null expectation for medium orders, and greater than null expectation for higher orders. Implying species are segregated more often than would be expected by chance in sites with few species, but co-occur more often than by chance in sites with many species, depicting a community structured by two different community assembly processes (Lagat *et al.,* 2021a). |
| `A8` | The observed is less than null expectation for `i = 2` but within null expectation for $i \geq 3$. This means a pairwise metric detects a non-random pattern of the community structure, but when higher order species are considered, a random pattern is produced. This is typical of a community whose species are segregated more often than by chance in sites with few species than in sites with many species (Lagat *et al.,* 2021a). |
| `A9` | The joint occupancy value of the observed community matrix (dark solid line) is below the null model. This means the null hypothesis should be rejected, confirming the presence of a mechanism of interest being tested (Lagat *et al.,* 2021a). It is typical of a community structured by inter-specific competition or limiting similarity, though predation might also generate similar patterns (Hein et al. 2014). |

#### Note

`Arch_schem` is not a generic function which can take in any dataset and give the outputs, but a path to a schematic diagram saved in this package. A representational figure from empirical, simulated or any known `.csv` binary data matrices can be accessed with Jo.plots function.

---

`cross_valid` *Cross validation of the generalised B-spline model*

---

#### Description

This function implements four different cross-validation techniques to evaluate the predictive ability of the generalised B-spline model (*sensu* Lagat *et al.,* 2021b). The four different techniques implemented are:

- validation set approach;
- k -fold;
- Leave-one-out-cross-validation (LOOCV), and
- Repeated k-fold.

#### Usage

```
cross_valid(gbsm_obj, type = "k-fold", p, k, k_fold.repeats)
```

#### Arguments

| | |
|---|---|
| `gbsm_obj` | An object of `class "gbsm"` (i.e., assigned to gbsm function). |
| `type` | The type of the cross-validation approach used. It must be ∈ {"validation.set", "k-fold", "LOOCV", "repeated.k-fold" } |
| `p` | The percentage (in decimal form) of data used in training the model. The value is used if the cross-validation approach implemented is "validation.set". |
| `k` | The value of `k` used in both "k-fold" and "repeated.k-fold" types of cross-validation. This value represents the number of subsets or groups that a given sample of data is to be split into. A value of 5 or 10 is used in practice, as it leads to an ideal bias-variance trade-off (Lagat *et al.,* 2021b). |
| `k_fold.repeats` | The number of replicates used in "repeated.k-fold" type of cross-validation. |

#### Details

The k-fold cross-validation approach is highly recommended due to its computational efficiency and an acceptable bias-variance trade-off, subject to the value of `k` chosen to be either 5 or 10 (Lagat *et al.,* 2021b). For more details on the other cross-validation approaches, see Lagat *et al.* (2021c).

#### Value

Depending on the type of cross-validation approach implemented, the `cross_valid` function returns:

- a `data.frame` with the following test errors (for "validation.set"):
  - `RMSE`: A root mean squared error;
  - `R_squared`: the Pearson's $r^2$, and
  - `MAE`: the mean absolute error.
- an `array` with test errors as above including the type of the regression model used, size of the samples, number of predictors, type of cross-validation performed, and summary of sample sizes (for "k-fold", "LOOCV", and "repeated.k-fold").

#### Examples

```
## Not run:

my.path <- system.file("extdata/gsmdat", package = "msco")
setwd(my.path)
s.data <- get(load("s.data.csv")) ## Species-by-site matrix
t.data <- get(load("t.data.csv")) ## Species-by-trait matrix
p.d.mat <- get(load("p.d.mat.csv")) ## Species-by-species phylogenetic distance matrix

gbsm_obj <- msco::gbsm(s.data, t.data, p.d.mat, metric= "Simpson_eqn", d.f=4,
 order.jo=3, degree=3, n=1000, b.plots=FALSE, bsplines="single", scat.plot=FALSE,
  response.curves=FALSE, leg=1, start=seq(-0.1, 0, length.out=(ncol(t.data)+2)*4+1))

val.set <- msco::cross_valid(gbsm_obj, type="validation.set", p=0.8)
val.set

kfold <- msco::cross_valid(gbsm_obj, type="k-fold", k=5)
kfold

loocv <- msco::cross_valid(gbsm_obj, type="LOOCV")
loocv

repeated.kfold <- msco::cross_valid(gbsm_obj, type="repeated.k-fold", k=5, k_fold.repeats=100)
repeated.kfold

## End(Not run)
```

gbsm *A generalised B-spline modelling for a set of neutral and trait-based variables*

#### Description

This function implements the generalised B-spline model (*sensu* Lagat *et al.*, 2021b) for dissecting the effects of neutral encounter versus functional traits on multi-order species interactions and co-occurrence. Generalized linear model (*sensu* Hastie and Tibshirani, 1986) with binomial variance distribution and log link functions employed, with predictors transformed using a linear combination of B-splines (*sensu* Curry and Schoenberg, 1988).

#### Usage

```
gbsm(
  s.data,
  t.data,
  p.d.mat,
  metric = "Simpson_eqn",
  d.f = 4,
  order.jo = 3,
  degree = 3,
  n = 1000,
  b.plots = TRUE,
  bsplines = "single",
  scat.plot = TRUE,
  response.curves = TRUE,
  leg = 1,
  start = seq(-0.1, 0, length.out = (ncol(t.data) + 2) * 4 + 1)
)
```

#### Arguments

| | |
|---|---|
| `s.data` | A species-by-site presence/absence `data.frame` with entries indicating occurrence (1) and non-occurrence (0) of species in a site. |
| `t.data` | A `data.frame` with traits as columns and species as rows. The species must be the same as in `s.data`. |
| `p.d.mat` | A symmetric `matrix` with dimension names as species and entries indicating the phylogenetic distance between any two of them (species). |
| `metric` | The type of rescaling applied to the joint occupancy metric. Available options are: `Simpson_eqn` for Simpson equivalent, `Sorensen_eqn` for Sorensen equivalent, and `raw` for the raw form of index without rescaling. |
| `d.f` | Degrees of freedom for B-splines. |
| `order.jo` | Specific number of species for which the joint occupancy is computed. To implement generalised B-spline modelling for multiple orders, see gbsm_m.orders function. |
| `degree` | Degree of the B-splines. |

n — Number of samples for which the joint occupancy is computed. These samples are non-overlapping. I.e., sampling is done without replacement. If the total number of combinations of i species chosen from the total species pool m, i.e. choose(m,i), is less than this value (n), choose(m,i) is used as the (maximum) number of samples one can set. Otherwise sampling without replacement is performed to select just the n samples.

b.plots — Boolean value indicating if B-spline basis functions should be plotted.

bsplines — This parameter indicates if a single or all B-spline curves should be plotted. If b.plots=TRUE and bsplines="single", the B-spline curves for the first predictor in t.data will be plotted. Any other value for bsplines (other than "single") results in the B-spline curves for all predictors being plotted.

scat.plot — Boolean value indicating if scatter plots between joint occupancy and its predicted values should be plotted.

response.curves — A boolean value indicating if all response curves should be plotted.

leg — Boolean value indicating if the legend of the gbsm outputs should be included in the plots. This parameter is added to help control the appearance of plots in gbsm_m.orders function.

start — Starting values for glm regression.

#### Value

gbsm function returns a list containing the following outputs:

order.jo — Order of joint occupancy

Predictors — Predictor variables used in GLM regression with binomial variance distribution function and log link function.

Responses — Response variables from GLM regression with binomial variance distribution function and log link function.

coeff — Coefficients of the generalized linear model used.

glm_obj — Generalized linear model used.

j.occs — Observed joint occupancies.

pred.j.occs — Predicted joint occupancies.

bs_pred — B-spline-transformed Predictors.

start — Starting values for the generalized linear model used.

var.expld — Amount of variation in joint occupancy explained by the Predictors. I.e., it is the Pearson's $r^2$ between the observed and predicted values of joint occupancy.

summary — summary of the regression results

#### Examples

```
## Not run:
 my.path <- system.file("extdata/gsmdat", package = "msco")
 setwd(my.path)
 s.data <- get(load("s.data.csv")) ## Species-by-site matrix
 t.data <- get(load("t.data.csv")) ## Species-by-trait matrix
 p.d.mat <- get(load("p.d.mat.csv")) ## Species-by-species phylogenetic distance matrix

 my.gbsm <- msco::gbsm(s.data, t.data, p.d.mat, metric = "Simpson_eqn",
  d.f=4, order.jo=3, degree=3, n=1000, b.plots=TRUE, scat.plot=TRUE,
   bsplines="single", response.curves=TRUE, leg=1,
    start=seq(-0.1, 0, length.out=(ncol(t.data)+2)*4+1))

 my.gbsm$bs_pred
 my.gbsm$Predictors
 my.gbsm$Responses
 my.gbsm$order.jo
 my.gbsm$var.expld

## End(Not run)
```

---

gbsm.res *Results on generalised B-spline modelling (presented in Lagat et al., 2021b)*

---

#### Description

This function allows the replication of the results on generalised B-spline modelling, presented in Lagat *et al.* (2021b). Executing gbsm.res() therefore gives these outputs that are saved as .RDS files in msco. If the codes that produced these (saved) outcomes are desired, the codes below are made available.

#### Usage

```
gbsm.res()
```

#### Value

Returns all the results presented in Lagat *et al.* (2021b). To replicate

- **Figs. 1**, **3**, **4**, **5**, and **Tables 1** and **S1**, execute the following code:

```
my.path <- system.file("extdata/gsmdat", package = "msco")
setwd(my.path)
s.data <- get(load("s.data.csv")) ##Species-by-site matrix
t.data <- get(load("t.data.csv")) ##Species-by-trait matrix
p.d.mat <- get(load("p.d.mat.csv")) ##Species-by-species phylogenetic distance matrix
RNGkind(sample.kind = "Rejection")
set.seed(0)
gb.res <- msco::gbsm_m.orders(s.data,
            t.data,
            p.d.mat,
```

```
                metric = "Simpson_eqn",
                orders = c(2:5, 8, 10, 15),
                d.f = 4,
                degree = 3,
                n = 1000,
                k = 5,
                p = 0.8,
                type = "k-fold",
                scat.plots = TRUE,
                response.curves = TRUE,
                j.occs.distrbn = TRUE,
                mp.plots = TRUE,
                start = seq(-0.1, 0, length.out=(ncol(t.data)+2)*4+1)
              )

gb.res$contbn_table$`order 3`  ## Table 1
gb.res$model.validation.table  ## Table S1
```

- **Figs. S1, 2,** and **S2**, execute the following codes:
  - **Fig. S1**:

```
remotes::install_github("jinyizju/V.PhyloMaker", force = TRUE)
library(V.PhyloMaker)
my.path <- system.file("extdata/gsmdat", package = "msco")
setwd(my.path)
s.data <- get(load("s.data.csv")) ##Species-by-site matrix
taxa <- get(load("taxa.levels.csv")) ##Species taxa
my.phylo.plot <- msco::s.phylo(s.data,
                          database = "ncbi",
                          obs.taxa = FALSE,
                          taxa.levels = taxa,
                          Obs.data = FALSE,
                          phy.d.mat = FALSE,
                          phylo.plot = TRUE)
```

  - **Fig. 2**:

```
my.path <- system.file("extdata/gsmdat", package = "msco")
setwd(my.path)
s.data <- get(load("s.data.csv")) ##Species-by-site matrix
t.data <- get(load("t.data.csv")) ##Species-by-trait matrix
p.d.mat <- get(load("p.d.mat.csv")) ##Species-by-species phylogenetic distance matrix
my.gbsm <- msco::gbsm(s.data,
                  t.data,
                  p.d.mat,
                  metric = "Simpson_eqn",
                  d.f = 4,
                  order.jo = 3,
                  degree = 3,
                  n = 1000,
                  b.plots = TRUE,
```

```
                              bsplines = "single",
                              scat.plot = FALSE,
                              response.curves = FALSE,
                              leg = 1,
                              start = seq(-0.1, 0, length.out=(ncol(t.data)+2)*4+1)
                            )
```

– **Fig. S2**:

```
my.path <- system.file("extdata/gsmdat", package = "msco")
setwd(my.path)
s.data <- get(load("s.data.csv")) ##Species-by-site matrix
t.data <- get(load("t.data.csv")) ##Species-by-trait matrix
p.d.mat <- get(load("p.d.mat.csv")) ##Species-by-species phylogenetic distance matrix
RNGkind(sample.kind = "Rejection")
set.seed(0)
pe <- msco::pred.error.bands(s.data,
                    t.data,
                    p.d.mat,
                    metric = "Simpson_eqn",
                    d.f = 4,
                    simm = 10,
                    orders = c(2:5, 8, 10, 15),
                    degree = 3,
                    n = 1000,
                    start = seq(-0.1, 0, length.out=(ncol(t.data)+2)*4+1)
                  )
```

**Caveat:** The above codes can collectively take approximately 7 minutes to execute (with prediction uncertainty plot taking 6 minutes alone). It took 7.3895 minutes to run (and output results) on a 64 bit system with 8 GB RAM and 3.60 GHz CPU.

#### Note

The function gbsm.res is not for general use. We included it in this package to help the readers of Lagat *et al.* (2021b) paper, who may want to get a deeper understanding of how the results presented in this paper were arrived at. It also allows deeper scrutiny of Lagat *et al.* (2021b)'s findings.

#### Examples

```
## Not run:

gbs.res <- msco::gbsm.res()
gbs.res$contbn_table$`order 3`
gbs.res$model.validation.table

## End(Not run)
```

`gbsm_m.orders` *Predictor's contribution and model performance assessment from the results on multiple orders of joint occupancy*

#### Description

This function implements the generalised B-spline model (gbsm; *sensu* Lagat et al., 2021b) for dissecting the effects of neutral encounter versus functional traits on multi-order species interactions and cooccurrence. Unlike gbsm that performs gbsm for a single order of species, gbsm_m.orders takes into account multiple orders of joint occupancy. In particular: for multiple joint occupancy orders, this function computes:

- each predictor's contribution to the explained variation in joint occupancy,
- the goodness-of-fit and model performance from cross-validation, and
- plots the:
  - response curves,
  - scatter plots (between the observed and predicted joint occupancy values),
  - histograms of the joint occupancy frequency distribution, and
  - model performance plots.

#### Usage

```
gbsm_m.orders(
  s.data,
  t.data,
  p.d.mat,
  metric = "Simpson_eqn",
  orders,
  d.f = 4,
  degree = 3,
  n = 1000,
  k = 5,
  p = 0.8,
  type = "k-fold",
  scat.plots = FALSE,
  response.curves = TRUE,
  j.occs.distrbn = FALSE,
  mp.plots = FALSE,
  start = seq(-0.1, 0, length.out = (ncol(t.data) + 2) * 4 + 1)
)
```

#### Arguments

`s.data` A species-by-site presence/absence `data.frame` with entries indicating occurrence (1) and non-occurrence (0) of species in a site.

`t.data` A `data.frame` with traits as columns and species as rows. The species must be the same as in `s.data`.

`p.d.mat` A symmetric `matrix` with `dimnames` as species and entries indicating the phylogenetic distance between any two of them (species).

| | |
|---|---|
| `metric` | As for gbsm. |
| `orders` | Specific number of species for which the joint occupancy is computed. |
| `d.f` | As for gbsm. |
| `degree` | As for gbsm. |
| `n` | As for gbsm. |
| `k` | As for cross_valid. |
| `p` | As for cross_valid. |
| `type` | As for cross_valid. |
| `scat.plots` | Boolean value indicating if scatter plots between joint occupancy and its predicted values should be plotted. |
| `response.curves` | A boolean value indicating if all response curves for all joint occupancy orders (`jo.orders`) should be plotted. |
| `j.occs.distrbn` | A boolean value indicating if the histograms of the frequency distribution of observed joint occupancy should be output. |
| `mp.plots` | A boolean value indicating if the model performance plots should be output. |
| `start` | As for gbsm. |

#### Value

`gbsm_m.orders` function returns a list containing the following outputs:

- `jo.orders`: A set of joint occupancy orders.
- `contrbn_table`: A `list` of `data.frames` consisting of:
  - `predictor`: A column of predictors.
  - `var.expld_M1`: A column of goodness-of-fit (I.e., the Pearson's $r^2$ between the observed and predicted values of joint occupancy when all predictors are used in the model.
  - `var.expld_M2`: The Pearson's $r^2$ between the observed and the predicted values of joint occupancy when all predictors except the predictor whose contribution is to be determined, are used in the model.
  - `contribution`: Each predictor's proportion of contribution in explaining joint occupancy. This value is given by:
    `contribution` = $\frac{var.expld_{M1} - var.expld_{M2}}{var.expld_{M1}}$
- `model.validation.table`: A `data.frame` with:
  - `orders`: Orders of joint occupancy used.
  - `Rsquared_gf`: Goodness-of-fit of the model. I.e., it is the Pearson's $r^2$ between the observed and predicted values of joint occupancy, for different orders.
  - `Rsquared_cv`: Model performance from cross-validation.
- `metric`: As for gbsm.
- `d.f`: As for gbsm.
- `n`: As for gbsm.
- `degree`: As for gbsm.
- `jo.orders`: Orders of joint occupancy used.

##### Examples

```
## Not run:
 my.path <- system.file("extdata/gsmdat", package = "msco")
 setwd(my.path)
 s.data <- get(load("s.data.csv")) ## Species-by-site matrix
 t.data <- get(load("t.data.csv")) ## Species-by-Trait matrix
 p.d.mat <- get(load("p.d.mat.csv")) ## Species-by-species phylogenetic distance matrix

 RNGkind(sample.kind = "Rejection")
 set.seed(0)
 jp <- msco::gbsm_m.orders(s.data, t.data, p.d.mat,
  metric="Simpson_eqn", orders = c(3:5, 8, 10, 15, 20), d.f=4,
   degree=3, n=1000, k=5, p=0.8, type="k-fold", scat.plots=TRUE,
    response.curves=TRUE, j.occs.distrbn=TRUE, mp.plots=TRUE,
     start=seq(-0.1, 0, length.out=(ncol(t.data)+2)*4+1))

 jp$contbn_table[[1]]
 jp$model.validation.table
 jp$jo.orders

 ## Close the open plots.gbsm.pdf file before running the 2nd example
 RNGkind(sample.kind = "Rejection")
 set.seed(0)
 jp2 <- msco::gbsm_m.orders(s.data, t.data, p.d.mat,
  metric="Sorensen_eqn", orders = c(3:5, 8, 10, 15, 20), d.f=4,
   degree=3, n=1000, k=5, p=0.8, type="k-fold", scat.plots=TRUE,
    response.curves=TRUE, j.occs.distrbn=TRUE, mp.plots=TRUE,
     start=seq(-0.1, 0, length.out=(ncol(t.data)+2)*4+1))

 jp2$contbn_table[[1]]
 jp2$model.validation.table
 jp2$jo.orders

## Close the open plots.gbsm.pdf file before running the 3rd example
 RNGkind(sample.kind = "Rejection")
 set.seed(0)
 jp3 <- msco::gbsm_m.orders(s.data, t.data, p.d.mat,
  metric="raw", orders = c(3:5, 8, 10, 15, 20), d.f=4,
   degree=3, n=1000, k=5, p=0.8, type="k-fold", scat.plots=TRUE,
    response.curves=TRUE, j.occs.distrbn=TRUE, mp.plots=TRUE,
     start=seq(-0.1, 0, length.out=(ncol(t.data)+2)*4+1))
```

```
 jp3$contbn_table[[1]]
 jp3$model.validation.table
 jp3$jo.orders

## End(Not run)
```

---

j.occ *Expected value of joint occupancy for order $i$ and its standard deviation*

---

#### Description

This function computes joint occupancy (the average number of sites harbouring a given number of $i$ species simultaneously), and its standard deviation.

#### Usage

```
j.occ(s.data, order, metric = "raw")
```

#### Arguments

| | |
|---|---|
| s.data | A species-by-site presence/absence matrix with entries indicating occurrence (1) and non-occurrence (0) of species in a site. |
| order | Specific number of species for which joint occupancy and its standard deviation is computed. |
| metric | The type of rescaling applied to the joint occupancy metric. Available options are: Simpson_eqn for Simpson equivalent, Sorensen_eqn for Sorensen equivalent, and raw for the raw form of index without rescaling. |

#### Value

Returns a list with the following outputs:

| | |
|---|---|
| jo.val | Joint occupancy value. |
| jo.sd | The standard deviation of jo.val. |

#### Examples

```
ex.data <- read.csv(system.file("extdata", "274.csv", package = "msco"))
jo <- msco::j.occ(ex.data, order = 3, metric = "raw")
jo

ex.data2 <- read.csv(system.file("extdata", "65.csv", package = "msco"))
jo2 <- msco::j.occ(ex.data2, order = 3, metric = "raw")
jo2
```

`j.occs` *Expected value of joint occupancy and its standard deviation for a range of orders*

#### Description

This function computes joint occupancy (the average number of sites harbouring a given number of $i$ species simultaneously) and its standard deviation for a range of orders (number of species).

#### Usage

```
j.occs(s.data, orders = 1:nrow(s.data), metric = "raw")
```

#### Arguments

| | |
|---|---|
| `s.data` | A species-by-site presence/absence matrix with entries indicating occurrence (1) and non-occurrence (0) of species in a site. |
| `orders` | Range number of species for which joint occupancy and its standard deviation is computed. |
| `metric` | The type of rescaling applied to the joint occupancy metric. Available options are: `Simpson_eqn` for Simpson equivalent, `Sorensen_eqn` for Sorensen equivalent, and `raw` for the raw form of index without rescaling. |

#### Value

Returns a `list` with the following outputs:

| | |
|---|---|
| `jo.vals` | A vector of joint occupancy values for a range number of species (in `orders`). |
| `jo.sds` | A vector of standard deviations of `jo.vals`. |

#### Examples

```
ex.data <- read.csv(system.file("extdata", "274.csv", package = "msco"))
jos <- msco::j.occs(ex.data, orders = 1:nrow(ex.data), metric = "raw")
jos

ex.data2 <- read.csv(system.file("extdata", "65.csv", package = "msco"))
jos2 <- msco::j.occs(ex.data2, orders = 1:nrow(ex.data), metric = "raw")
jos2
```

Jo.eng *Joint occupancy model engine*

#### Description

This function is the engine behind the null model testing of species co-occurrence patterns, and analyses of the joint occupancy decline and the parametric forms of this decline, for one particular community. In particular, Jo.eng:

- computes the joint occupancy (i.e. the number of sites or assemblages harbouring multiple species simultaneously);
- performs a null model test using the same index;
- fits the three regression models (exponential, power law and exponential-power law) to joint occupancy decline (*sensu* Lagat *et al.,* 2021a) with order (number of species);
- estimates the parameter values of these models;
- determines the best model among the three using AIC values;
- quantifies the performance of the fitted models using the Pearson's $r^2$;
- plots the joint occupancy decline regression and null models, and
- ascertains the archetypes of the patterns of species co-occurrences (from null model test) from which inferences on the type of drivers structuralising ecological communities can be made.

#### Usage

```
Jo.eng(
  s.data,
  algo = "sim2",
  metric = "raw",
  nReps = 999,
  dig = 3,
  s.dplot = FALSE,
  All.plots = TRUE,
  Jo.coeff = TRUE,
  my.AIC = TRUE,
  my.rsq = TRUE,
  Exp_Reg = TRUE,
  P.law_Reg = TRUE,
  Exp_p.l_Reg = TRUE,
  Obs.data = FALSE,
  Sim.data = FALSE,
  Jo_val.sim = FALSE,
  C.I_Jo_val.sim = FALSE,
  Jo_val.obs = TRUE,
  Metric = TRUE,
  Algorithm = TRUE,
  S.order = TRUE,
  nmod_stats = TRUE,
  Pt_Arch_Vals = TRUE,
  Atype = TRUE,
  p.n.plot = FALSE,
```

```
  trans = FALSE,
  lab = FALSE,
  leg = FALSE,
  m.n.plot = FALSE
)
```

#### Arguments

| | |
|---|---|
| `s.data` | A species-by-site presence/absence matrix with entries indicating occurrence (1) and non-occurrence (0) of species in a site. |
| `algo` | Randomisation algorithm used for the comparison with the null model. The possible options to choose from are: `sim1`, `sim2`, `sim3`, `sim4`, `sim5`, `sim6`, `sim7`, `sim8`, and `sim9`, all from Gotelli (2000). `sim2` is highly recommended (see Lagat *et al.,* 2021a). |
| `metric` | The type of rescaling applied to the joint occupancy metric. Available options are: `Simpson_eqn` for Simpson equivalent, `Sorensen_eqn` for Sorensen equivalent, and `raw` for the raw form of index without rescaling. |
| `nReps` | Number of simulations used in the null model test. |
| `dig` | The number of decimal places of the joint occupancy values (y axis) in the plots. The default is 3. |
| `s.dplot` | A Boolean indicating whether the standard deviation of multi-species co-occurrence index should be included in the plots of joint occupancy decline or not. |
| `All.plots` | A Boolean indicating whether joint occupancy decline regression and null model plots should be output. |
| `Jo.coeff` | A Boolean indicating if coefficient estimates of the joint occupancy decline regression models should be printed. |
| `my.AIC` | A Boolean indicating whether Akaike Information Criterion of the joint occupancy decline regression models should be output or not. |
| `my.rsq` | A Boolean indicating whether square of correlation coefficient between the observed and predicted values of joint occupancy should be output. |
| `Exp_Reg` | A Boolean indicating if exponential regression parametric model should be printed. |
| `P.law_Reg` | A Boolean indicating if power law regression parametric model should be printed. |
| `Exp_p.l_Reg` | A Boolean indicating if exponential-power law regression parametric model should be printed. |
| `Obs.data` | A Boolean indicating if observed/empirical data should be output. |
| `Sim.data` | A Boolean indicating if simulated/random data produced using any of the simulation algorithms should be output. |
| `Jo_val.sim` | A Boolean indicating if joint occupancy values of the simulated species-by-site presence/absence matrices should be output. |
| `C.I_Jo_val.sim` | A Boolean indicating if 95% confidence interval of the joint occupancy values of the simulated data should be printed. This interval is the area under the null model. |
| `Jo_val.obs` | A Boolean indicating if joint occupancy values of the observed species-by-site presence/absence matrices should be output. |
| `Metric` | A Boolean indicating if metric used should be printed. |
| `Algorithm` | A Boolean indicating if simulation algorithm used should be printed. |

| | |
|---|---|
| S.order | A Boolean indicating if the number of species whose joint occupancy is computed should be printed. |
| nmod_stats | A Boolean indicating whether the summary statistics for the null model test should be output. |
| Pt_Arch_Vals | A Boolean indicating if character strings indicating the location of joint occupancy value of the observed data relative to the critical values of the 95% closed confidence interval for every order (number of species), should be printed. |
| Atype | A Boolean indicating if a character string indicating the overall archetype of joint occupancy decline should be printed.This value must be $\in$ {"A1", "A2", "A3", "A4", "A5", "A6", "A7", "A8", "A9"} or "NA". "NA" could be the combinations of two or more of the nine expected archetypes. |
| p.n.plot | A Boolean indicating whether null model plot produced using the pairwise natural metric should be output. |
| trans | A Boolean indicating if the observed and simulated values used in p.n.plot should be transformed by raising them to (1/100). This can be done to compare p.n.plot with All.plots at a point where the order, i = 2. |
| lab | A Boolean indicating if the plot labels should be added to the m.n.plot. This parameter helps to control the appearance of plots in this function. |
| leg | A Boolean indicating if the legend should be added to the m.n.plot. This parameter helps to control the appearance of plots in this function. |
| m.n.plot | A Boolean indicating whether null model plot produced using joint occupancy metrics should be output. The default is FALSE. |

#### Value

Jo.eng function returns a list containing the following outputs:

| | |
|---|---|
| all.plots | Joint occupancy decline regression and null model plots. |
| jo.coeff | Coefficient estimates of the joint occupancy decline regression models. |
| AIC | Akaike information criterion of the joint occupancy decline regression models. |
| r2 | Square of correlation coefficient between the observed and predicted values of joint occupancy. |
| Exp_reg | Exponential regression parametric model. |
| P.law_reg | Power law regression parametric model. |
| Exp_p.l_reg | Exponential-power law regression parametric model. |
| Obs.data | Observed/empirical data. |
| Sim.data | Simulated/random data produced using any of the simulation algorithms. |
| jo.val.sim | Joint occupancy value of the simulated species-by-site presence/absence matrices. |
| C.I_Jo_val.sim | 95% confidence interval of the joint occupancy value of the simulated data. |
| jo.val.obs | joint occupancy value of the observed species-by-site presence/absence matrices. |
| Metric | Metric used. It must be "j.occ". |
| Algorithm | Simulation algorithm used. |
| nReps | Number of simulations performed. This value together with the joint occupancy value of the observed data, constitutes the sampling distribution. |

| | |
|---|---|
| s.order | Number of species whose joint occupancy is computed. |
| Pt_Arch_vals | Character strings indicating the location of joint occupancy value of the observed data relative to the critical values of the 95% closed confidence interval of the simulated data, for every order (number of species). |
| Archetype | A character string indicating the overall archetype from Pt_Arch_vals. It must be ∈ {"A1", "A2", "A3", "A4", "A5", "A6", "A7", "A8", "A9"} or "NA". "NA" could be the combinations of two or more of the nine expected archetypes (see Arch_schem). |

#### Examples

```
ex.data <- read.csv(system.file("extdata", "274.csv", package = "msco"))
j.en <- msco::Jo.eng(ex.data, algo="sim2", metric = "raw", nReps = 999,
           dig = 3, s.dplot = FALSE, All.plots = TRUE, Jo.coeff = TRUE,
           my.AIC = TRUE, my.rsq = TRUE, Exp_Reg = TRUE, P.law_Reg = TRUE,
           Exp_p.l_Reg = TRUE, Obs.data = FALSE, Sim.data = FALSE,
           Jo_val.sim = FALSE, C.I_Jo_val.sim = FALSE, Jo_val.obs = TRUE,
           Metric = TRUE, Algorithm = TRUE, S.order = TRUE,
           nmod_stats = TRUE, Pt_Arch_Vals = TRUE, Atype = TRUE,
           p.n.plot = FALSE, trans = FALSE, lab=FALSE, leg=FALSE, m.n.plot = FALSE)
j.en
```

---

### Jo.plots — *Joint occupancy parametric and null model plots*

---

#### Description

Plots the null model and joint occupancy decline with order (number of species) and fits the decline to exponential, power law and exponential-power law parametric models, respectively.

#### Usage

```
Jo.plots(jo_Obj)
```

#### Arguments

| | |
|---|---|
| jo_Obj | A joint occupancy model object returned by the function Jo.eng. |

#### Details

This function provides a visualization of the forms of joint occupancy decline and null model test. It offers information on:

- the outcomes of the null model test (through the appended archetype value on (e) plot) and
- the comparisons between the forms of joint occupancy decline (through the affixed AIC and rsq values on (b), (c) and (d) plots, respectively).

##### Value

Produces a figure consisting of the following plots:

- `(a)` Joint occupancy decline.
- `(b)` Exponential regression of the joint occupancy decline.
- `(c)` Power law regression of the joint occupancy decline.
- `(d)` Exponential-power law regression of the joint occupancy decline.
- `(e)` Null model test.

##### Examples

```
## Not run:

ex.data <- read.csv(system.file("extdata", "274.csv", package = "msco"))
jo_Obj <- msco::Jo.eng(ex.data, nReps = 999, All.plots = TRUE, s.dplot = FALSE, dig = 3)
jplots <- msco::Jo.plots(jo_Obj)
jplots

ex.data2 <- read.csv(system.file("extdata", "22.csv", package = "msco"))
jo_Obj2 <- msco::Jo.eng(ex.data2, nReps = 999, All.plots = TRUE, s.dplot = FALSE, dig = 3)
jplots2 <- msco::Jo.plots(jo_Obj2)
jplots2

ex.data3 <- read.csv(system.file("extdata", "78.csv", package = "msco"))
jo_Obj3 <- msco::Jo.eng(ex.data3, nReps = 999, All.plots = TRUE, s.dplot = FALSE, dig = 3)
jplots3 <- msco::Jo.plots(jo_Obj3)
jplots3

ex.data4 <- read.csv(system.file("extdata", "65.csv", package = "msco"))
jo_Obj4 <- msco::Jo.eng(ex.data4, nReps = 999, All.plots = TRUE, s.dplot = FALSE, dig = 3)
jplots4 <- msco::Jo.plots(jo_Obj4)
jplots4

## End(Not run)
```

---

#### `Jo.res` *Results on joint occupancy index (presented in Lagat et al., 2021a)*

---

##### Description

This function allows the replication of the results presented in Lagat *et al.* (2021a). Executing `Jo.res()` therefore gives these outputs that are saved as `.RDS` files in `msco`. If the codes that produced these (saved) outcomes are desired, the codes below are made available.

##### Usage

```
Jo.res()
```

##### Value

Returns all the results presented in Lagat *et al.* (2021a). To replicate these results, execute the following code:

```
RNGkind(sample.kind = "Rejection")
set.seed(39)
my.path <- system.file("extdata/myCSVs", package = "msco")
setwd(my.path)
my.files <- gtools::mixedsort(list.files(path = my.path, pattern = "*.csv"))
Lag.res <- msco::mJo.eng(my.files,
                 algo = "sim2",
                 metric = "raw",
                 m.Jo.plots = TRUE,
                 Archetypes = FALSE,
                 AICs = FALSE,
                 params = FALSE,
                 my.r2 = FALSE,
                 my.r2.s = TRUE,
                 best.mod2 = TRUE,
                 best.mod3 = TRUE,
                 params_c.i = TRUE
                );Lag.res
```

**Caveat:** The above code can take approximately 10 minutes to execute. It took 10.39014 minutes to run (and output results) on a 64 bit system with 8 GB RAM and 3.60 GHz CPU.

- **Fig. 3** can be replicated using:

```
path <- system.file("ms", package = "msco")
grDevices::pdf(file = paste0(path, "/real.arch.plots.pdf"),
 paper = "a4r", height = 8.27, width = 11.69)
msco:::nullmod_archs()
grDevices::dev.off()
system(paste0('open "', paste0(path, "/real.arch.plots.pdf"), '"'))
```

- **Fig. S1** can be replicated using:

```
my.path <- system.file("extdata/myCSVs", package = "msco")
setwd(my.path)
my.files <- gtools::mixedsort(list.files(path = my.path, pattern = "*.csv"))
grDevices::pdf(file = paste0(my.path, "/richns_cv.archs.pdf"),
 paper = "a4r", height = 5, width = 5)
richn.cv <- msco:::richness.variances(my.files)
grDevices::dev.off()
system(paste0('open "', paste0(my.path, "/richns_cv.archs.pdf"), '"'))
```

##### Note

The function Jo.res is not for general use. We included it in this package to help the readers of Lagat *et al.* (2021a) paper, who may want to get a deeper understanding of how the results presented in this paper were arrived at. It also allows deeper scrutiny of Lagat *et al.* (2021a)'s findings.

##### Examples

```
## Not run:

ms.res <- msco::Jo.res()
ms.res$r2.s
ms.res$best.mod2
ms.res$best.mod3
ms.res$params_c.i

## End(Not run)
```

---

mJo.eng *Joint occupancy model engine for multiple communities*

---

##### Description

This function is the engine behind the null model testing of species co-occurrence patterns, and analyses of the joint occupancy decline and the parametric forms of this decline, for multiple communities. In particular:

- It performs the null model testing of species co-occurrence patterns and generates the archetypes depicting how joint occupancy declines with the number of species (the order of msco) based on species-by-site presence/absence `.csv` data matrices. From these archetypes, inferences can be made according to the implemented null models;
- Determines the robustness of the exponential, power law and exponential-power law forms of joint occupancy decline by computing the Pearson's $r^2$ between the joint occupancy values of the observed data and predicted data, for all orders of species;
- Gives a summary of the total number of communities (under each and for all archetypes) whose forms of joint occupancy decline have $r^2 > 0.95$;
- Computes the AIC and Delta AIC of joint occupancy decline regression models for all communities;
- Computes the total number of communities:
  - with exponential as the best form of joint occupancy decline than power law and vice versa;
  - with either of the three regression models (exponential, power law and exponential-power law) having the best form of the joint occupancy decline;
- Estimates the parameters of:
  1. **exponential:** $j^{\{i\}} = a \times exp(b \times i)$;
  2. **power law:** $j^{\{i\}} = a \times i^b$; and
  3. **exponential-power law:** $j^{\{i\}} = a \times exp(b \times i) \times i^c$

  forms of joint occupancy decline, respectively, and their 95% confidence interval.

##### Usage

```
mJo.eng(
  my.files,
  algo = "sim2",
  metric = "raw",
  nReps = 999,
  Archetypes = FALSE,
  AICs = FALSE,
  params = FALSE,
  best.mod2 = FALSE,
  best.mod3 = FALSE,
  params_c.i = FALSE,
  my.r2 = FALSE,
  my.r2.s = FALSE,
  m.Jo.plots = FALSE
)
```

##### Arguments

| | |
|---|---|
| `my.files` | A vector containing names of species-by-site presence/absence `.csv` data matrices. The data matrices should be saved in the working directory. |
| `algo` | Simulation algorithm used. The possible options to choose from are: `sim1`, `sim2`, `sim3`, `sim4`, `sim5`, `sim6`, `sim7`, `sim8`, and `sim9`, all from Gotelli (2000). `sim2` is highly recommended (see Lagat *et al.,* 2021a). |
| `metric` | The type of rescaling applied to the joint occupancy metric. Available options are: `Simpson_eqn` for Simpson equivalent, `Sorensen_eqn` for Sorensen equivalent, and `raw` for the raw form of index without rescaling. |
| `nReps` | Number of simulations used in the null model test. |
| `Archetypes` | A Boolean indicating if the archetypes of the patterns of species co-occurrences in multiple communities should be included in the output. |
| `AICs` | A Boolean indicating whether the akaike information criterion (AIC) and Delta AIC of joint occupancy decline regression models for all communities should be included in the output. |
| `params` | A Boolean indicating whether parameter estimates of the joint occupancy decline regression models should be included in the output. |
| `best.mod2` | A Boolean indicating if exponential and power law regression model comparisons should be included in the output. |
| `best.mod3` | A Boolean indicating if exponential, power law and exponential-power law regression model comparisons should be included in the output. |
| `params_c.i` | A Boolean indicating if 95% C.I of the parameter estimates of the joint occupancy decline regression models should be included in the output. |
| `my.r2` | A Boolean indicating if the robustness of joint occupancy decline regression models should be computed and output. |
| `my.r2.s` | A Boolean indicating if the robustness summary values of joint occupancy decline regression models should be computed and output. |
| `m.Jo.plots` | A Boolean indicating whether joint occupancy parametric and null model plots for multiple communities should be included in the output. |

#### Details

mJo.eng function is useful when analyzing multiple species-by-site presence/absence data matrices at once. If one community matrix is analyzed, the outputs of the function Jo.eng should suffice.

#### Value

mJo.eng function returns a list containing the following outputs:

$Archs

For every community, a list consisting of:

- $nmod_stats: A data frame with the summary statistics for the null model test; and
- $Archetype: Archetypes of the patterns of species co-occurrences in ecological communities/matrices (my.files). These archetypes must be ∈ {"A1", "A2", "A3", "A4", "A5", "A6", "A7", "A8", "A9"} or "NA". "NA" could be the combinations of two or more of the nine expected archetypes.

$all.AICs

A list of data.frames containig the following components:

| | |
|---|---|
| df | The number of parameters in each of the three (exponential, power law and exponential-power law) joint occupancy decline regression models. |
| aic | The aic values for each of the three joint occupancy decline regression models. |
| delta_aic3 | The delta_aic values for each of the three joint occupancy decline regression models. |
| delta_aic2 | The delta_aic values for exponential and power law forms of joint occupancy decline regression models. |

$params

A data.frame consisting of:

| | |
|---|---|
| arch | The archetypes of the patterns of species co-occurrences in each of the species-by-site presence/absence .csv data matrices. |
| a.ex | The a parameter estimate of the exponential form of joint occupancy decline. |
| b.ex | The b parameter estimate of the exponential form of joint occupancy decline. |
| a.pl | The a parameter estimate of the power law form of joint occupancy decline. |
| b.pl | The b parameter estimate of the power law form of joint occupancy decline. |
| a.expl | The a parameter estimate of the exponential-power law form of joint occupancy decline. |
| b.expl | The b parameter estimate of the exponential-power law form of joint occupancy decline. |
| c.expl | The c parameter estimate of the exponential-power law form of joint occupancy decline. |

$best.mod2

Atable containig the following components:

| | |
|---|---|
| n | The number of ecological communities represented by species-by-site presence/absence .csv data matrices. |
| n.lwst_aic | The number of communities with exponential as the best form of joint occupancy decline than power law. |

n.delta_aic — The number of communities whose exponential and power law forms of joint occupancy decline have delta_aic = 0, respectively. This number must be equal to n.lwst_aic.

% — The percentage of n.lwst_aic (or n.delta_aic) relative to the total number of communities (n) analyzed.

$best.mod3

A table containig the following components:

n — The number of ecological communities represented by species-by-site presence/absence .csv data matrices.

n.lwst_aic — The number of communities with exponential or power law or exponential-power law as the best form of joint occupancy decline among the three (exponential, power law and exponential-power law) regression models.

n.delta_aic — The number of communities whose exponential, power law and exponential-power law forms of joint occupancy decline, respectively, have delta_aic = 0. This number must be equal to n.lwst_aic.

% — The percentage of n.lwst_aic (or n.delta_aic) relative to the total number of communities (n) analyzed.

$params_c.i

A data.frame consisting of:

arch — The archetypes of the patterns of species co-occurrences in each of the species-by-site presence/absence .csv data matrices.

n — The number of communities under every archetype.

ex_% — The percentages of the number of communities (under every archetype) where exponential form of joint occupancy decline fitted better than power law.

a.ex — The 95% closed confidence interval of the a parameter estimates of the exponential form of joint occupancy decline, under every archetype.

b.ex — The 95% closed confidence interval of the b parameter estimates of the exponential form of joint occupancy decline, under every archetype.

p.l_% — The percentages of the number of communities (under every archetype) where power law form of joint occupancy decline fitted better than exponential.

a.pl — The 95% closed confidence interval of the a parameter estimates of the power law form of joint occupancy decline, under every archetype.

b.pl — The 95% closed confidence interval of the b parameter estimates of the power law form of joint occupancy decline, under every archetype.

ex.pl_% — The percentages of the number of communities (under every archetype) where exponential-power law form of joint occupancy decline fitted better than both the exponential and power law forms.

a.expl — The 95% closed confidence interval of the a parameter estimates of the exponential-power law form of joint occupancy decline, under every archetype.

b.expl — The 95% closed confidence interval of the b parameter estimates of the exponential-power law form of joint occupancy decline, under every archetype.

c.expl — The 95% closed confidence interval of the c parameter estimates of the exponential-power law form of joint occupancy decline, under every archetype.

`$r2`

A `list` of `data.frames` containig the following components:

| | |
|---|---|
| `rsq.ex` | $r^2$ for the exponential form of joint occupancy decline. |
| `rsq.pl` | $r^2$ for the power law form of joint occupancy decline. |
| `rsq.ex.pl` | $r^2$ for the exponential-power law form of joint occupancy decline. |

`$r2.s`

- A `list` containig the following components:
  `$rsq.per.Archs`
  - `Archs`: Archetypes of the patterns of species co-occurrences in each of the species-by-site presence/absence .csv data matrices.
  - `n.a`: Number of communities under each archetype.
  - `rsq.ex`: Number of communities under each archetype whose exponential forms of joint occupancy decline have $r^2 > 0.95$.
  - `rsq.pl`: Number of communities under each archetype whose power law forms of joint occupancy decline have $r^2 > 0.95$.
  - `rsq.ex-pl`: Number of communities under each archetype whose exponential-power law forms of joint occupancy decline have $r^2 > 0.95$.

  `$rsq.all.Communities`
  - `n`: Number of all communities analyzed
  - `ex`: Number of communities whose exponential forms of joint occupancy decline have $r^2 > 0.95$
  - `pl`: Number of communities whose power law forms of joint occupancy decline have $r^2 > 0.95$
  - `ex.pl`: Number of communities whose exponential-power law forms of joint occupancy decline have $r^2 > 0.95$

`$m.Jo.plots`

Produces a `.pdf` file with multiple figures each consisting of the following plots:

| | |
|---|---|
| `(a)` | as for Jo.plots |
| `(b)` | as for Jo.plots |
| `(c)` | as for Jo.plots |
| `(d)` | as for Jo.plots |
| `(e)` | as for Jo.plots |

#### Examples

```
## Not run:

my.path <- system.file("extdata", package = "msco")
setwd(my.path)
my.files <- gtools::mixedsort(list.files(path = my.path, pattern = "*.csv"))
my.res <- msco::mJo.eng(my.files = my.files, algo = "sim2", Archetypes = TRUE,
             metric = "raw", nReps = 999, AICs = FALSE, params = FALSE,
             best.mod2 = FALSE, best.mod3 = FALSE, params_c.i = FALSE,
             my.r2 = FALSE, my.r2.s = FALSE, m.Jo.plots = FALSE)
my.res$Archs$`252.csv`

my.path2 <- system.file("extdata/myCSVs", package = "msco")
setwd(my.path2)
my.files2 <- gtools::mixedsort(list.files(path = my.path2, pattern = "*.csv"))
my.res2 <- msco::mJo.eng(my.files = my.files2[250:255], algo = "sim2", Archetypes = FALSE,
              metric = "raw", nReps = 999, AICs = FALSE, params = TRUE,
              best.mod2 = FALSE, best.mod3 = FALSE, params_c.i = FALSE,
              my.r2 = FALSE, my.r2.s = FALSE, m.Jo.plots = FALSE)
my.res2

my.path2 <- system.file("extdata/myCSVs", package = "msco")
setwd(my.path2)
my.files2 <- gtools::mixedsort(list.files(path = my.path2, pattern = "*.csv"))
my.res3 <- msco::mJo.eng(my.files = my.files2[250:255], algo = "sim2", Archetypes = FALSE,
              metric = "raw", nReps = 999, AICs = FALSE, params = FALSE,
              best.mod2 = FALSE, best.mod3 = FALSE, params_c.i = TRUE,
              my.r2 = FALSE, my.r2.s = FALSE, m.Jo.plots = TRUE)
my.res3

## End(Not run)
```

---

msco.res *Results on* msco *illustration (presented in Lagat et al., 2021c)*

---

#### Description

This function allows the replication of the results on msco R package illustration paper presented in Lagat *et al.* (2021c). Executing msco.res() therefore gives these outputs that are saved as .RDS files in msco. If the codes that produced these (saved) outcomes are desired, the codes below are made available.

#### Usage

```
msco.res()
```

#### Value

Returns all the results presented in Lagat *et al.* (2021c). To replicate

- **Figs. 1**, **2** and **Table 2**, execute the following code:

```
RNGkind(sample.kind = "Rejection")
set.seed(14)
ex.data <- read.csv(system.file("extdata", "251.csv", package = "msco"))
j.en <- msco::Jo.eng(ex.data,
            algo = "sim2",
            metric = "raw",
            nReps = 999,
            dig = 3,
            s.dplot = FALSE,
            All.plots = TRUE,
            Jo.coeff = TRUE,
            my.AIC = TRUE,
            my.rsq = TRUE,
            Exp_Reg = TRUE,
            P.law_Reg = TRUE,
            Exp_p.l_Reg = TRUE,
            Obs.data = FALSE,
            Sim.data = FALSE,
            Jo_val.sim = FALSE,
            lab = FALSE,
            leg = FALSE,
            C.I_Jo_val.sim = FALSE,
            Jo_val.obs = TRUE,
            Metric = TRUE,
            Algorithm = TRUE,
            S.order = TRUE,
            nmod_stats = TRUE,
            Pt_Arch_Vals = TRUE,
            Atype = TRUE,
            p.n.plot = TRUE,
            trans = FALSE,
            m.n.plot = FALSE)

j.en$nmod_stats ## Table 2
grDevices::pdf(file = paste0(system.file("ms", package = "msco"),
 "/aJo.plots.pdf"), paper = "a4r", height = 8.27, width = 11.69)
j.en$all.plots
grDevices::dev.off()
system(paste0('open "', paste0(system.file("ms", package = "msco"), ## Fig. 2
 "/aJo.plots.pdf"), '"'))
```

- **Fig. 4**, execute the following code:

```
RNGkind(sample.kind = "Rejection")
set.seed(14)
grDevices::pdf(file = paste0(system.file("ms", package = "msco"),
 "/real.arch.plots2.pdf"), paper = "a4r", height = 8.27, width = 11.69)
msco:::nullmod_archs2()
grDevices::dev.off()
system(paste0('open "', paste0(system.file("ms", package = "msco"),
 "/real.arch.plots2.pdf"), '"'))
```

- **Fig. 5**, execute the following code:

```
my.path <- system.file("extdata/gsmdat", package = "msco")
setwd(my.path)
s.data <- get(load("s.data.csv")) #Species-by-site matrix
t.data <- get(load("t.data.csv")) #Species-by-trait matrix
p.d.mat <- get(load("p.d.mat.csv")) #Species-by-species phylogenetic distance matrix
RNGkind(sample.kind = "Rejection")
set.seed(0)
gb.res <- msco::gbsm_m.orders(s.data,
            t.data,
            p.d.mat,
            metric = "Simpson_eqn",
            orders = c(3:5, 8, 10, 15, 20),
            d.f = 4,
            degree = 3,
            n = 1000,
            k = 5,
            p = 0.8,
            type = "k-fold",
            scat.plots = FALSE,
            response.curves = TRUE,
            j.occs.distrbn = FALSE,
            mp.plots = FALSE,
            start = seq(-0.1, 0, length.out=(ncol(t.data)+2)*4+1)
          )
```

#### Note

The function msco.res is not for general use. We included it in this package to help the readers of Lagat *et al.* (2021c) paper, who may want to get a deeper understanding of how the results presented in this paper were arrived at. It also allows deeper scrutiny of Lagat *et al.* (2021c)'s findings, and broader understanding of the main functionalities of `msco` R package.

#### Examples

```
## Not run:

ms.res <- msco::msco.res()
ms.res$nmod_stats ## Table 2
```

```
## End(Not run)
```

---

`pred.error.bands` *Prediction uncertainty*

---

#### Description

This function plots the response curves showing the effect of the predictors (i.e. trait-based and neutral forces) on joint occupancy as the response variable, with prediction error bands (as the standard deviation from the mean of the response variable) for all orders of joint occupancy.

#### Usage

```
pred.error.bands(
  s.data,
  t.data,
  p.d.mat,
  metric = "Simpson_eqn",
  d.f = 4,
  simm = 10,
  orders,
  degree = 3,
  n = 1000,
  start = seq(-0.1, 0, length.out = (ncol(t.data) + 2) * 4 + 1)
)
```

#### Arguments

| | |
|---|---|
| `s.data` | A species-by-site presence/absence `data.frame` with entries indicating occurrence (1) and non-occurrence (0) of species in a site. |
| `t.data` | A `data.frame` with traits as columns and species as rows. The species must be the same as in `s.data`. |
| `p.d.mat` | A symmetric `matrix` with `dimnames` as species and entries indicating the phylogenetic distance between any two of them (species). |
| `metric` | As for gbsm_m.orders. |
| `d.f` | As for gbsm_m.orders. |
| `simm` | Number of Monte Carlo simulations performed |
| `orders` | As for gbsm_m.orders |
| `degree` | As for gbsm_m.orders. |
| `n` | As for gbsm_m.orders. |
| `start` | As for gbsm_m.orders. |

#### Value

`pred.error.bands` function returns:

| | |
|---|---|
| `predictors` | a `data.frame` of predictors |
| `responses` | a `data.frame` of response values of predictors |

responses.sim_stats
: a data.frame of the reponses' mean and standard deviation (from simm replicates), and

- the response curves with prediction error bands for all orders of joint occupancy

#### Examples

```
## Not run:
 my.path <- system.file("extdata/gsmdat", package = "msco")
 setwd(my.path)
 s.data <- get(load("s.data.csv")) ## Species-by-site matrix
 t.data <- get(load("t.data.csv")) ## Species-by-Trait matrix
 p.d.mat <- get(load("p.d.mat.csv")) ## Species-by-species phylogenetic distance matrix

 RNGkind(sample.kind = "Rejection")
 set.seed(0)
pe <- msco::pred.error.bands(s.data, t.data, p.d.mat, metric="Simpson_eqn", d.f=4, simm=10,
  orders = c(2:5, 8, 10, 15), degree=3, n=1000,
   start=seq(-0.1, 0, length.out=(ncol(t.data)+2)*4+1))

 pe$predictors$`order 2`
 pe$responses$`order 2`
 pe$responses.sim_stats$`order 2`

 pe$predictors$`order 3`
 pe$responses$`order 3`
 pe$responses.sim_stats$`order 3`

 pe$predictors$`order 10`
 pe$responses$`order 10`
 pe$responses.sim_stats$`order 10`

## End(Not run)
```

---

s.phylo *Species phylogeny generator*

---

#### Description

This function generates the phylogeney of species and plots the phylogenetic tree. In particular, given a species-by-site matrix (community), s.phylo:

- uses the tax_name function to obtain (from the NCBI or ITIS online databases) the genus and family taxa levels of species in the community. If NCBI is used, getting an API key is recommended. See tax_name for more information. NCBI is used as default in this function;
- uses the `phylo.maker` function to obtain the phylogeny (an object of `class: "phylo"`) of species in the community using taxa obtained above;
- computes the phylogenetic distance matrix using the cophenetic.phylo function and the phylogeny obtained above as input;
- plots the phylogenetic tree using the phylogeny obtained above.

#### Usage

```
s.phylo(
  s.data,
  database = "ncbi",
  obs.taxa = TRUE,
  taxa.levels = NULL,
  Obs.data = TRUE,
  phy.d.mat = TRUE,
  phylo.plot = TRUE
)
```

#### Arguments

| | |
|---|---|
| `s.data` | A species-by-site presence/absence `data.frame` with entries indicating occurrence (1) and non-occurrence (0) of species in a site. The rows should have species' scientific names following binomial nomenclature, with no initials. |
| `database` | The online database used to obtain the taxonomic names (species, genus and family) for a given rank (species list in this function). The options are "ncbi" (default) or "itis". |
| `obs.taxa` | A Boolean indicating if `taxa.levels` should be included in the returned list. |
| `taxa.levels` | Species taxa (i.e. a `data.frame` with species, genus, and family as `colnames`) used in extracting phylogenetic distance matrix between species. If supplied, `taxa.levels` won't be computed from online repositories. Taxa provision is highly recommended. |
| `Obs.data` | A Boolean indicating if `s.data` should be included in the returned list. |
| `phy.d.mat` | A Boolean indicating if phylogenetic distance matrix should be in the returned list. |
| `phylo.plot` | Boolean value indicating if the phylogenetic tree (cluster dendrogram) should be plotted. |

#### Value

Returns a `list` with the following outputs:

- `s.data`: A `data.frame` with sites as columns and species as rows.
- `taxa.levels`: A `data.frame` with the following columns:
  - `species`: Species names in `s.data`.
  - `genus`: Genus names of species in `s.data`.
  - `family`: Family names of species in `s.data`.

- p.d.matrix: A symmetric matrix with dimension names as species and entries indicating the phylogenetic distance between any two of them (species).
- phylo.plot: A phylogenetic tree (cluster dendrogram) of species in s.data

#### Examples

```
## Not run:

remotes::install_github("jinyizju/V.PhyloMaker", force = TRUE)
library(V.PhyloMaker)
my.path <- system.file("extdata/gsmdat", package = "msco")
setwd(my.path)
s.data <- get(load("s.data.csv"))
taxa <- get(load("taxa.levels.csv"))

my.s.phylo <- msco::s.phylo(s.data, database = "ncbi", obs.taxa=TRUE,
 taxa.levels = taxa, Obs.data=TRUE, phy.d.mat=TRUE, phylo.plot = TRUE)

my.s.data <- my.s.phylo$s.data
my.s.data

my.taxa <- my.s.phylo$taxa.levels
my.taxa

my.p.d.mat <- my.s.phylo$phylogenetic.distance.matrix
my.p.d.mat

## End(Not run)
```

### Index

Arch_schem, 2, *18*

cophenetic.phylo, *31*
cross_valid, 4, *11*

gbsm, *4*, 5, *10–12*
gbsm.res, 8, *10*
gbsm_m.orders, *6, 7*, *10*, 10, *30*

j.occ, 13
j.occs, 14
Jo.eng, *15*, 15, *19*, *23*
Jo.plots, *3*, 19, *26*
Jo.res, 20, *21*

mJo.eng, 22
msco.res, 27, *29*

pred.error.bands, 29

s.phylo, *31*, 31

tax_name, *31*
