## Appendix S2: Figure S1 for "A multi-species co-occurrence index to avoid type II errors in null model testing"

**Appendix S2: Richness vs Archetype variation**

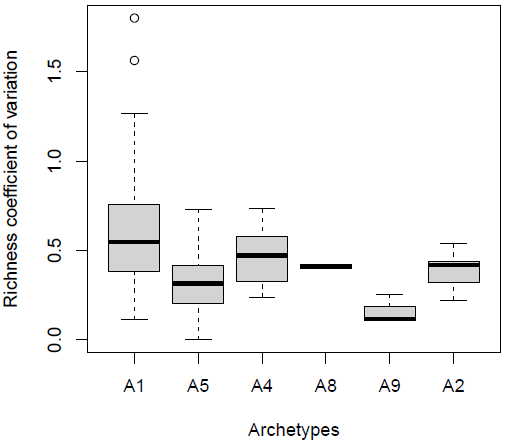

**Figure S1.** Richness coefficient of variation boxplots for different archetypes using empirical data (from Atmar and Patterson, 1995).
