## Appendix S3: Results from 289 communities analyzed for "A multi-species co-occurrence index to avoid type II errors in null model testing"

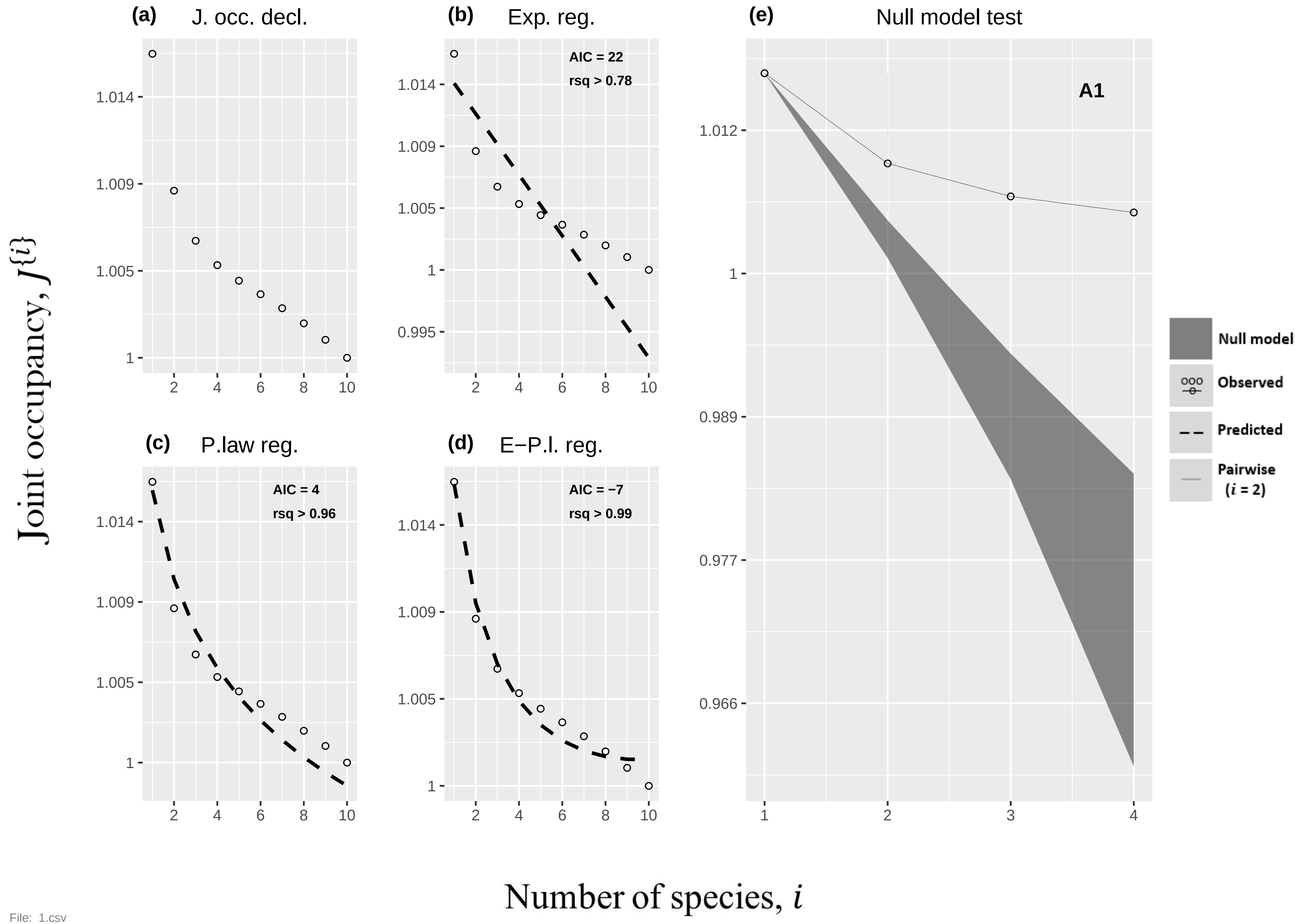

(a) J. occ. decl.
(b) Exp. reg.
AIC = 22
rsq > 0.78
(c) P.law reg.
AIC = 4
rsq > 0.96
(d) E−P.l. reg.
AIC = −7
rsq > 0.99
(e) Null model test
A1
Null model
Observed
Predicted
Pairwise (i = 2)
Joint occupancy, J{i}
Number of species, i


File: 1.csv

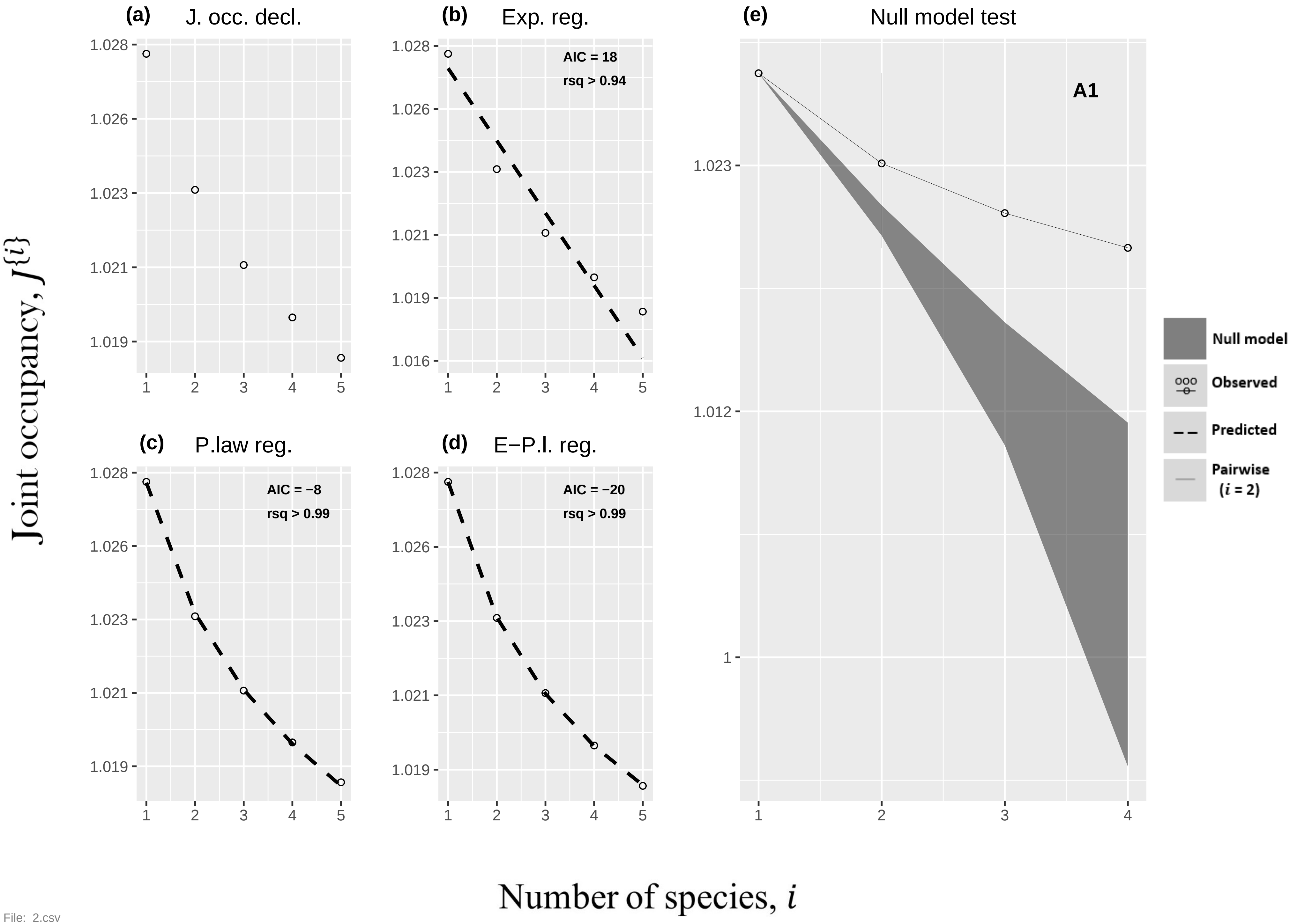
(a) J. occ. decl.
(b) Exp. reg.
AIC = 18
rsq > 0.94
(c) P.law reg.
AIC = −8
rsq > 0.99
(d) E−P.l. reg.
AIC = −20
rsq > 0.99
(e) Null model test
A1
Null model
Observed
Predicted
Pairwise (i = 2)
Joint occupancy, J{i}
Number of species, i

File: 2.csv

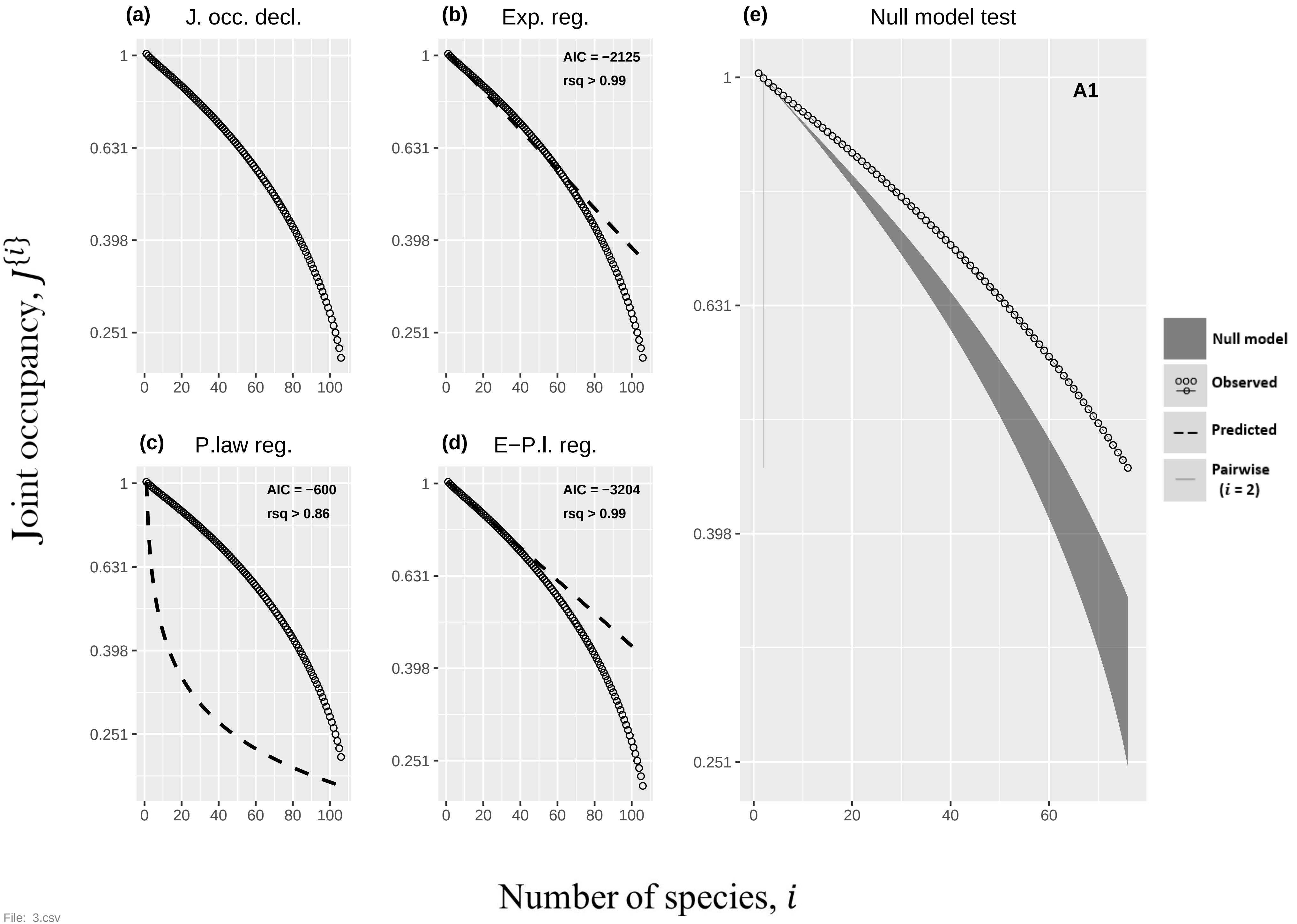
(a) J. occ. decl.
(b) Exp. reg.
AIC = −2125
rsq > 0.99
(c) P.law reg.
AIC = −600
rsq > 0.86
(d) E−P.l. reg.
AIC = −3204
rsq > 0.99
(e) Null model test
A1
Null model
Observed
Predicted
Pairwise (i = 2)
Joint occupancy, J{i}
Number of species, i
1
0.631
0.398
0.251
0
20
40
60
80
100

File: 3.csv

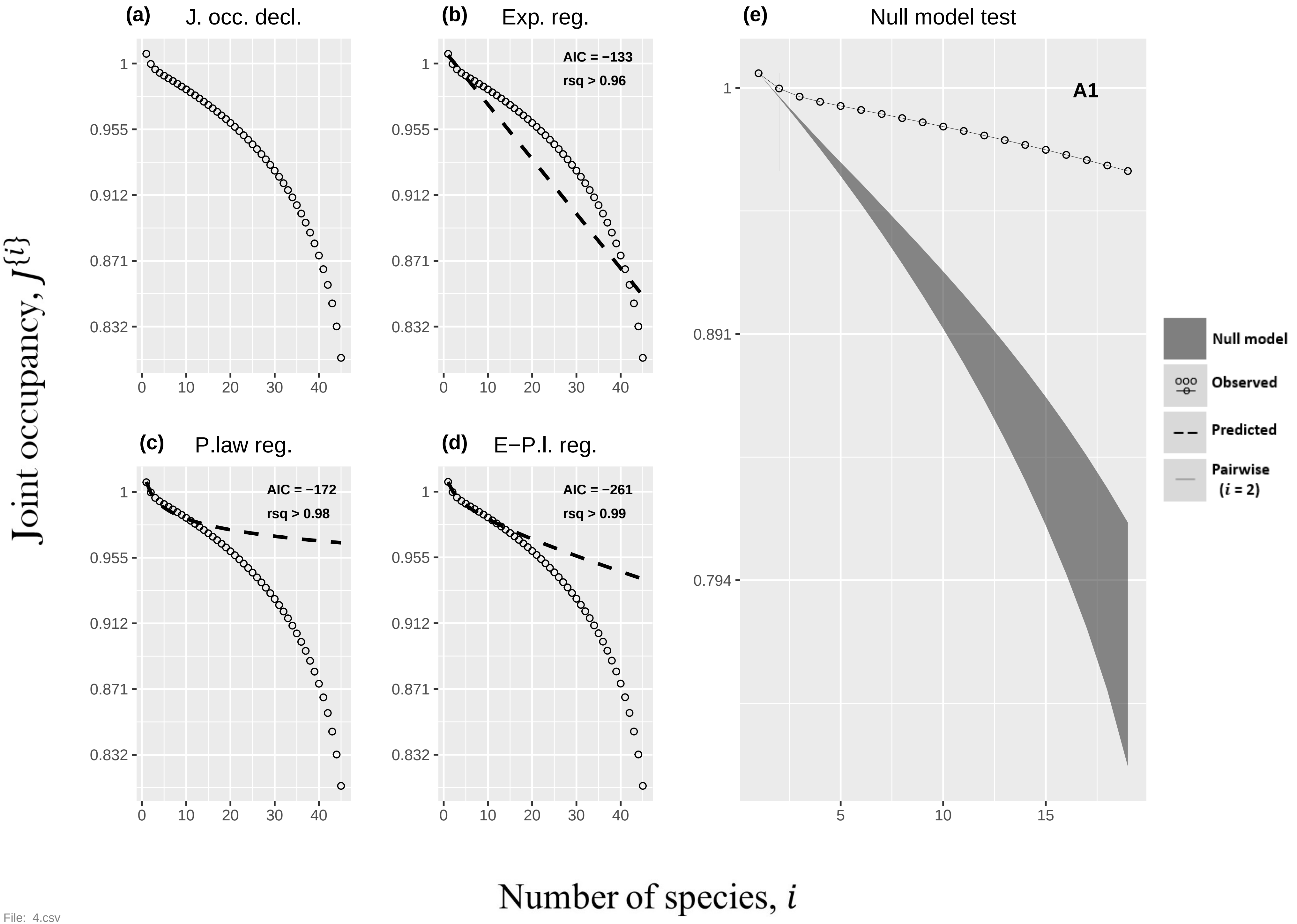

(a) J. occ. decl.
(b) Exp. reg.
AIC = −133
rsq > 0.96
(c) P.law reg.
AIC = −172
rsq > 0.98
(d) E−P.l. reg.
AIC = −261
rsq > 0.99
(e) Null model test
A1
Null model
Observed
Predicted
Pairwise ($i$ = 2)
1
0.955
0.912
0.871
0.832
0
10
20
30
40
0.891
0.794
5
15
Joint occupancy, $J^{\{i\}}$
Number of species, $i$


File: 4.csv

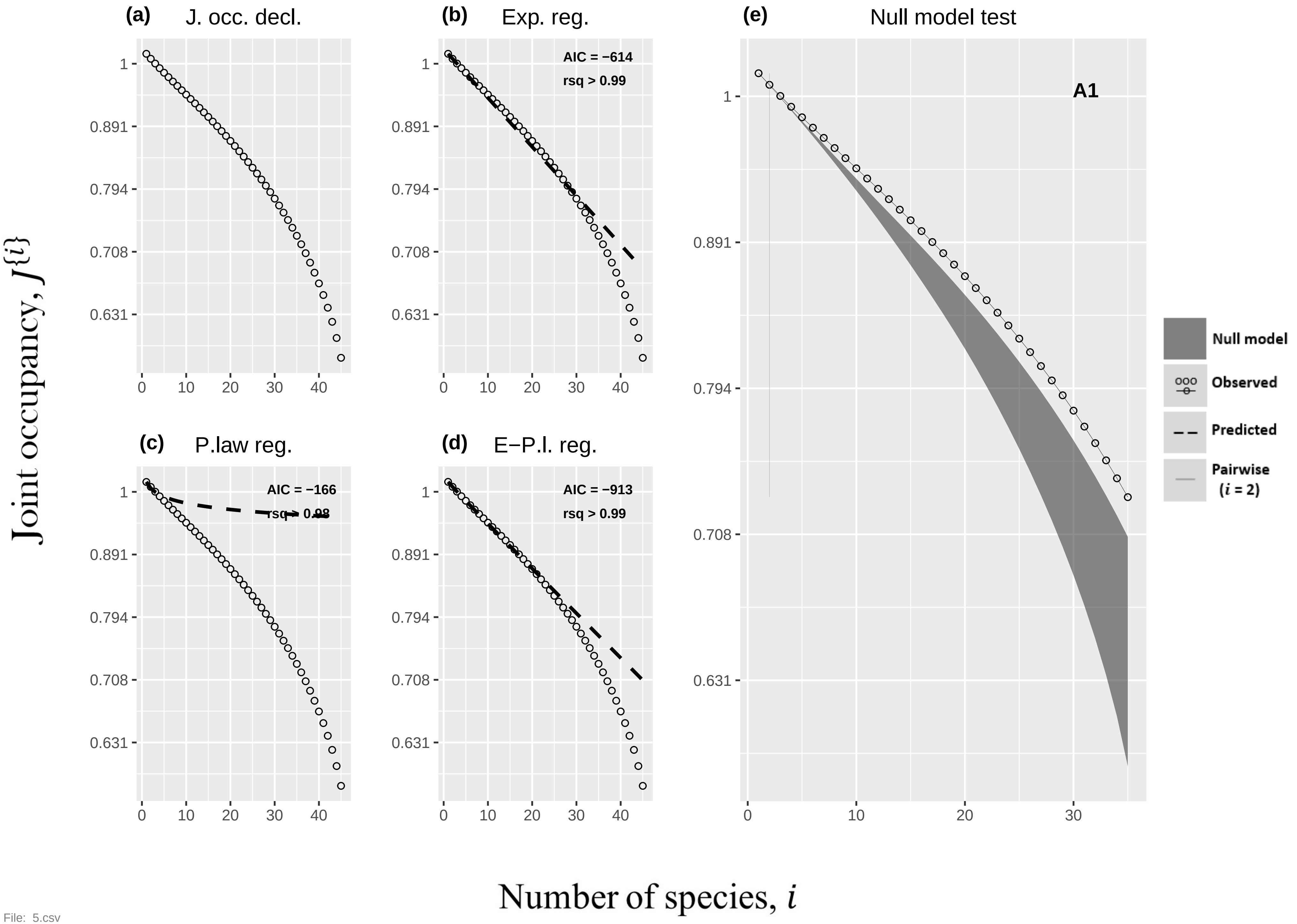

(a) J. occ. decl.
(b) Exp. reg.
AIC = −614
rsq > 0.99
(c) P.law reg.
AIC = −166
rsq > 0.98
(d) E−P.l. reg.
AIC = −913
rsq > 0.99
(e) Null model test
A1
Null model
Observed
Predicted
Pairwise
(i = 2)
Joint occupancy, J{i}
Number of species, i
1
0.891
0.794
0.708
0.631
0
10
20
30
40


File: 5.csv

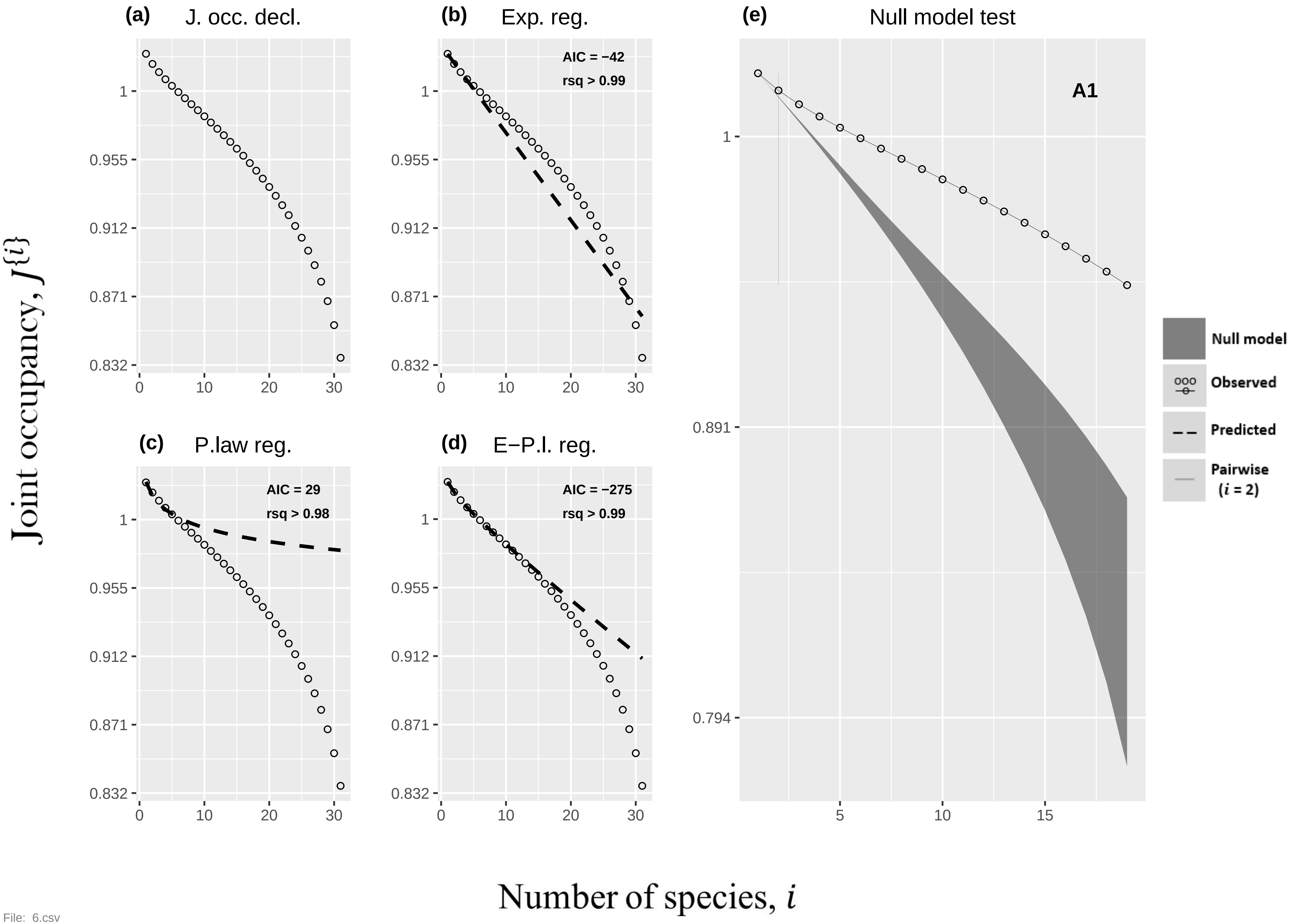

(a) J. occ. decl.
(b) Exp. reg.
AIC = −42
rsq > 0.99
(c) P.law reg.
AIC = 29
rsq > 0.98
(d) E−P.l. reg.
AIC = −275
rsq > 0.99
(e) Null model test
A1
Null model
Observed
Predicted
Pairwise (i = 2)
1
0.955
0.912
0.871
0.832
0.891
0.794
0
5
10
15
20
30
Joint occupancy, J{i}
Number of species, i


File: 6.csv

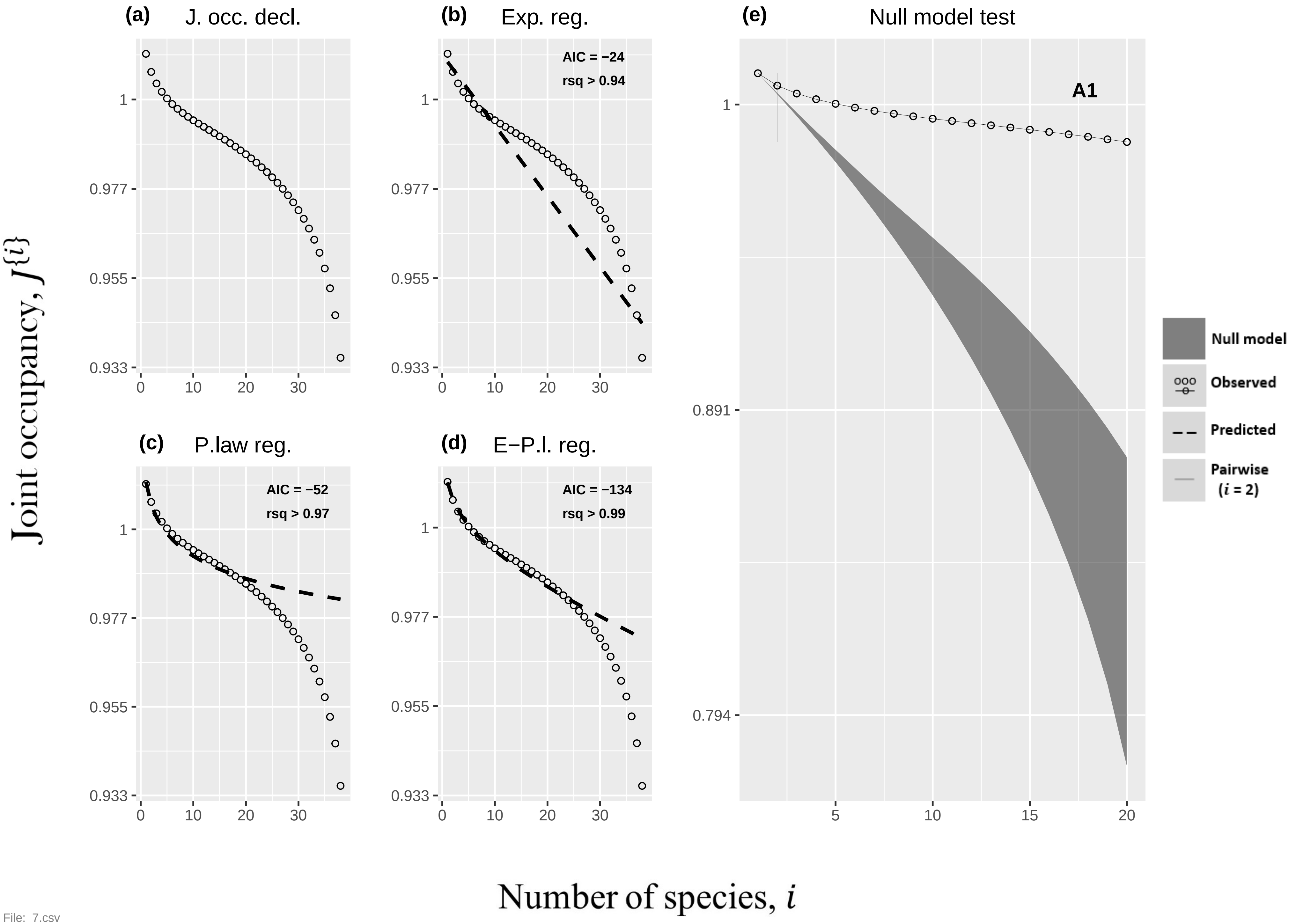
(a) J. occ. decl.
(b) Exp. reg.
AIC = −24
rsq > 0.94
(c) P.law reg.
AIC = −52
rsq > 0.97
(d) E−P.l. reg.
AIC = −134
rsq > 0.99
(e) Null model test
A1
Null model
Observed
Predicted
Pairwise (i = 2)
Joint occupancy, J{i}
Number of species, i
1
0.977
0.955
0.933
0.891
0.794
0
10
20
30
5
15

File: 7.csv

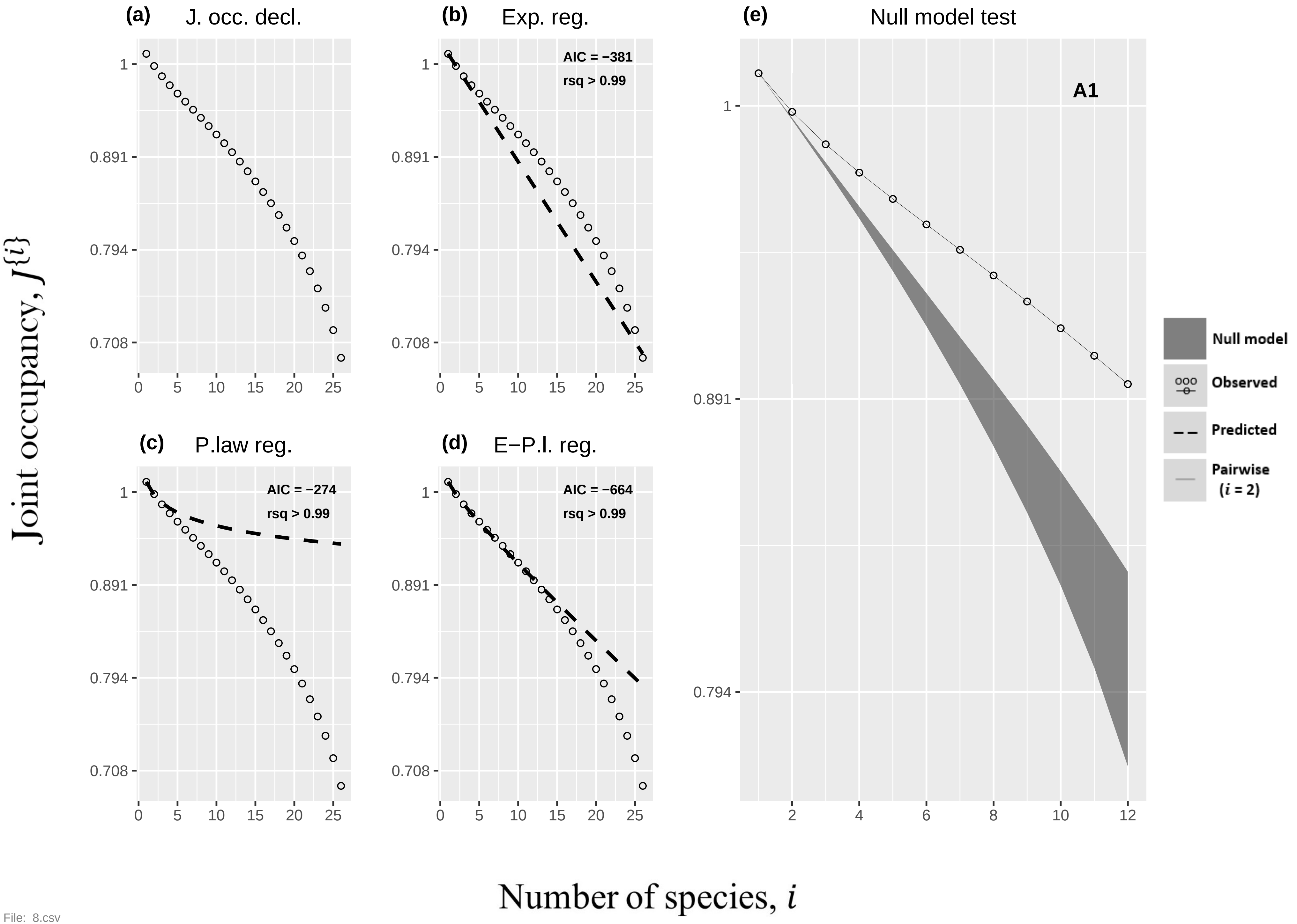
(a) J. occ. decl.
(b) Exp. reg.
AIC = −381
rsq > 0.99
(c) P.law reg.
AIC = −274
rsq > 0.99
(d) E−P.l. reg.
AIC = −664
rsq > 0.99
(e) Null model test
A1
Null model
Observed
Predicted
Pairwise ($i$ = 2)
Joint occupancy, $J\{i\}$
Number of species, $i$
1
0.891
0.794
0.708
0
5
10
15
20
25
2
4
6
8
10
12

File: 8.csv

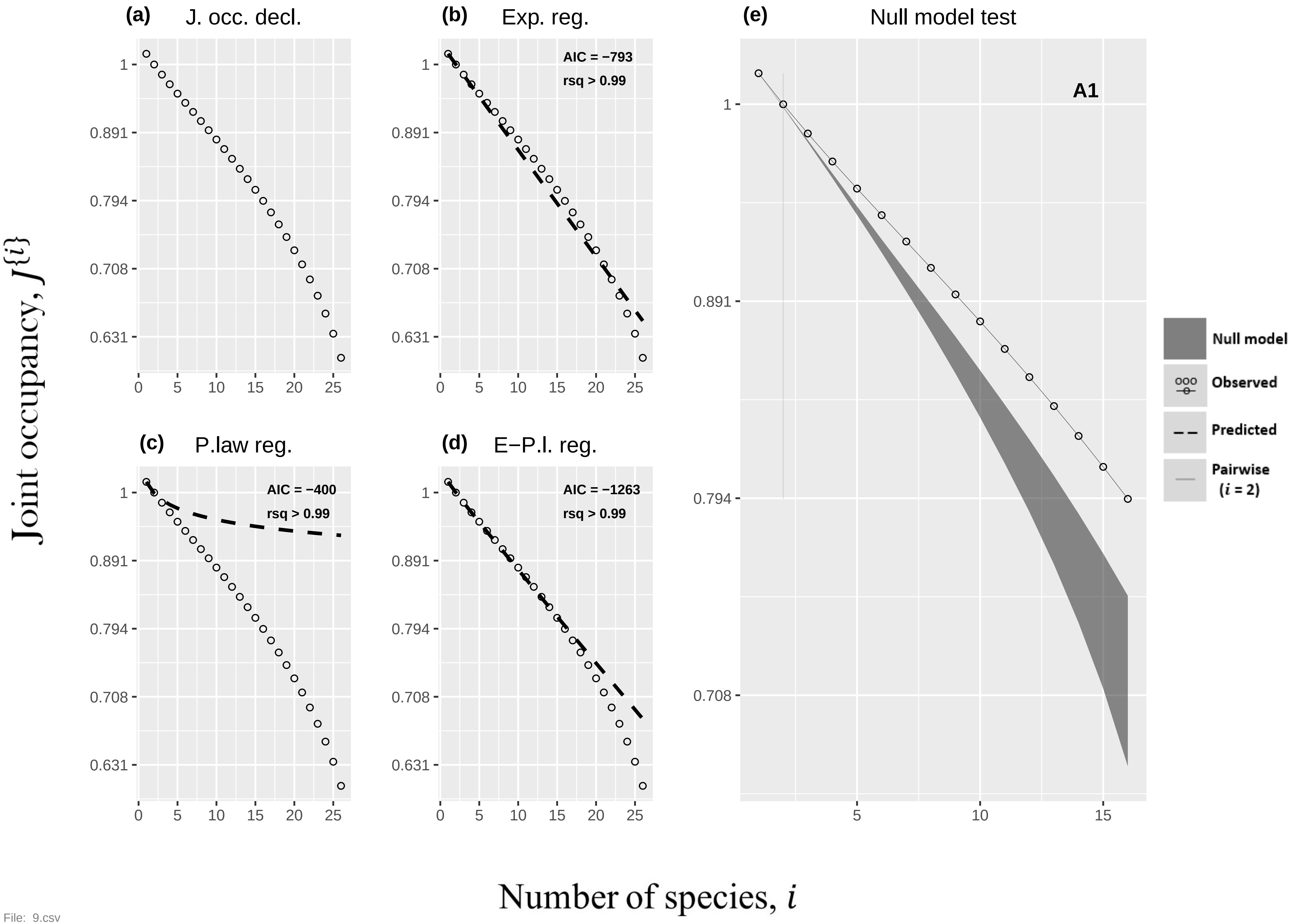

(a) J. occ. decl.
(b) Exp. reg.
AIC = −793
rsq > 0.99
(c) P.law reg.
AIC = −400
rsq > 0.99
(d) E−P.l. reg.
AIC = −1263
rsq > 0.99
(e) Null model test
A1
Null model
Observed
Predicted
Pairwise (i = 2)
Joint occupancy, J{i}
Number of species, i
1
0.891
0.794
0.708
0.631
0
5
10
15
20
25


File: 9.csv

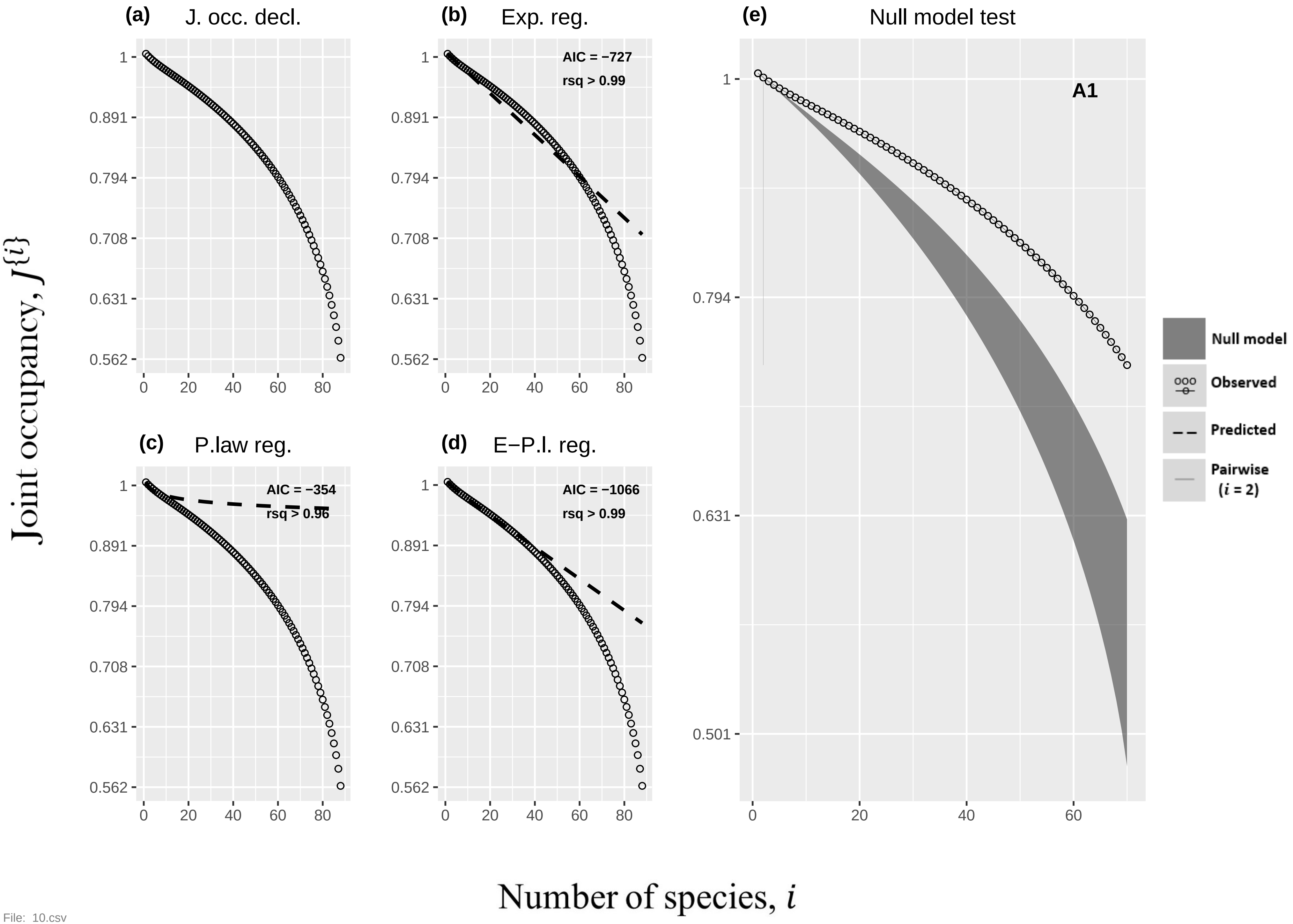

(a) J. occ. decl.
(b) Exp. reg.
AIC = −727
rsq > 0.99
(c) P.law reg.
AIC = −354
rsq > 0.96
(d) E−P.l. reg.
AIC = −1066
rsq > 0.99
(e) Null model test
A1
Null model
Observed
Predicted
Pairwise (i = 2)
Joint occupancy, J{i}
Number of species, i
1
0.891
0.794
0.708
0.631
0.562
0.501
0
20
40
60
80


File: 10.csv

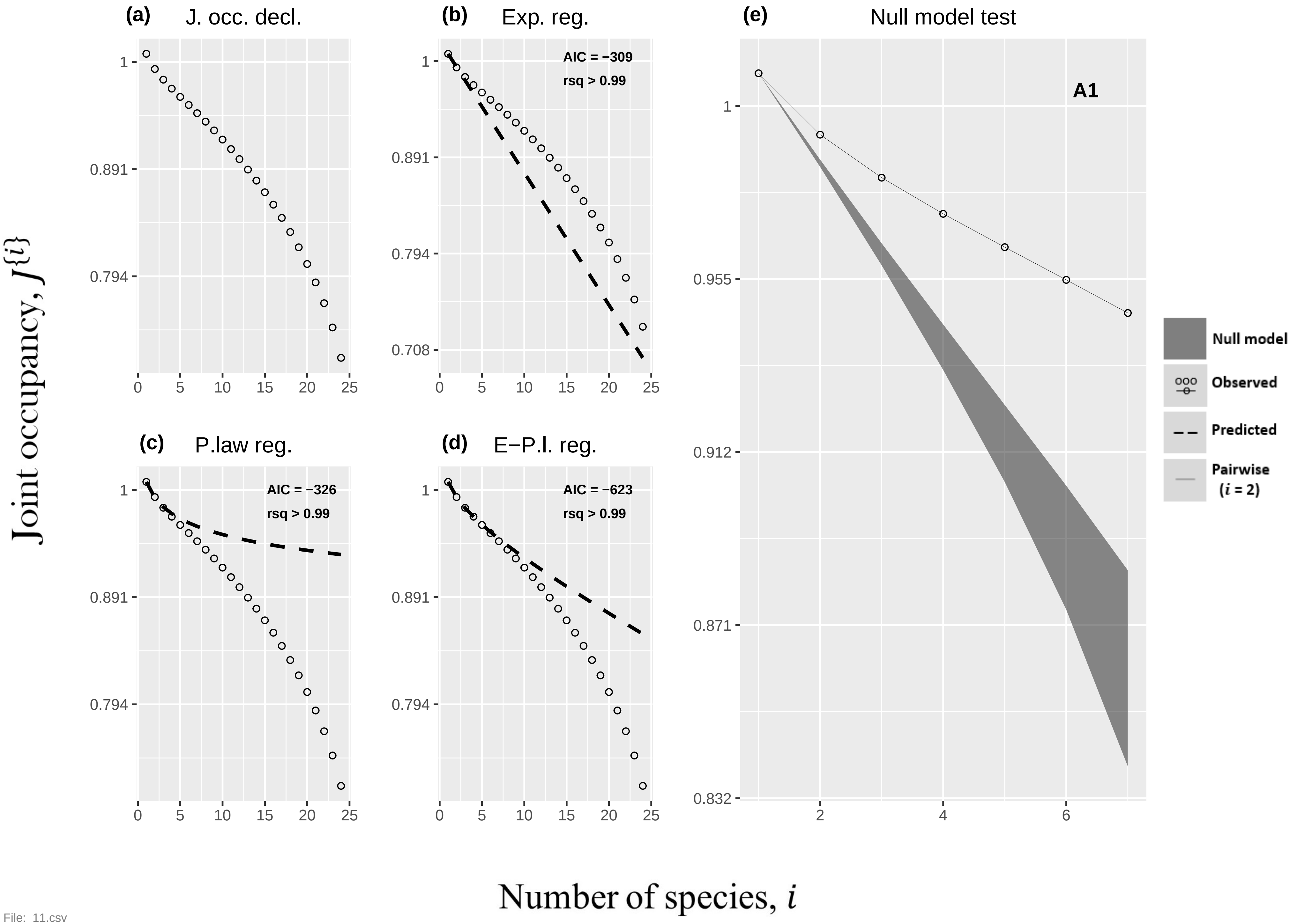
(a) J. occ. decl.
(b) Exp. reg.
AIC = −309
rsq > 0.99
(c) P.law reg.
AIC = −326
rsq > 0.99
(d) E−P.l. reg.
AIC = −623
rsq > 0.99
(e) Null model test
A1
Null model
Observed
Predicted
Pairwise (i = 2)
Joint occupancy, J{i}
Number of species, i
1
0.891
0.794
0.708
0.955
0.912
0.871
0.832
0 5 10 15 20 25
2 4 6

File: 11.csv

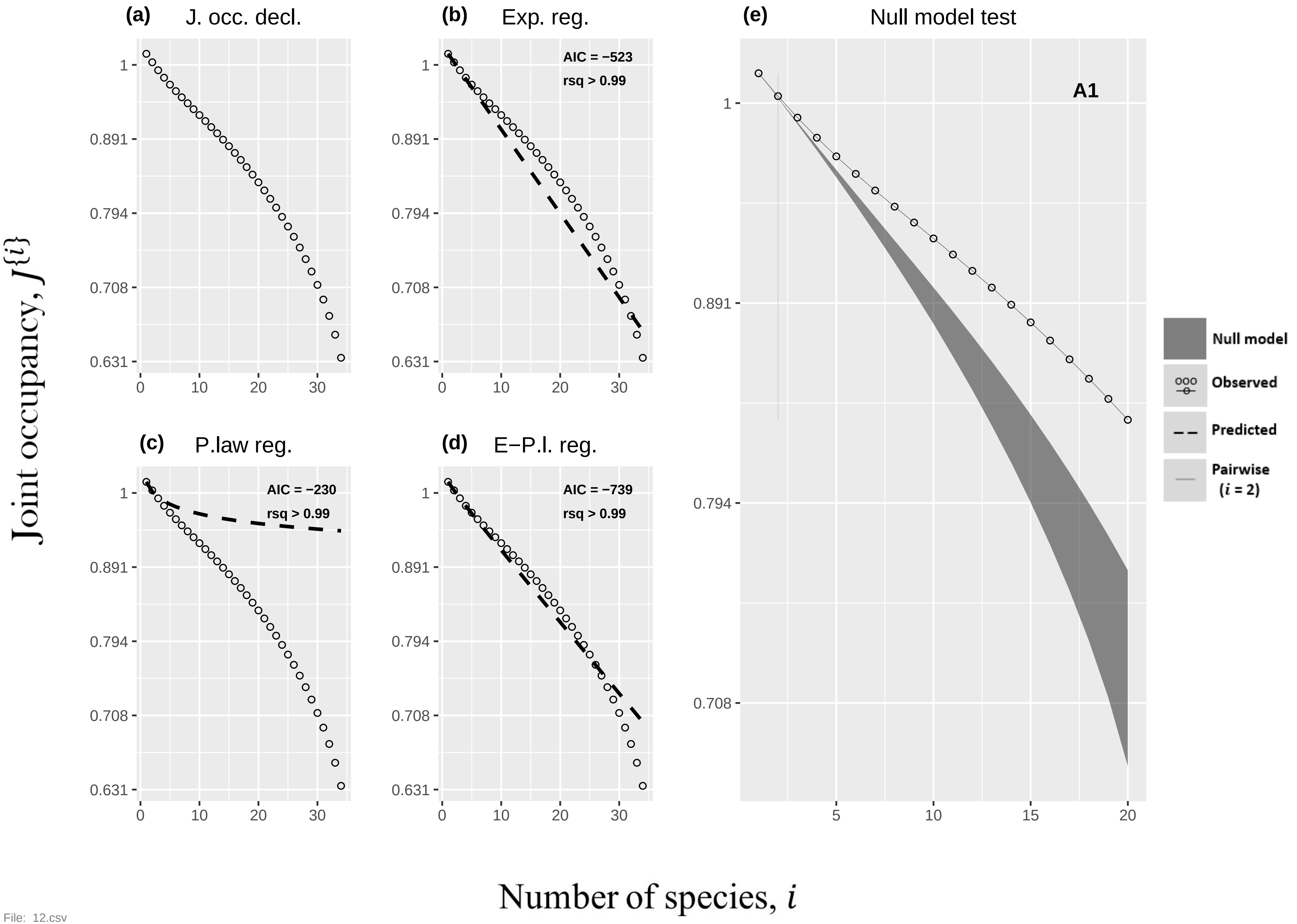
(a) J. occ. decl.
(b) Exp. reg.
AIC = −523
rsq > 0.99
(c) P.law reg.
AIC = −230
rsq > 0.99
(d) E−P.l. reg.
AIC = −739
rsq > 0.99
(e) Null model test
A1
Null model
Observed
Predicted
Pairwise (i = 2)
Joint occupancy, J{i}
Number of species, i
1
0.891
0.794
0.708
0.631
0
10
20
30
5
15

File: 12.csv

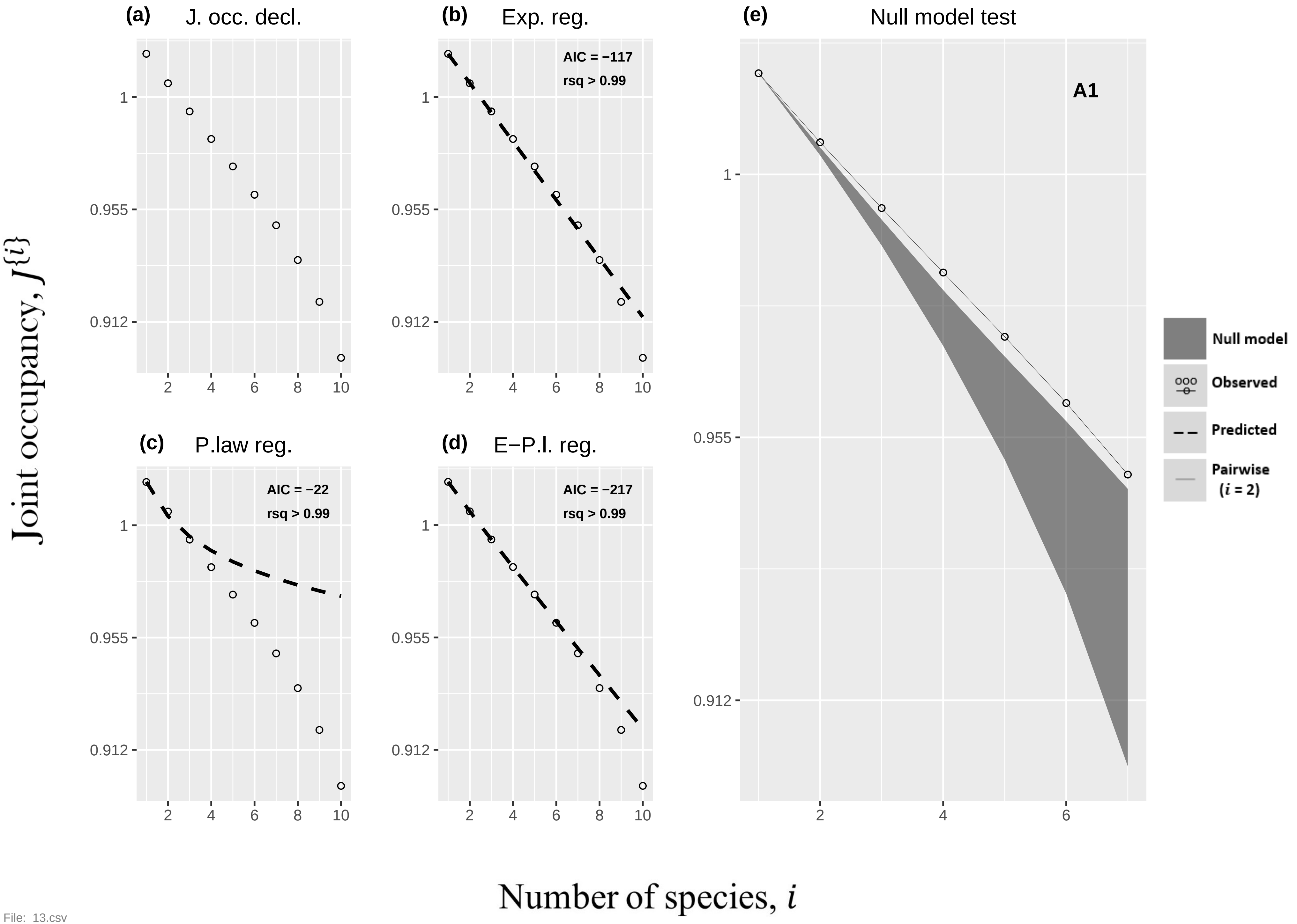

(a) J. occ. decl.
(b) Exp. reg.
AIC = −117
rsq > 0.99
(c) P.law reg.
AIC = −22
rsq > 0.99
(d) E−P.l. reg.
AIC = −217
rsq > 0.99
(e) Null model test
A1
Null model
Observed
Predicted
Pairwise
(i = 2)
Joint occupancy, J{i}
Number of species, i
1
0.955
0.912
2
4
6
8
10

File: 13.csv

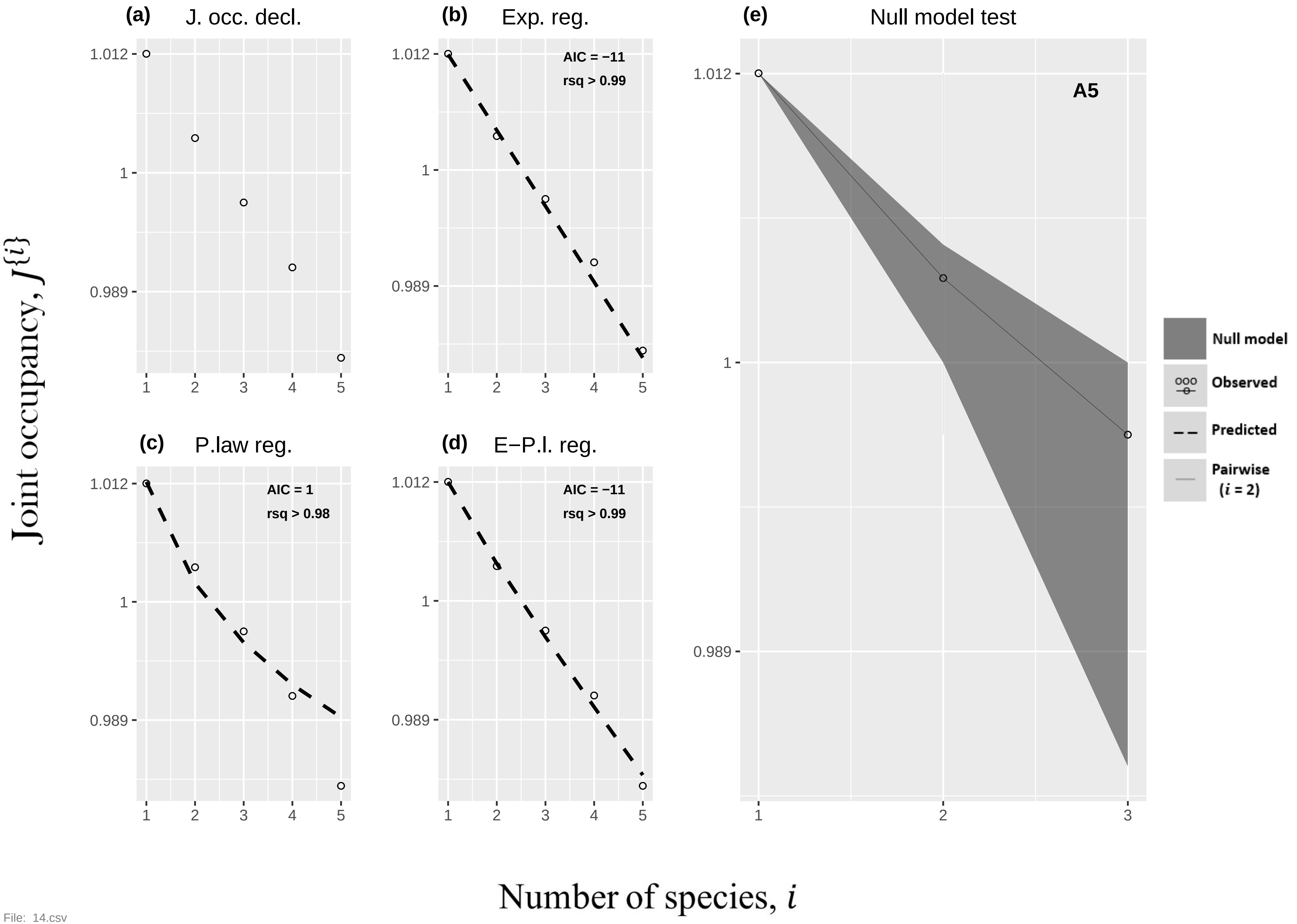

(a) J. occ. decl.
(b) Exp. reg.
AIC = −11
rsq > 0.99
(c) P.law reg.
AIC = 1
rsq > 0.98
(d) E−P.l. reg.
AIC = −11
rsq > 0.99
(e) Null model test
A5
Null model
Observed
Predicted
Pairwise (i = 2)
1.012
1
0.989
Joint occupancy, J{i}
Number of species, i


File: 14.csv

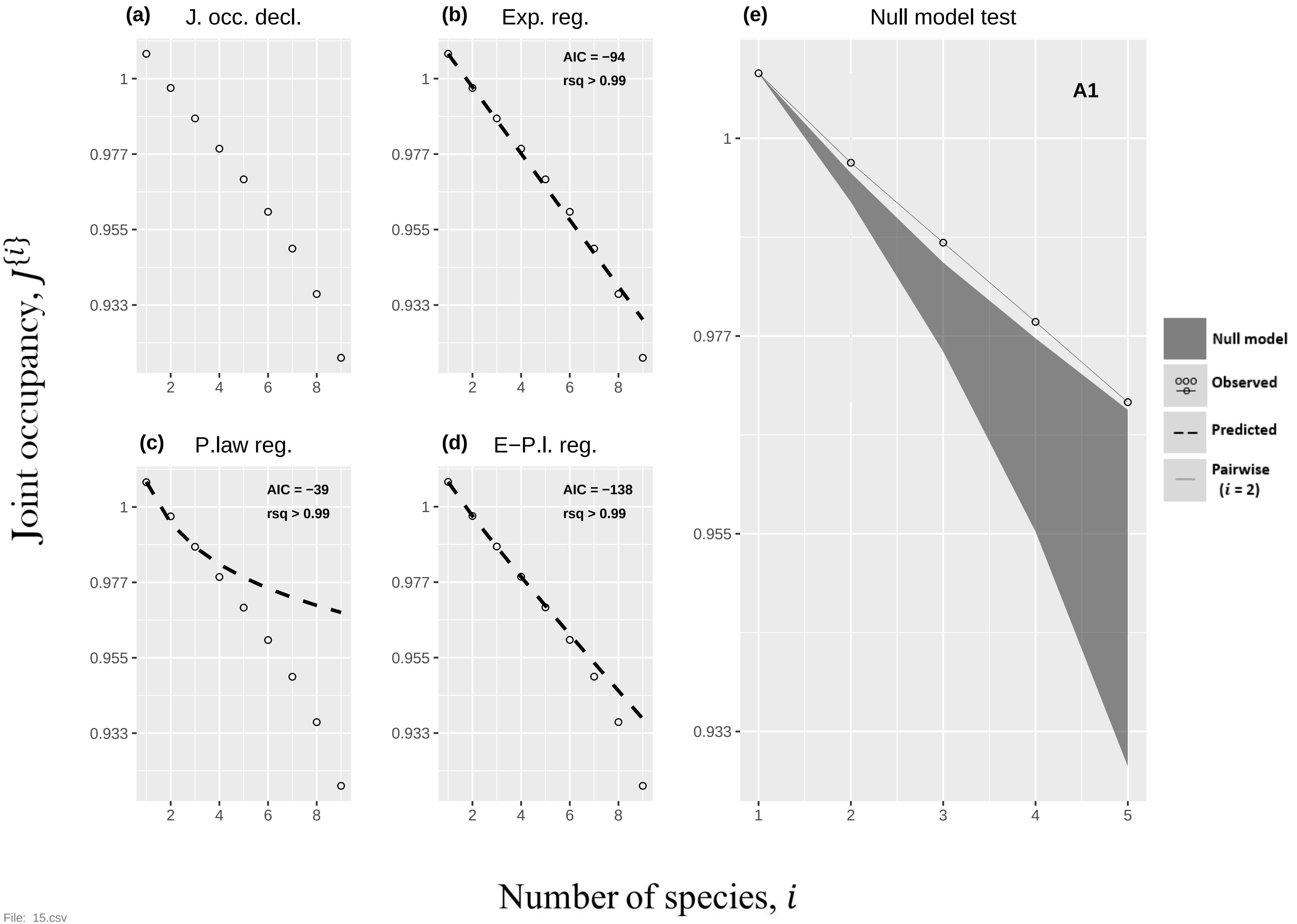

(a) J. occ. decl.
(b) Exp. reg.
AIC = −94
rsq > 0.99
(e) Null model test
A1
(c) P.law reg.
AIC = −39
rsq > 0.99
(d) E−P.l. reg.
AIC = −138
rsq > 0.99
Null model
Observed
Predicted
Pairwise (i = 2)
Joint occupancy, J{i}
Number of species, i
1
0.977
0.955
0.933
2
4
6
8
1
2
3
4
5


File: 15.csv

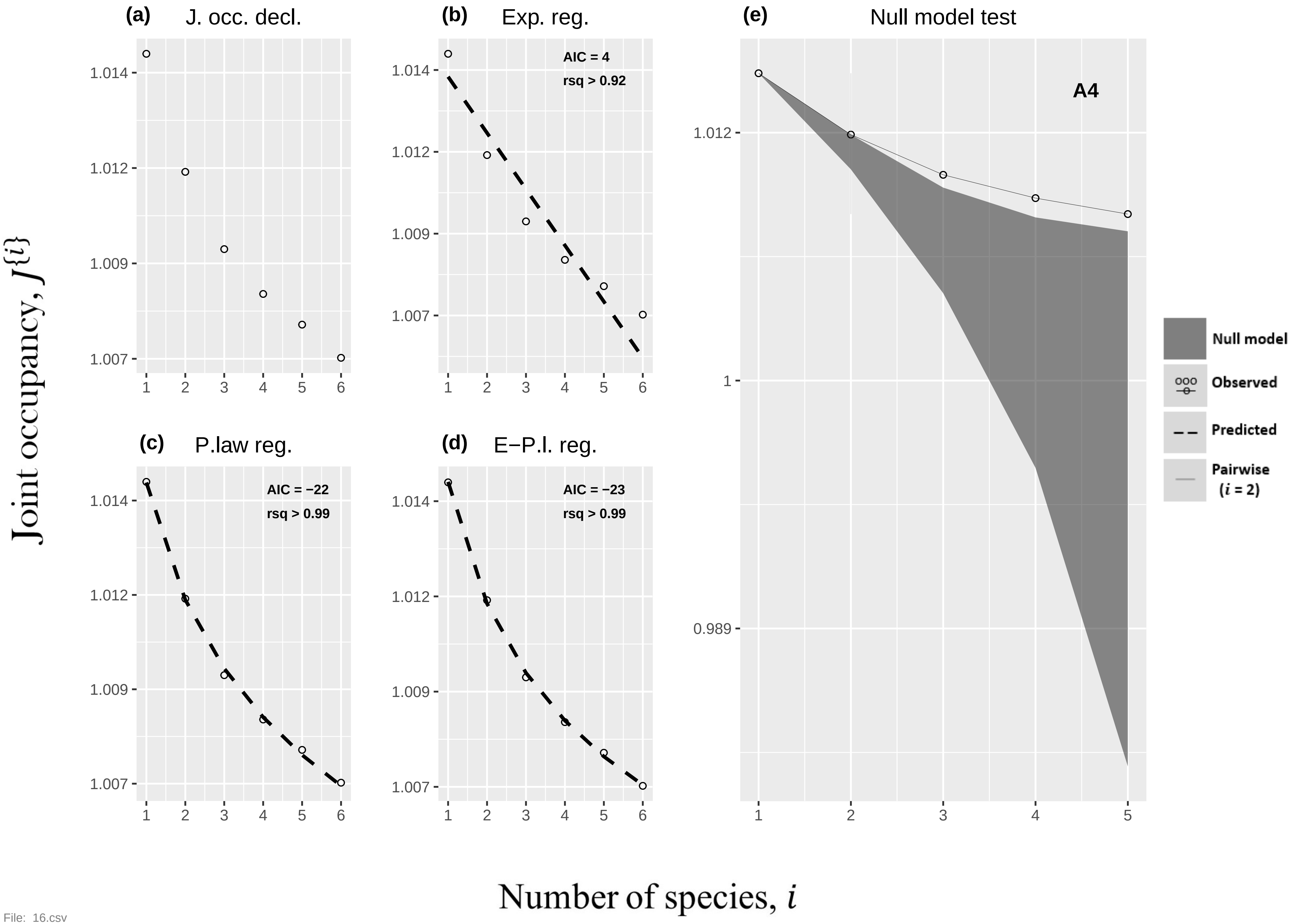

(a)
J. occ. decl.
(b)
Exp. reg.
AIC = 4
rsq > 0.92
(c)
P.law reg.
AIC = −22
rsq > 0.99
(d)
E−P.l. reg.
AIC = −23
rsq > 0.99
(e)
Null model test
A4
Null model
Observed
Predicted
Pairwise
(i = 2)
1.014
1.012
1.009
1.007
1
0.989
1 2 3 4 5 6
Joint occupancy, J{i}
Number of species, i


File: 16.csv

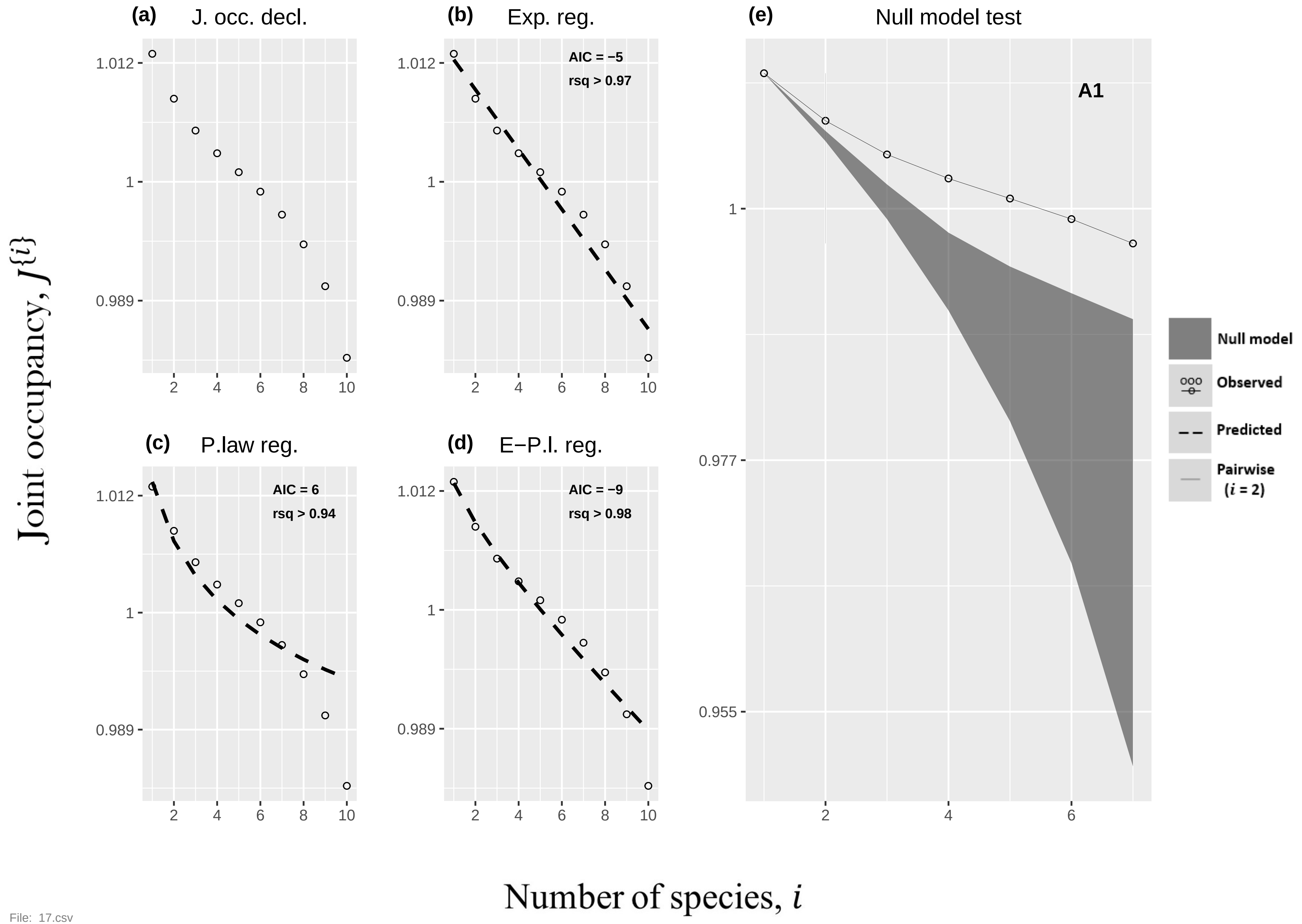
(a) J. occ. decl.
(b) Exp. reg.
AIC = −5
rsq > 0.97
(c) P.law reg.
AIC = 6
rsq > 0.94
(d) E−P.l. reg.
AIC = −9
rsq > 0.98
(e) Null model test
A1
Null model
Observed
Predicted
Pairwise (i = 2)
1.012
1
0.989
0.977
0.955
2
4
6
8
10
Joint occupancy, J{i}
Number of species, i

File: 17.csv

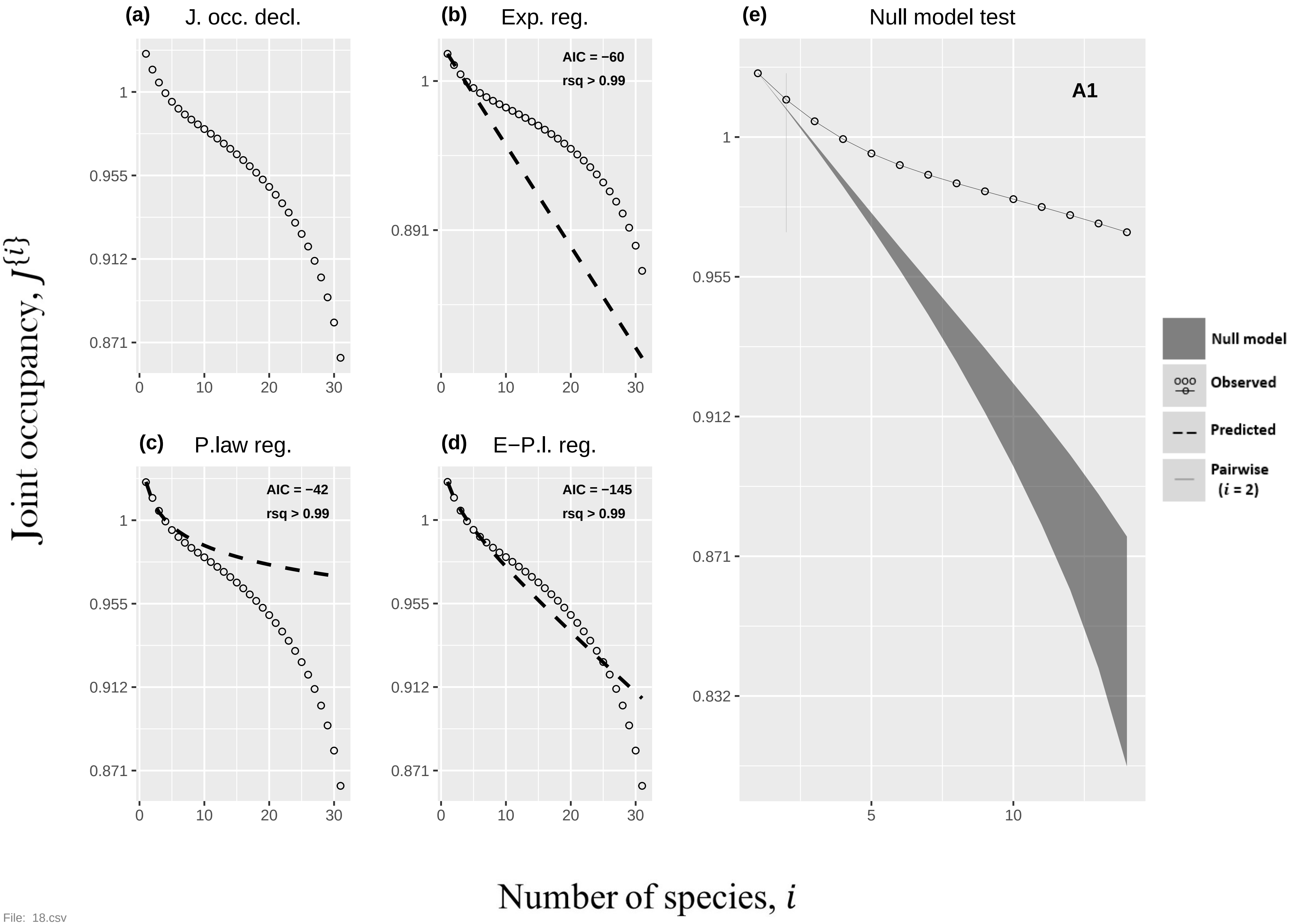
(a) J. occ. decl.
(b) Exp. reg.
AIC = −60
rsq > 0.99
(c) P.law reg.
AIC = −42
rsq > 0.99
(d) E−P.l. reg.
AIC = −145
rsq > 0.99
(e) Null model test
A1
Null model
Observed
Predicted
Pairwise (i = 2)
Joint occupancy, J{i}
Number of species, i

File: 18.csv

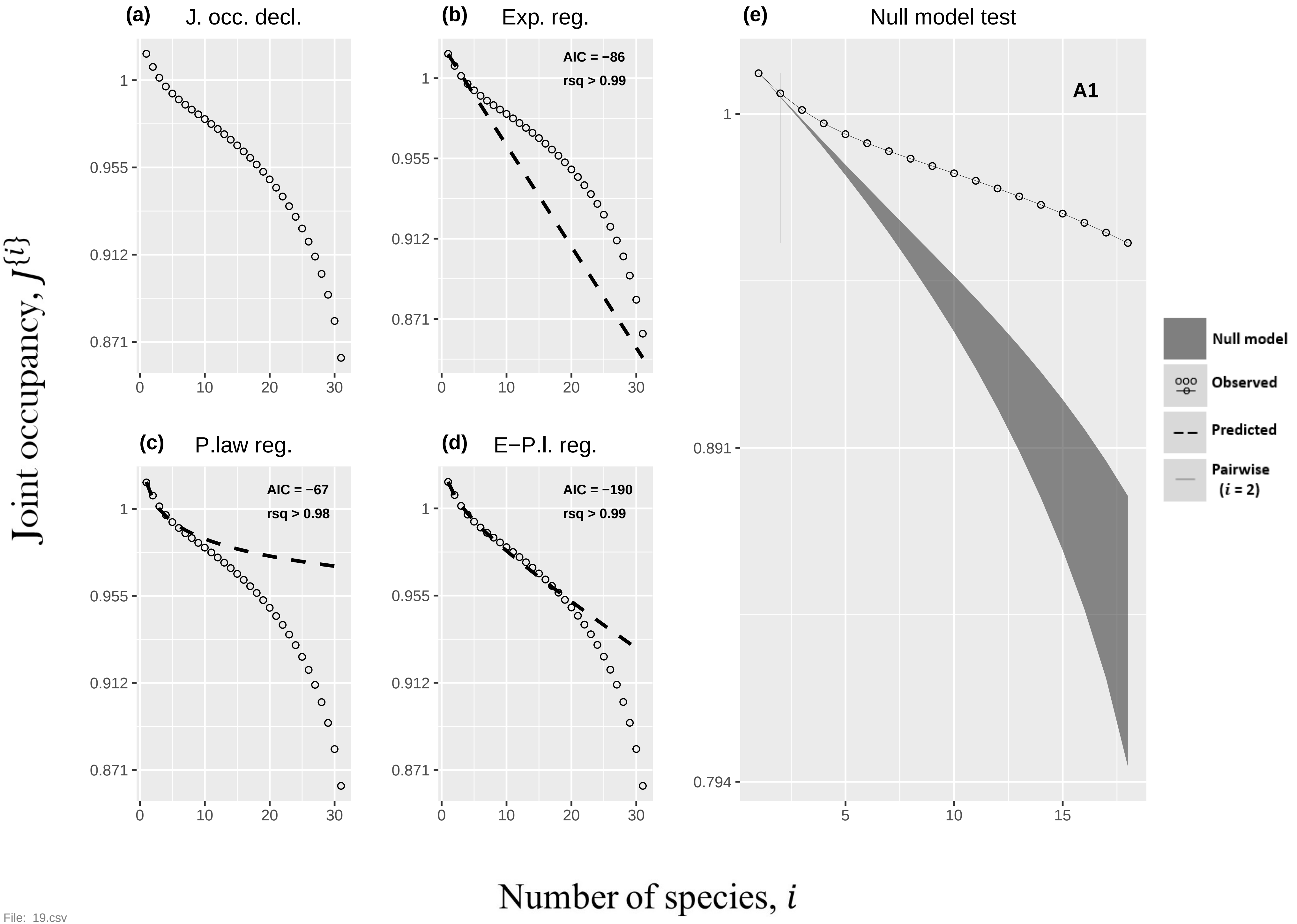

(a) J. occ. decl.
(b) Exp. reg.
AIC = −86
rsq > 0.99
(c) P.law reg.
AIC = −67
rsq > 0.98
(d) E−P.l. reg.
AIC = −190
rsq > 0.99
(e) Null model test
A1
Null model
Observed
Predicted
Pairwise (i = 2)
Joint occupancy, J{i}
Number of species, i


File: 19.csv

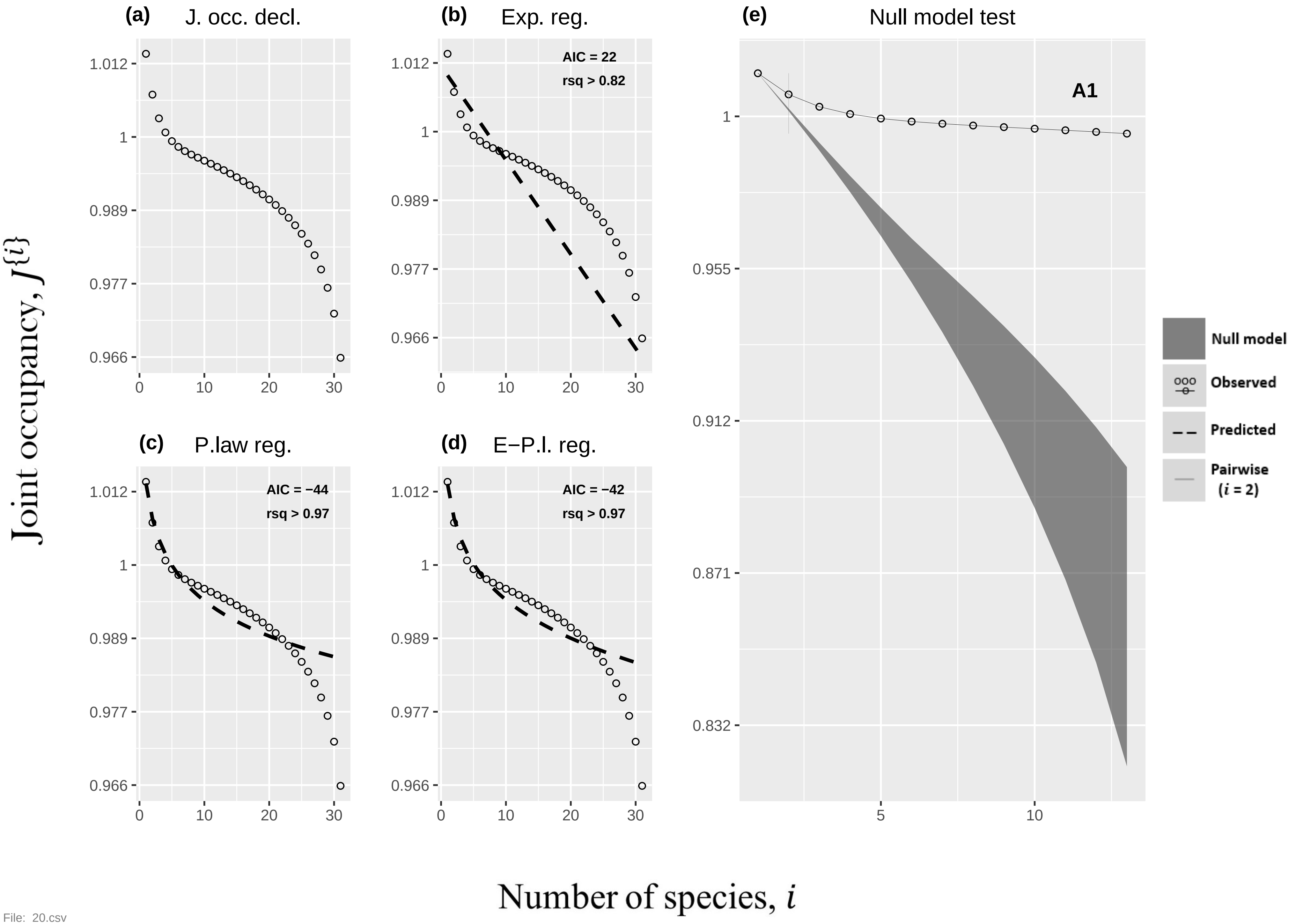
(a)
J. occ. decl.
(b)
Exp. reg.
AIC = 22
rsq > 0.82
(c)
P.law reg.
AIC = −44
rsq > 0.97
(d)
E−P.l. reg.
AIC = −42
rsq > 0.97
(e)
Null model test
A1
Null model
Observed
Predicted
Pairwise
(i = 2)
Joint occupancy, J{i}
Number of species, i
1.012
1
0.989
0.977
0.966
0
10
20
30
0.955
0.912
0.871
0.832
5
10

File: 20.csv

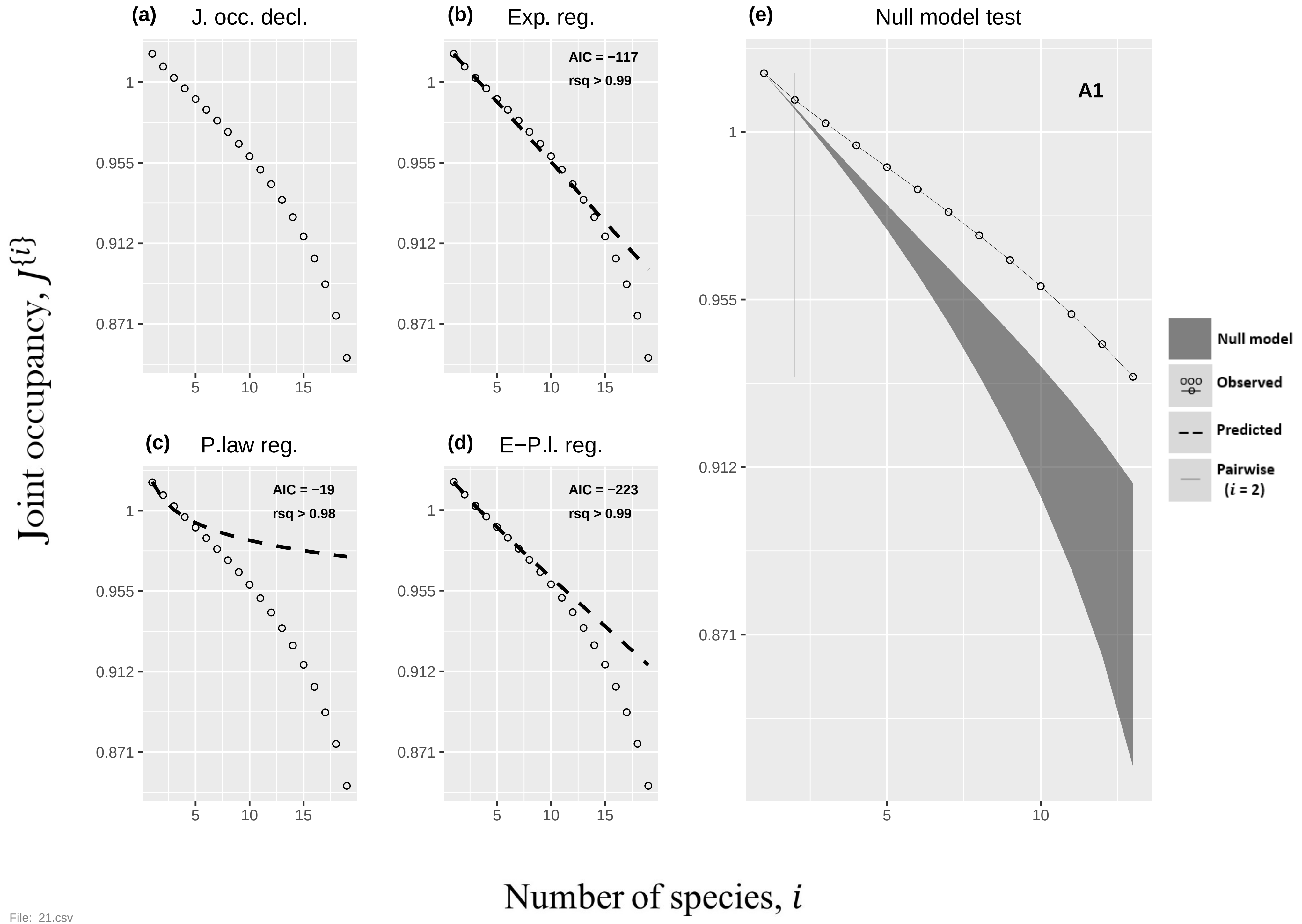

(a) J. occ. decl.
(b) Exp. reg.
AIC = −117
rsq > 0.99
(c) P.law reg.
AIC = −19
rsq > 0.98
(d) E−P.l. reg.
AIC = −223
rsq > 0.99
(e) Null model test
A1
Null model
Observed
Predicted
Pairwise
($i$ = 2)
1
0.955
0.912
0.871
5
10
15
Joint occupancy, $J\{i\}$
Number of species, $i$


File: 21.csv

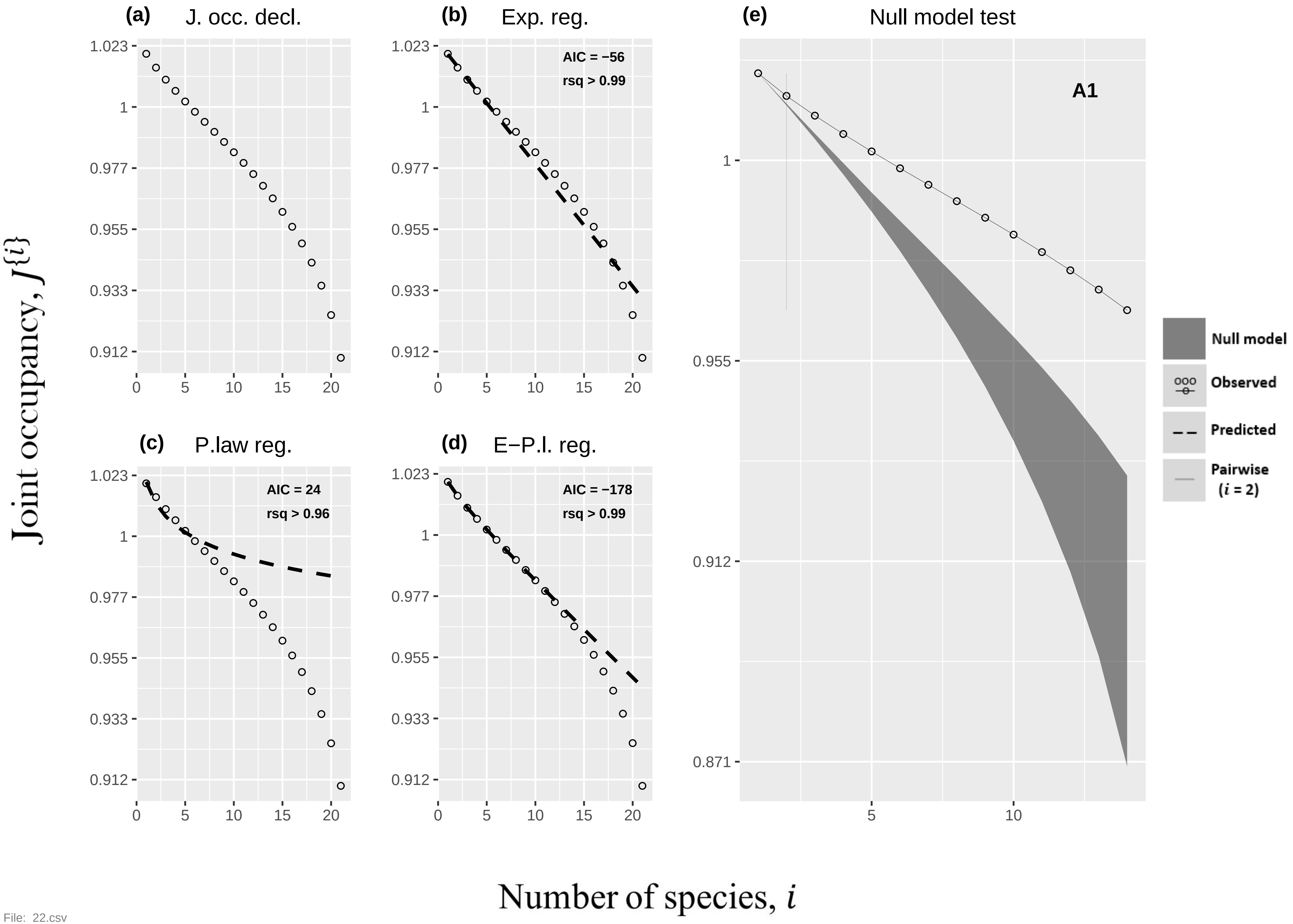

(a) J. occ. decl.
(b) Exp. reg.
AIC = −56
rsq > 0.99
(c) P.law reg.
AIC = 24
rsq > 0.96
(d) E−P.l. reg.
AIC = −178
rsq > 0.99
(e) Null model test
A1
Null model
Observed
Predicted
Pairwise ($i$ = 2)
Joint occupancy, $J^{\{i\}}$
Number of species, $i$
1.023
1
0.977
0.955
0.933
0.912
0.871
0
5
10
15
20


File: 22.csv

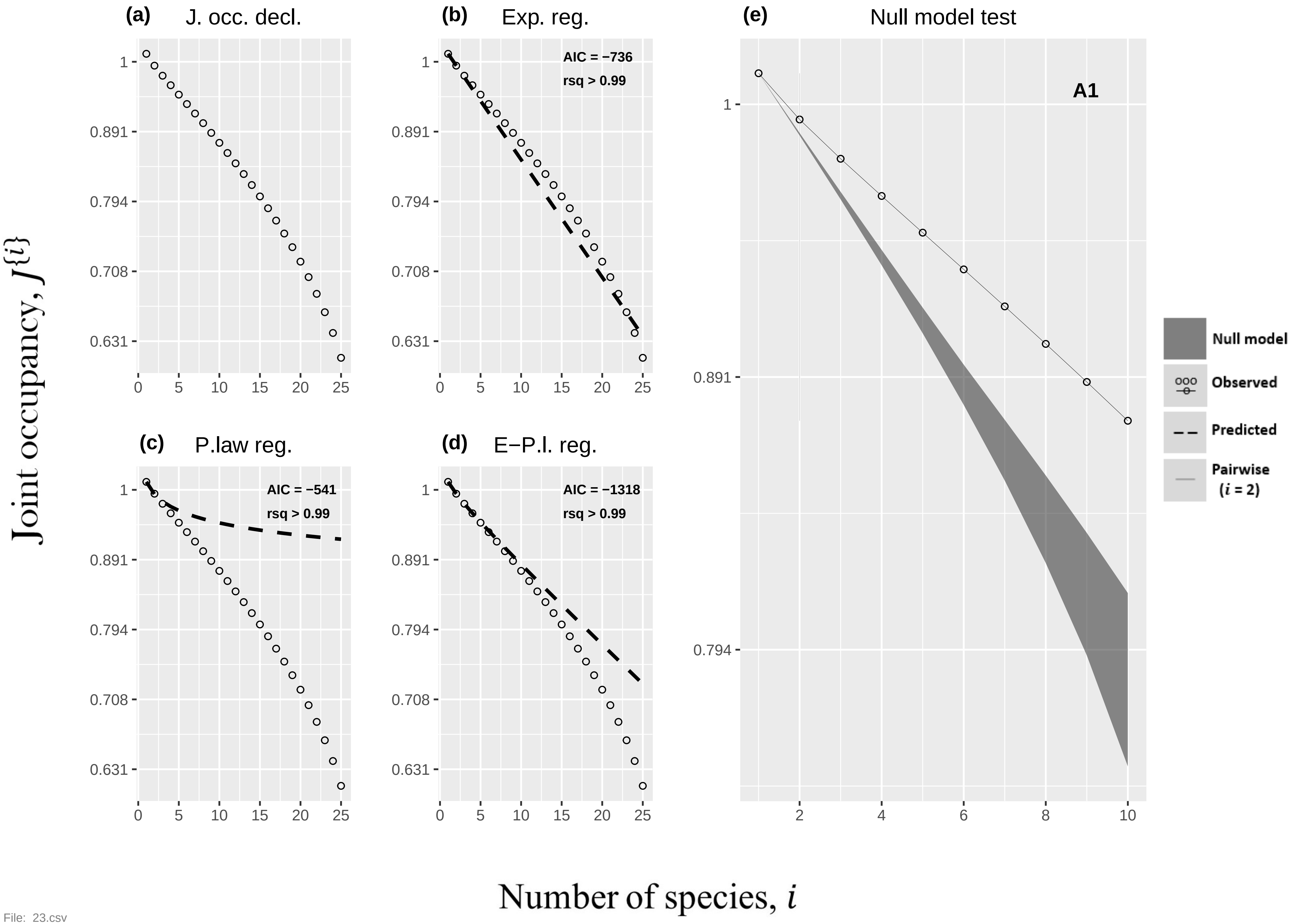

(a) J. occ. decl.
(b) Exp. reg.
AIC = −736
rsq > 0.99
(c) P.law reg.
AIC = −541
rsq > 0.99
(d) E−P.l. reg.
AIC = −1318
rsq > 0.99
(e) Null model test
A1
Null model
Observed
Predicted
Pairwise (i = 2)
Joint occupancy, J{i}
Number of species, i
1
0.891
0.794
0.708
0.631
0 5 10 15 20 25
2 4 6 8 10

File: 23.csv

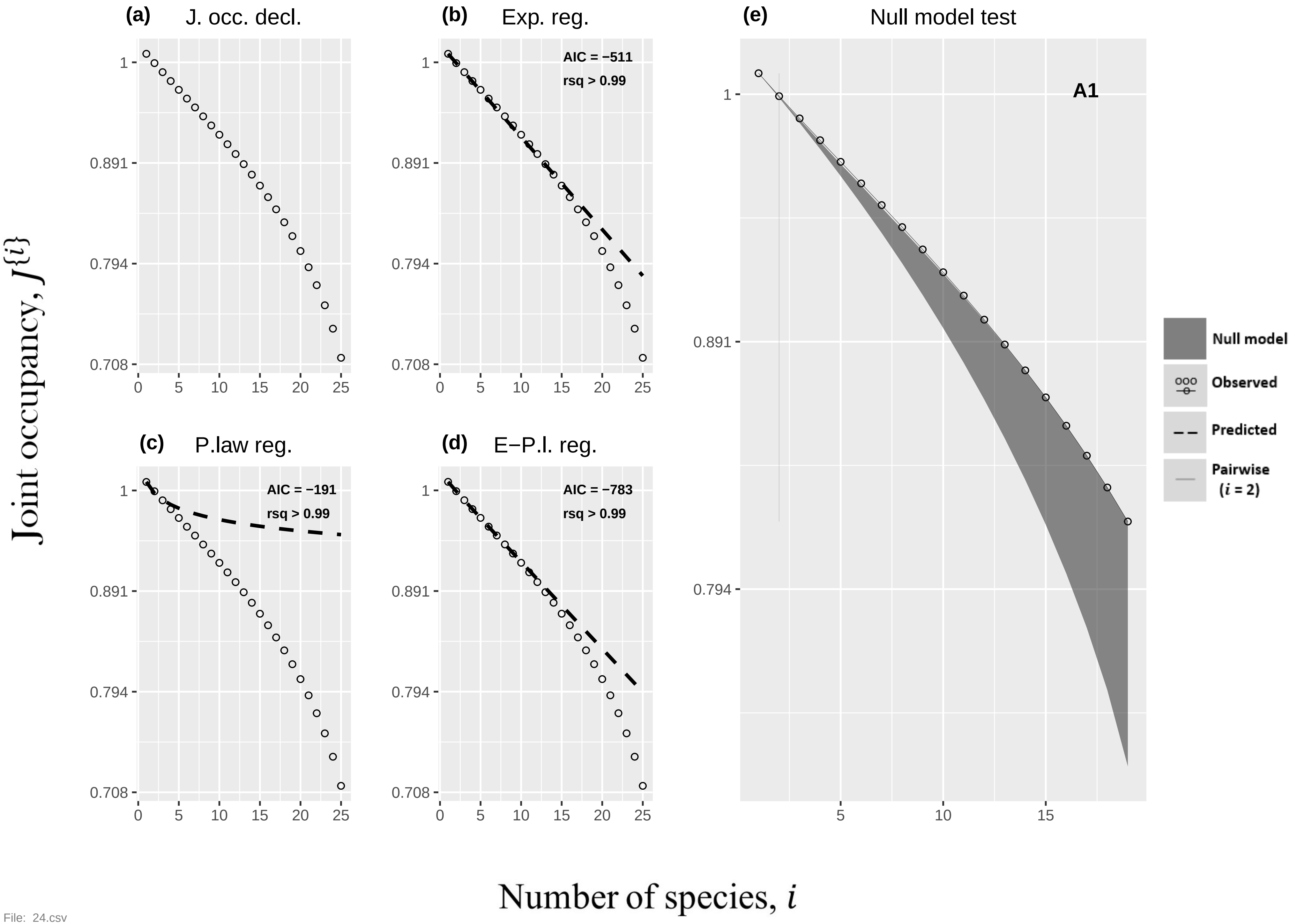
(a) J. occ. decl.
(b) Exp. reg.
AIC = −511
rsq > 0.99
(c) P.law reg.
AIC = −191
rsq > 0.99
(d) E−P.l. reg.
AIC = −783
rsq > 0.99
(e) Null model test
A1
Null model
Observed
Predicted
Pairwise (i = 2)
Joint occupancy, J{i}
Number of species, i
1
0.891
0.794
0.708
0
5
10
15
20
25

File: 24.csv

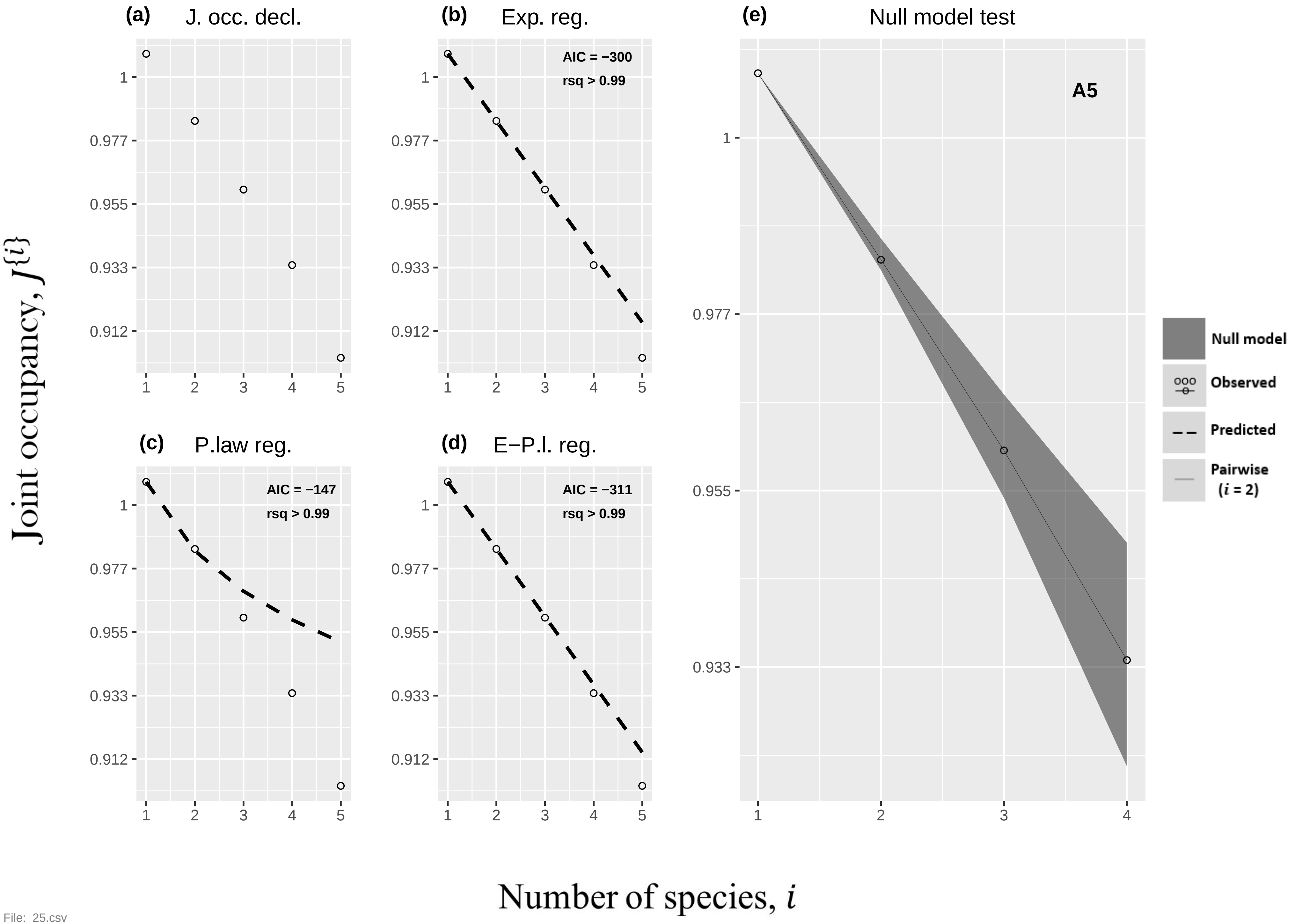
(a) J. occ. decl.
(b) Exp. reg.
AIC = −300
rsq > 0.99
(c) P.law reg.
AIC = −147
rsq > 0.99
(d) E−P.l. reg.
AIC = −311
rsq > 0.99
(e) Null model test
A5
Null model
Observed
Predicted
Pairwise
(i = 2)
Joint occupancy, J{i}
Number of species, i
1
0.977
0.955
0.933
0.912
1 2 3 4 5


File: 25.csv

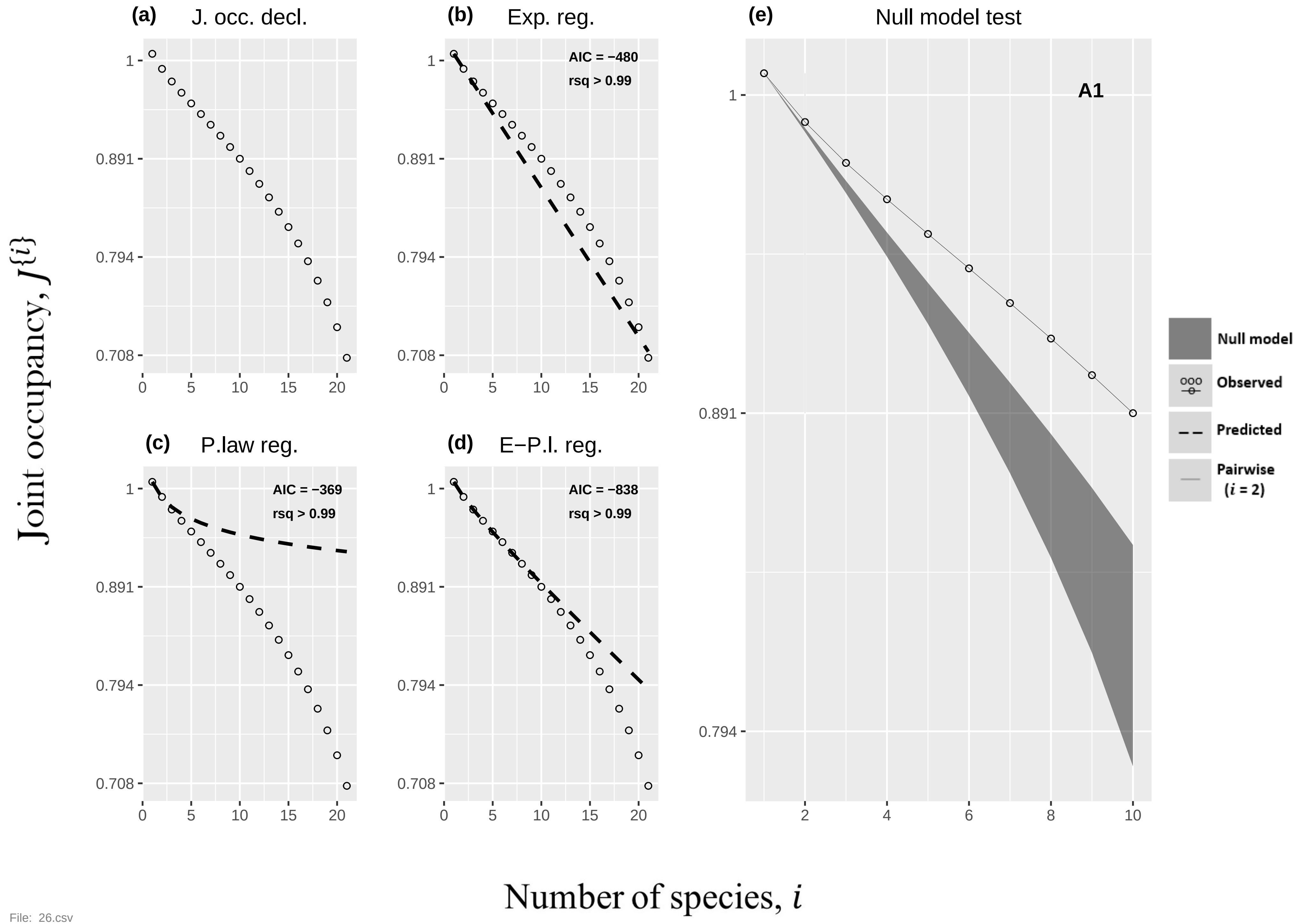

(a) J. occ. decl.
(b) Exp. reg.
AIC = −480
rsq > 0.99
(c) P.law reg.
AIC = −369
rsq > 0.99
(d) E−P.l. reg.
AIC = −838
rsq > 0.99
(e) Null model test
A1
Null model
Observed
Predicted
Pairwise ($i$ = 2)
1
0.891
0.794
0.708
0 5 10 15 20
2 4 6 8 10
Joint occupancy, $J\{i\}$
Number of species, $i$


File: 26.csv

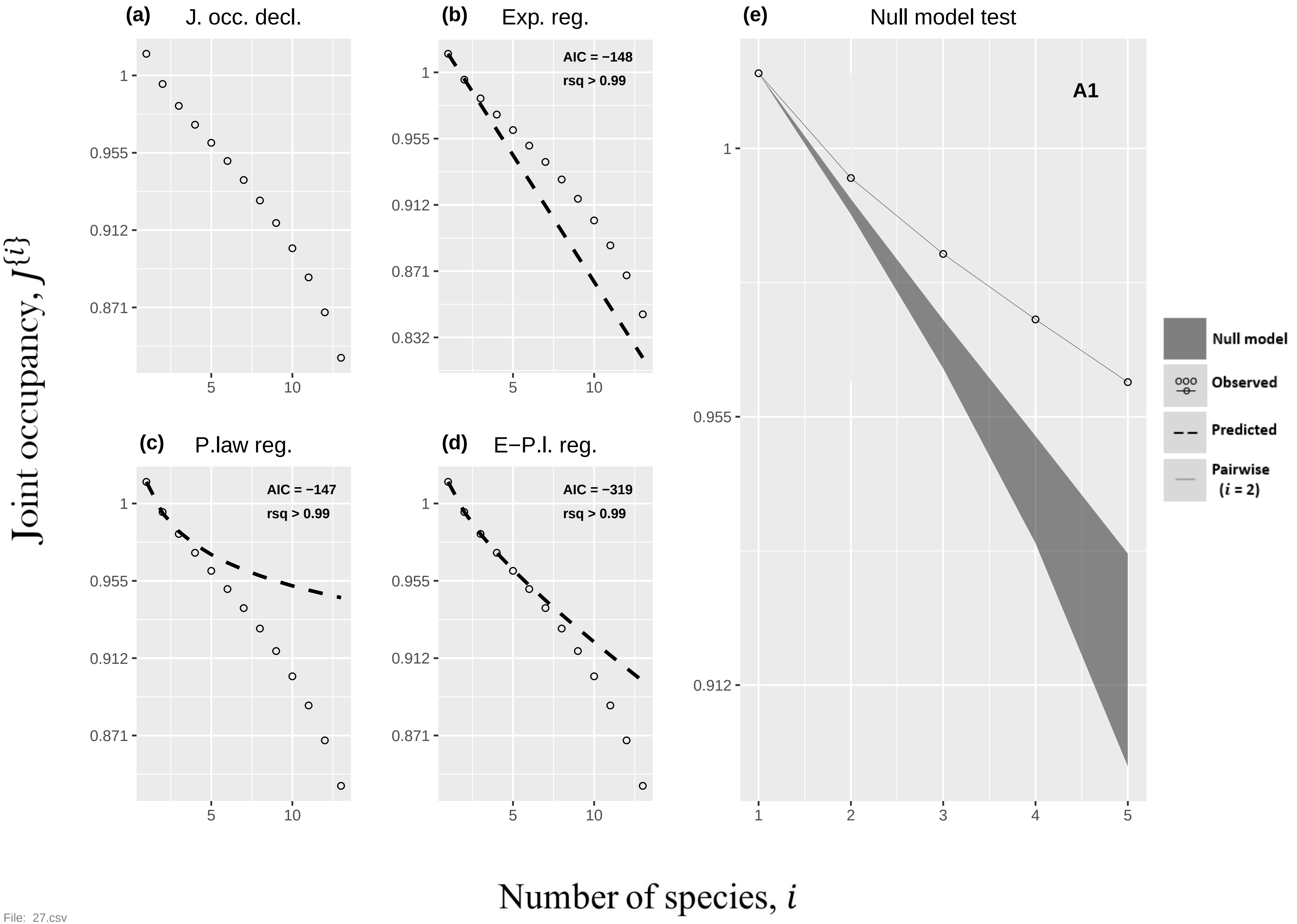

(a) J. occ. decl.
(b) Exp. reg.
AIC = −148
rsq > 0.99
(c) P.law reg.
AIC = −147
rsq > 0.99
(d) E−P.l. reg.
AIC = −319
rsq > 0.99
(e) Null model test
A1
Null model
Observed
Predicted
Pairwise
(i = 2)
1
0.955
0.912
0.871
0.832
5
10
1
2
3
4
5
Joint occupancy, J{i}
Number of species, i


File: 27.csv

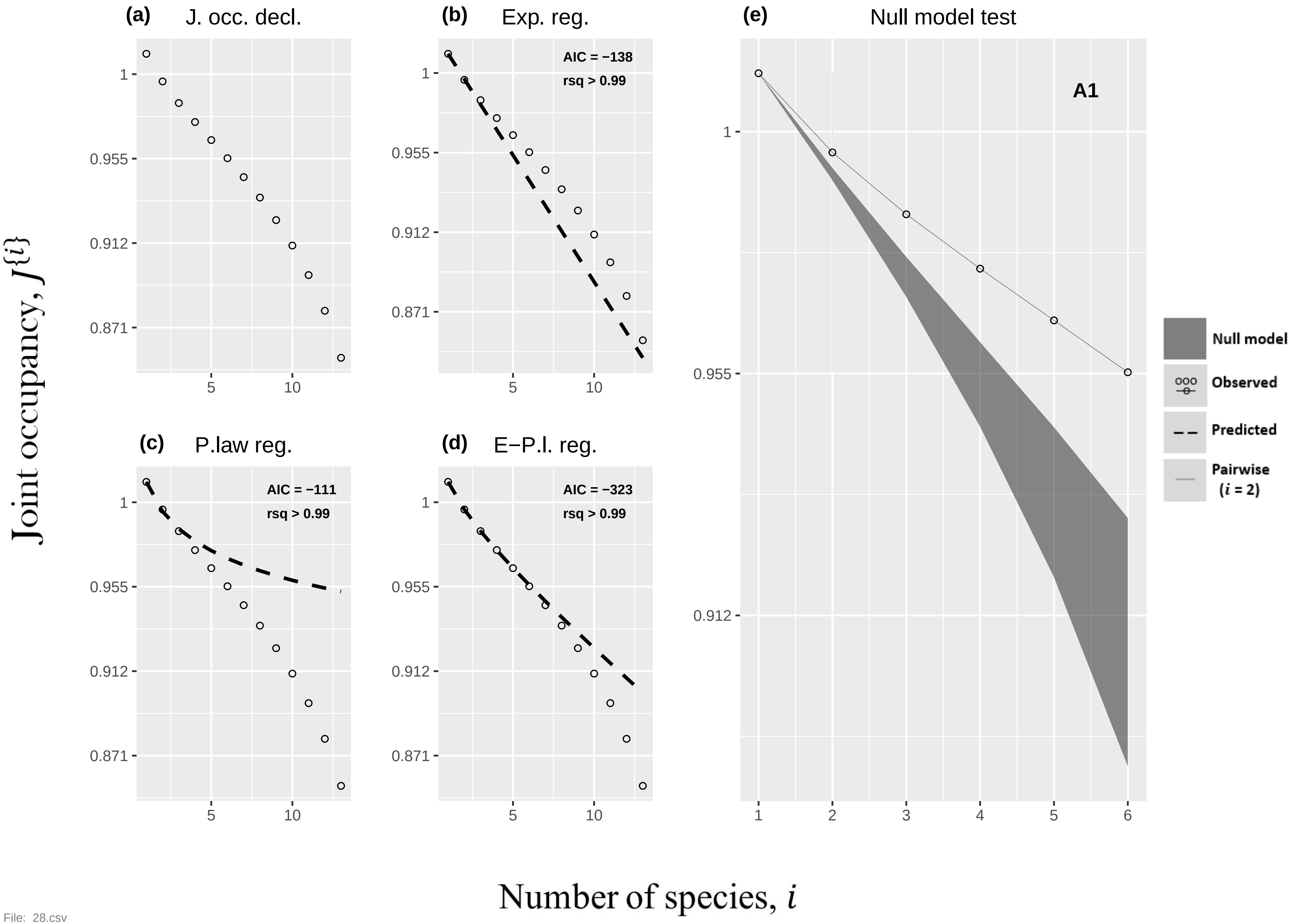

(a) J. occ. decl.
(b) Exp. reg.
AIC = −138
rsq > 0.99
(c) P.law reg.
AIC = −111
rsq > 0.99
(d) E−P.l. reg.
AIC = −323
rsq > 0.99
(e) Null model test
A1
Null model
Observed
Predicted
Pairwise
(i = 2)
1
0.955
0.912
0.871
5
10
1
2
3
4
5
6
Joint occupancy, J{i}
Number of species, i


File: 28.csv

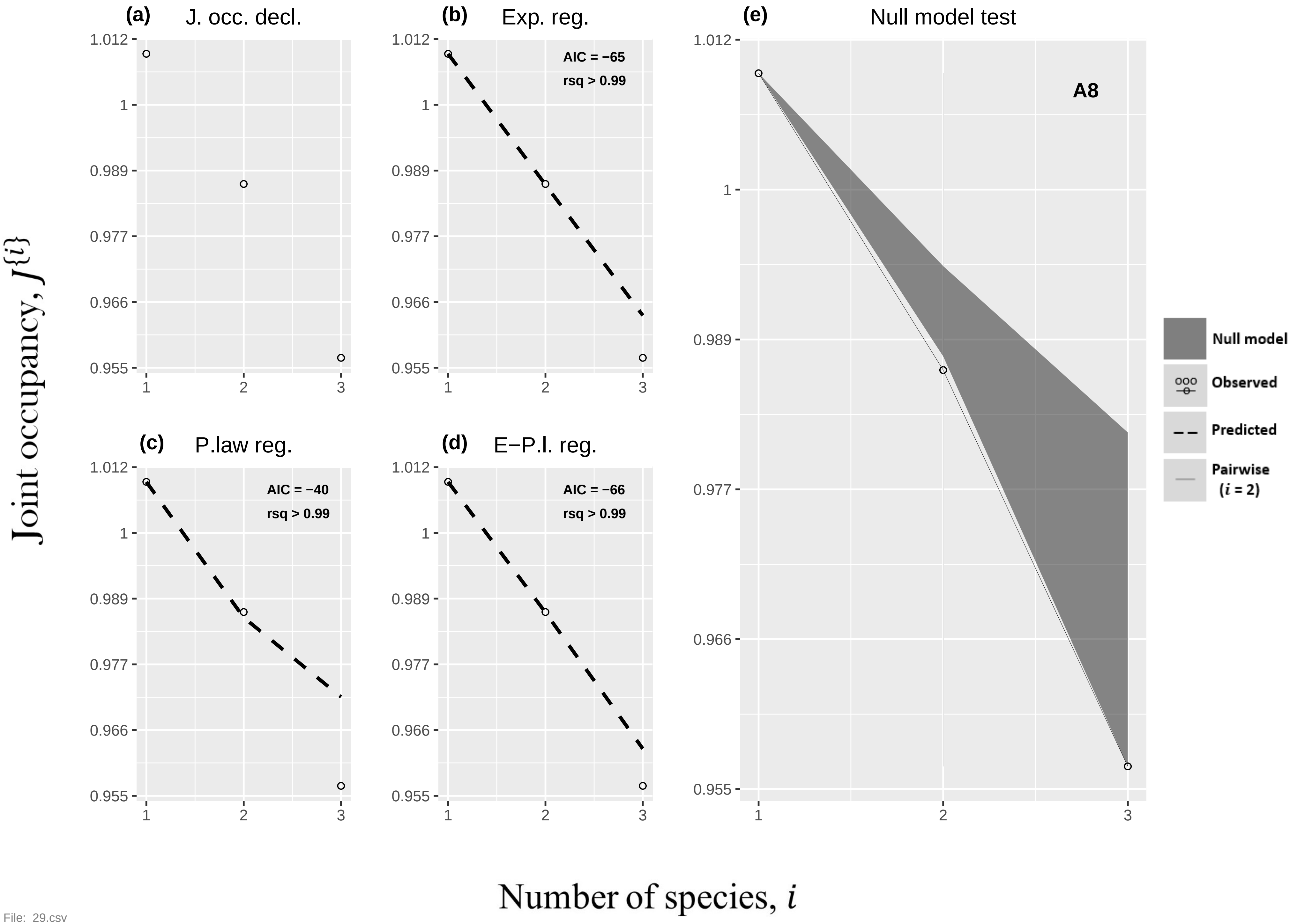

(a) J. occ. decl.
(b) Exp. reg.
AIC = −65
rsq > 0.99
(c) P.law reg.
AIC = −40
rsq > 0.99
(d) E−P.l. reg.
AIC = −66
rsq > 0.99
(e) Null model test
A8
Null model
Observed
Predicted
Pairwise (i = 2)
1.012
1
0.989
0.977
0.966
0.955
1
2
3
Joint occupancy, J{i}
Number of species, i


File: 29.csv

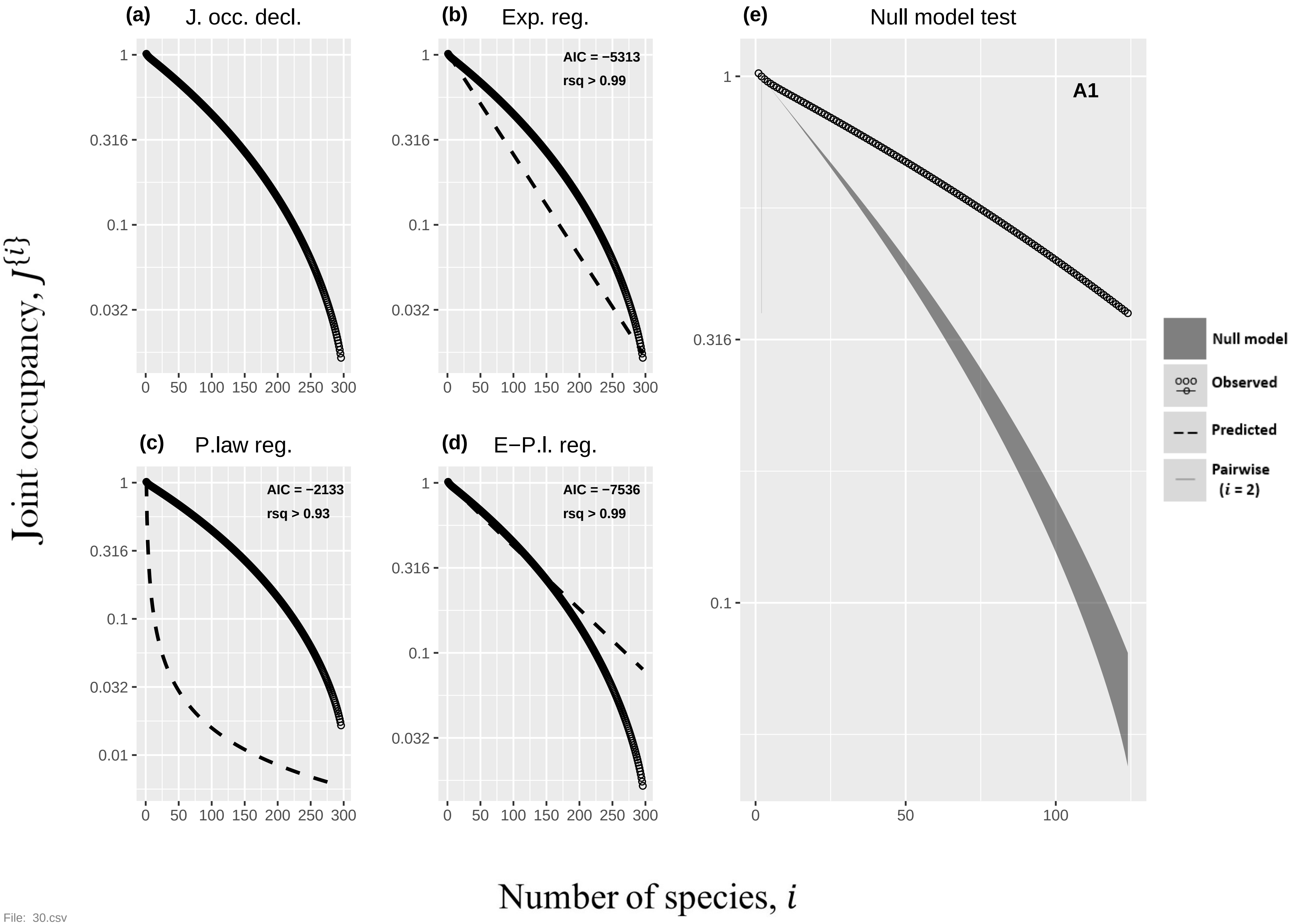

(a) J. occ. decl.
(b) Exp. reg.
AIC = −5313
rsq > 0.99
(c) P.law reg.
AIC = −2133
rsq > 0.93
(d) E−P.l. reg.
AIC = −7536
rsq > 0.99
(e) Null model test
A1
Null model
Observed
Predicted
Pairwise ($i$ = 2)
Joint occupancy, $J\{i\}$
Number of species, $i$
1
0.316
0.1
0.032
0.01
0
50
100
150
200
250
300


File: 30.csv

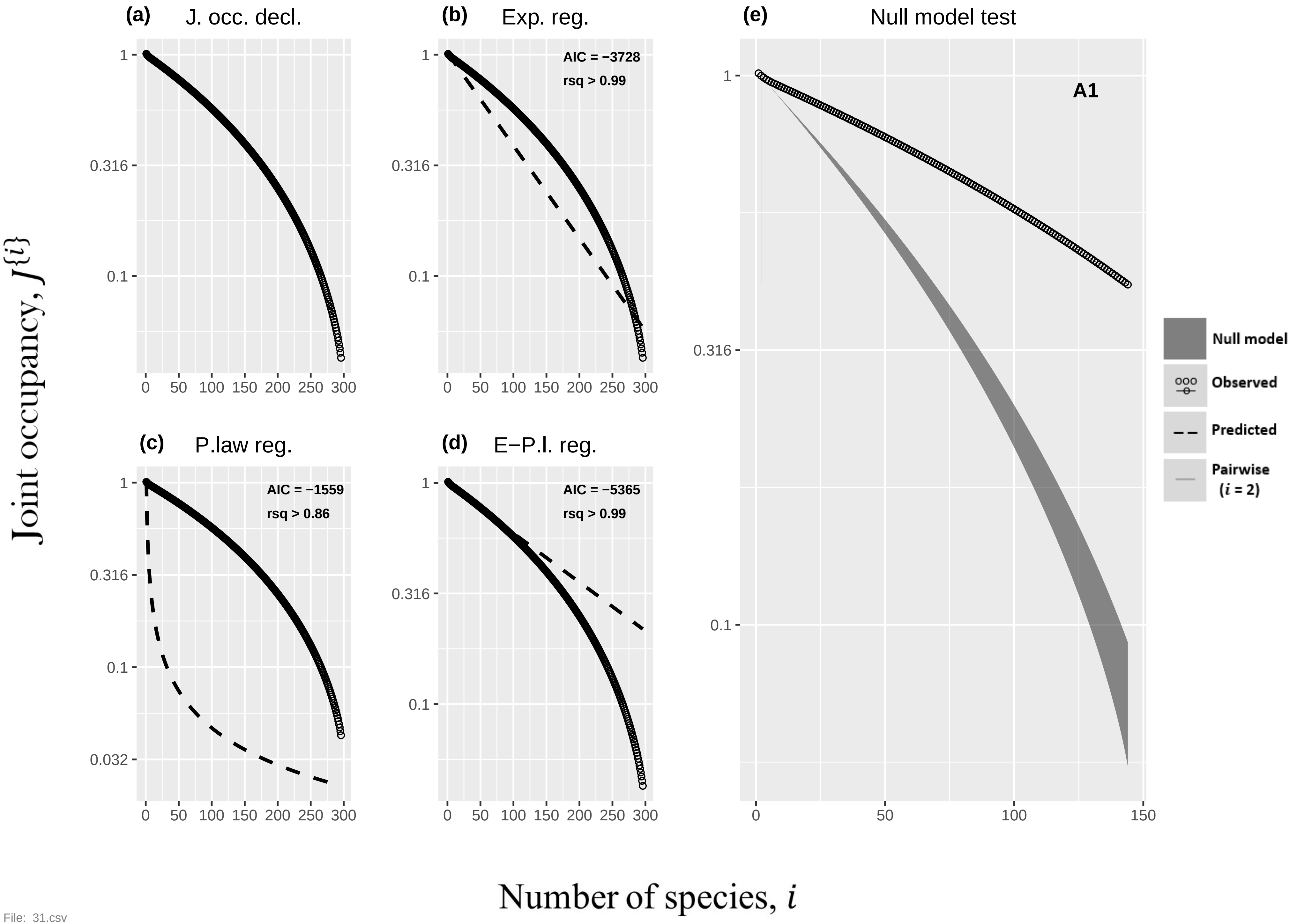

(a)
J. occ. decl.
(b)
Exp. reg.
AIC = −3728
rsq > 0.99
(e)
Null model test
A1
(c)
P.law reg.
AIC = −1559
rsq > 0.86
(d)
E−P.l. reg.
AIC = −5365
rsq > 0.99
Null model
Observed
Predicted
Pairwise
(i = 2)
Joint occupancy, J{i}
Number of species, i


File: 31.csv

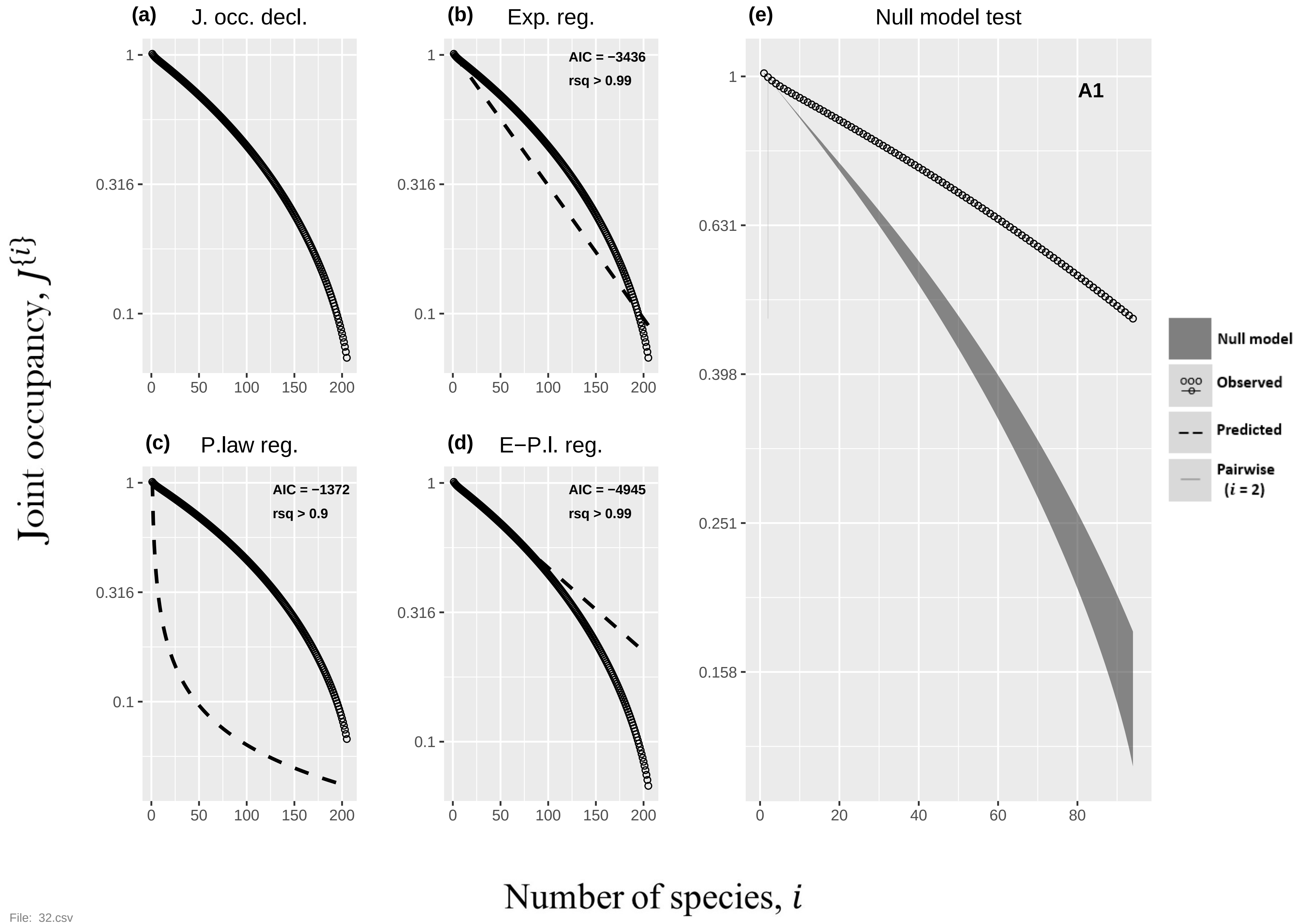
(a) J. occ. decl.
(b) Exp. reg.
AIC = −3436
rsq > 0.99
(c) P.law reg.
AIC = −1372
rsq > 0.9
(d) E−P.l. reg.
AIC = −4945
rsq > 0.99
(e) Null model test
A1
Null model
Observed
Predicted
Pairwise (i = 2)
Joint occupancy, J{i}
Number of species, i
1
0.316
0.1
0.631
0.398
0.251
0.158
0
20
40
60
80
50
100
150
200

File: 32.csv

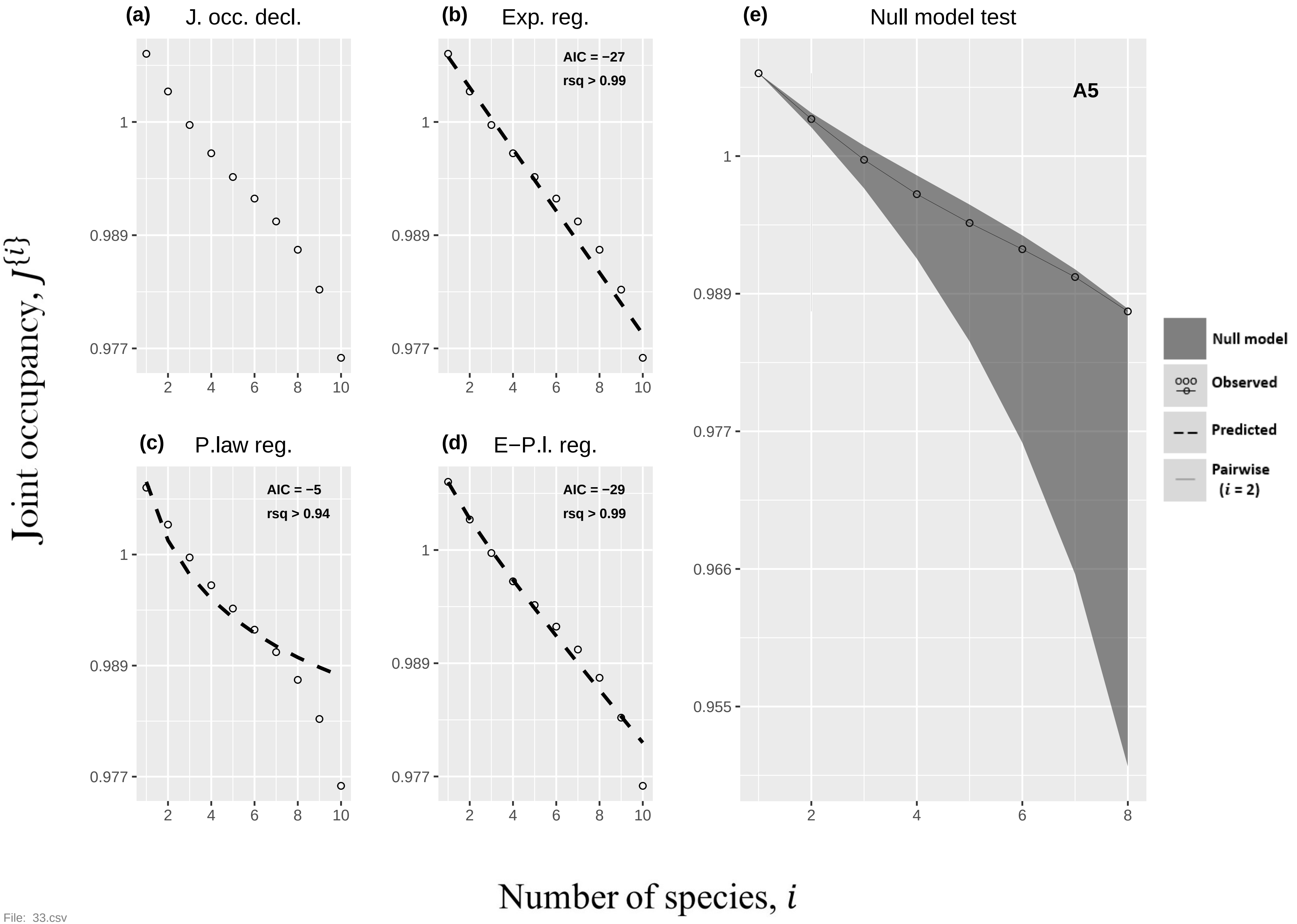
(a) J. occ. decl.
(b) Exp. reg.
AIC = −27
rsq > 0.99
(c) P.law reg.
AIC = −5
rsq > 0.94
(d) E−P.l. reg.
AIC = −29
rsq > 0.99
(e) Null model test
A5
Null model
Observed
Predicted
Pairwise (i = 2)
Joint occupancy, J{i}
Number of species, i

File: 33.csv

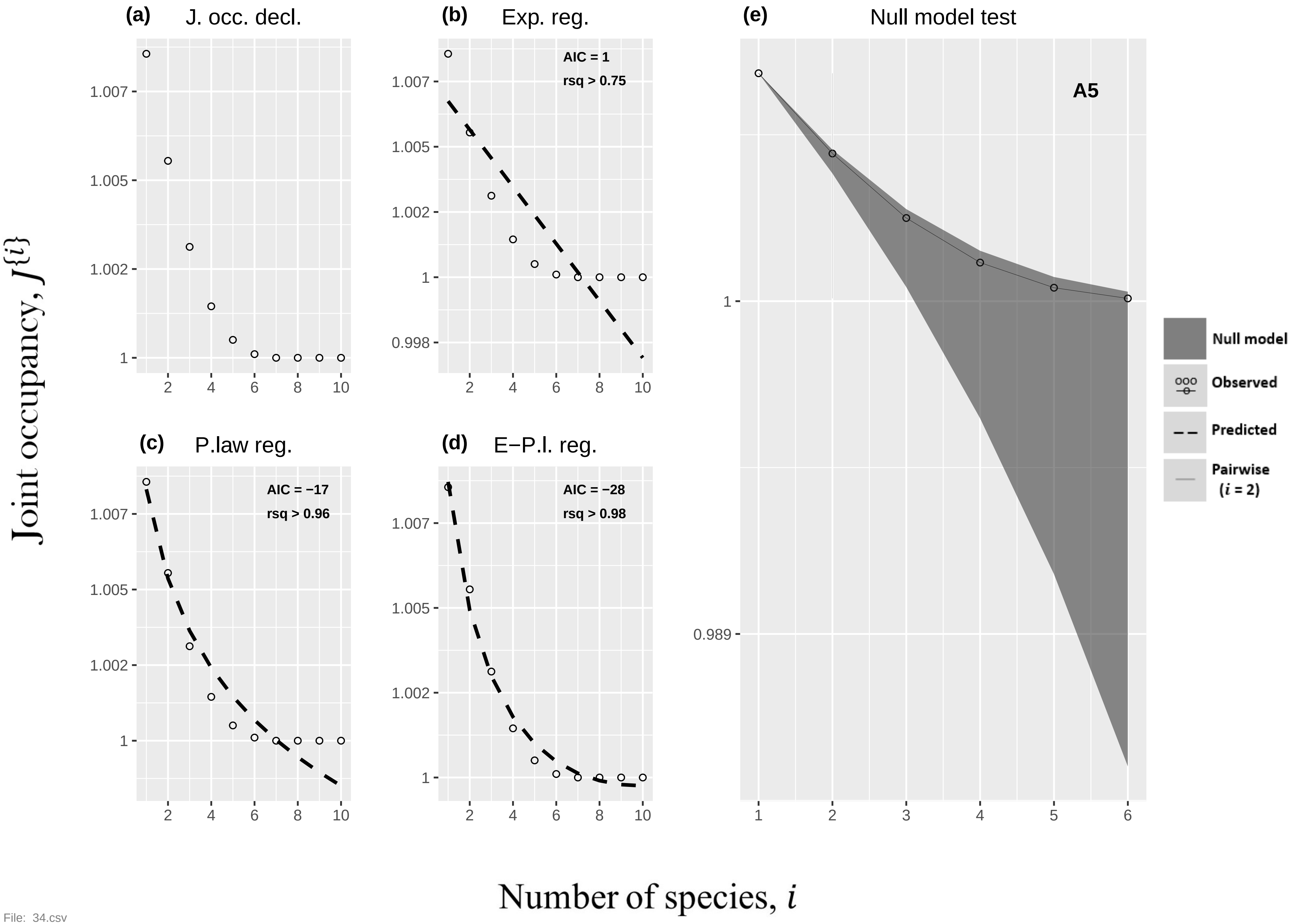
(a) J. occ. decl.
(b) Exp. reg.
AIC = 1
rsq > 0.75
(c) P.law reg.
AIC = −17
rsq > 0.96
(d) E−P.l. reg.
AIC = −28
rsq > 0.98
(e) Null model test
A5
Null model
Observed
Predicted
Pairwise
(i = 2)
Joint occupancy, J{i}
Number of species, i

File: 34.csv

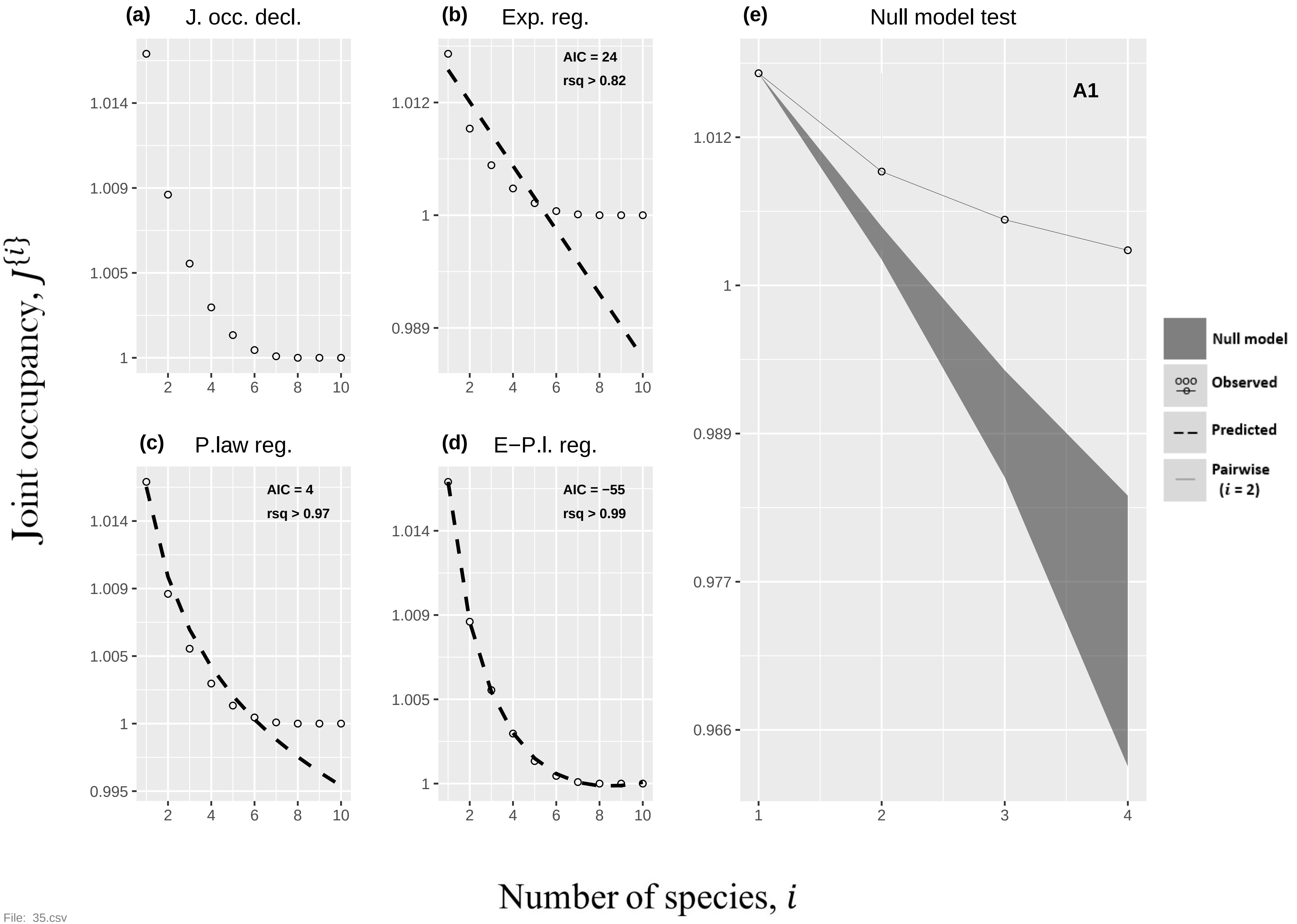
(a)
J. occ. decl.
(b)
Exp. reg.
AIC = 24
rsq > 0.82
(c)
P.law reg.
AIC = 4
rsq > 0.97
(d)
E−P.l. reg.
AIC = −55
rsq > 0.99
(e)
Null model test
A1
Null model
Observed
Predicted
Pairwise
(i = 2)
Joint occupancy, J{i}
Number of species, i

File: 35.csv

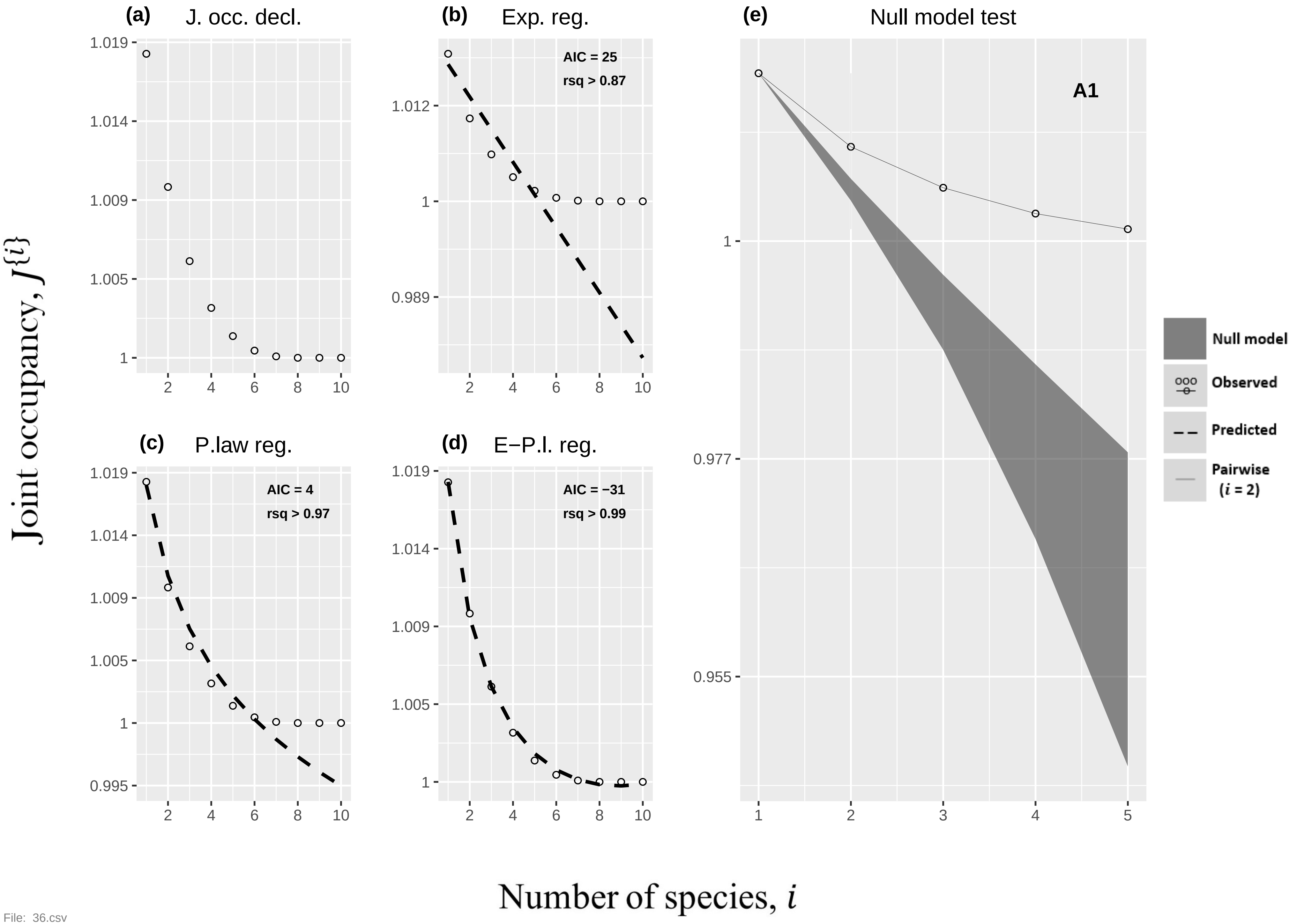

(a) J. occ. decl.
(b) Exp. reg.
AIC = 25
rsq > 0.87
(c) P.law reg.
AIC = 4
rsq > 0.97
(d) E−P.l. reg.
AIC = −31
rsq > 0.99
(e) Null model test
A1
Null model
Observed
Predicted
Pairwise (i = 2)
Joint occupancy, J{i}
Number of species, i


File: 36.csv

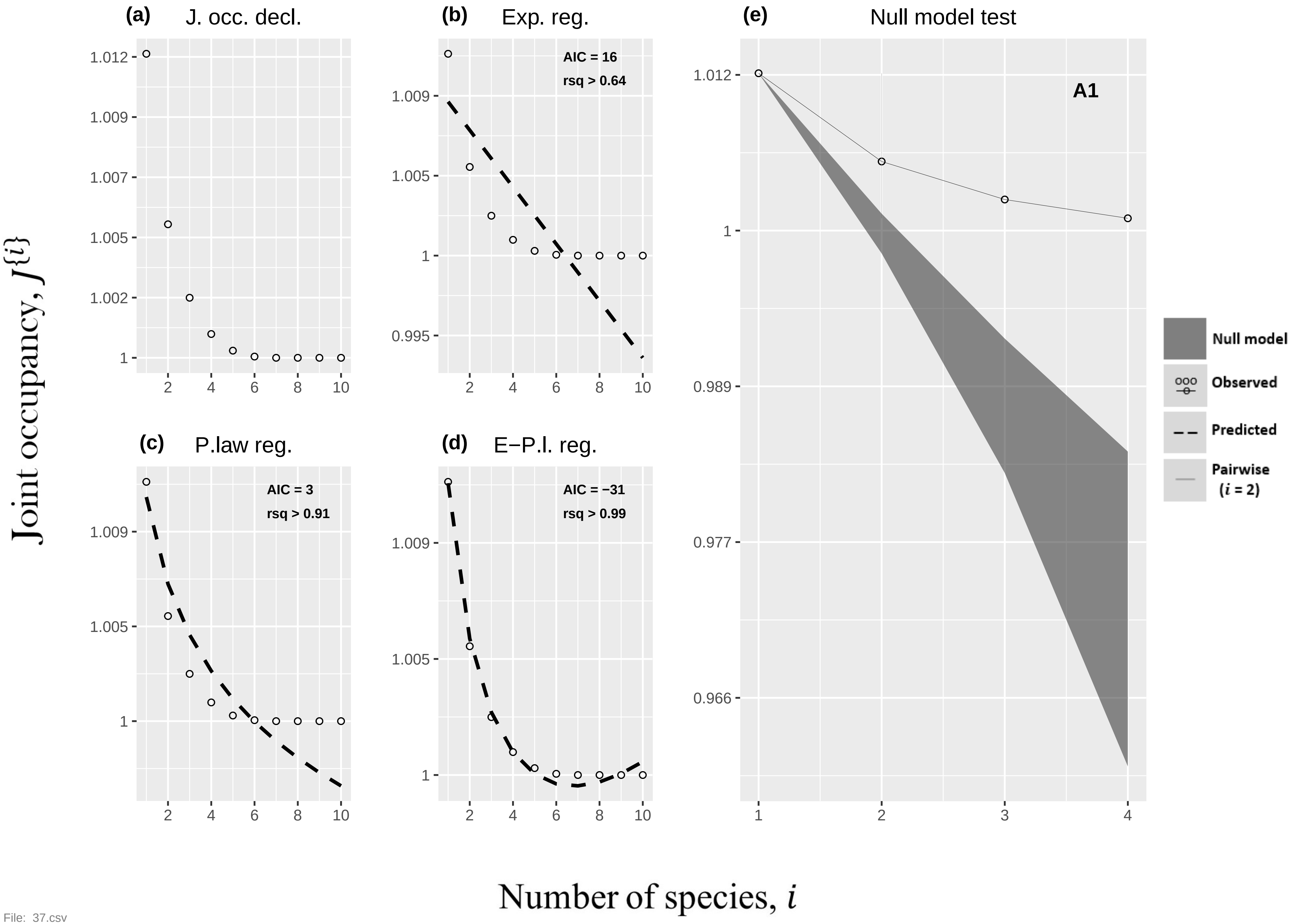
(a) J. occ. decl.
(b) Exp. reg.
AIC = 16
rsq > 0.64
(e) Null model test
A1
(c) P.law reg.
AIC = 3
rsq > 0.91
(d) E−P.l. reg.
AIC = −31
rsq > 0.99
Null model
Observed
Predicted
Pairwise (i = 2)
Joint occupancy, J{i}
Number of species, i

File: 37.csv

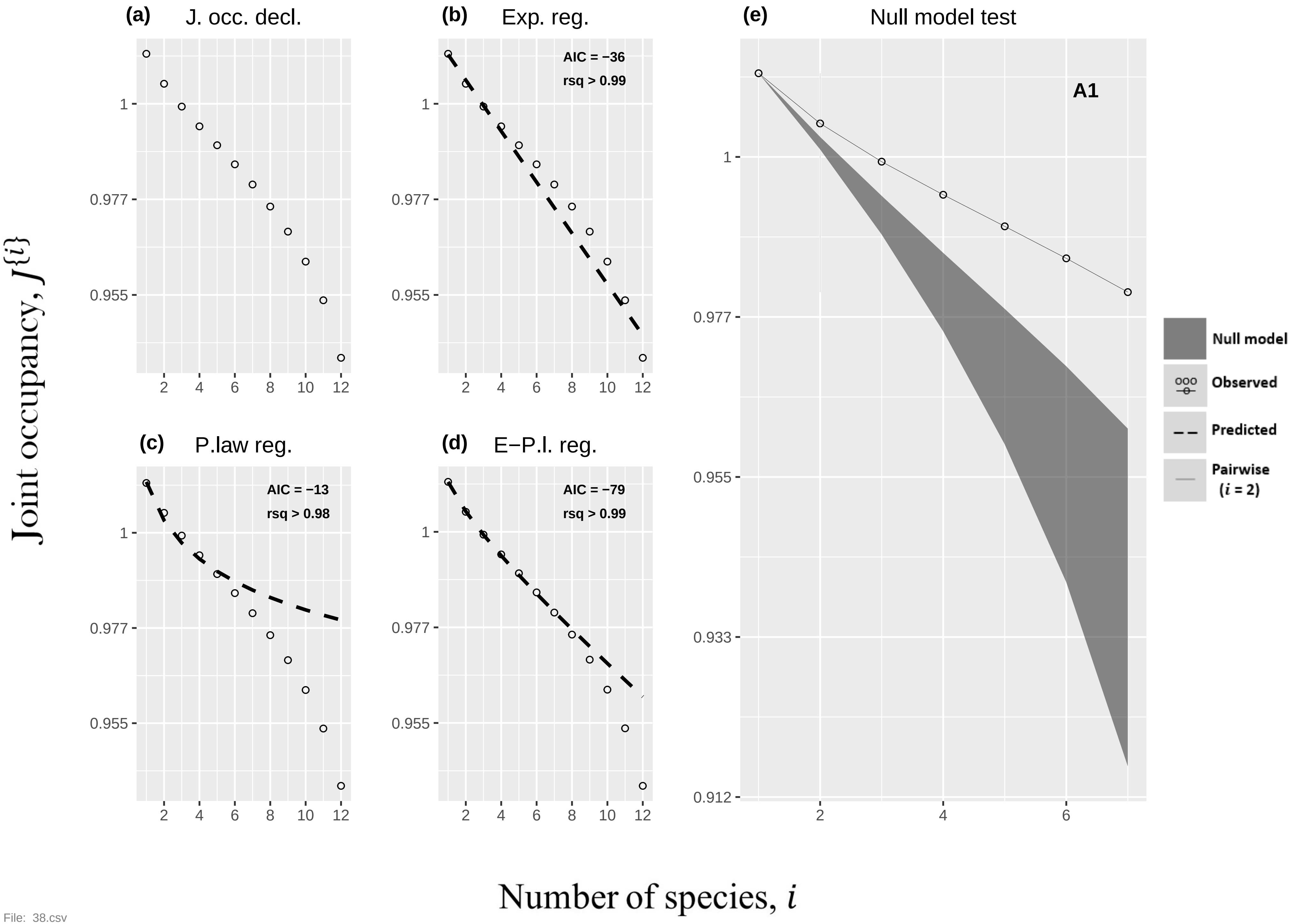

(a) J. occ. decl.
(b) Exp. reg.
AIC = −36
rsq > 0.99
(c) P.law reg.
AIC = −13
rsq > 0.98
(d) E−P.l. reg.
AIC = −79
rsq > 0.99
(e) Null model test
A1
Null model
Observed
Predicted
Pairwise (i = 2)
1
0.977
0.955
0.933
0.912
2
4
6
8
10
12
Joint occupancy, J{i}
Number of species, i


File: 38.csv

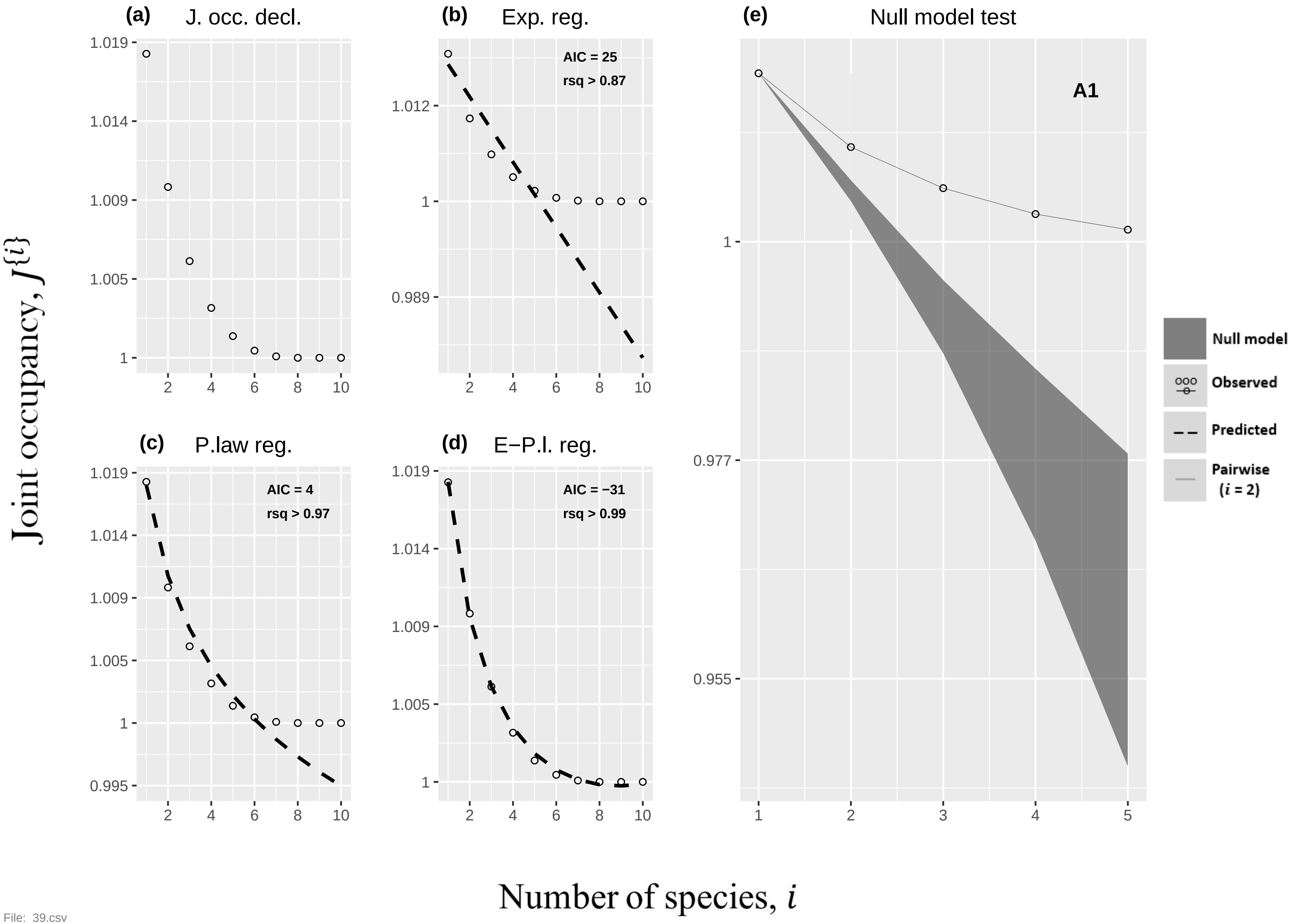

(a) J. occ. decl.
(b) Exp. reg.
AIC = 25
rsq > 0.87
(c) P.law reg.
AIC = 4
rsq > 0.97
(d) E−P.l. reg.
AIC = −31
rsq > 0.99
(e) Null model test
A1
Null model
Observed
Predicted
Pairwise (i = 2)
Joint occupancy, J{i}
Number of species, i


File: 39.csv

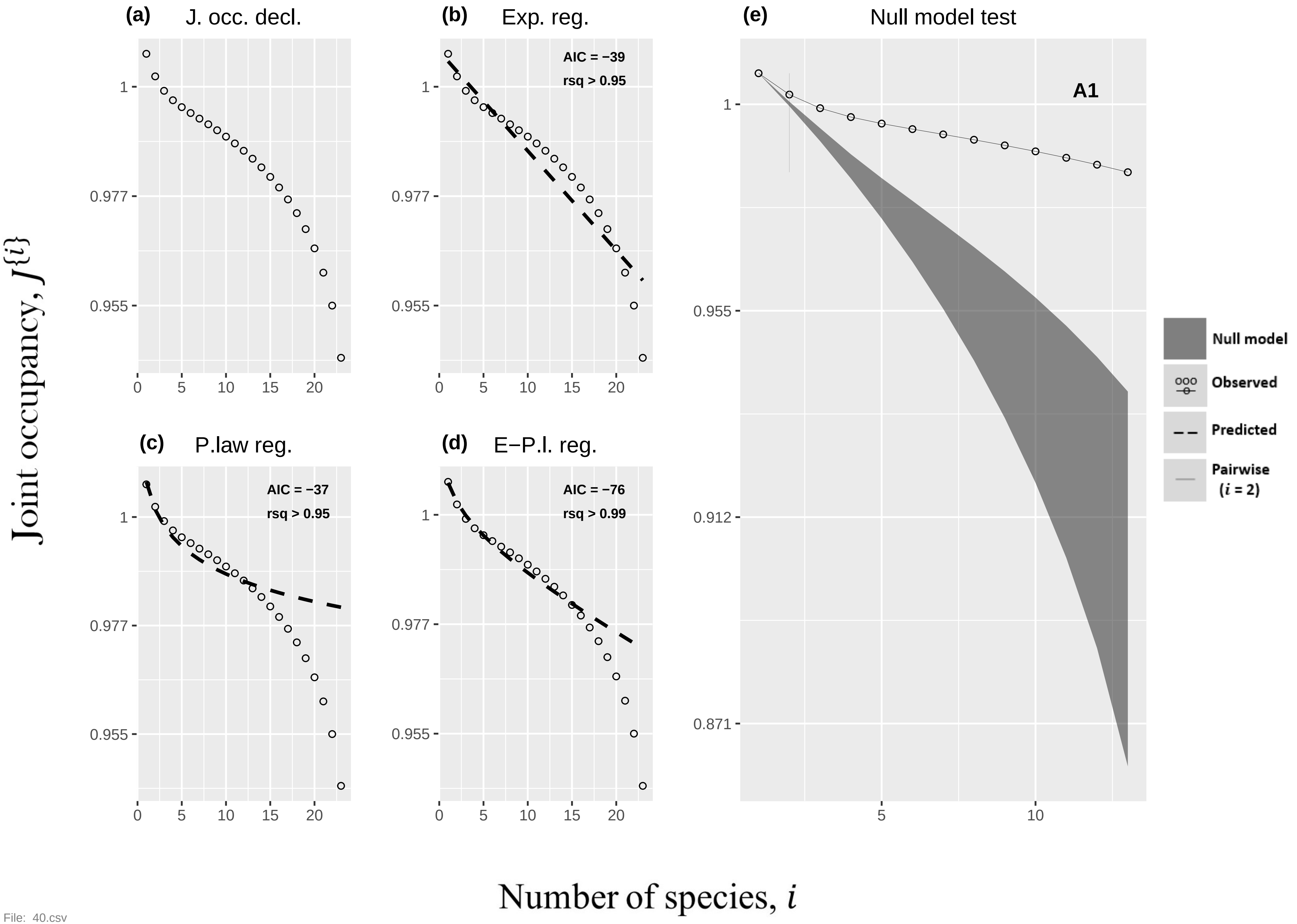

(a)
J. occ. decl.
(b)
Exp. reg.
AIC = −39
rsq > 0.95
(c)
P.law reg.
AIC = −37
rsq > 0.95
(d)
E−P.l. reg.
AIC = −76
rsq > 0.99
(e)
Null model test
A1
Null model
Observed
Predicted
Pairwise
(i = 2)
Joint occupancy, J{i}
Number of species, i
1
0.977
0.955
0.912
0.871
0
5
10
15
20

File: 40.csv

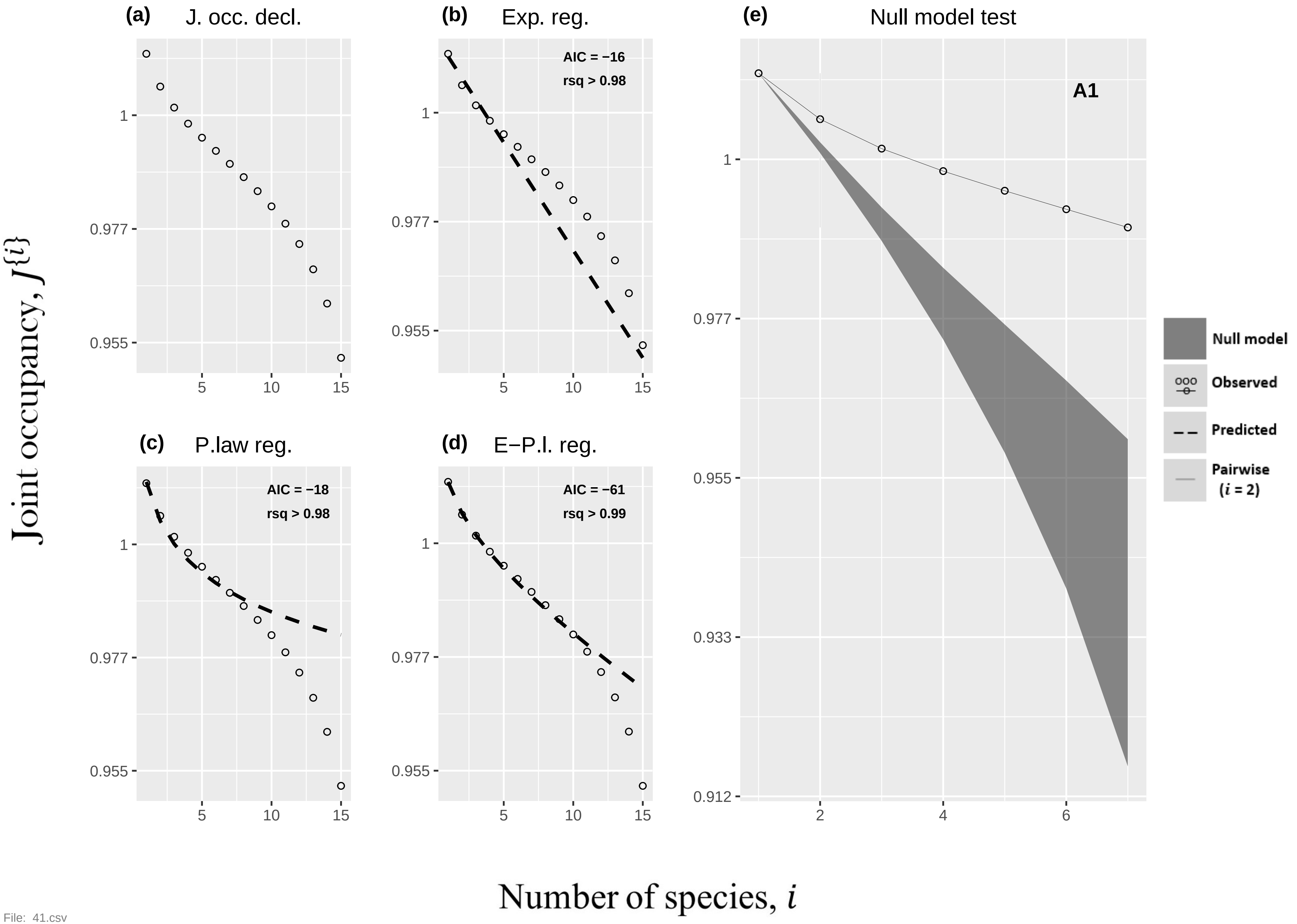
(a) J. occ. decl.
(b) Exp. reg.
AIC = −16
rsq > 0.98
(c) P.law reg.
AIC = −18
rsq > 0.98
(d) E−P.l. reg.
AIC = −61
rsq > 0.99
(e) Null model test
A1
Null model
Observed
Predicted
Pairwise (i = 2)
Joint occupancy, J{i}
Number of species, i
1
0.977
0.955
0.933
0.912
5
10
15
2
4
6
File: 41.csv

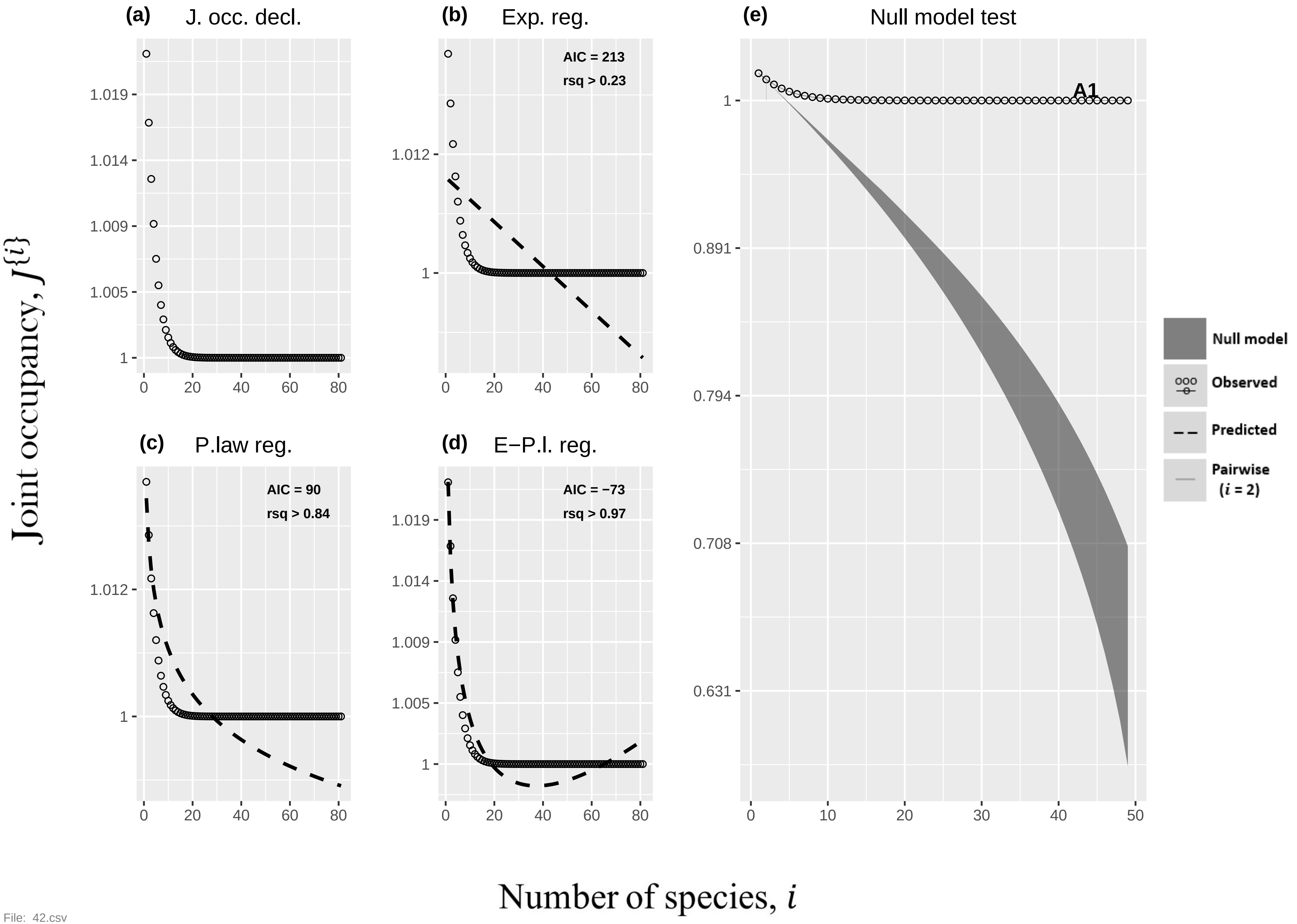

(a)
J. occ. decl.
(b)
Exp. reg.
AIC = 213
rsq > 0.23
(c)
P.law reg.
AIC = 90
rsq > 0.84
(d)
E−P.l. reg.
AIC = −73
rsq > 0.97
(e)
Null model test
A1
Null model
Observed
Predicted
Pairwise
(i = 2)
Joint occupancy, J{i}
Number of species, i


File: 42.csv

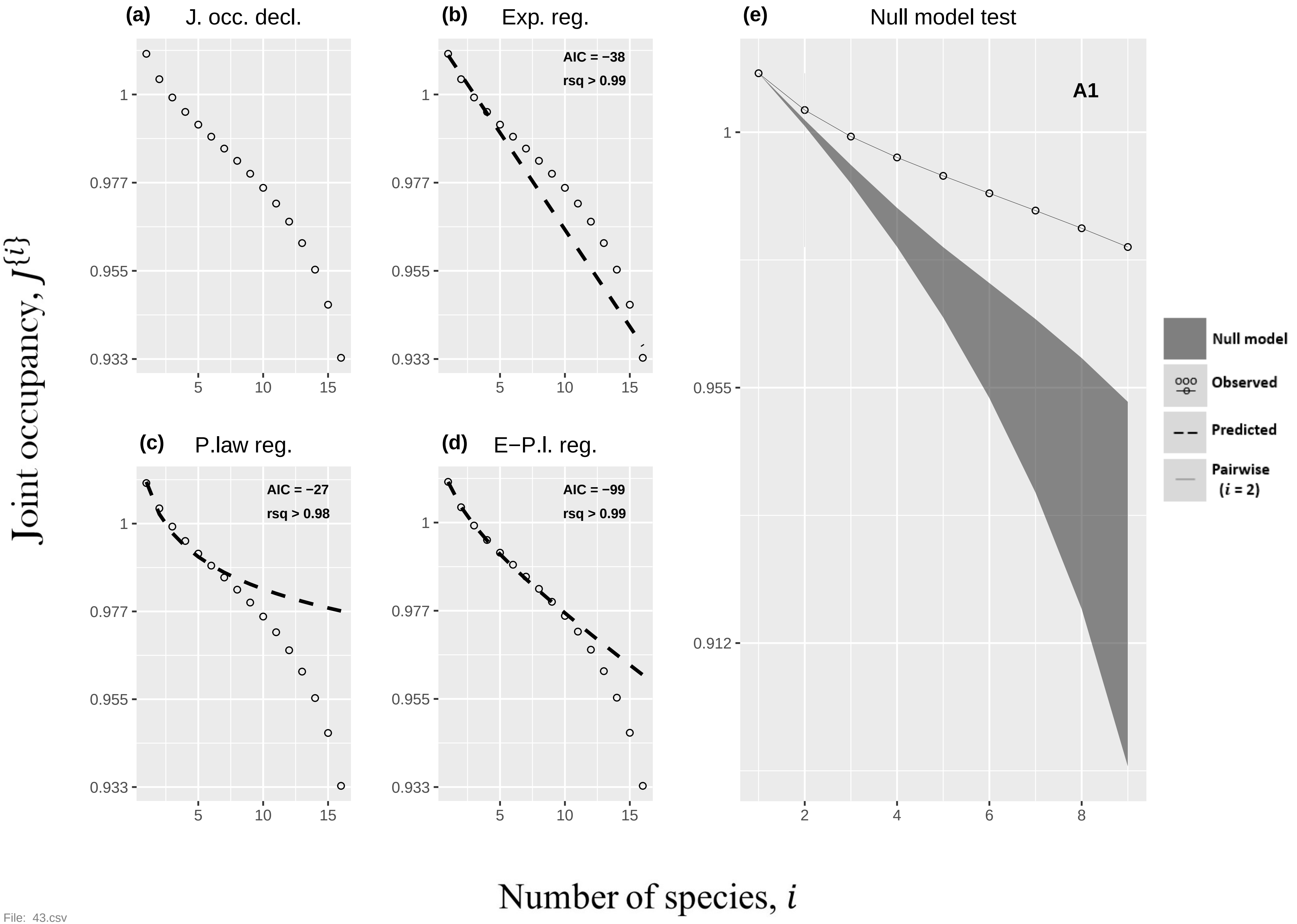

(a) J. occ. decl.
(b) Exp. reg.
AIC = −38
rsq > 0.99
(e) Null model test
A1
(c) P.law reg.
AIC = −27
rsq > 0.98
(d) E−P.l. reg.
AIC = −99
rsq > 0.99
Null model
Observed
Predicted
Pairwise
(i = 2)
1
0.977
0.955
0.933
0.912
5
10
15
2
4
6
8
Joint occupancy, J{i}
Number of species, i


File: 43.csv

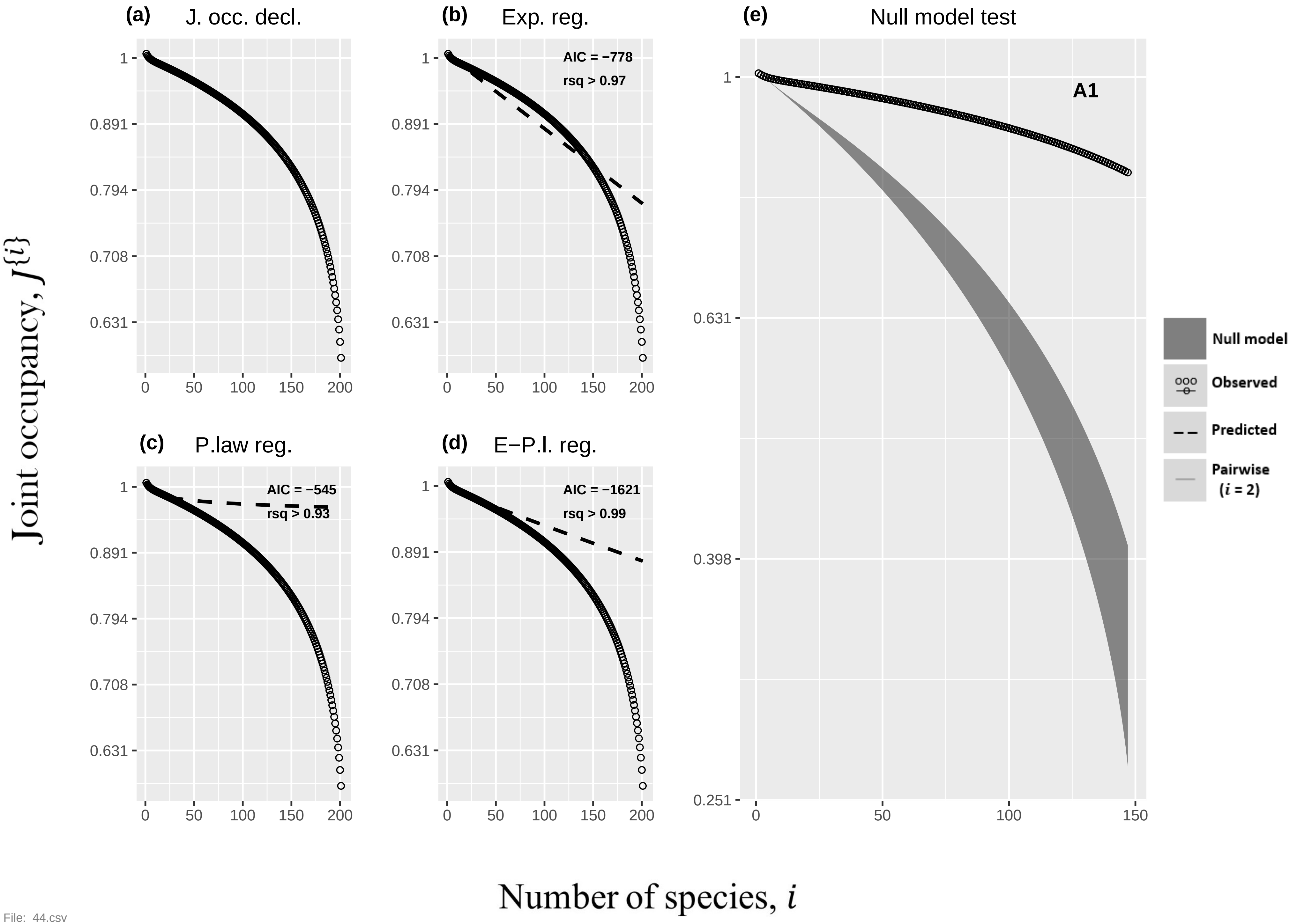

(a)
J. occ. decl.
(b)
Exp. reg.
AIC = −778
rsq > 0.97
(c)
P.law reg.
AIC = −545
rsq > 0.93
(d)
E−P.l. reg.
AIC = −1621
rsq > 0.99
(e)
Null model test
A1
1
0.891
0.794
0.708
0.631
0.398
0.251
0
50
100
150
200
Null model
Observed
Predicted
Pairwise
(i = 2)
Joint occupancy, J{i}
Number of species, i


File: 44.csv

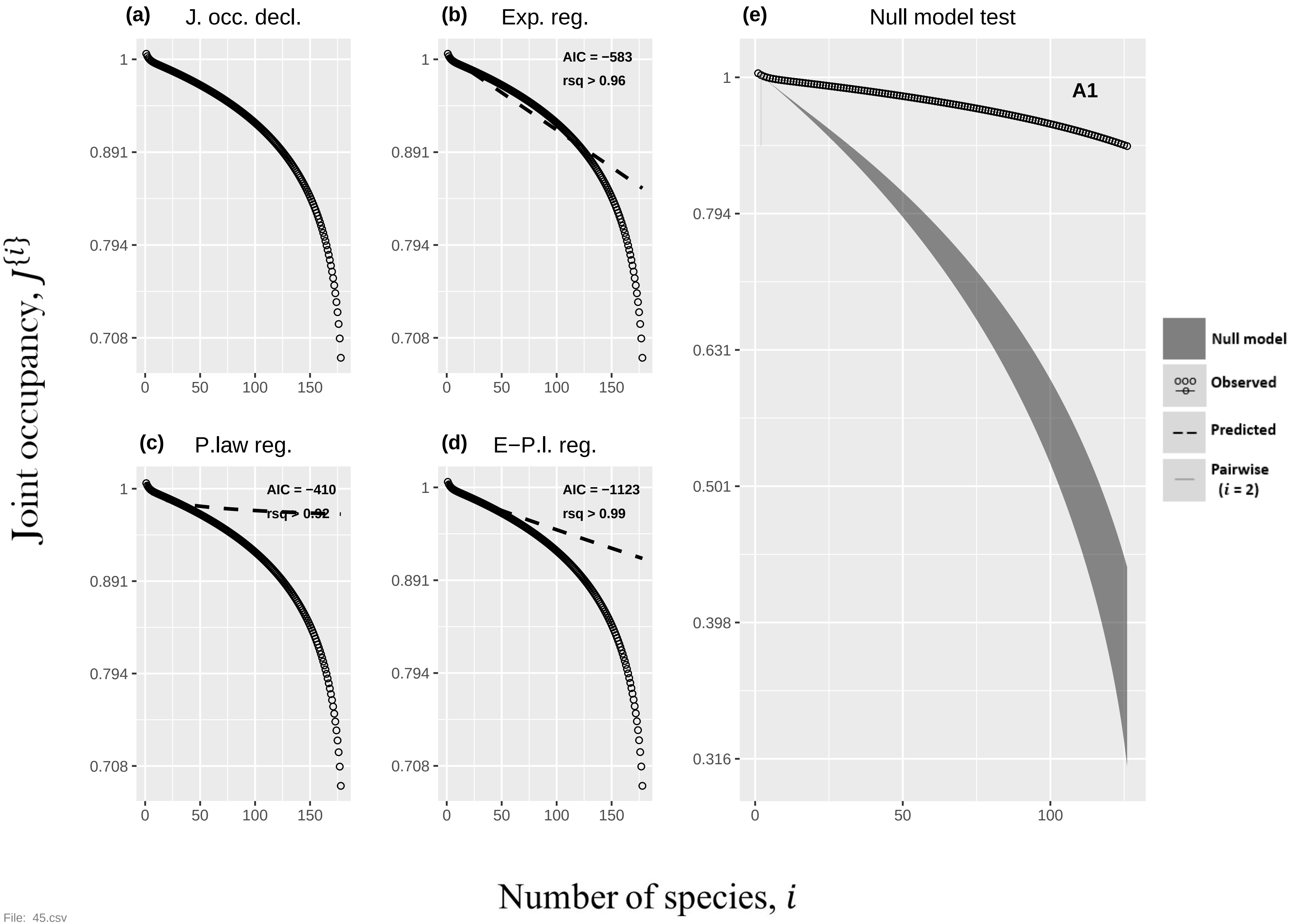

(a) J. occ. decl.
(b) Exp. reg.
AIC = −583
rsq > 0.96
(c) P.law reg.
AIC = −410
rsq > 0.92
(d) E−P.l. reg.
AIC = −1123
rsq > 0.99
(e) Null model test
A1
Null model
Observed
Predicted
Pairwise (i = 2)
1
0.891
0.794
0.708
0.631
0.501
0.398
0.316
0
50
100
150
Joint occupancy, J{i}
Number of species, i


File: 45.csv

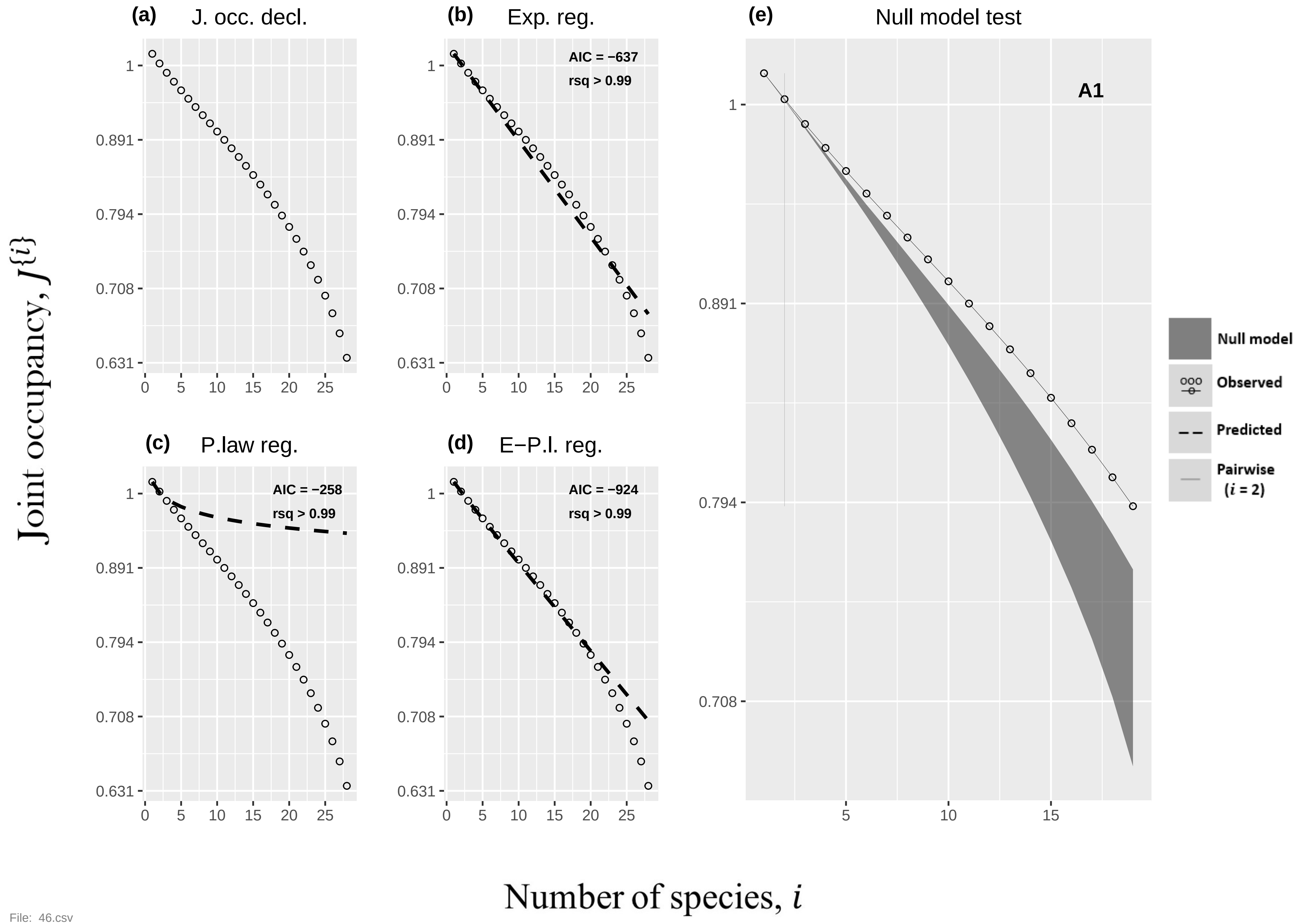
(a) J. occ. decl.
(b) Exp. reg.
AIC = −637
rsq > 0.99
(c) P.law reg.
AIC = −258
rsq > 0.99
(d) E−P.l. reg.
AIC = −924
rsq > 0.99
(e) Null model test
A1
Null model
Observed
Predicted
Pairwise ($i$ = 2)
Joint occupancy, $J^{\{i\}}$
Number of species, $i$
1
0.891
0.794
0.708
0.631
0
5
10
15
20
25

File: 46.csv

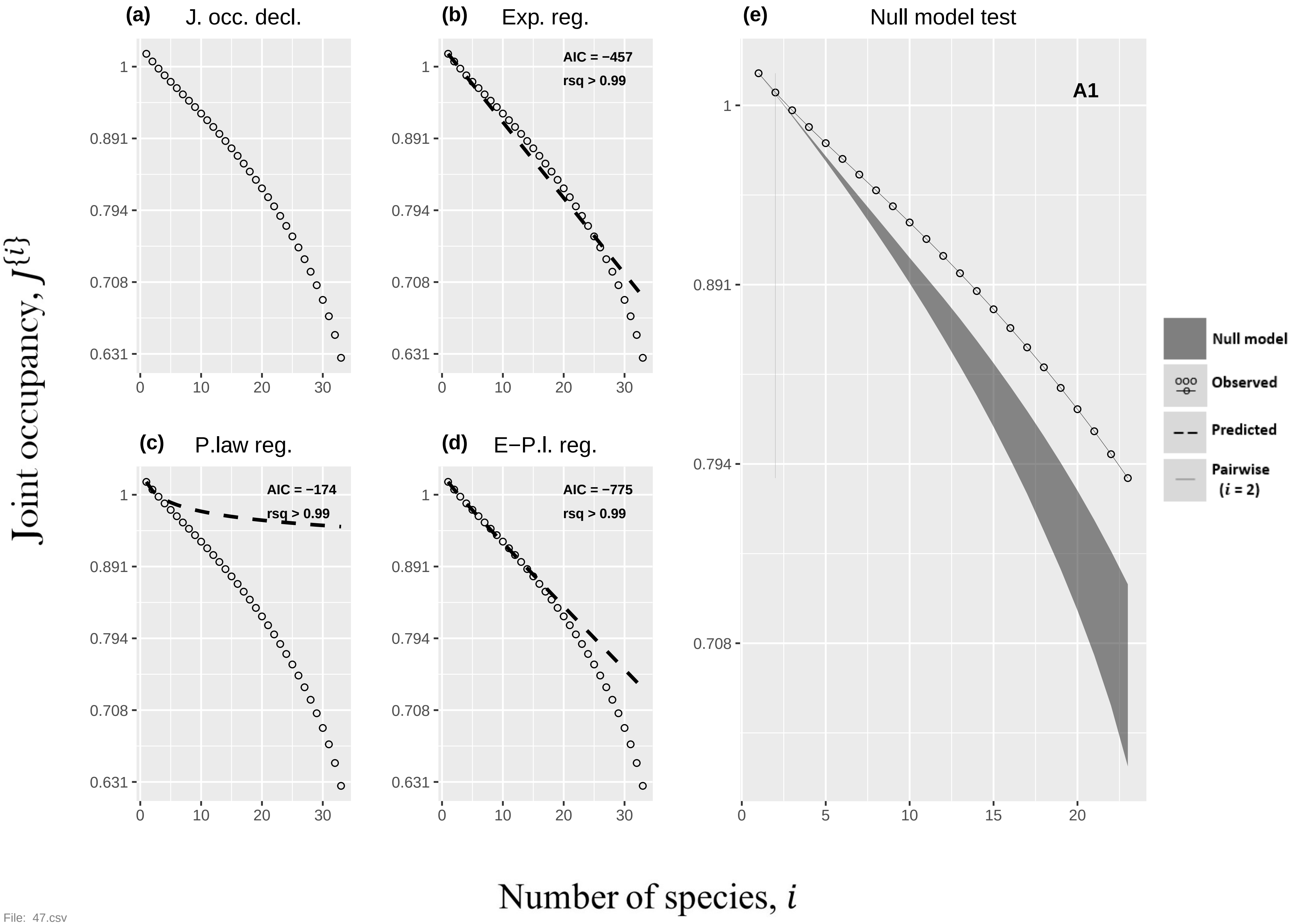
(a) J. occ. decl.
(b) Exp. reg.
AIC = −457
rsq > 0.99
(c) P.law reg.
AIC = −174
rsq > 0.99
(d) E−P.l. reg.
AIC = −775
rsq > 0.99
(e) Null model test
A1
Null model
Observed
Predicted
Pairwise ($i$ = 2)
Joint occupancy, $J^{\{i\}}$
Number of species, $i$

File: 47.csv

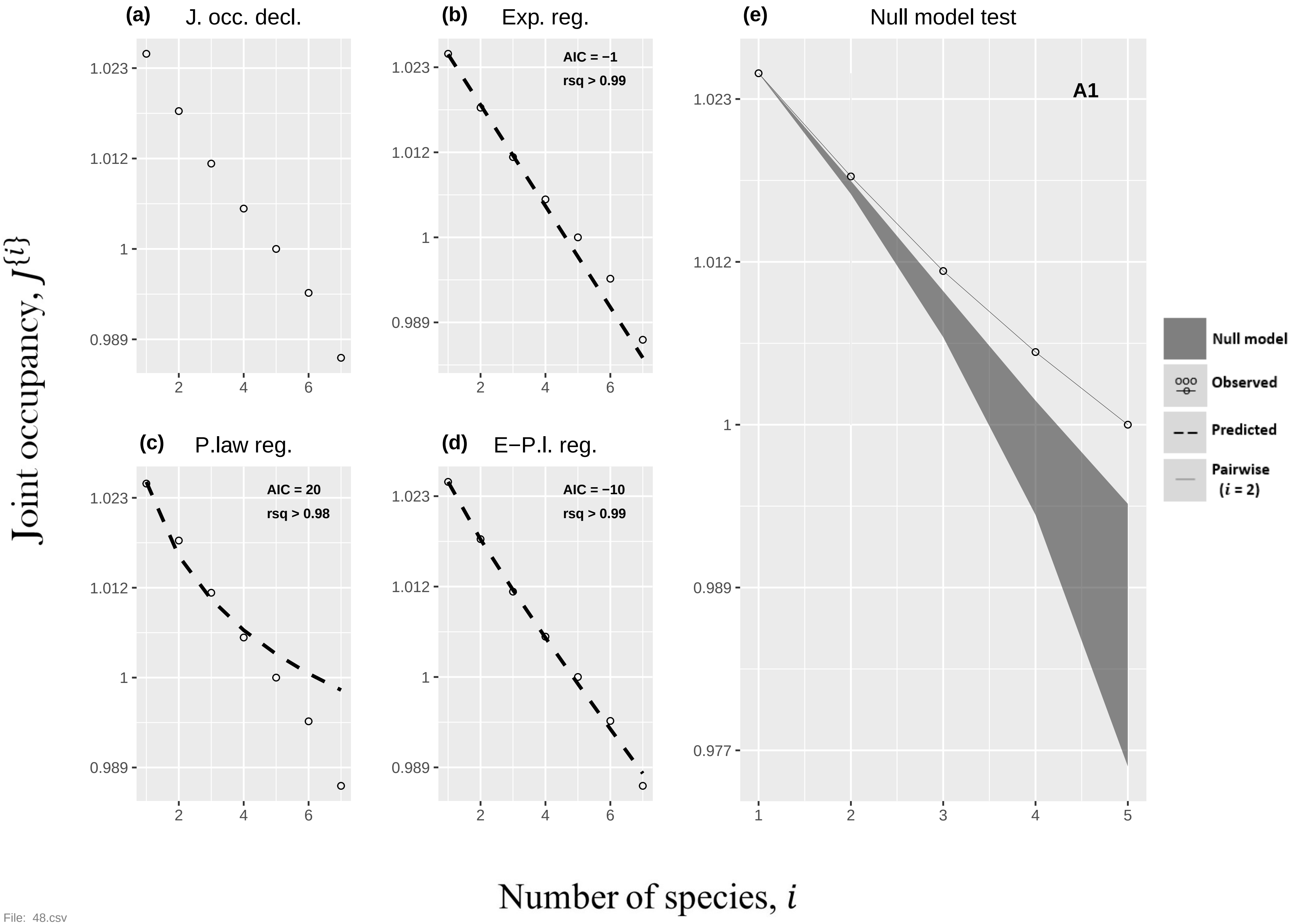

(a) J. occ. decl.
(b) Exp. reg.
AIC = −1
rsq > 0.99
(e) Null model test
A1
(c) P.law reg.
AIC = 20
rsq > 0.98
(d) E−P.l. reg.
AIC = −10
rsq > 0.99
Null model
Observed
Predicted
Pairwise (i = 2)
Joint occupancy, J{i}
Number of species, i


File: 48.csv

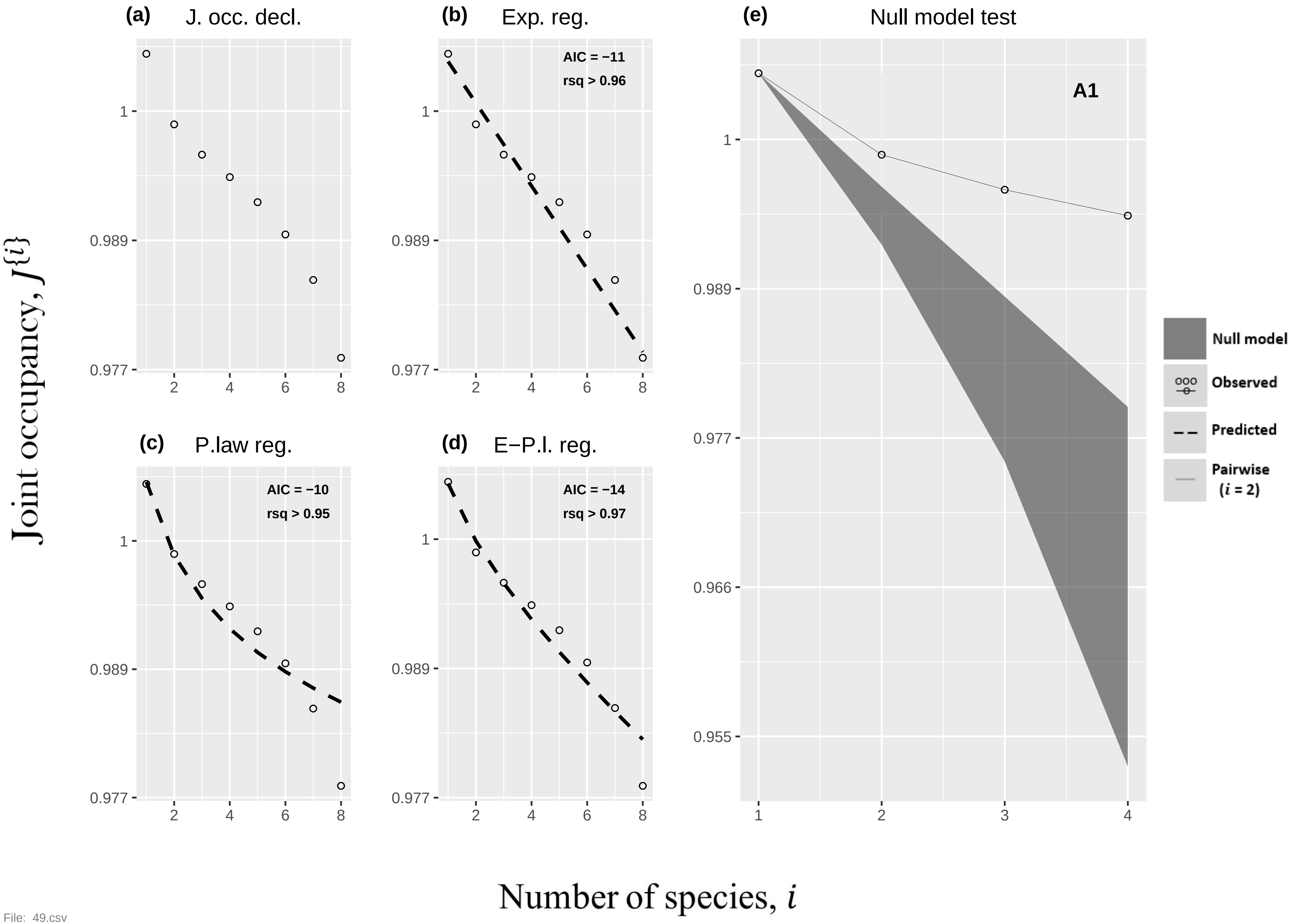

(a) J. occ. decl.
(b) Exp. reg.
AIC = −11
rsq > 0.96
(c) P.law reg.
AIC = −10
rsq > 0.95
(d) E−P.l. reg.
AIC = −14
rsq > 0.97
(e) Null model test
A1
Null model
Observed
Predicted
Pairwise (i = 2)
1
0.989
0.977
0.966
0.955
2
4
6
8
Joint occupancy, J{i}
Number of species, i


File: 49.csv

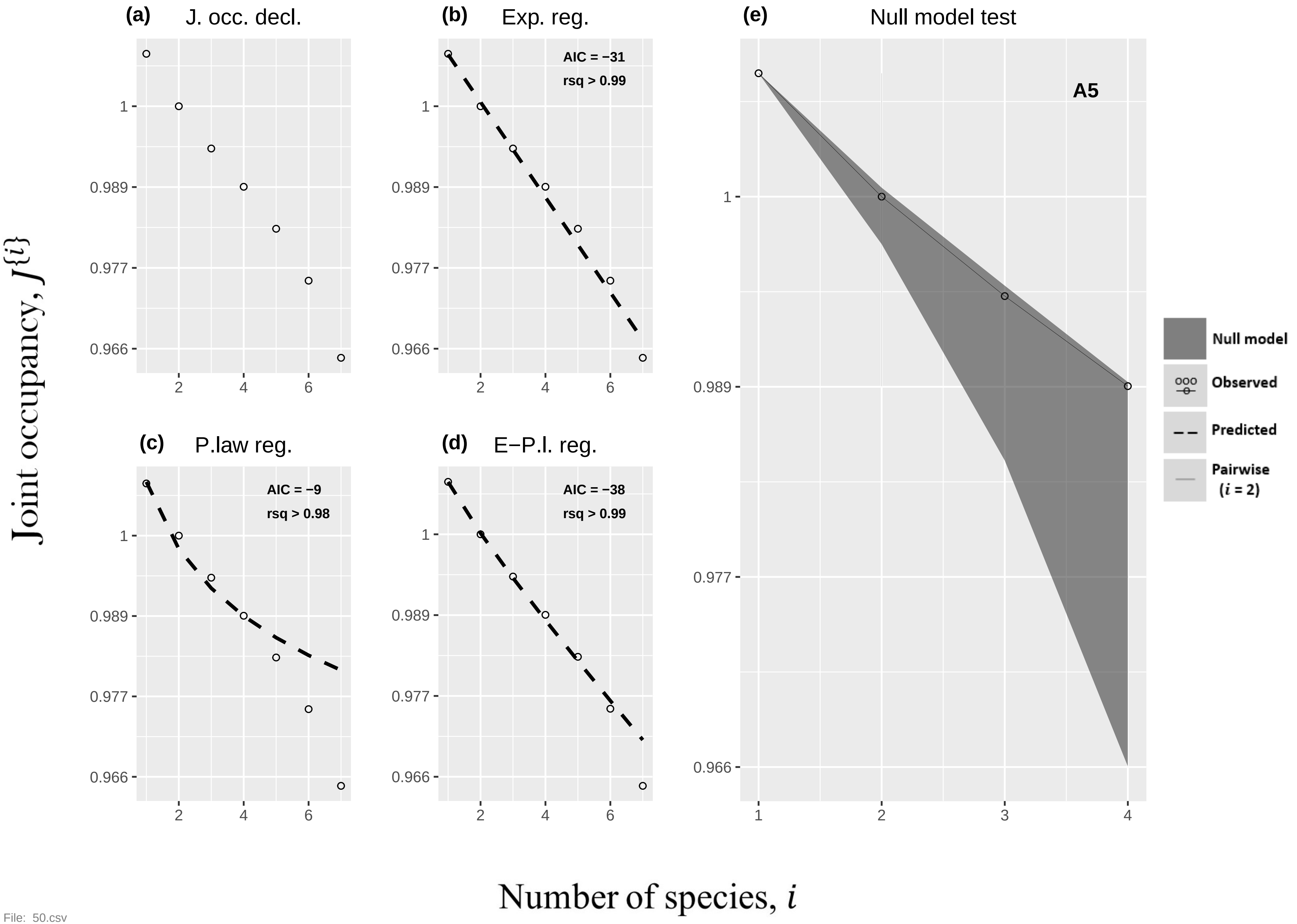

(a)
J. occ. decl.
(b)
Exp. reg.
AIC = −31
rsq > 0.99
(c)
P.law reg.
AIC = −9
rsq > 0.98
(d)
E−P.l. reg.
AIC = −38
rsq > 0.99
(e)
Null model test
A5
Null model
Observed
Predicted
Pairwise (i = 2)
Joint occupancy, J{i}
Number of species, i
1
0.989
0.977
0.966
2
4
6
3


File: 50.csv

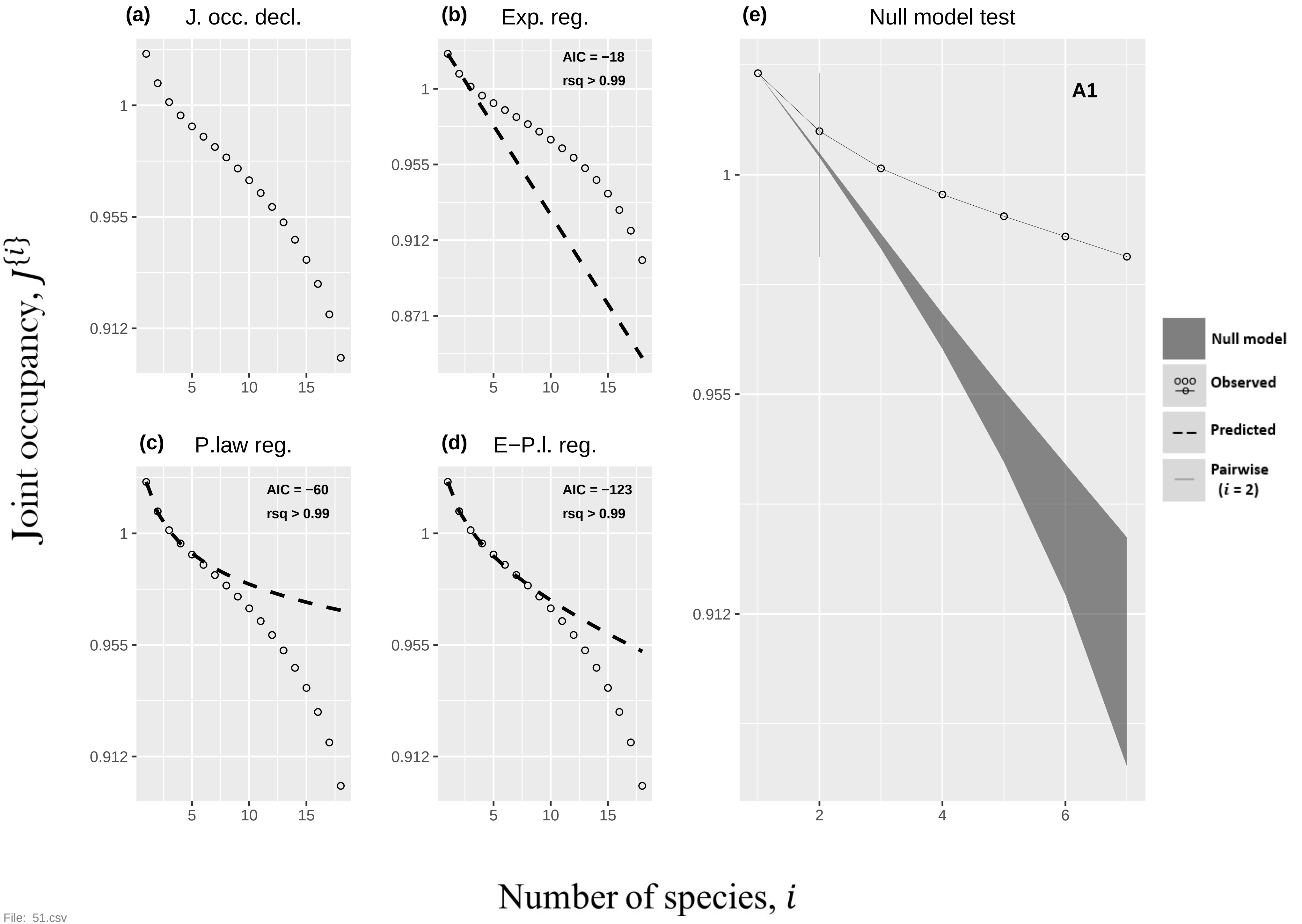

(a) J. occ. decl.
(b) Exp. reg.
AIC = −18
rsq > 0.99
(c) P.law reg.
AIC = −60
rsq > 0.99
(d) E−P.l. reg.
AIC = −123
rsq > 0.99
(e) Null model test
A1
Null model
Observed
Predicted
Pairwise (i = 2)
Joint occupancy, J{i}
Number of species, i


File: 51.csv

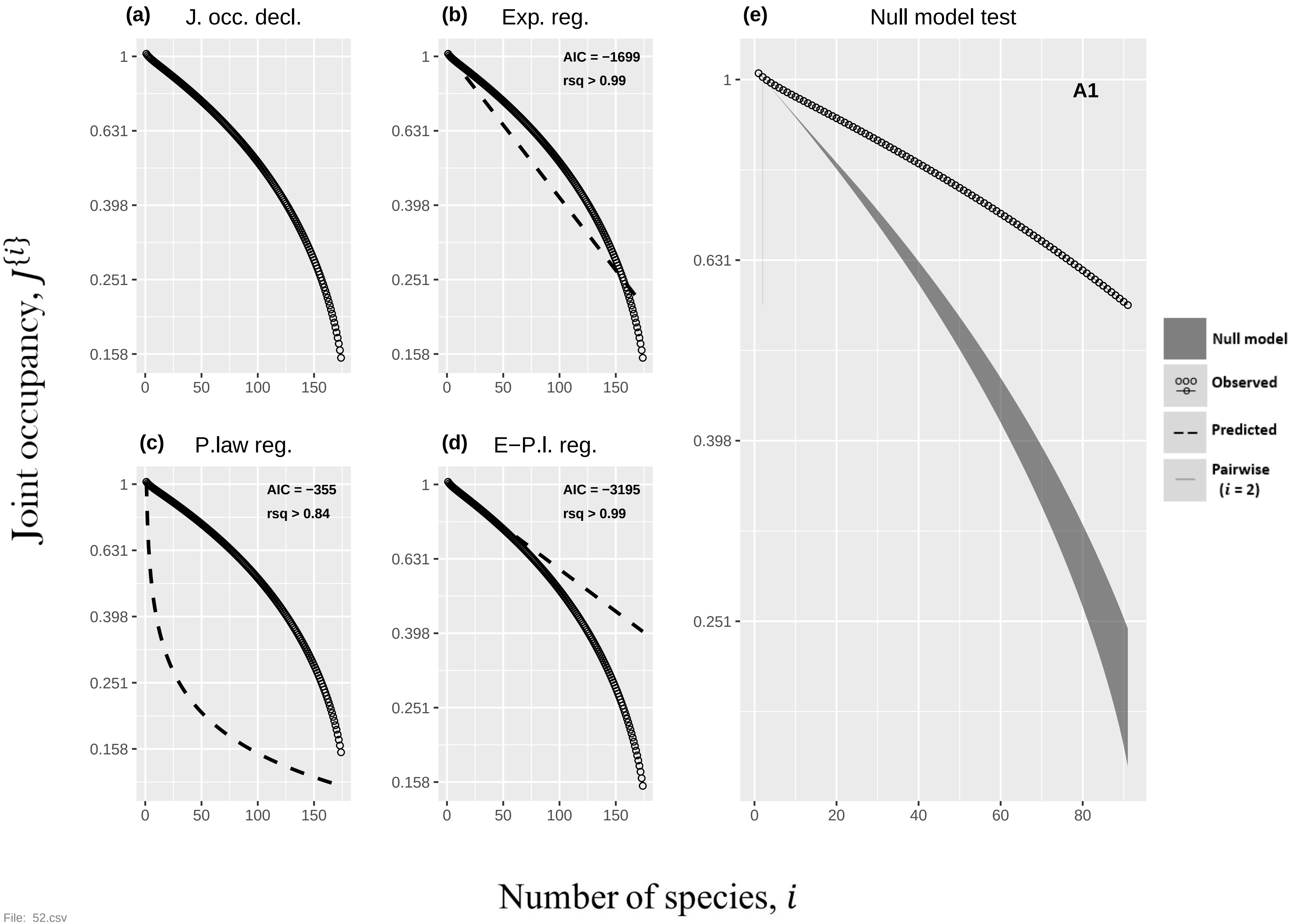

(a) J. occ. decl.
(b) Exp. reg.
AIC = −1699
rsq > 0.99
(c) P.law reg.
AIC = −355
rsq > 0.84
(d) E−P.l. reg.
AIC = −3195
rsq > 0.99
(e) Null model test
A1
Null model
Observed
Predicted
Pairwise
(i = 2)
1
0.631
0.398
0.251
0.158
0
50
100
150
20
40
60
80
Joint occupancy, J{i}
Number of species, i


File: 52.csv

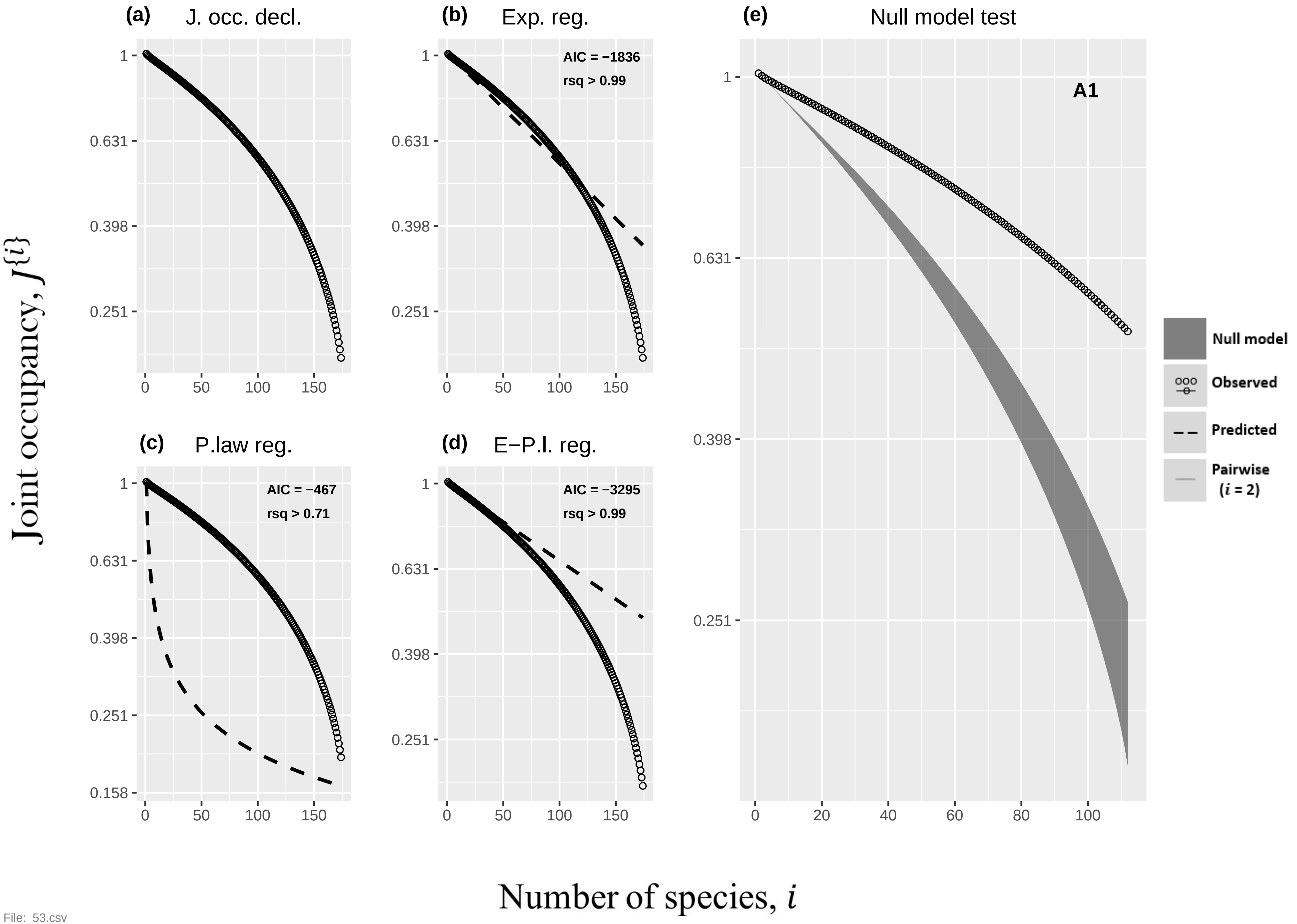
(a) J. occ. decl.
(b) Exp. reg.
AIC = −1836
rsq > 0.99
(e) Null model test
A1
(c) P.law reg.
AIC = −467
rsq > 0.71
(d) E−P.l. reg.
AIC = −3295
rsq > 0.99
Null model
Observed
Predicted
Pairwise (i = 2)
Joint occupancy, J{i}
Number of species, i


File: 53.csv

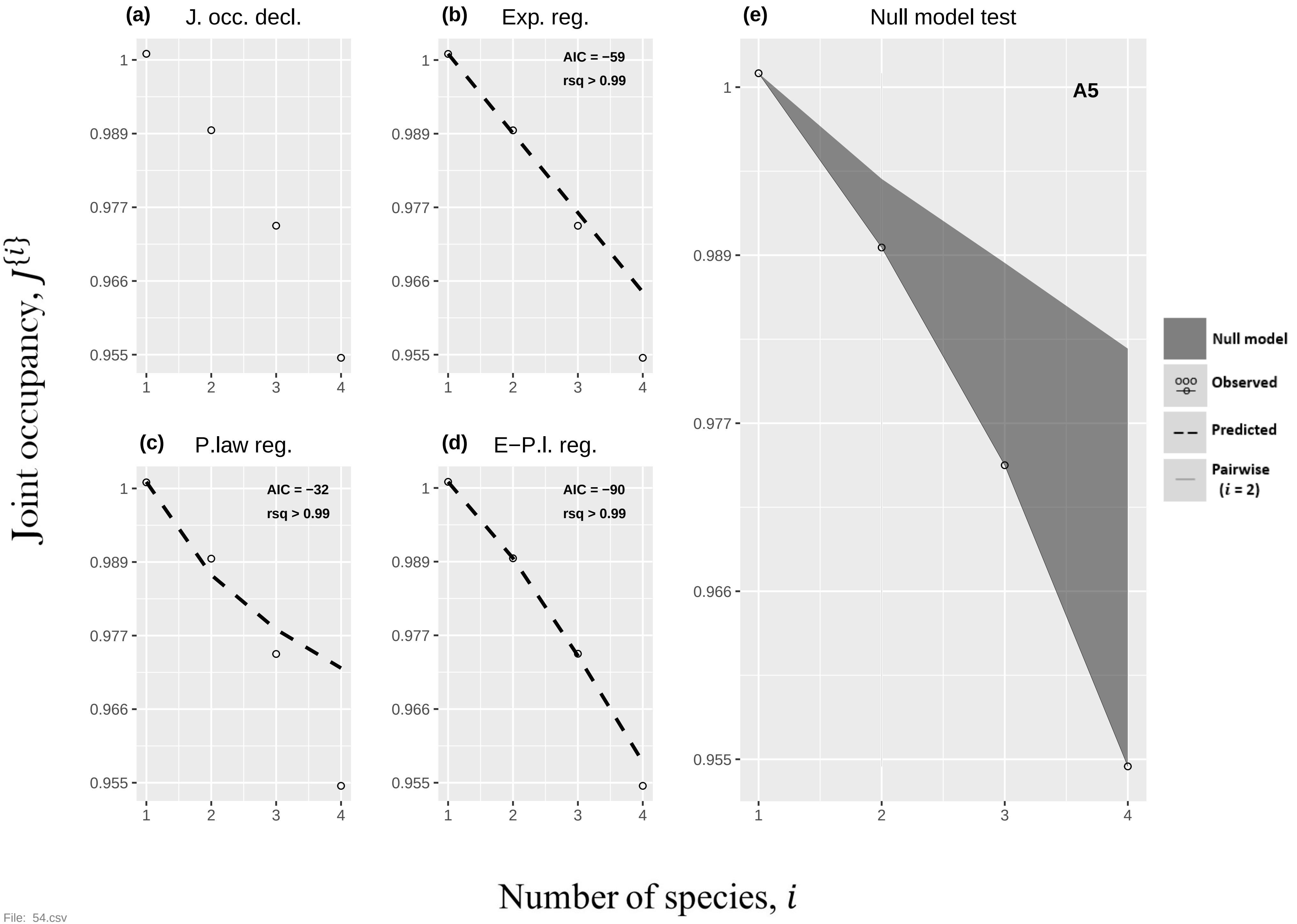

(a)
J. occ. decl.
(b)
Exp. reg.
AIC = −59
rsq > 0.99
(c)
P.law reg.
AIC = −32
rsq > 0.99
(d)
E−P.l. reg.
AIC = −90
rsq > 0.99
(e)
Null model test
A5
Null model
Observed
Predicted
Pairwise (i = 2)
1
0.989
0.977
0.966
0.955
1
2
3
4
Joint occupancy, J{i}
Number of species, i

File: 54.csv

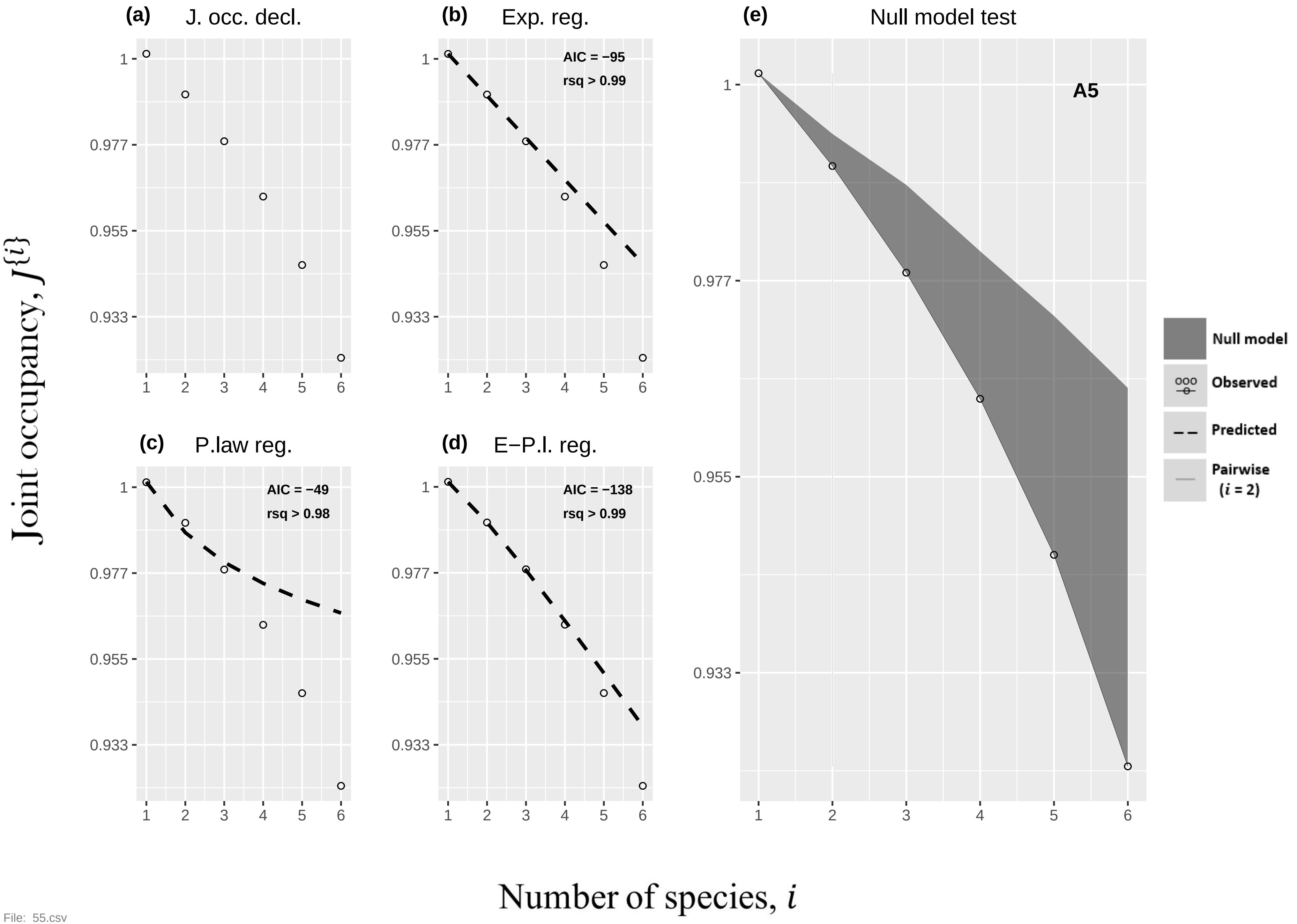
(a) J. occ. decl.
(b) Exp. reg.
AIC = −95
rsq > 0.99
(c) P.law reg.
AIC = −49
rsq > 0.98
(d) E−P.l. reg.
AIC = −138
rsq > 0.99
(e) Null model test
A5
Null model
Observed
Predicted
Pairwise ($i$ = 2)
Joint occupancy, $J^{\{i\}}$
Number of species, $i$
1
0.977
0.955
0.933
1 2 3 4 5 6

File: 55.csv

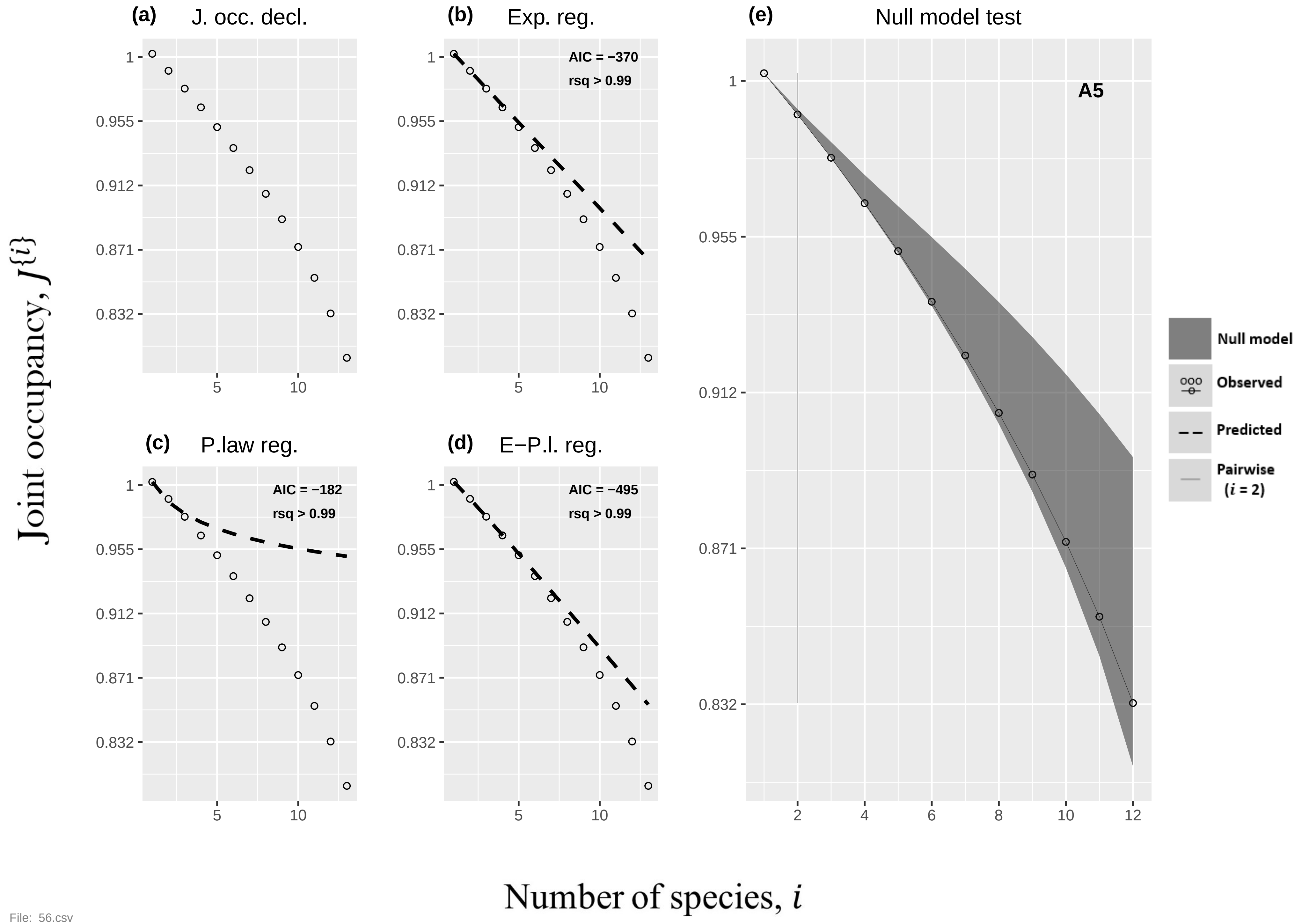
(a) J. occ. decl.
(b) Exp. reg.
AIC = −370
rsq > 0.99
(c) P.law reg.
AIC = −182
rsq > 0.99
(d) E−P.l. reg.
AIC = −495
rsq > 0.99
(e) Null model test
A5
1
0.955
0.912
0.871
0.832
5
10
2
4
6
8
12
Null model
Observed
Predicted
Pairwise
($i$ = 2)
Joint occupancy, $J\{i\}$
Number of species, $i$

File: 56.csv

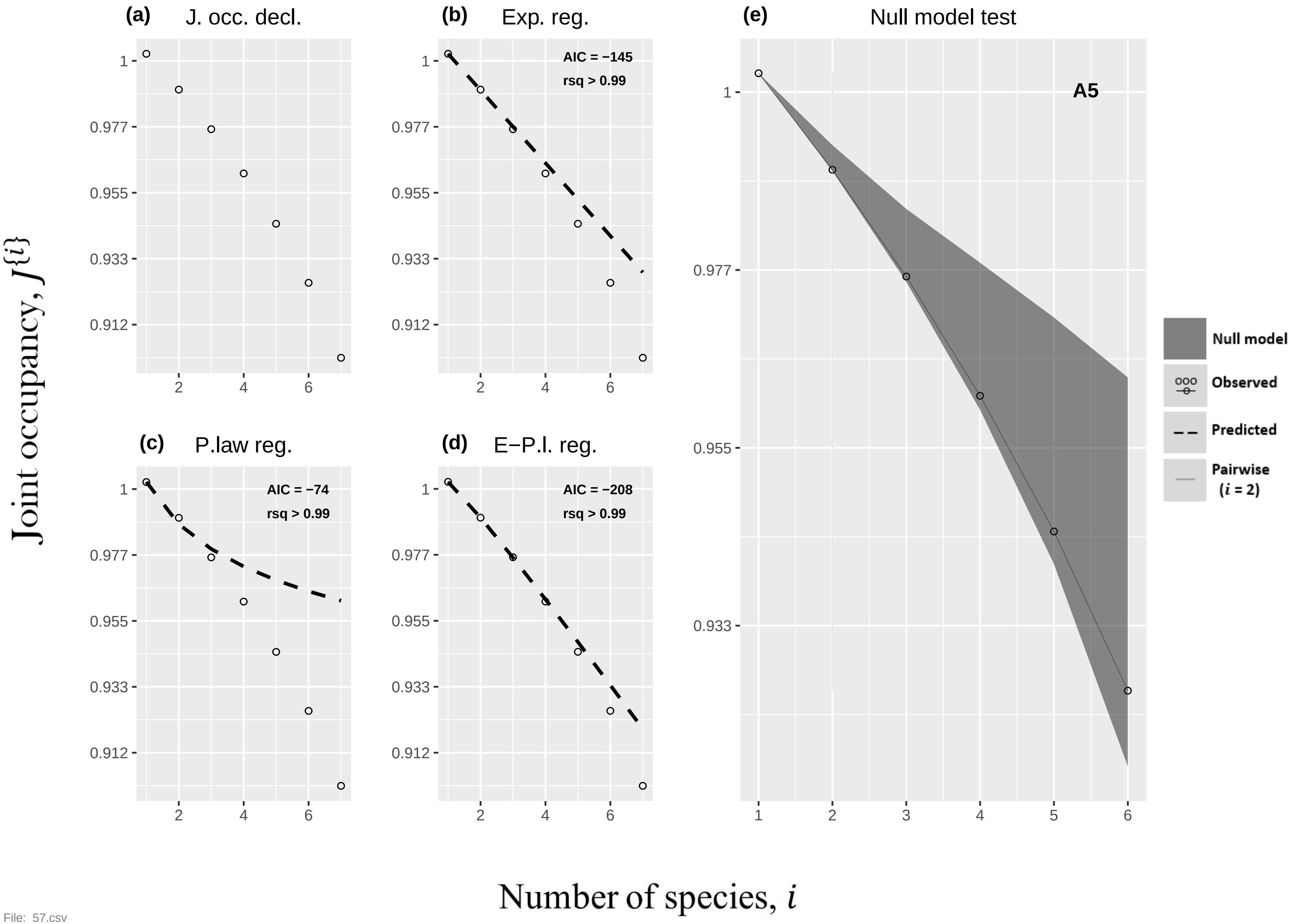
(a)
J. occ. decl.
(b)
Exp. reg.
AIC = −145
rsq > 0.99
(c)
P.law reg.
AIC = −74
rsq > 0.99
(d)
E−P.l. reg.
AIC = −208
rsq > 0.99
(e)
Null model test
A5
Null model
Observed
Predicted
Pairwise
(i = 2)
1
0.977
0.955
0.933
0.912
2
4
6
3
5
Joint occupancy, J{i}
Number of species, i

File: 57.csv

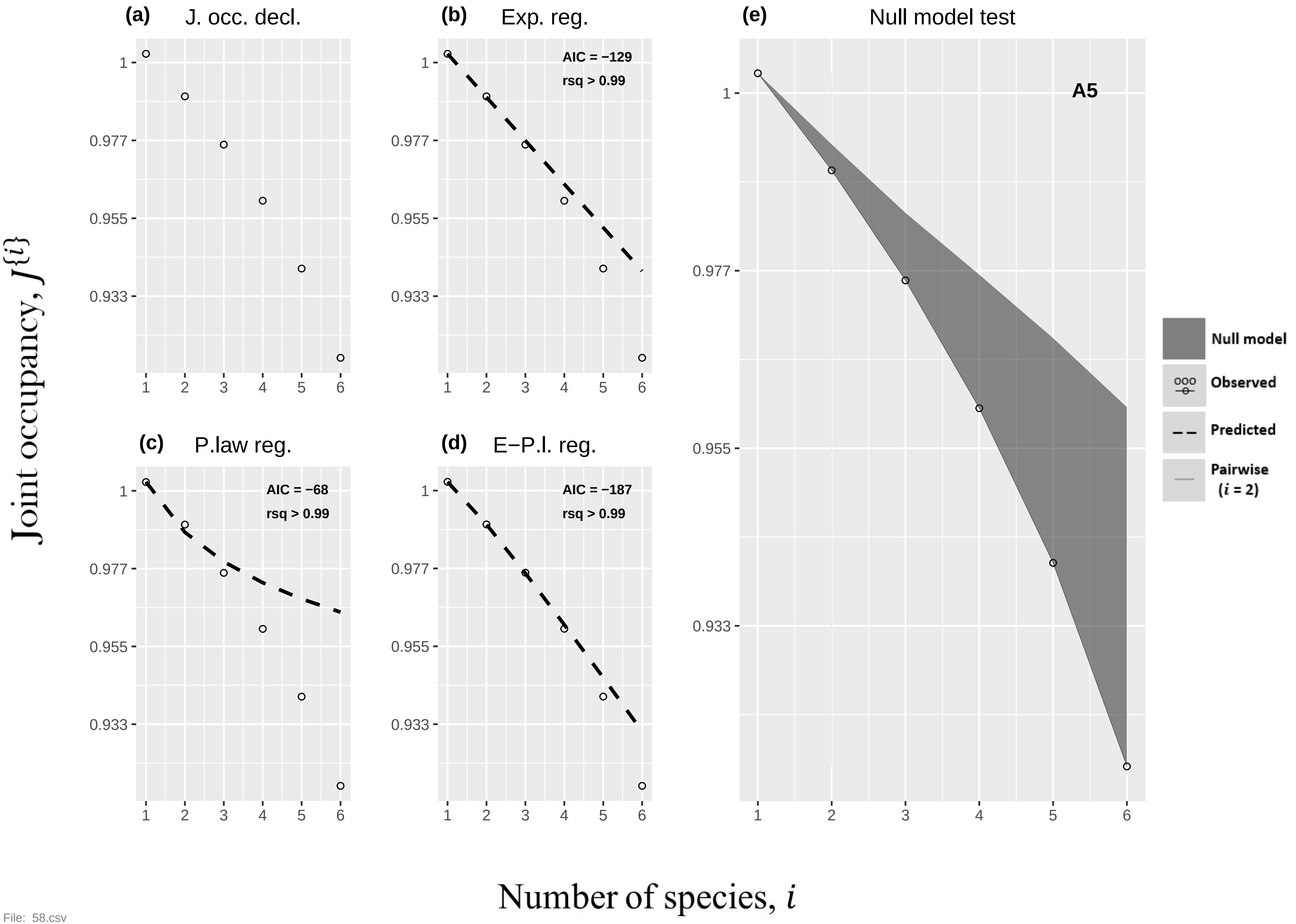

(a) J. occ. decl.
(b) Exp. reg.
AIC = −129
rsq > 0.99
(e) Null model test
A5
(c) P.law reg.
AIC = −68
rsq > 0.99
(d) E−P.l. reg.
AIC = −187
rsq > 0.99
Null model
Observed
Predicted
Pairwise (i = 2)
Joint occupancy, J{i}
Number of species, i


File: 58.csv

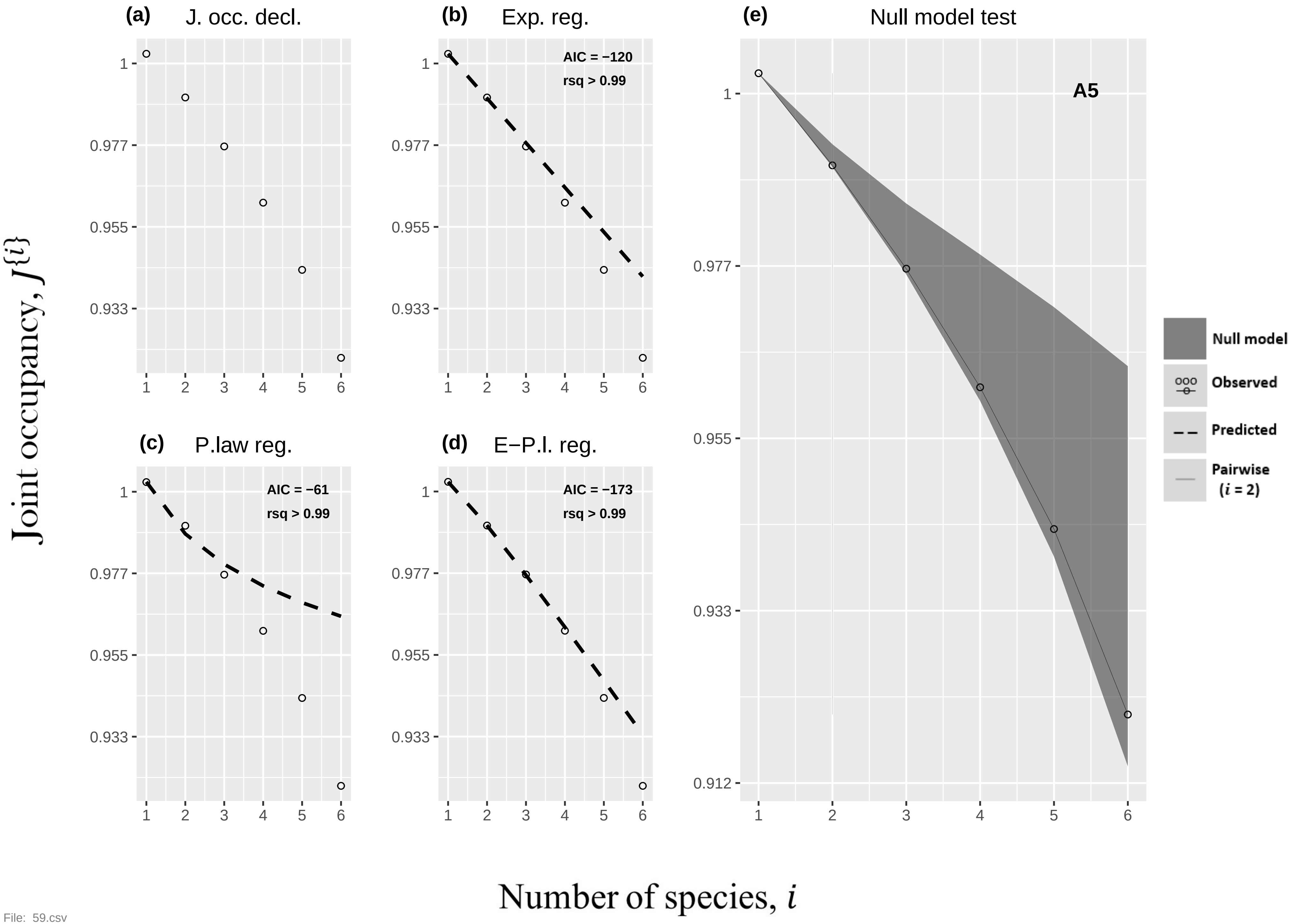

(a)
J. occ. decl.
(b)
Exp. reg.
AIC = −120
rsq > 0.99
(e)
Null model test
A5
(c)
P.law reg.
AIC = −61
rsq > 0.99
(d)
E−P.l. reg.
AIC = −173
rsq > 0.99
Null model
Observed
Predicted
Pairwise
(i = 2)
Joint occupancy, J{i}
Number of species, i


File: 59.csv

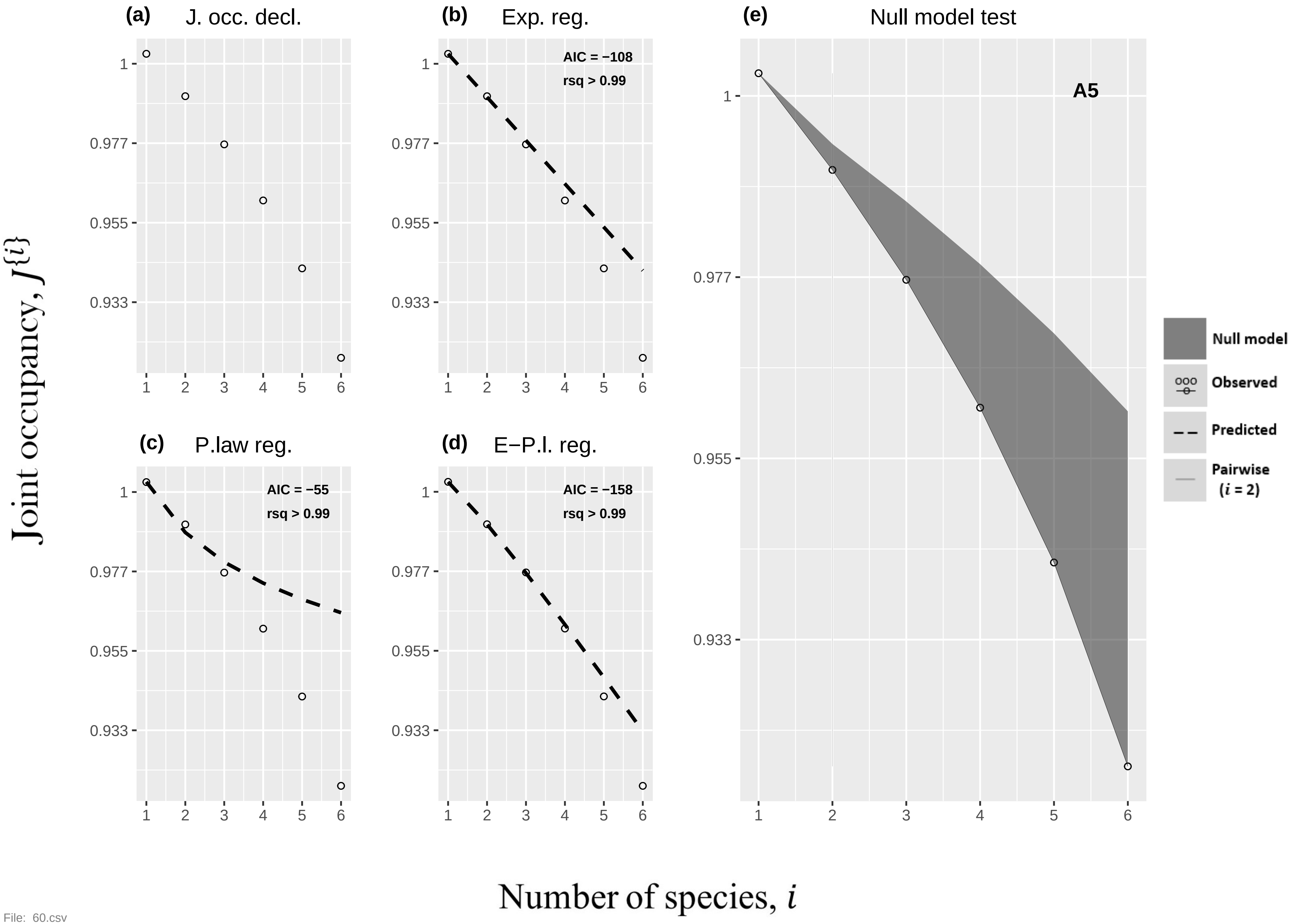

(a) J. occ. decl.
(b) Exp. reg.
AIC = −108
rsq > 0.99
(c) P.law reg.
AIC = −55
rsq > 0.99
(d) E−P.l. reg.
AIC = −158
rsq > 0.99
(e) Null model test
A5
Null model
Observed
Predicted
Pairwise (i = 2)
1
0.977
0.955
0.933
1 2 3 4 5 6
Joint occupancy, J{i}
Number of species, i


File: 60.csv

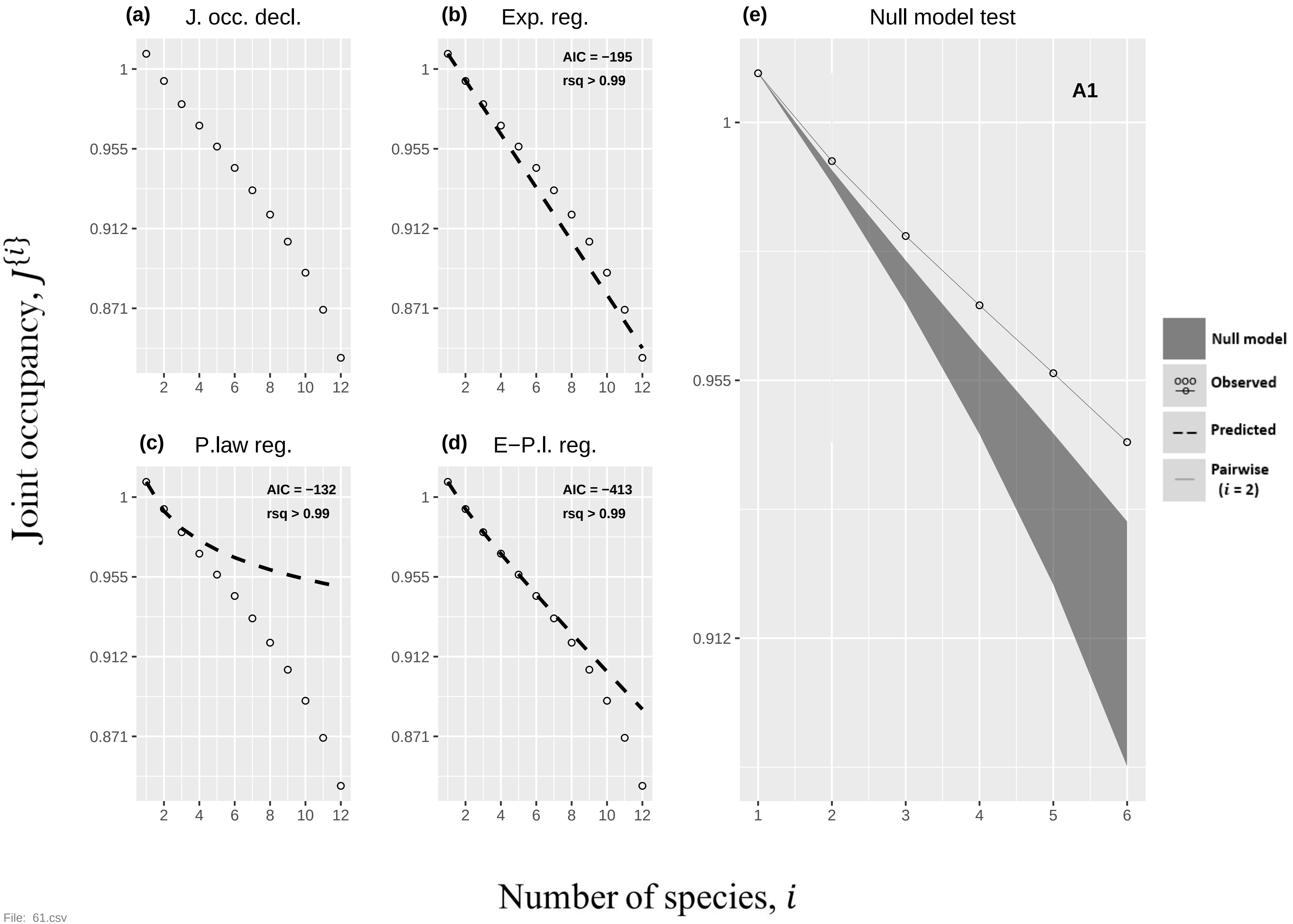

(a) J. occ. decl.
(b) Exp. reg.
AIC = −195
rsq > 0.99
(c) P.law reg.
AIC = −132
rsq > 0.99
(d) E−P.l. reg.
AIC = −413
rsq > 0.99
(e) Null model test
A1
Null model
Observed
Predicted
Pairwise (i = 2)
1
0.955
0.912
0.871
2 4 6 8 10 12
1 2 3 4 5 6
Joint occupancy, J{i}
Number of species, i


File: 61.csv

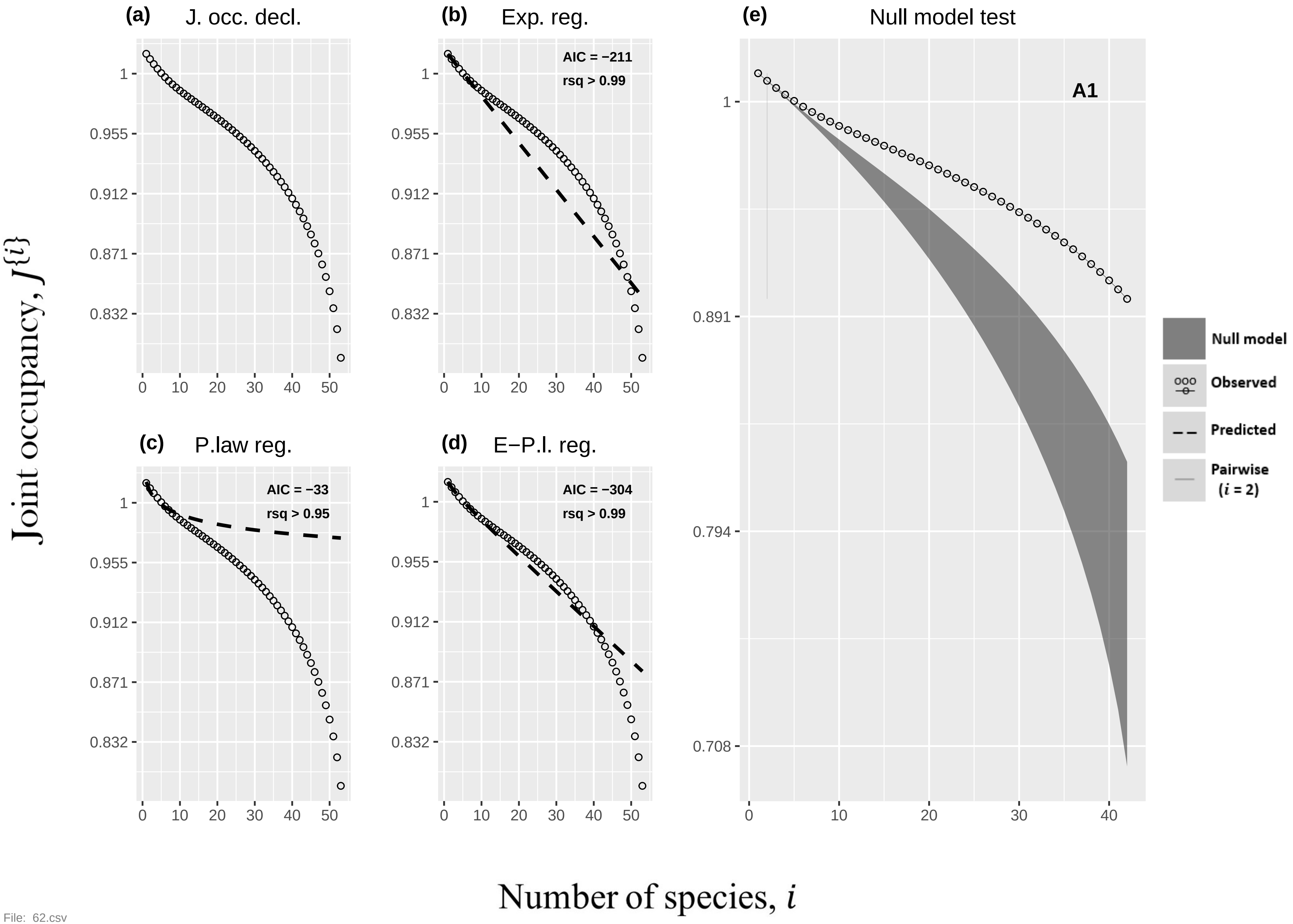

(a)
J. occ. decl.
(b)
Exp. reg.
AIC = −211
rsq > 0.99
(c)
P.law reg.
AIC = −33
rsq > 0.95
(d)
E−P.l. reg.
AIC = −304
rsq > 0.99
(e)
Null model test
A1
1
0.955
0.912
0.871
0.832
0.891
0.794
0.708
0
10
20
30
40
50
Null model
Observed
Predicted
Pairwise (i = 2)
Joint occupancy, J{i}
Number of species, i


File: 62.csv

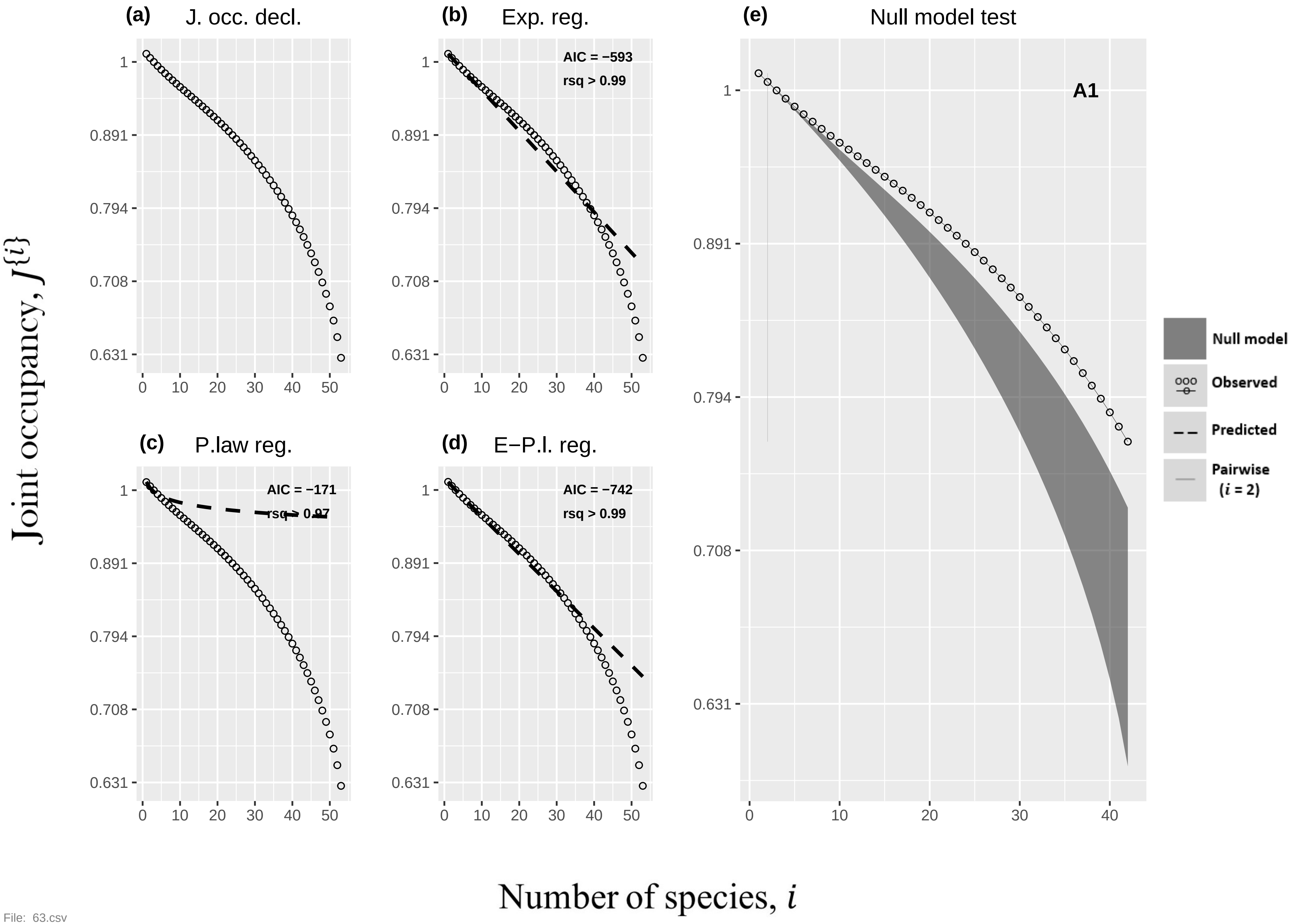
(a) J. occ. decl.
(b) Exp. reg.
AIC = −593
rsq > 0.99
(c) P.law reg.
AIC = −171
rsq > 0.97
(d) E−P.l. reg.
AIC = −742
rsq > 0.99
(e) Null model test
A1
Null model
Observed
Predicted
Pairwise (i = 2)
Joint occupancy, J{i}
Number of species, i
1
0.891
0.794
0.708
0.631
0
10
20
30
40
50

File: 63.csv

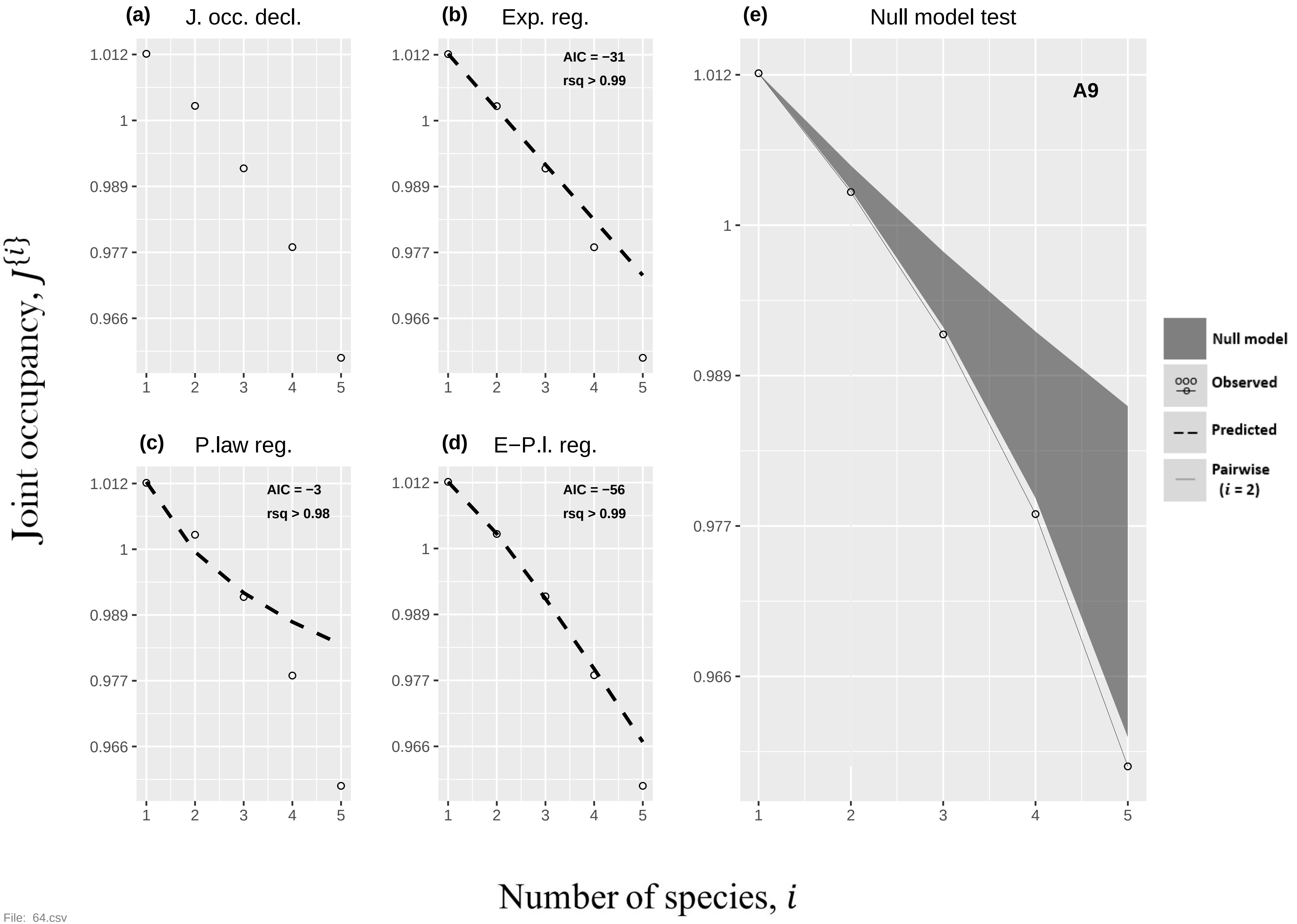
(a)
J. occ. decl.
(b)
Exp. reg.
AIC = −31
rsq > 0.99
(c)
P.law reg.
AIC = −3
rsq > 0.98
(d)
E−P.l. reg.
AIC = −56
rsq > 0.99
(e)
Null model test
A9
1.012
1
0.989
0.977
0.966
1
2
3
4
5
Null model
Observed
Predicted
Pairwise
(i = 2)
Joint occupancy, J{i}
Number of species, i

File: 64.csv

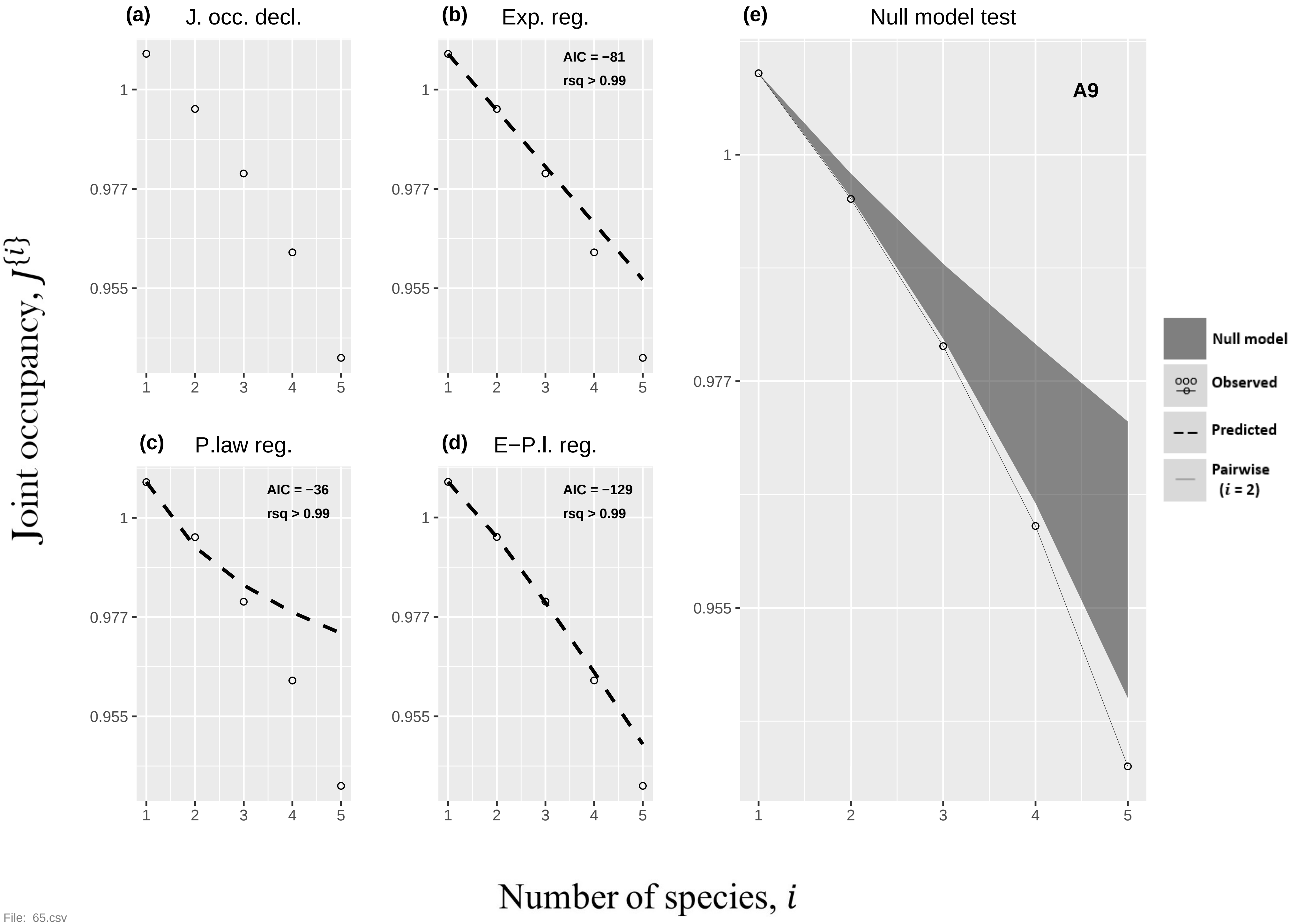

(a) J. occ. decl.
(b) Exp. reg.
AIC = −81
rsq > 0.99
(c) P.law reg.
AIC = −36
rsq > 0.99
(d) E−P.l. reg.
AIC = −129
rsq > 0.99
(e) Null model test
A9
Null model
Observed
Predicted
Pairwise (i = 2)
Joint occupancy, J{i}
Number of species, i


File: 65.csv

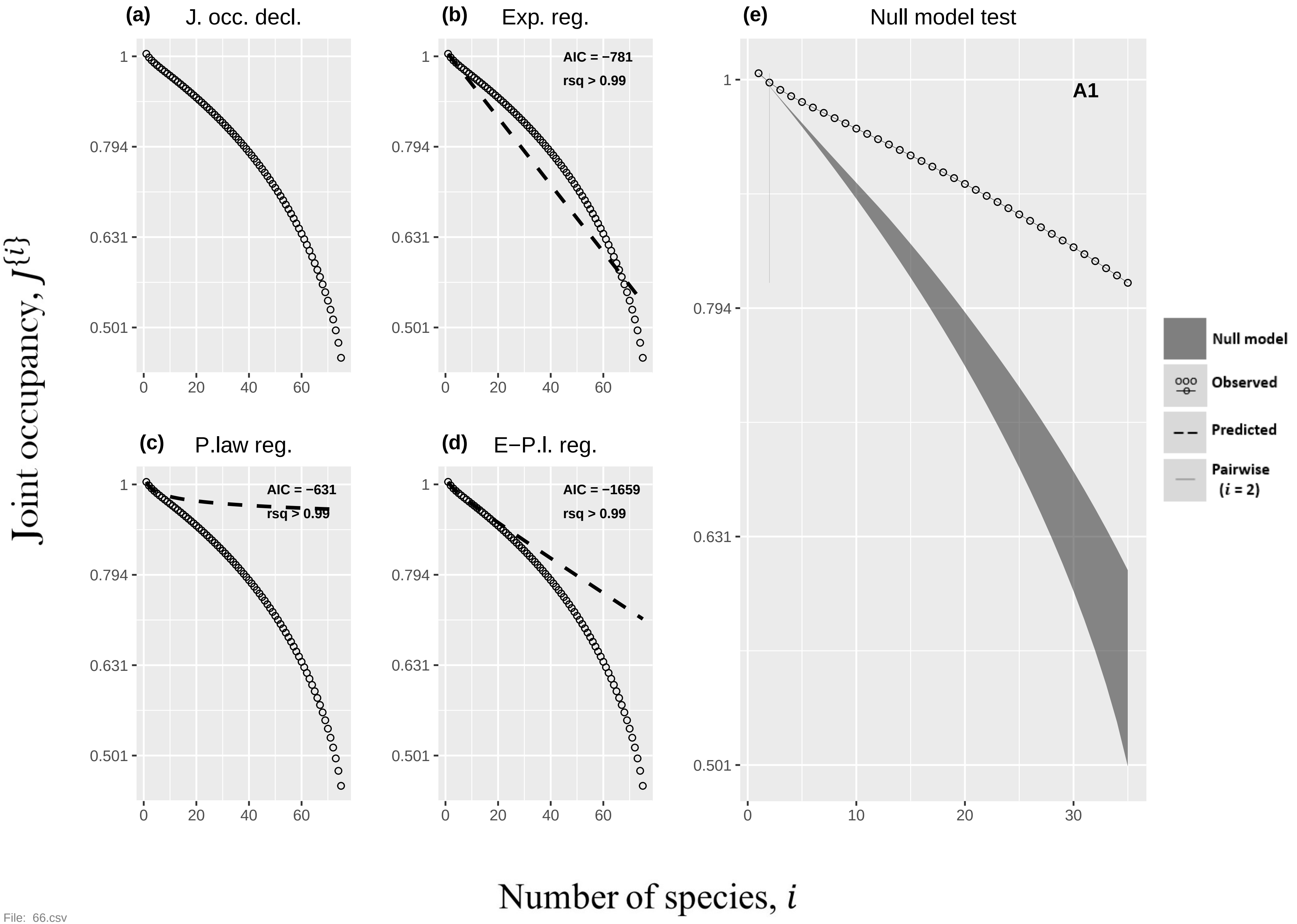
(a) J. occ. decl.
(b) Exp. reg.
AIC = −781
rsq > 0.99
(c) P.law reg.
AIC = −631
rsq > 0.99
(d) E−P.l. reg.
AIC = −1659
rsq > 0.99
(e) Null model test
A1
Null model
Observed
Predicted
Pairwise
(i = 2)
Joint occupancy, J{i}
Number of species, i
1
0.794
0.631
0.501
0
20
40
60
10
30

File: 66.csv

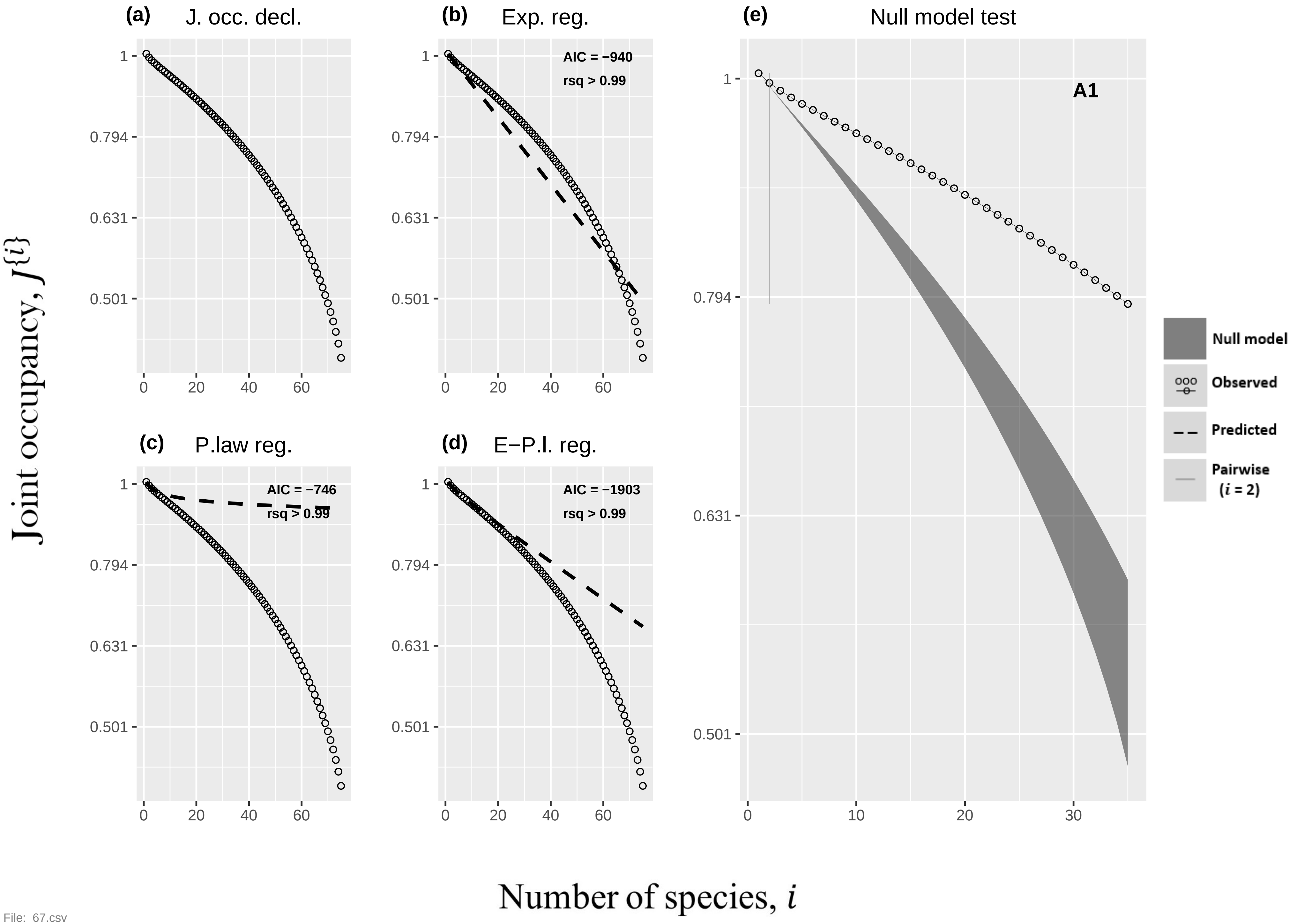

(a) J. occ. decl.
(b) Exp. reg.
AIC = −940
rsq > 0.99
(c) P.law reg.
AIC = −746
rsq > 0.99
(d) E−P.l. reg.
AIC = −1903
rsq > 0.99
(e) Null model test
A1
Null model
Observed
Predicted
Pairwise (i = 2)
Joint occupancy, J{i}
Number of species, i
1
0.794
0.631
0.501
0
20
40
60
10
30


File: 67.csv

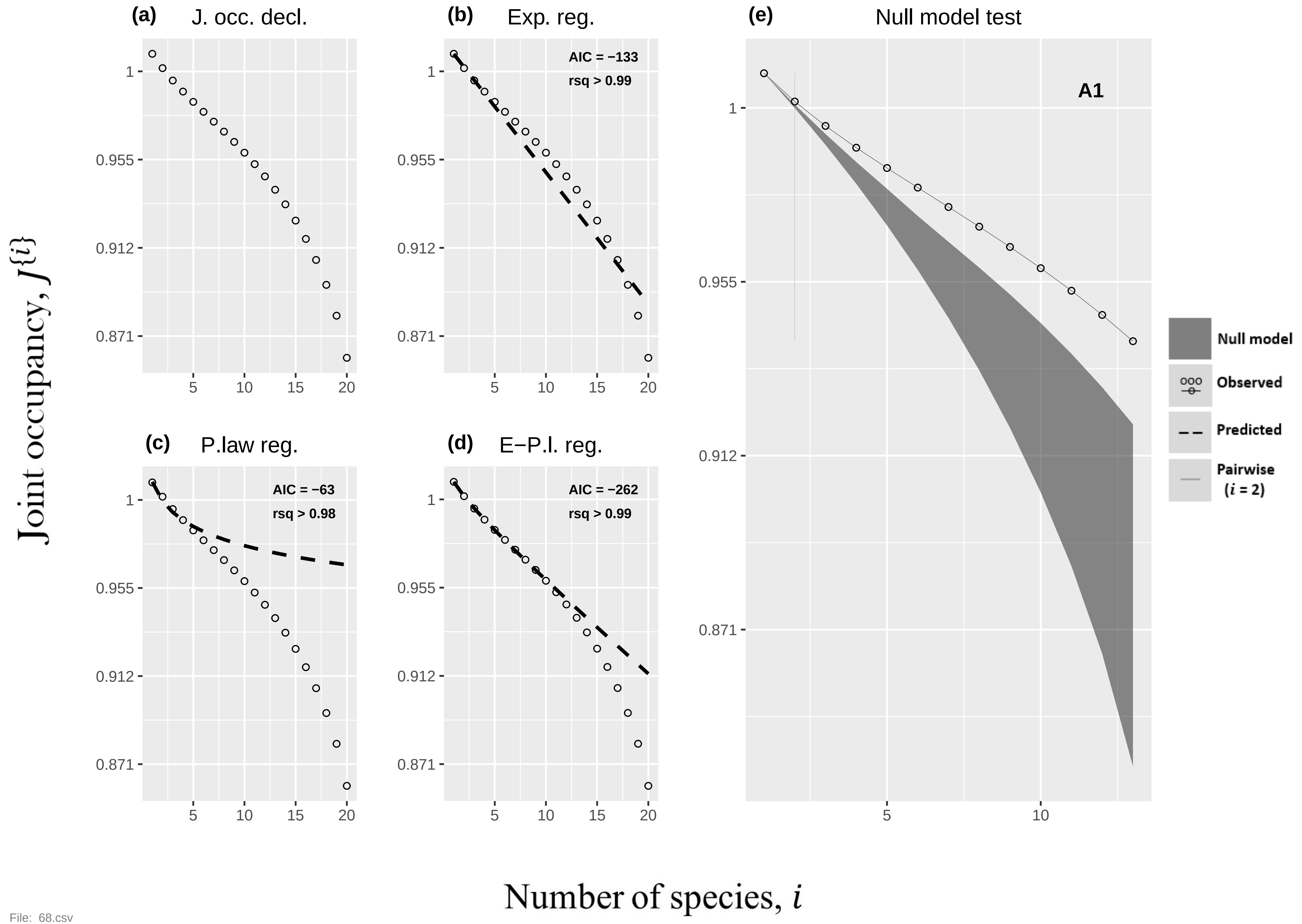

(a) J. occ. decl.
(b) Exp. reg.
AIC = −133
rsq > 0.99
(c) P.law reg.
AIC = −63
rsq > 0.98
(d) E−P.l. reg.
AIC = −262
rsq > 0.99
(e) Null model test
A1
Null model
Observed
Predicted
Pairwise (i = 2)
1
0.955
0.912
0.871
5
10
15
20
Joint occupancy, J{i}
Number of species, i


File: 68.csv

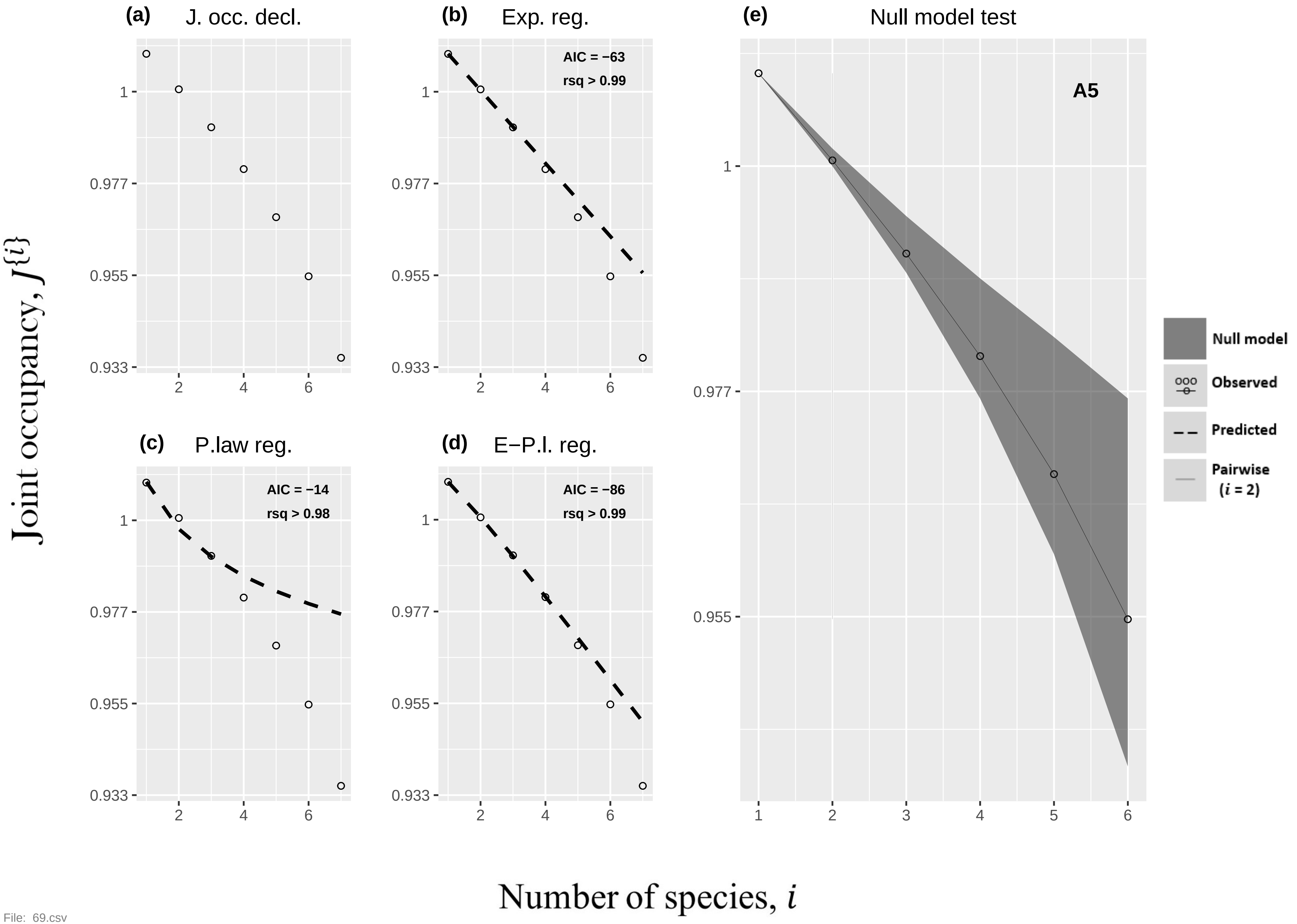

(a) J. occ. decl.
(b) Exp. reg.
AIC = −63
rsq > 0.99
(c) P.law reg.
AIC = −14
rsq > 0.98
(d) E−P.l. reg.
AIC = −86
rsq > 0.99
(e) Null model test
A5
Null model
Observed
Predicted
Pairwise
(i = 2)
1
0.977
0.955
0.933
2
4
6
1
2
3
4
5
6
Joint occupancy, J{i}
Number of species, i


File: 69.csv

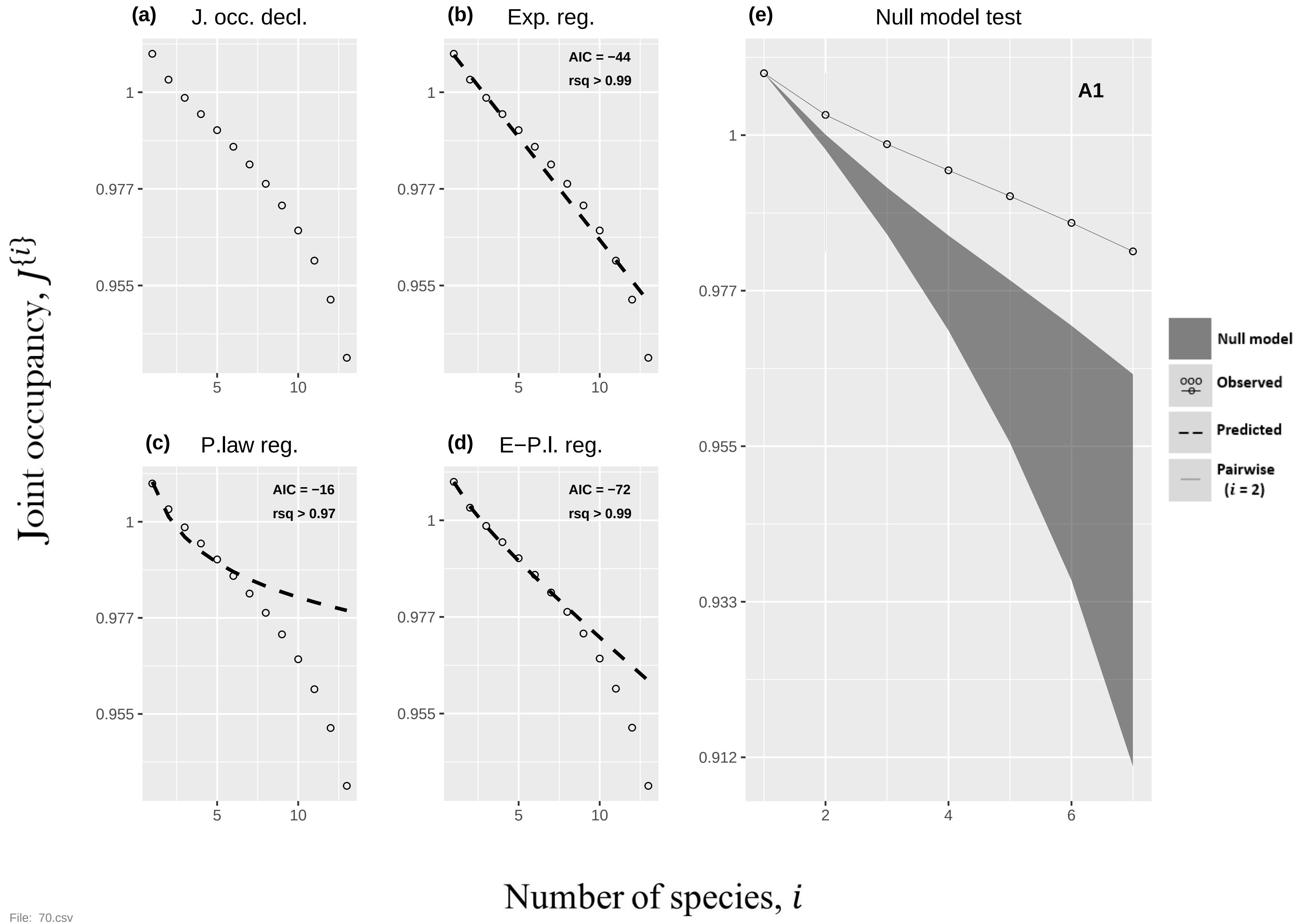

(a) J. occ. decl.
(b) Exp. reg.
AIC = −44
rsq > 0.99
(c) P.law reg.
AIC = −16
rsq > 0.97
(d) E−P.l. reg.
AIC = −72
rsq > 0.99
(e) Null model test
A1
Null model
Observed
Predicted
Pairwise (i = 2)
1
0.977
0.955
0.933
0.912
5
10
2
4
6
Joint occupancy, J{i}
Number of species, i


File: 70.csv

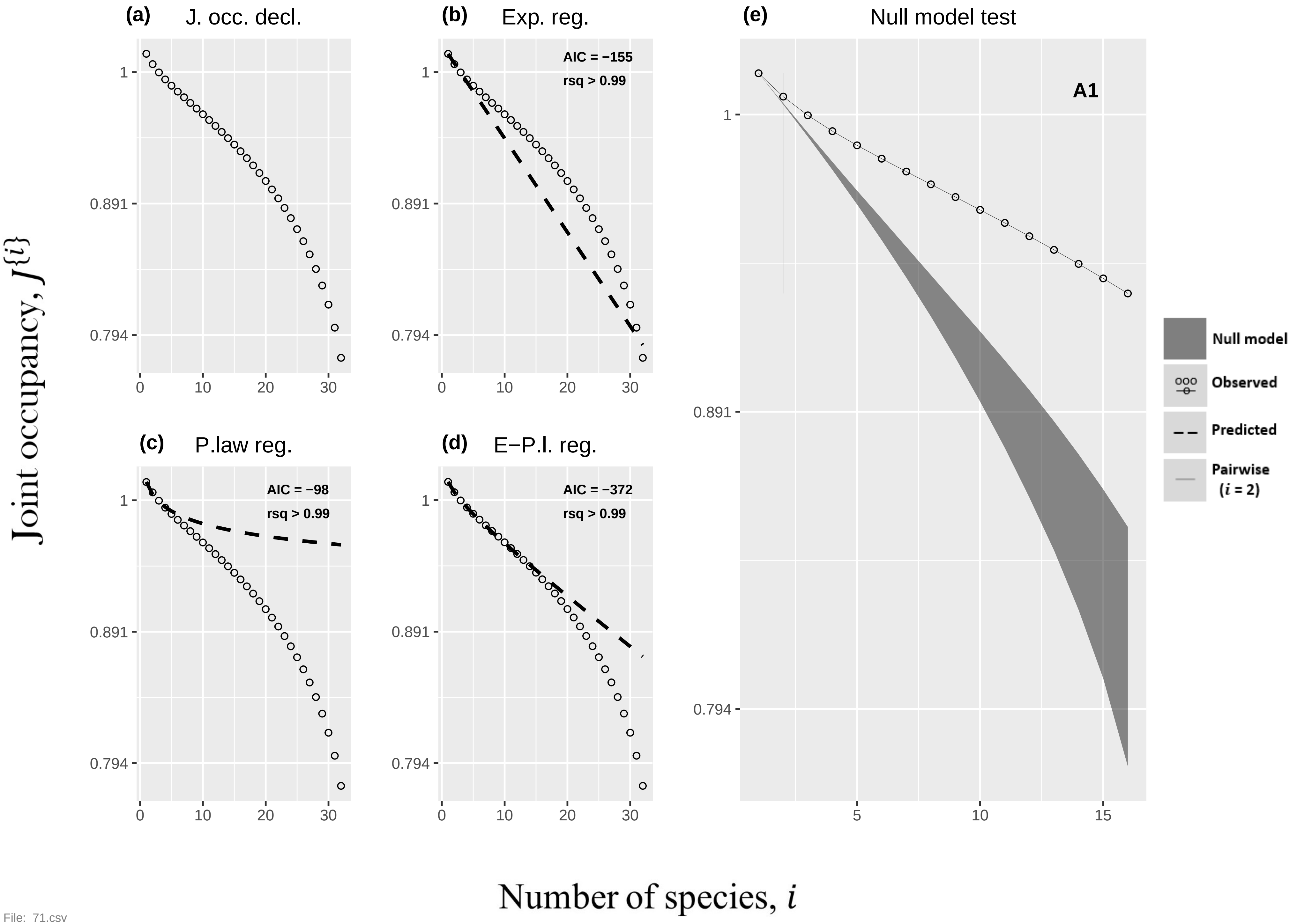

(a) J. occ. decl.
(b) Exp. reg.
AIC = −155
rsq > 0.99
(c) P.law reg.
AIC = −98
rsq > 0.99
(d) E−P.l. reg.
AIC = −372
rsq > 0.99
(e) Null model test
A1
Null model
Observed
Predicted
Pairwise (i = 2)
1
0.891
0.794
0
10
20
30
5
15
Joint occupancy, J{i}
Number of species, i

File: 71.csv

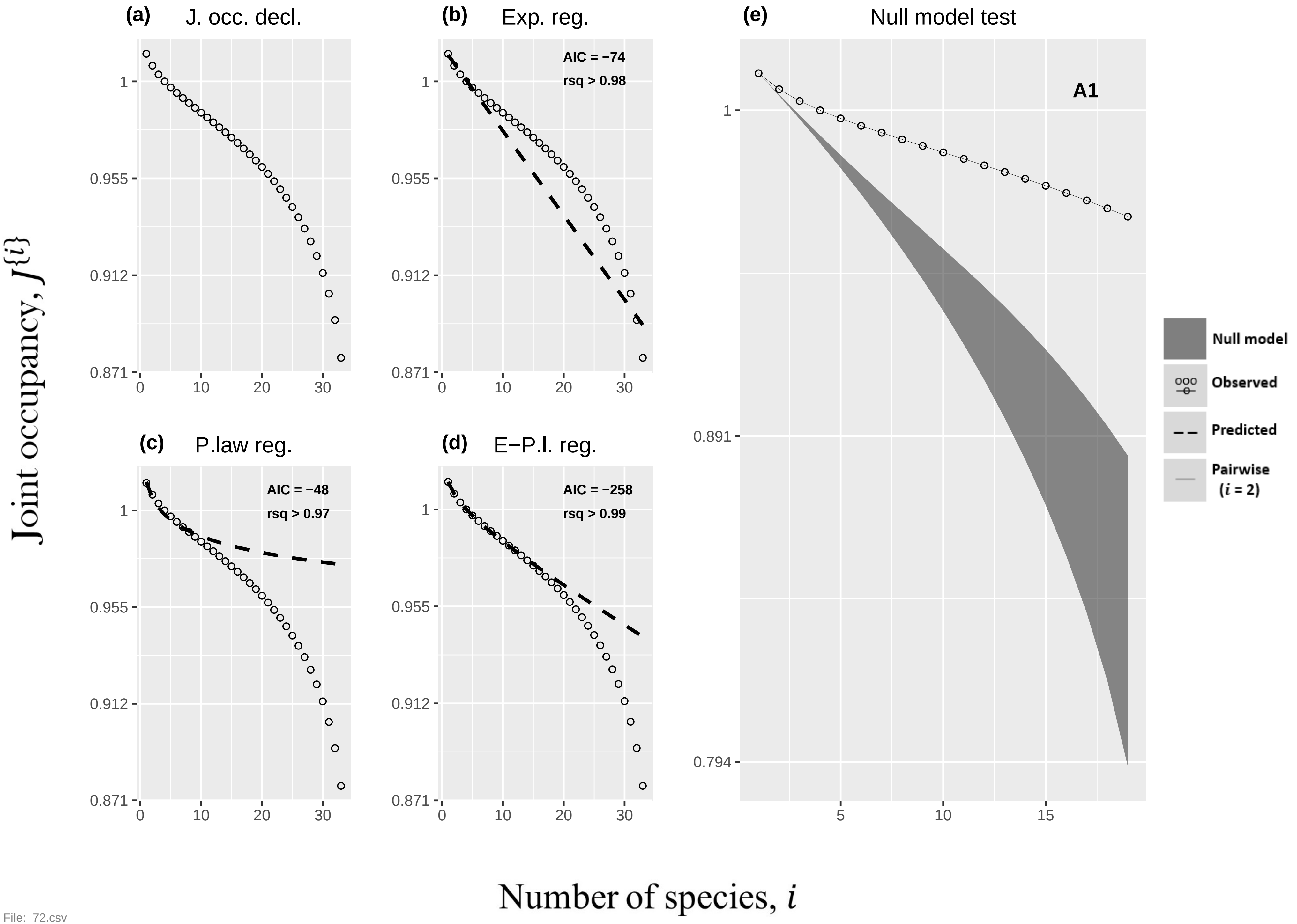

(a)
J. occ. decl.
(b)
Exp. reg.
AIC = −74
rsq > 0.98
(c)
P.law reg.
AIC = −48
rsq > 0.97
(d)
E−P.l. reg.
AIC = −258
rsq > 0.99
(e)
Null model test
A1
Null model
Observed
Predicted
Pairwise (i = 2)
1
0.955
0.912
0.871
0.891
0.794
0
10
20
30
5
15
Joint occupancy, J{i}
Number of species, i


File: 72.csv

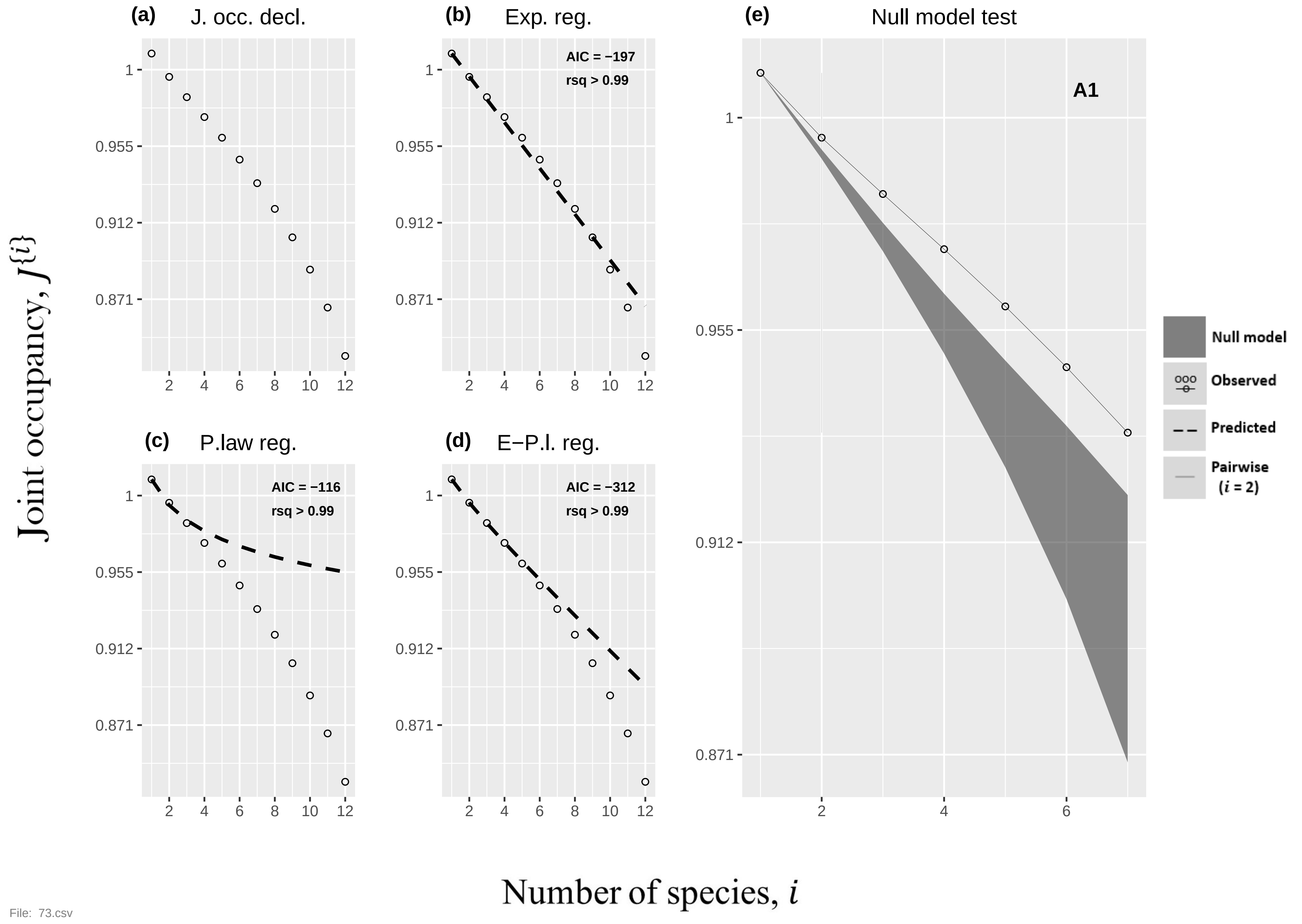
(a) J. occ. decl.
(b) Exp. reg.
AIC = −197
rsq > 0.99
(c) P.law reg.
AIC = −116
rsq > 0.99
(d) E−P.l. reg.
AIC = −312
rsq > 0.99
(e) Null model test
A1
Null model
Observed
Predicted
Pairwise (i = 2)
Joint occupancy, J{i}
Number of species, i
1
0.955
0.912
0.871
2
4
6
8
10
12
File: 73.csv

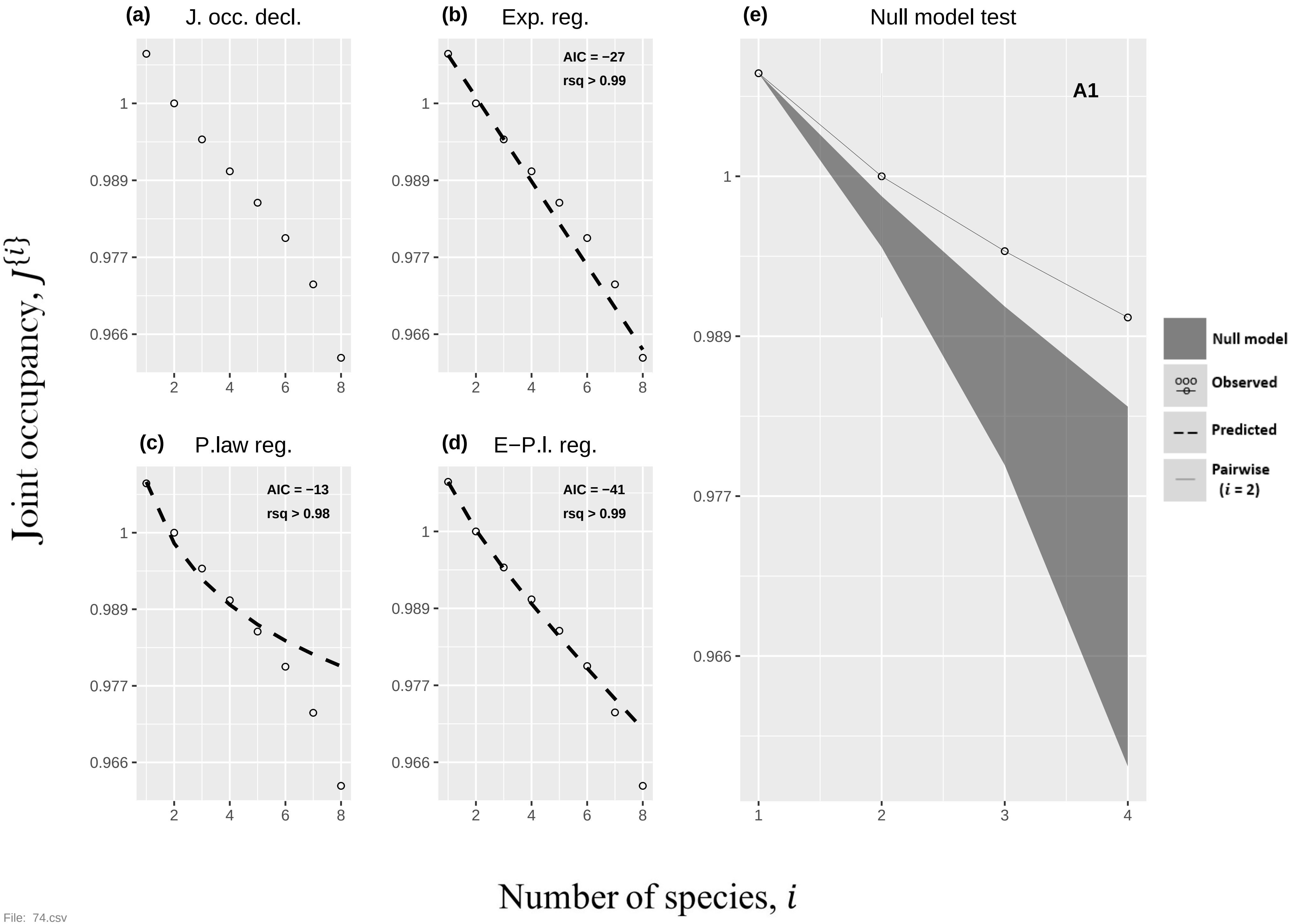
(a)
J. occ. decl.
(b)
Exp. reg.
AIC = −27
rsq > 0.99
(c)
P.law reg.
AIC = −13
rsq > 0.98
(d)
E−P.l. reg.
AIC = −41
rsq > 0.99
(e)
Null model test
A1
Null model
Observed
Predicted
Pairwise
(i = 2)
Joint occupancy, J{i}
Number of species, i
1
0.989
0.977
0.966
2
4
6
8
3

File: 74.csv

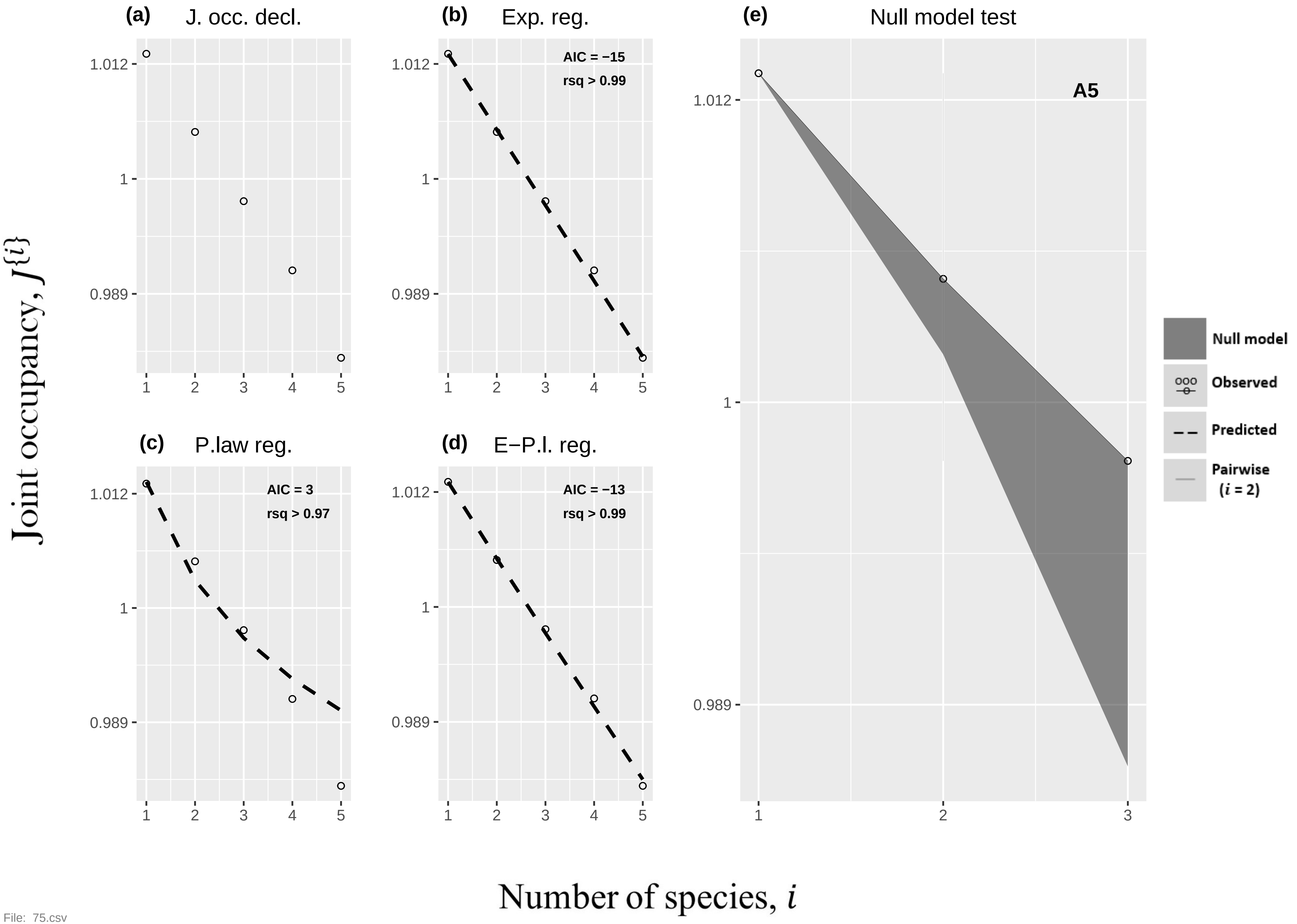
(a)
J. occ. decl.
(b)
Exp. reg.
AIC = −15
rsq > 0.99
(e)
Null model test
A5
(c)
P.law reg.
AIC = 3
rsq > 0.97
(d)
E−P.l. reg.
AIC = −13
rsq > 0.99
1.012
1
0.989
1
2
3
4
5
Null model
Observed
Predicted
Pairwise
(i = 2)
Joint occupancy, J{i}
Number of species, i

File: 75.csv

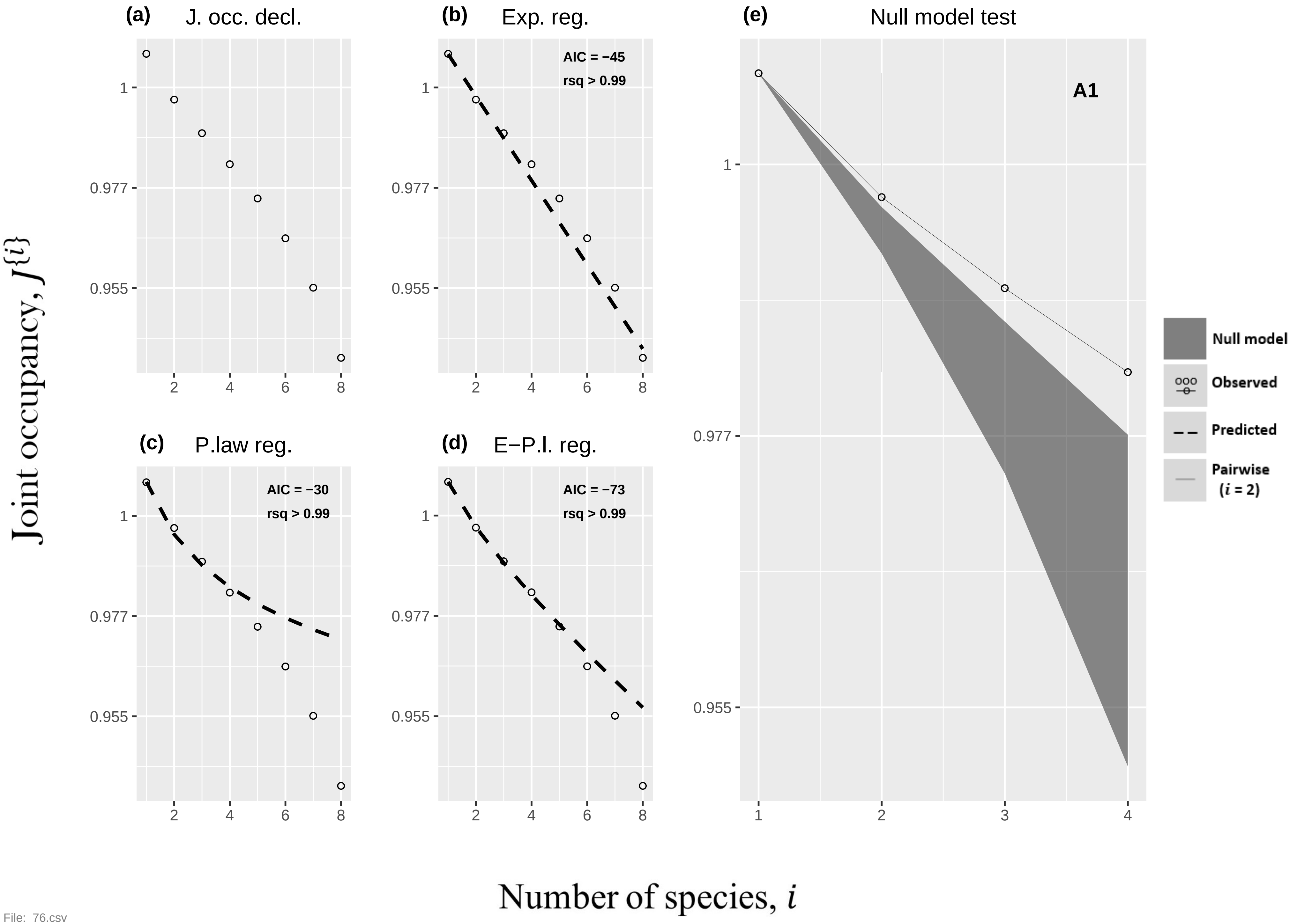

(a) J. occ. decl.
(b) Exp. reg.
AIC = −45
rsq > 0.99
(c) P.law reg.
AIC = −30
rsq > 0.99
(d) E−P.l. reg.
AIC = −73
rsq > 0.99
(e) Null model test
A1
Null model
Observed
Predicted
Pairwise (i = 2)
1
0.977
0.955
Joint occupancy, J{i}
Number of species, i


File: 76.csv

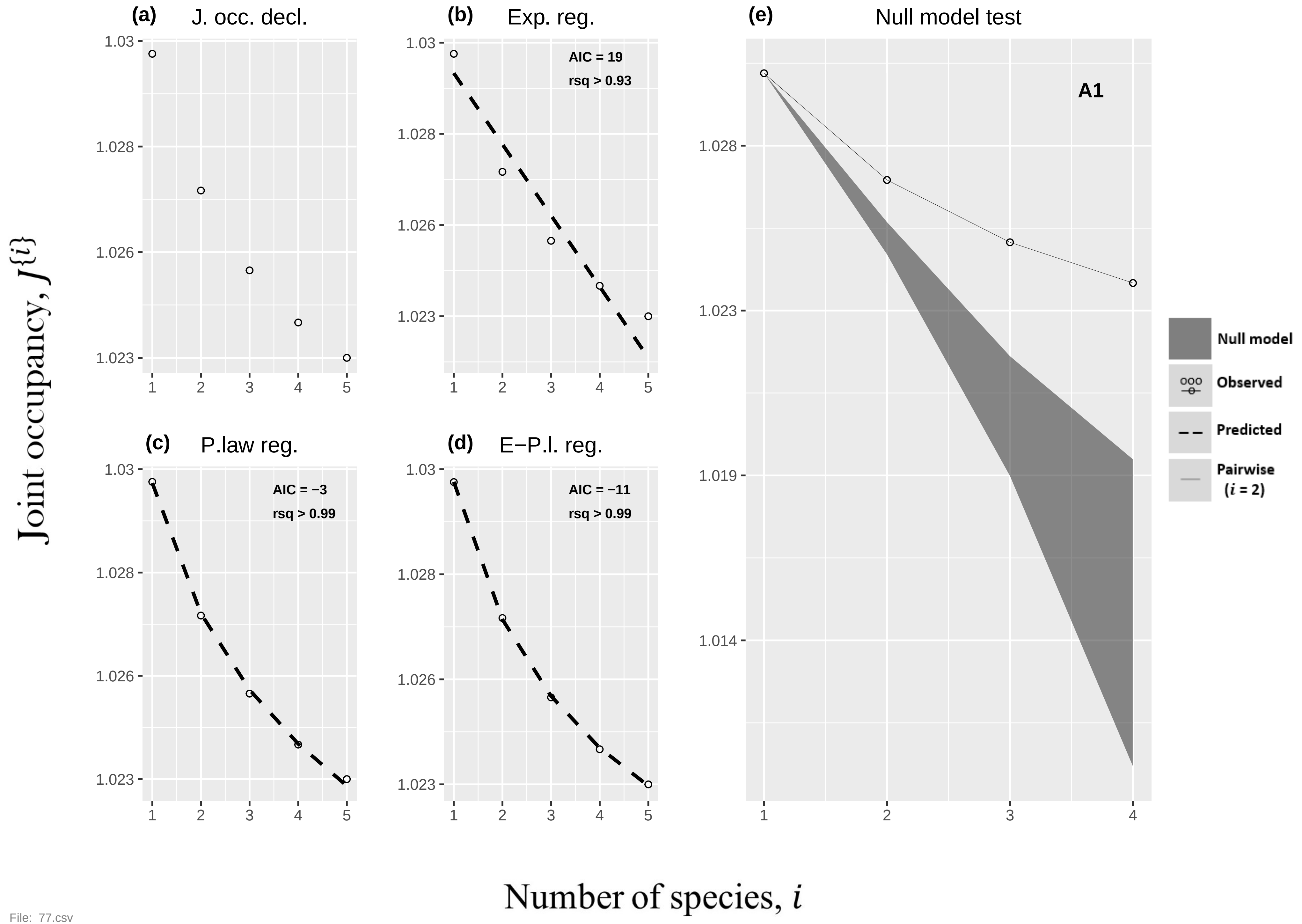

(a)
J. occ. decl.
(b)
Exp. reg.
AIC = 19
rsq > 0.93
(c)
P.law reg.
AIC = −3
rsq > 0.99
(d)
E−P.l. reg.
AIC = −11
rsq > 0.99
(e)
Null model test
A1
Null model
Observed
Predicted
Pairwise
(i = 2)
Joint occupancy, J{i}
Number of species, i


File: 77.csv

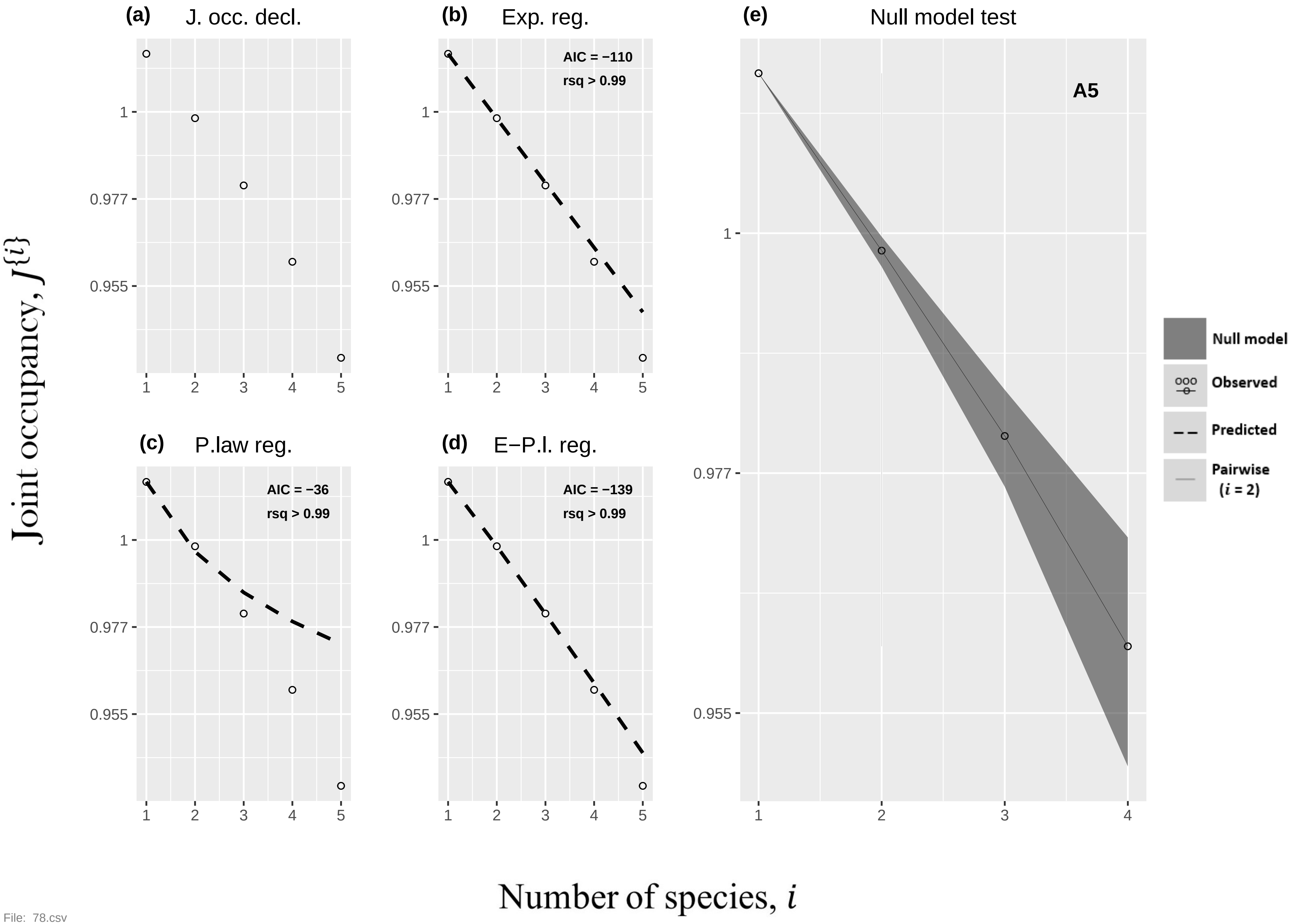

(a) J. occ. decl.
(b) Exp. reg.
AIC = −110
rsq > 0.99
(e) Null model test
A5
(c) P.law reg.
AIC = −36
rsq > 0.99
(d) E−P.l. reg.
AIC = −139
rsq > 0.99
Null model
Observed
Predicted
Pairwise ($i$ = 2)
1
0.977
0.955
1 2 3 4 5
1 2 3 4
Joint occupancy, $J^{\{i\}}$
Number of species, $i$


File: 78.csv

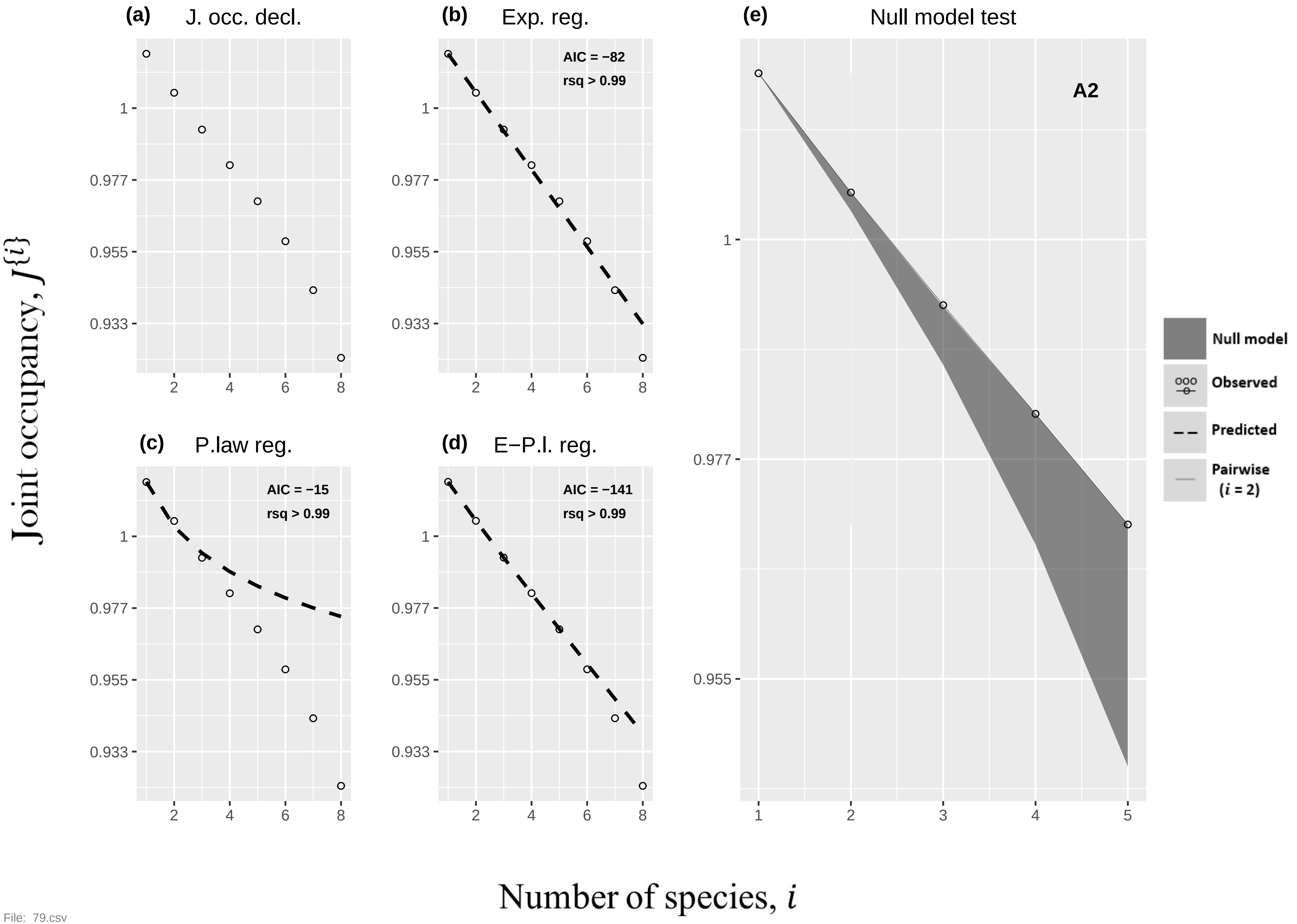
(a)
J. occ. decl.
(b)
Exp. reg.
AIC = −82
rsq > 0.99
(e)
Null model test
A2
(c)
P.law reg.
AIC = −15
rsq > 0.99
(d)
E−P.l. reg.
AIC = −141
rsq > 0.99
Null model
Observed
Predicted
Pairwise
(i = 2)
Joint occupancy, J{i}
Number of species, i
1
0.977
0.955
0.933
2
4
6
8
3
5

File: 79.csv

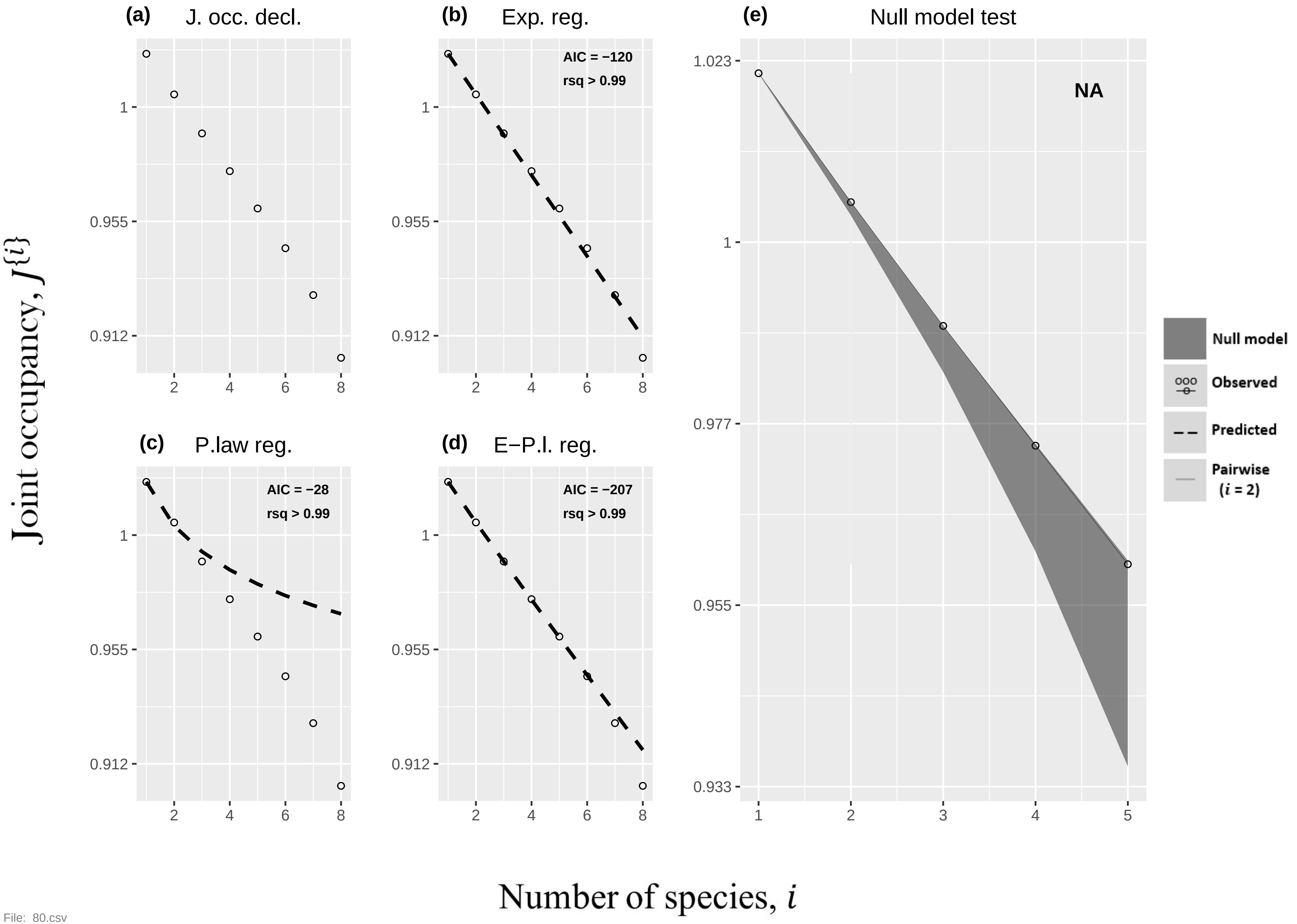

(a) J. occ. decl.
(b) Exp. reg.
AIC = −120
rsq > 0.99
(c) P.law reg.
AIC = −28
rsq > 0.99
(d) E−P.l. reg.
AIC = −207
rsq > 0.99
(e) Null model test
NA
1.023
1
0.977
0.955
0.933
0.955
0.912
1
2
3
4
5
6
8
Null model
Observed
Predicted
Pairwise
(i = 2)
Joint occupancy, J{i}
Number of species, i


File: 80.csv

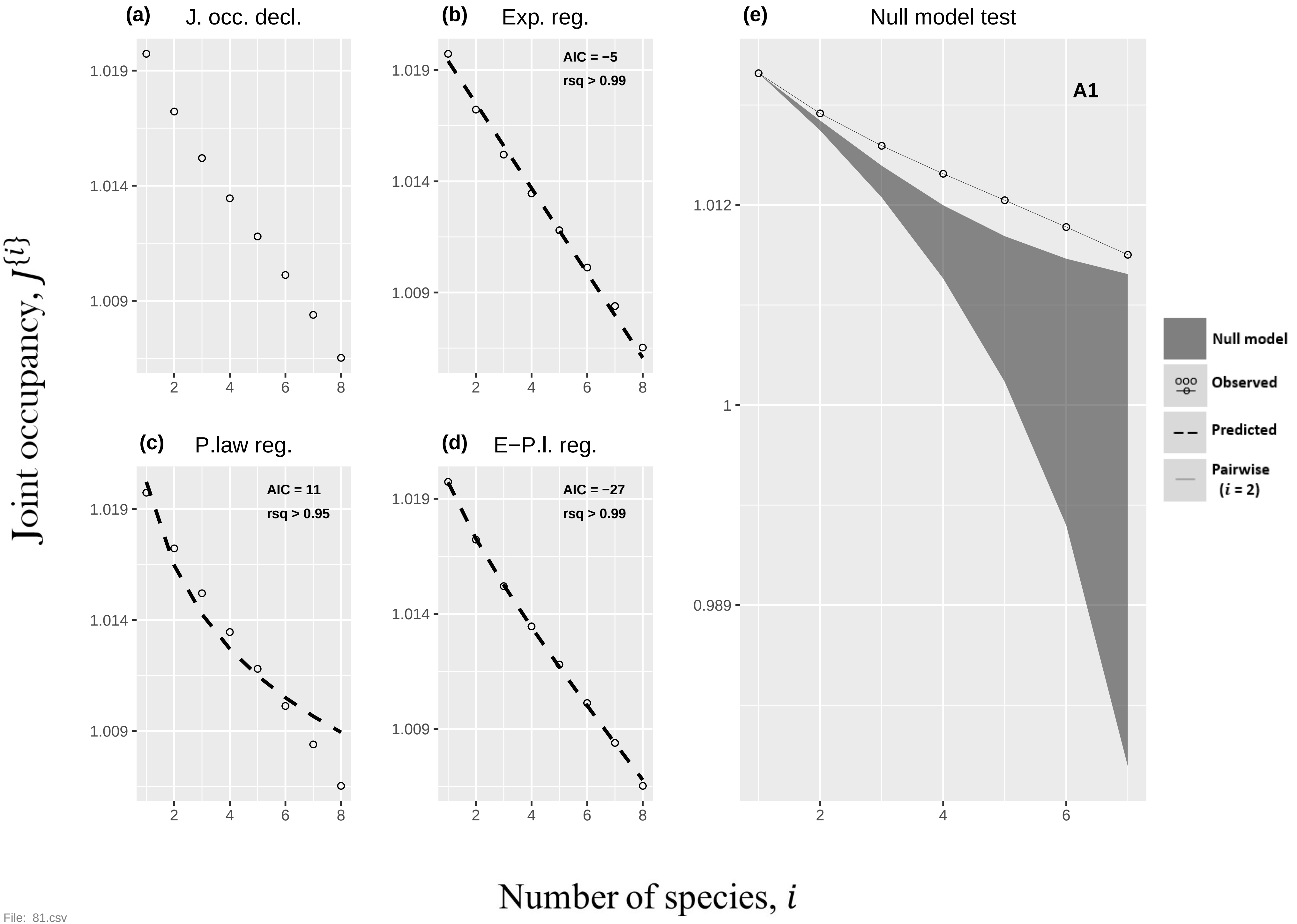
(a) J. occ. decl.
(b) Exp. reg.
AIC = −5
rsq > 0.99
(c) P.law reg.
AIC = 11
rsq > 0.95
(d) E−P.l. reg.
AIC = −27
rsq > 0.99
(e) Null model test
A1
Null model
Observed
Predicted
Pairwise (i = 2)
Joint occupancy, J{i}
Number of species, i
1.019
1.014
1.009
1.012
1
0.989
2
4
6
8
File: 81.csv

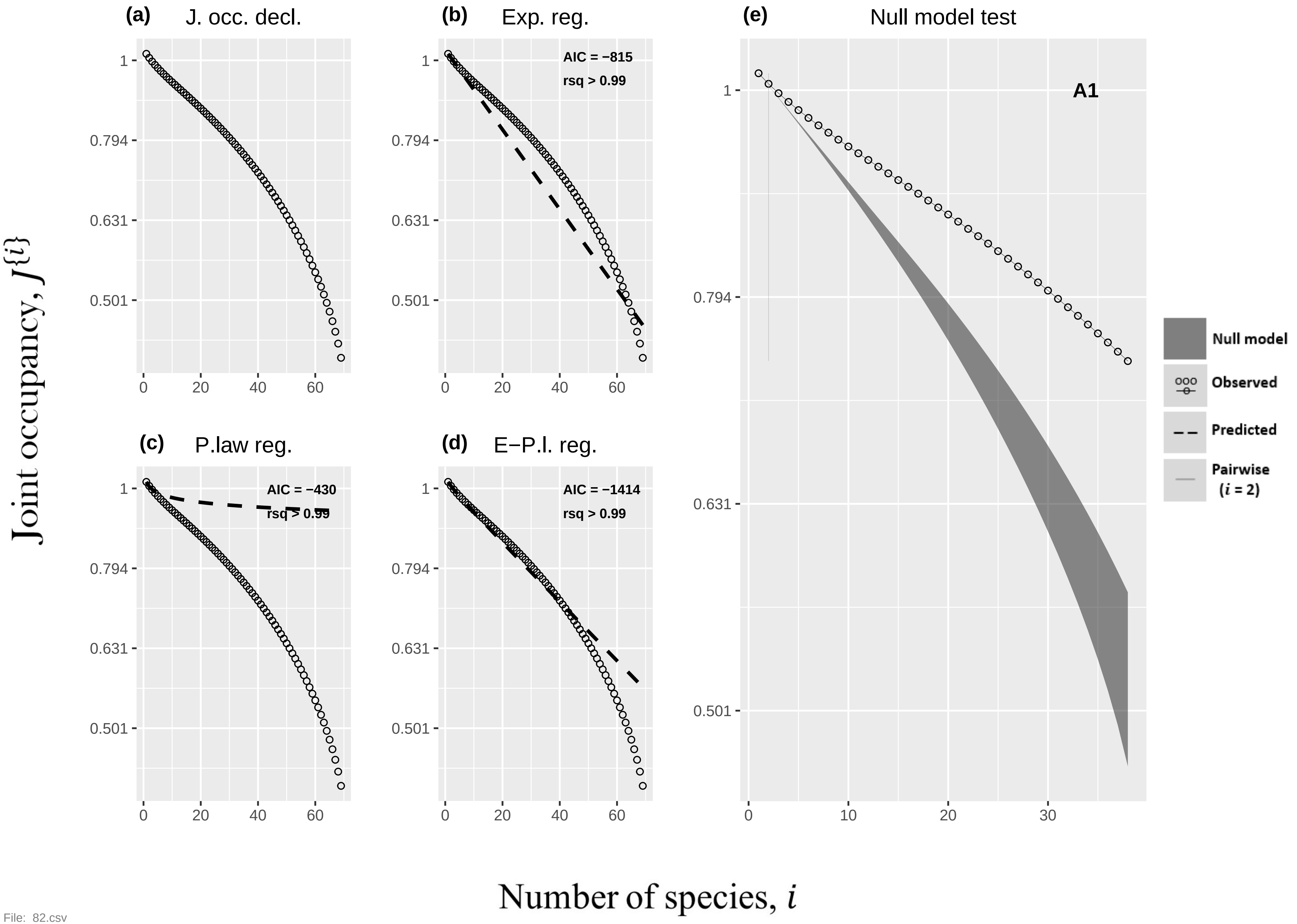

(a) J. occ. decl.
(b) Exp. reg.
AIC = −815
rsq > 0.99
(c) P.law reg.
AIC = −430
rsq > 0.99
(d) E−P.l. reg.
AIC = −1414
rsq > 0.99
(e) Null model test
A1
Null model
Observed
Predicted
Pairwise (i = 2)
Joint occupancy, J{i}
Number of species, i


File: 82.csv

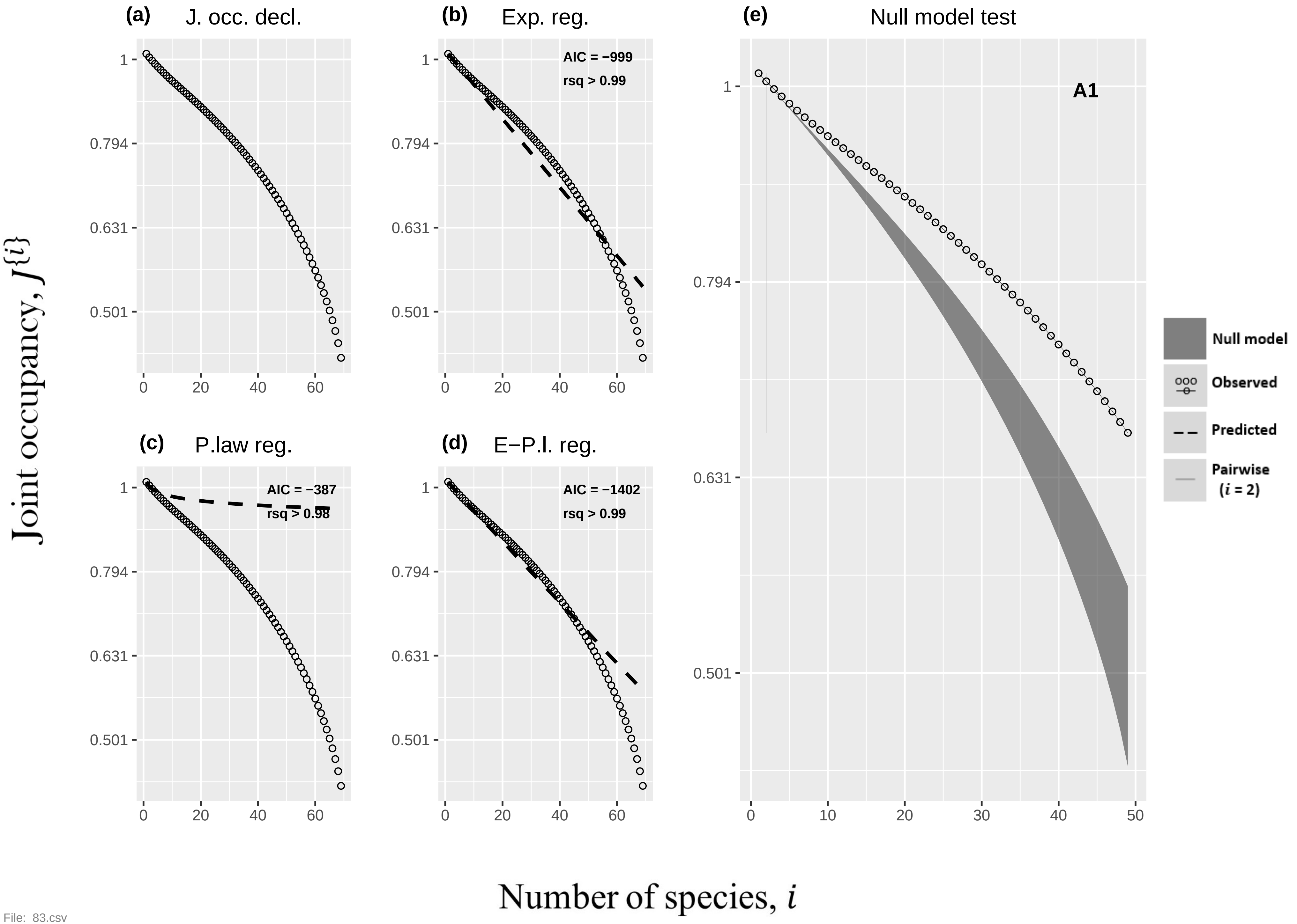
(a) J. occ. decl.
(b) Exp. reg.
AIC = −999
rsq > 0.99
(e) Null model test
A1
(c) P.law reg.
AIC = −387
rsq > 0.98
(d) E−P.l. reg.
AIC = −1402
rsq > 0.99
Null model
Observed
Predicted
Pairwise (i = 2)
Joint occupancy, J{i}
Number of species, i
1
0.794
0.631
0.501
0
20
40
60
10
30
50

File: 83.csv

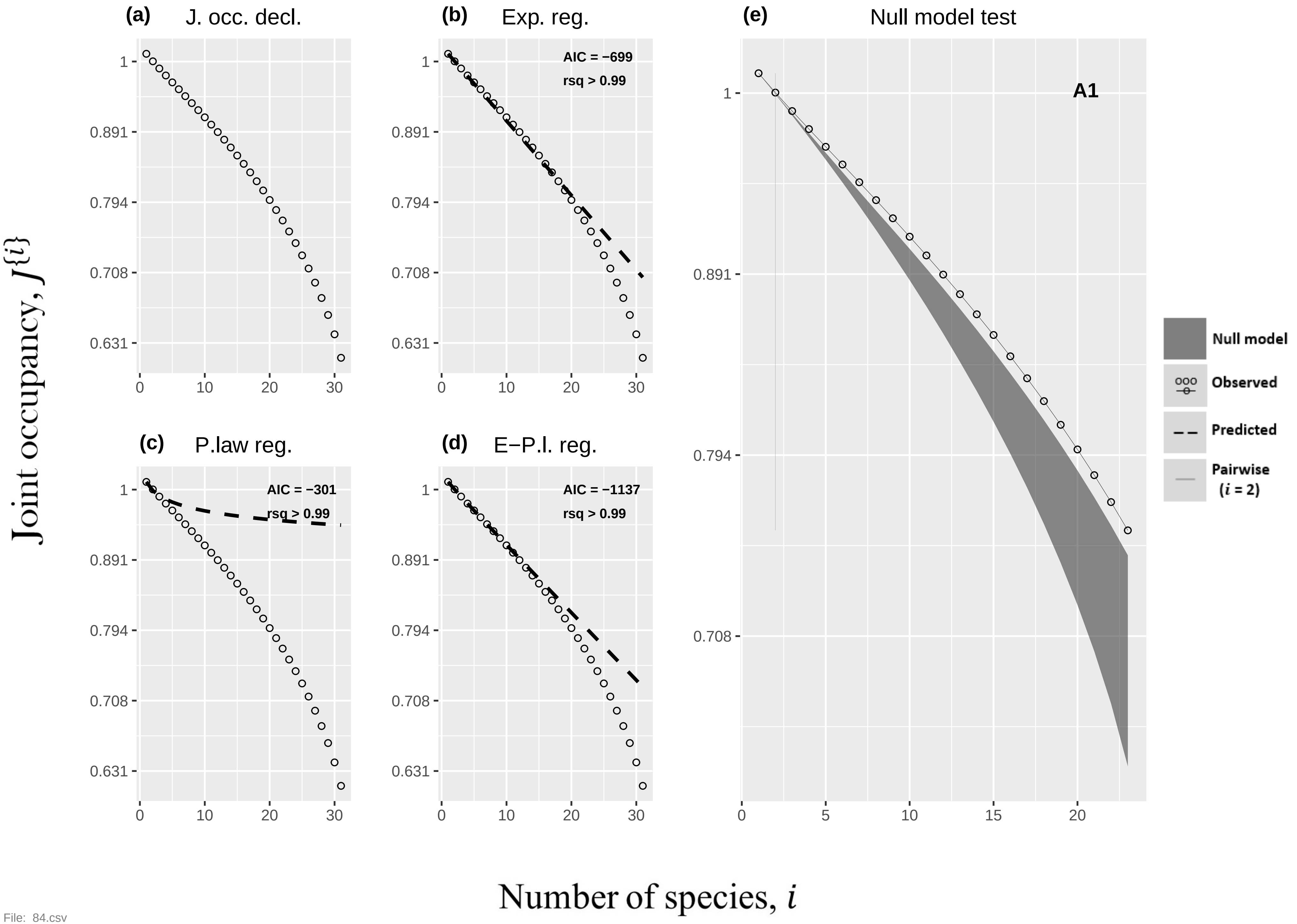

(a) J. occ. decl.
(b) Exp. reg.
AIC = −699
rsq > 0.99
(e) Null model test
A1
(c) P.law reg.
AIC = −301
rsq > 0.99
(d) E−P.l. reg.
AIC = −1137
rsq > 0.99
Null model
Observed
Predicted
Pairwise (i = 2)
Joint occupancy, J{i}
Number of species, i
1
0.891
0.794
0.708
0.631
0
10
20
30
5
15


File: 84.csv

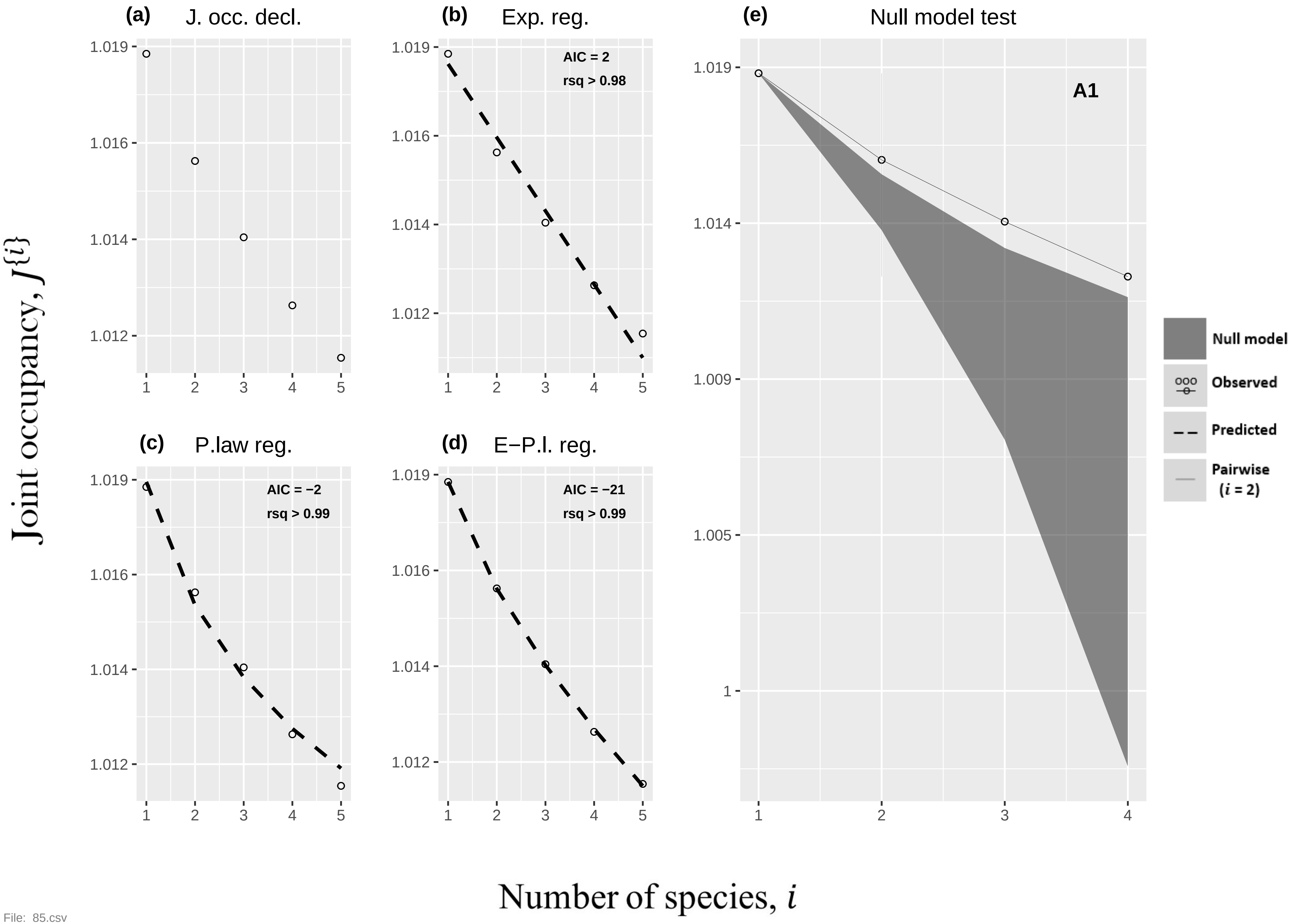
(a) J. occ. decl.
(b) Exp. reg.
AIC = 2
rsq > 0.98
(c) P.law reg.
AIC = −2
rsq > 0.99
(d) E−P.l. reg.
AIC = −21
rsq > 0.99
(e) Null model test
A1
Null model
Observed
Predicted
Pairwise (i = 2)
Joint occupancy, J{i}
Number of species, i
1.019
1.016
1.014
1.012
1.009
1.005
1
1
2
3
4
5

File: 85.csv

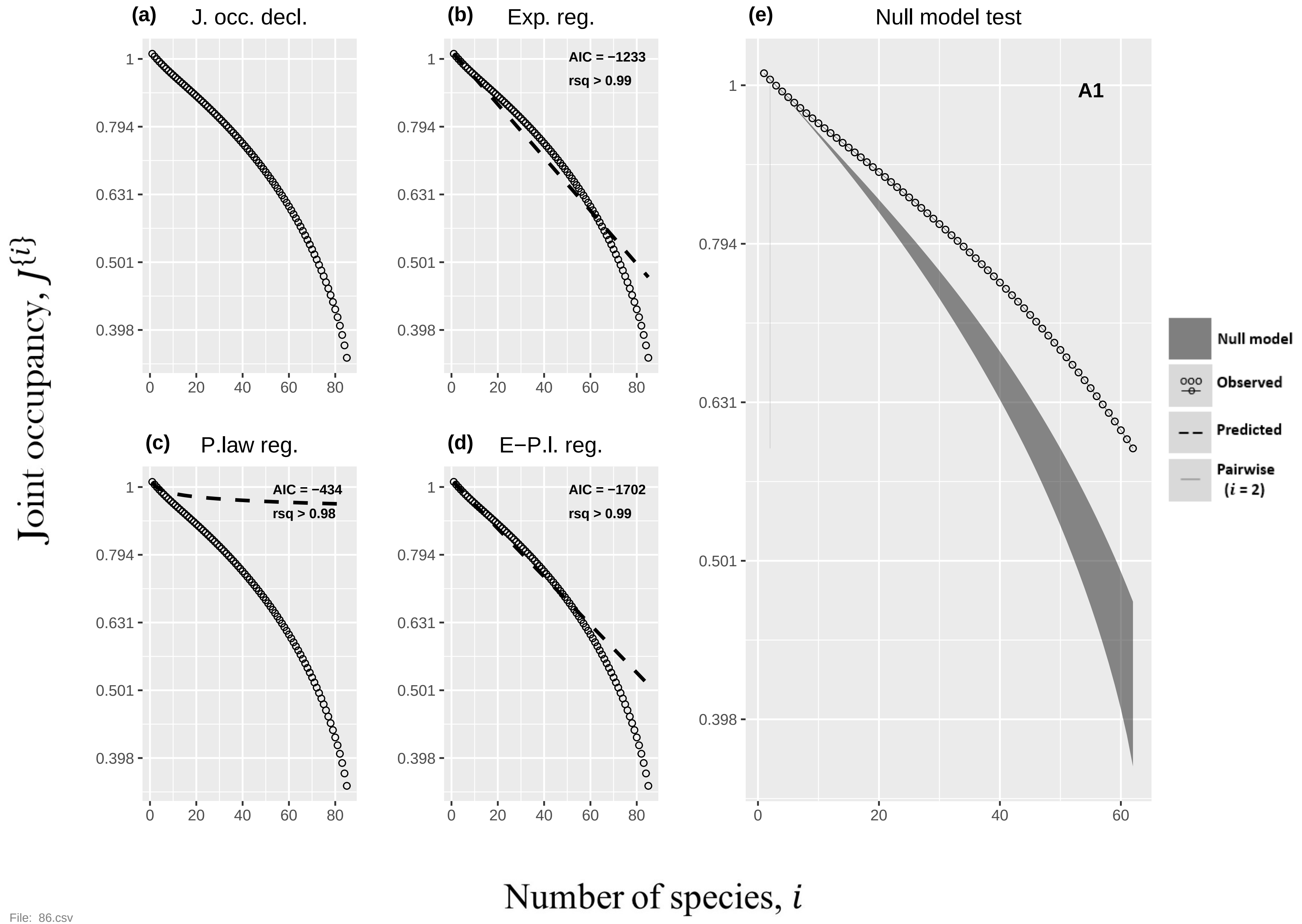

(a)
J. occ. decl.
(b)
Exp. reg.
AIC = −1233
rsq > 0.99
(c)
P.law reg.
AIC = −434
rsq > 0.98
(d)
E−P.l. reg.
AIC = −1702
rsq > 0.99
(e)
Null model test
A1
Null model
Observed
Predicted
Pairwise
(i = 2)
1
0.794
0.631
0.501
0.398
0
20
40
60
80
Joint occupancy, J{i}
Number of species, i


File: 86.csv

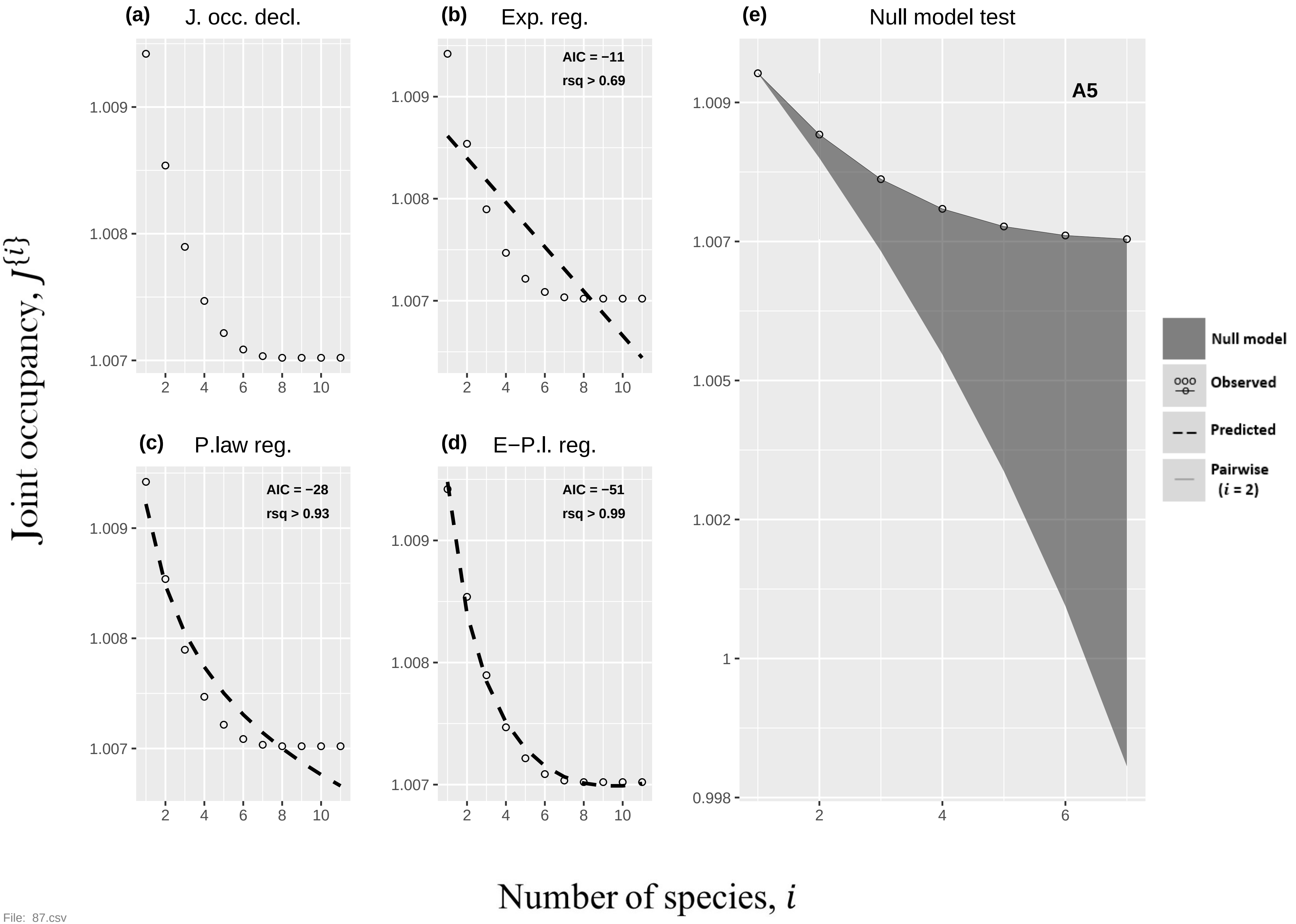
(a) J. occ. decl.
(b) Exp. reg.
AIC = −11
rsq > 0.69
(c) P.law reg.
AIC = −28
rsq > 0.93
(d) E−P.l. reg.
AIC = −51
rsq > 0.99
(e) Null model test
A5
Null model
Observed
Predicted
Pairwise
(i = 2)
Joint occupancy, J{i}
Number of species, i
1.009
1.008
1.007
1.005
1.002
1
0.998
2
4
6
8
10
File: 87.csv

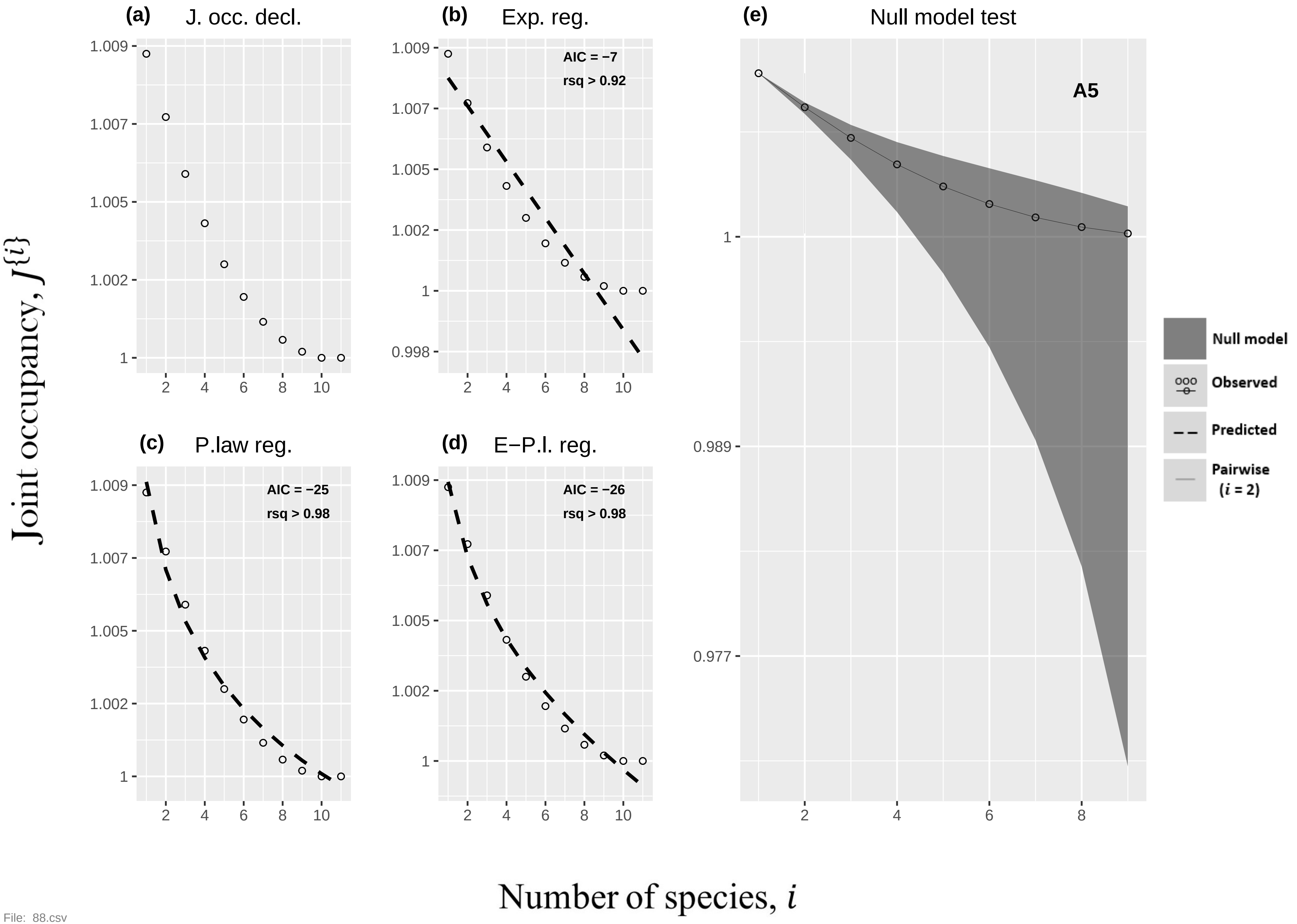
(a) J. occ. decl.
(b) Exp. reg.
AIC = −7
rsq > 0.92
(c) P.law reg.
AIC = −25
rsq > 0.98
(d) E−P.l. reg.
AIC = −26
rsq > 0.98
(e) Null model test
A5
Null model
Observed
Predicted
Pairwise (i = 2)
Joint occupancy, J{i}
Number of species, i

File: 88.csv

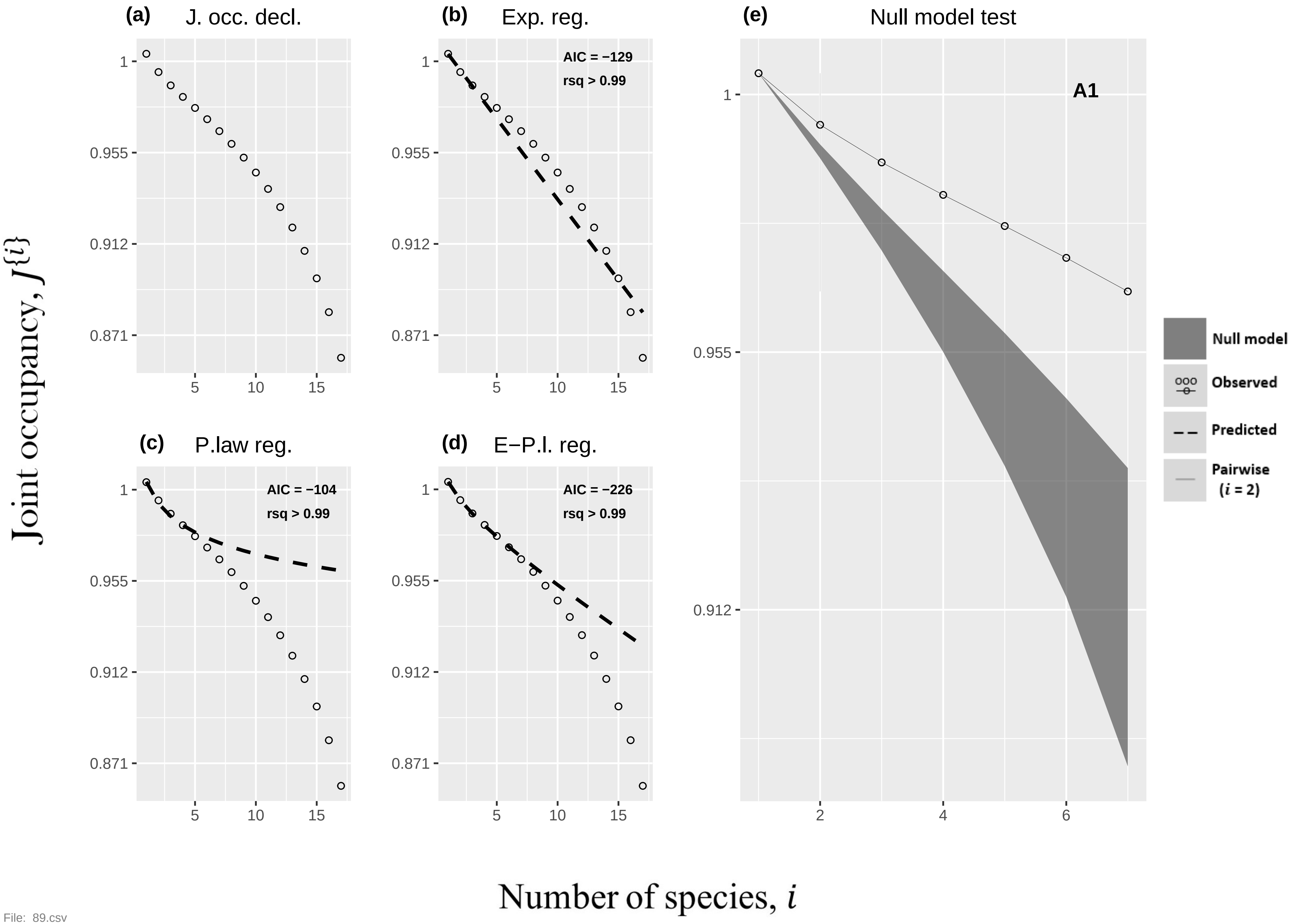

(a)
J. occ. decl.
(b)
Exp. reg.
AIC = −129
rsq > 0.99
(c)
P.law reg.
AIC = −104
rsq > 0.99
(d)
E−P.l. reg.
AIC = −226
rsq > 0.99
(e)
Null model test
A1
Null model
Observed
Predicted
Pairwise (i = 2)
1
0.955
0.912
0.871
5
10
15
2
4
6
Joint occupancy, J{i}
Number of species, i

File: 89.csv

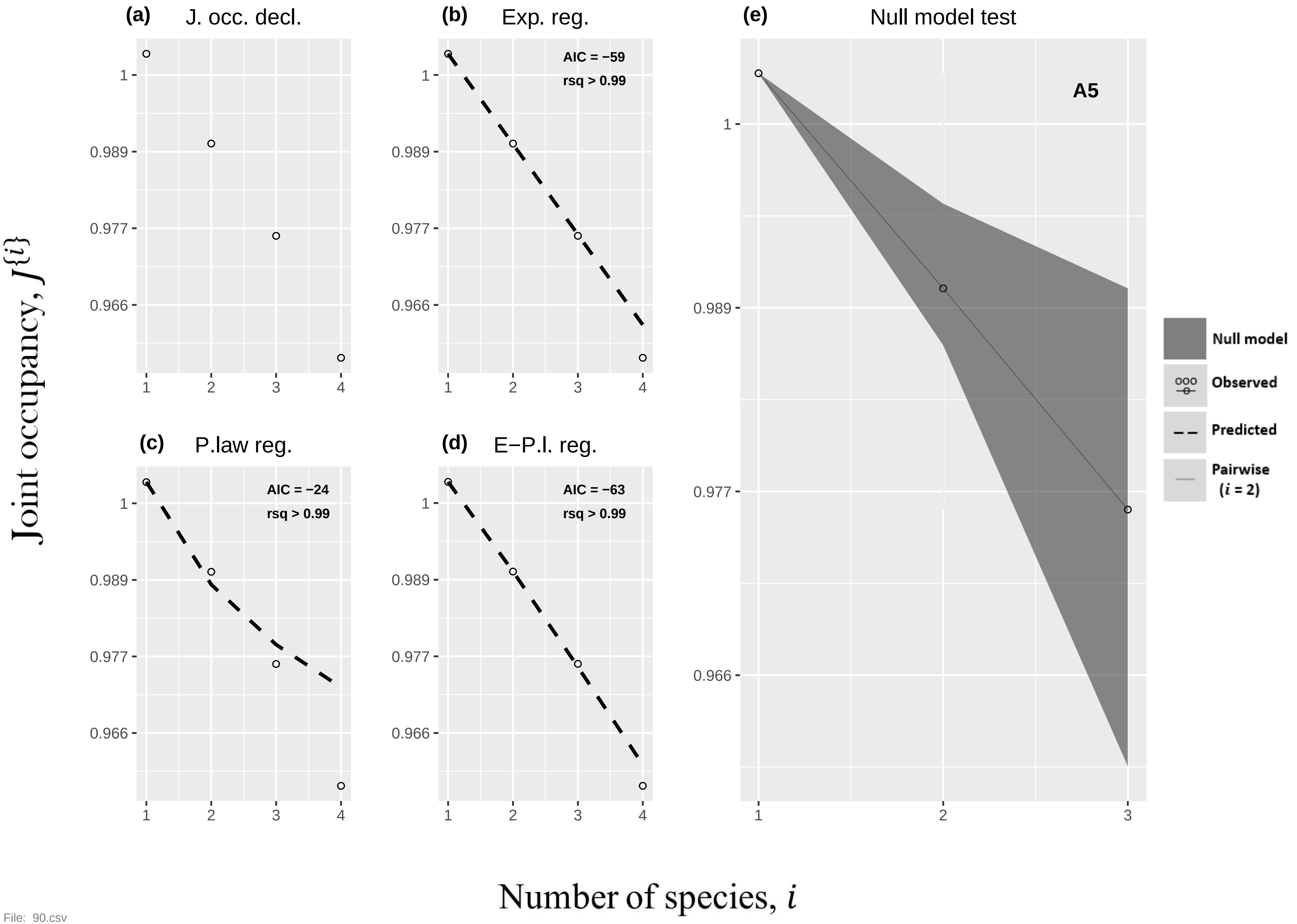

(a)
J. occ. decl.
(b)
Exp. reg.
AIC = −59
rsq > 0.99
(c)
P.law reg.
AIC = −24
rsq > 0.99
(d)
E−P.l. reg.
AIC = −63
rsq > 0.99
(e)
Null model test
A5
Null model
Observed
Predicted
Pairwise
(i = 2)
1
0.989
0.977
0.966
1
2
3
4
Joint occupancy, J{i}
Number of species, i


File: 90.csv

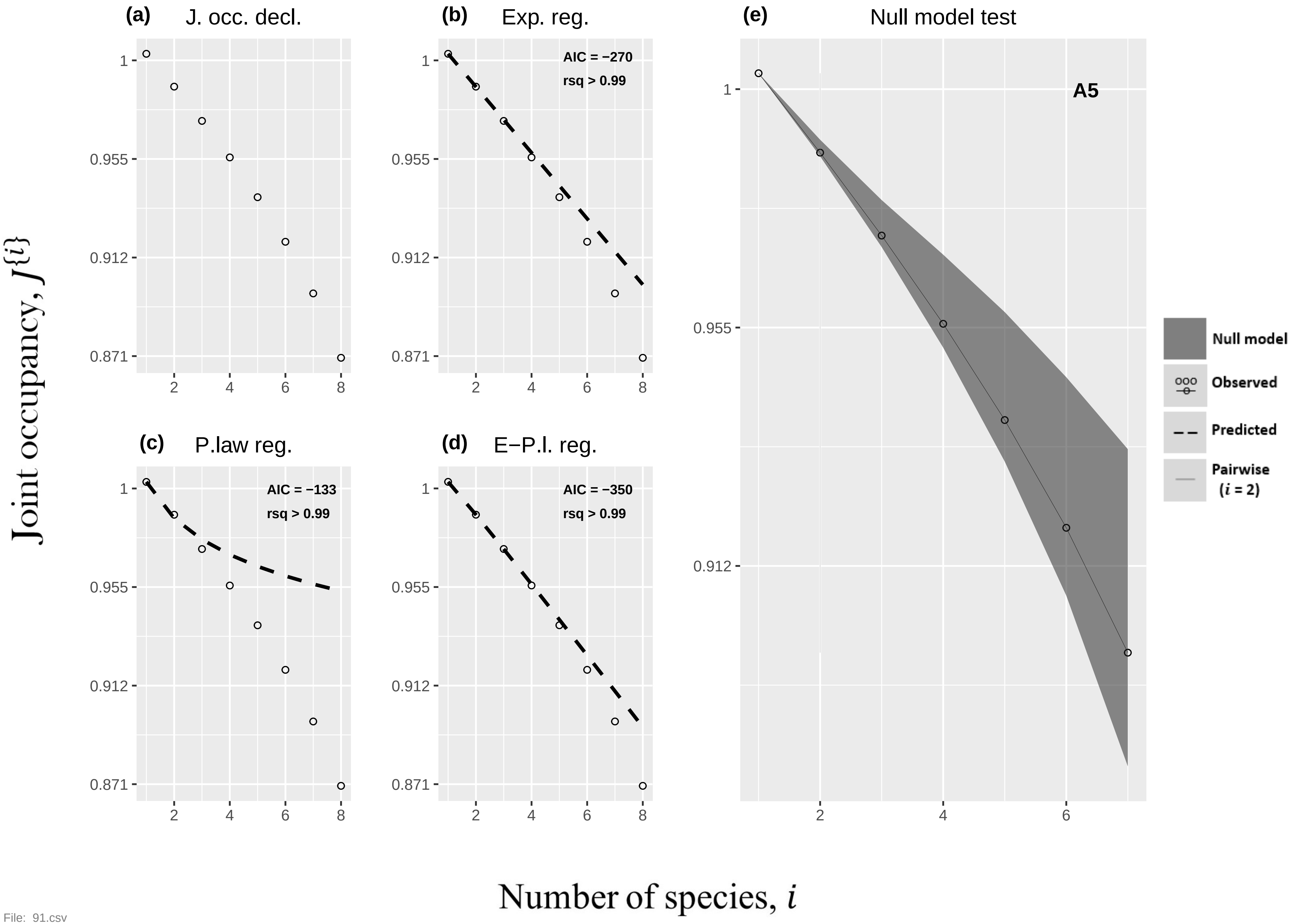

(a) J. occ. decl.
(b) Exp. reg.
AIC = −270
rsq > 0.99
(c) P.law reg.
AIC = −133
rsq > 0.99
(d) E−P.l. reg.
AIC = −350
rsq > 0.99
(e) Null model test
A5
Null model
Observed
Predicted
Pairwise (i = 2)
1
0.955
0.912
0.871
2
4
6
8
Joint occupancy, J{i}
Number of species, i


File: 91.csv

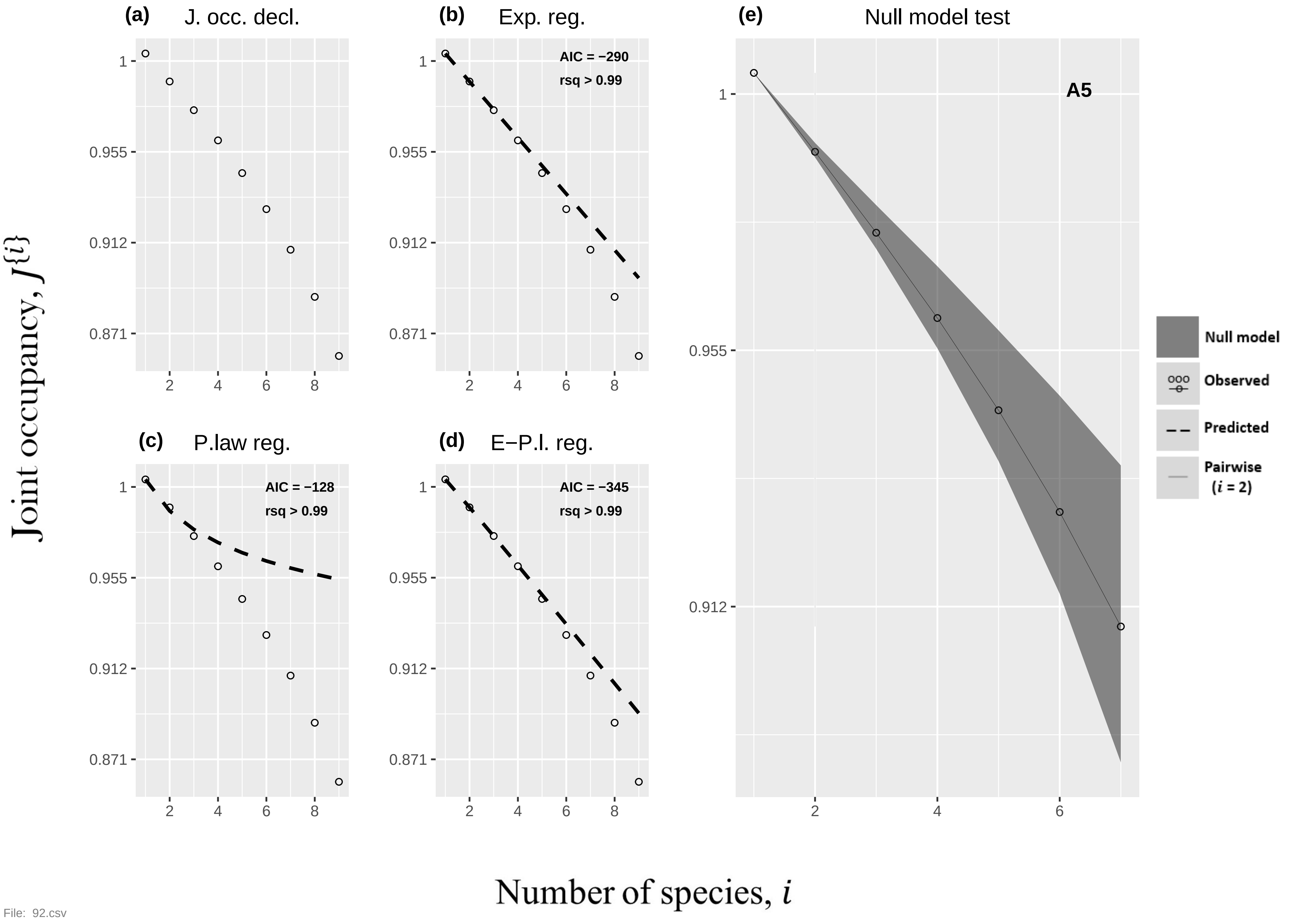
(a) J. occ. decl.
(b) Exp. reg.
AIC = −290
rsq > 0.99
(c) P.law reg.
AIC = −128
rsq > 0.99
(d) E−P.l. reg.
AIC = −345
rsq > 0.99
(e) Null model test
A5
Null model
Observed
Predicted
Pairwise (i = 2)
Joint occupancy, J{i}
Number of species, i
1
0.955
0.912
0.871
2
4
6
8
File: 92.csv

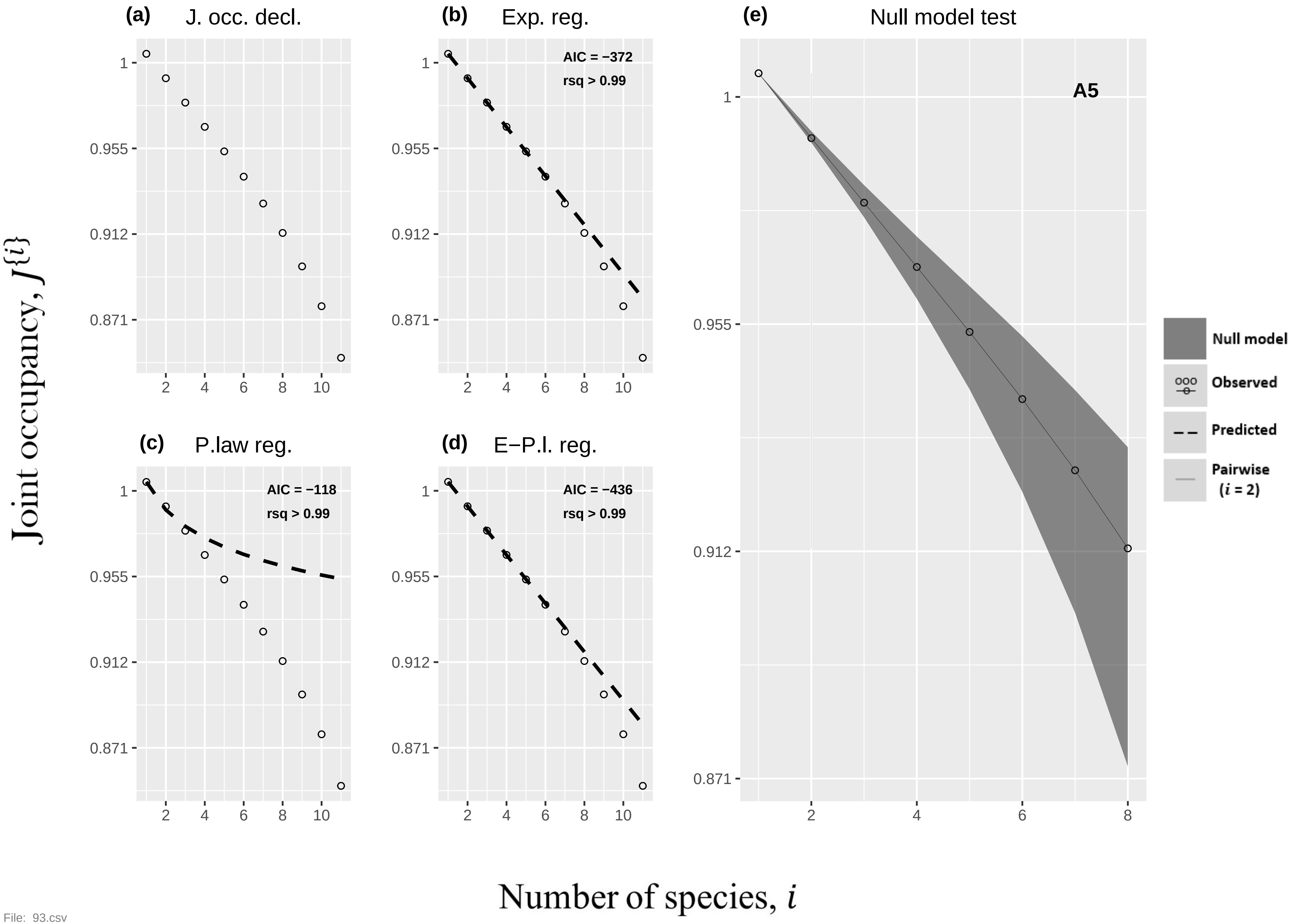

(a)
J. occ. decl.
(b)
Exp. reg.
AIC = −372
rsq > 0.99
(c)
P.law reg.
AIC = −118
rsq > 0.99
(d)
E−P.l. reg.
AIC = −436
rsq > 0.99
(e)
Null model test
A5
Null model
Observed
Predicted
Pairwise
(i = 2)
1
0.955
0.912
0.871
2
4
6
8
10
Joint occupancy, J{i}
Number of species, i


File: 93.csv

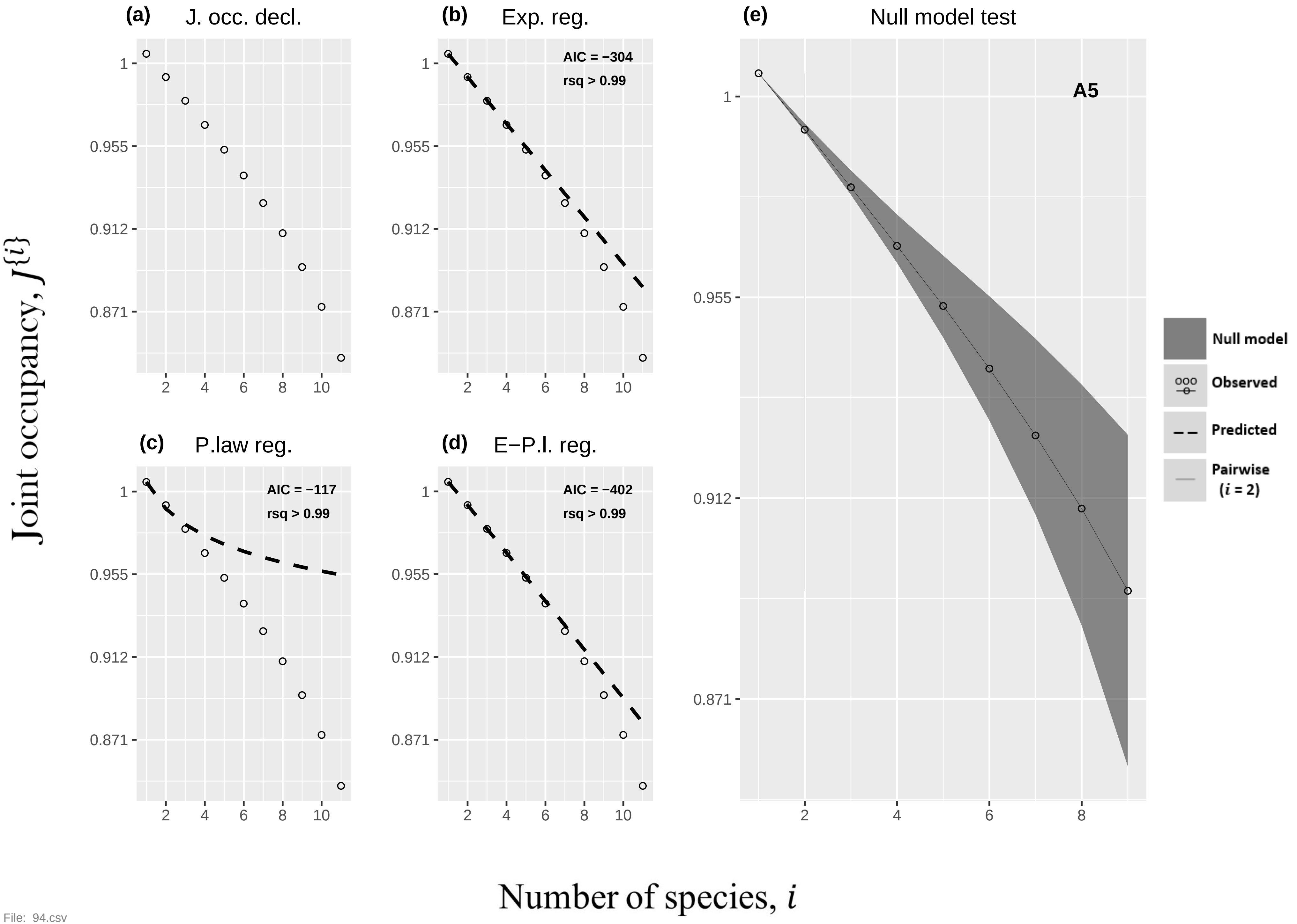
(a) J. occ. decl.
(b) Exp. reg.
AIC = −304
rsq > 0.99
(c) P.law reg.
AIC = −117
rsq > 0.99
(d) E−P.l. reg.
AIC = −402
rsq > 0.99
(e) Null model test
A5
Null model
Observed
Predicted
Pairwise (i = 2)
Joint occupancy, J{i}
Number of species, i
1
0.955
0.912
0.871
2
4
6
8
10

File: 94.csv

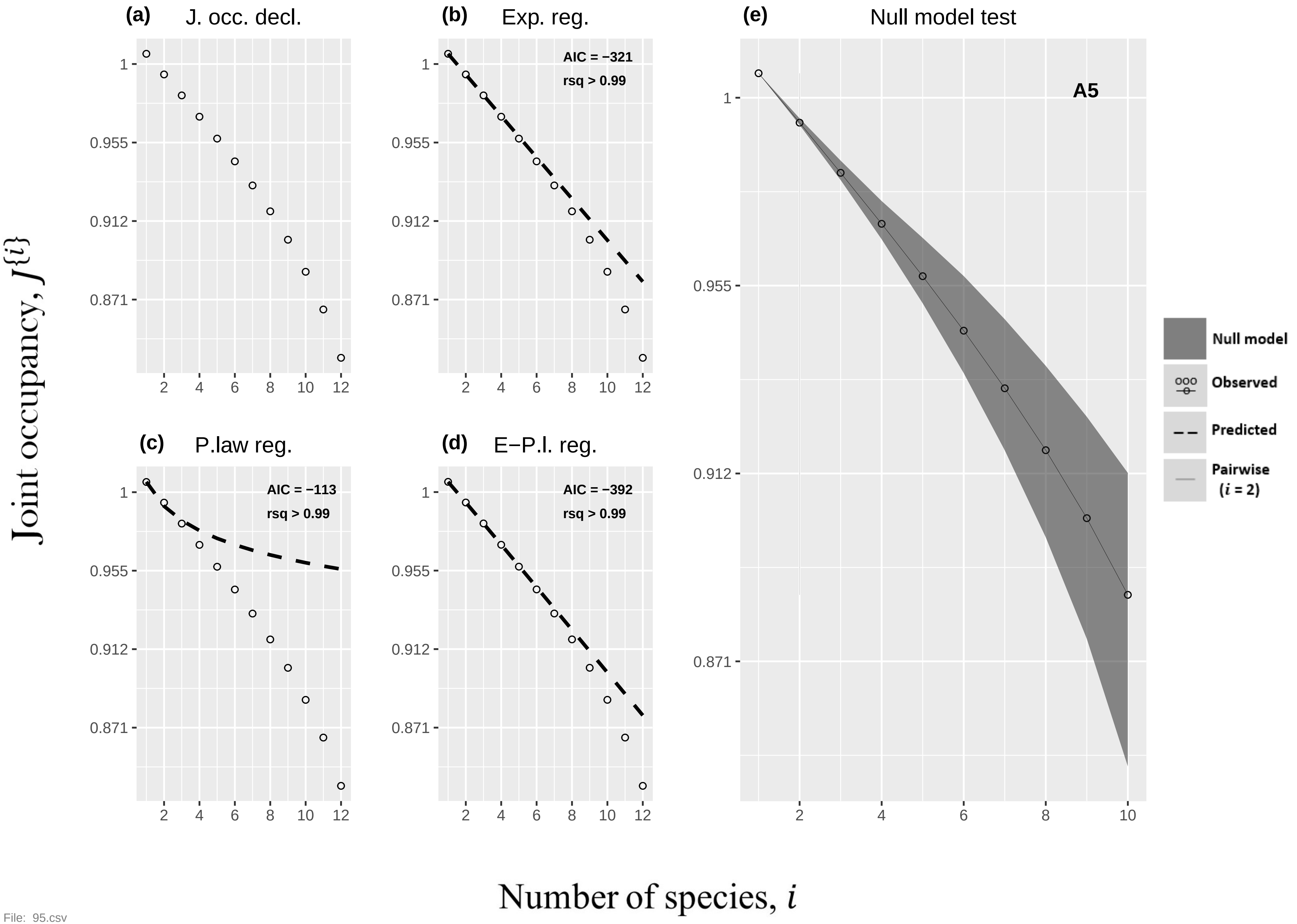

(a) J. occ. decl.
(b) Exp. reg.
AIC = −321
rsq > 0.99
(c) P.law reg.
AIC = −113
rsq > 0.99
(d) E−P.l. reg.
AIC = −392
rsq > 0.99
(e) Null model test
A5
Null model
Observed
Predicted
Pairwise (i = 2)
1
0.955
0.912
0.871
2 4 6 8 10 12
2 4 6 8 10
Joint occupancy, J{i}
Number of species, i


File: 95.csv

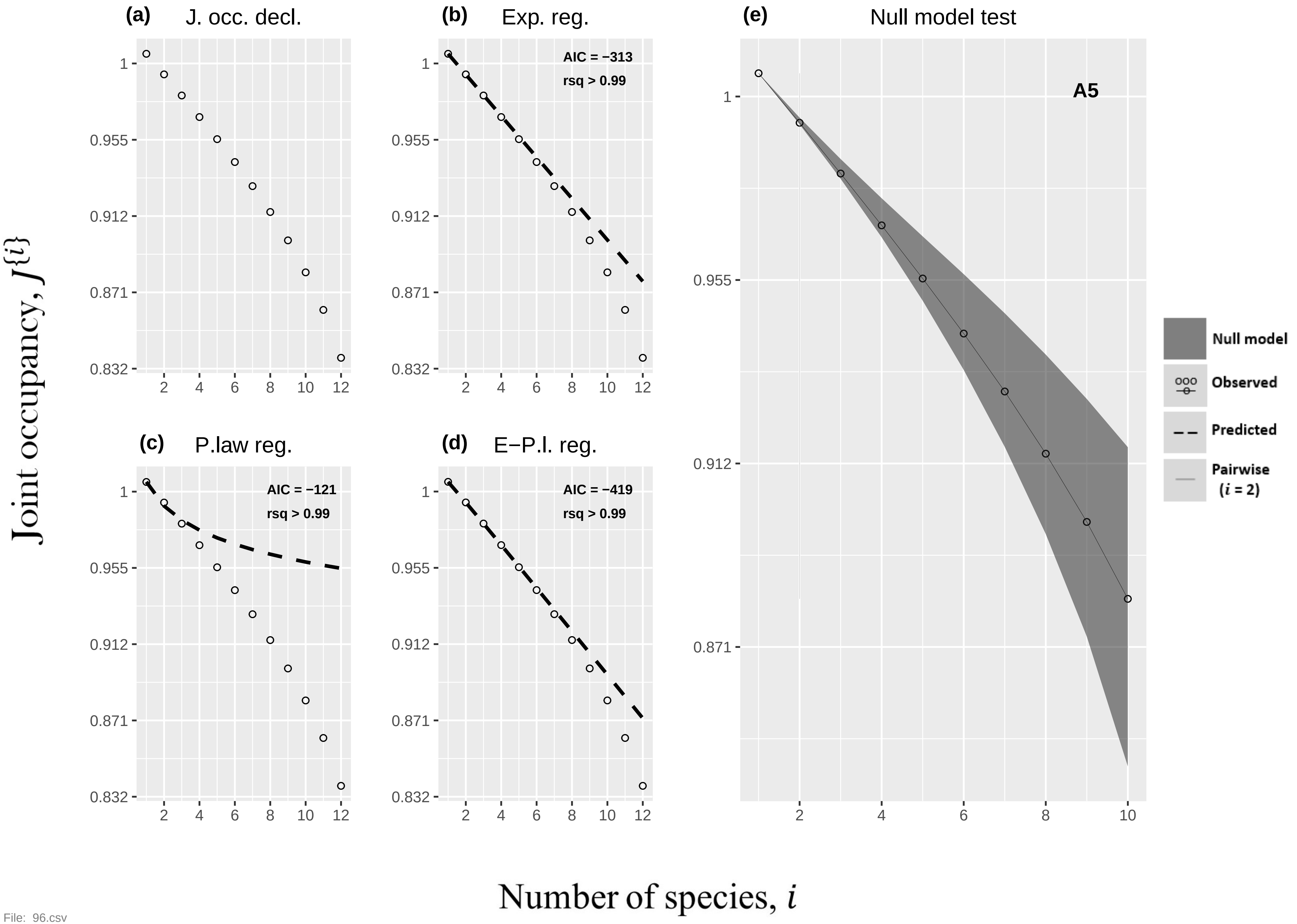
(a)
J. occ. decl.
(b)
Exp. reg.
AIC = −313
rsq > 0.99
(e)
Null model test
A5
(c)
P.law reg.
AIC = −121
rsq > 0.99
(d)
E−P.l. reg.
AIC = −419
rsq > 0.99
Null model
Observed
Predicted
Pairwise (i = 2)
Joint occupancy, J{i}
Number of species, i
1
0.955
0.912
0.871
0.832
2
4
6
8
10
12
File: 96.csv

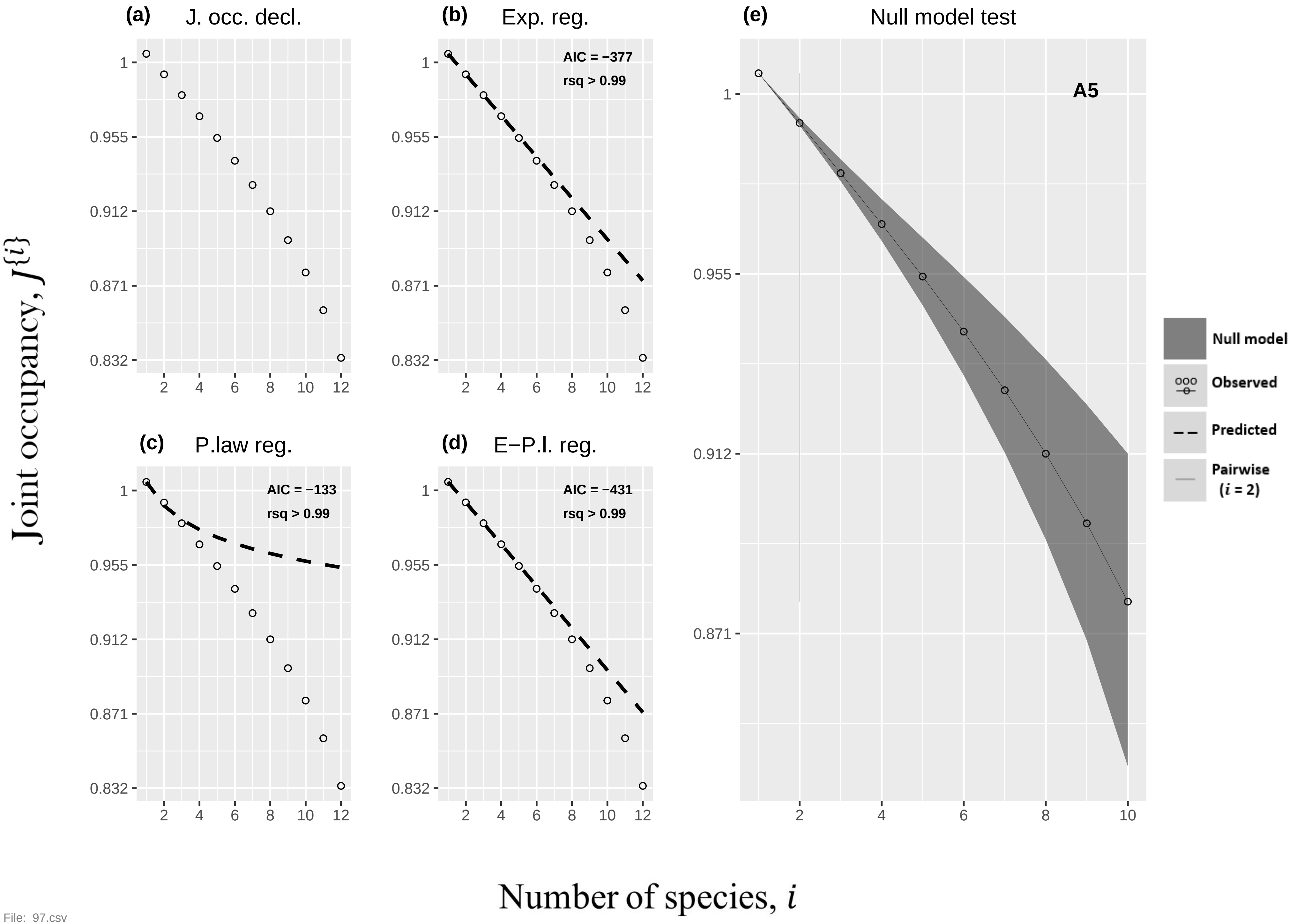
(a) J. occ. decl.
(b) Exp. reg.
AIC = −377
rsq > 0.99
(c) P.law reg.
AIC = −133
rsq > 0.99
(d) E−P.l. reg.
AIC = −431
rsq > 0.99
(e) Null model test
A5
Null model
Observed
Predicted
Pairwise (i = 2)
Joint occupancy, J{i}
Number of species, i

File: 97.csv

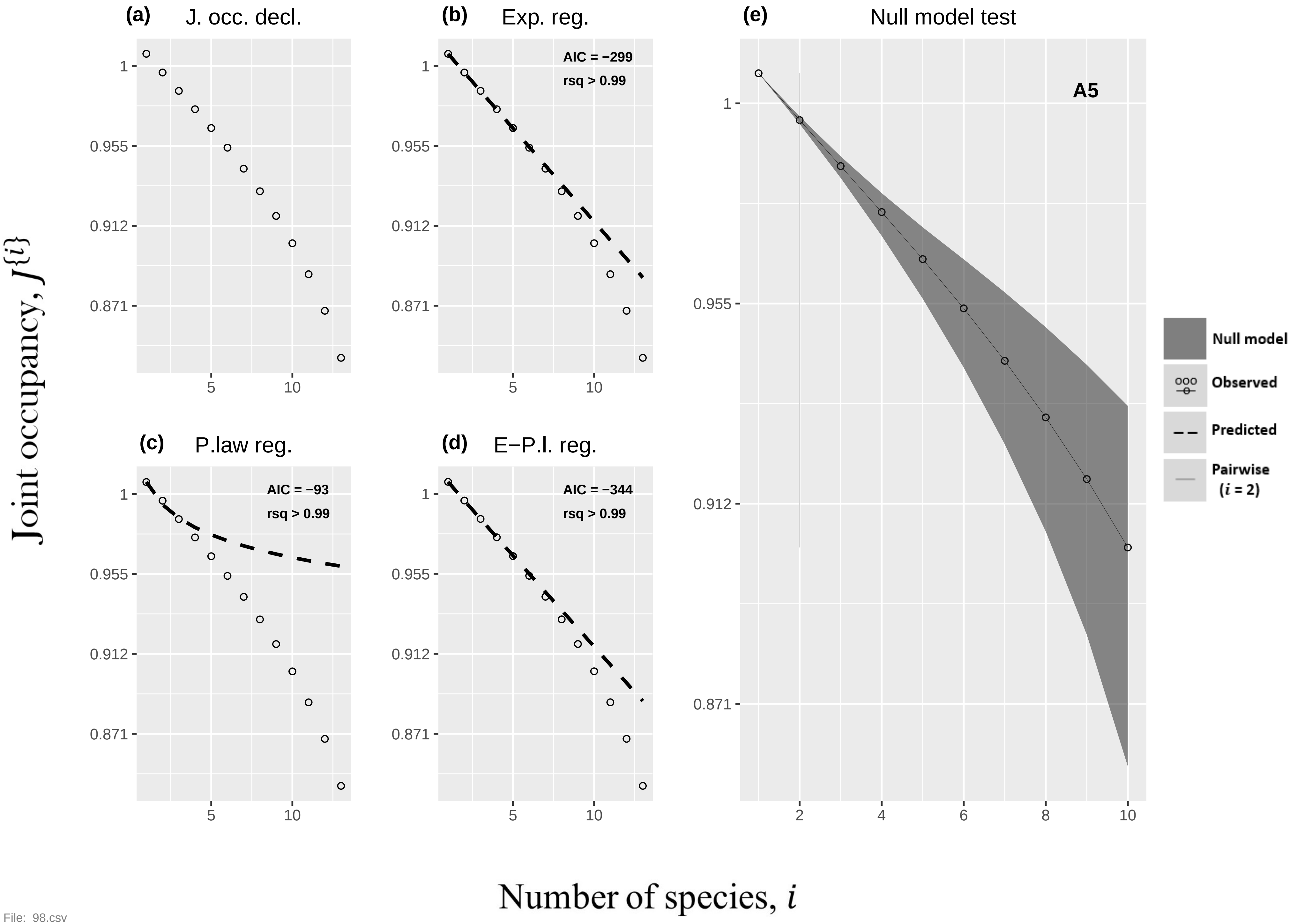
(a)
J. occ. decl.
(b)
Exp. reg.
AIC = −299
rsq > 0.99
(c)
P.law reg.
AIC = −93
rsq > 0.99
(d)
E−P.l. reg.
AIC = −344
rsq > 0.99
(e)
Null model test
A5
Null model
Observed
Predicted
Pairwise
(i = 2)
1
0.955
0.912
0.871
5
10
2
4
6
8
Joint occupancy, J{i}
Number of species, i

File: 98.csv

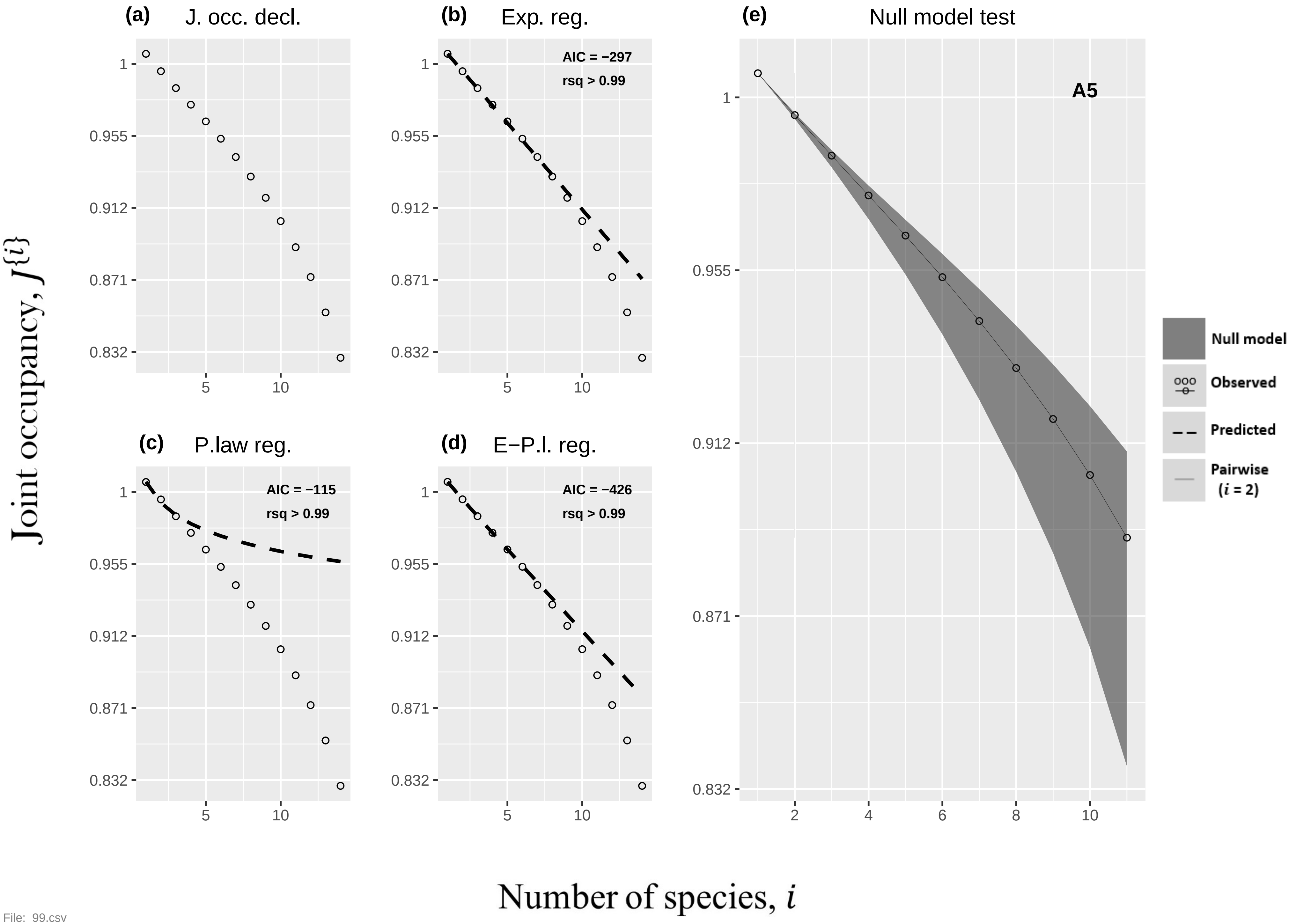
(a) J. occ. decl.
(b) Exp. reg.
AIC = −297
rsq > 0.99
(c) P.law reg.
AIC = −115
rsq > 0.99
(d) E−P.l. reg.
AIC = −426
rsq > 0.99
(e) Null model test
A5
Null model
Observed
Predicted
Pairwise (i = 2)
Joint occupancy, J{i}
Number of species, i
1
0.955
0.912
0.871
0.832
5
10
2
4
6
8

File: 99.csv

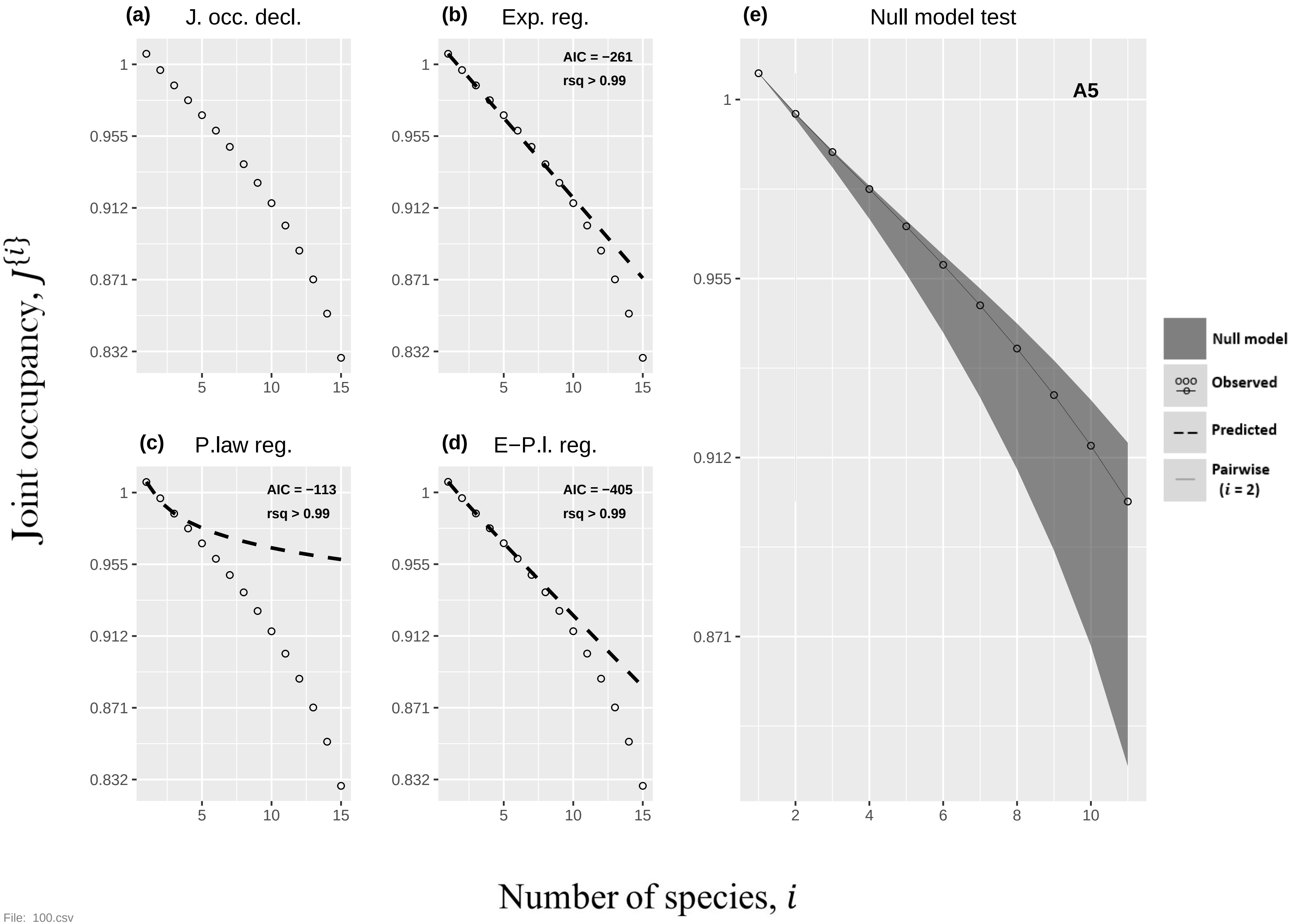
(a)
J. occ. decl.
(b)
Exp. reg.
AIC = −261
rsq > 0.99
(c)
P.law reg.
AIC = −113
rsq > 0.99
(d)
E−P.l. reg.
AIC = −405
rsq > 0.99
(e)
Null model test
A5
Null model
Observed
Predicted
Pairwise
(i = 2)
Joint occupancy, J{i}
Number of species, i
1
0.955
0.912
0.871
0.832
5
10
15
2
4
6
8
10

File: 100.csv

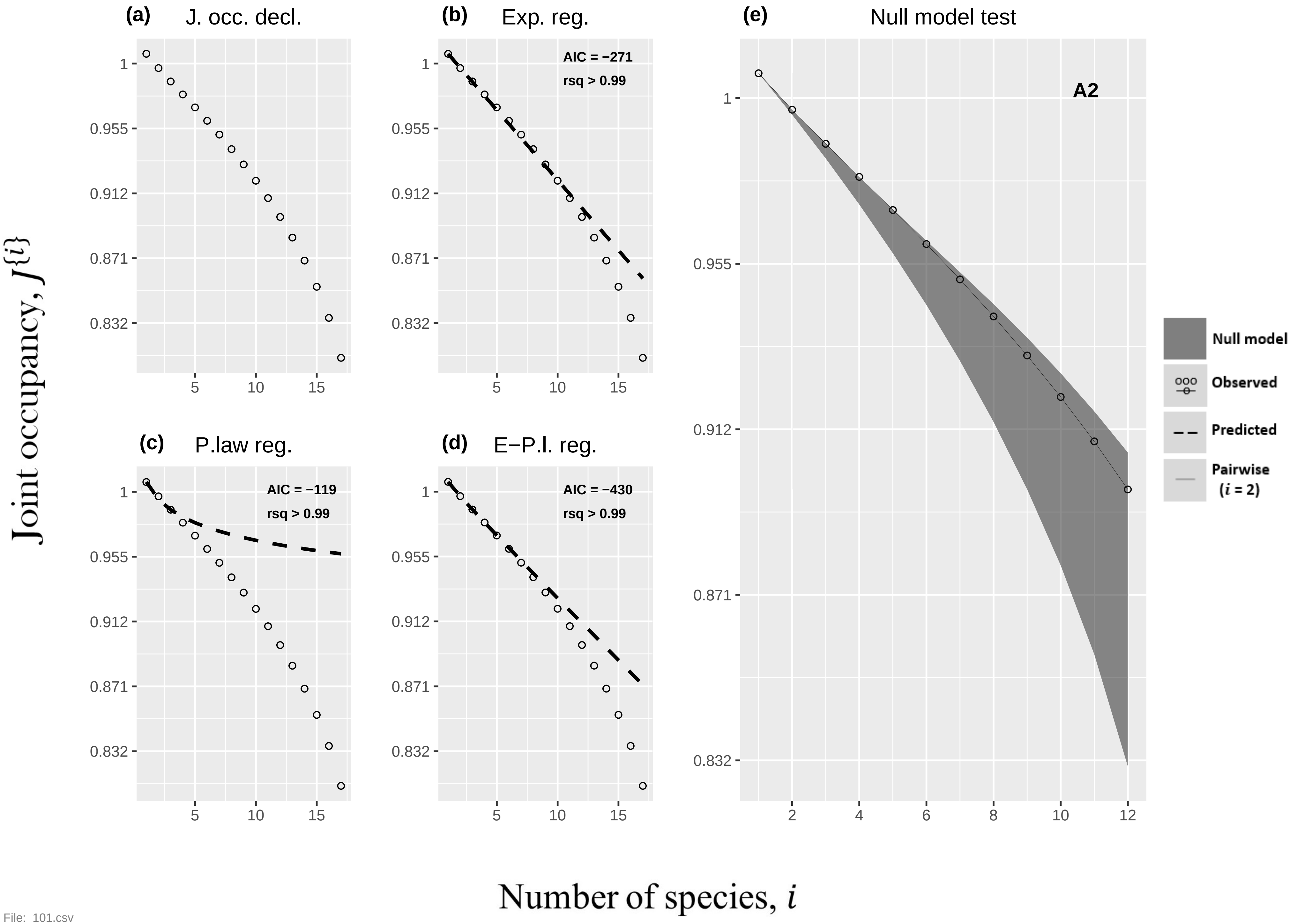

(a)
J. occ. decl.
(b)
Exp. reg.
AIC = −271
rsq > 0.99
(c)
P.law reg.
AIC = −119
rsq > 0.99
(d)
E−P.l. reg.
AIC = −430
rsq > 0.99
(e)
Null model test
A2
Null model
Observed
Predicted
Pairwise
(i = 2)
Joint occupancy, J{i}
Number of species, i
1
0.955
0.912
0.871
0.832
5
10
15
2
4
6
8
10
12


File: 101.csv

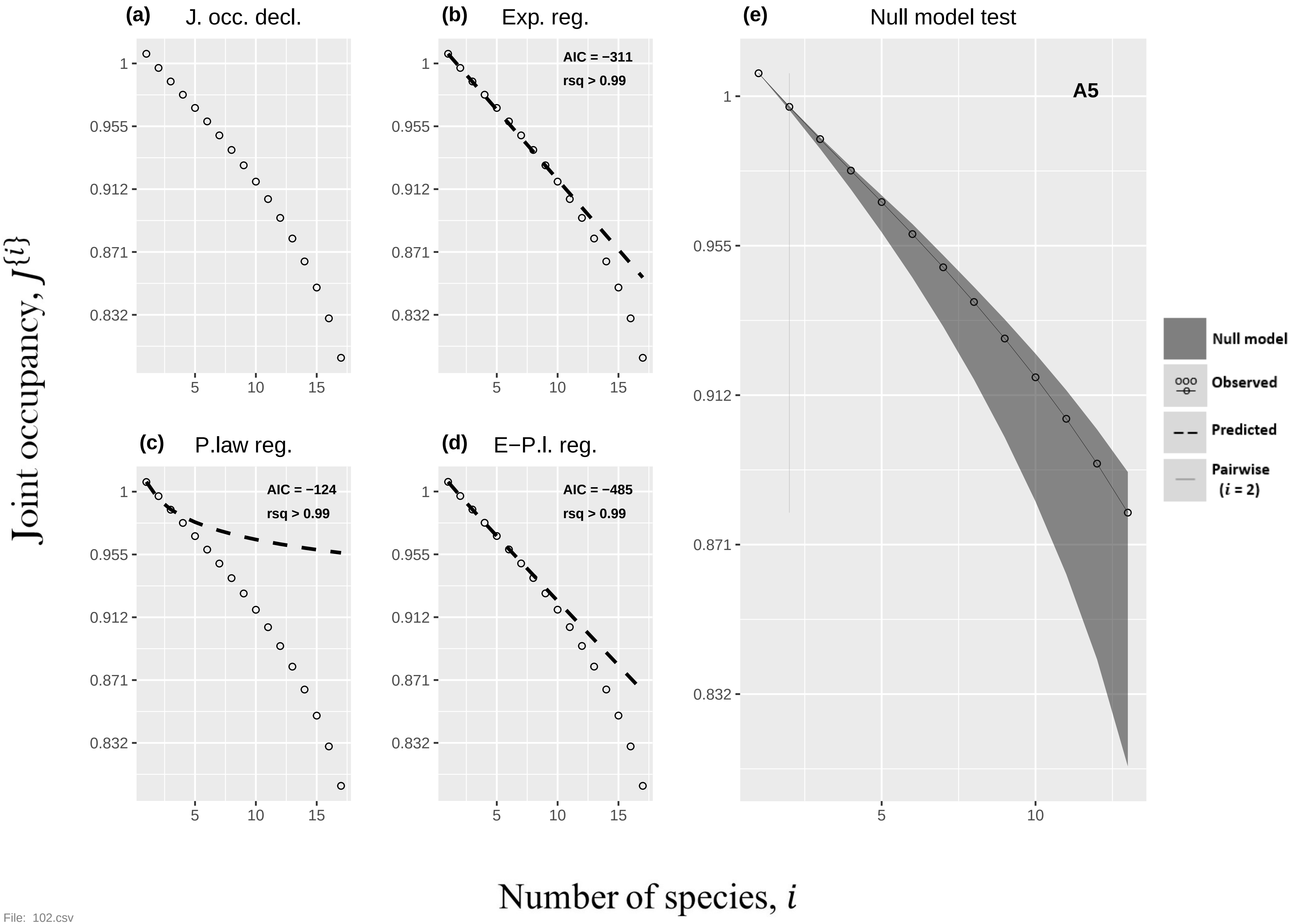

(a) J. occ. decl.
(b) Exp. reg.
AIC = −311
rsq > 0.99
(c) P.law reg.
AIC = −124
rsq > 0.99
(d) E−P.l. reg.
AIC = −485
rsq > 0.99
(e) Null model test
A5
Null model
Observed
Predicted
Pairwise (i = 2)
1
0.955
0.912
0.871
0.832
5
10
15
Joint occupancy, J{i}
Number of species, i


File: 102.csv

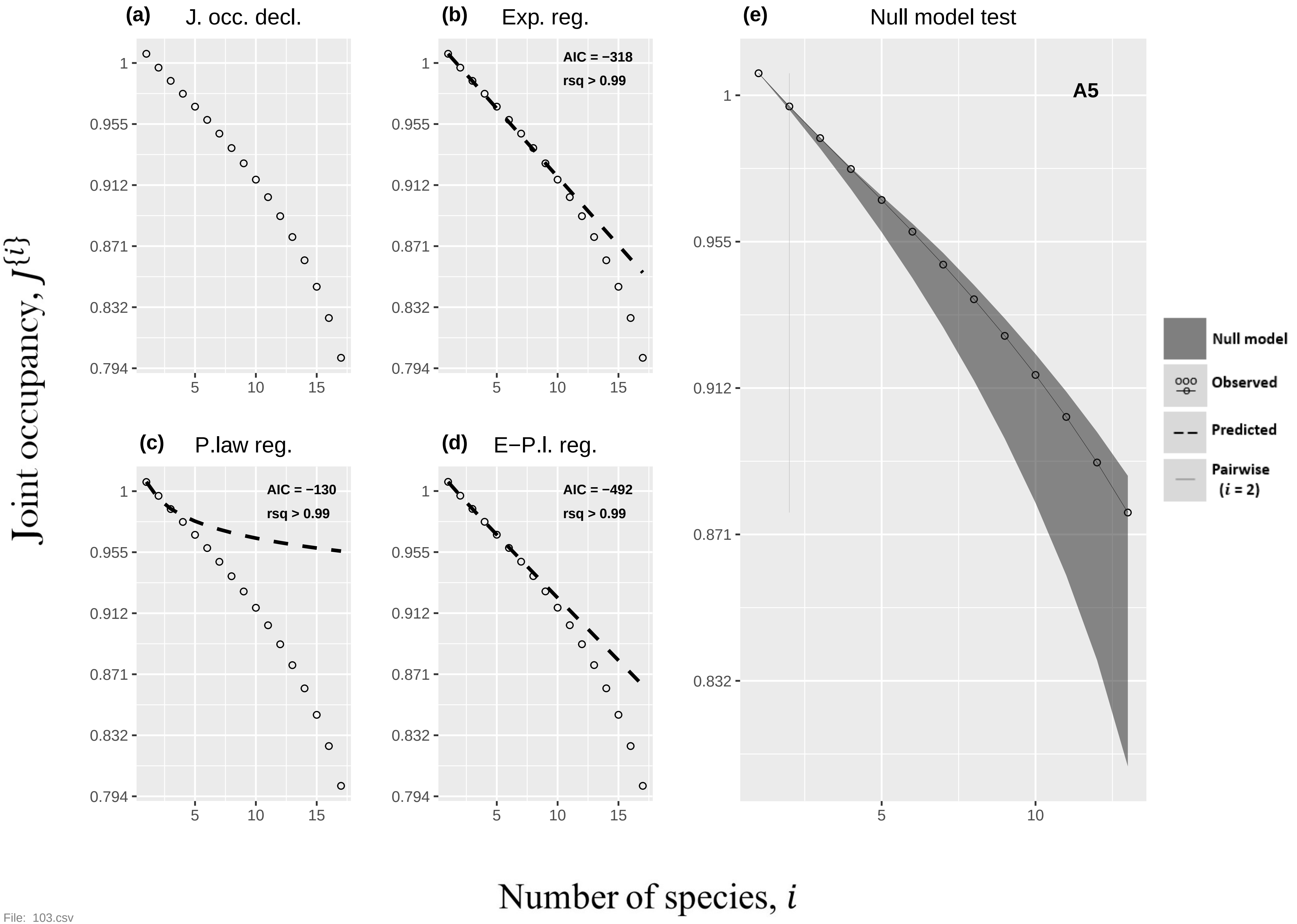

(a) J. occ. decl.
(b) Exp. reg.
AIC = −318
rsq > 0.99
(c) P.law reg.
AIC = −130
rsq > 0.99
(d) E−P.l. reg.
AIC = −492
rsq > 0.99
(e) Null model test
A5
Null model
Observed
Predicted
Pairwise ($i$ = 2)
1
0.955
0.912
0.871
0.832
0.794
5
10
15
Joint occupancy, $J\{i\}$
Number of species, $i$


File: 103.csv

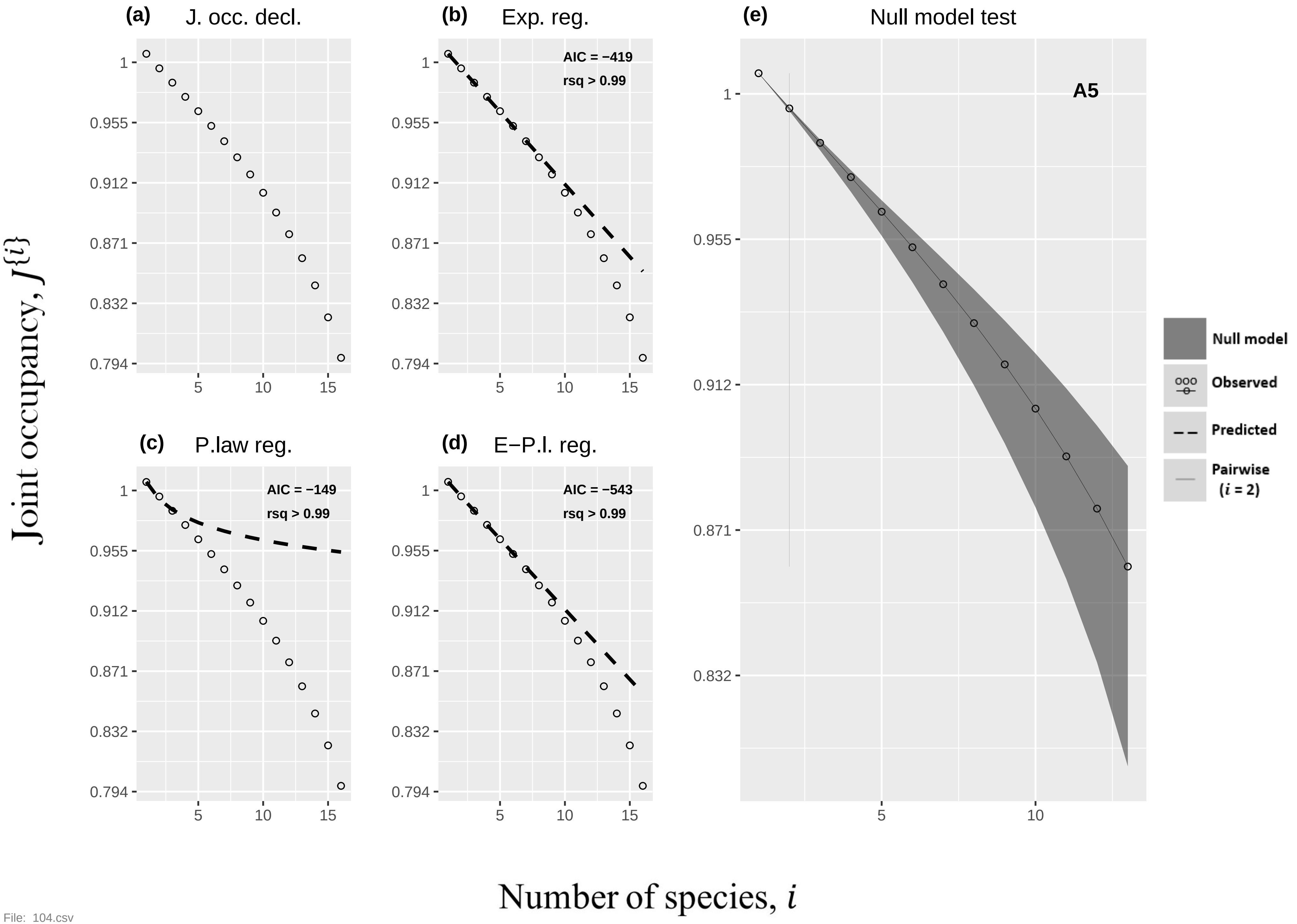

(a)
J. occ. decl.
(b)
Exp. reg.
AIC = −419
rsq > 0.99
(e)
Null model test
A5
(c)
P.law reg.
AIC = −149
rsq > 0.99
(d)
E−P.l. reg.
AIC = −543
rsq > 0.99
1
0.955
0.912
0.871
0.832
0.794
5
10
15
Null model
Observed
Predicted
Pairwise
(i = 2)
Joint occupancy, J{i}
Number of species, i


File: 104.csv

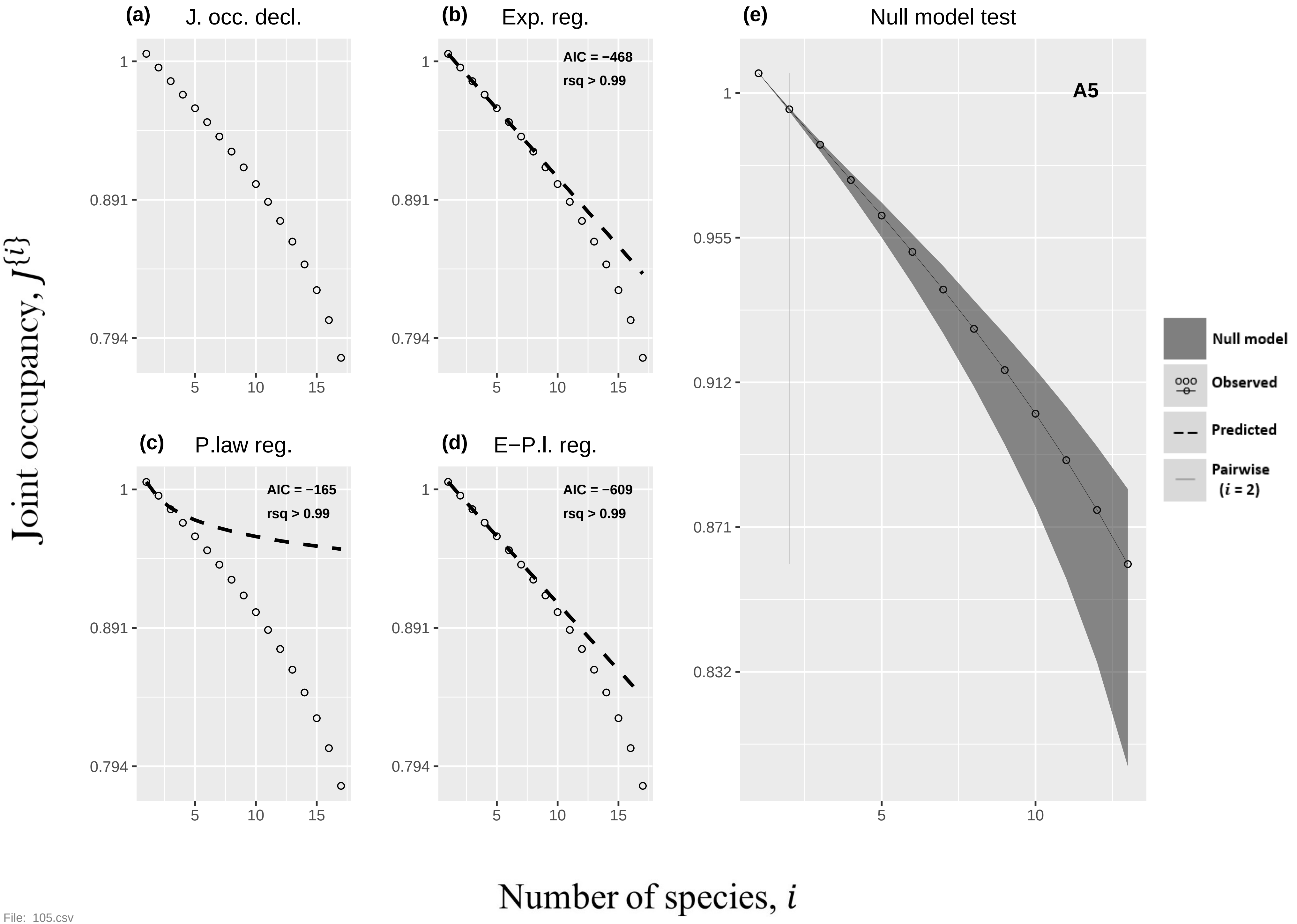

(a) J. occ. decl.
(b) Exp. reg.
AIC = −468
rsq > 0.99
(c) P.law reg.
AIC = −165
rsq > 0.99
(d) E−P.l. reg.
AIC = −609
rsq > 0.99
(e) Null model test
A5
Null model
Observed
Predicted
Pairwise (i = 2)
1
0.891
0.794
0.955
0.912
0.871
0.832
5
10
15
Joint occupancy, J{i}
Number of species, i


File: 105.csv

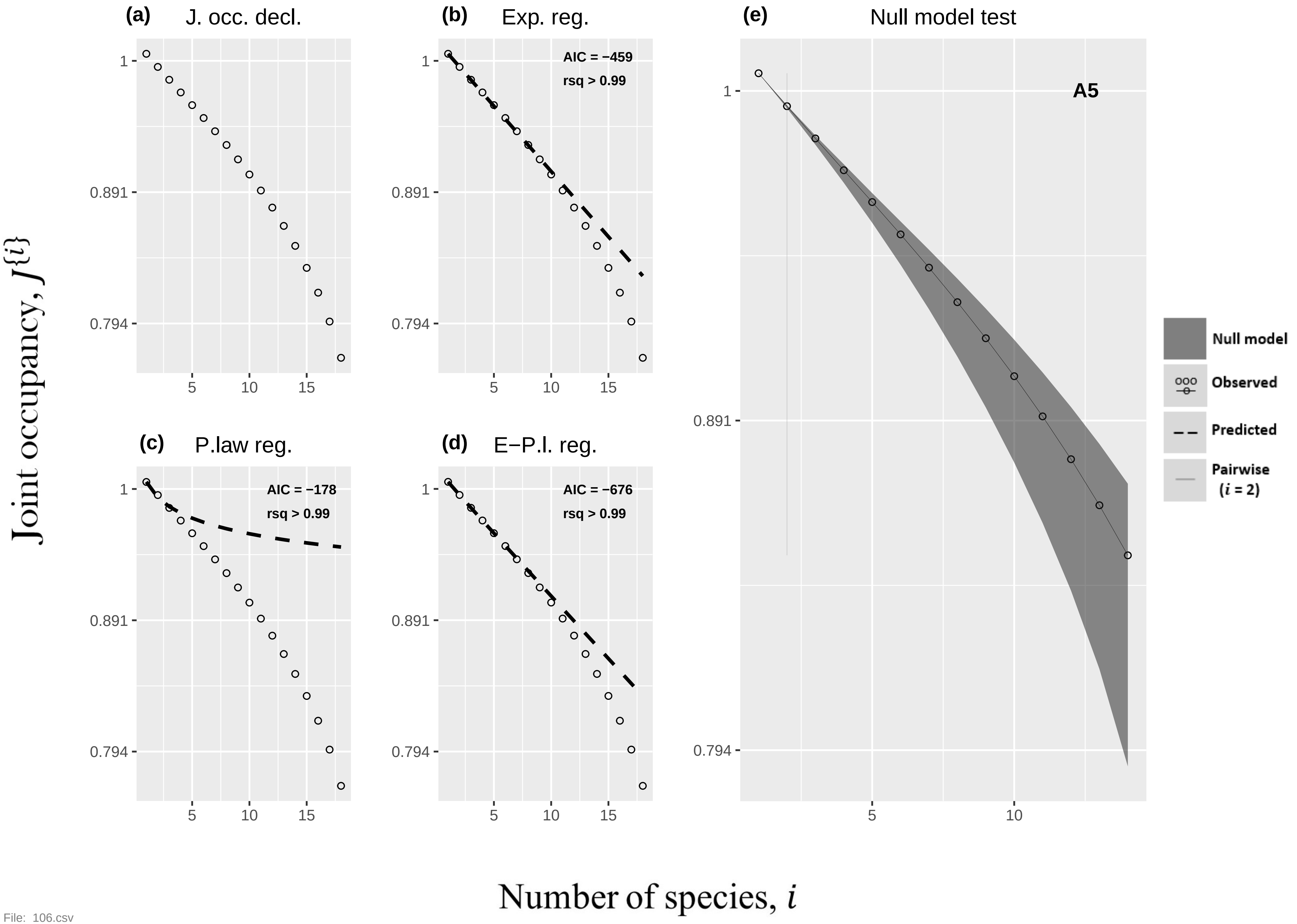

(a) J. occ. decl.
(b) Exp. reg.
AIC = −459
rsq > 0.99
(c) P.law reg.
AIC = −178
rsq > 0.99
(d) E−P.l. reg.
AIC = −676
rsq > 0.99
(e) Null model test
A5
Null model
Observed
Predicted
Pairwise (i = 2)
1
0.891
0.794
5
10
15
Joint occupancy, J{i}
Number of species, i


File: 106.csv

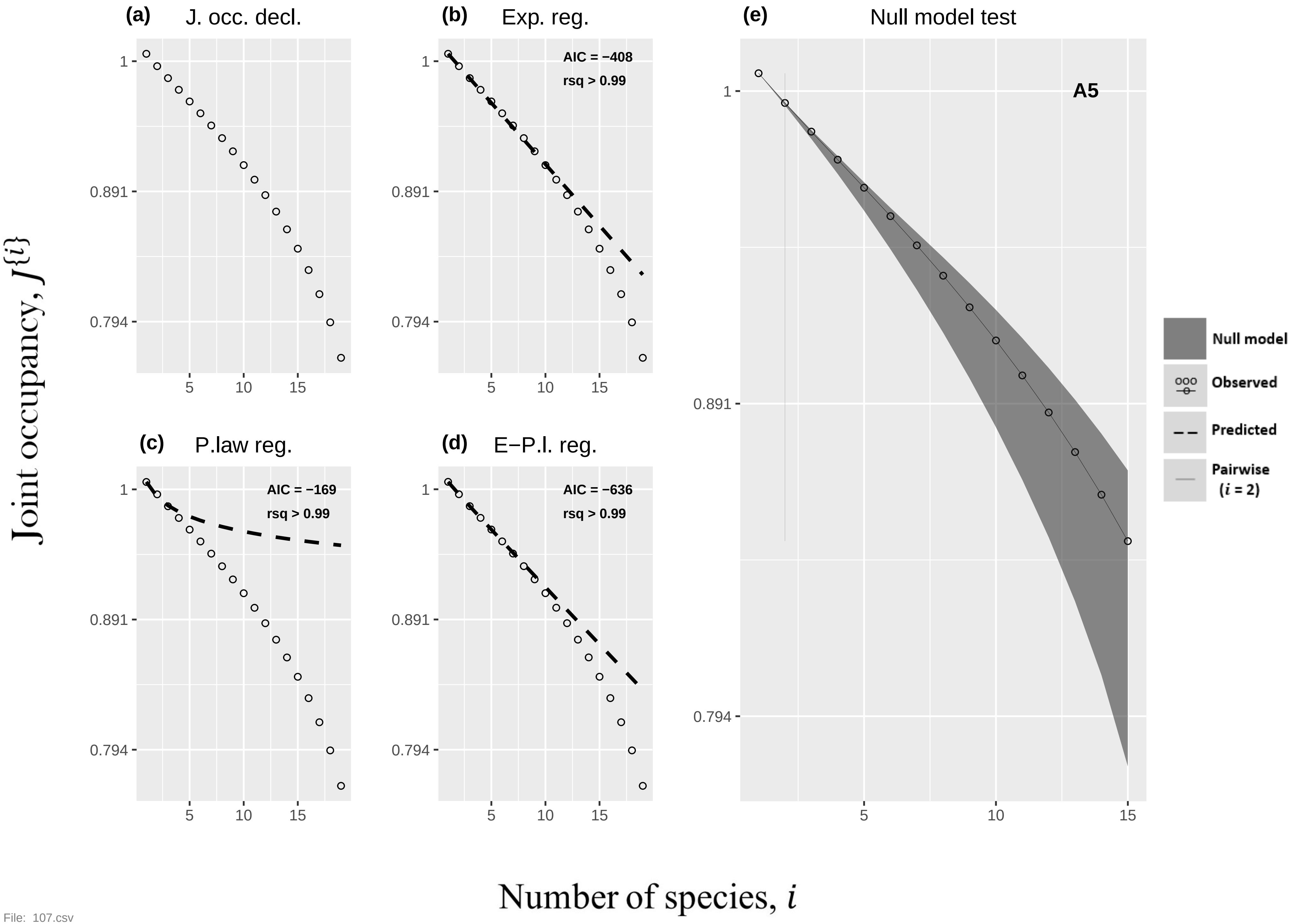

(a)
J. occ. decl.
(b)
Exp. reg.
AIC = −408
rsq > 0.99
(c)
P.law reg.
AIC = −169
rsq > 0.99
(d)
E−P.l. reg.
AIC = −636
rsq > 0.99
(e)
Null model test
A5
Null model
Observed
Predicted
Pairwise
(i = 2)
1
0.891
0.794
5
10
15
Joint occupancy, J{i}
Number of species, i


File: 107.csv

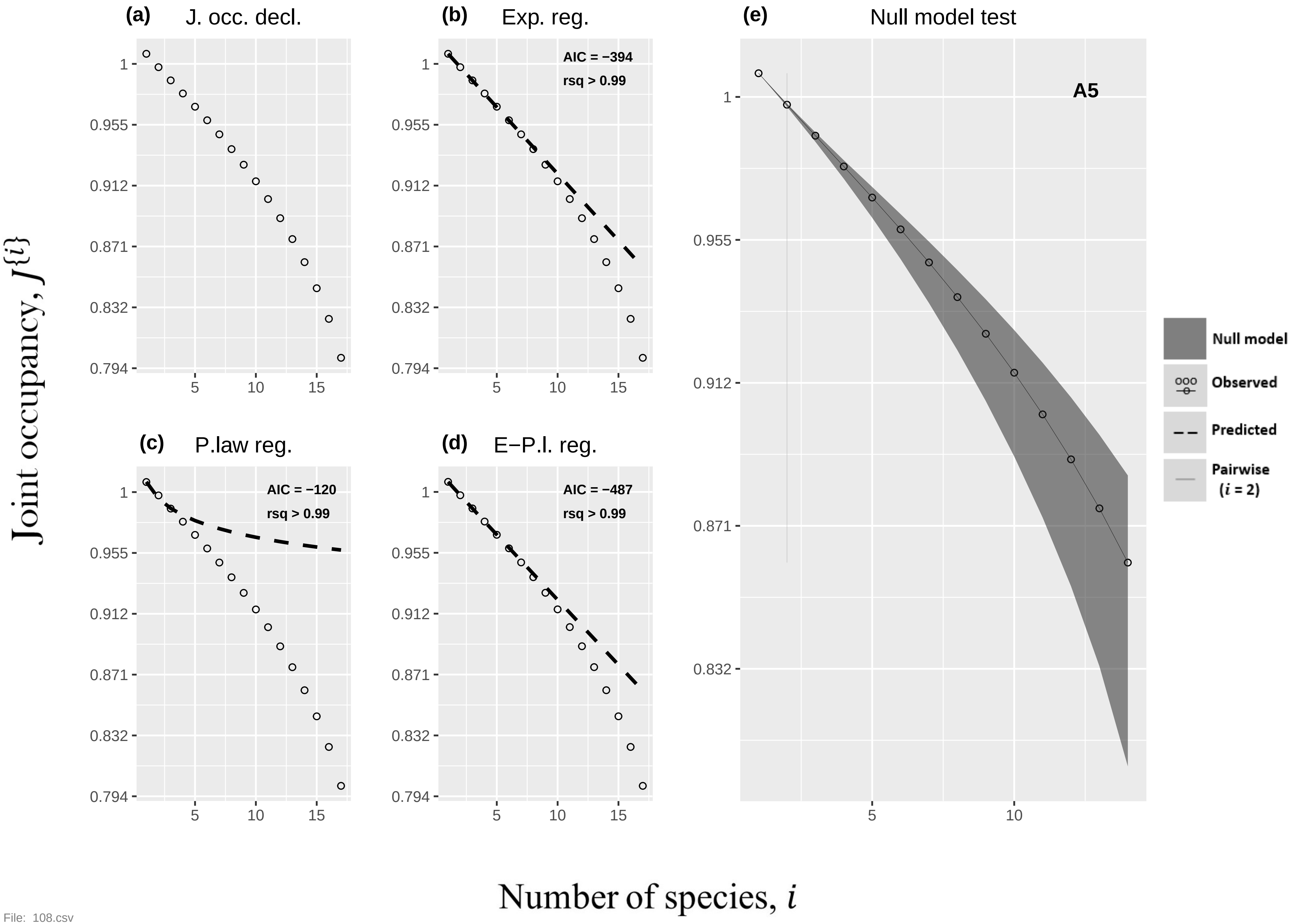
(a) J. occ. decl.
(b) Exp. reg.
AIC = −394
rsq > 0.99
(c) P.law reg.
AIC = −120
rsq > 0.99
(d) E−P.l. reg.
AIC = −487
rsq > 0.99
(e) Null model test
A5
Null model
Observed
Predicted
Pairwise (i = 2)
1
0.955
0.912
0.871
0.832
0.794
5
10
15
Joint occupancy, J{i}
Number of species, i

File: 108.csv

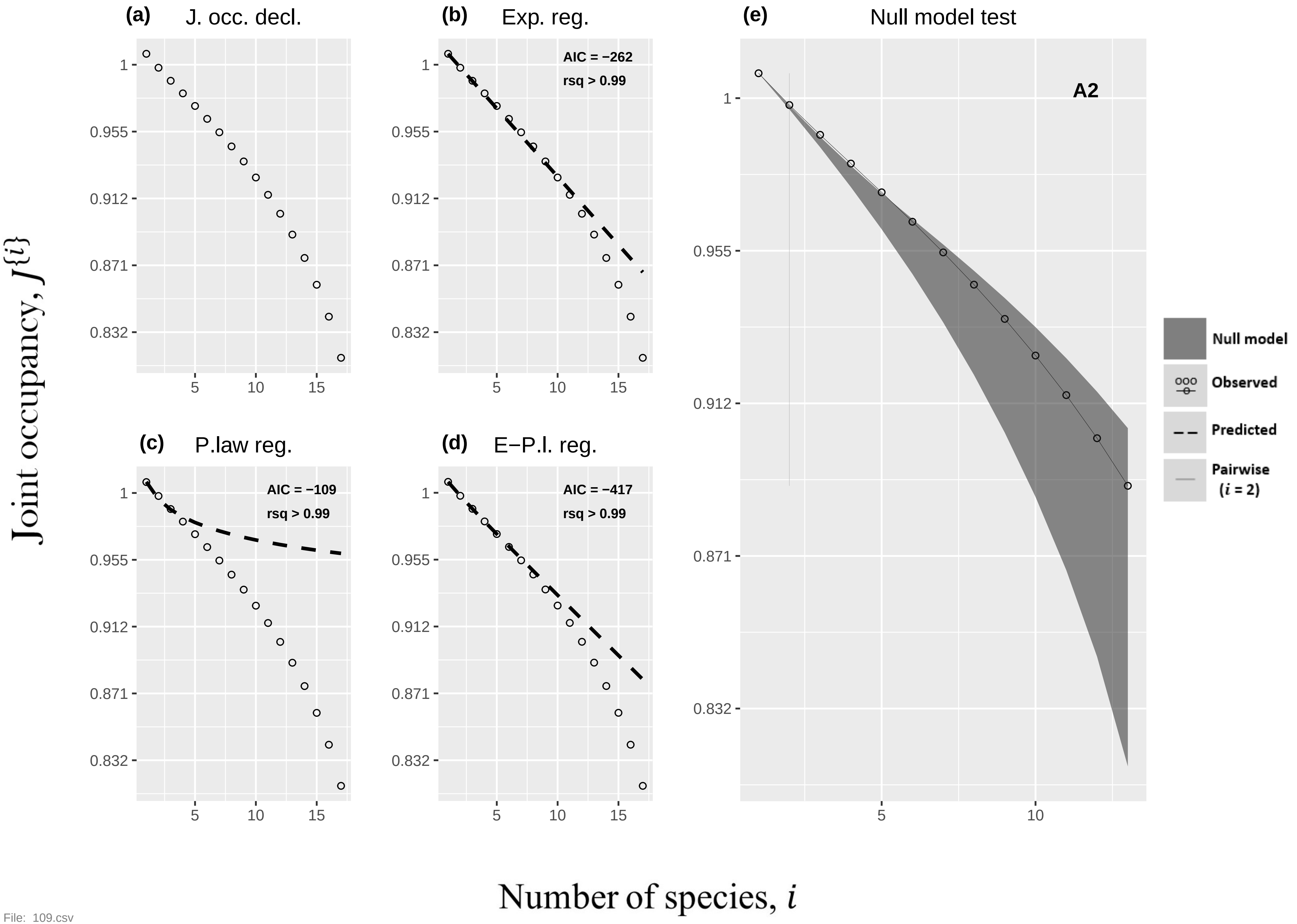

(a)
J. occ. decl.
(b)
Exp. reg.
AIC = −262
rsq > 0.99
(c)
P.law reg.
AIC = −109
rsq > 0.99
(d)
E−P.l. reg.
AIC = −417
rsq > 0.99
(e)
Null model test
A2
1
0.955
0.912
0.871
0.832
5
10
15
Null model
Observed
Predicted
Pairwise (i = 2)
Joint occupancy, J{i}
Number of species, i


File: 109.csv

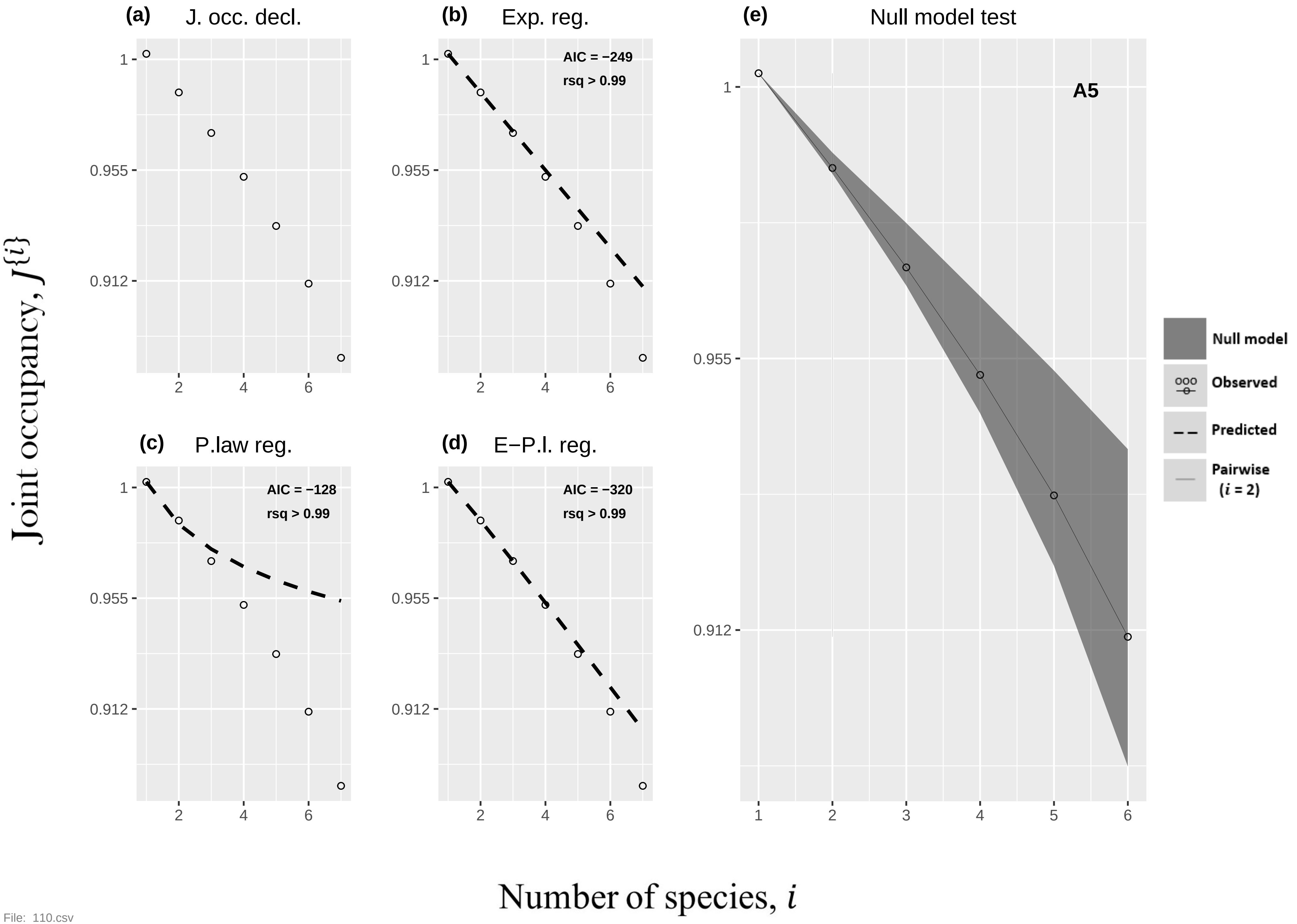

(a) J. occ. decl.
(b) Exp. reg.
AIC = −249
rsq > 0.99
(c) P.law reg.
AIC = −128
rsq > 0.99
(d) E−P.l. reg.
AIC = −320
rsq > 0.99
(e) Null model test
A5
Null model
Observed
Predicted
Pairwise (i = 2)
Joint occupancy, J{i}
Number of species, i
1
0.955
0.912
2
4
6
1
2
3
4
5
6

File: 110.csv

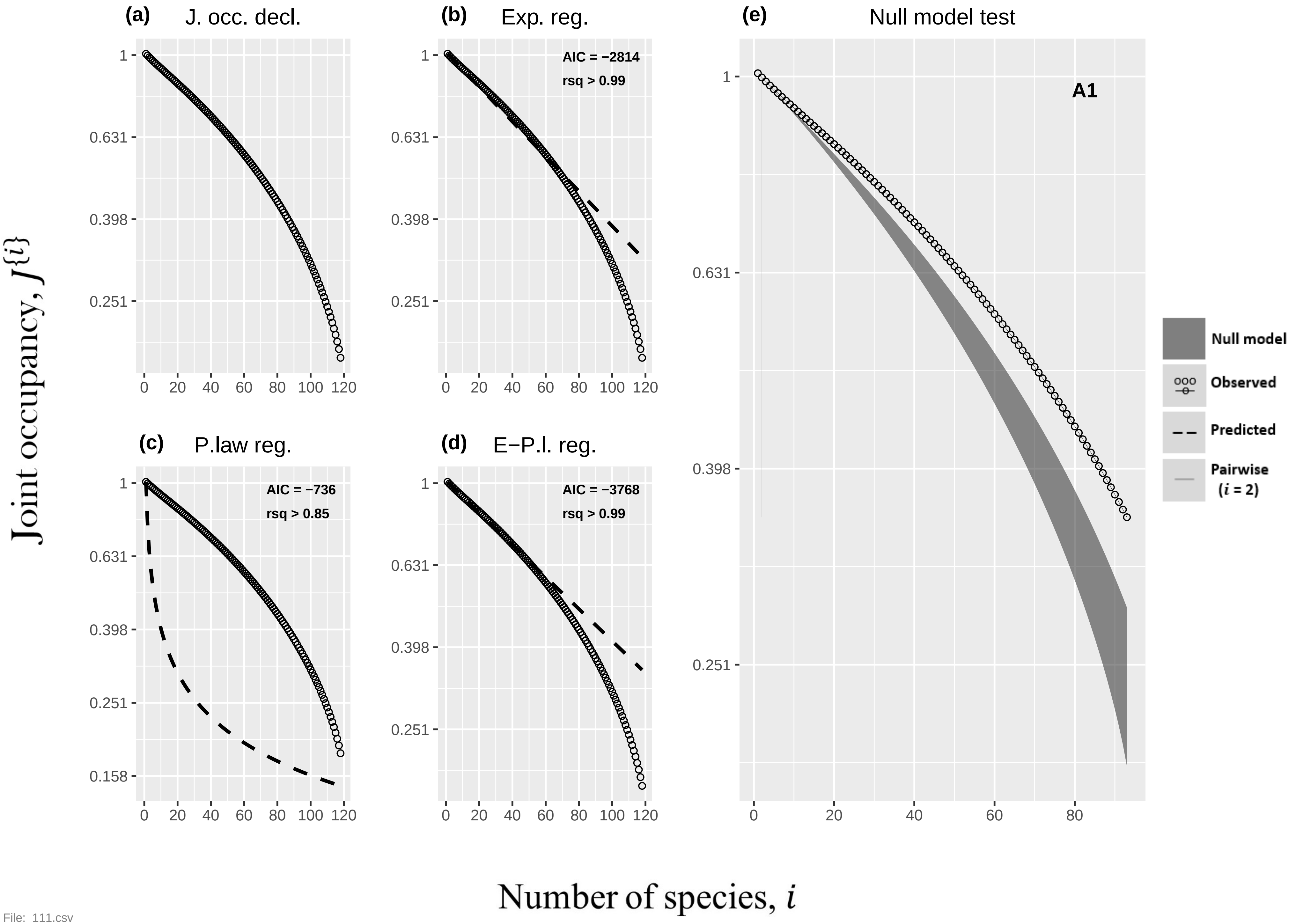

(a)
J. occ. decl.
(b)
Exp. reg.
AIC = −2814
rsq > 0.99
(c)
P.law reg.
AIC = −736
rsq > 0.85
(d)
E−P.l. reg.
AIC = −3768
rsq > 0.99
(e)
Null model test
A1
Null model
Observed
Predicted
Pairwise (i = 2)
Joint occupancy, J{i}
Number of species, i


File: 111.csv

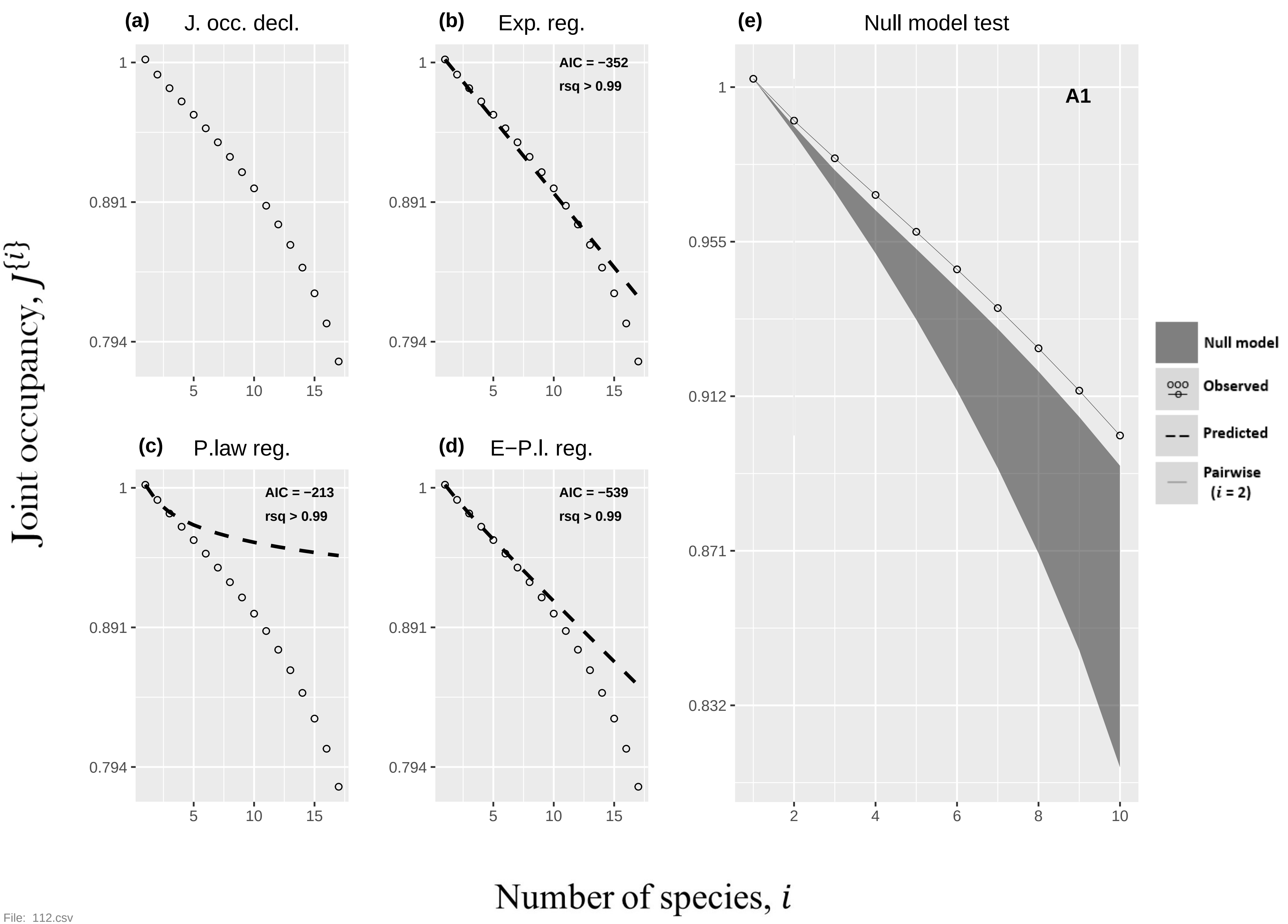

(a) J. occ. decl.
(b) Exp. reg.
AIC = −352
rsq > 0.99
(c) P.law reg.
AIC = −213
rsq > 0.99
(d) E−P.l. reg.
AIC = −539
rsq > 0.99
(e) Null model test
A1
Null model
Observed
Predicted
Pairwise ($i$ = 2)
1
0.955
0.912
0.871
0.832
0.891
0.794
5
10
15
2
4
6
8
Joint occupancy, $J\{i\}$
Number of species, $i$


File: 112.csv

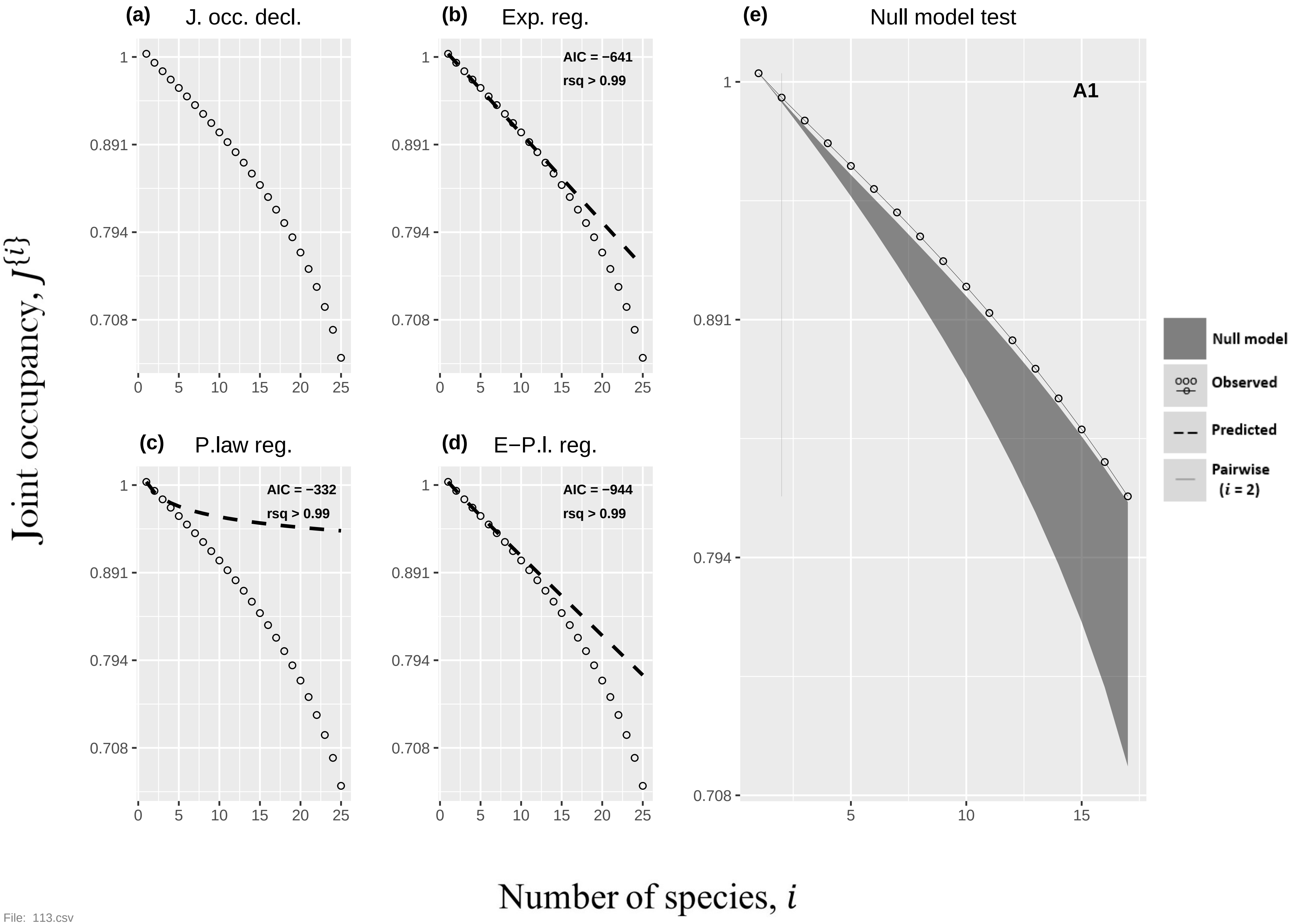

(a) J. occ. decl.
(b) Exp. reg.
AIC = −641
rsq > 0.99
(c) P.law reg.
AIC = −332
rsq > 0.99
(d) E−P.l. reg.
AIC = −944
rsq > 0.99
(e) Null model test
A1
1
0.891
0.794
0.708
0
5
10
15
20
25
Null model
Observed
Predicted
Pairwise
(i = 2)
Joint occupancy, J{i}
Number of species, i


File: 113.csv

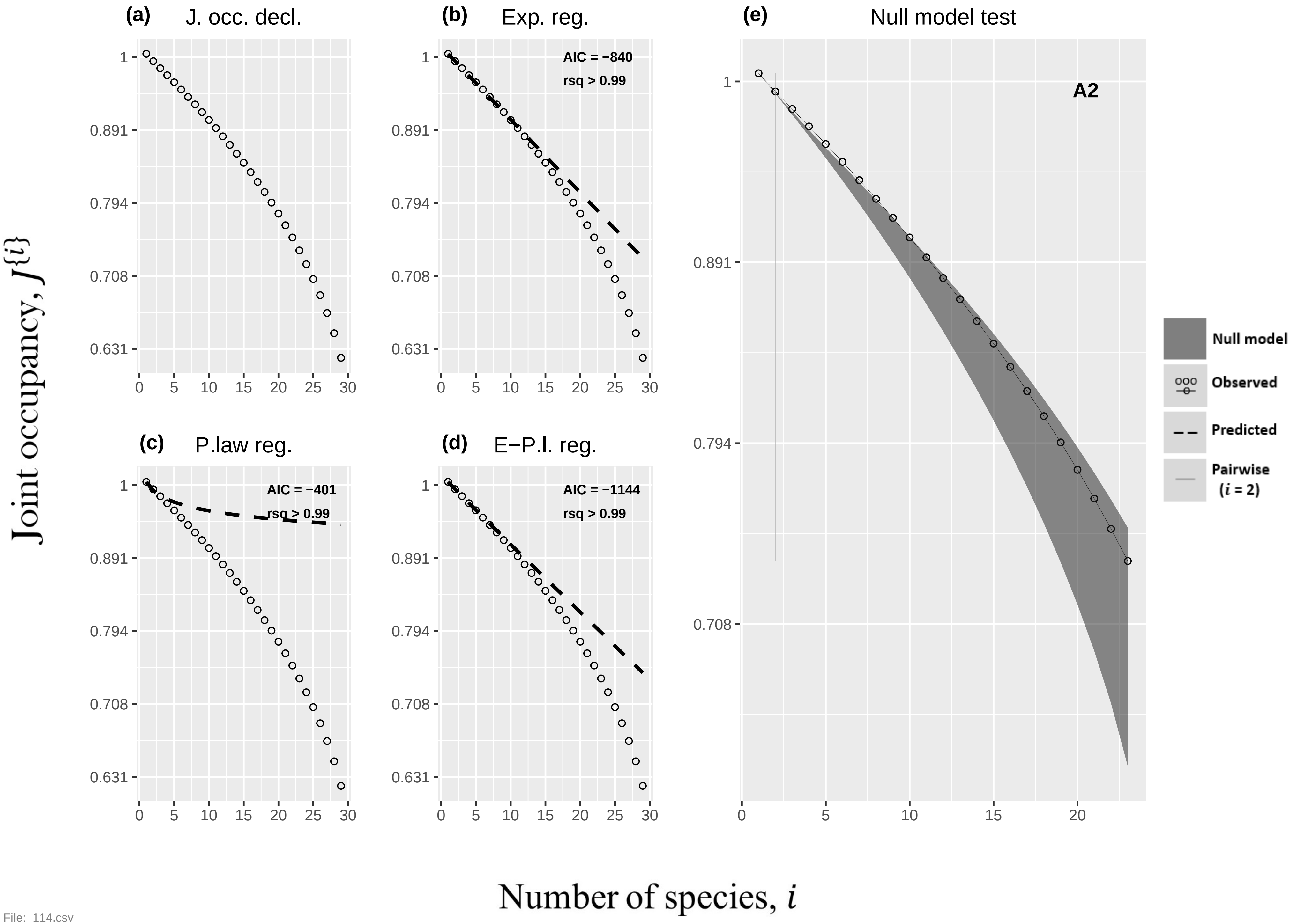

(a) J. occ. decl.
(b) Exp. reg.
AIC = −840
rsq > 0.99
(c) P.law reg.
AIC = −401
rsq > 0.99
(d) E−P.l. reg.
AIC = −1144
rsq > 0.99
(e) Null model test
A2
Null model
Observed
Predicted
Pairwise (i = 2)
1
0.891
0.794
0.708
0.631
0 5 10 15 20 25 30
Joint occupancy, J{i}
Number of species, i


File: 114.csv

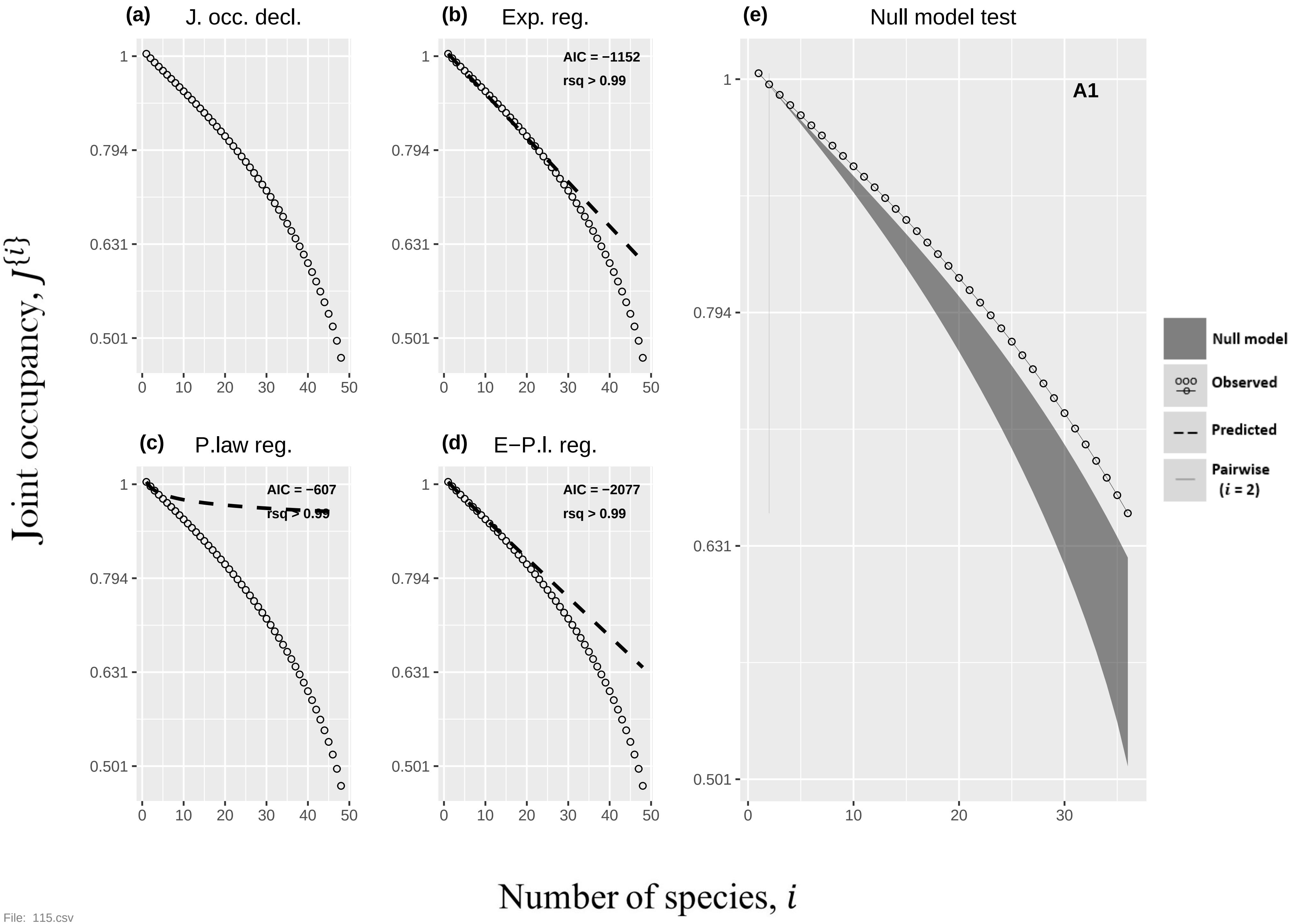

(a) J. occ. decl.
(b) Exp. reg.
AIC = −1152
rsq > 0.99
(c) P.law reg.
AIC = −607
rsq > 0.99
(d) E−P.l. reg.
AIC = −2077
rsq > 0.99
(e) Null model test
A1
Null model
Observed
Predicted
Pairwise (i = 2)
1
0.794
0.631
0.501
0
10
20
30
40
50
Joint occupancy, J{i}
Number of species, i


File: 115.csv

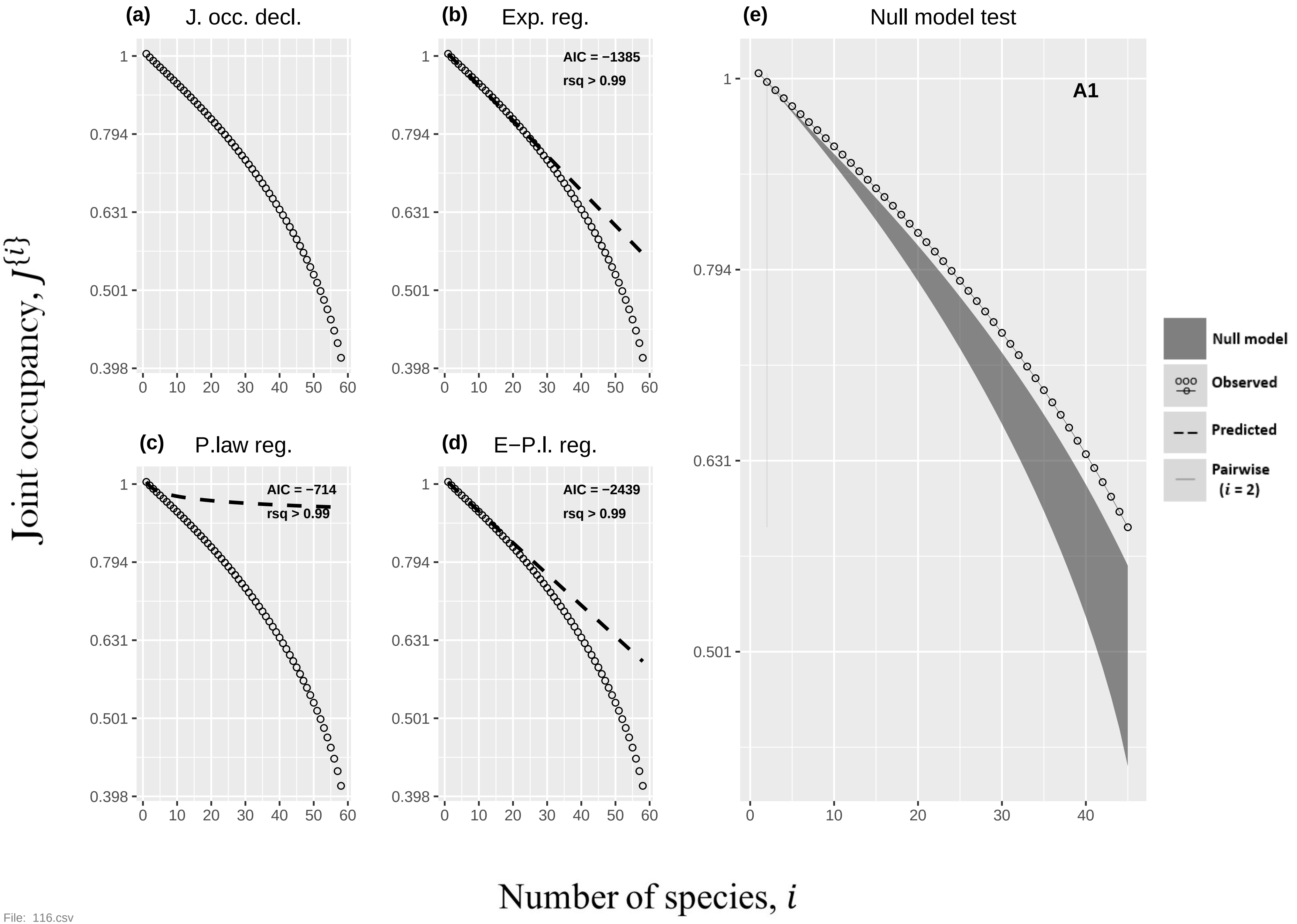

(a) J. occ. decl.
(b) Exp. reg.
AIC = −1385
rsq > 0.99
(c) P.law reg.
AIC = −714
rsq > 0.99
(d) E−P.l. reg.
AIC = −2439
rsq > 0.99
(e) Null model test
A1
Null model
Observed
Predicted
Pairwise ($i$ = 2)
Joint occupancy, $J\{i\}$
Number of species, $i$


File: 116.csv

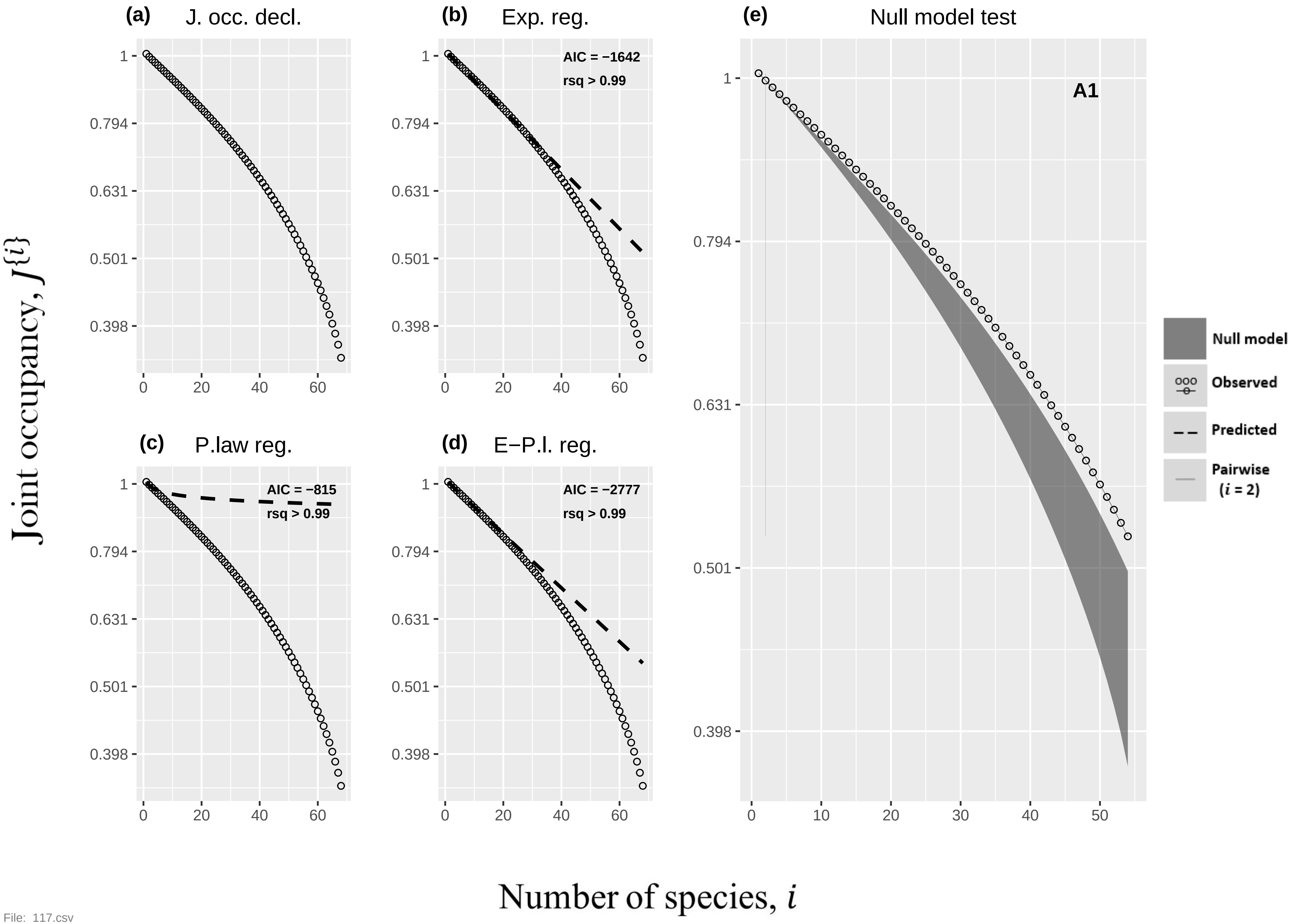
(a) J. occ. decl.
(b) Exp. reg.
AIC = −1642
rsq > 0.99
(c) P.law reg.
AIC = −815
rsq > 0.99
(d) E−P.l. reg.
AIC = −2777
rsq > 0.99
(e) Null model test
A1
Null model
Observed
Predicted
Pairwise (i = 2)
Joint occupancy, J{i}
Number of species, i
1
0.794
0.631
0.501
0.398
0
10
20
30
40
50
60

File: 117.csv

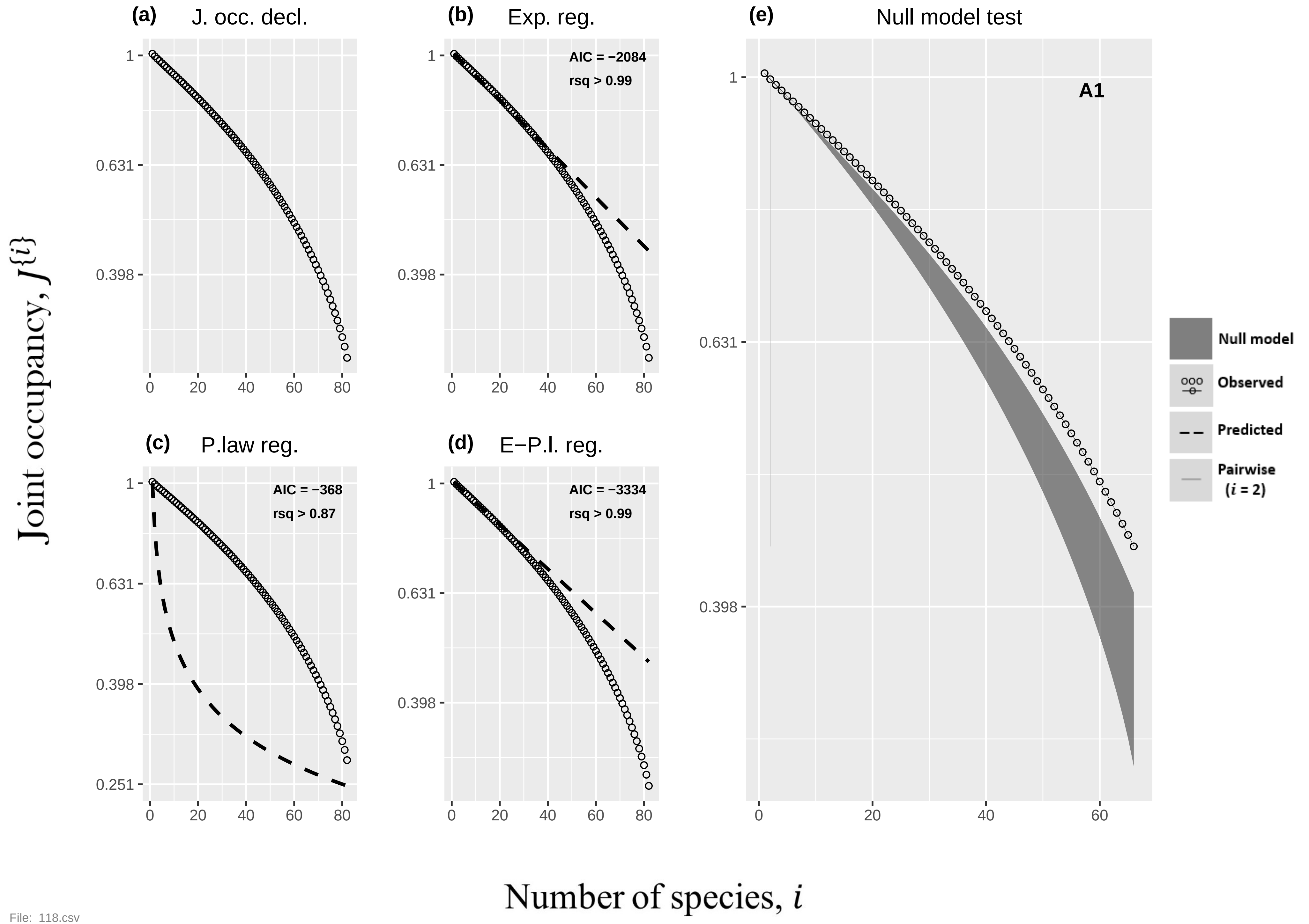

(a) J. occ. decl.
(b) Exp. reg.
AIC = −2084
rsq > 0.99
(c) P.law reg.
AIC = −368
rsq > 0.87
(d) E−P.l. reg.
AIC = −3334
rsq > 0.99
(e) Null model test
A1
Null model
Observed
Predicted
Pairwise
($i$ = 2)
Joint occupancy, $J^{\{i\}}$
Number of species, $i$


File: 118.csv

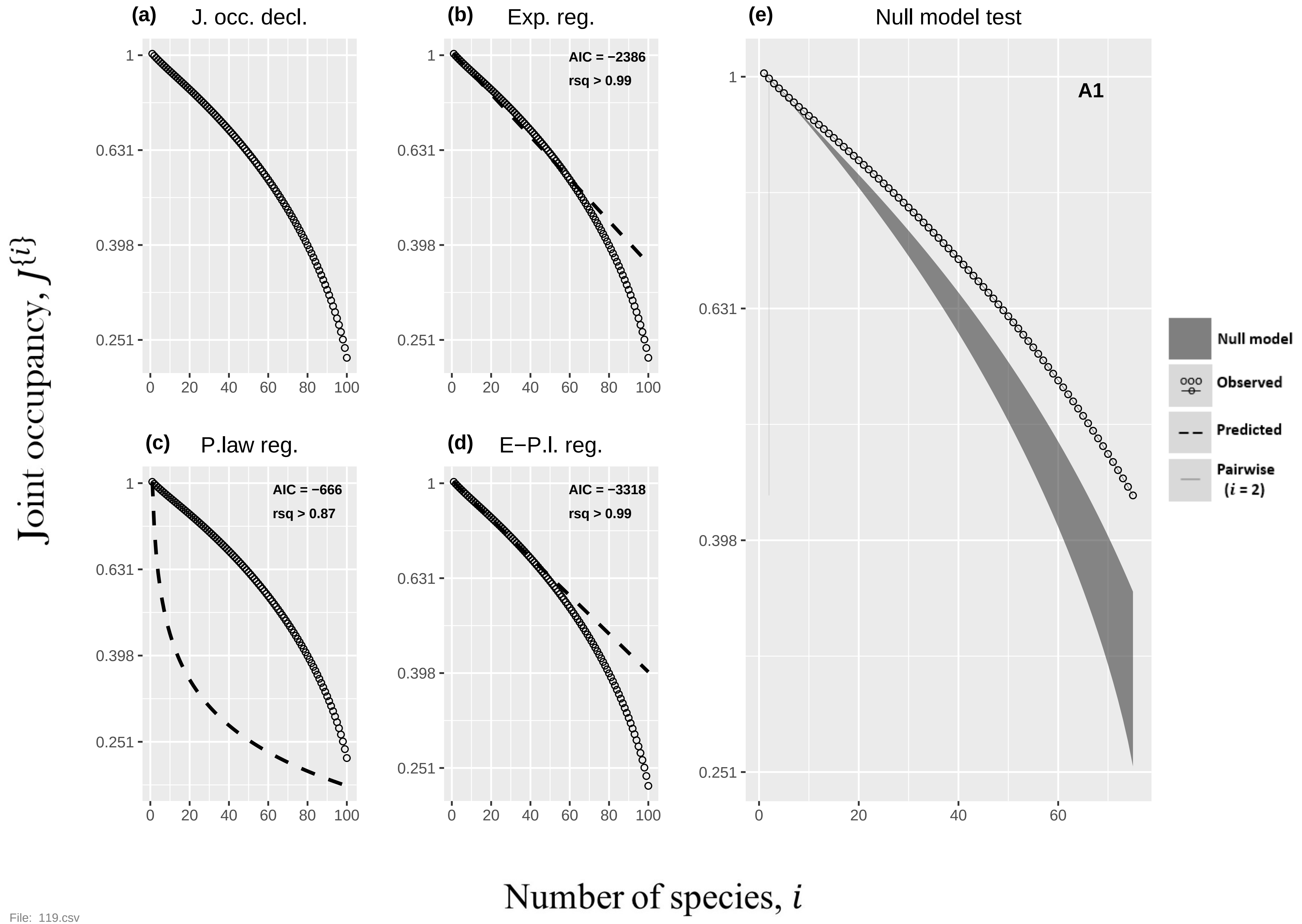
(a) J. occ. decl.
(b) Exp. reg.
AIC = −2386
rsq > 0.99
(e) Null model test
A1
(c) P.law reg.
AIC = −666
rsq > 0.87
(d) E−P.l. reg.
AIC = −3318
rsq > 0.99
Null model
Observed
Predicted
Pairwise ($i$ = 2)
Joint occupancy, $J^{\{i\}}$
Number of species, $i$
1
0.631
0.398
0.251
0
20
40
60
80
100

File: 119.csv

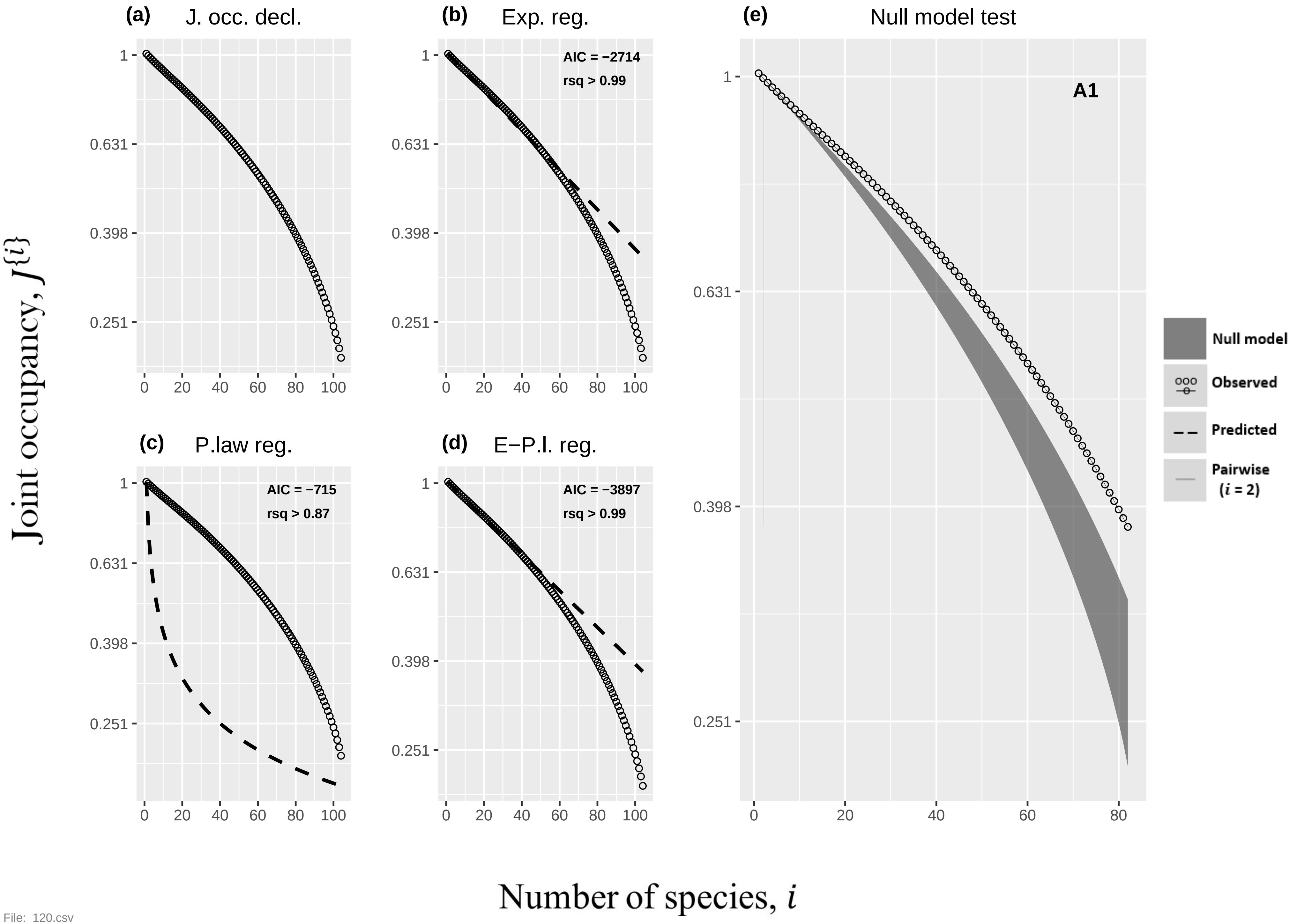

(a) J. occ. decl.
(b) Exp. reg.
AIC = −2714
rsq > 0.99
(c) P.law reg.
AIC = −715
rsq > 0.87
(d) E−P.l. reg.
AIC = −3897
rsq > 0.99
(e) Null model test
A1
Null model
Observed
Predicted
Pairwise
(i = 2)
Joint occupancy, J{i}
Number of species, i
1
0.631
0.398
0.251
0
20
40
60
80
100


File: 120.csv

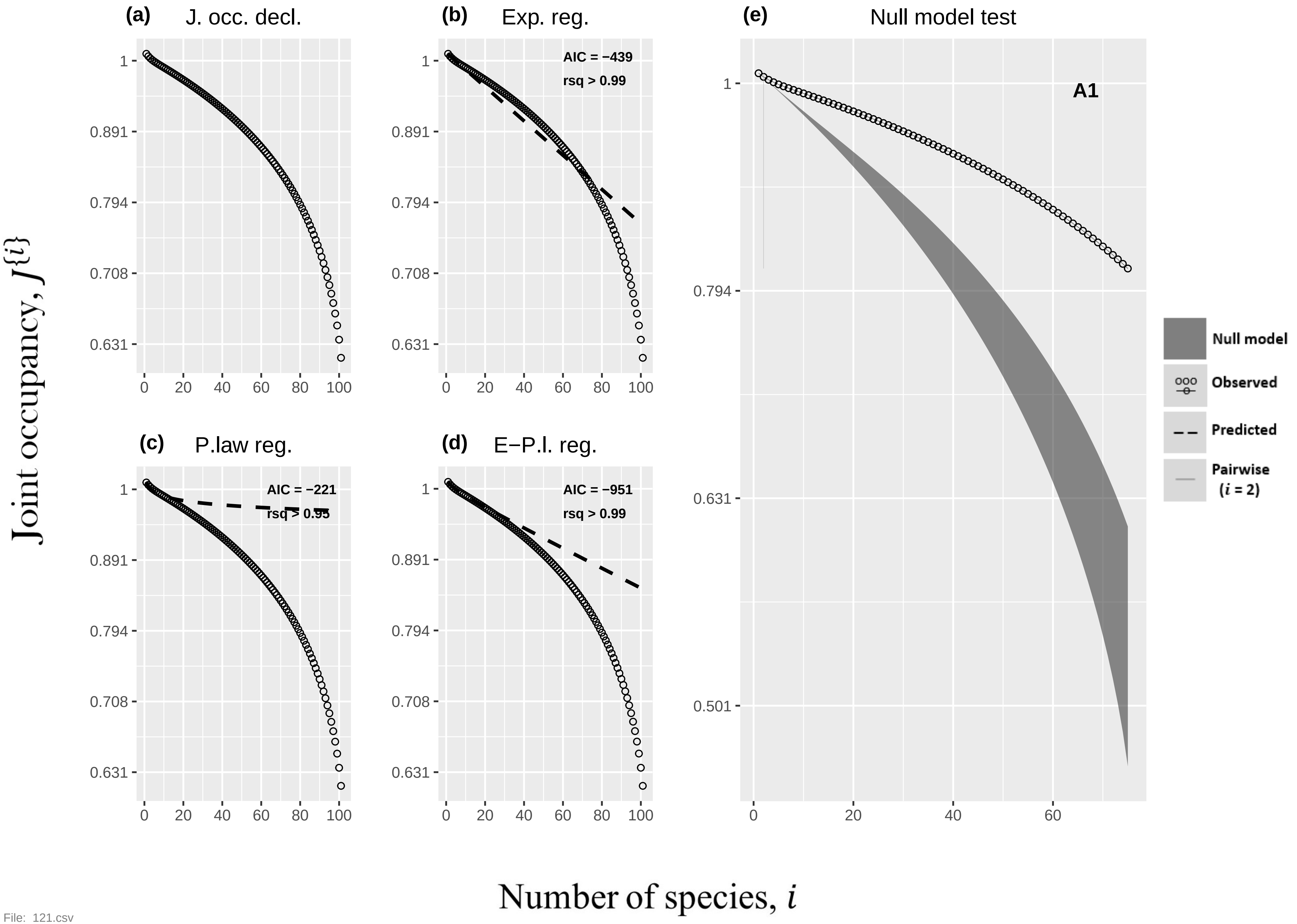

(a) J. occ. decl.
(b) Exp. reg.
AIC = −439
rsq > 0.99
(c) P.law reg.
AIC = −221
rsq > 0.95
(d) E−P.l. reg.
AIC = −951
rsq > 0.99
(e) Null model test
A1
Null model
Observed
Predicted
Pairwise (i = 2)
1
0.891
0.794
0.708
0.631
0.501
0 20 40 60 80 100
Joint occupancy, J{i}
Number of species, i


File: 121.csv

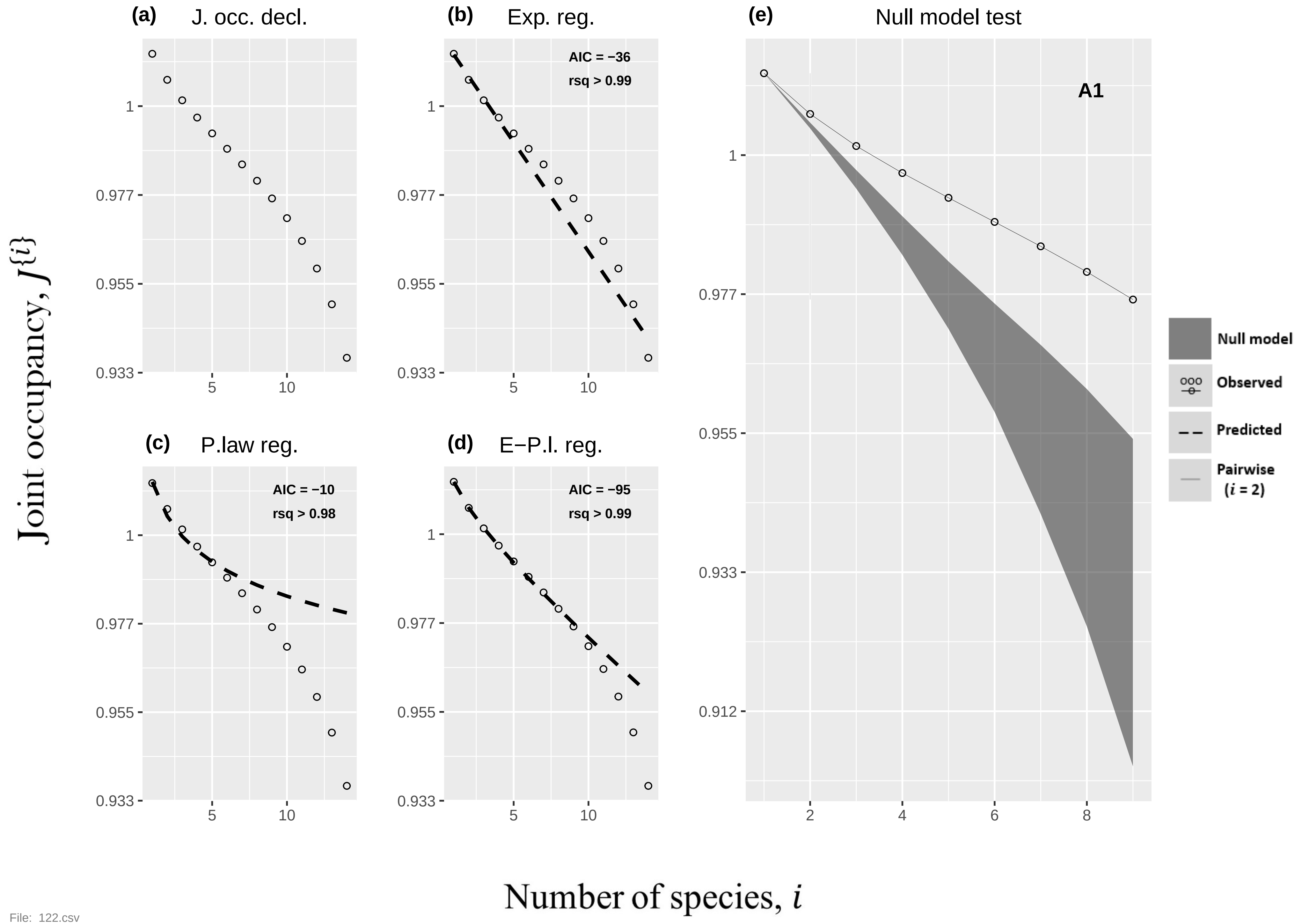

(a)
J. occ. decl.
(b)
Exp. reg.
AIC = −36
rsq > 0.99
(c)
P.law reg.
AIC = −10
rsq > 0.98
(d)
E−P.l. reg.
AIC = −95
rsq > 0.99
(e)
Null model test
A1
Null model
Observed
Predicted
Pairwise
(i = 2)
1
0.977
0.955
0.933
0.912
5
10
2
4
6
8
Joint occupancy, J{i}
Number of species, i


File: 122.csv

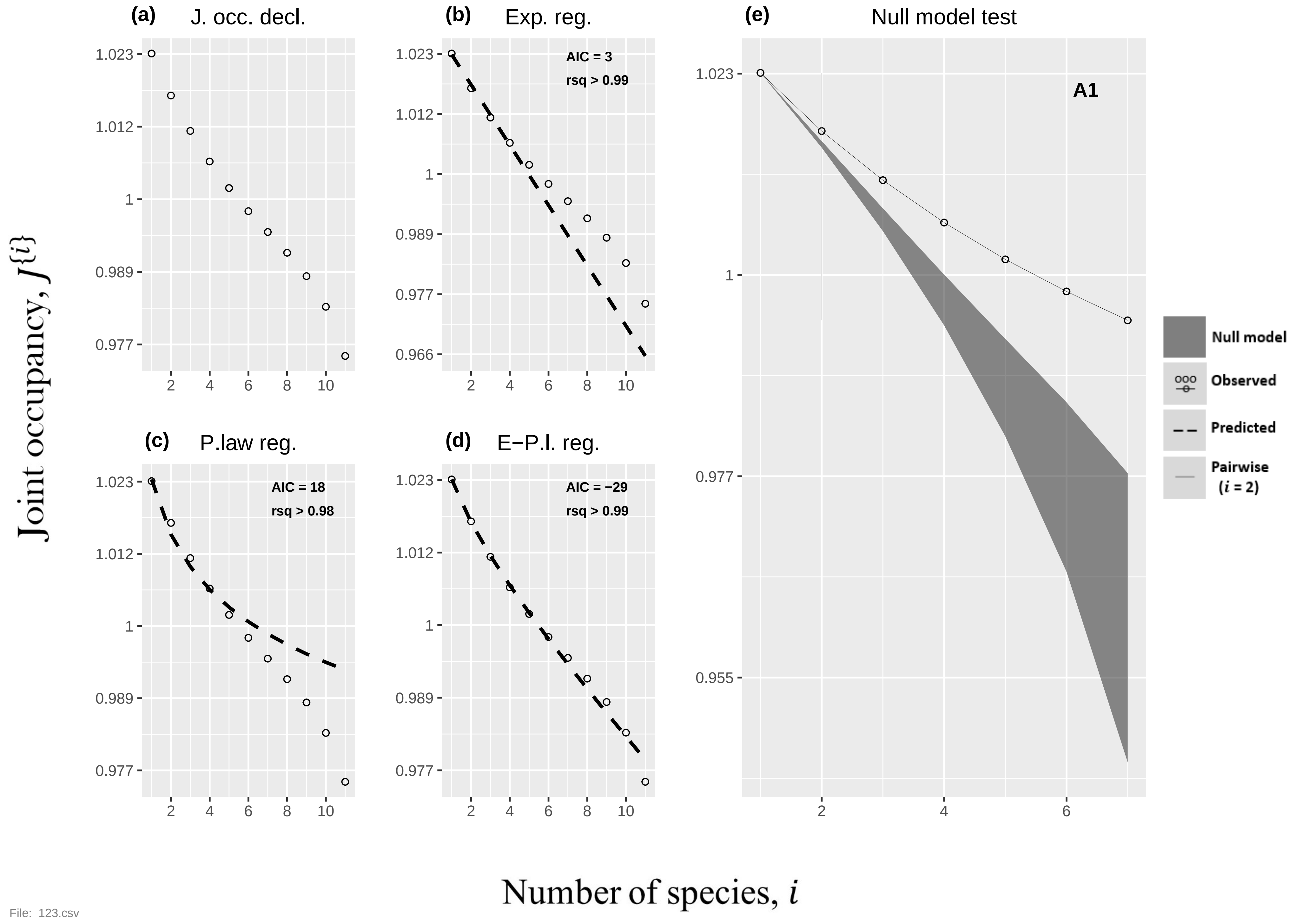
(a) J. occ. decl.
(b) Exp. reg.
AIC = 3
rsq > 0.99
(c) P.law reg.
AIC = 18
rsq > 0.98
(d) E−P.l. reg.
AIC = −29
rsq > 0.99
(e) Null model test
A1
Null model
Observed
Predicted
Pairwise
($i$ = 2)
Joint occupancy, $J^{\{i\}}$
Number of species, $i$
File: 123.csv

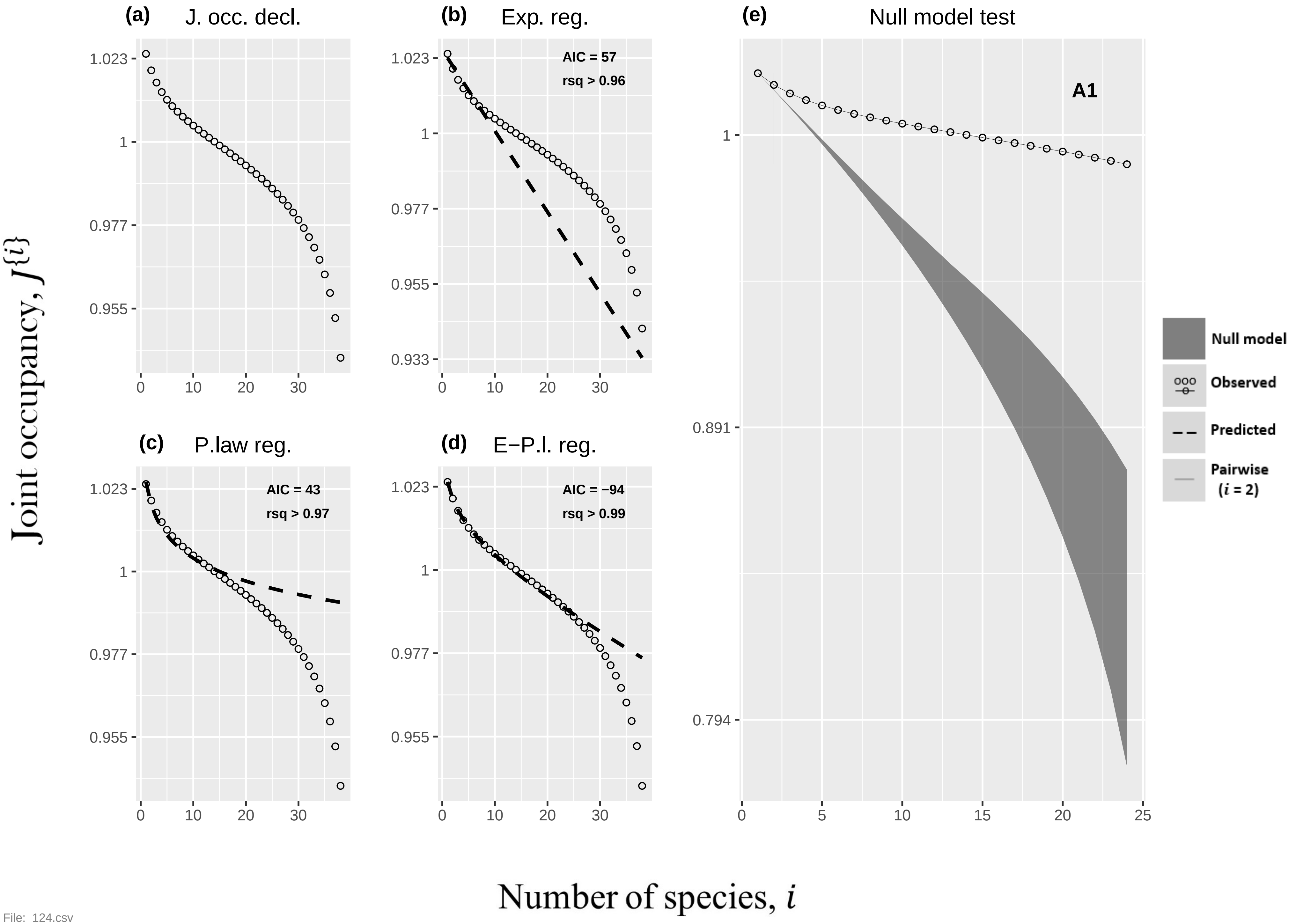

(a) J. occ. decl.
(b) Exp. reg.
AIC = 57
rsq > 0.96
(e) Null model test
A1
(c) P.law reg.
AIC = 43
rsq > 0.97
(d) E−P.l. reg.
AIC = −94
rsq > 0.99
Null model
Observed
Predicted
Pairwise (i = 2)
Joint occupancy, J{i}
Number of species, i
1.023
1
0.977
0.955
0.933
0.891
0.794
0
5
10
15
20
25
30


File: 124.csv

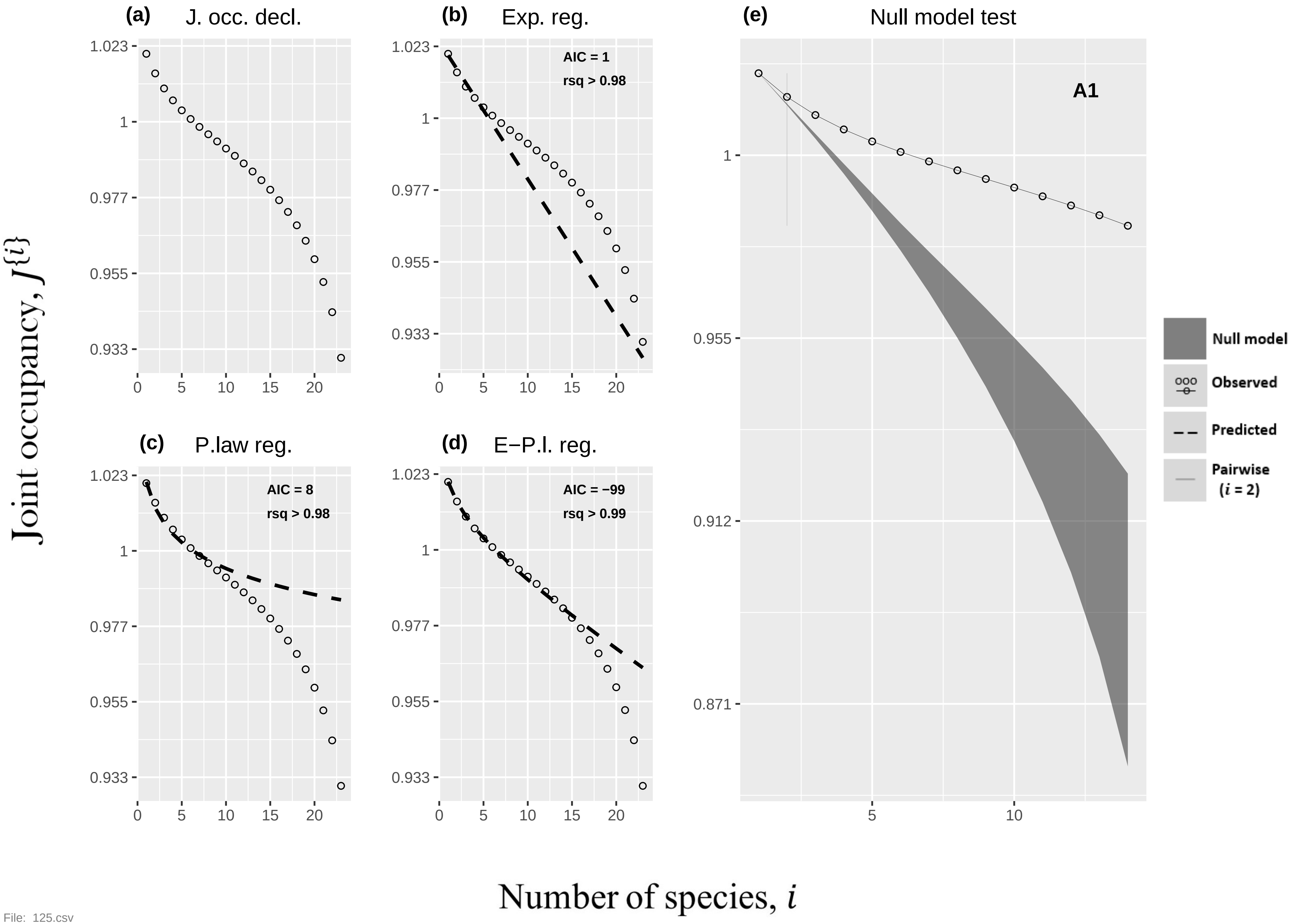
(a)
J. occ. decl.
(b)
Exp. reg.
AIC = 1
rsq > 0.98
(c)
P.law reg.
AIC = 8
rsq > 0.98
(d)
E−P.l. reg.
AIC = −99
rsq > 0.99
(e)
Null model test
A1
Null model
Observed
Predicted
Pairwise
(i = 2)
1.023
1
0.977
0.955
0.933
0.912
0.871
0
5
10
15
20
Joint occupancy, J{i}
Number of species, i

File: 125.csv

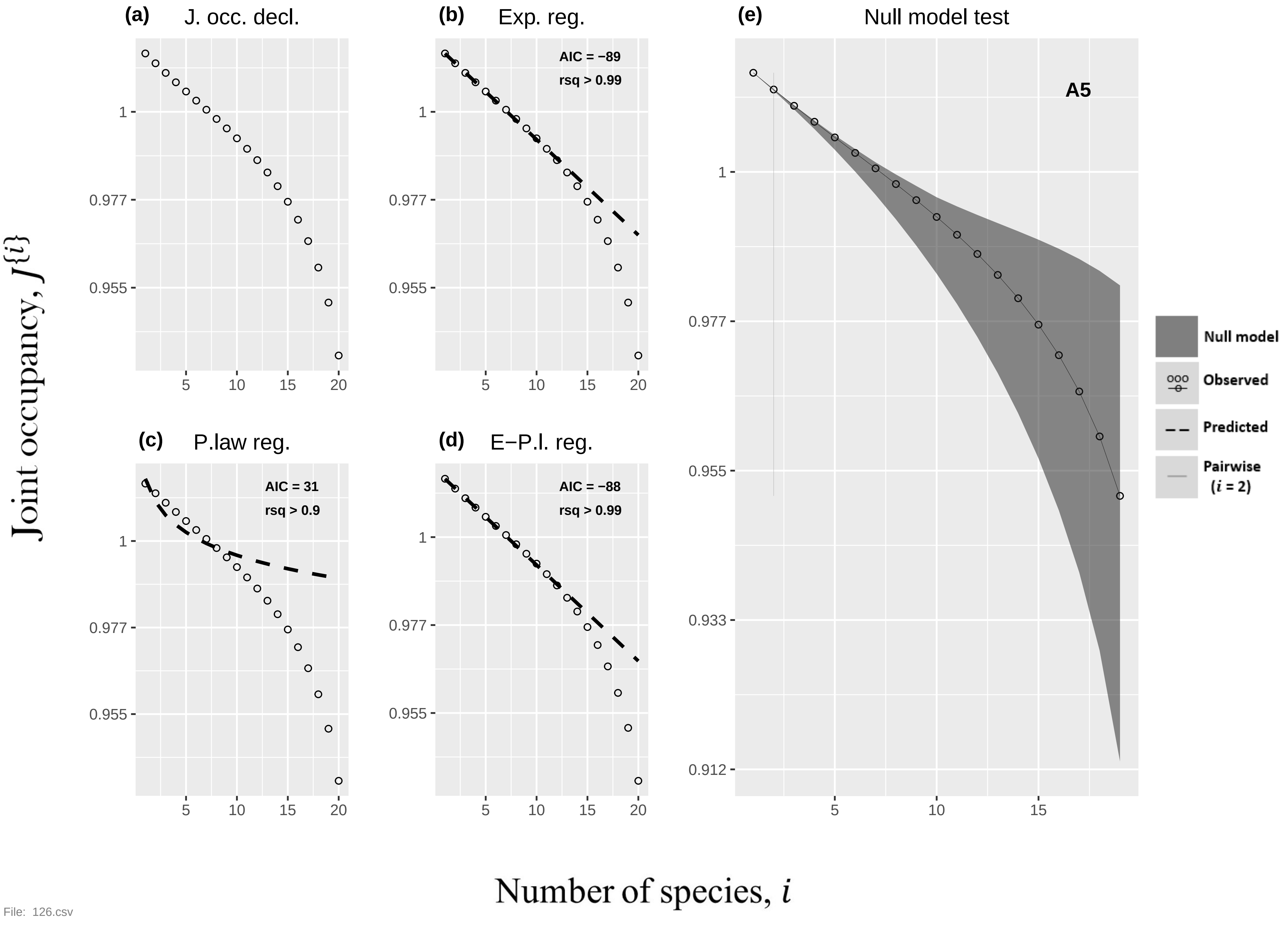
(a)
J. occ. decl.
(b)
Exp. reg.
AIC = −89
rsq > 0.99
(c)
P.law reg.
AIC = 31
rsq > 0.9
(d)
E−P.l. reg.
AIC = −88
rsq > 0.99
(e)
Null model test
A5
Null model
Observed
Predicted
Pairwise (i = 2)
1
0.977
0.955
0.933
0.912
5
10
15
20
Joint occupancy, J{i}
Number of species, i
File: 126.csv

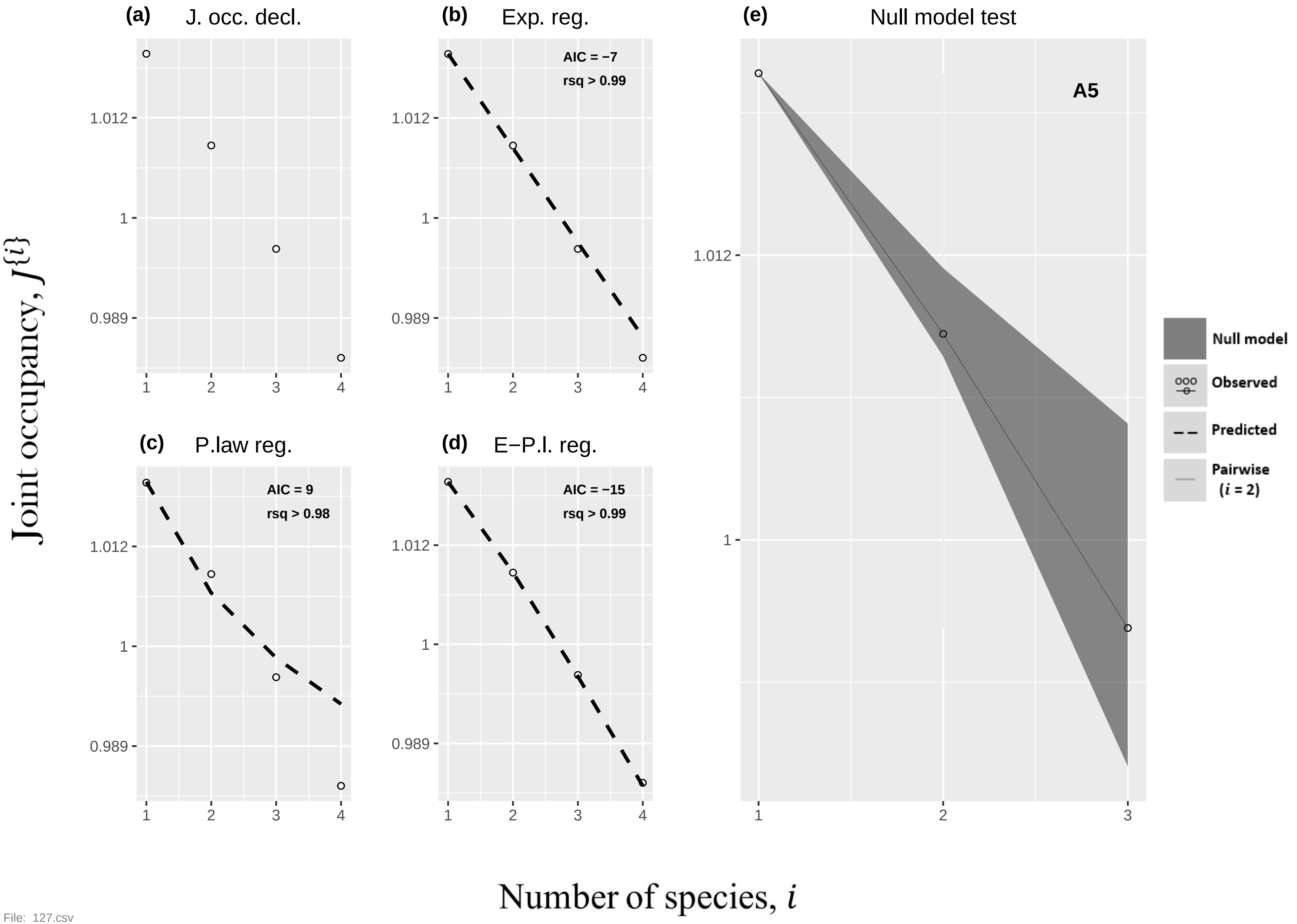

(a) J. occ. decl.
(b) Exp. reg.
AIC = −7
rsq > 0.99
(c) P.law reg.
AIC = 9
rsq > 0.98
(d) E−P.l. reg.
AIC = −15
rsq > 0.99
(e) Null model test
A5
Null model
Observed
Predicted
Pairwise (i = 2)
1.012
1
0.989
1
2
3
4
Joint occupancy, J{i}
Number of species, i


File: 127.csv

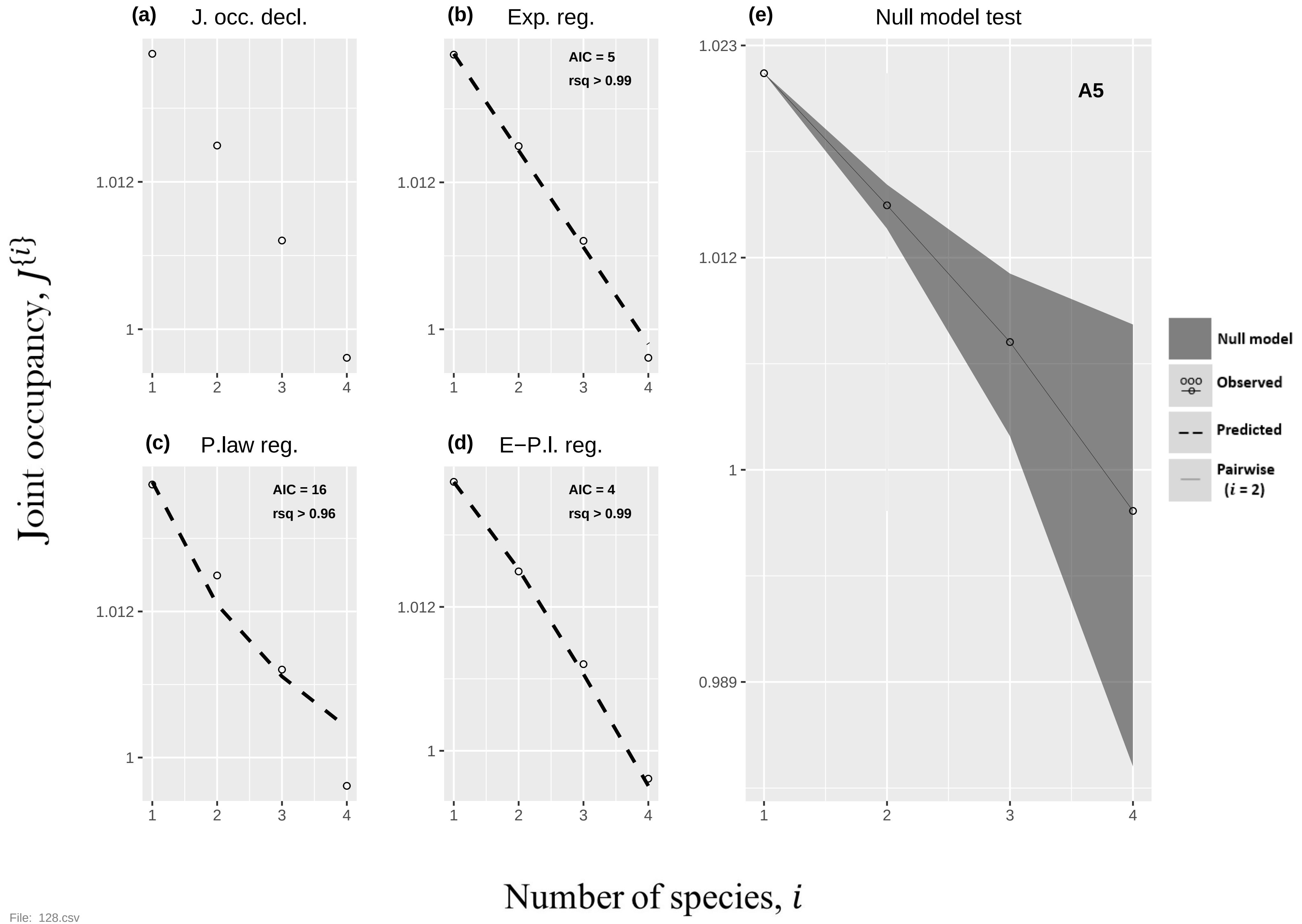
(a) J. occ. decl.
(b) Exp. reg.
AIC = 5
rsq > 0.99
(c) P.law reg.
AIC = 16
rsq > 0.96
(d) E−P.l. reg.
AIC = 4
rsq > 0.99
(e) Null model test
A5
Null model
Observed
Predicted
Pairwise (i = 2)
1.023
1.012
1
0.989
1 2 3 4
Joint occupancy, J{i}
Number of species, i

File: 128.csv

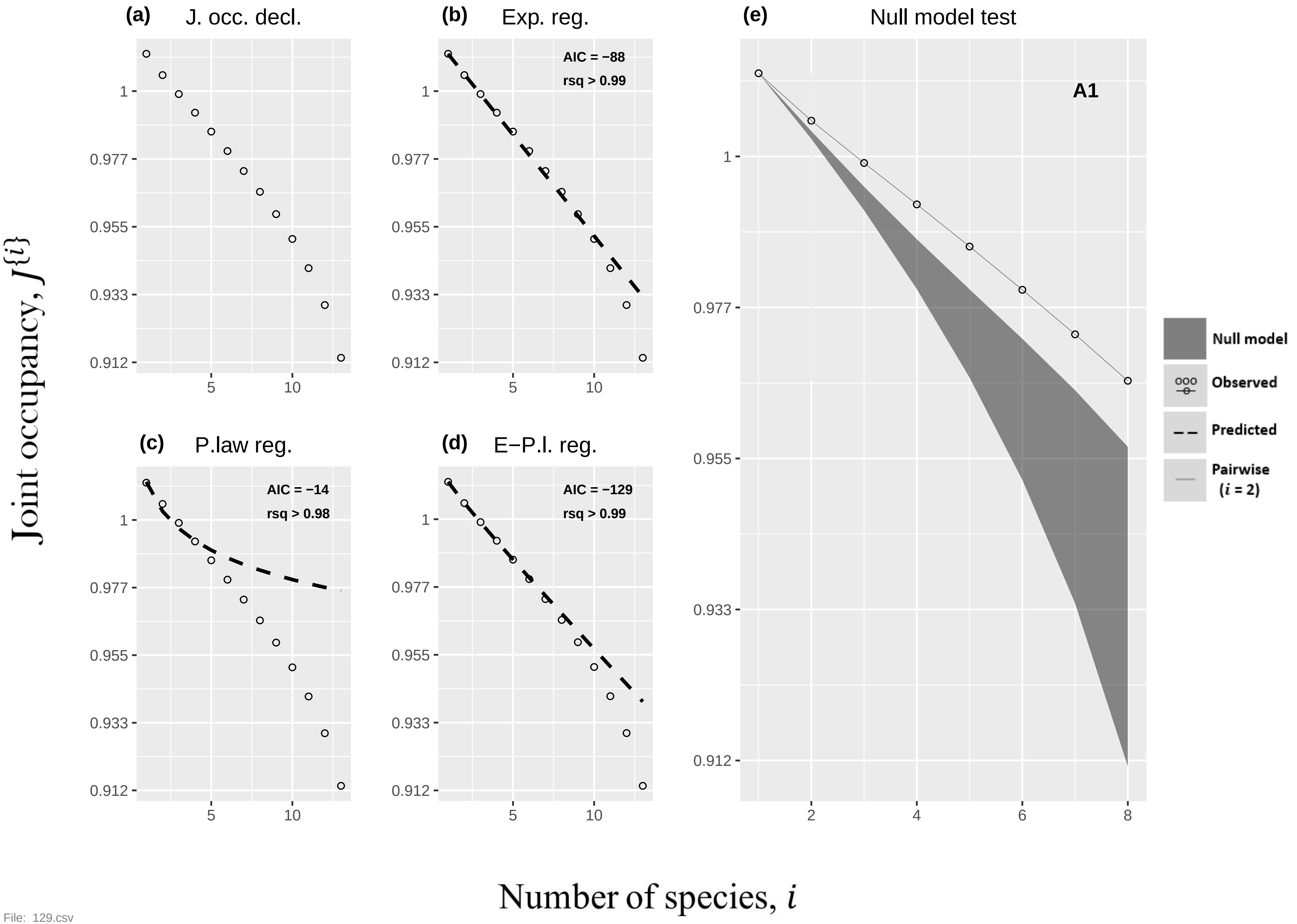
(a) J. occ. decl.
(b) Exp. reg.
AIC = −88
rsq > 0.99
(e) Null model test
A1
(c) P.law reg.
AIC = −14
rsq > 0.98
(d) E−P.l. reg.
AIC = −129
rsq > 0.99
Null model
Observed
Predicted
Pairwise (i = 2)
Joint occupancy, J{i}
Number of species, i
1
0.977
0.955
0.933
0.912
5
10
2
4
6
8

File: 129.csv

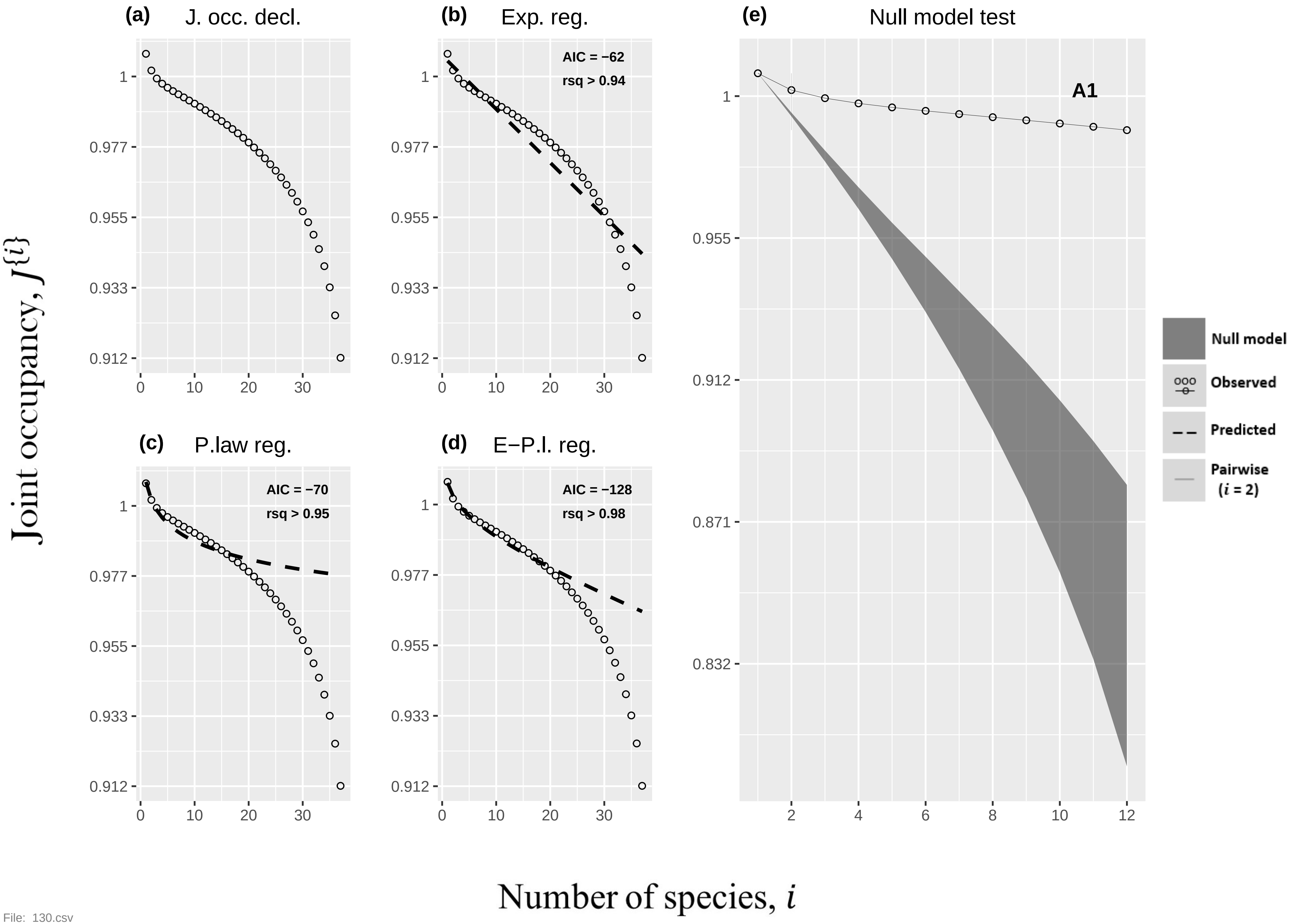

(a) J. occ. decl.
(b) Exp. reg.
AIC = −62
rsq > 0.94
(c) P.law reg.
AIC = −70
rsq > 0.95
(d) E−P.l. reg.
AIC = −128
rsq > 0.98
(e) Null model test
A1
Null model
Observed
Predicted
Pairwise (i = 2)
Joint occupancy, J{i}
Number of species, i


File: 130.csv

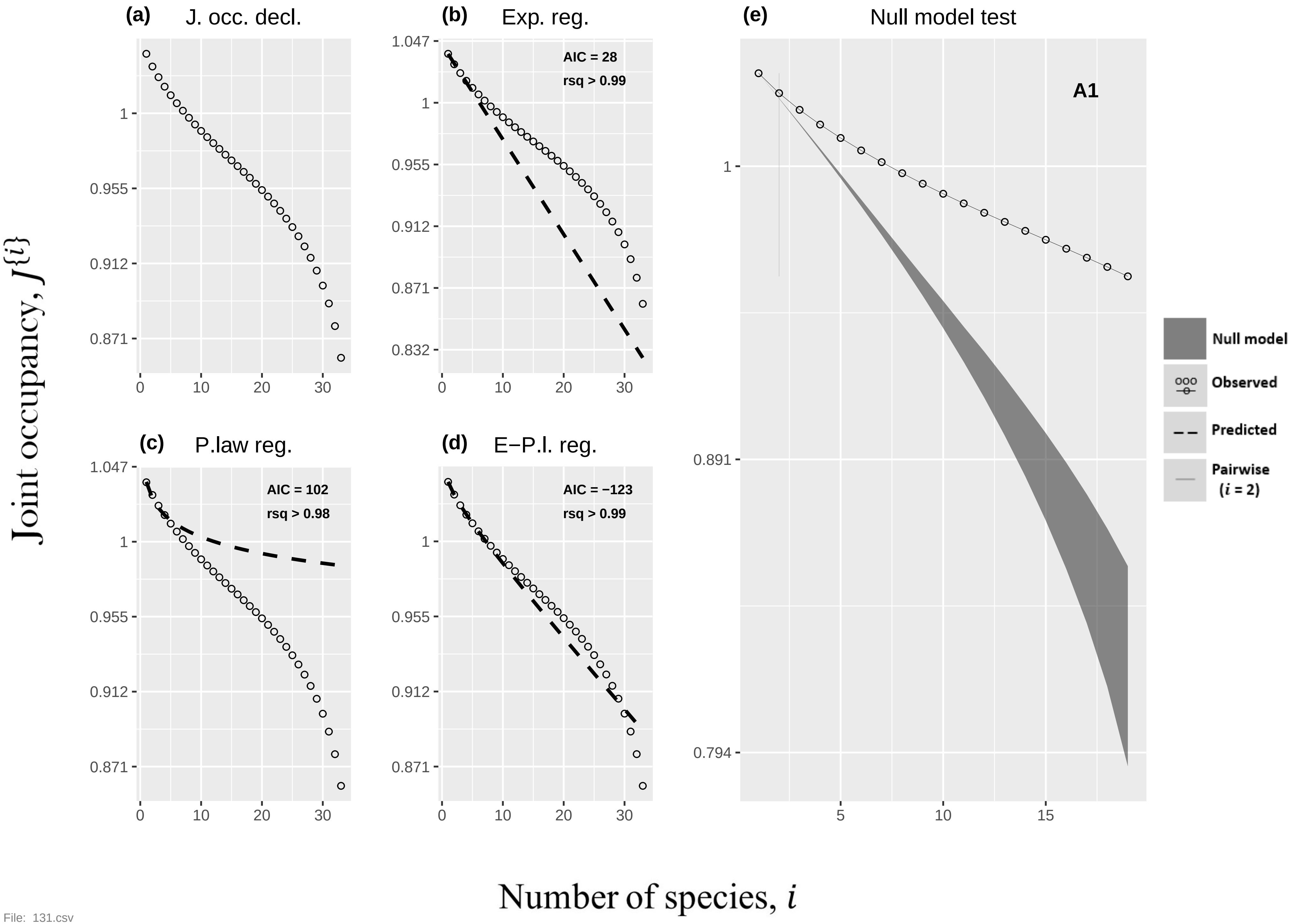

(a) J. occ. decl.
(b) Exp. reg.
AIC = 28
rsq > 0.99
(c) P.law reg.
AIC = 102
rsq > 0.98
(d) E−P.l. reg.
AIC = −123
rsq > 0.99
(e) Null model test
A1
Null model
Observed
Predicted
Pairwise (i = 2)
1.047
1
0.955
0.912
0.871
0.832
0.891
0.794
0
10
20
30
5
15
Joint occupancy, J{i}
Number of species, i


File: 131.csv

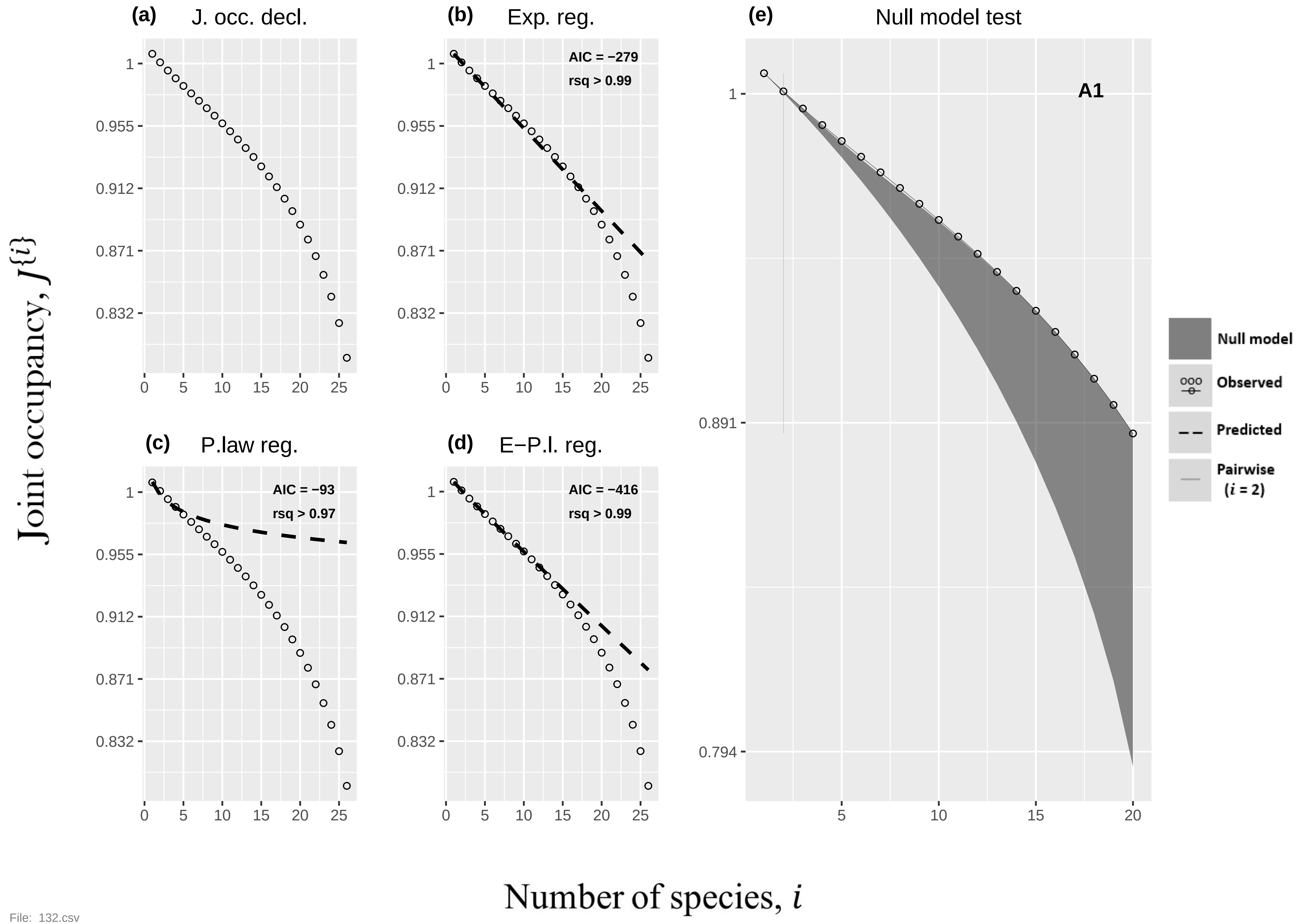

(a) J. occ. decl.
(b) Exp. reg.
AIC = −279
rsq > 0.99
(c) P.law reg.
AIC = −93
rsq > 0.97
(d) E−P.l. reg.
AIC = −416
rsq > 0.99
(e) Null model test
A1
Null model
Observed
Predicted
Pairwise (i = 2)
Joint occupancy, J{i}
Number of species, i
1
0.955
0.912
0.871
0.832
0.891
0.794
0
5
10
15
20
25


File: 132.csv

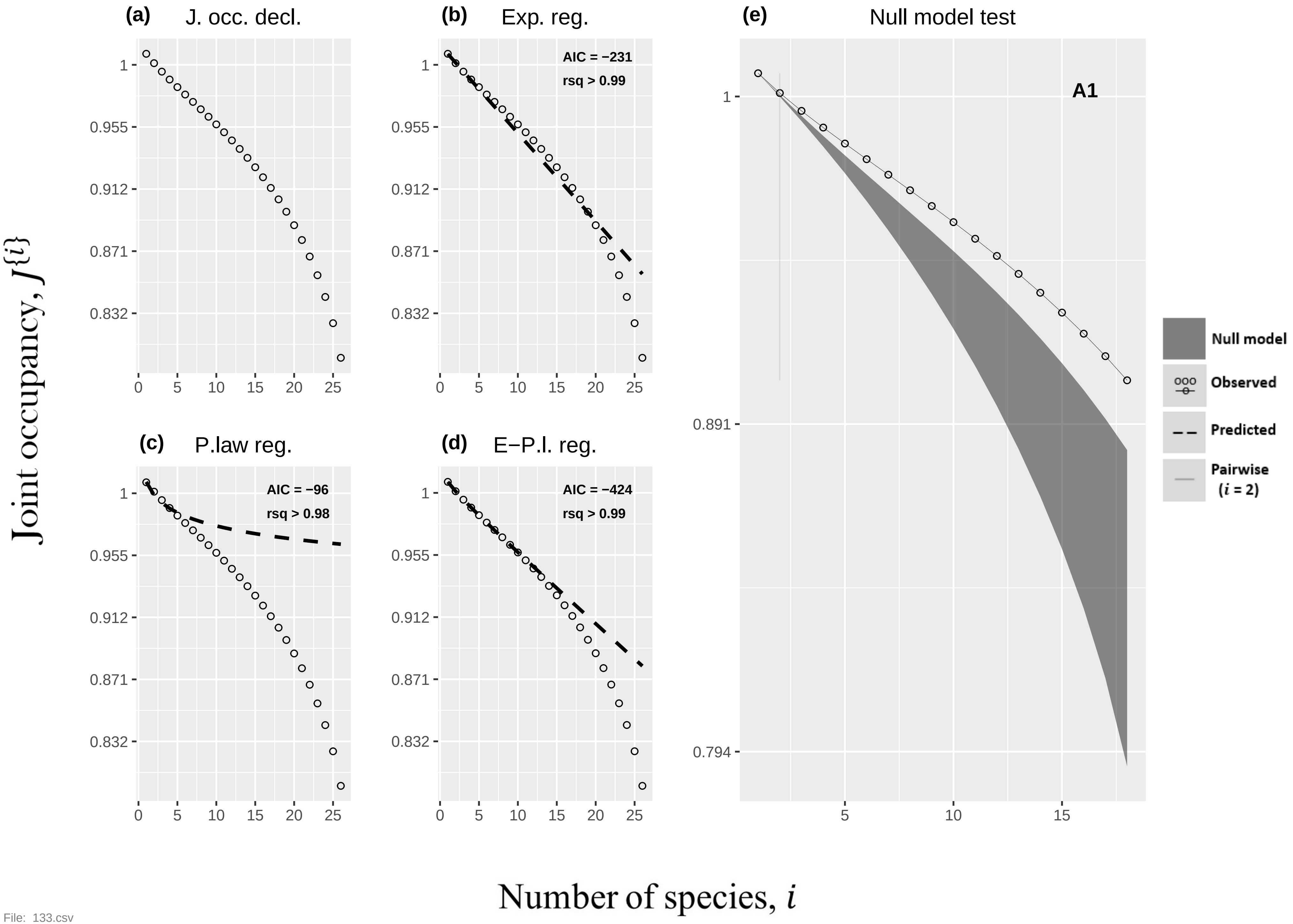
(a)
J. occ. decl.
(b)
Exp. reg.
AIC = −231
rsq > 0.99
(c)
P.law reg.
AIC = −96
rsq > 0.98
(d)
E−P.l. reg.
AIC = −424
rsq > 0.99
(e)
Null model test
A1
Null model
Observed
Predicted
Pairwise
(i = 2)
Joint occupancy, J{i}
Number of species, i

File: 133.csv

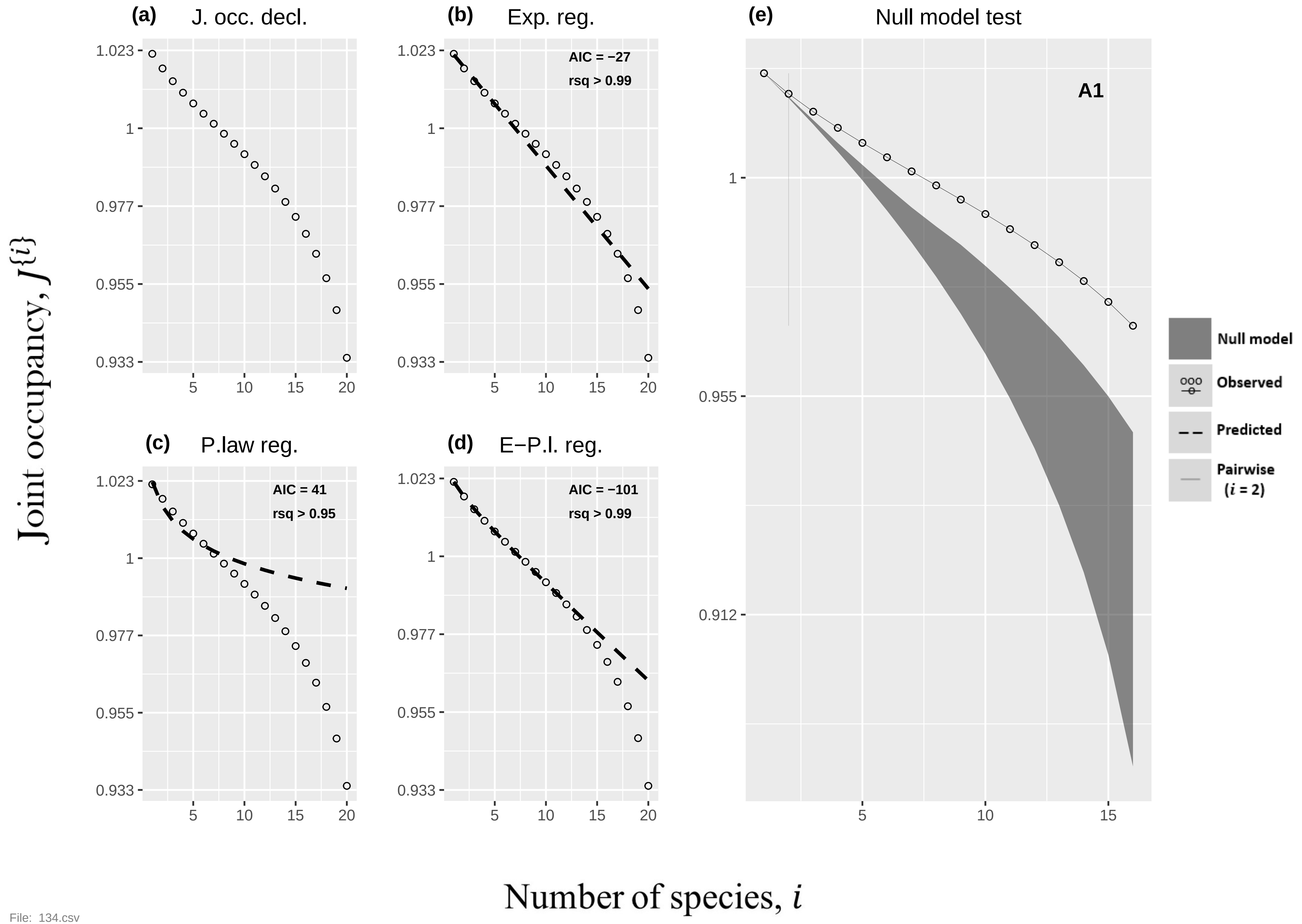
(a) J. occ. decl.
(b) Exp. reg.
AIC = −27
rsq > 0.99
(c) P.law reg.
AIC = 41
rsq > 0.95
(d) E−P.l. reg.
AIC = −101
rsq > 0.99
(e) Null model test
A1
Null model
Observed
Predicted
Pairwise (i = 2)
Joint occupancy, J{i}
Number of species, i
1.023
1
0.977
0.955
0.933
0.912
5
10
15
20

File: 134.csv

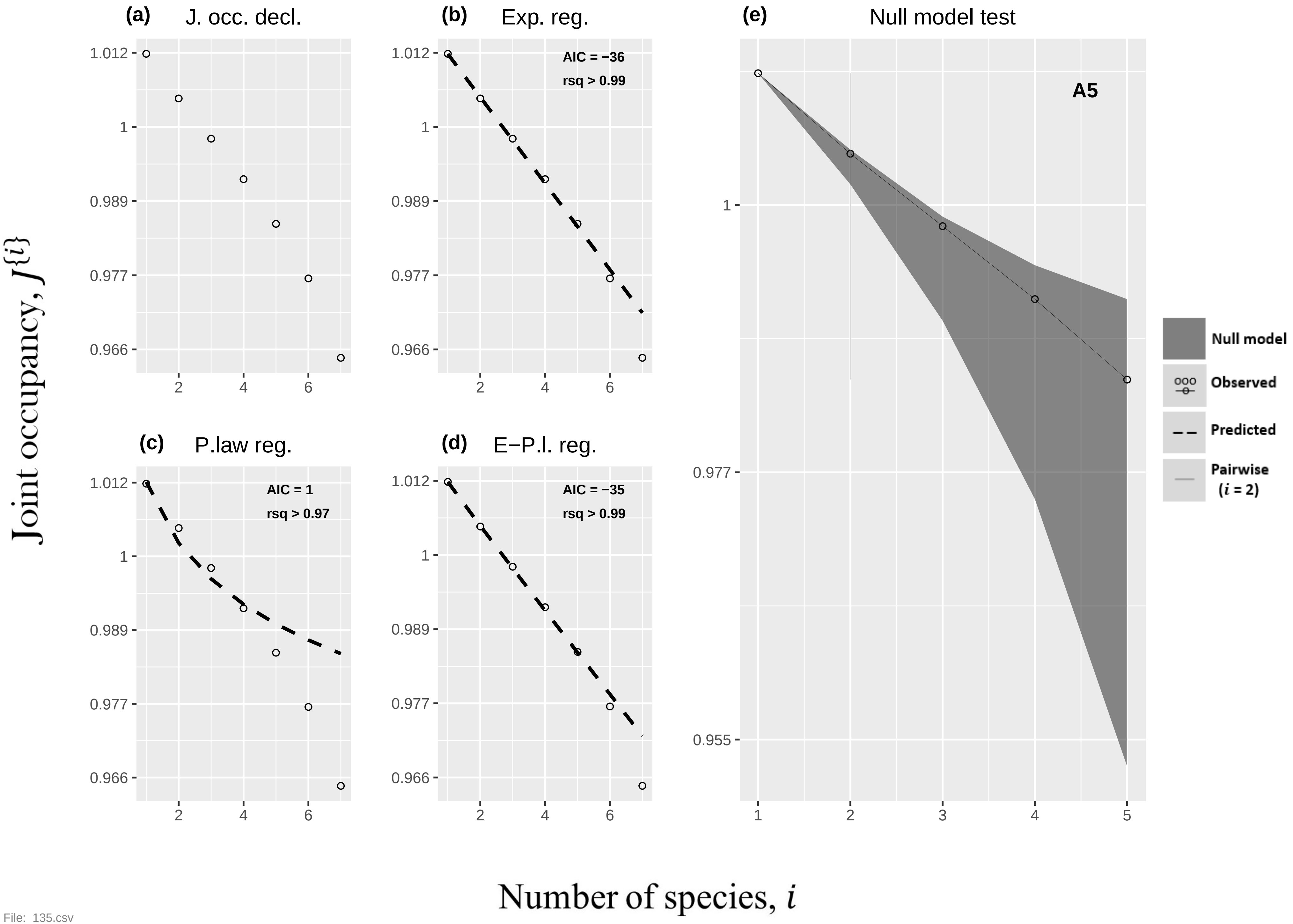
(a) J. occ. decl.
(b) Exp. reg.
AIC = −36
rsq > 0.99
(c) P.law reg.
AIC = 1
rsq > 0.97
(d) E−P.l. reg.
AIC = −35
rsq > 0.99
(e) Null model test
A5
Null model
Observed
Predicted
Pairwise (i = 2)
Joint occupancy, J{i}
Number of species, i
1.012
1
0.989
0.977
0.966
0.955
2
4
6
1
3
5

File: 135.csv

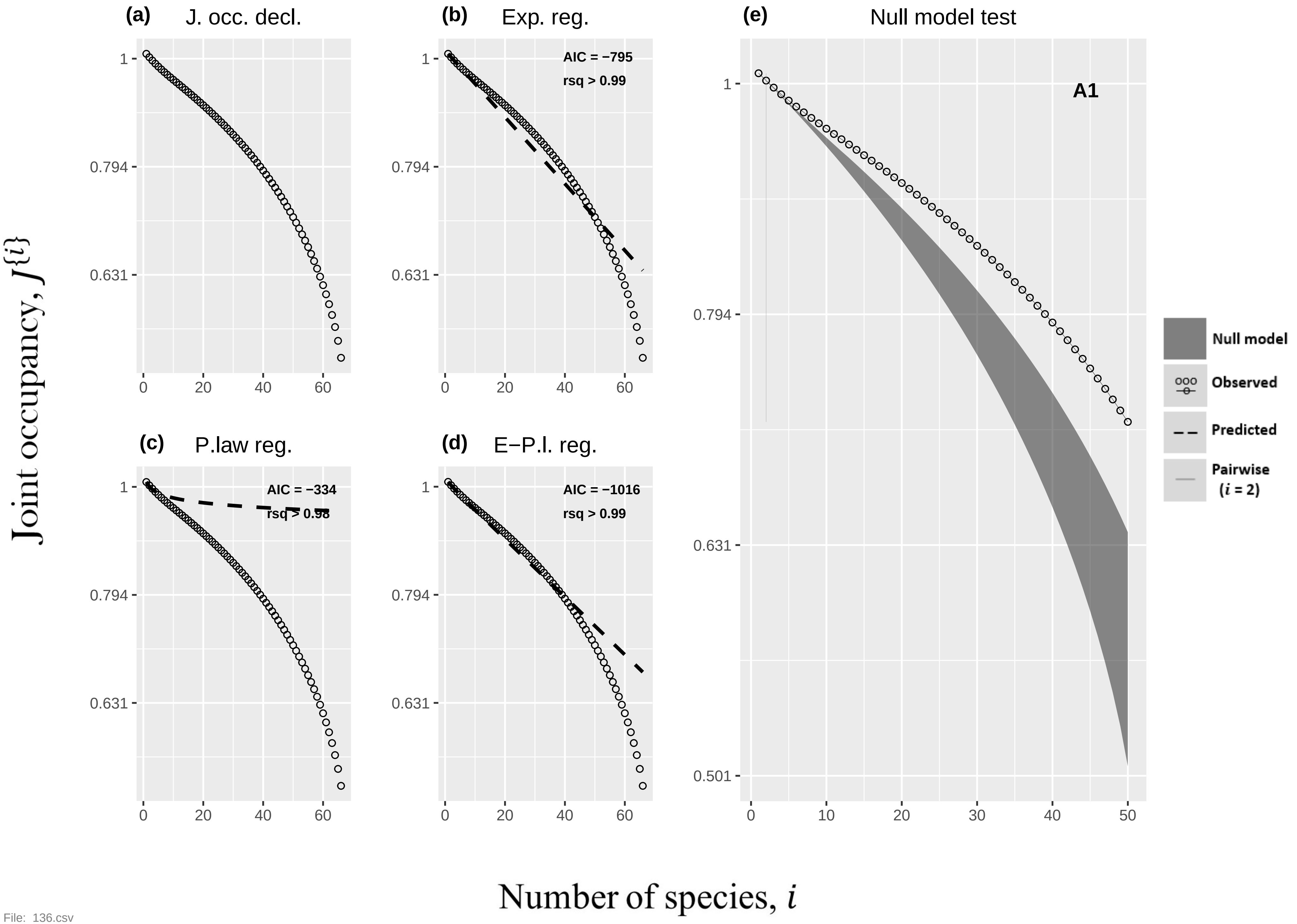

(a) J. occ. decl.
(b) Exp. reg.
AIC = −795
rsq > 0.99
(c) P.law reg.
AIC = −334
rsq > 0.98
(d) E−P.l. reg.
AIC = −1016
rsq > 0.99
(e) Null model test
A1
Null model
Observed
Predicted
Pairwise (i = 2)
Joint occupancy, J{i}
Number of species, i
1
0.794
0.631
0.501
0
20
40
60
10
30
50


File: 136.csv

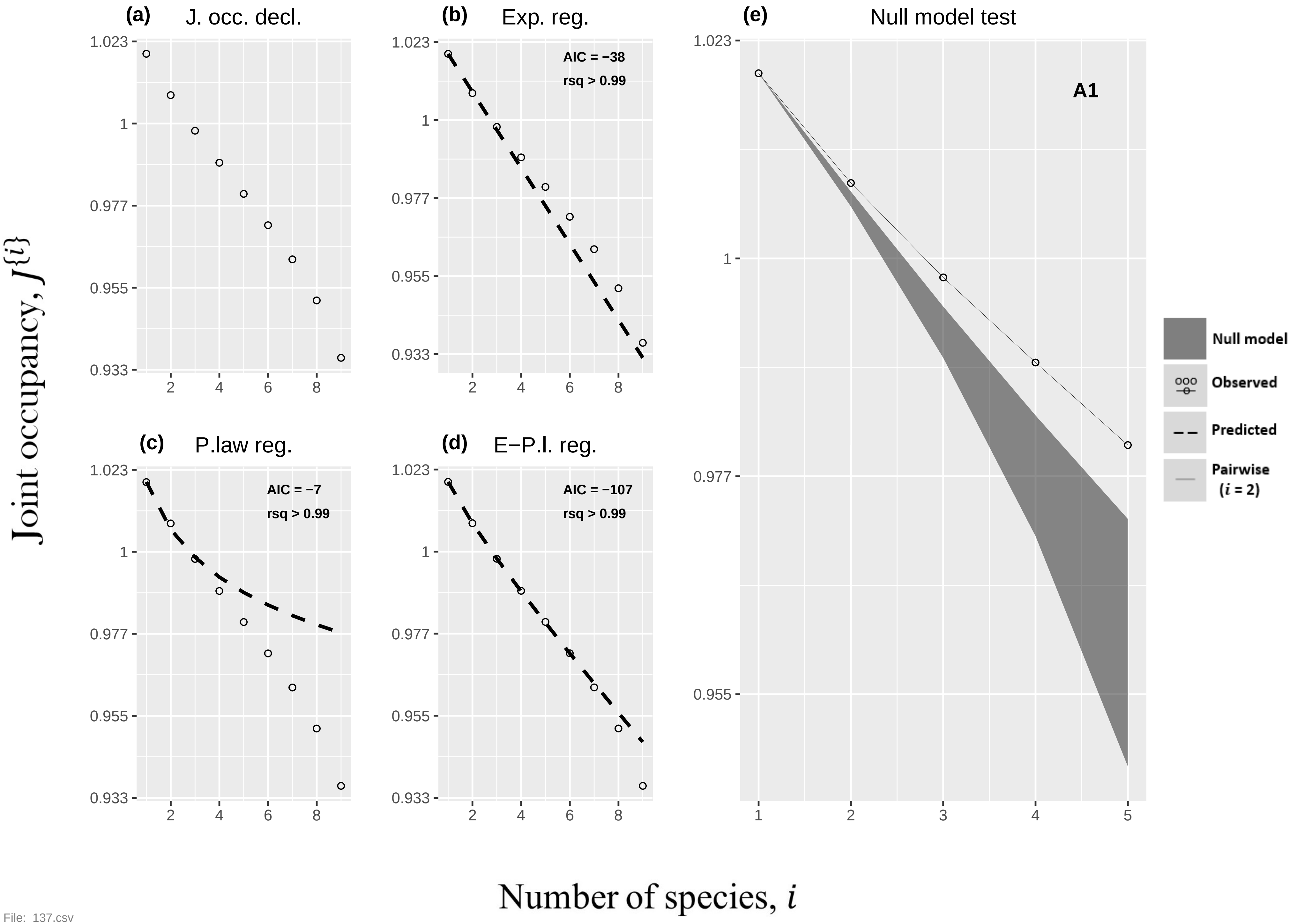

(a) J. occ. decl.
(b) Exp. reg.
AIC = −38
rsq > 0.99
(c) P.law reg.
AIC = −7
rsq > 0.99
(d) E−P.l. reg.
AIC = −107
rsq > 0.99
(e) Null model test
A1
Null model
Observed
Predicted
Pairwise
(i = 2)
Joint occupancy, J{i}
Number of species, i


File: 137.csv

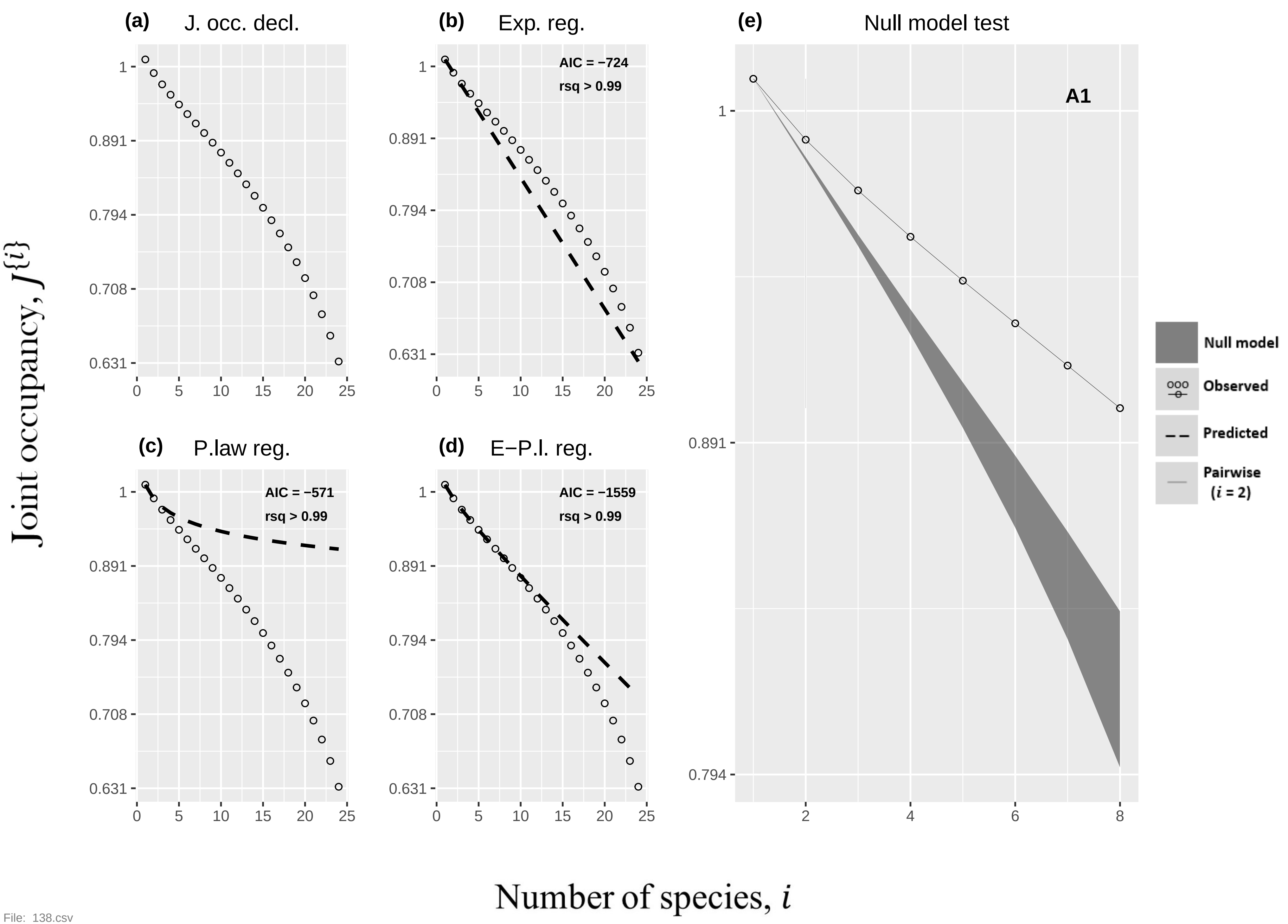
(a) J. occ. decl.
(b) Exp. reg.
AIC = −724
rsq > 0.99
(c) P.law reg.
AIC = −571
rsq > 0.99
(d) E−P.l. reg.
AIC = −1559
rsq > 0.99
(e) Null model test
A1
Null model
Observed
Predicted
Pairwise (i = 2)
1
0.891
0.794
0.708
0.631
0 5 10 15 20 25
2 4 6 8
Joint occupancy, J{i}
Number of species, i

File: 138.csv

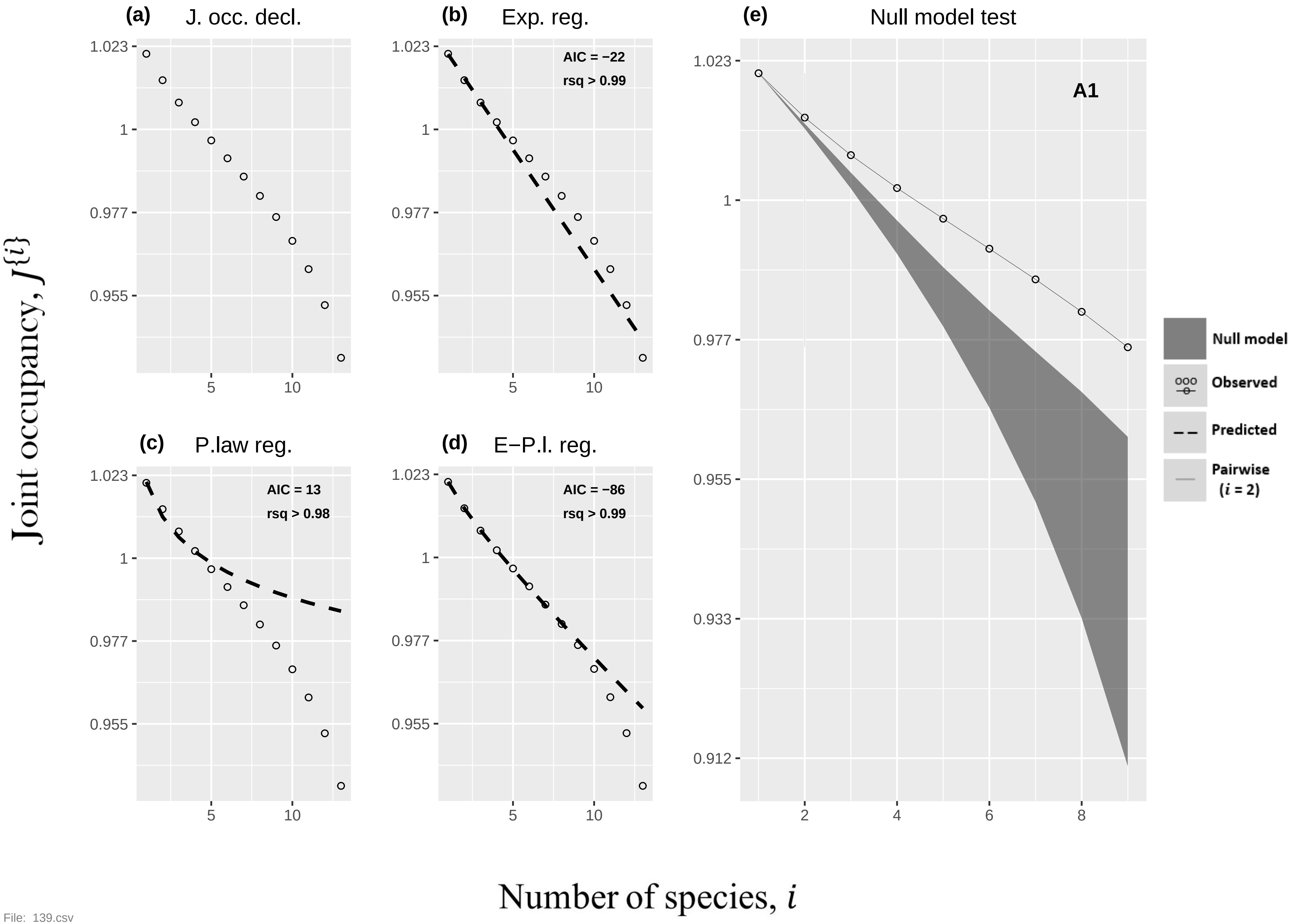

(a) J. occ. decl.
(b) Exp. reg.
AIC = −22
rsq > 0.99
(c) P.law reg.
AIC = 13
rsq > 0.98
(d) E−P.l. reg.
AIC = −86
rsq > 0.99
(e) Null model test
A1
Null model
Observed
Predicted
Pairwise (i = 2)
Joint occupancy, J{i}
Number of species, i


File: 139.csv

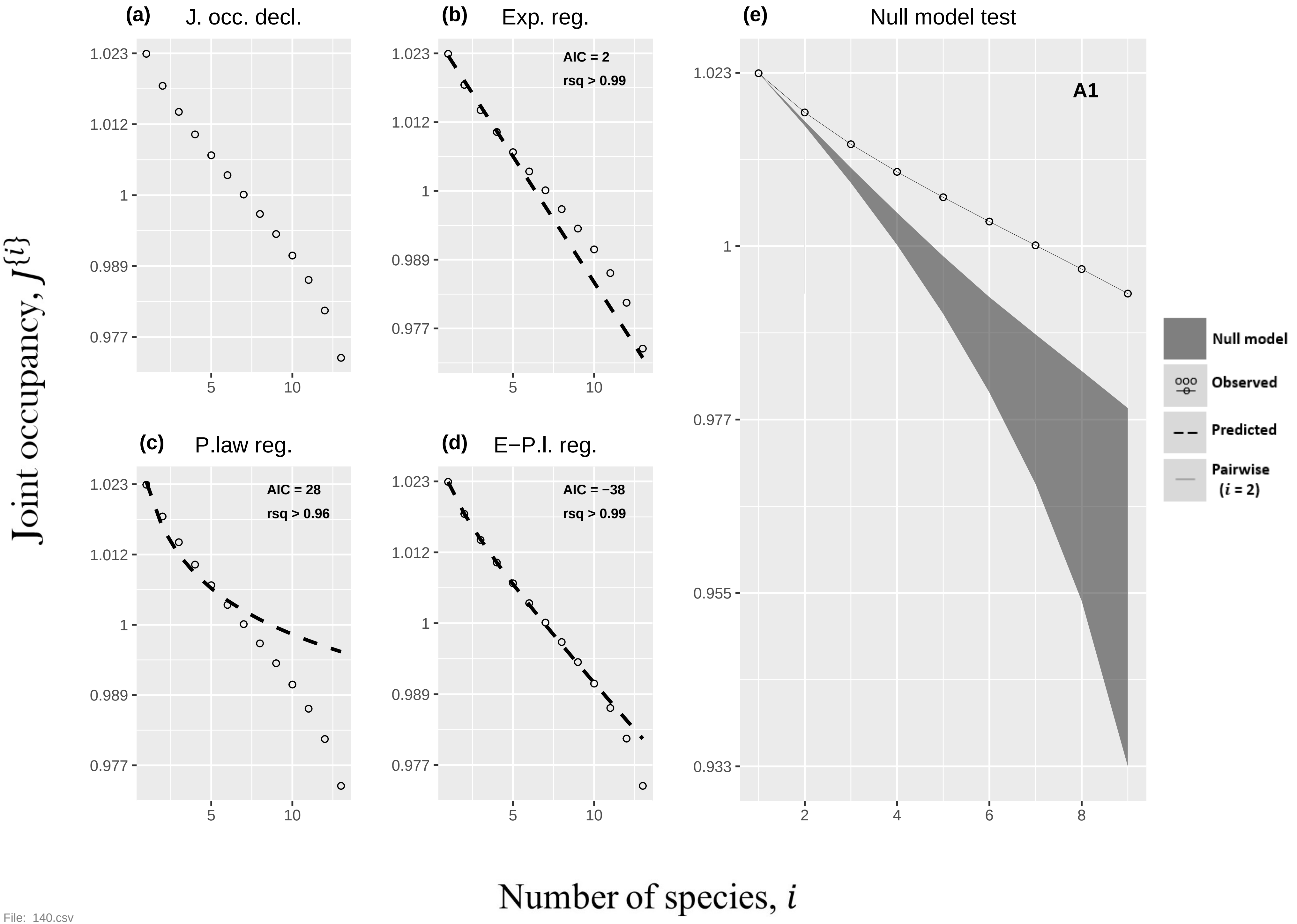
(a) J. occ. decl.
(b) Exp. reg.
AIC = 2
rsq > 0.99
(c) P.law reg.
AIC = 28
rsq > 0.96
(d) E−P.l. reg.
AIC = −38
rsq > 0.99
(e) Null model test
A1
Null model
Observed
Predicted
Pairwise (i = 2)
Joint occupancy, J{i}
Number of species, i
1.023
1.012
1
0.989
0.977
0.955
0.933
5
10
2
4
6
8

File: 140.csv

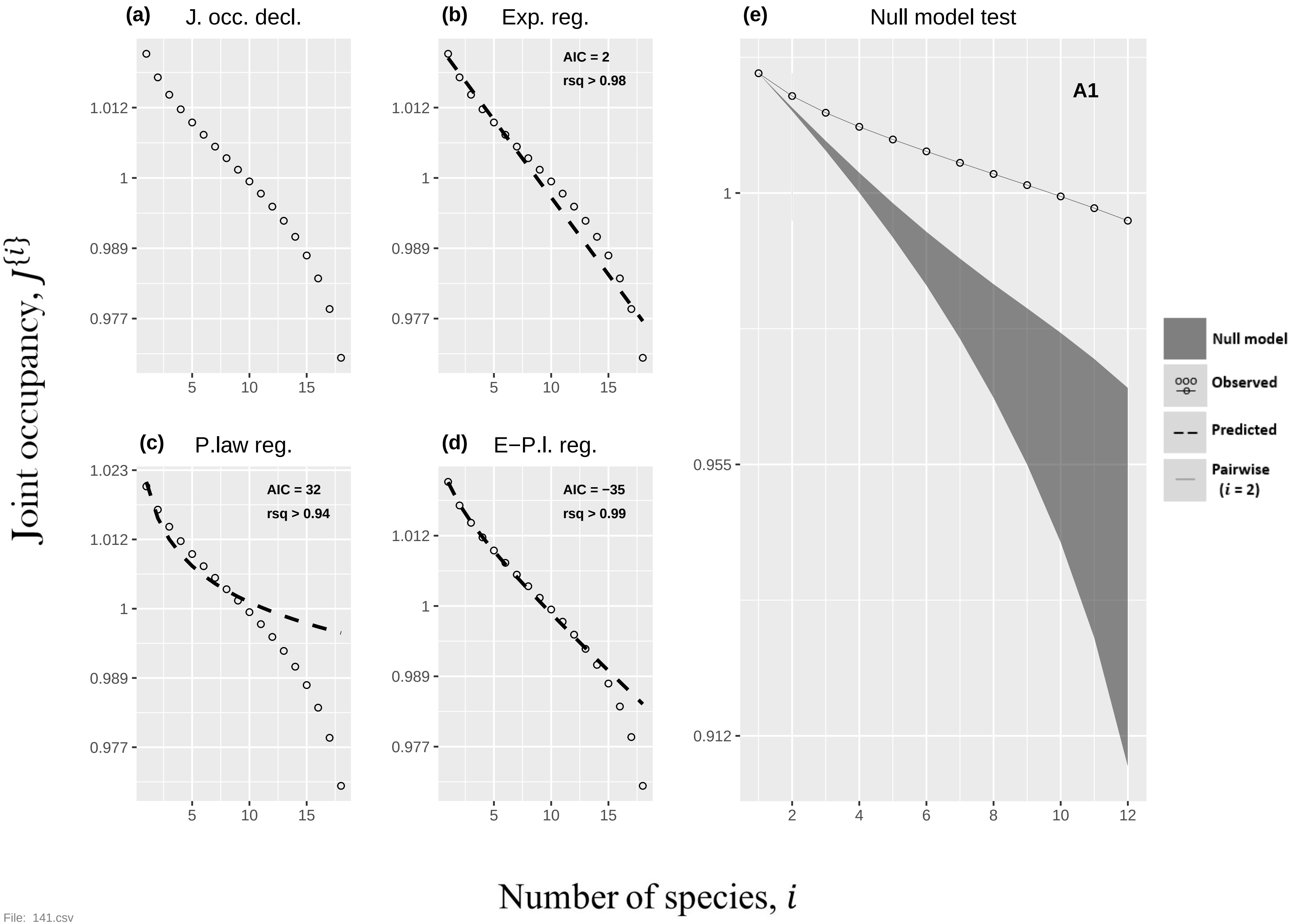
(a) J. occ. decl.
(b) Exp. reg.
AIC = 2
rsq > 0.98
(c) P.law reg.
AIC = 32
rsq > 0.94
(d) E−P.l. reg.
AIC = −35
rsq > 0.99
(e) Null model test
A1
Null model
Observed
Predicted
Pairwise (i = 2)
1.023
1.012
1
0.989
0.977
0.955
0.912
5
10
15
2
4
6
8
10
12
Joint occupancy, J{i}
Number of species, i

File: 141.csv

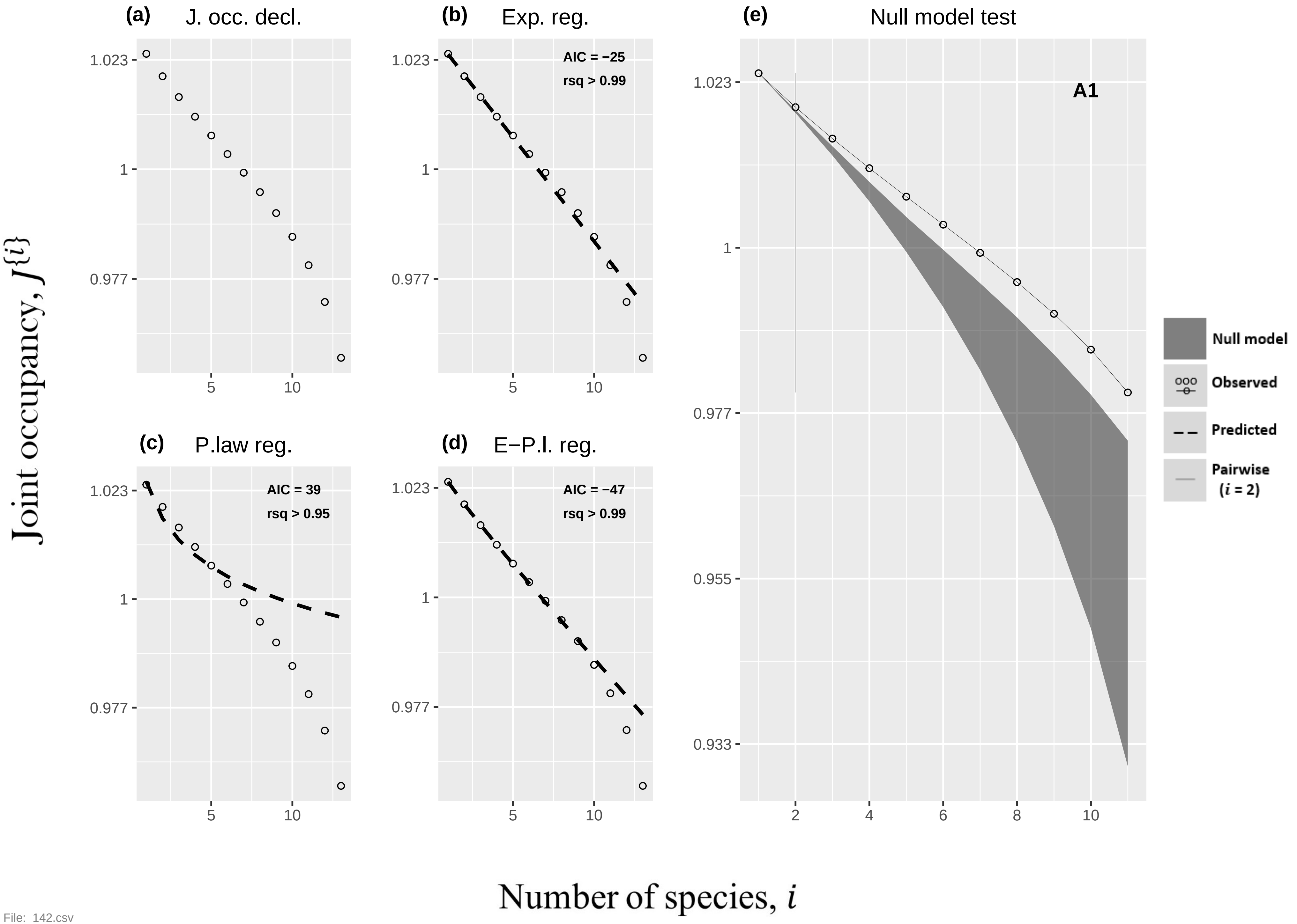

(a) J. occ. decl.
(b) Exp. reg.
AIC = −25
rsq > 0.99
(c) P.law reg.
AIC = 39
rsq > 0.95
(d) E−P.l. reg.
AIC = −47
rsq > 0.99
(e) Null model test
A1
Null model
Observed
Predicted
Pairwise
(i = 2)
Joint occupancy, J{i}
Number of species, i
1.023
1
0.977
0.955
0.933


File: 142.csv

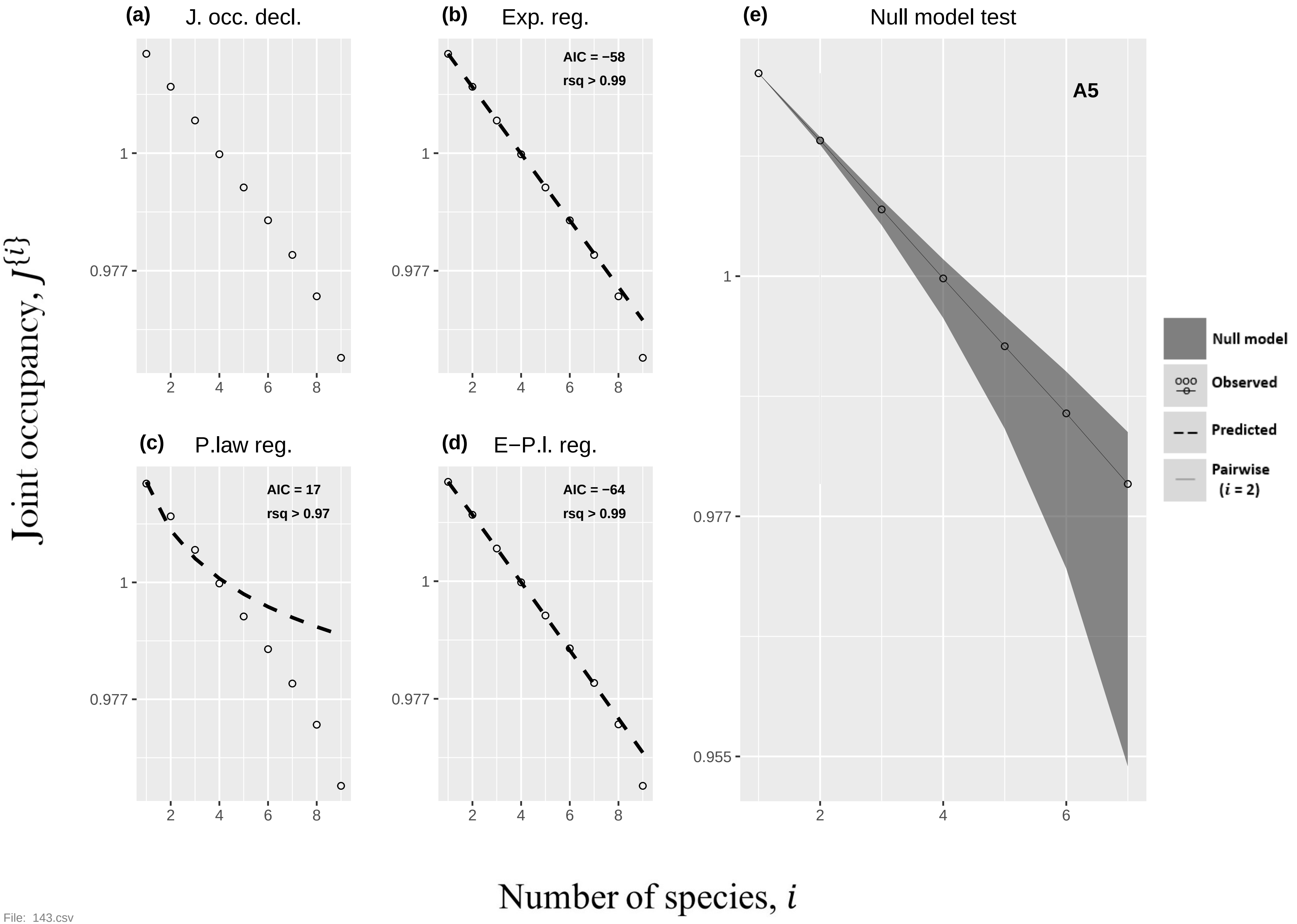

(a) J. occ. decl.
(b) Exp. reg.
AIC = −58
rsq > 0.99
(c) P.law reg.
AIC = 17
rsq > 0.97
(d) E−P.l. reg.
AIC = −64
rsq > 0.99
(e) Null model test
A5
Null model
Observed
Predicted
Pairwise
(i = 2)
1
0.977
0.955
2
4
6
8
Joint occupancy, J{i}
Number of species, i

File: 143.csv

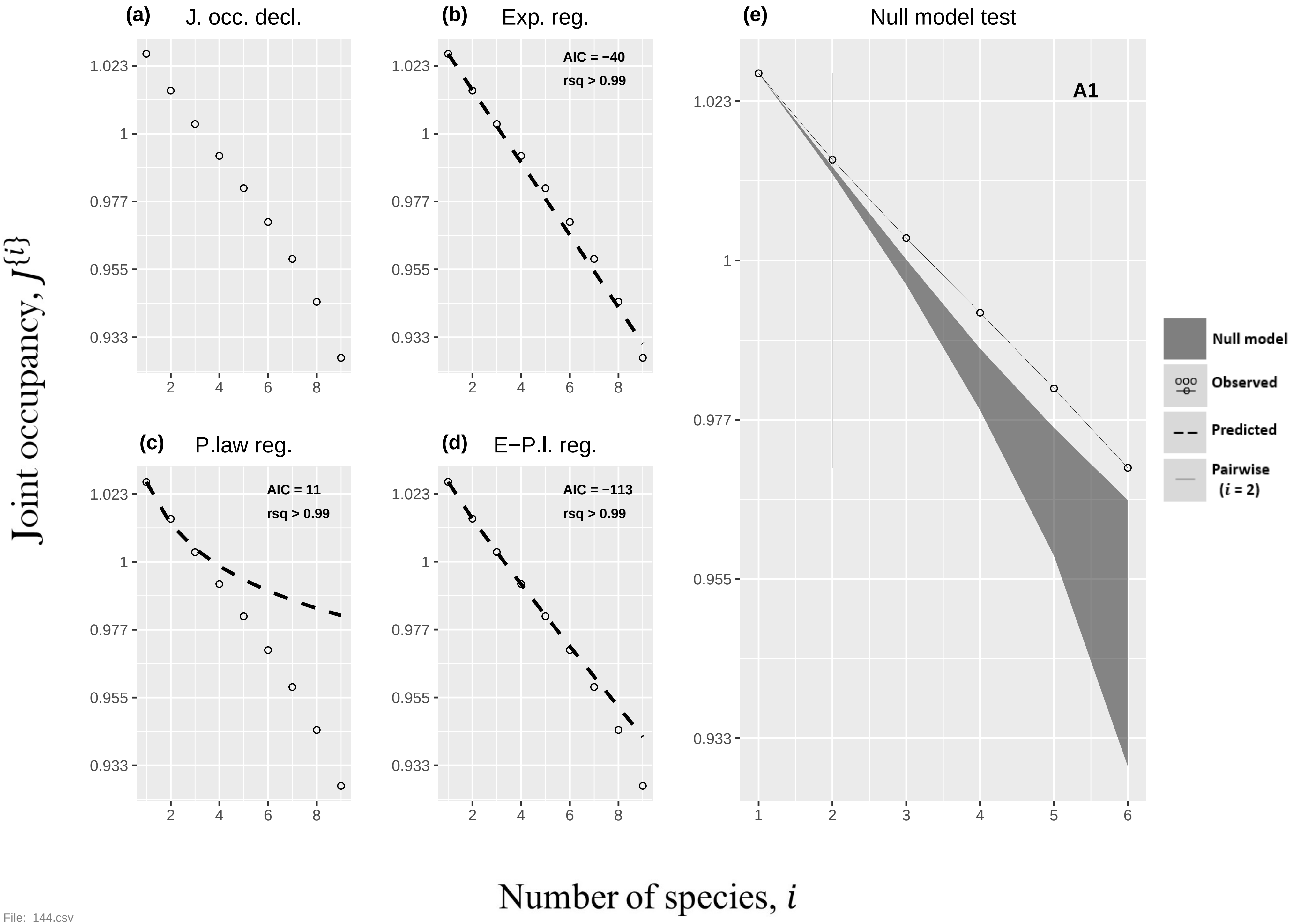

(a) J. occ. decl.
(b) Exp. reg.
AIC = −40
rsq > 0.99
(c) P.law reg.
AIC = 11
rsq > 0.99
(d) E−P.l. reg.
AIC = −113
rsq > 0.99
(e) Null model test
A1
Null model
Observed
Predicted
Pairwise ($i$ = 2)
Joint occupancy, $J^{\{i\}}$
Number of species, $i$


File: 144.csv

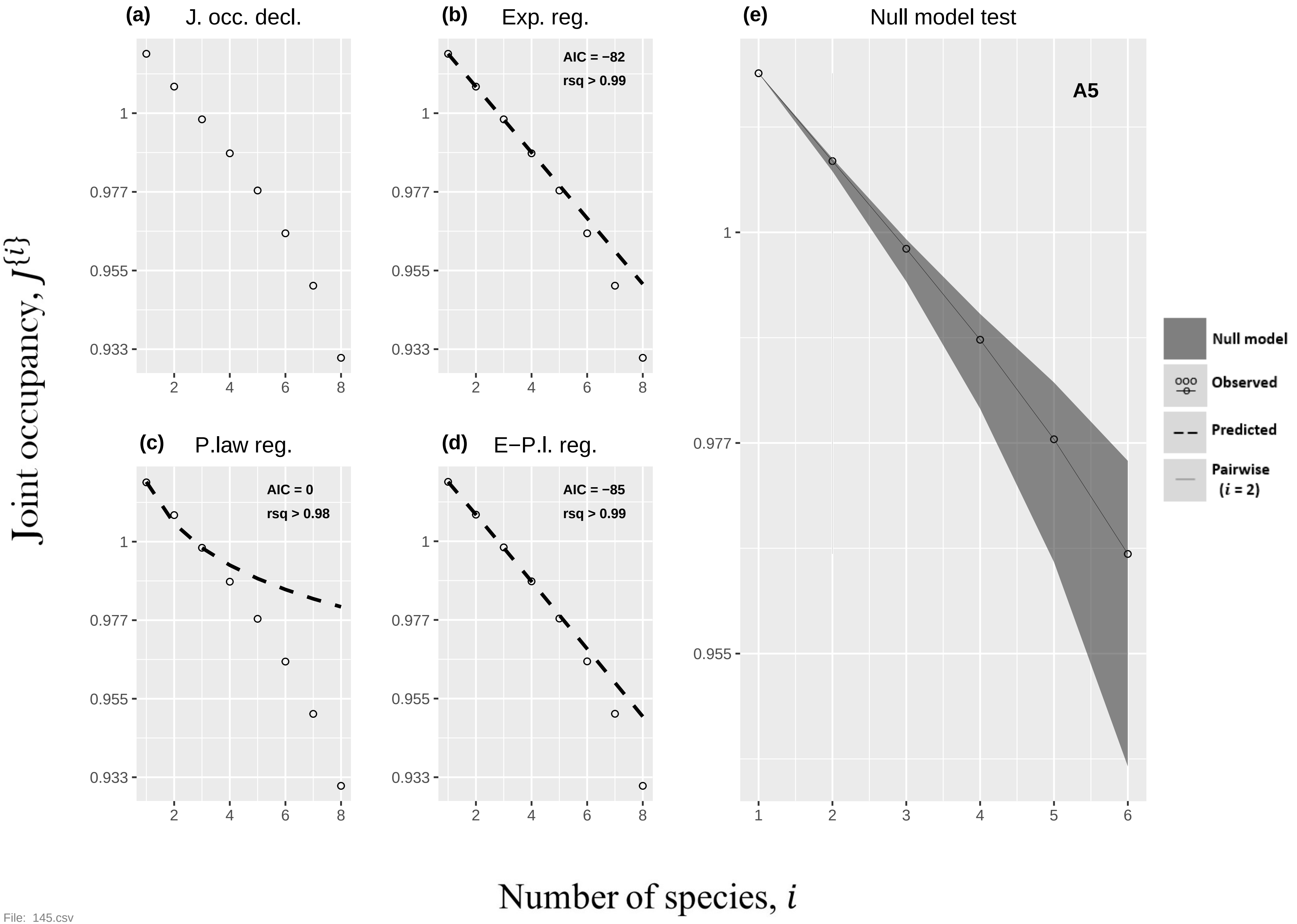

(a) J. occ. decl.
(b) Exp. reg.
AIC = −82
rsq > 0.99
(c) P.law reg.
AIC = 0
rsq > 0.98
(d) E−P.l. reg.
AIC = −85
rsq > 0.99
(e) Null model test
A5
Null model
Observed
Predicted
Pairwise
($i$ = 2)
1
0.977
0.955
0.933
2
4
6
8
1
2
3
4
5
6
Joint occupancy, $J^{\{i\}}$
Number of species, $i$

File: 145.csv

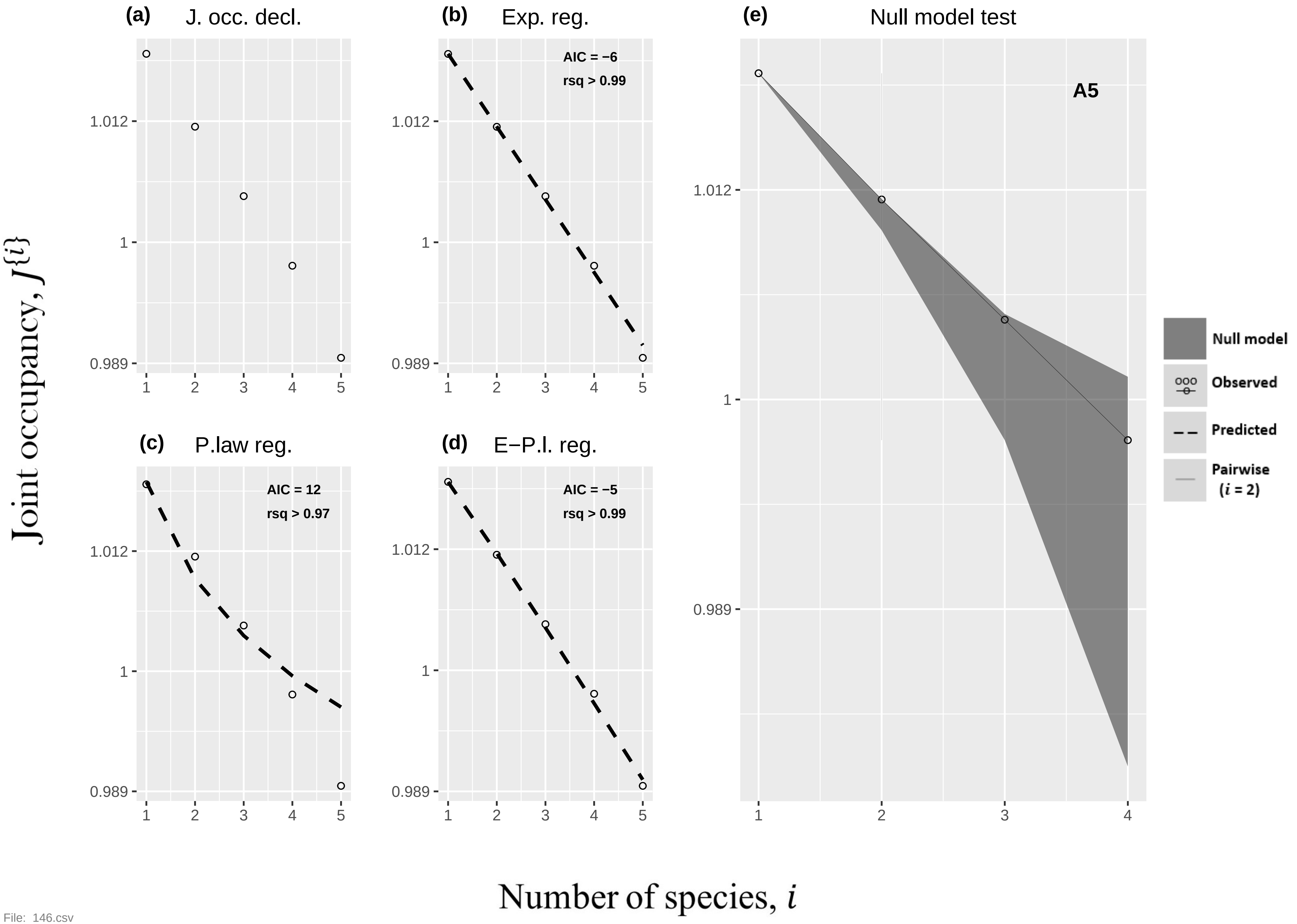
(a)
J. occ. decl.
(b)
Exp. reg.
AIC = −6
rsq > 0.99
(c)
P.law reg.
AIC = 12
rsq > 0.97
(d)
E−P.l. reg.
AIC = −5
rsq > 0.99
(e)
Null model test
A5
Null model
Observed
Predicted
Pairwise
(i = 2)
1.012
1
0.989
1
2
3
4
5
Joint occupancy, J{i}
Number of species, i

File: 146.csv

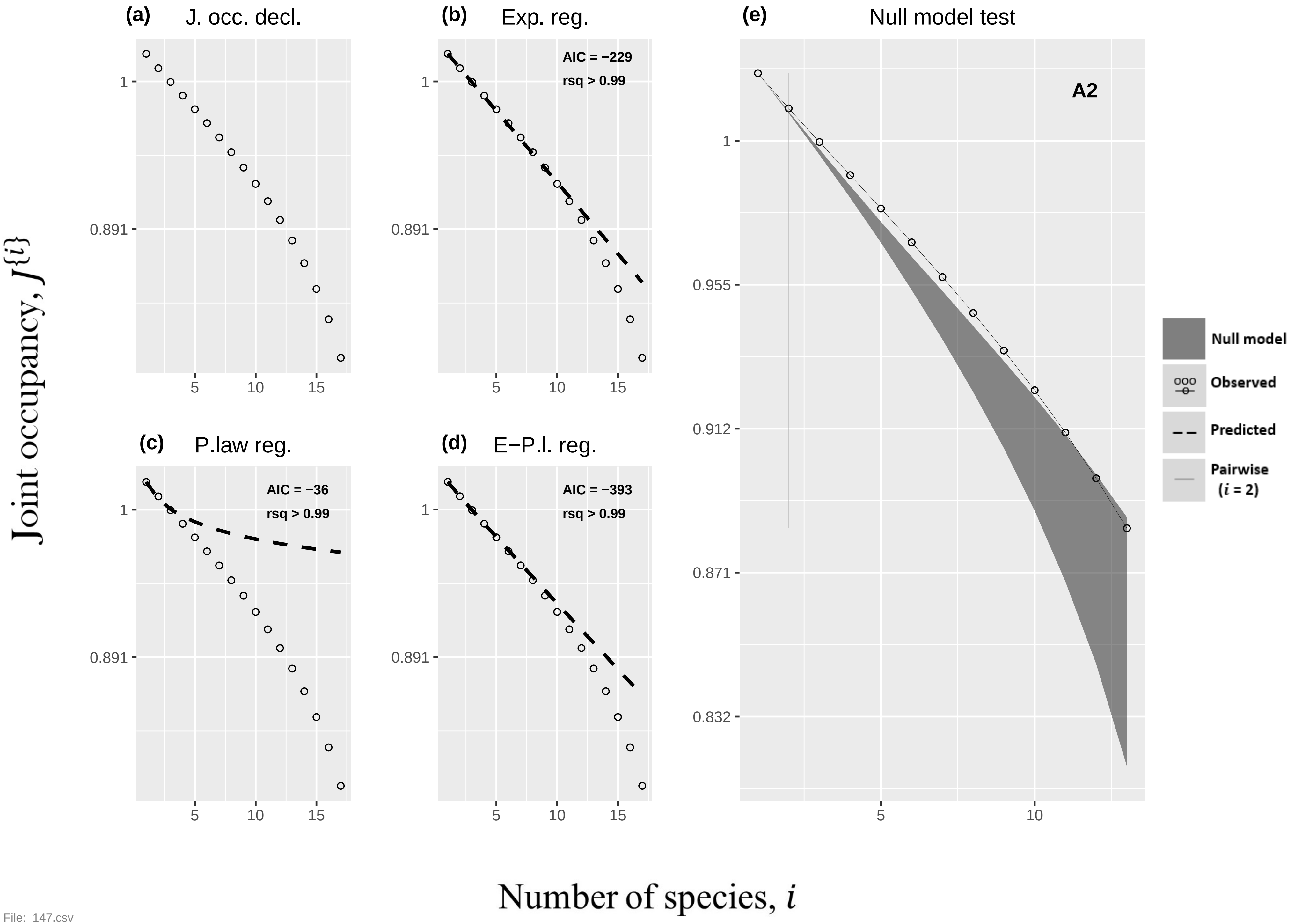

(a) J. occ. decl.
(b) Exp. reg.
AIC = −229
rsq > 0.99
(e) Null model test
A2
(c) P.law reg.
AIC = −36
rsq > 0.99
(d) E−P.l. reg.
AIC = −393
rsq > 0.99
Null model
Observed
Predicted
Pairwise (i = 2)
Joint occupancy, J{i}
Number of species, i
1
0.891
0.955
0.912
0.871
0.832
5
10
15


File: 147.csv

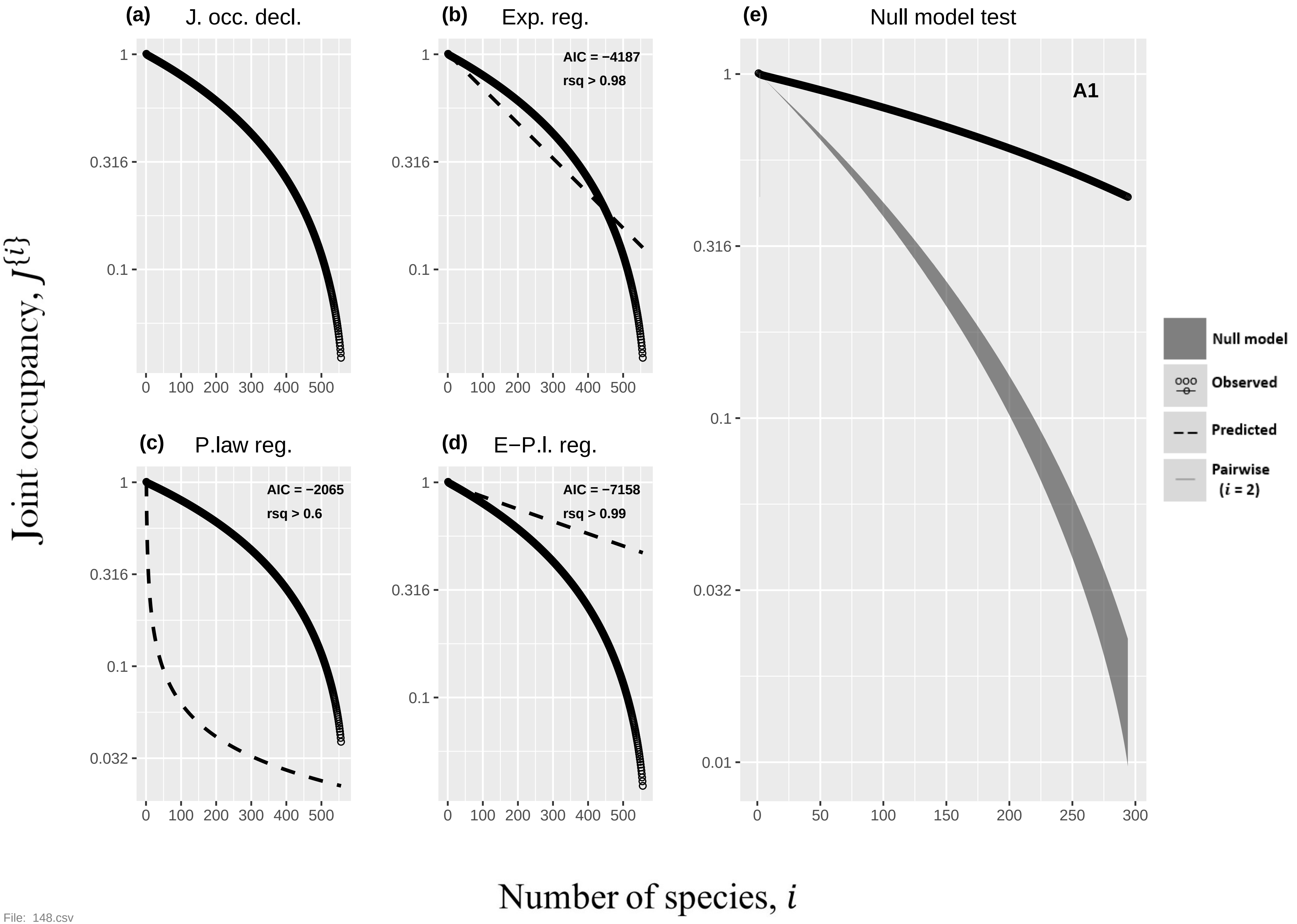

(a) J. occ. decl.
(b) Exp. reg.
AIC = −4187
rsq > 0.98
(c) P.law reg.
AIC = −2065
rsq > 0.6
(d) E−P.l. reg.
AIC = −7158
rsq > 0.99
(e) Null model test
A1
Null model
Observed
Predicted
Pairwise
(i = 2)
Joint occupancy, J{i}
Number of species, i


File: 148.csv

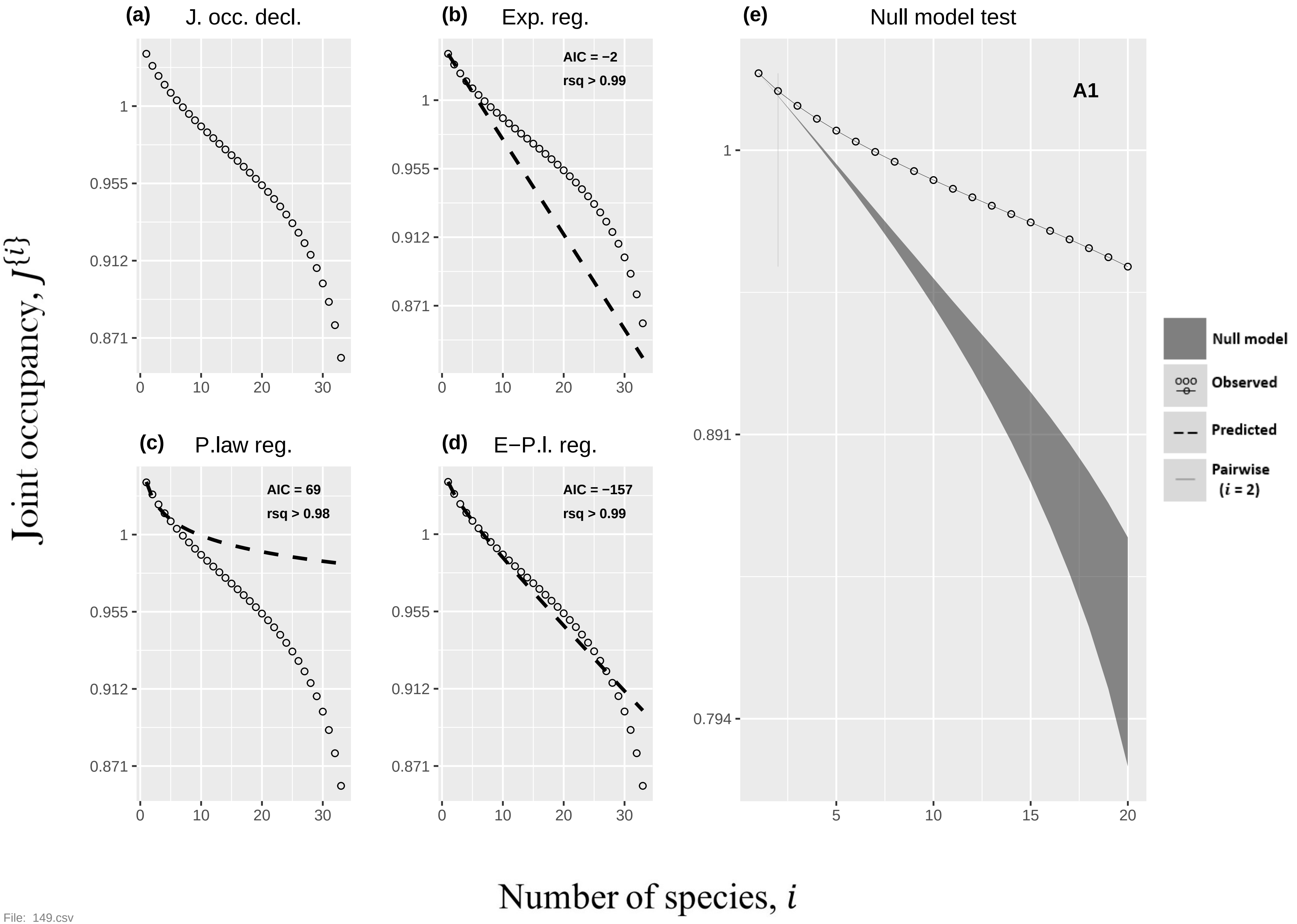

(a) J. occ. decl.
(b) Exp. reg.
AIC = −2
rsq > 0.99
(c) P.law reg.
AIC = 69
rsq > 0.98
(d) E−P.l. reg.
AIC = −157
rsq > 0.99
(e) Null model test
A1
Null model
Observed
Predicted
Pairwise
(i = 2)
1
0.955
0.912
0.871
0.891
0.794
0
10
20
30
5
15
Joint occupancy, J{i}
Number of species, i


File: 149.csv

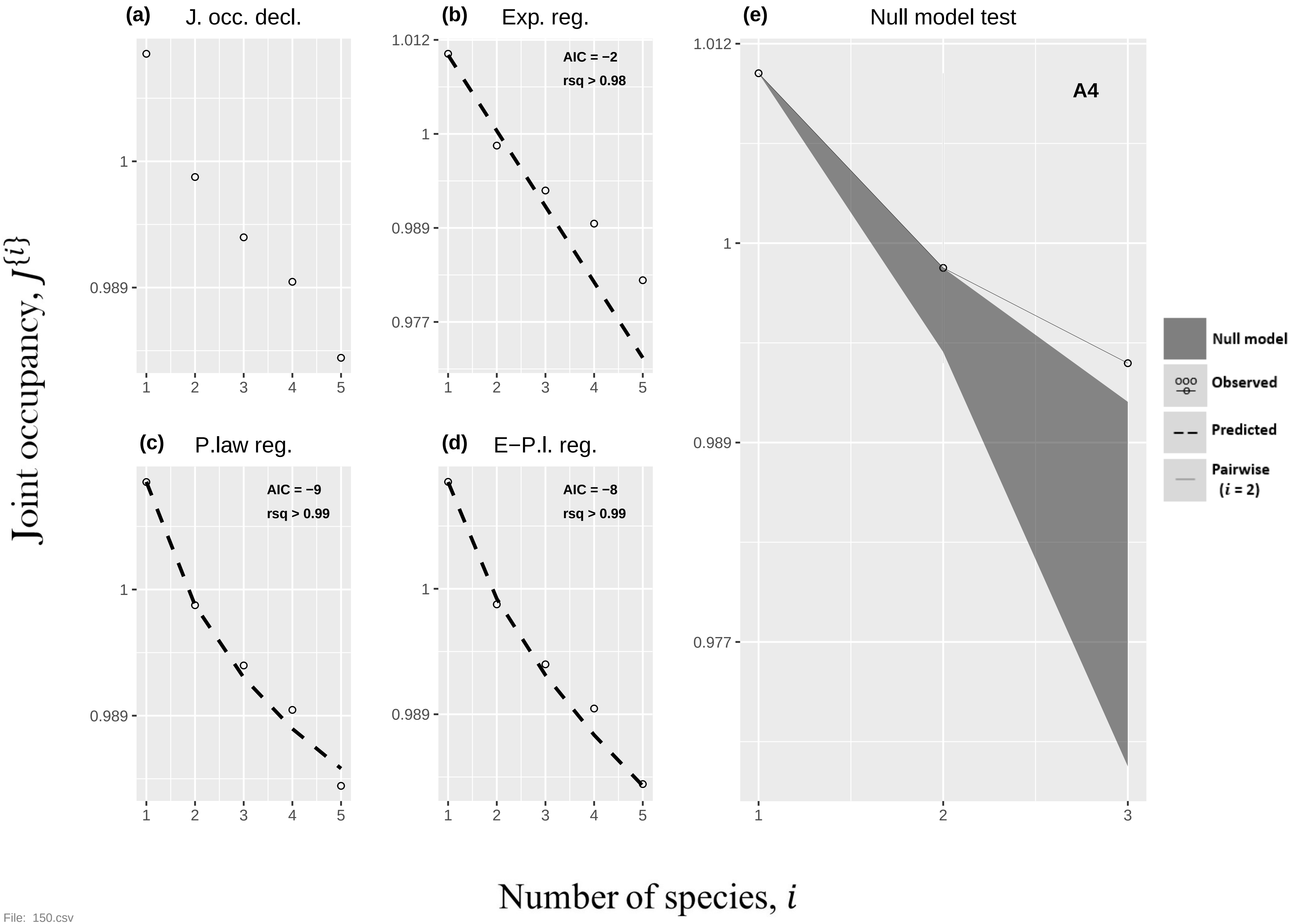
(a) J. occ. decl.
(b) Exp. reg.
AIC = −2
rsq > 0.98
(e) Null model test
A4
(c) P.law reg.
AIC = −9
rsq > 0.99
(d) E−P.l. reg.
AIC = −8
rsq > 0.99
Null model
Observed
Predicted
Pairwise
(i = 2)
1.012
1
0.989
0.977
1
2
3
4
5
Joint occupancy, J{i}
Number of species, i

File: 150.csv

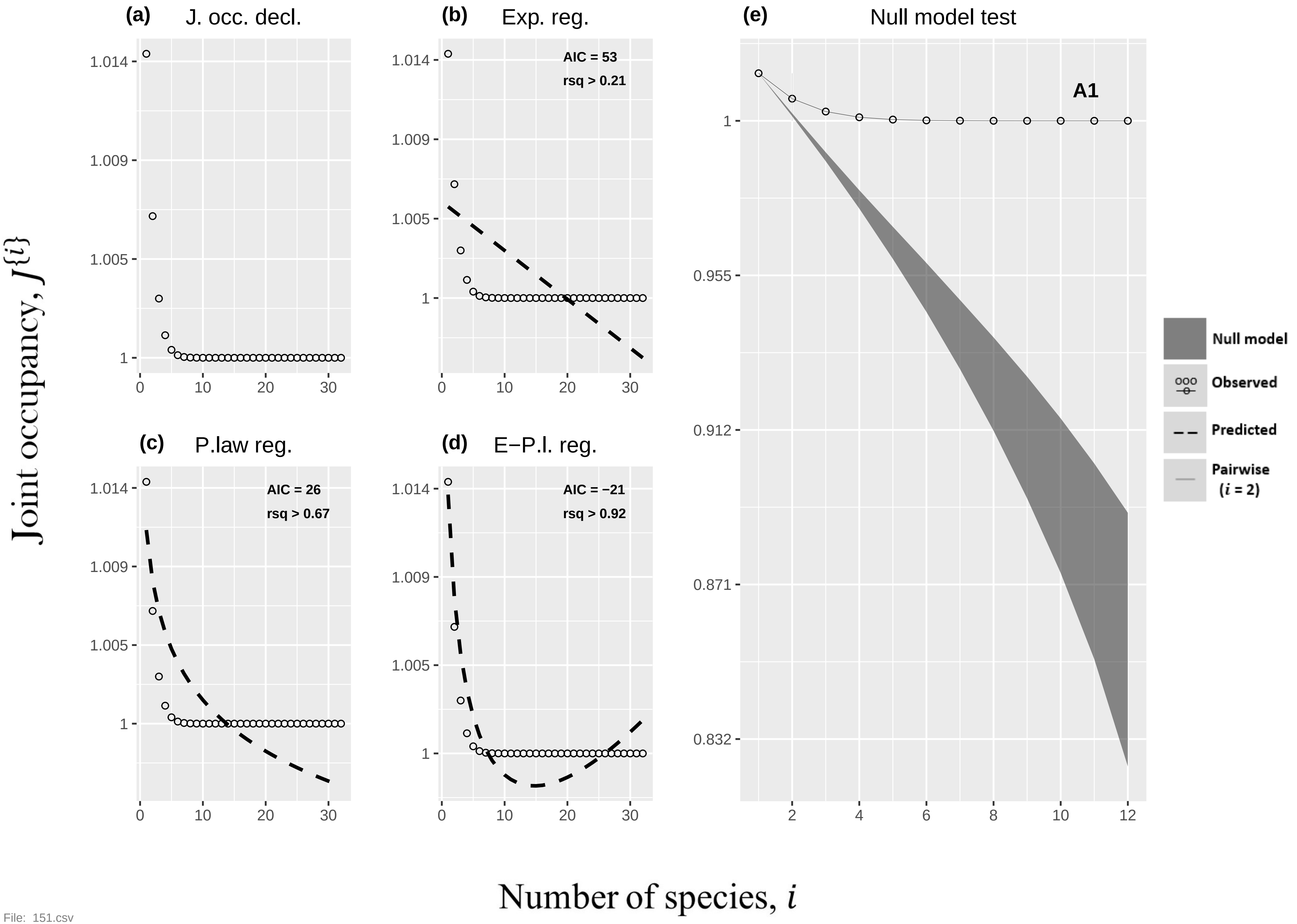
(a)
J. occ. decl.
(b)
Exp. reg.
AIC = 53
rsq > 0.21
(c)
P.law reg.
AIC = 26
rsq > 0.67
(d)
E−P.l. reg.
AIC = −21
rsq > 0.92
(e)
Null model test
A1
Null model
Observed
Predicted
Pairwise (i = 2)
Joint occupancy, J{i}
Number of species, i

File: 151.csv

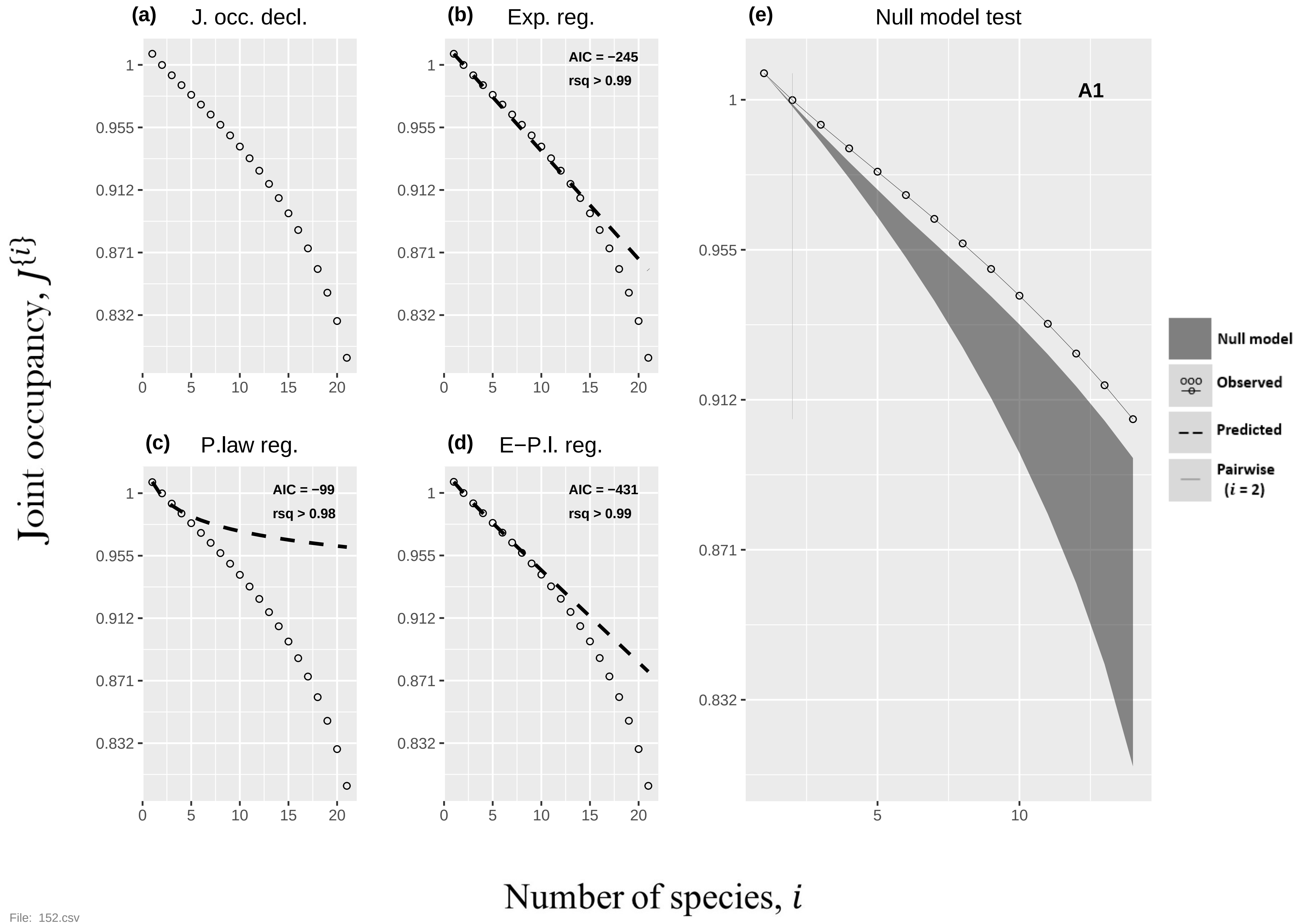

(a) J. occ. decl.
(b) Exp. reg.
AIC = −245
rsq > 0.99
(c) P.law reg.
AIC = −99
rsq > 0.98
(d) E−P.l. reg.
AIC = −431
rsq > 0.99
(e) Null model test
A1
Null model
Observed
Predicted
Pairwise (i = 2)
Joint occupancy, J{i}
Number of species, i
1
0.955
0.912
0.871
0.832
0
5
10
15
20

File: 152.csv

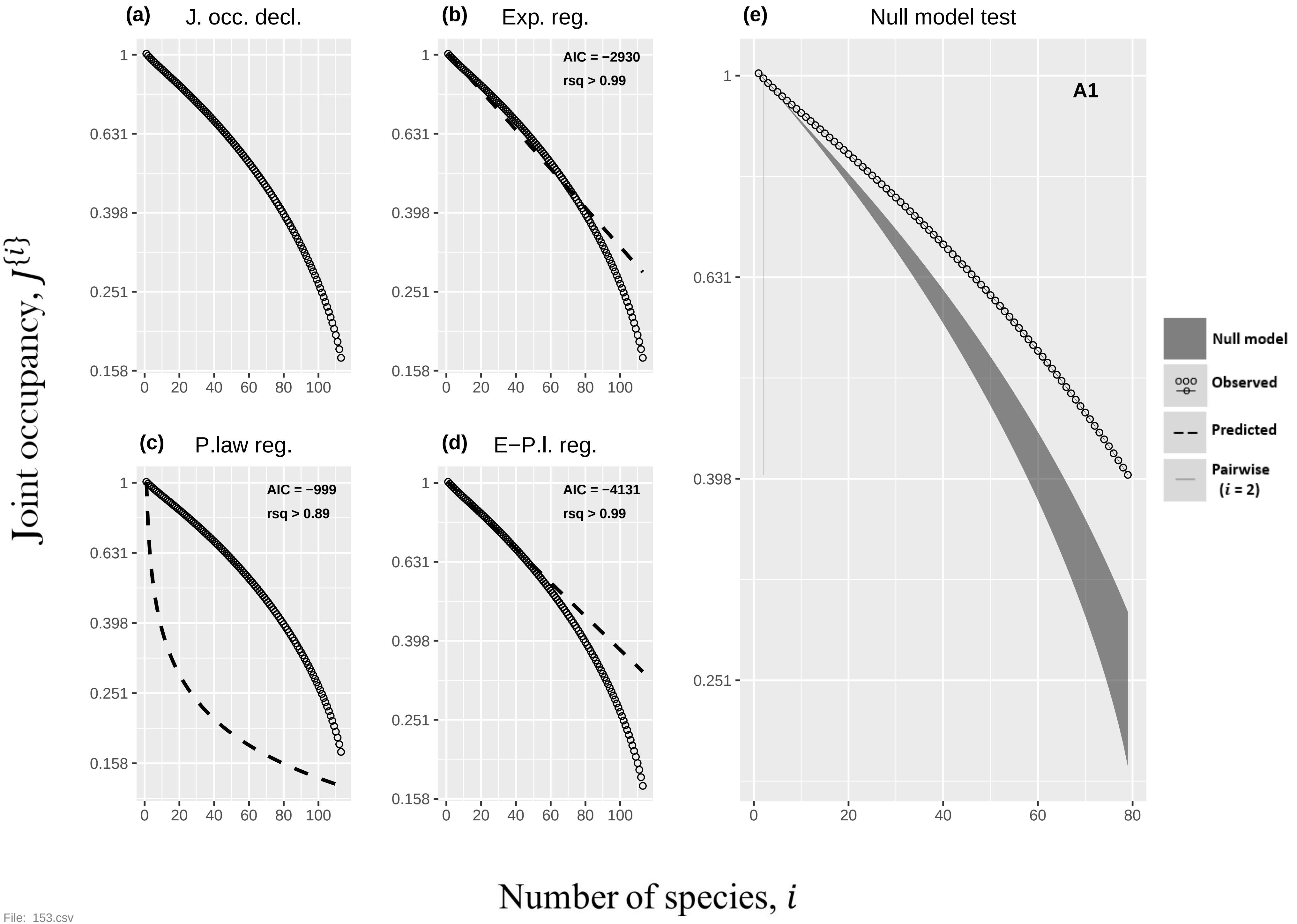

(a)
J. occ. decl.
(b)
Exp. reg.
AIC = −2930
rsq > 0.99
(c)
P.law reg.
AIC = −999
rsq > 0.89
(d)
E−P.l. reg.
AIC = −4131
rsq > 0.99
(e)
Null model test
A1
Null model
Observed
Predicted
Pairwise
(i = 2)
1
0.631
0.398
0.251
0.158
0
20
40
60
80
100
Joint occupancy, J{i}
Number of species, i

File: 153.csv

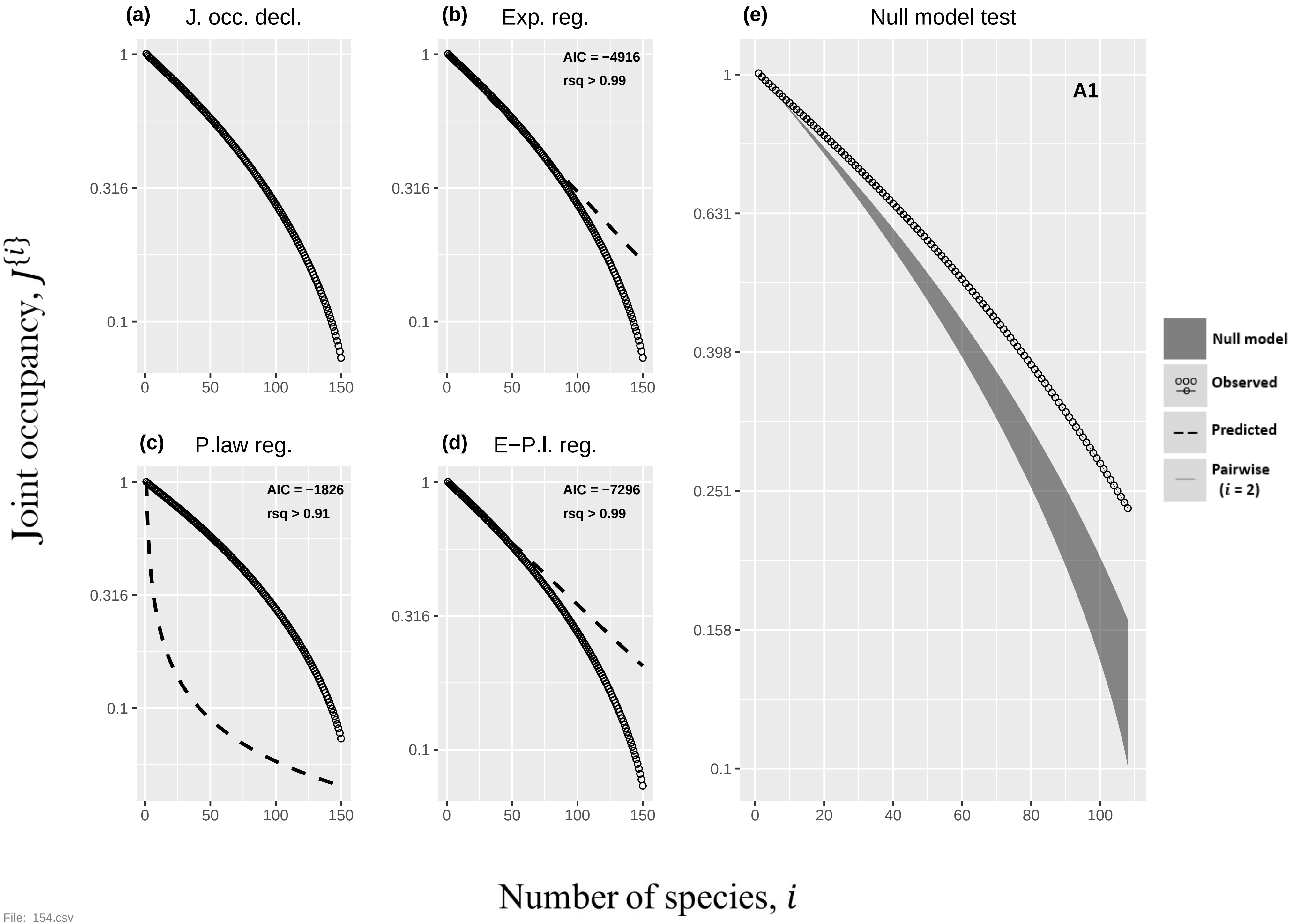

(a) J. occ. decl.
(b) Exp. reg.
AIC = −4916
rsq > 0.99
(c) P.law reg.
AIC = −1826
rsq > 0.91
(d) E−P.l. reg.
AIC = −7296
rsq > 0.99
(e) Null model test
A1
1
0.631
0.398
0.251
0.158
0.1
0.316
0 50 100 150
0 20 40 60 80 100
Null model
Observed
Predicted
Pairwise ($i$ = 2)
Joint occupancy, $J^{\{i\}}$
Number of species, $i$


File: 154.csv

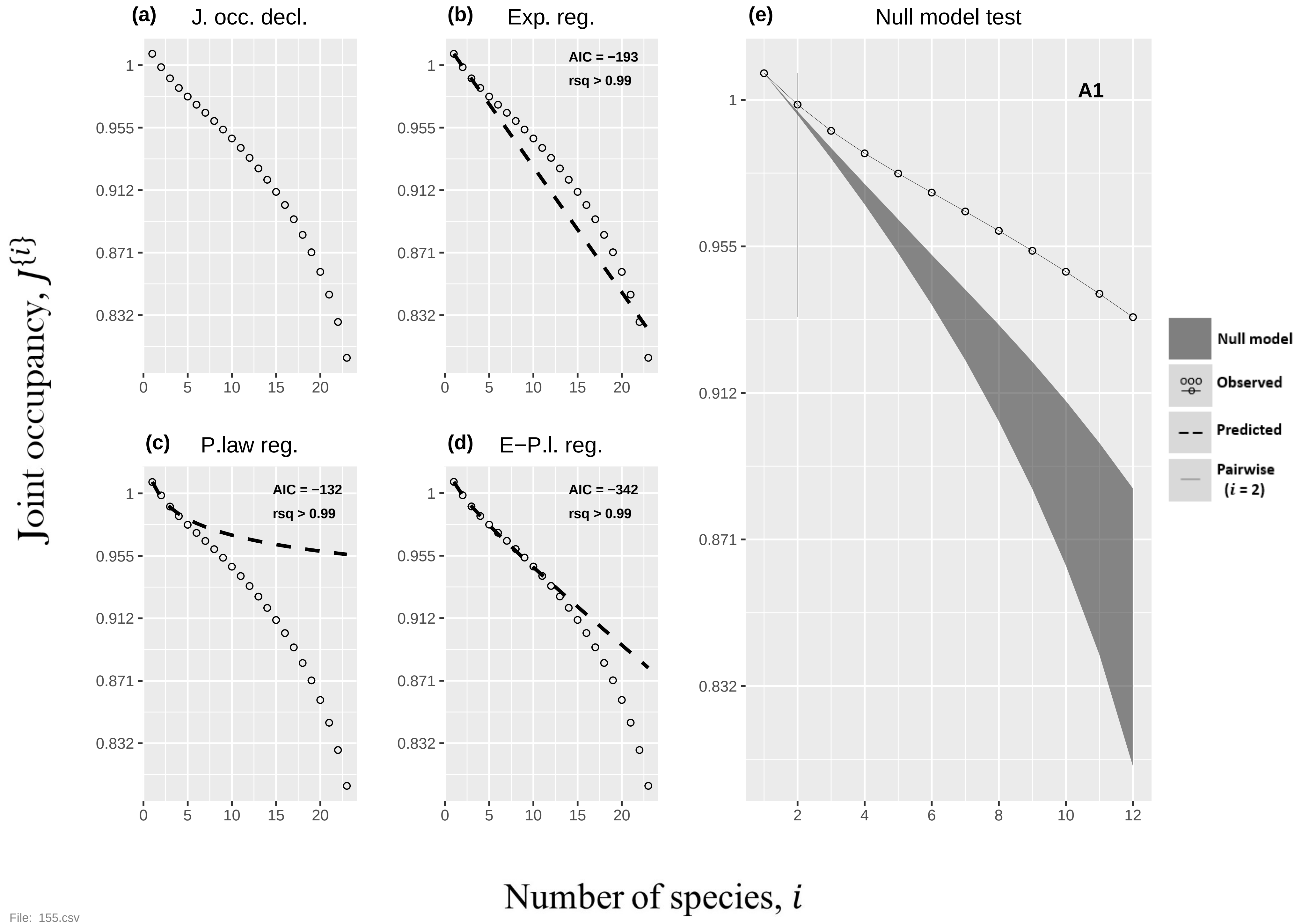
(a) J. occ. decl.
(b) Exp. reg.
AIC = −193
rsq > 0.99
(c) P.law reg.
AIC = −132
rsq > 0.99
(d) E−P.l. reg.
AIC = −342
rsq > 0.99
(e) Null model test
A1
Null model
Observed
Predicted
Pairwise (i = 2)
Joint occupancy, J{i}
Number of species, i
1
0.955
0.912
0.871
0.832
0
5
10
15
20
2
4
6
8
10
12

File: 155.csv

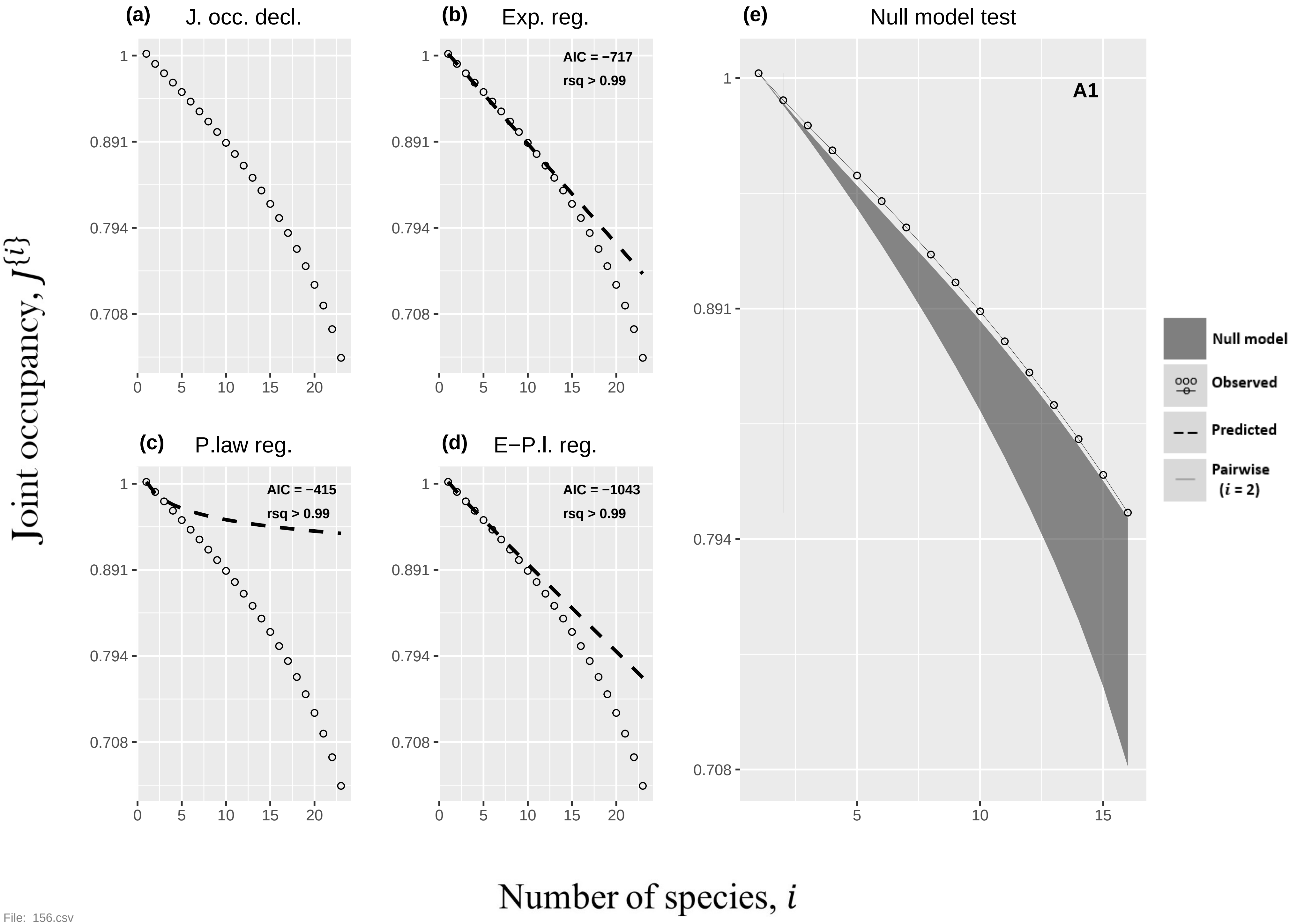

(a) J. occ. decl.
(b) Exp. reg.
AIC = −717
rsq > 0.99
(c) P.law reg.
AIC = −415
rsq > 0.99
(d) E−P.l. reg.
AIC = −1043
rsq > 0.99
(e) Null model test
A1
Null model
Observed
Predicted
Pairwise ($i$ = 2)
Joint occupancy, $J^{\{i\}}$
Number of species, $i$


File: 156.csv

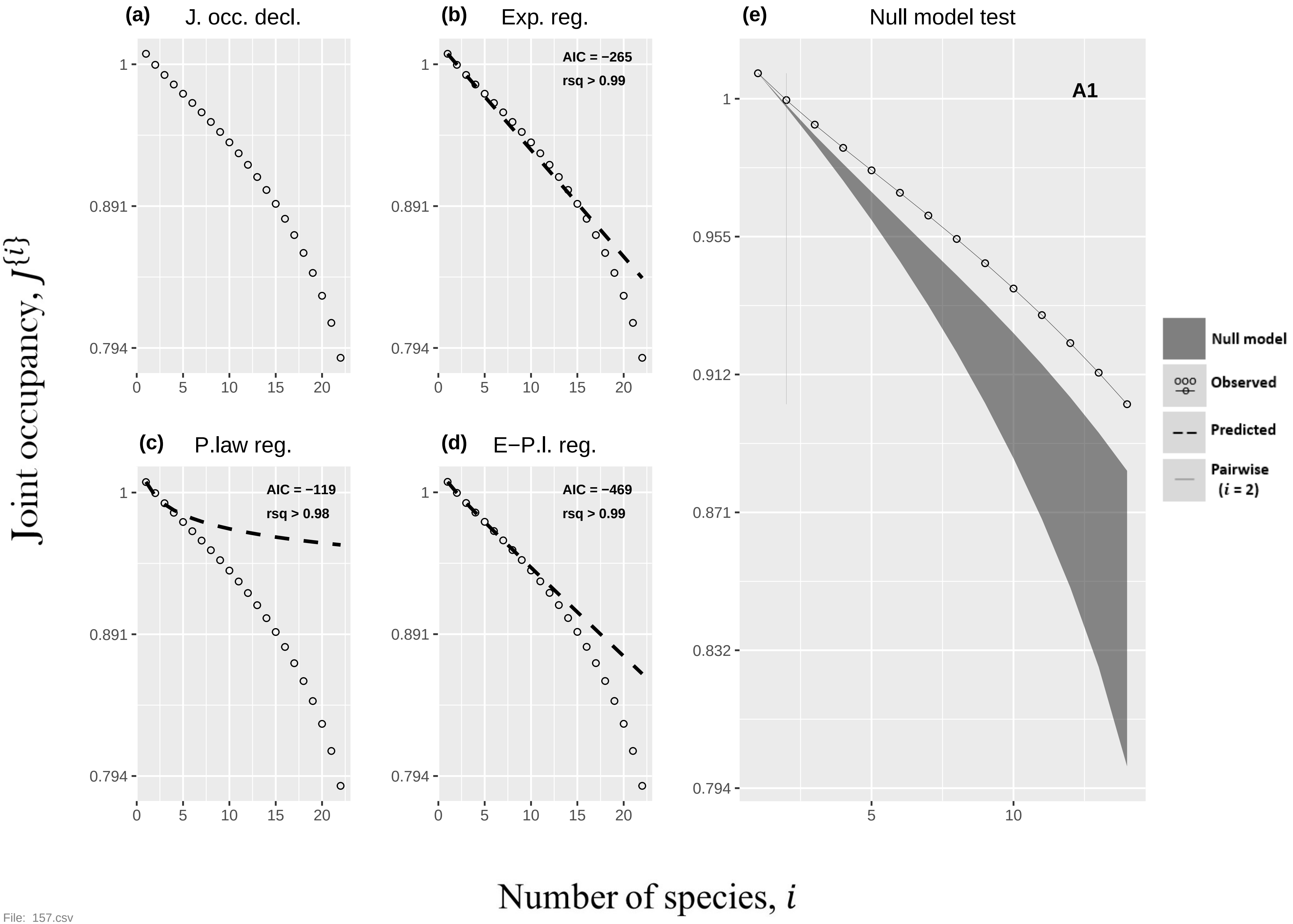

(a) J. occ. decl.
(b) Exp. reg.
AIC = −265
rsq > 0.99
(c) P.law reg.
AIC = −119
rsq > 0.98
(d) E−P.l. reg.
AIC = −469
rsq > 0.99
(e) Null model test
A1
Null model
Observed
Predicted
Pairwise
(i = 2)
Joint occupancy, J{i}
Number of species, i
1
0.891
0.794
0.955
0.912
0.871
0.832
0
5
10
15
20


File: 157.csv

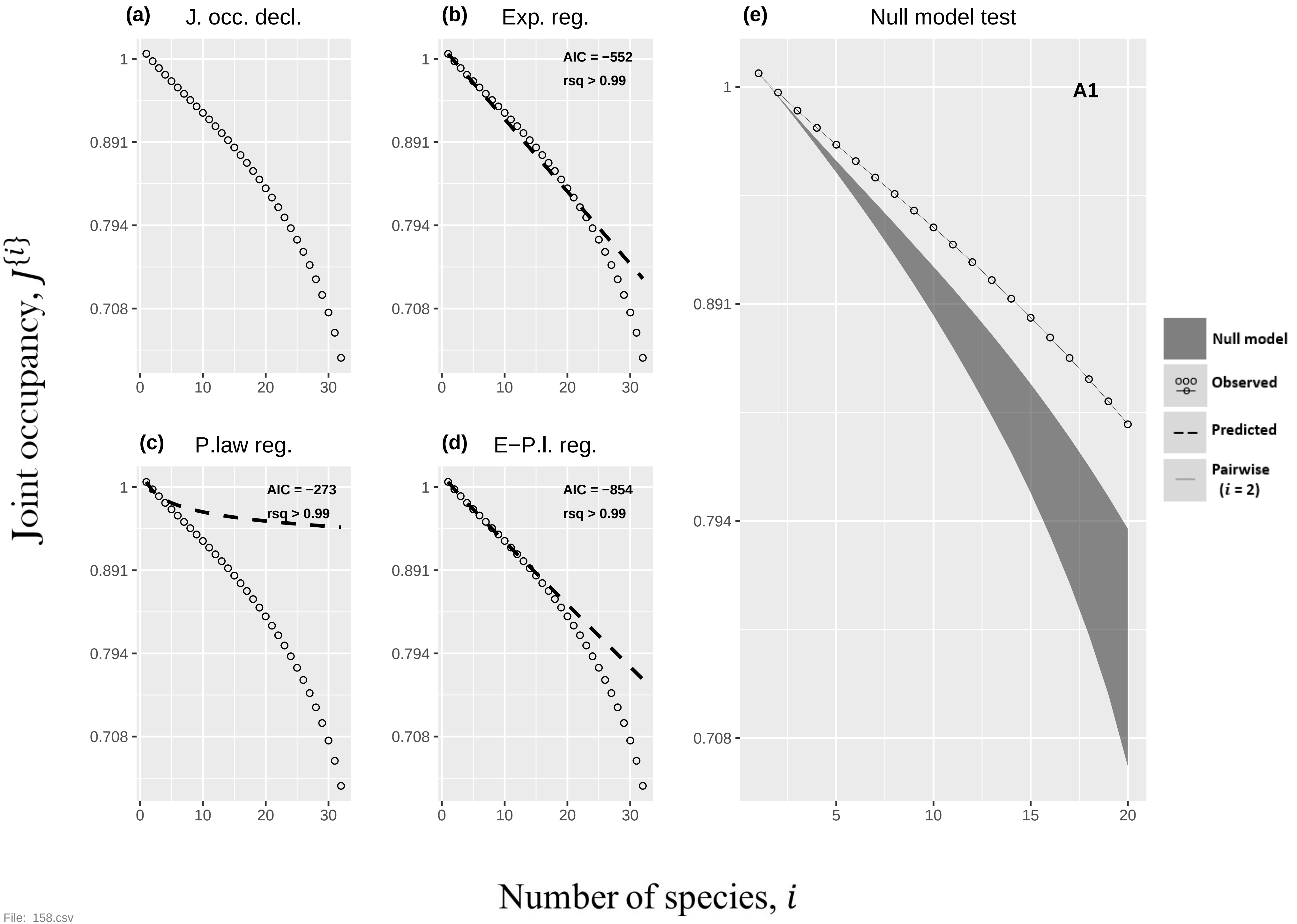
(a)
J. occ. decl.
(b)
Exp. reg.
AIC = −552
rsq > 0.99
(c)
P.law reg.
AIC = −273
rsq > 0.99
(d)
E−P.l. reg.
AIC = −854
rsq > 0.99
(e)
Null model test
A1
Null model
Observed
Predicted
Pairwise
(i = 2)
1
0.891
0.794
0.708
0
10
20
30
5
15
Joint occupancy, J{i}
Number of species, i

File: 158.csv

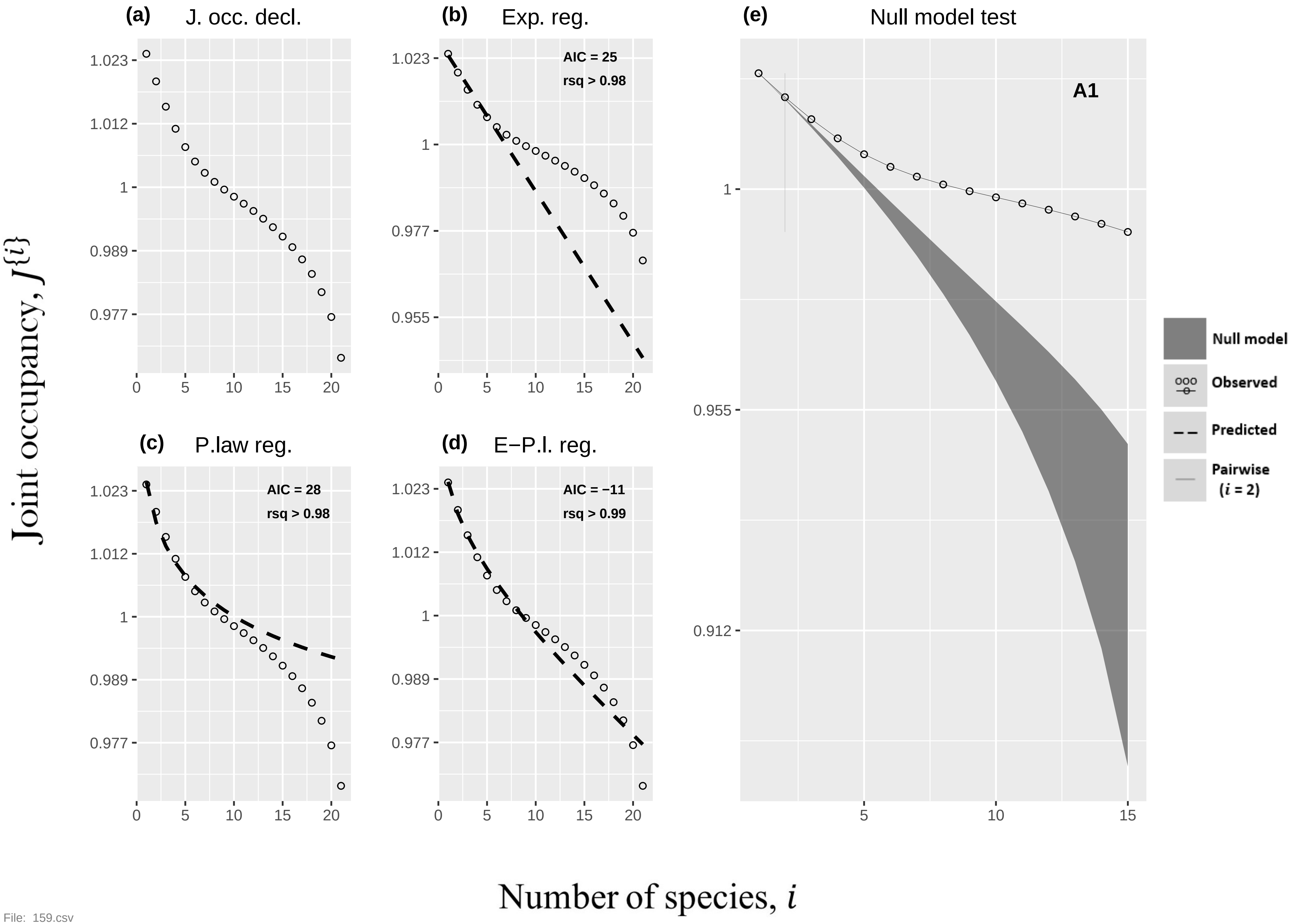

(a) J. occ. decl.
(b) Exp. reg.
AIC = 25
rsq > 0.98
(c) P.law reg.
AIC = 28
rsq > 0.98
(d) E−P.l. reg.
AIC = −11
rsq > 0.99
(e) Null model test
A1
Null model
Observed
Predicted
Pairwise ($i$ = 2)
Joint occupancy, $J\{i\}$
Number of species, $i$


File: 159.csv

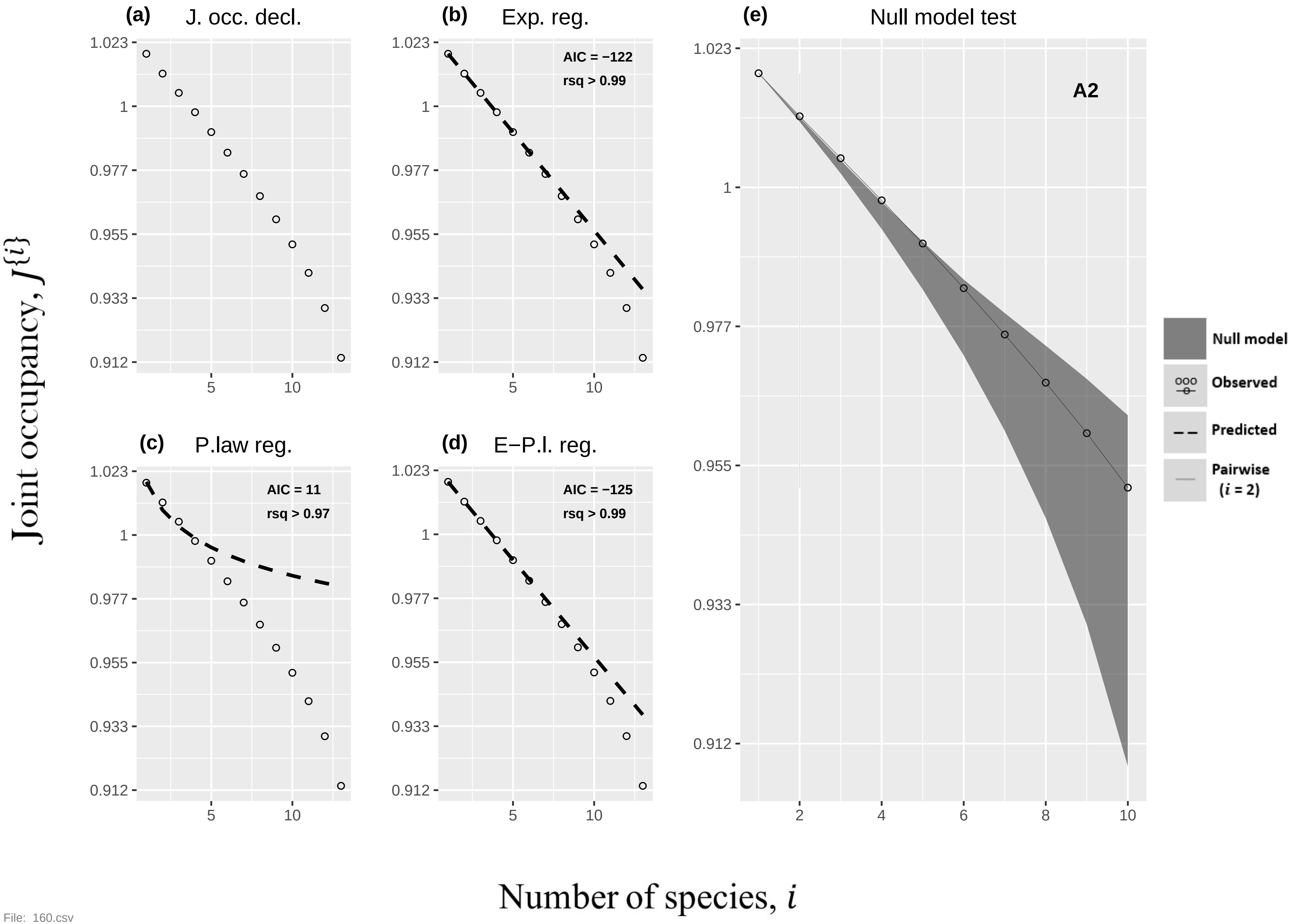

(a) J. occ. decl.
(b) Exp. reg.
AIC = −122
rsq > 0.99
(c) P.law reg.
AIC = 11
rsq > 0.97
(d) E−P.l. reg.
AIC = −125
rsq > 0.99
(e) Null model test
A2
Null model
Observed
Predicted
Pairwise
(i = 2)
1.023
1
0.977
0.955
0.933
0.912
5
10
2
4
6
8
Joint occupancy, J{i}
Number of species, i


File: 160.csv

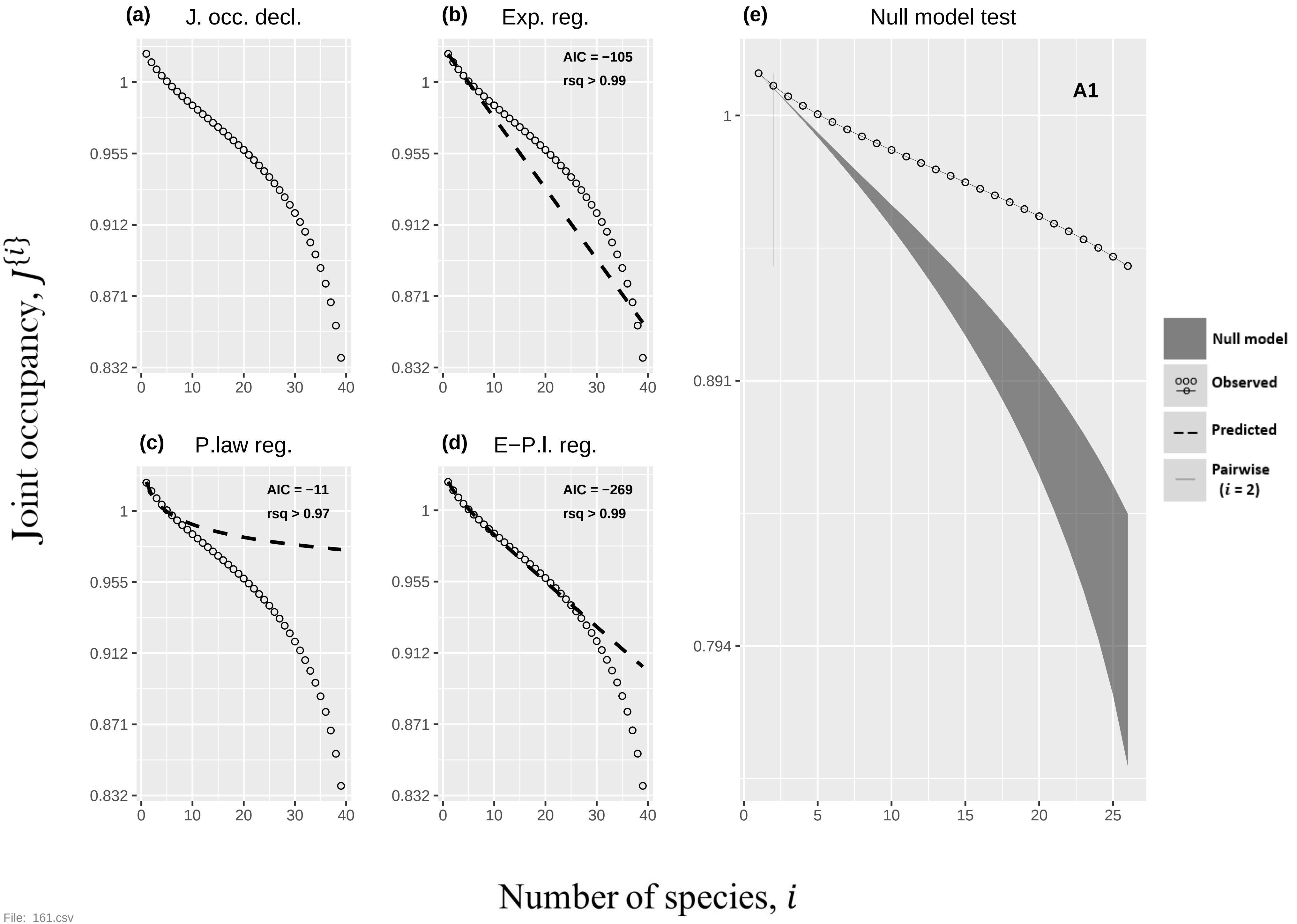

(a) J. occ. decl.
(b) Exp. reg.
AIC = −105
rsq > 0.99
(c) P.law reg.
AIC = −11
rsq > 0.97
(d) E−P.l. reg.
AIC = −269
rsq > 0.99
(e) Null model test
A1
Null model
Observed
Predicted
Pairwise (i = 2)
1
0.955
0.912
0.871
0.832
0.891
0.794
0
10
20
30
40
5
15
25
Joint occupancy, J{i}
Number of species, i


File: 161.csv

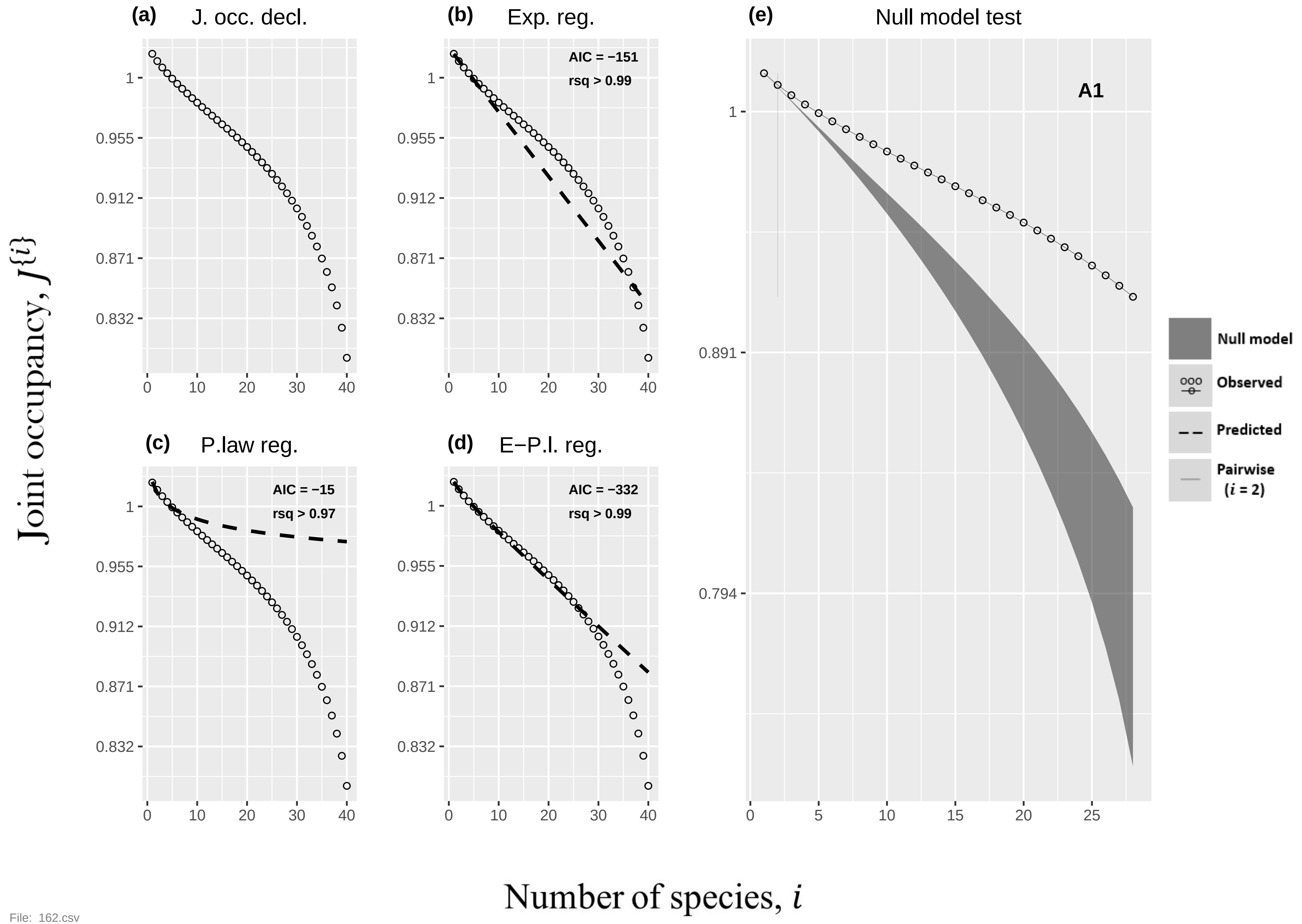
(a) J. occ. decl.
(b) Exp. reg.
AIC = −151
rsq > 0.99
(c) P.law reg.
AIC = −15
rsq > 0.97
(d) E−P.l. reg.
AIC = −332
rsq > 0.99
(e) Null model test
A1
Null model
Observed
Predicted
Pairwise (i = 2)
Joint occupancy, J{i}
Number of species, i
1
0.955
0.912
0.871
0.832
0.891
0.794
0
10
20
30
40
5
15
25


File: 162.csv

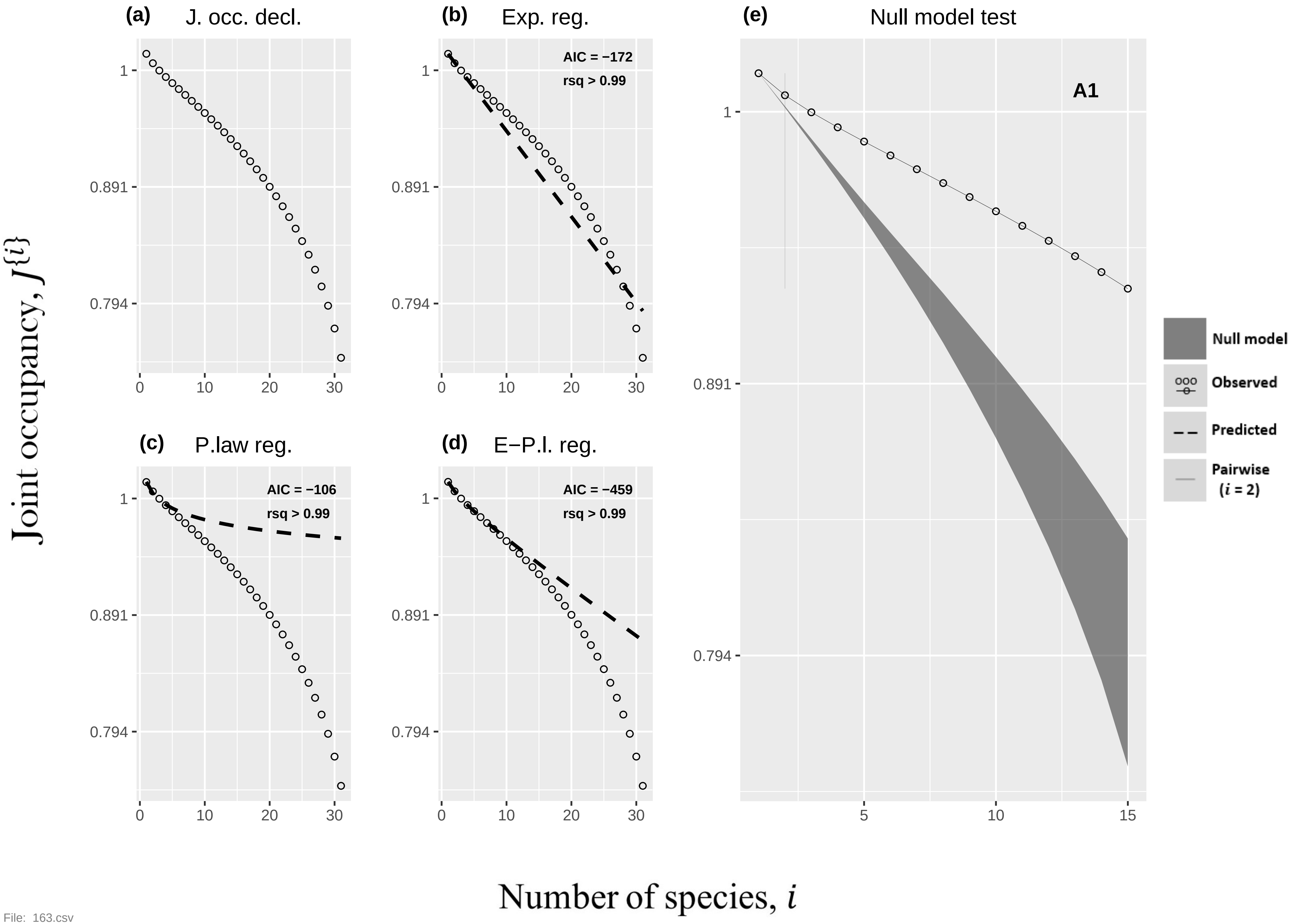

(a) J. occ. decl.
(b) Exp. reg.
AIC = −172
rsq > 0.99
(e) Null model test
A1
(c) P.law reg.
AIC = −106
rsq > 0.99
(d) E−P.l. reg.
AIC = −459
rsq > 0.99
Null model
Observed
Predicted
Pairwise (i = 2)
Joint occupancy, J{i}
Number of species, i


File: 163.csv

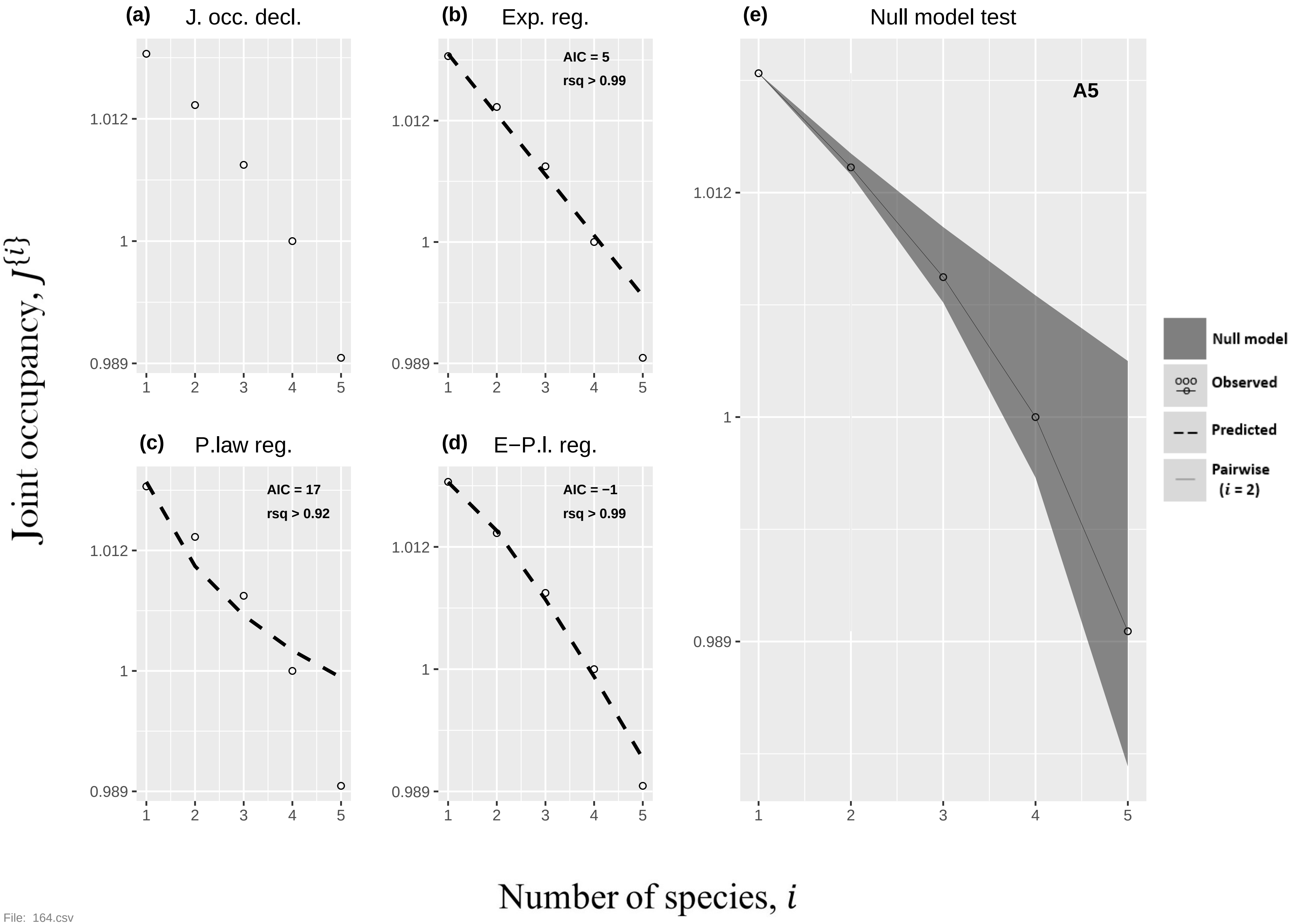
(a)
J. occ. decl.
(b)
Exp. reg.
AIC = 5
rsq > 0.99
(c)
P.law reg.
AIC = 17
rsq > 0.92
(d)
E−P.l. reg.
AIC = −1
rsq > 0.99
(e)
Null model test
A5
1.012
1
0.989
1 2 3 4 5
Null model
Observed
Predicted
Pairwise
(i = 2)
Joint occupancy, J{i}
Number of species, i

File: 164.csv

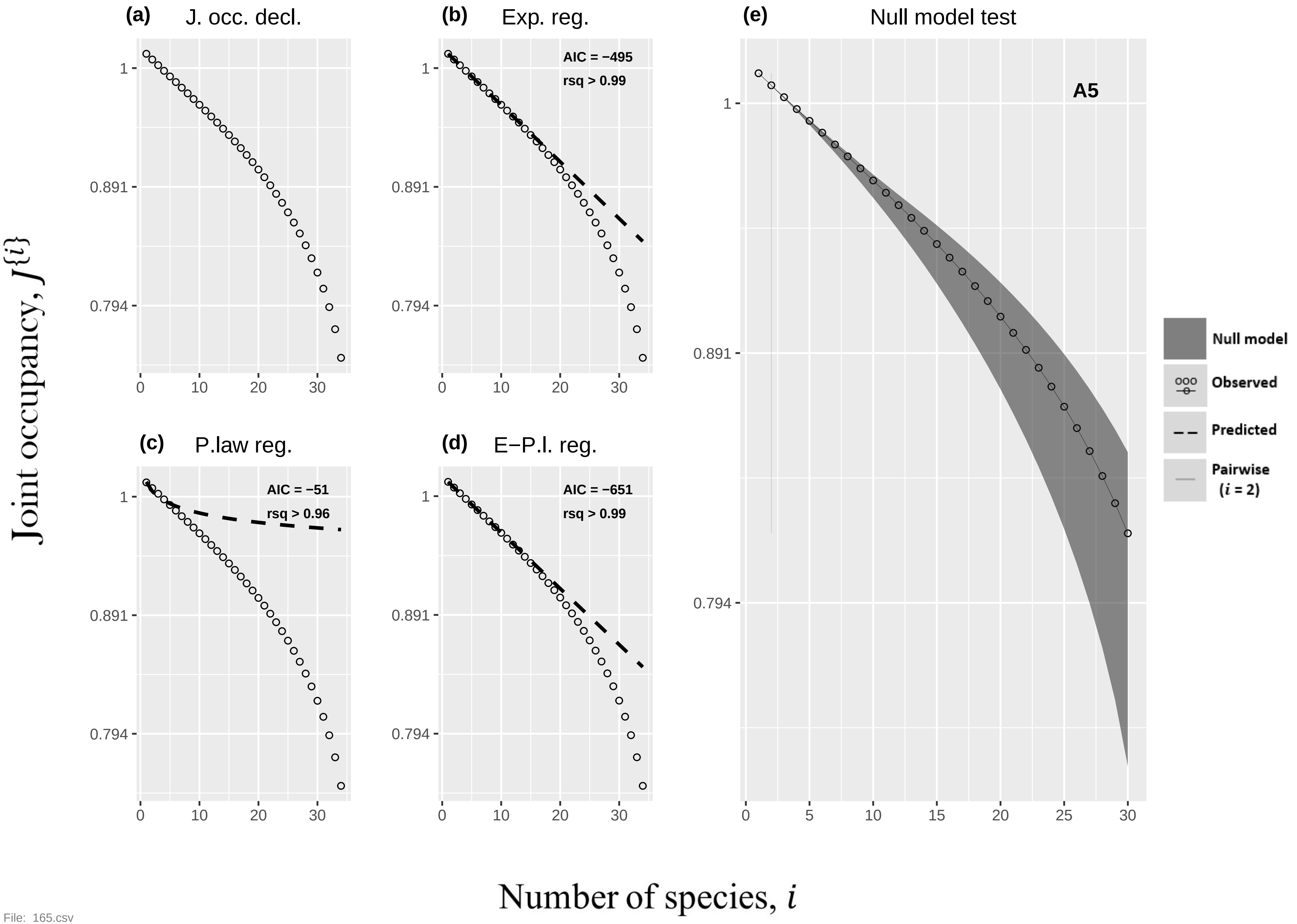

(a) J. occ. decl.
(b) Exp. reg.
AIC = −495
rsq > 0.99
(c) P.law reg.
AIC = −51
rsq > 0.96
(d) E−P.l. reg.
AIC = −651
rsq > 0.99
(e) Null model test
A5
Null model
Observed
Predicted
Pairwise (i = 2)
1
0.891
0.794
0
5
10
15
20
25
30
Joint occupancy, J{i}
Number of species, i


File: 165.csv

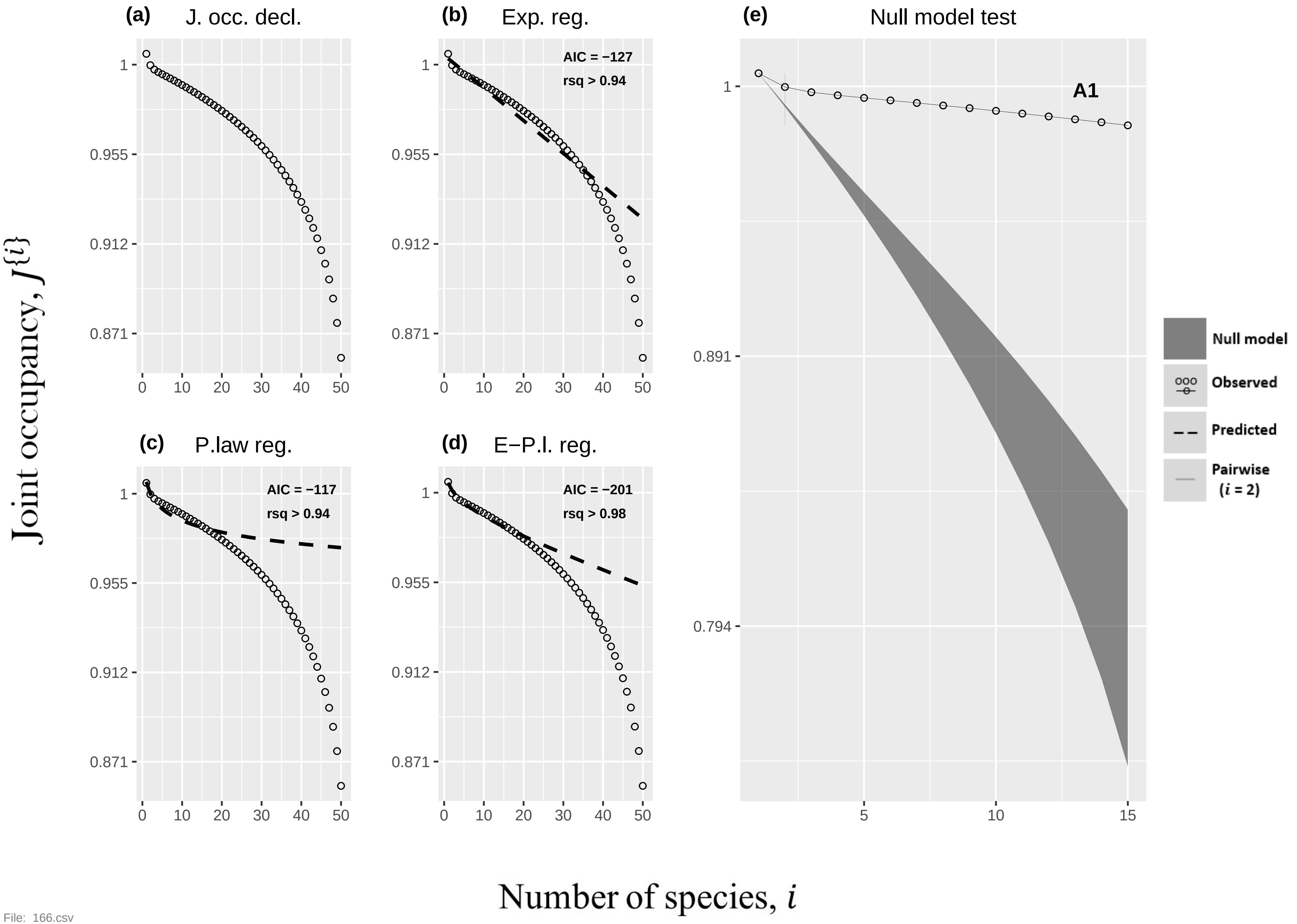

(a) J. occ. decl.
(b) Exp. reg.
AIC = −127
rsq > 0.94
(c) P.law reg.
AIC = −117
rsq > 0.94
(d) E−P.l. reg.
AIC = −201
rsq > 0.98
(e) Null model test
A1
Null model
Observed
Predicted
Pairwise ($i$ = 2)
1
0.955
0.912
0.871
0.891
0.794
0
10
20
30
40
50
5
15
Joint occupancy, $J\{i\}$
Number of species, $i$

File: 166.csv

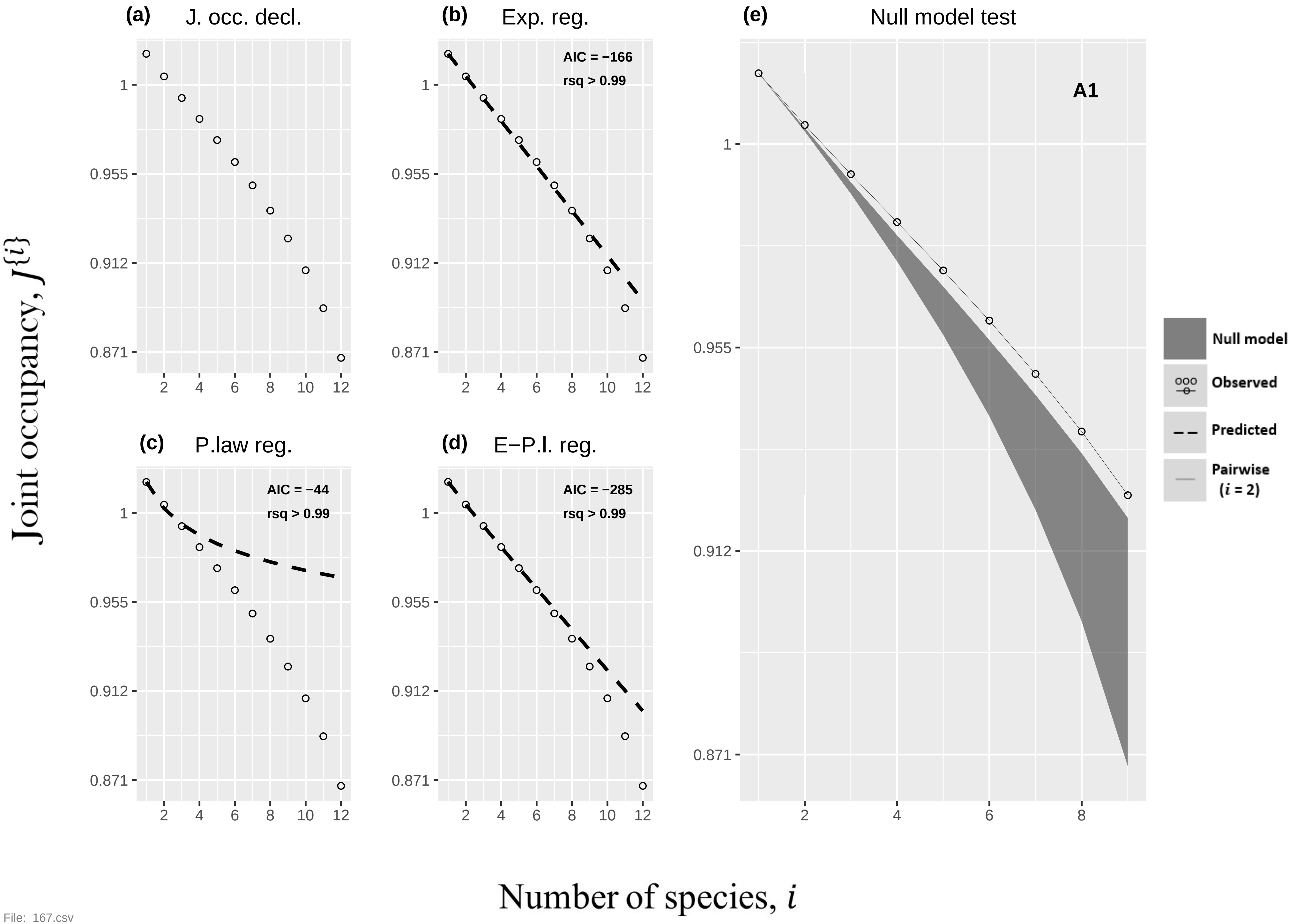

(a) J. occ. decl.
(b) Exp. reg.
AIC = −166
rsq > 0.99
(c) P.law reg.
AIC = −44
rsq > 0.99
(d) E−P.l. reg.
AIC = −285
rsq > 0.99
(e) Null model test
A1
Null model
Observed
Predicted
Pairwise (i = 2)
1
0.955
0.912
0.871
2
4
6
8
10
12
Joint occupancy, J{i}
Number of species, i


File: 167.csv

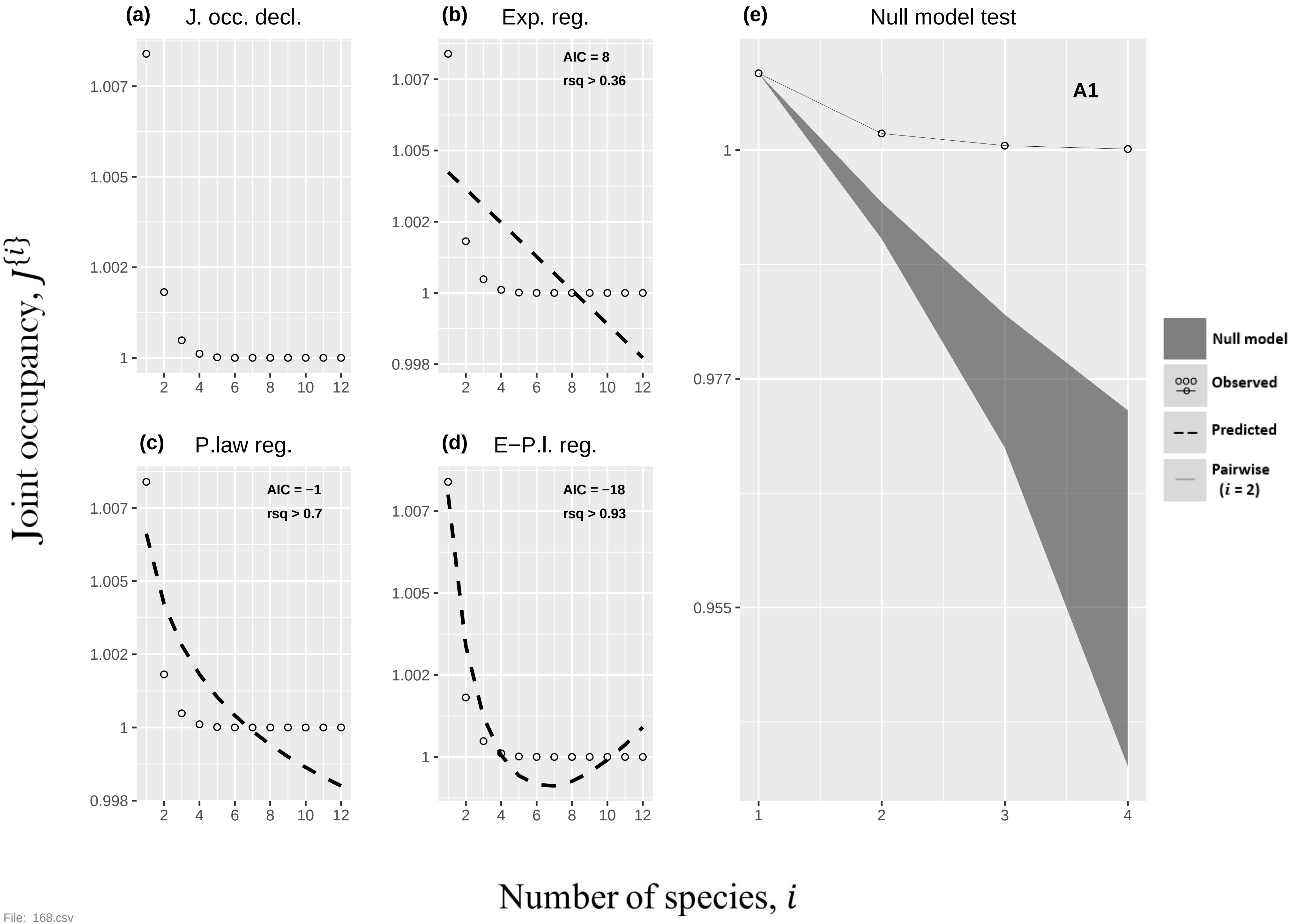
(a) J. occ. decl.
(b) Exp. reg.
AIC = 8
rsq > 0.36
(e) Null model test
A1
(c) P.law reg.
AIC = −1
rsq > 0.7
(d) E−P.l. reg.
AIC = −18
rsq > 0.93
Null model
Observed
Predicted
Pairwise (i = 2)
Joint occupancy, J{i}
Number of species, i

File: 168.csv

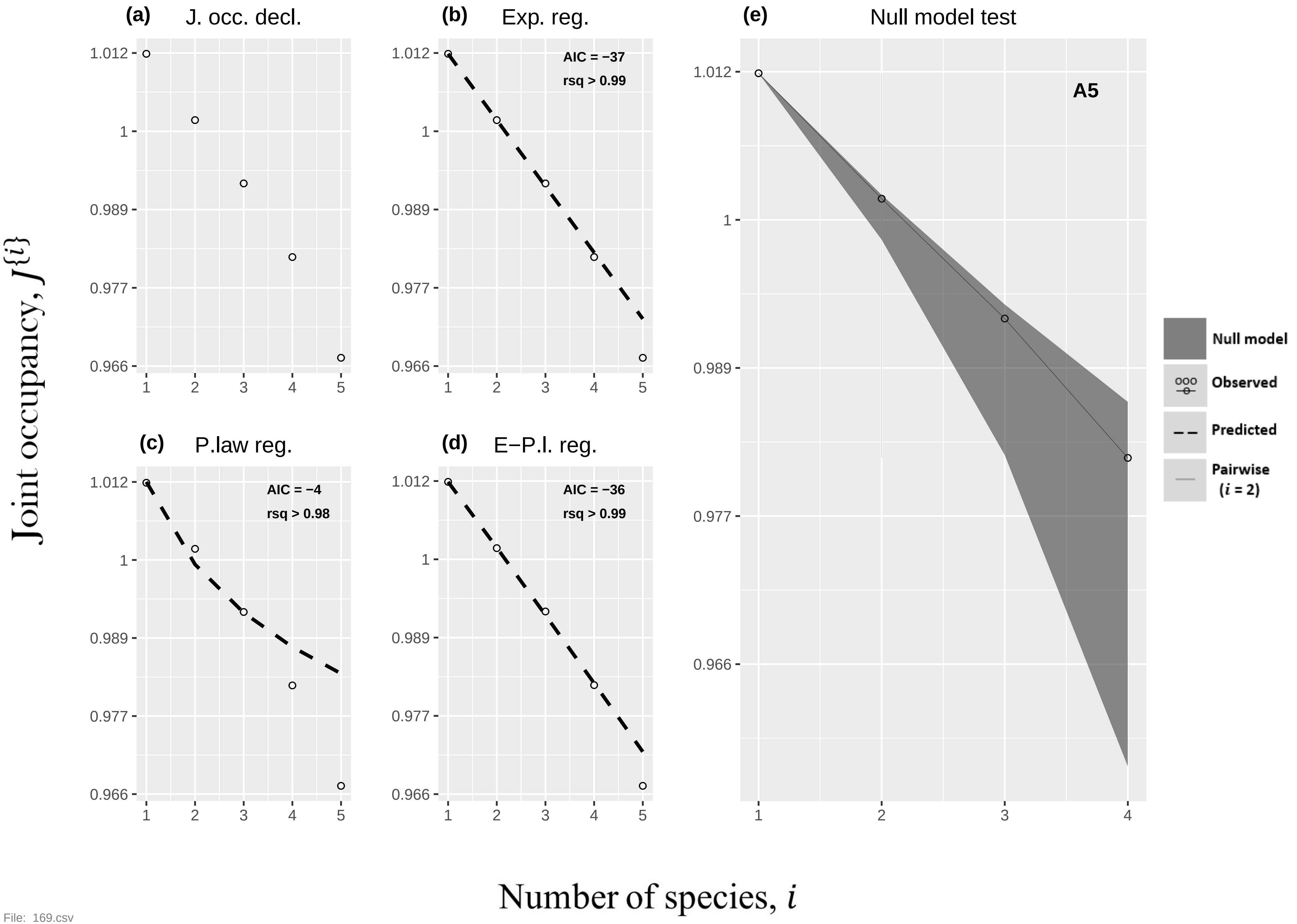

(a) J. occ. decl.
(b) Exp. reg.
AIC = −37
rsq > 0.99
(c) P.law reg.
AIC = −4
rsq > 0.98
(d) E−P.l. reg.
AIC = −36
rsq > 0.99
(e) Null model test
A5
Null model
Observed
Predicted
Pairwise (i = 2)
1.012
1
0.989
0.977
0.966
Joint occupancy, J{i}
Number of species, i


File: 169.csv

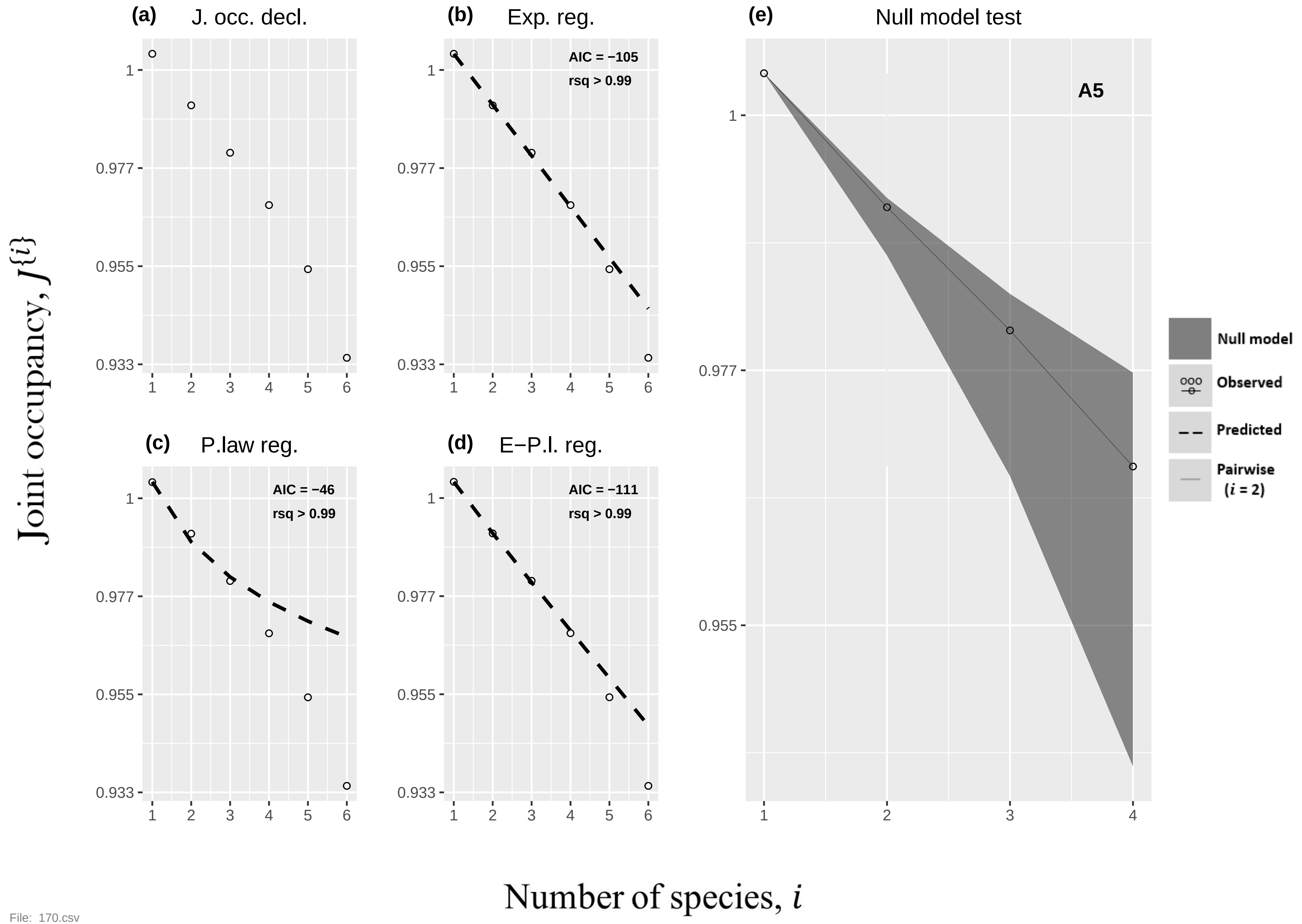

(a) J. occ. decl.
(b) Exp. reg.
AIC = −105
rsq > 0.99
(c) P.law reg.
AIC = −46
rsq > 0.99
(d) E−P.l. reg.
AIC = −111
rsq > 0.99
(e) Null model test
A5
Null model
Observed
Predicted
Pairwise (i = 2)
1
0.977
0.955
0.933
1 2 3 4 5 6
Joint occupancy, J{i}
Number of species, i


File: 170.csv

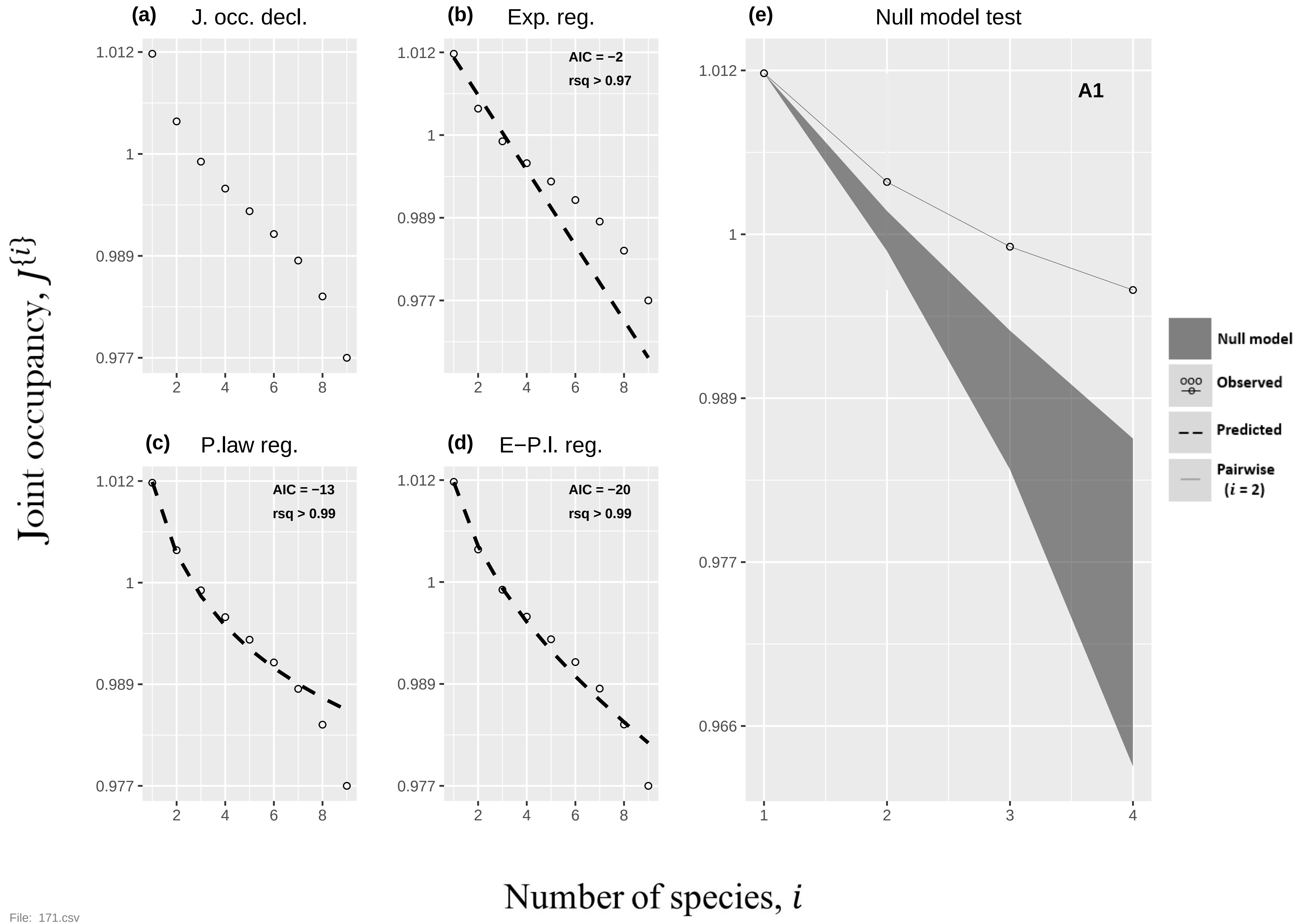

(a) J. occ. decl.
(b) Exp. reg.
AIC = −2
rsq > 0.97
(c) P.law reg.
AIC = −13
rsq > 0.99
(d) E−P.l. reg.
AIC = −20
rsq > 0.99
(e) Null model test
A1
Null model
Observed
Predicted
Pairwise (i = 2)
Joint occupancy, J{i}
Number of species, i
1.012
1
0.989
0.977
0.966


File: 171.csv

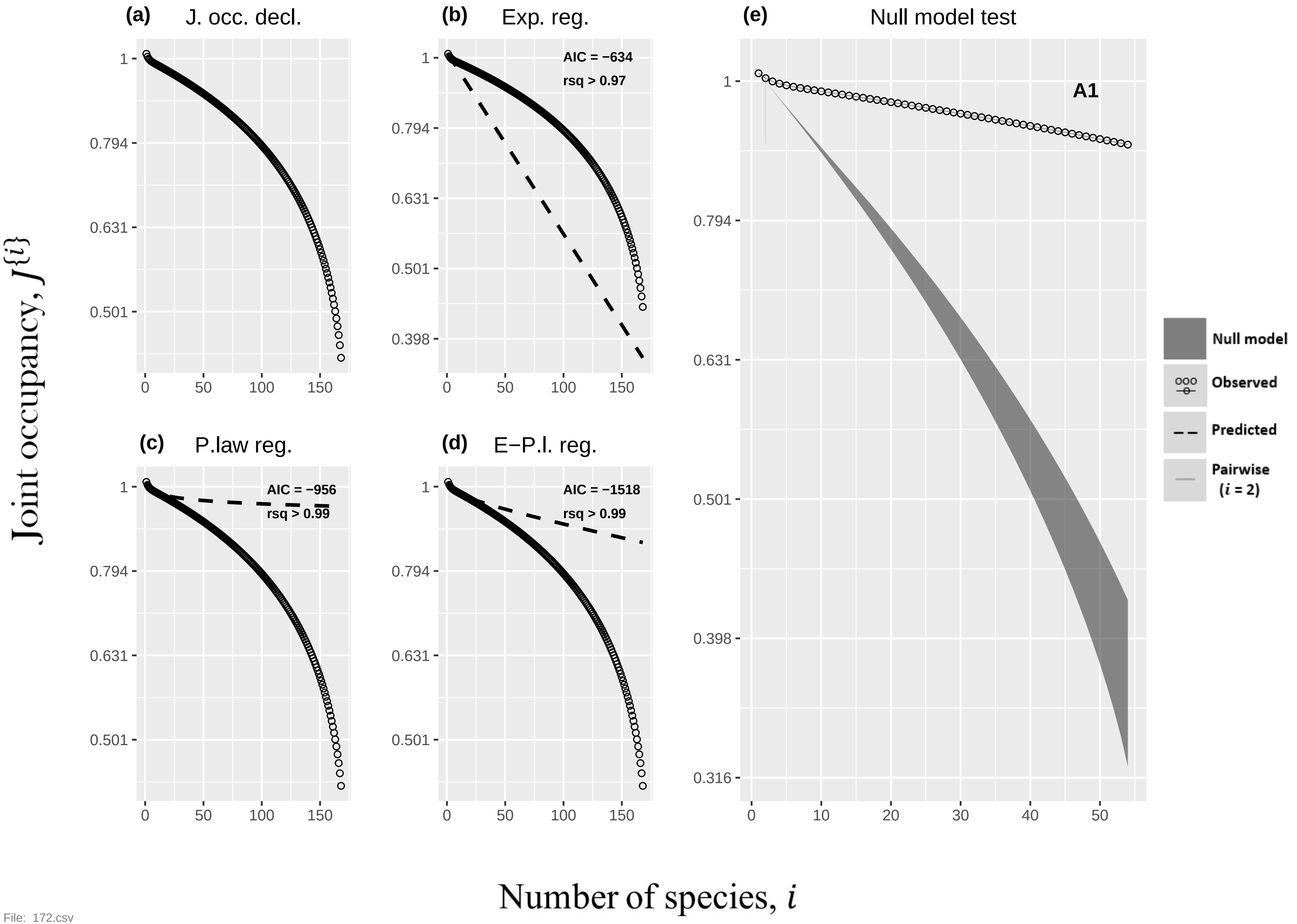
(a) J. occ. decl.
(b) Exp. reg.
AIC = −634
rsq > 0.97
(c) P.law reg.
AIC = −956
rsq > 0.99
(d) E−P.l. reg.
AIC = −1518
rsq > 0.99
(e) Null model test
A1
Null model
Observed
Predicted
Pairwise (i = 2)
Joint occupancy, J{i}
Number of species, i
1
0.794
0.631
0.501
0.398
0.316
0
50
100
150
10
20
30
40

File: 172.csv

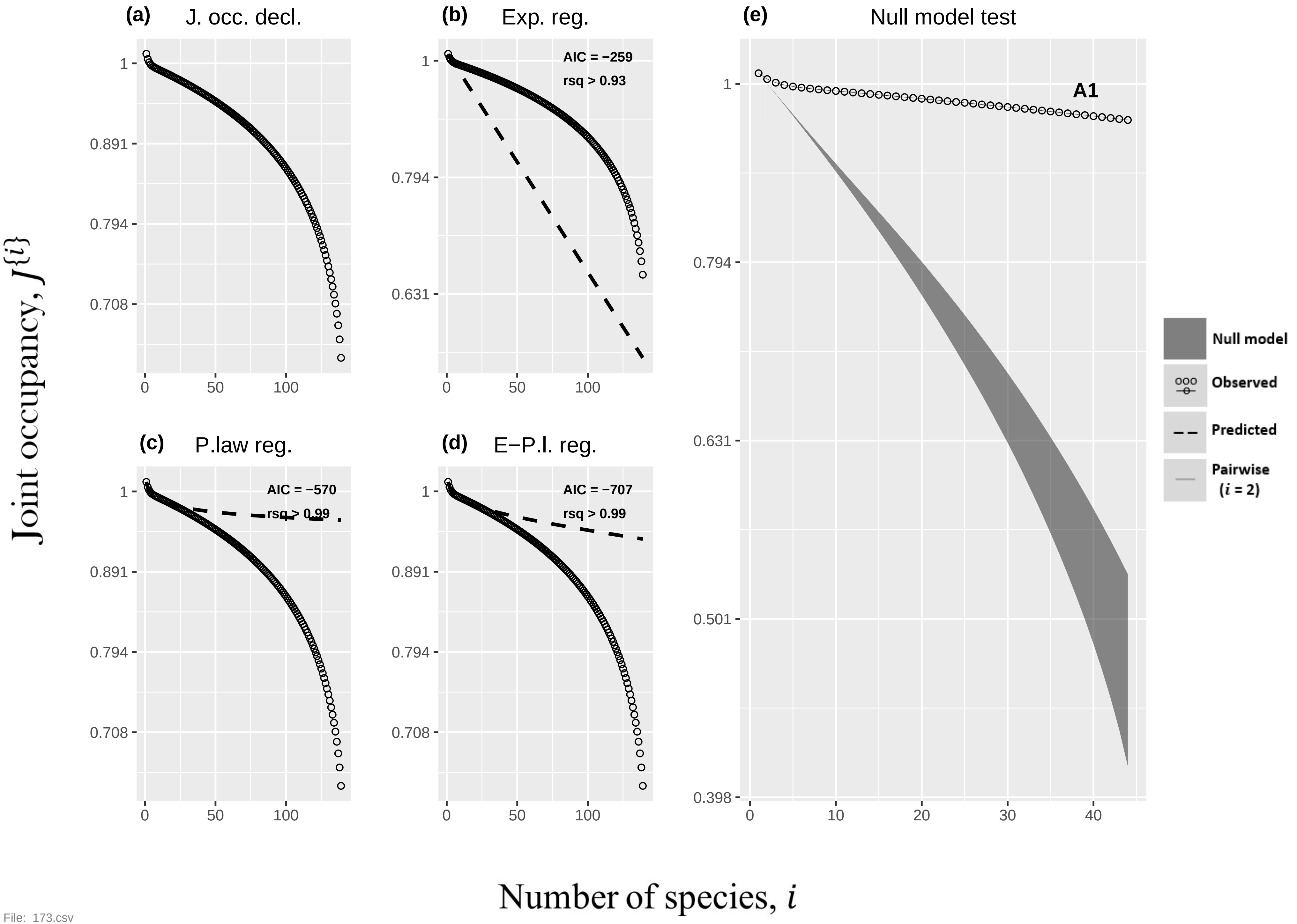

(a)
J. occ. decl.
(b)
Exp. reg.
AIC = −259
rsq > 0.93
(c)
P.law reg.
AIC = −570
rsq > 0.99
(d)
E−P.l. reg.
AIC = −707
rsq > 0.99
(e)
Null model test
A1
Null model
Observed
Predicted
Pairwise (i = 2)
Joint occupancy, J{i}
Number of species, i


File: 173.csv

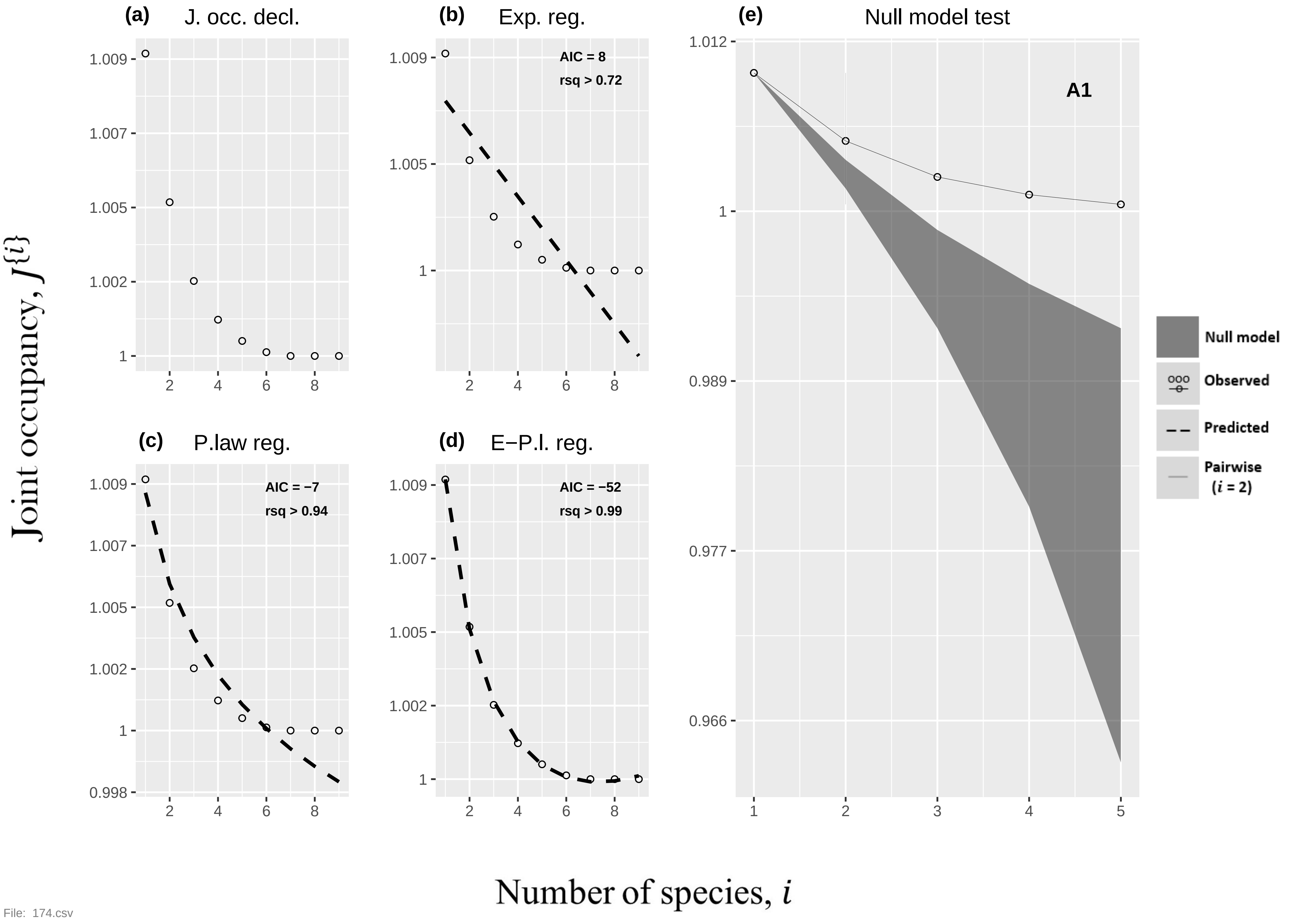

(a) J. occ. decl.
(b) Exp. reg.
AIC = 8
rsq > 0.72
(c) P.law reg.
AIC = −7
rsq > 0.94
(d) E−P.l. reg.
AIC = −52
rsq > 0.99
(e) Null model test
A1
Null model
Observed
Predicted
Pairwise (i = 2)
Joint occupancy, J{i}
Number of species, i
File: 174.csv

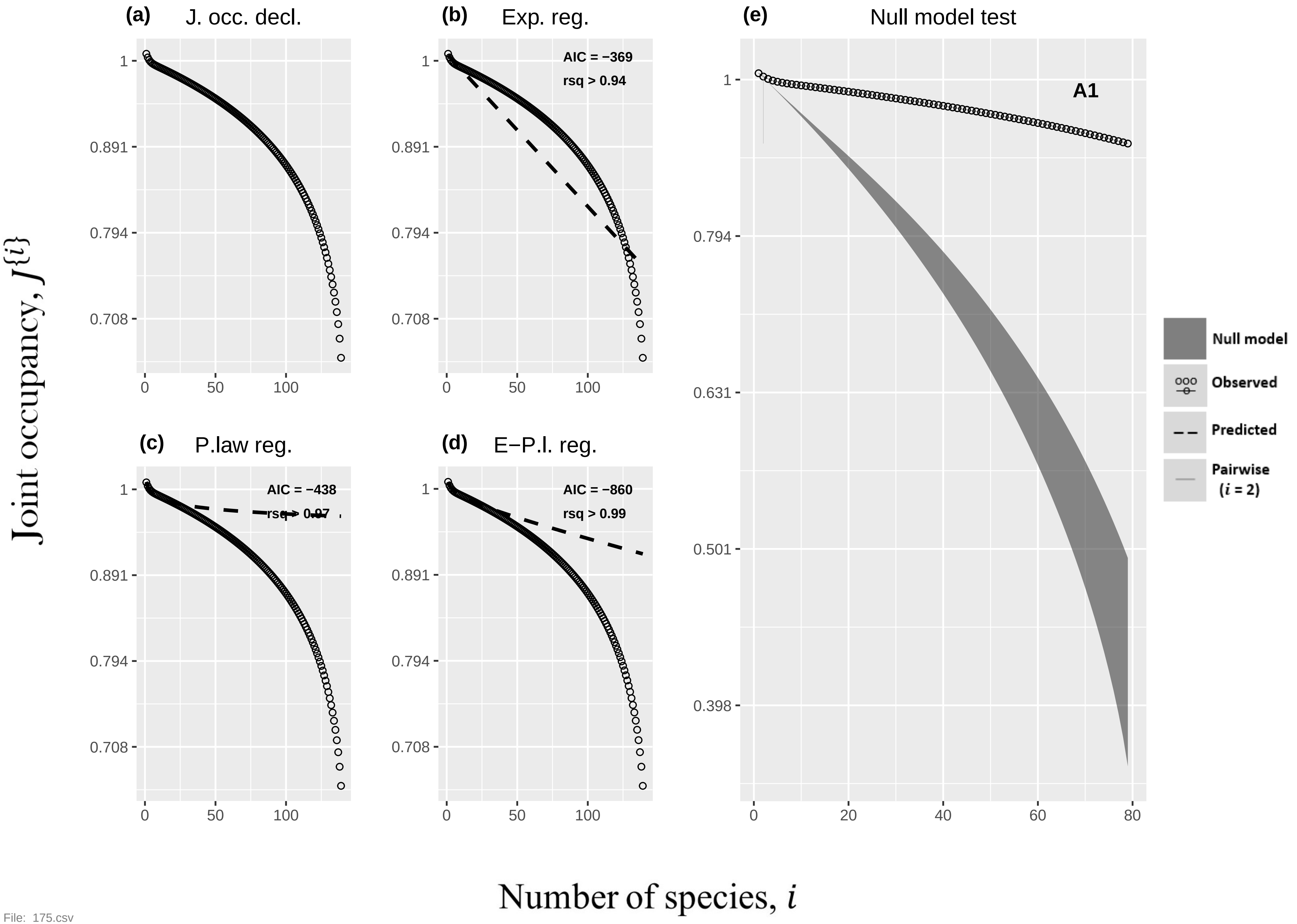
(a) J. occ. decl.
(b) Exp. reg.
AIC = −369
rsq > 0.94
(c) P.law reg.
AIC = −438
rsq > 0.97
(d) E−P.l. reg.
AIC = −860
rsq > 0.99
(e) Null model test
A1
Null model
Observed
Predicted
Pairwise ($i$ = 2)
Joint occupancy, $J\{i\}$
Number of species, $i$
1
0.891
0.794
0.708
0.631
0.501
0.398
0
50
100
20
40
60
80
File: 175.csv

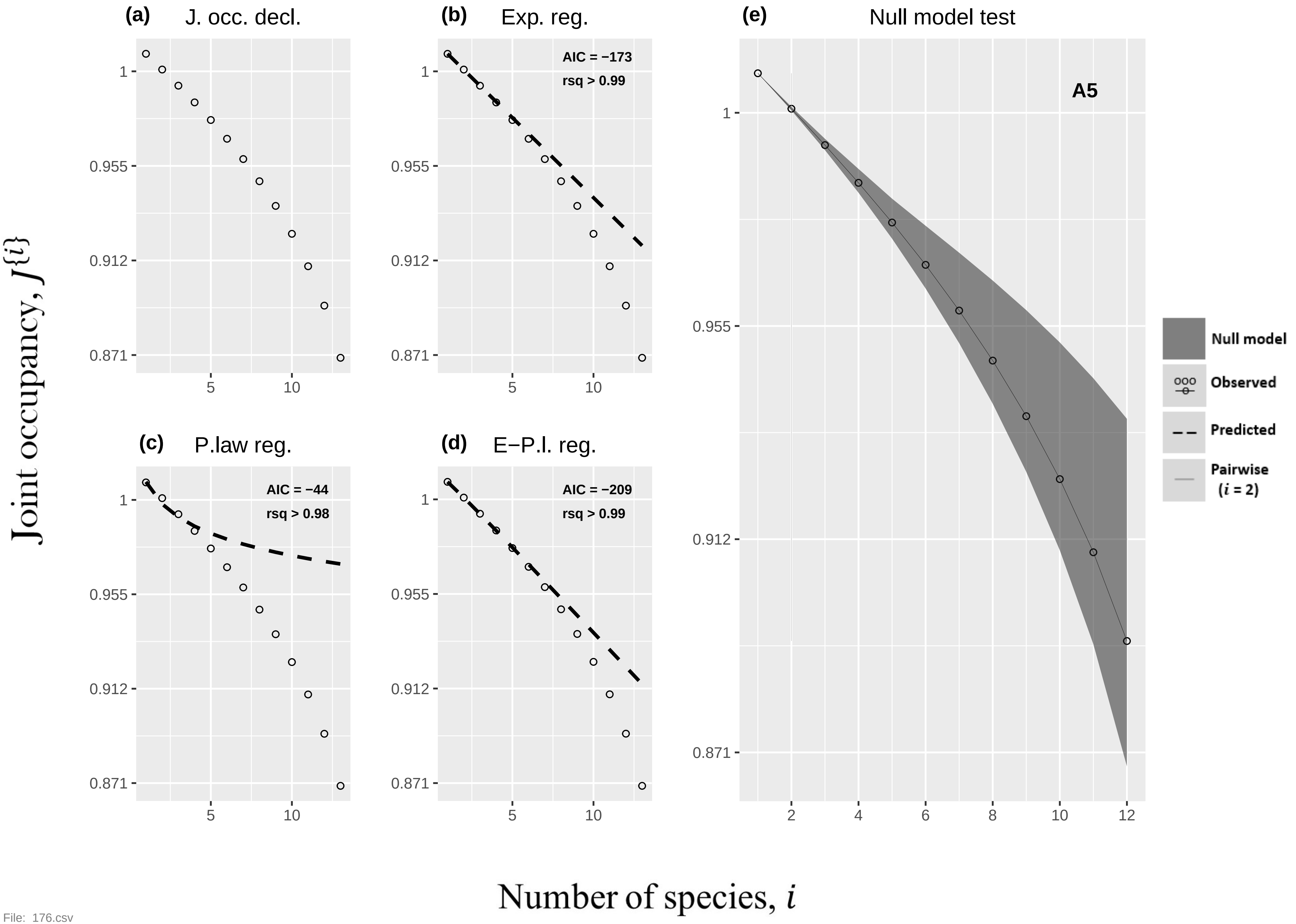
(a) J. occ. decl.
(b) Exp. reg.
AIC = −173
rsq > 0.99
(c) P.law reg.
AIC = −44
rsq > 0.98
(d) E−P.l. reg.
AIC = −209
rsq > 0.99
(e) Null model test
A5
Null model
Observed
Predicted
Pairwise (i = 2)
Joint occupancy, J{i}
Number of species, i
1
0.955
0.912
0.871
5
10
2
4
6
8
12

File: 176.csv

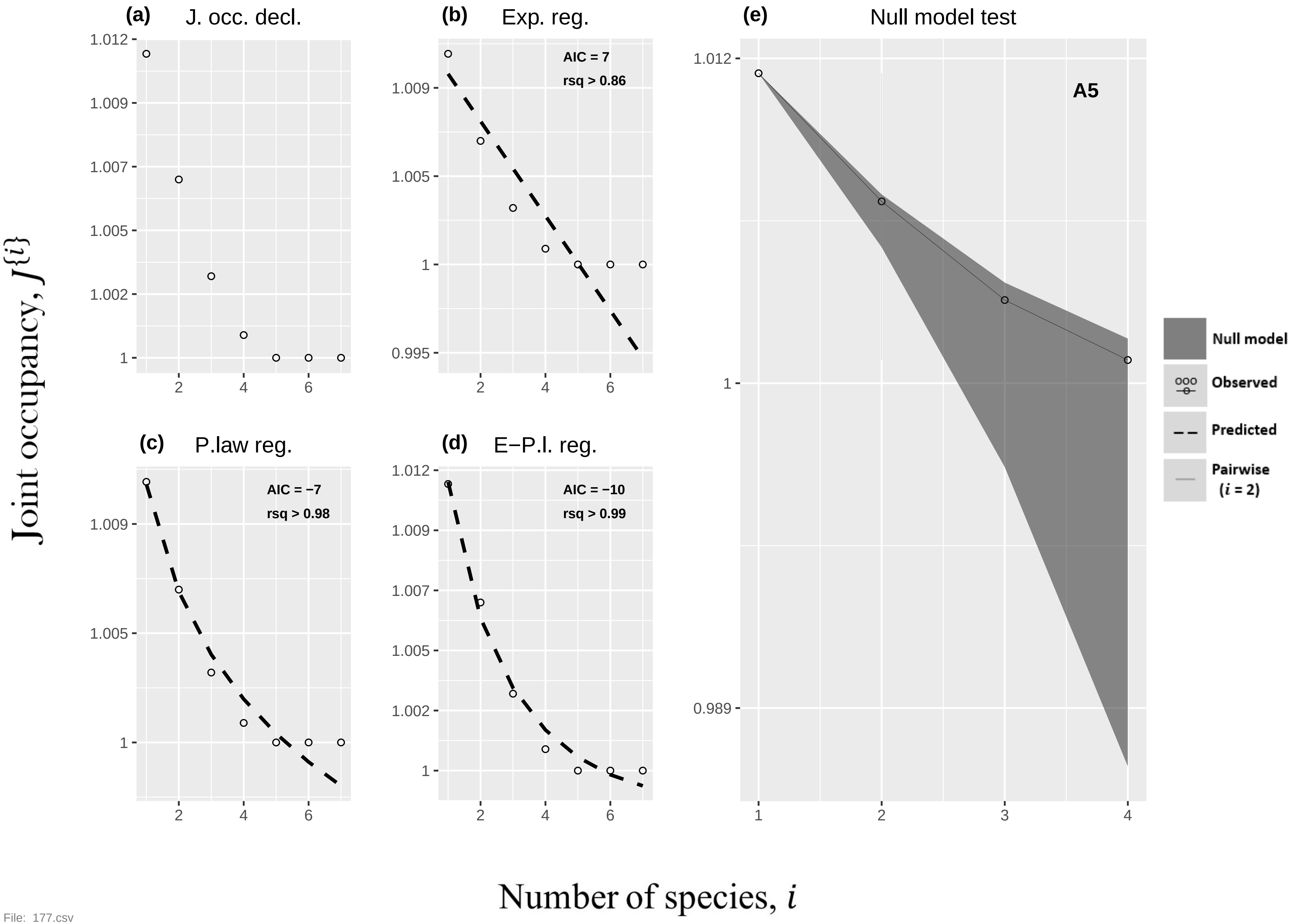

(a) J. occ. decl.
(b) Exp. reg.
AIC = 7
rsq > 0.86
(c) P.law reg.
AIC = −7
rsq > 0.98
(d) E−P.l. reg.
AIC = −10
rsq > 0.99
(e) Null model test
A5
Null model
Observed
Predicted
Pairwise (i = 2)
1.012
1.009
1.007
1.005
1.002
1
0.995
0.989
2
4
6
1
3
Joint occupancy, J{i}
Number of species, i

File: 177.csv

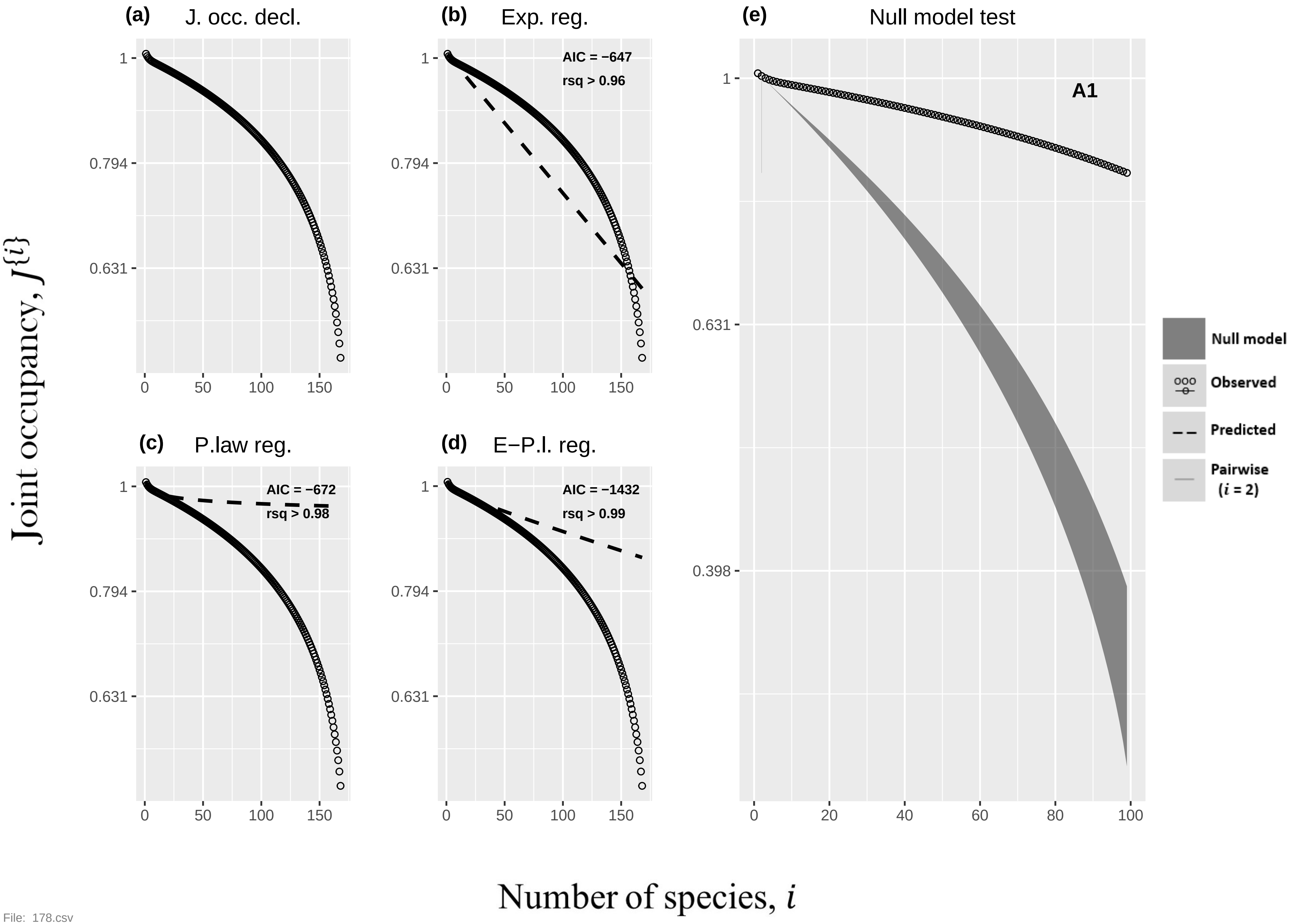
(a) J. occ. decl.
(b) Exp. reg.
AIC = −647
rsq > 0.96
(c) P.law reg.
AIC = −672
rsq > 0.98
(d) E−P.l. reg.
AIC = −1432
rsq > 0.99
(e) Null model test
A1
Null model
Observed
Predicted
Pairwise (i = 2)
Joint occupancy, J{i}
Number of species, i
1
0.794
0.631
0.398
0
50
100
150
20
40
60
80

File: 178.csv

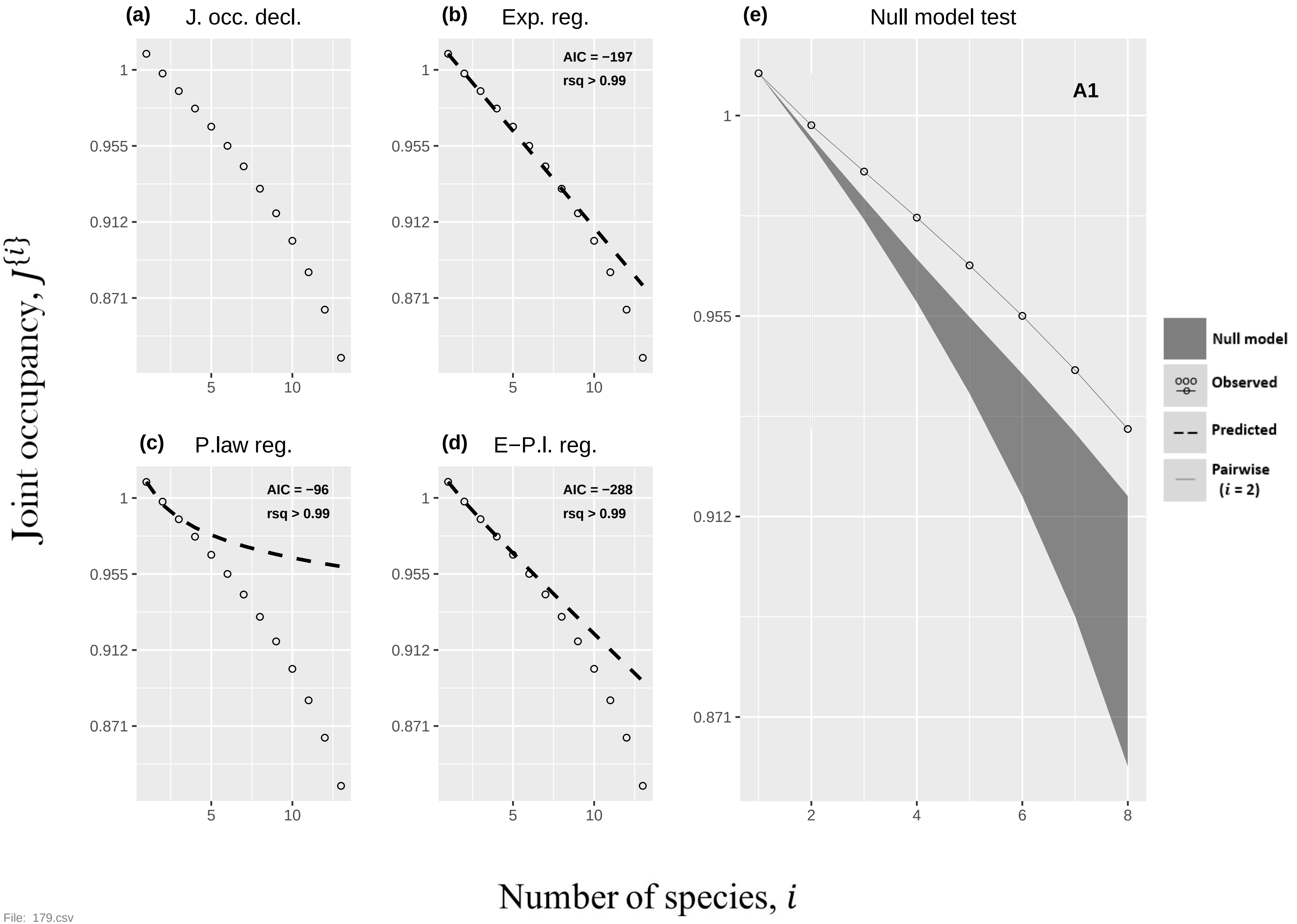

(a) J. occ. decl.
(b) Exp. reg.
AIC = −197
rsq > 0.99
(c) P.law reg.
AIC = −96
rsq > 0.99
(d) E−P.l. reg.
AIC = −288
rsq > 0.99
(e) Null model test
A1
Null model
Observed
Predicted
Pairwise (i = 2)
1
0.955
0.912
0.871
5
10
2
4
6
8
Joint occupancy, J{i}
Number of species, i


File: 179.csv

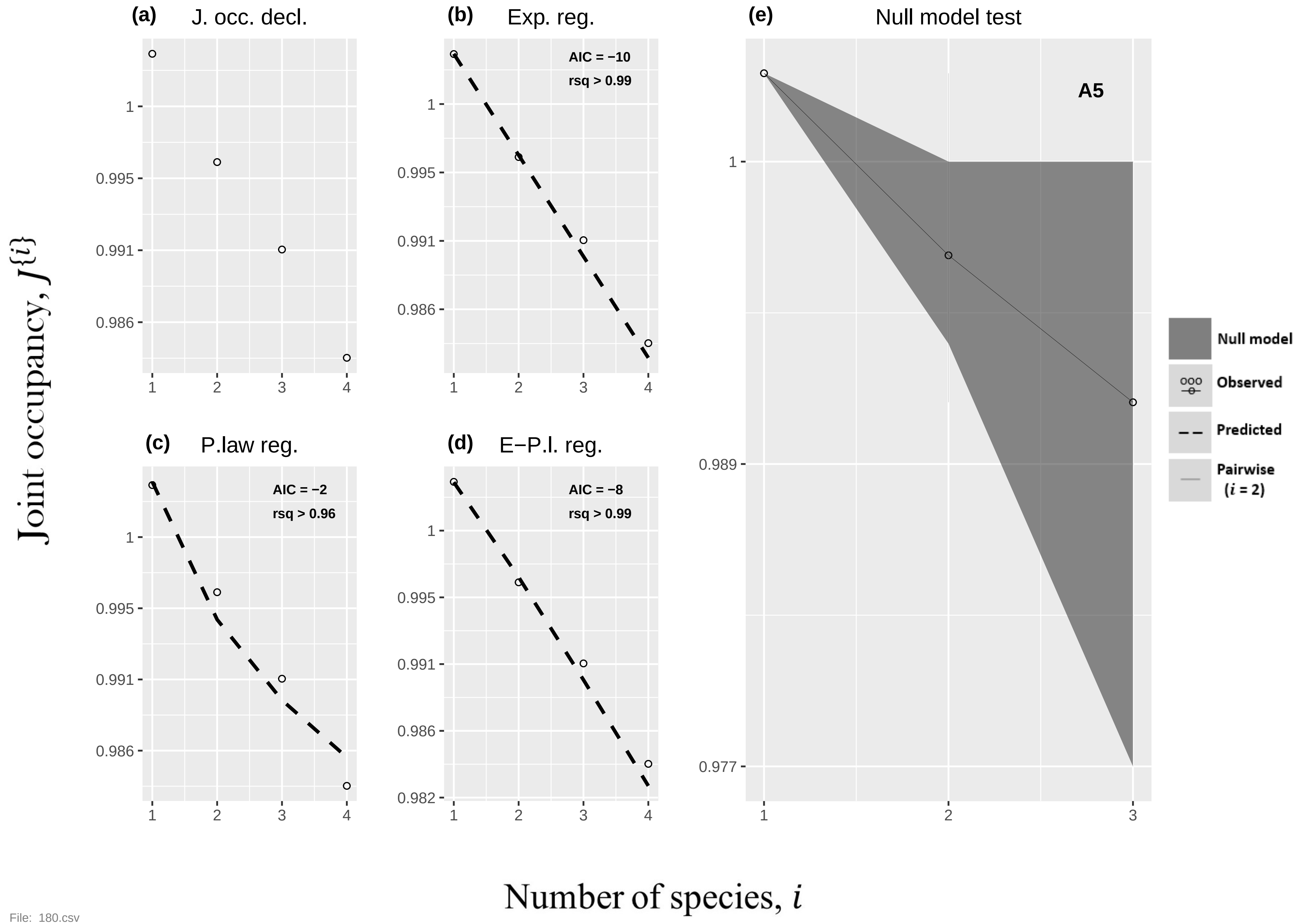
(a) J. occ. decl.
(b) Exp. reg.
AIC = −10
rsq > 0.99
(c) P.law reg.
AIC = −2
rsq > 0.96
(d) E−P.l. reg.
AIC = −8
rsq > 0.99
(e) Null model test
A5
Null model
Observed
Predicted
Pairwise (i = 2)
Joint occupancy, J{i}
Number of species, i

File: 180.csv

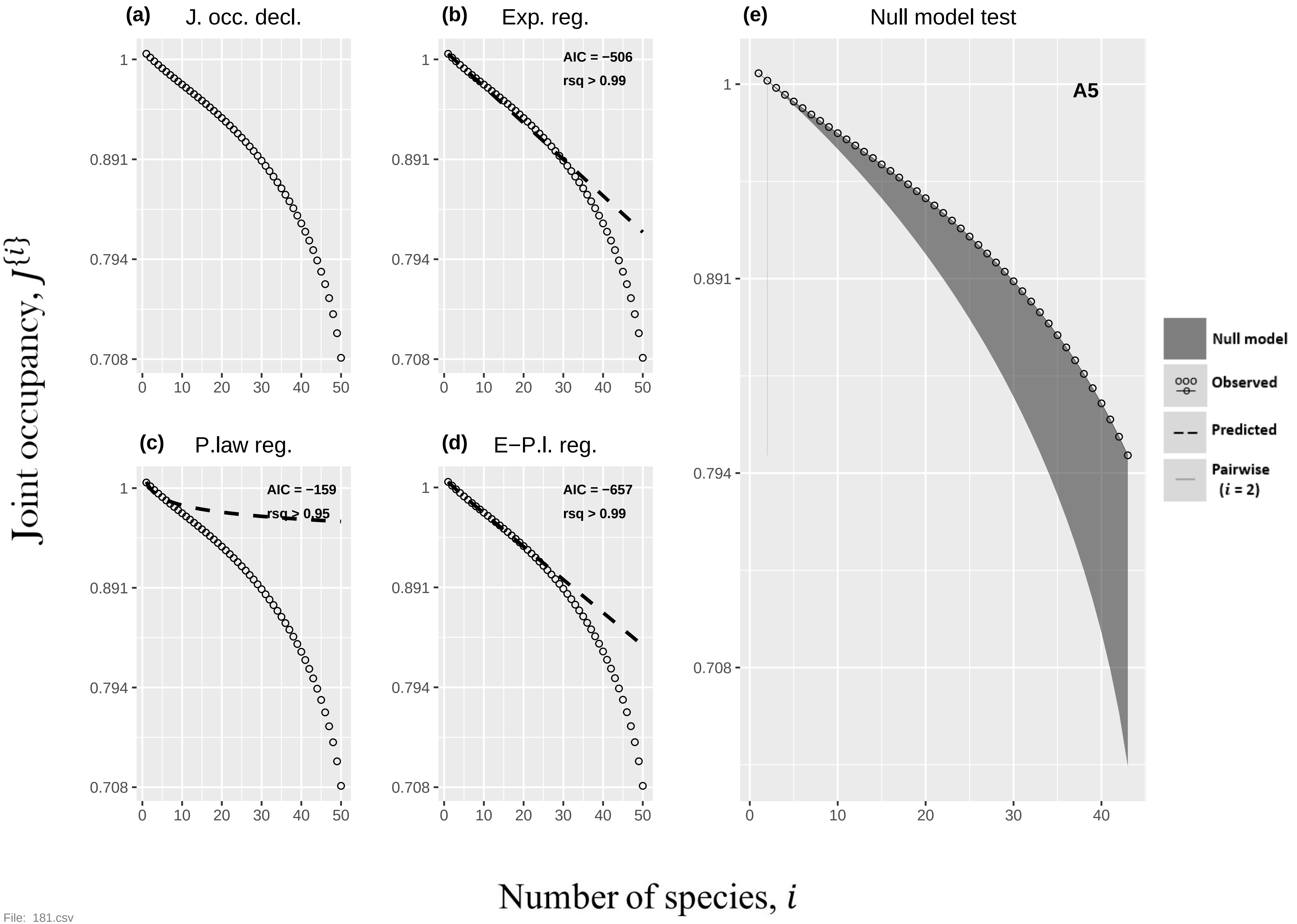
(a) J. occ. decl.
(b) Exp. reg.
AIC = −506
rsq > 0.99
(c) P.law reg.
AIC = −159
rsq > 0.95
(d) E−P.l. reg.
AIC = −657
rsq > 0.99
(e) Null model test
A5
Null model
Observed
Predicted
Pairwise
(i = 2)
Joint occupancy, J{i}
Number of species, i
1
0.891
0.794
0.708
0
10
20
30
40
50

File: 181.csv

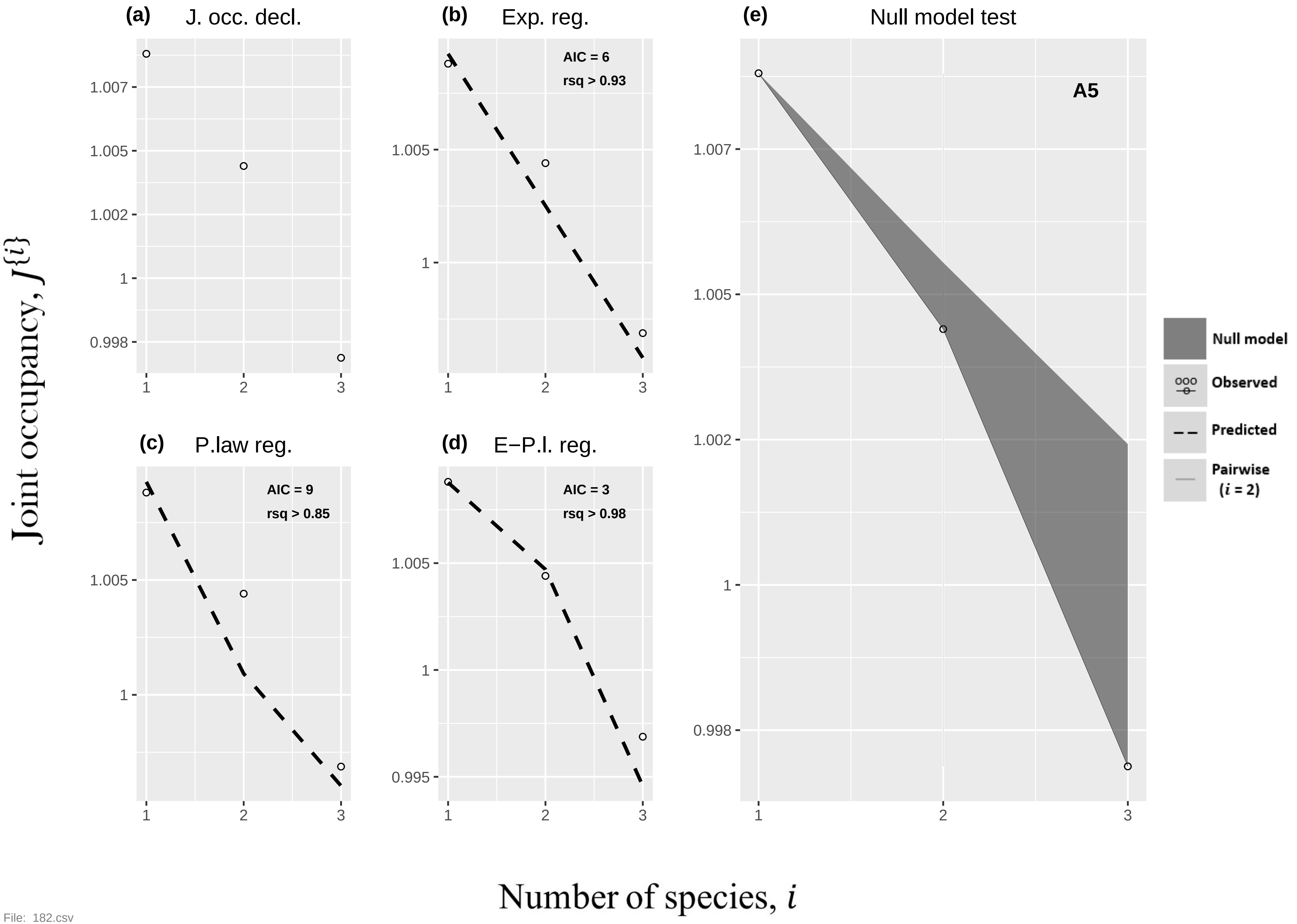

(a)
J. occ. decl.
(b)
Exp. reg.
AIC = 6
rsq > 0.93
(c)
P.law reg.
AIC = 9
rsq > 0.85
(d)
E−P.l. reg.
AIC = 3
rsq > 0.98
(e)
Null model test
A5
Null model
Observed
Predicted
Pairwise (i = 2)
Joint occupancy, J{i}
Number of species, i


File: 182.csv

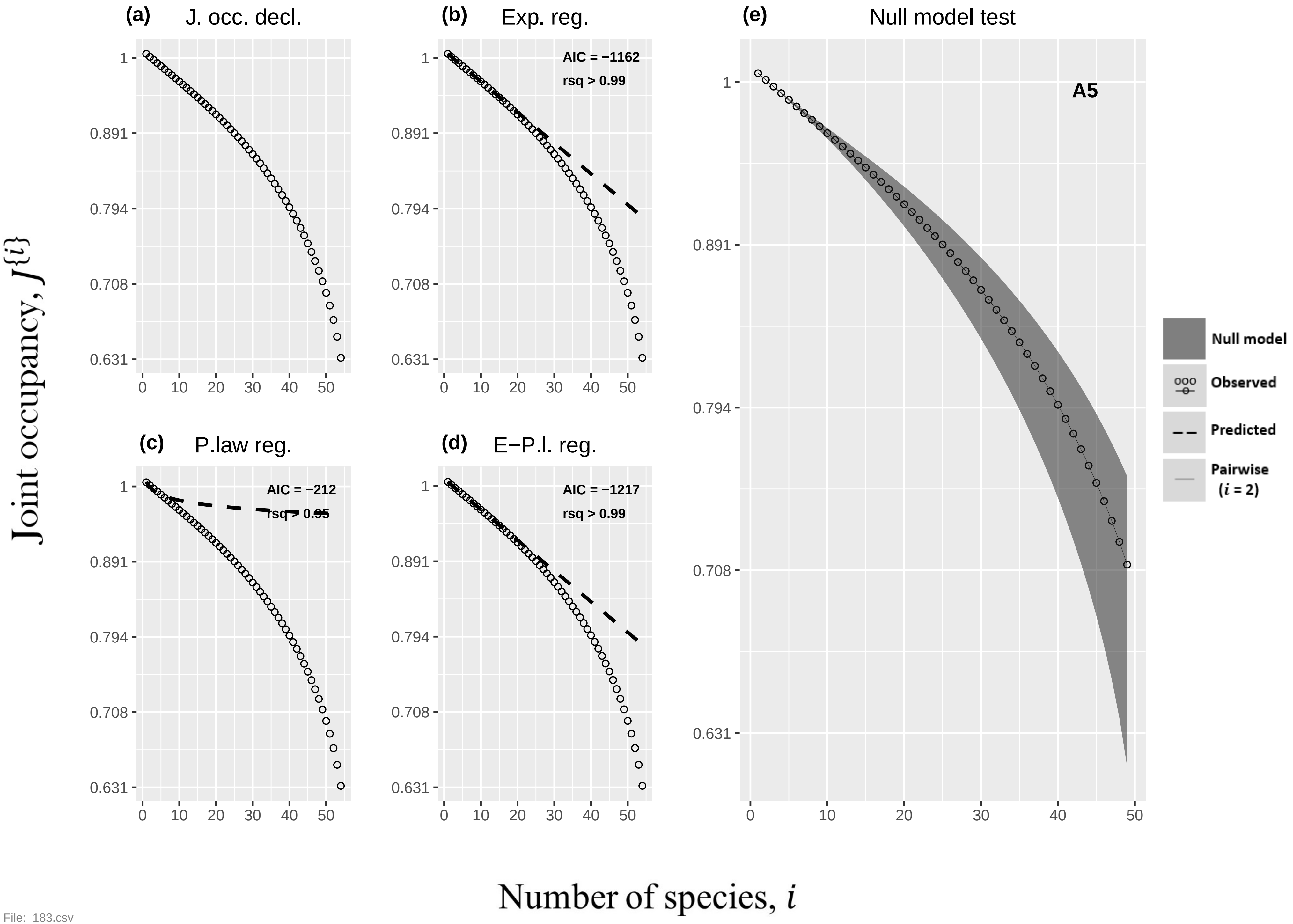

(a) J. occ. decl.
(b) Exp. reg.
AIC = −1162
rsq > 0.99
(c) P.law reg.
AIC = −212
rsq > 0.95
(d) E−P.l. reg.
AIC = −1217
rsq > 0.99
(e) Null model test
A5
Null model
Observed
Predicted
Pairwise (i = 2)
1
0.891
0.794
0.708
0.631
0
10
20
30
40
50
Joint occupancy, J{i}
Number of species, i


File: 183.csv

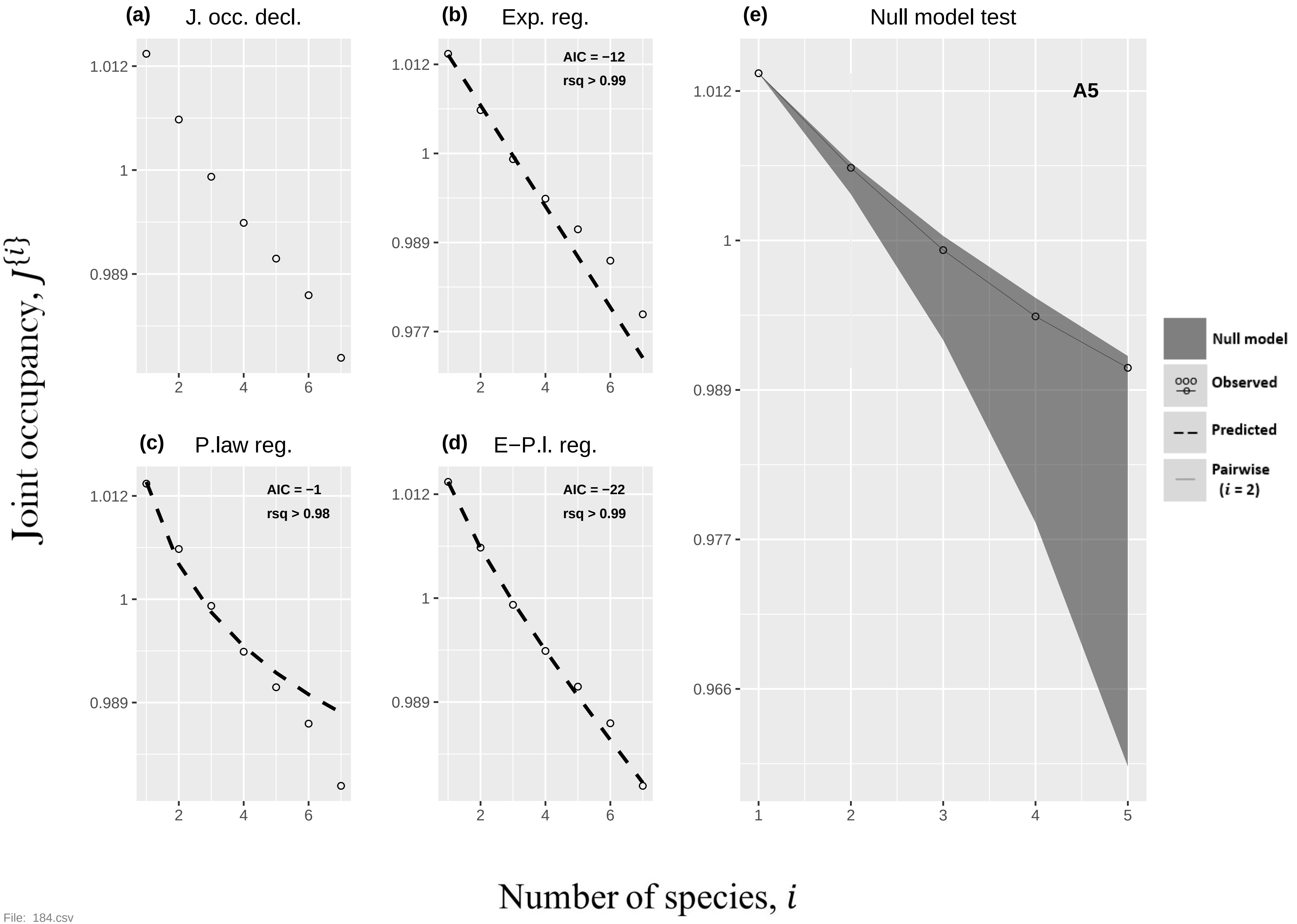
(a) J. occ. decl.
(b) Exp. reg.
AIC = −12
rsq > 0.99
(c) P.law reg.
AIC = −1
rsq > 0.98
(d) E−P.l. reg.
AIC = −22
rsq > 0.99
(e) Null model test
A5
Null model
Observed
Predicted
Pairwise (i = 2)
1.012
1
0.989
0.977
0.966
2
4
6
1
2
3
4
5
Joint occupancy, J{i}
Number of species, i

File: 184.csv

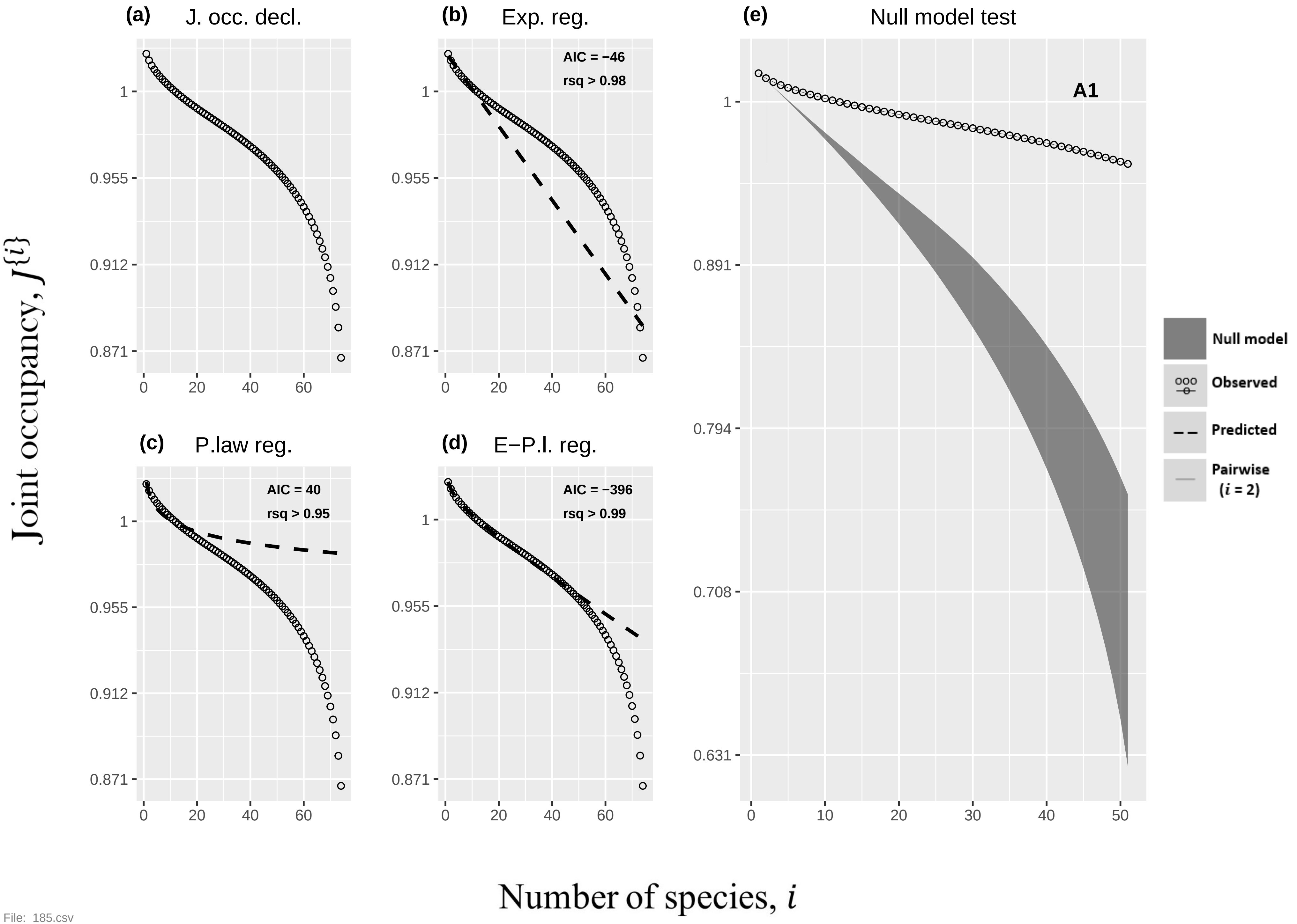

(a)
J. occ. decl.
(b)
Exp. reg.
AIC = −46
rsq > 0.98
(c)
P.law reg.
AIC = 40
rsq > 0.95
(d)
E−P.l. reg.
AIC = −396
rsq > 0.99
(e)
Null model test
A1
Null model
Observed
Predicted
Pairwise
(i = 2)
Joint occupancy, J{i}
Number of species, i
1
0.955
0.912
0.871
0.891
0.794
0.708
0.631
0
10
20
30
40
50
60


File: 185.csv

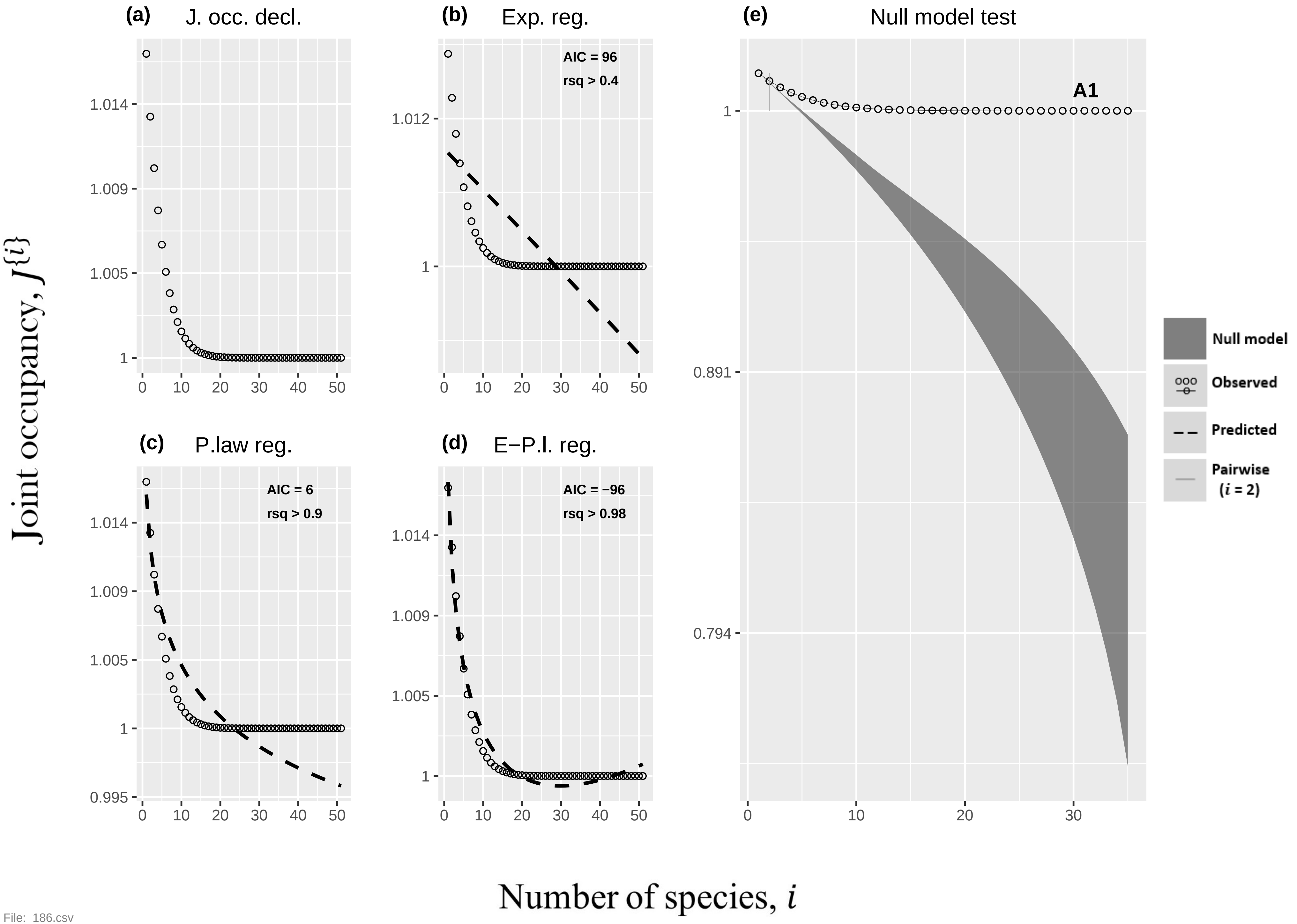

(a) J. occ. decl.
(b) Exp. reg.
AIC = 96
rsq > 0.4
(e) Null model test
A1
(c) P.law reg.
AIC = 6
rsq > 0.9
(d) E−P.l. reg.
AIC = −96
rsq > 0.98
Null model
Observed
Predicted
Pairwise (i = 2)
Joint occupancy, J{i}
Number of species, i


File: 186.csv

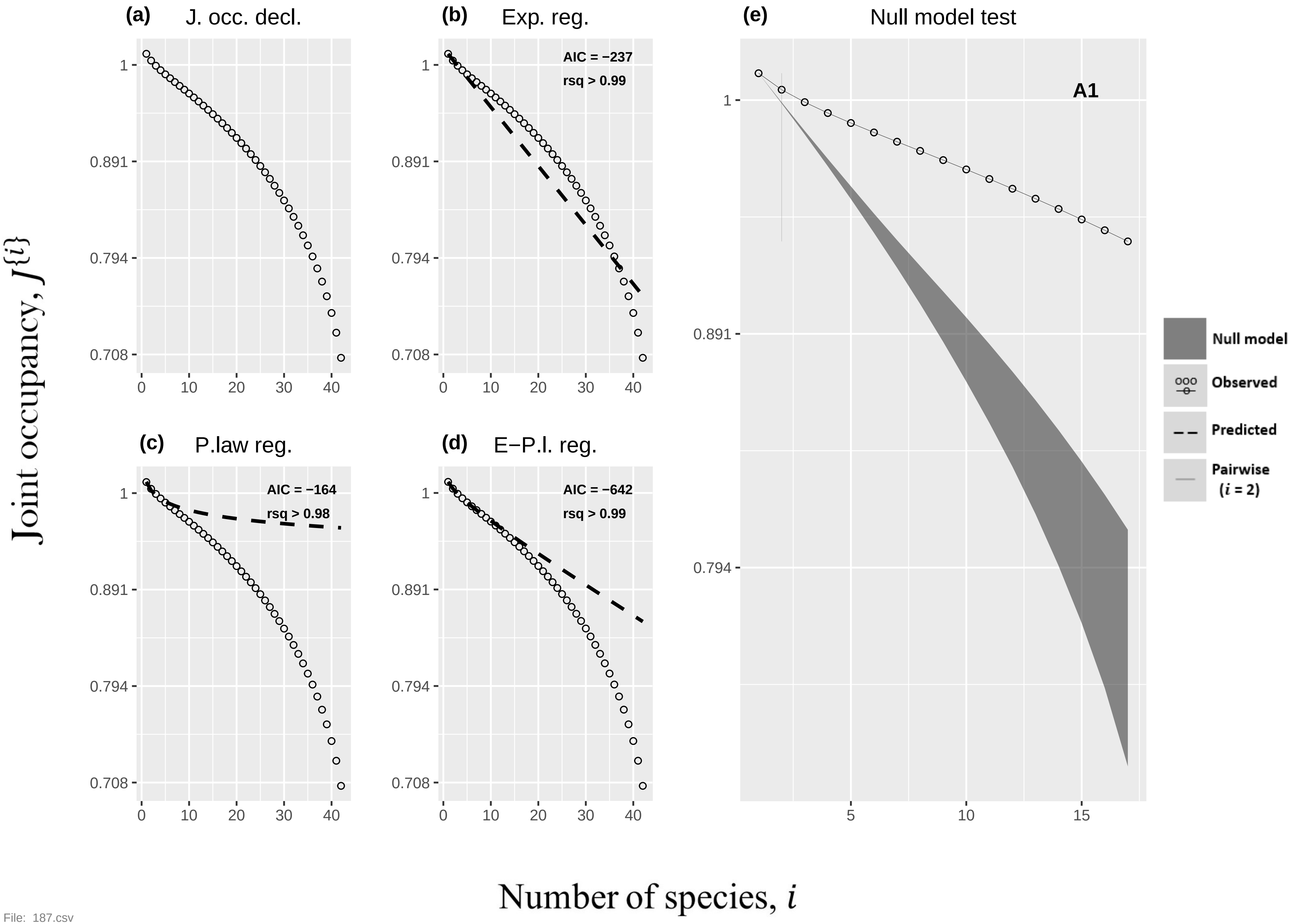
(a) J. occ. decl.
(b) Exp. reg.
AIC = −237
rsq > 0.99
(c) P.law reg.
AIC = −164
rsq > 0.98
(d) E−P.l. reg.
AIC = −642
rsq > 0.99
(e) Null model test
A1
Null model
Observed
Predicted
Pairwise ($i$ = 2)
1
0.891
0.794
0.708
0 10 20 30 40
5 10 15
Joint occupancy, $J^{\{i\}}$
Number of species, $i$

File: 187.csv

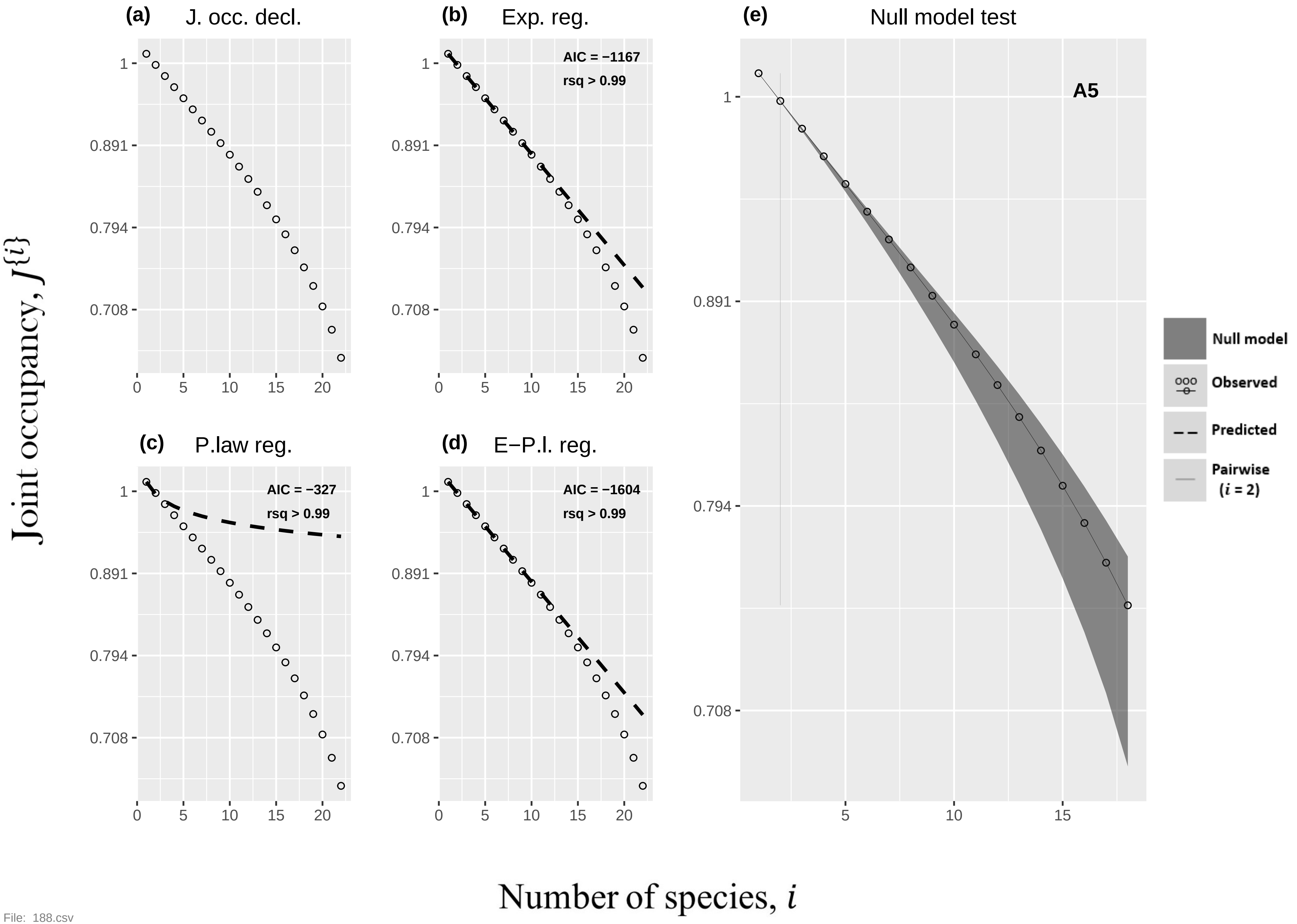
(a) J. occ. decl.
(b) Exp. reg.
AIC = −1167
rsq > 0.99
(c) P.law reg.
AIC = −327
rsq > 0.99
(d) E−P.l. reg.
AIC = −1604
rsq > 0.99
(e) Null model test
A5
Null model
Observed
Predicted
Pairwise (i = 2)
1
0.891
0.794
0.708
0
5
10
15
20
Joint occupancy, J{i}
Number of species, i

File: 188.csv

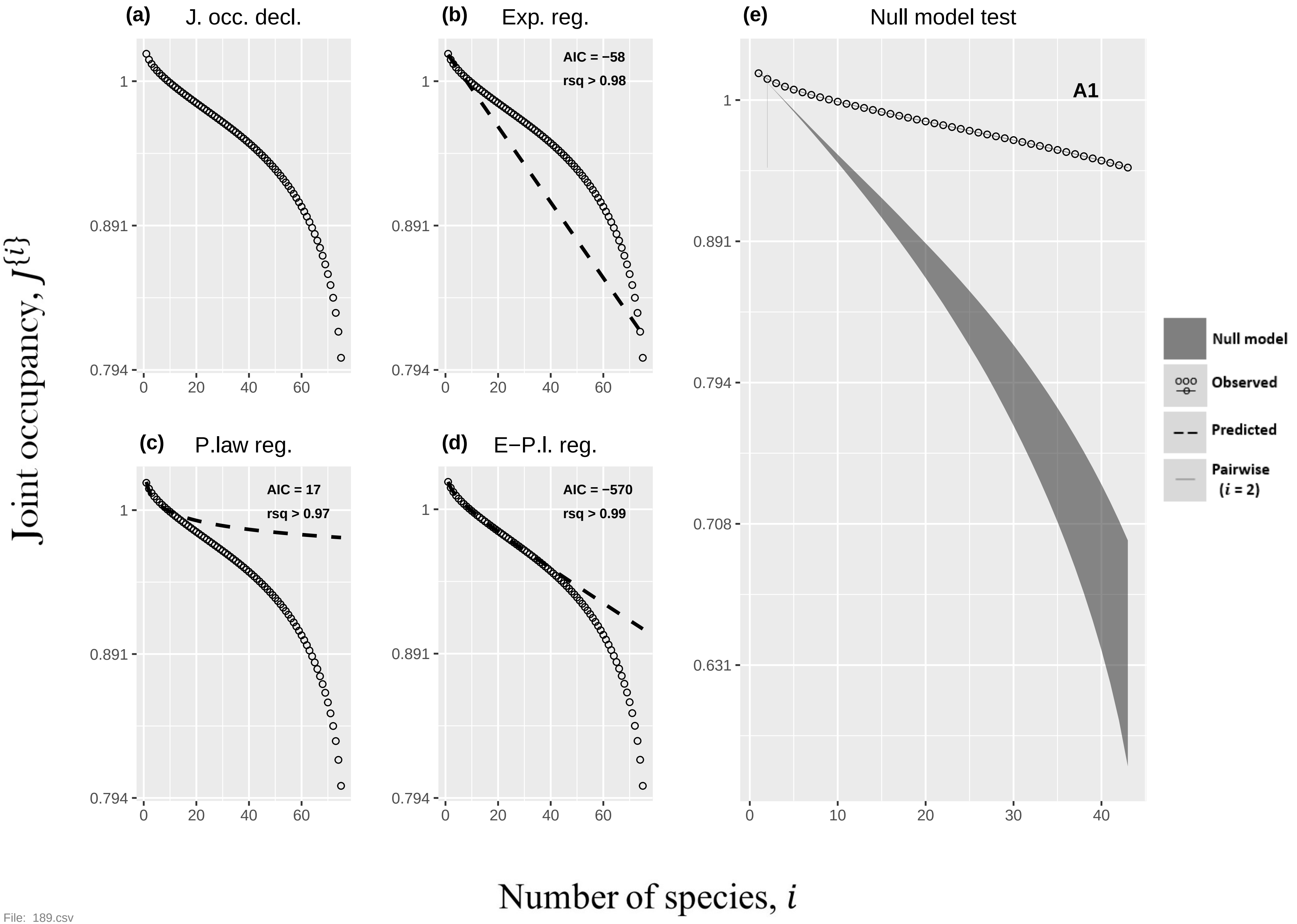

(a)
J. occ. decl.
(b)
Exp. reg.
AIC = −58
rsq > 0.98
(c)
P.law reg.
AIC = 17
rsq > 0.97
(d)
E−P.l. reg.
AIC = −570
rsq > 0.99
(e)
Null model test
A1
Null model
Observed
Predicted
Pairwise
(i = 2)
Joint occupancy, J{i}
Number of species, i
1
0.891
0.794
0.708
0.631
0
10
20
30
40
60


File: 189.csv

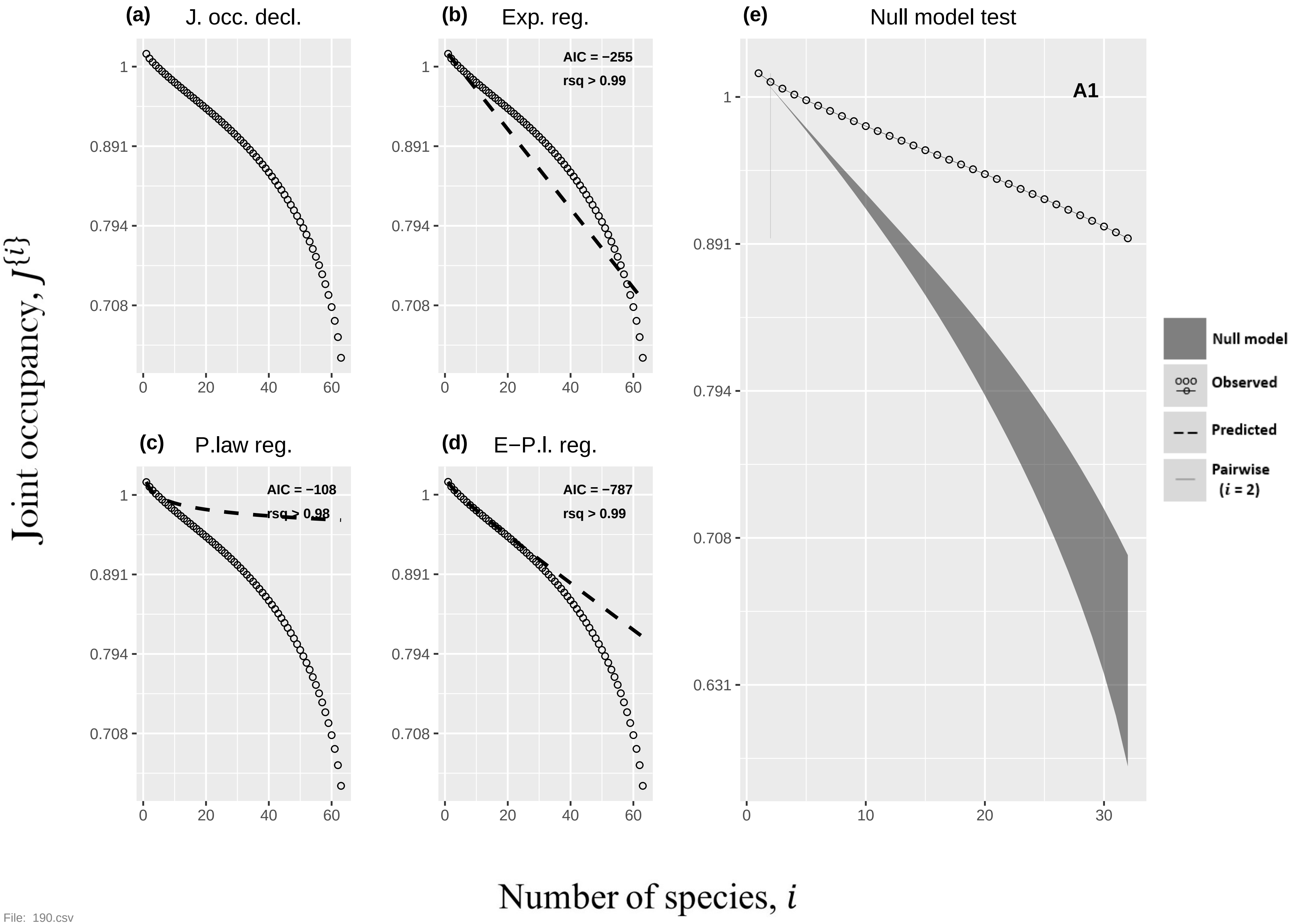
(a) J. occ. decl.
(b) Exp. reg.
AIC = −255
rsq > 0.99
(c) P.law reg.
AIC = −108
rsq > 0.98
(d) E−P.l. reg.
AIC = −787
rsq > 0.99
(e) Null model test
A1
Null model
Observed
Predicted
Pairwise (i = 2)
1
0.891
0.794
0.708
0.631
0
20
40
60
10
30
Joint occupancy, J{i}
Number of species, i

File: 190.csv

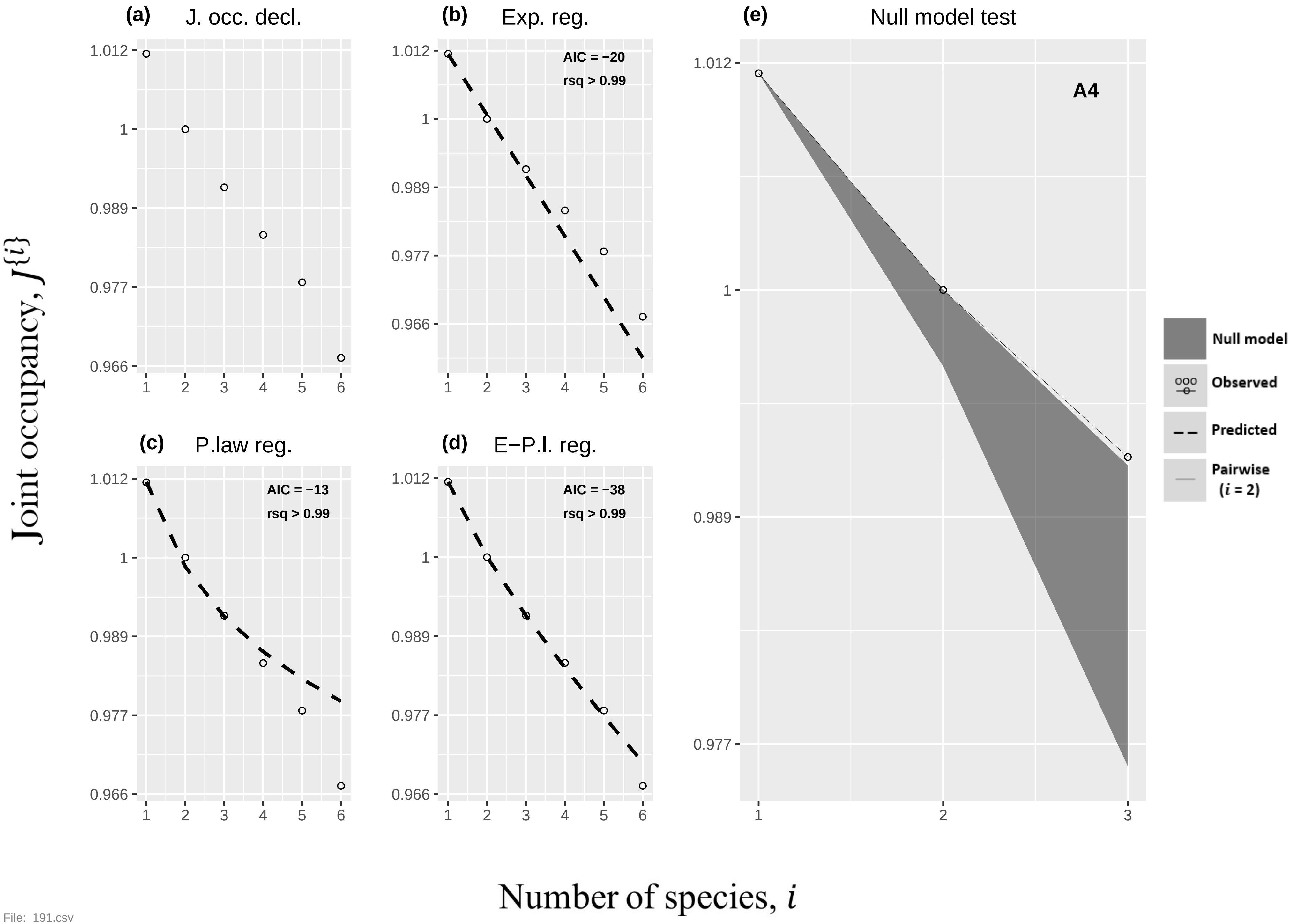

(a) J. occ. decl.
(b) Exp. reg.
AIC = −20
rsq > 0.99
(c) P.law reg.
AIC = −13
rsq > 0.99
(d) E−P.l. reg.
AIC = −38
rsq > 0.99
(e) Null model test
A4
Null model
Observed
Predicted
Pairwise ($i$ = 2)
1.012
1
0.989
0.977
0.966
Joint occupancy, $J\{i\}$
Number of species, $i$


File: 191.csv

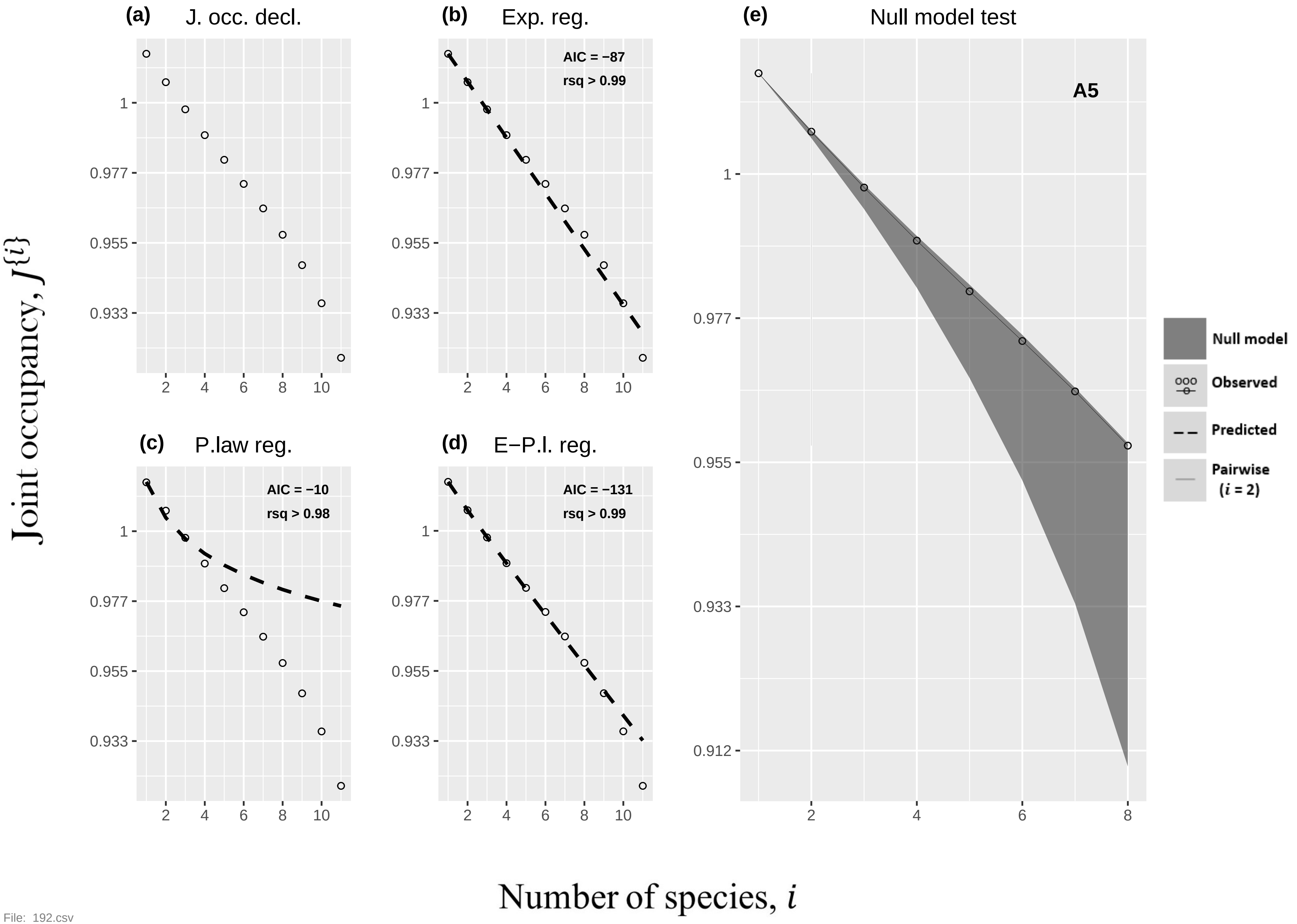

(a) J. occ. decl.
(b) Exp. reg.
AIC = −87
rsq > 0.99
(c) P.law reg.
AIC = −10
rsq > 0.98
(d) E−P.l. reg.
AIC = −131
rsq > 0.99
(e) Null model test
A5
Null model
Observed
Predicted
Pairwise (i = 2)
Joint occupancy, J{i}
Number of species, i


File: 192.csv

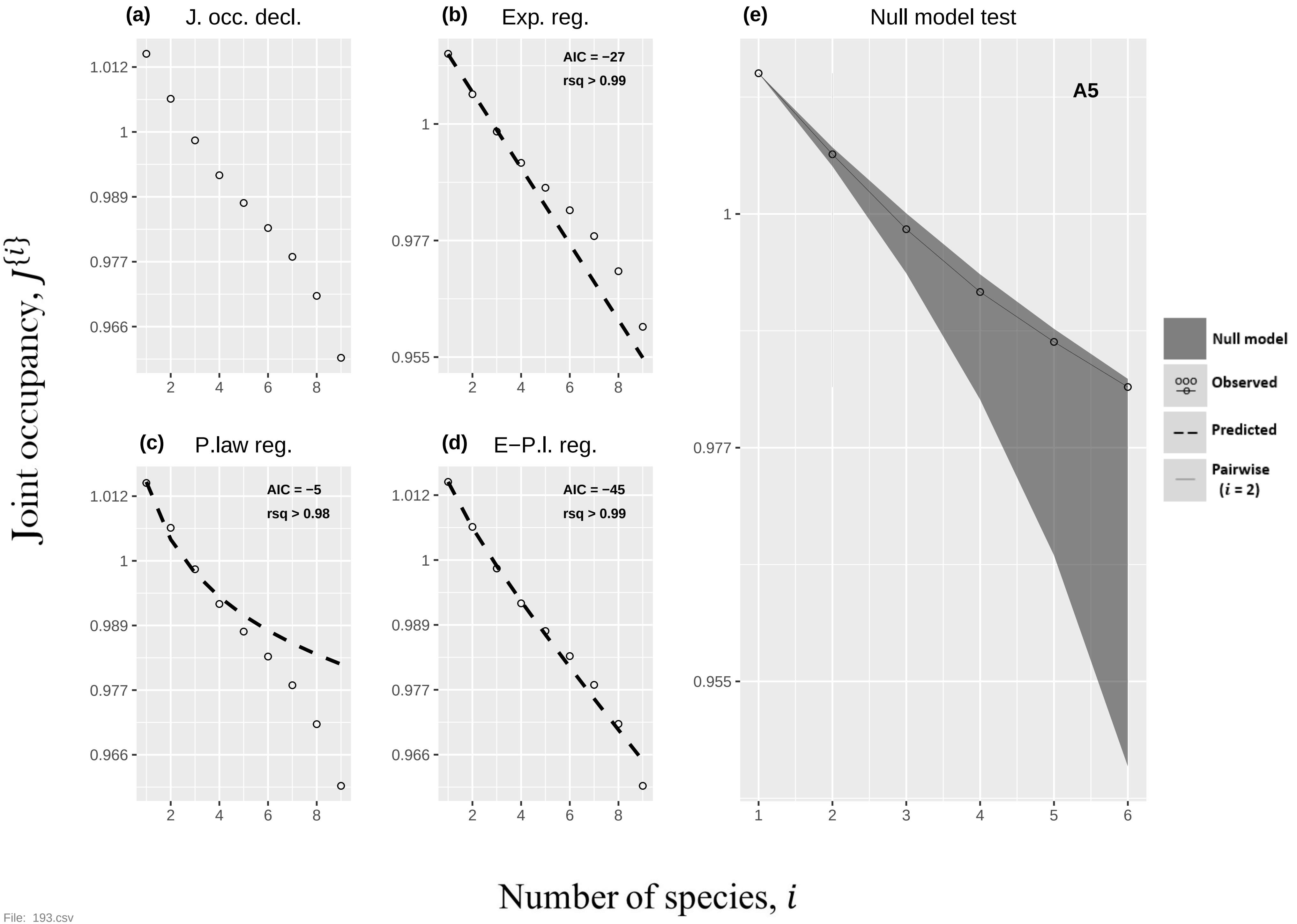
(a) J. occ. decl.
(b) Exp. reg.
AIC = −27
rsq > 0.99
(e) Null model test
A5
(c) P.law reg.
AIC = −5
rsq > 0.98
(d) E−P.l. reg.
AIC = −45
rsq > 0.99
Null model
Observed
Predicted
Pairwise
(i = 2)
Joint occupancy, J{i}
Number of species, i

File: 193.csv

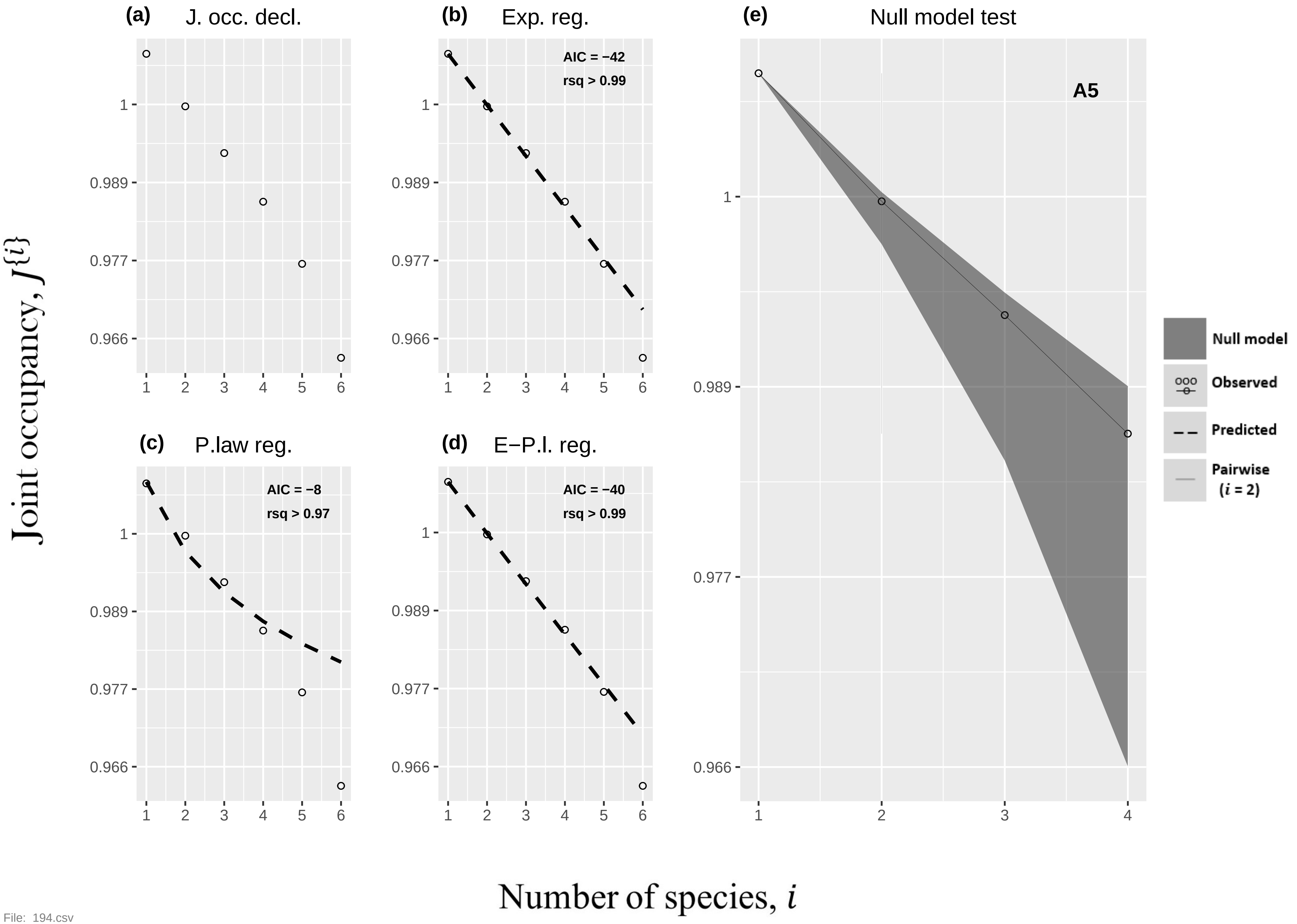
(a)
J. occ. decl.
(b)
Exp. reg.
AIC = −42
rsq > 0.99
(c)
P.law reg.
AIC = −8
rsq > 0.97
(d)
E−P.l. reg.
AIC = −40
rsq > 0.99
(e)
Null model test
A5
Null model
Observed
Predicted
Pairwise (i = 2)
Joint occupancy, J{i}
Number of species, i
1
0.989
0.977
0.966
1 2 3 4 5 6
1 2 3 4
File: 194.csv

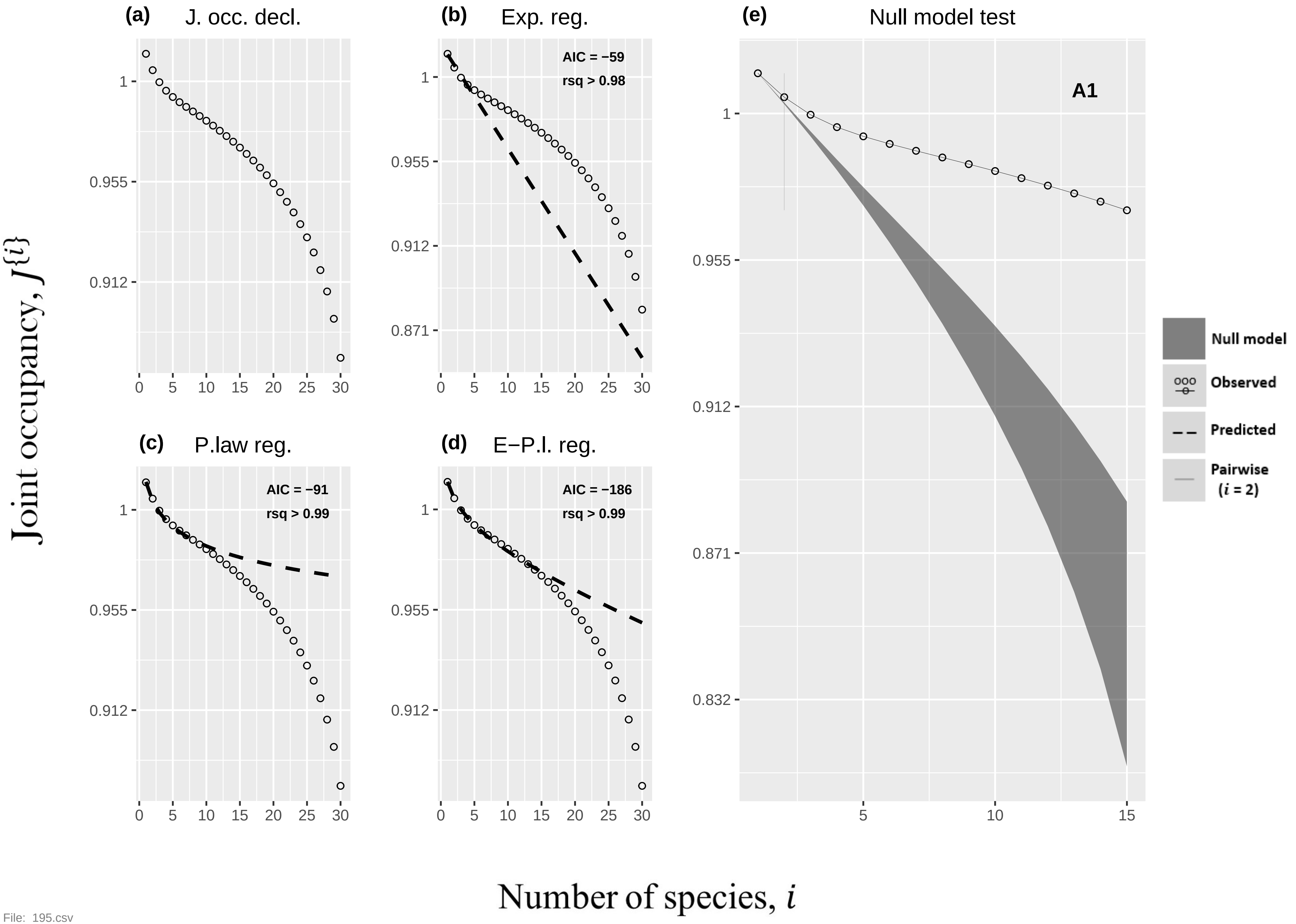

(a) J. occ. decl.
(b) Exp. reg.
AIC = −59
rsq > 0.98
(c) P.law reg.
AIC = −91
rsq > 0.99
(d) E−P.l. reg.
AIC = −186
rsq > 0.99
(e) Null model test
A1
Null model
Observed
Predicted
Pairwise (i = 2)
Joint occupancy, J{i}
Number of species, i
1
0.955
0.912
0.871
0.832
0
5
10
15
20
25
30


File: 195.csv

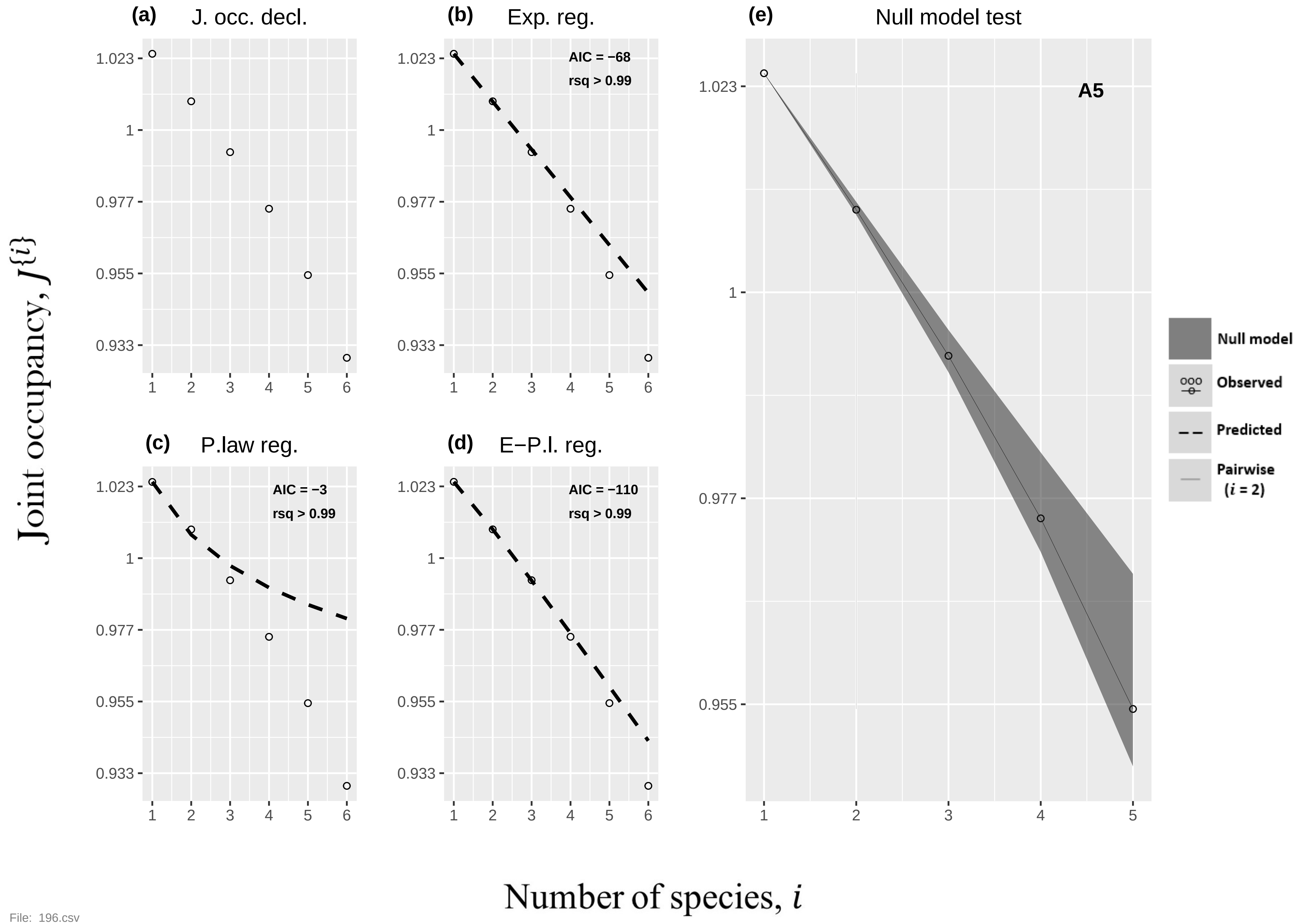
(a)
J. occ. decl.
(b)
Exp. reg.
AIC = −68
rsq > 0.99
(c)
P.law reg.
AIC = −3
rsq > 0.99
(d)
E−P.l. reg.
AIC = −110
rsq > 0.99
(e)
Null model test
A5
Null model
Observed
Predicted
Pairwise (i = 2)
Joint occupancy, J{i}
Number of species, i
1.023
1
0.977
0.955
0.933
1
2
3
4
5
6

File: 196.csv

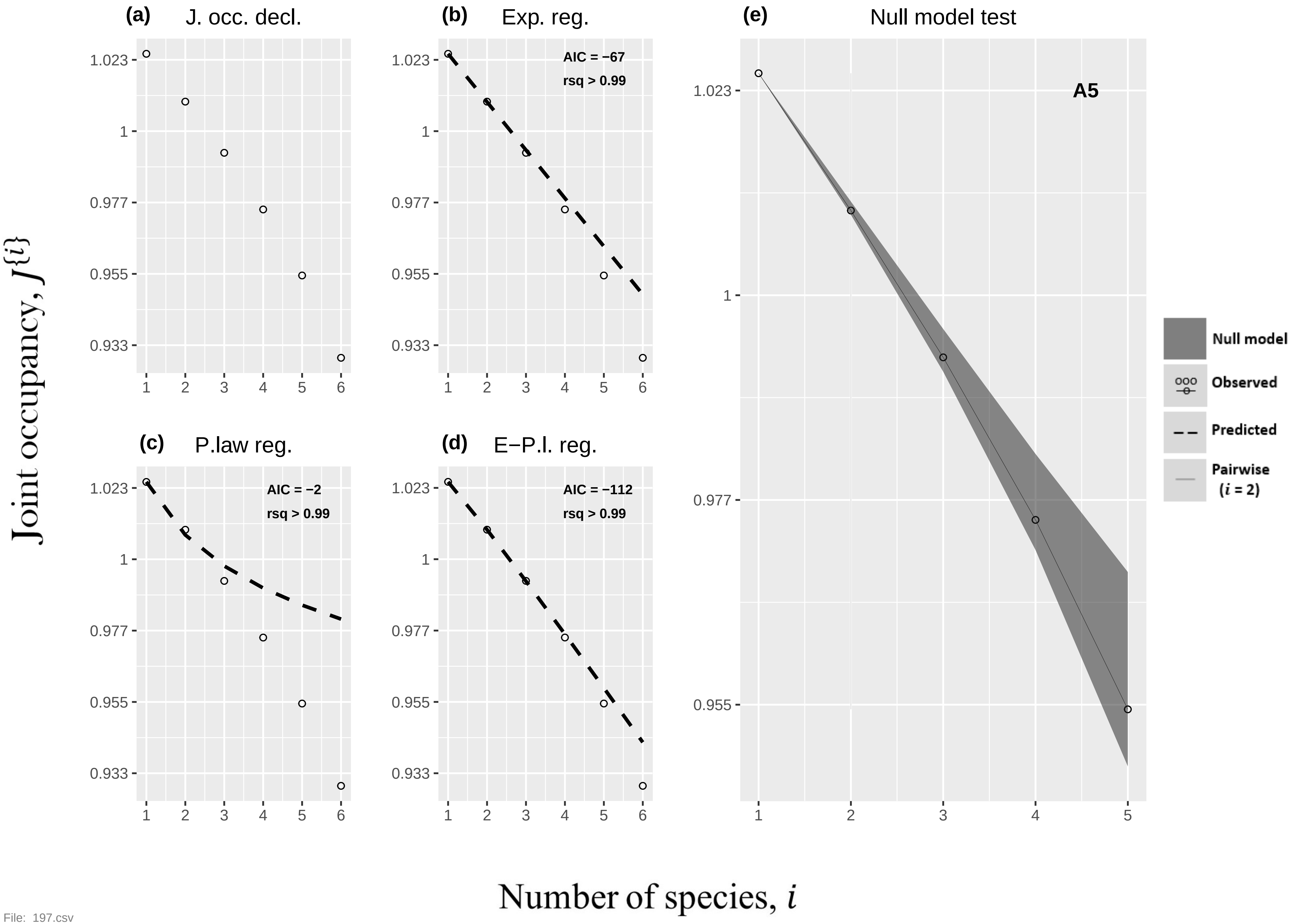

(a) J. occ. decl.
(b) Exp. reg.
AIC = −67
rsq > 0.99
(c) P.law reg.
AIC = −2
rsq > 0.99
(d) E−P.l. reg.
AIC = −112
rsq > 0.99
(e) Null model test
A5
Null model
Observed
Predicted
Pairwise ($i$ = 2)
1.023
1
0.977
0.955
0.933
1 2 3 4 5 6
Joint occupancy, $J^{\{i\}}$
Number of species, $i$


File: 197.csv

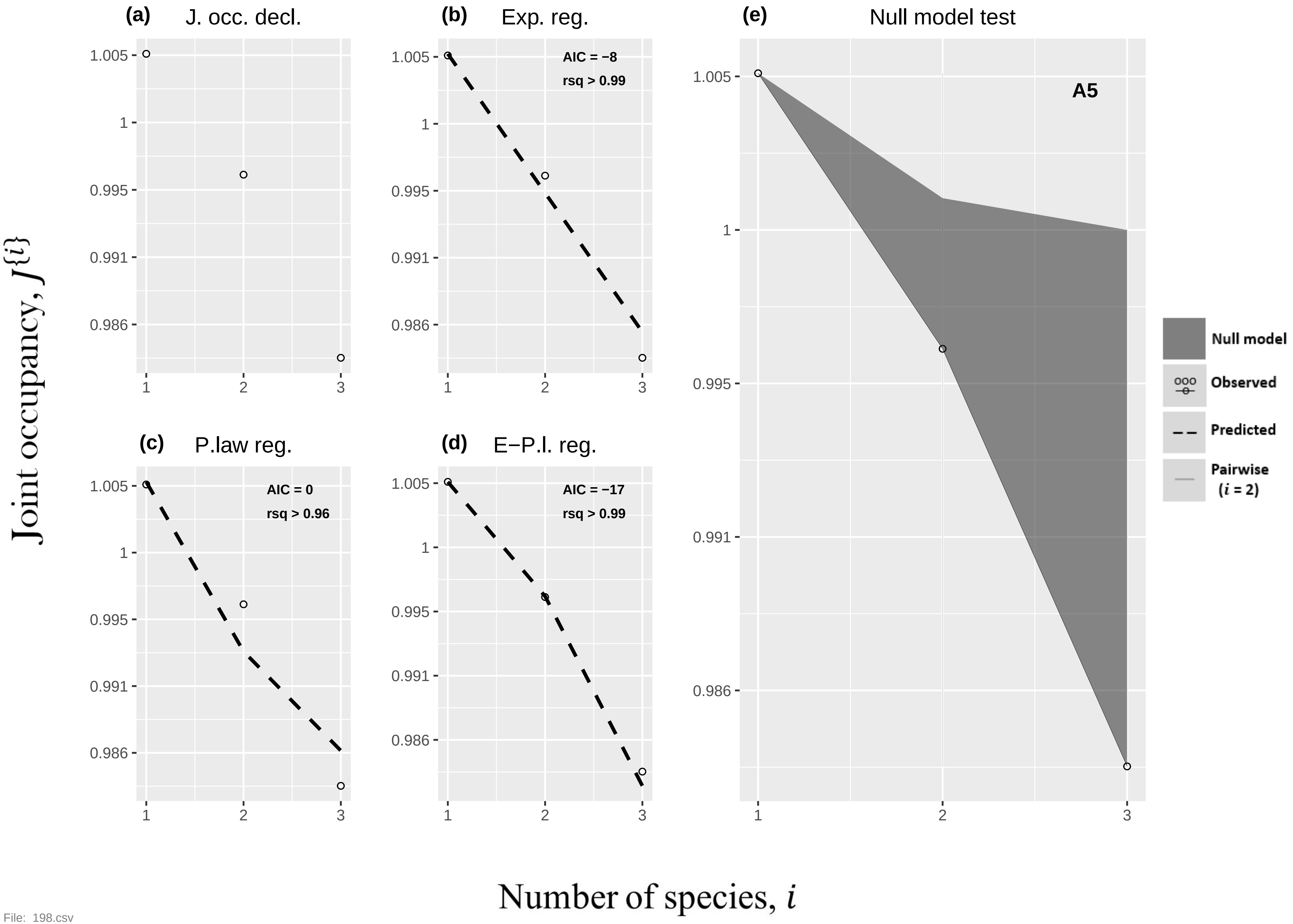

(a) J. occ. decl.
(b) Exp. reg.
AIC = −8
rsq > 0.99
(c) P.law reg.
AIC = 0
rsq > 0.96
(d) E−P.l. reg.
AIC = −17
rsq > 0.99
(e) Null model test
A5
Null model
Observed
Predicted
Pairwise (i = 2)
1.005
1
0.995
0.991
0.986
1
2
3
Joint occupancy, J{i}
Number of species, i


File: 198.csv

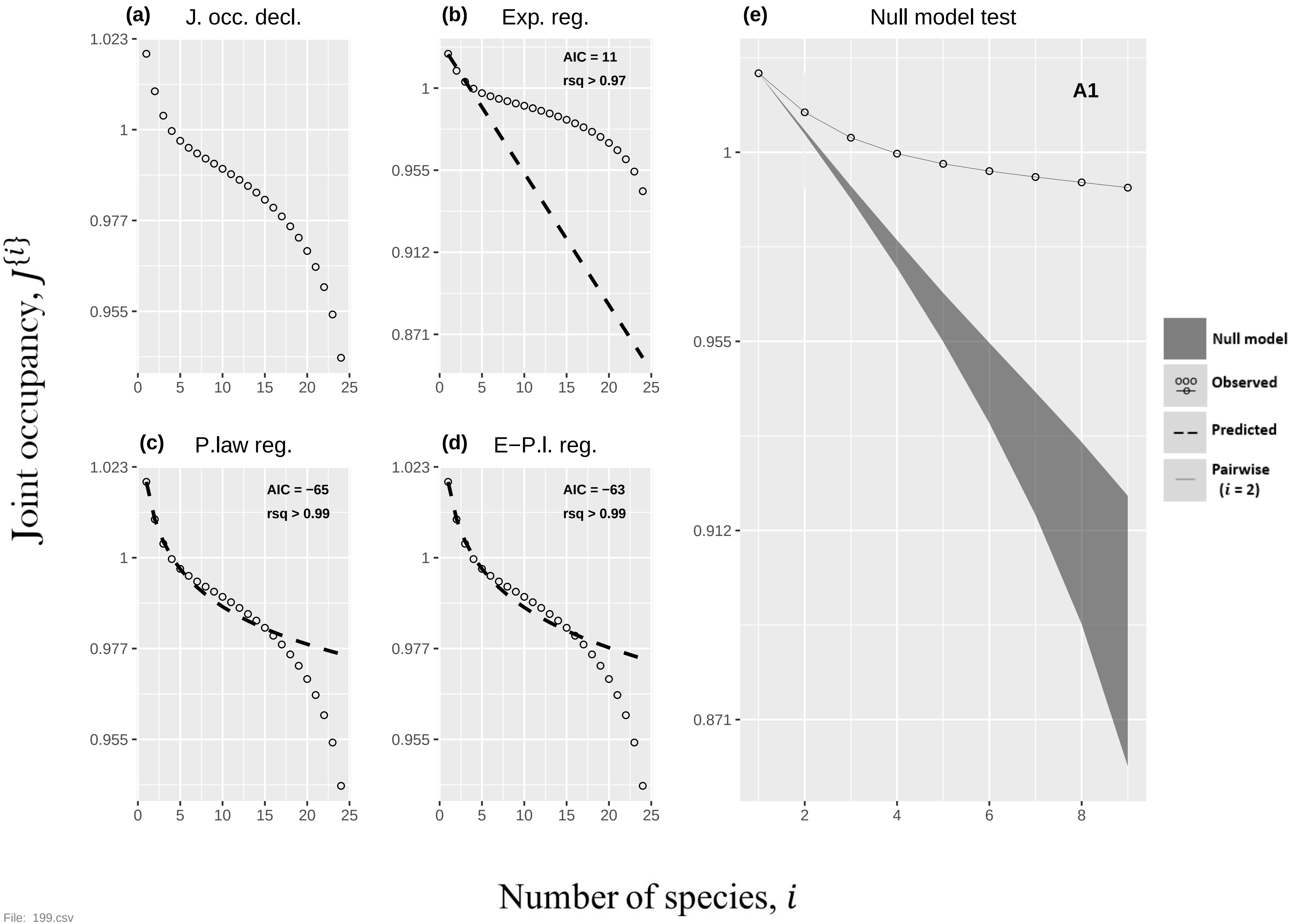
(a) J. occ. decl.
(b) Exp. reg.
AIC = 11
rsq > 0.97
(c) P.law reg.
AIC = −65
rsq > 0.99
(d) E−P.l. reg.
AIC = −63
rsq > 0.99
(e) Null model test
A1
Null model
Observed
Predicted
Pairwise ($i$ = 2)
1.023
1
0.977
0.955
0.912
0.871
0 5 10 15 20 25
2 4 6 8
Joint occupancy, $J^{\{i\}}$
Number of species, $i$

File: 199.csv

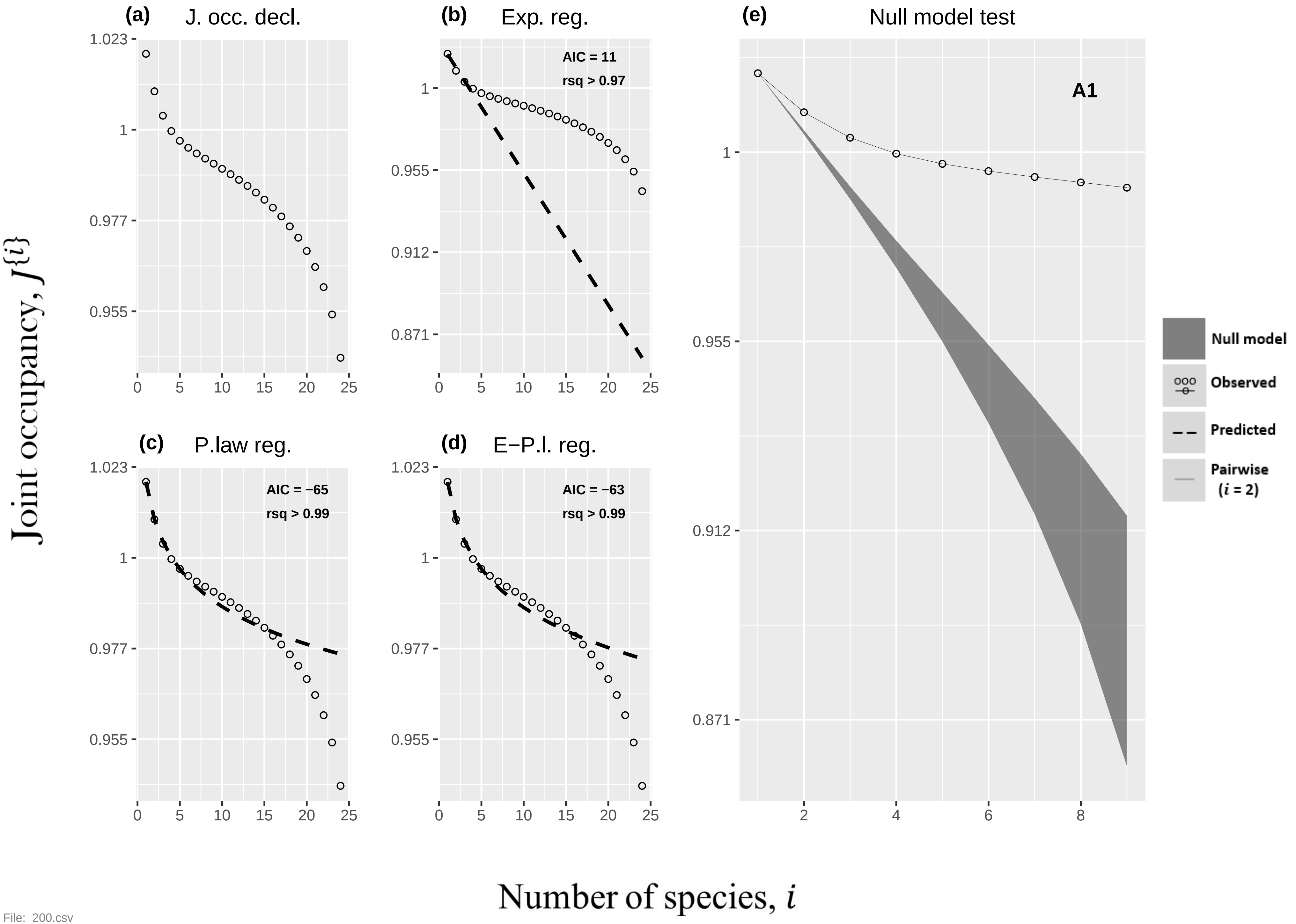

(a) J. occ. decl.
(b) Exp. reg.
AIC = 11
rsq > 0.97
(c) P.law reg.
AIC = −65
rsq > 0.99
(d) E−P.l. reg.
AIC = −63
rsq > 0.99
(e) Null model test
A1
Null model
Observed
Predicted
Pairwise
(i = 2)
Joint occupancy, J{i}
Number of species, i


File: 200.csv

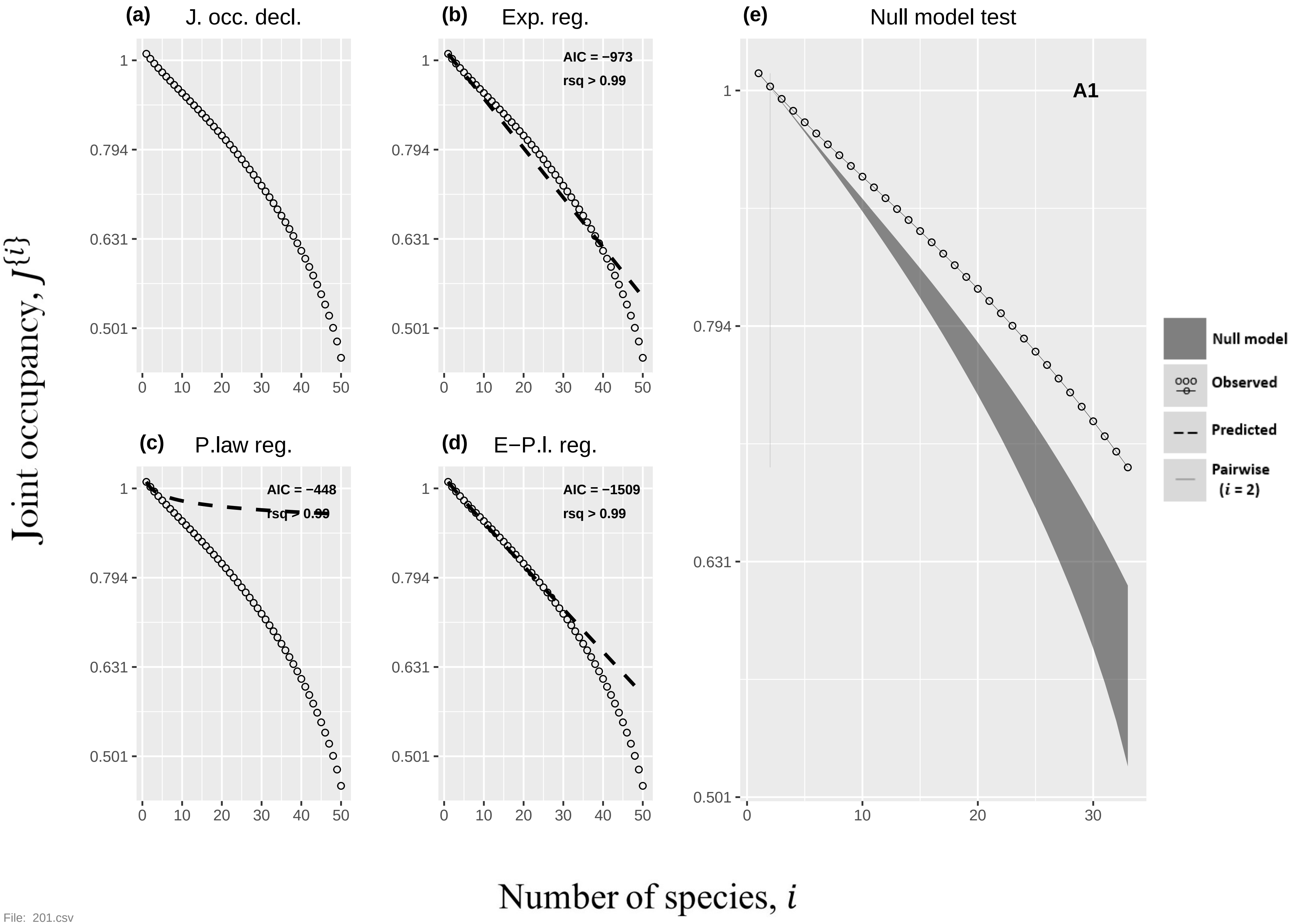

(a) J. occ. decl.
(b) Exp. reg.
AIC = −973
rsq > 0.99
(c) P.law reg.
AIC = −448
rsq > 0.99
(d) E−P.l. reg.
AIC = −1509
rsq > 0.99
(e) Null model test
A1
Null model
Observed
Predicted
Pairwise (i = 2)
Joint occupancy, J{i}
Number of species, i


File: 201.csv

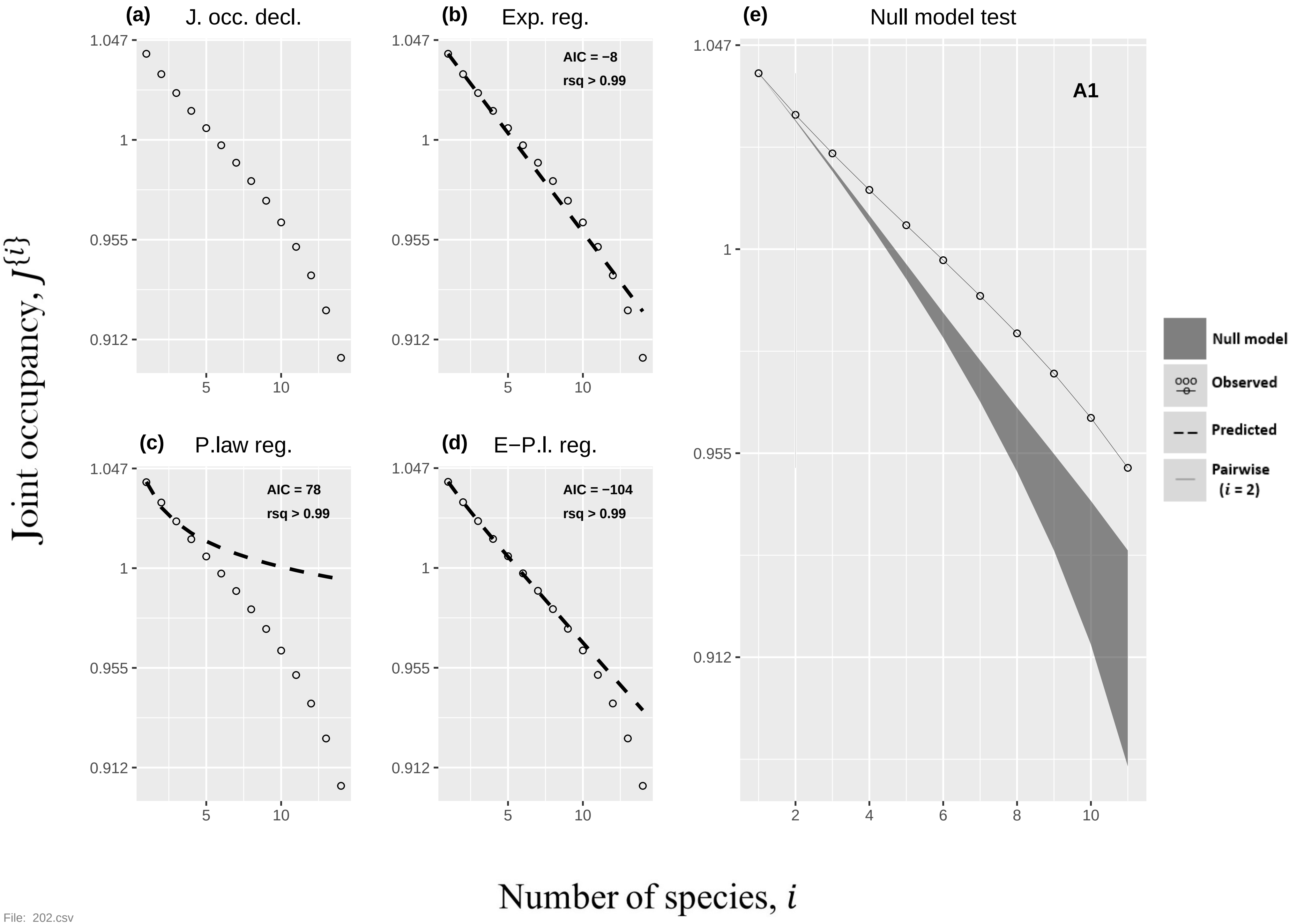
(a)
J. occ. decl.
(b)
Exp. reg.
AIC = −8
rsq > 0.99
(c)
P.law reg.
AIC = 78
rsq > 0.99
(d)
E−P.l. reg.
AIC = −104
rsq > 0.99
(e)
Null model test
A1
1.047
1
0.955
0.912
5
10
2
4
6
8
Null model
Observed
Predicted
Pairwise
(i = 2)
Joint occupancy, J{i}
Number of species, i
File: 202.csv

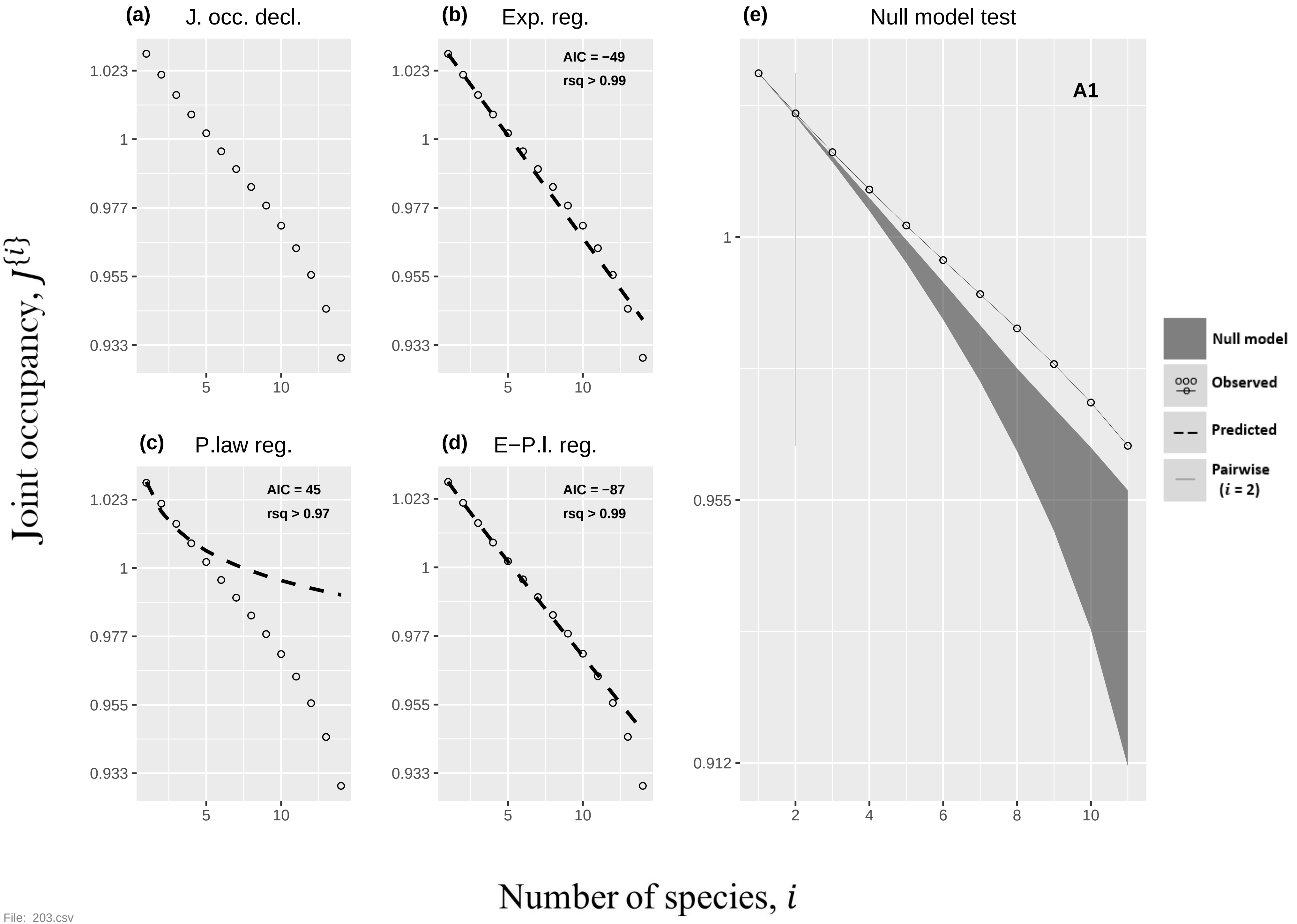
(a) J. occ. decl.
(b) Exp. reg.
AIC = −49
rsq > 0.99
(c) P.law reg.
AIC = 45
rsq > 0.97
(d) E−P.l. reg.
AIC = −87
rsq > 0.99
(e) Null model test
A1
Null model
Observed
Predicted
Pairwise (i = 2)
1.023
1
0.977
0.955
0.933
0.912
5
10
2
4
6
8
Joint occupancy, J{i}
Number of species, i

File: 203.csv

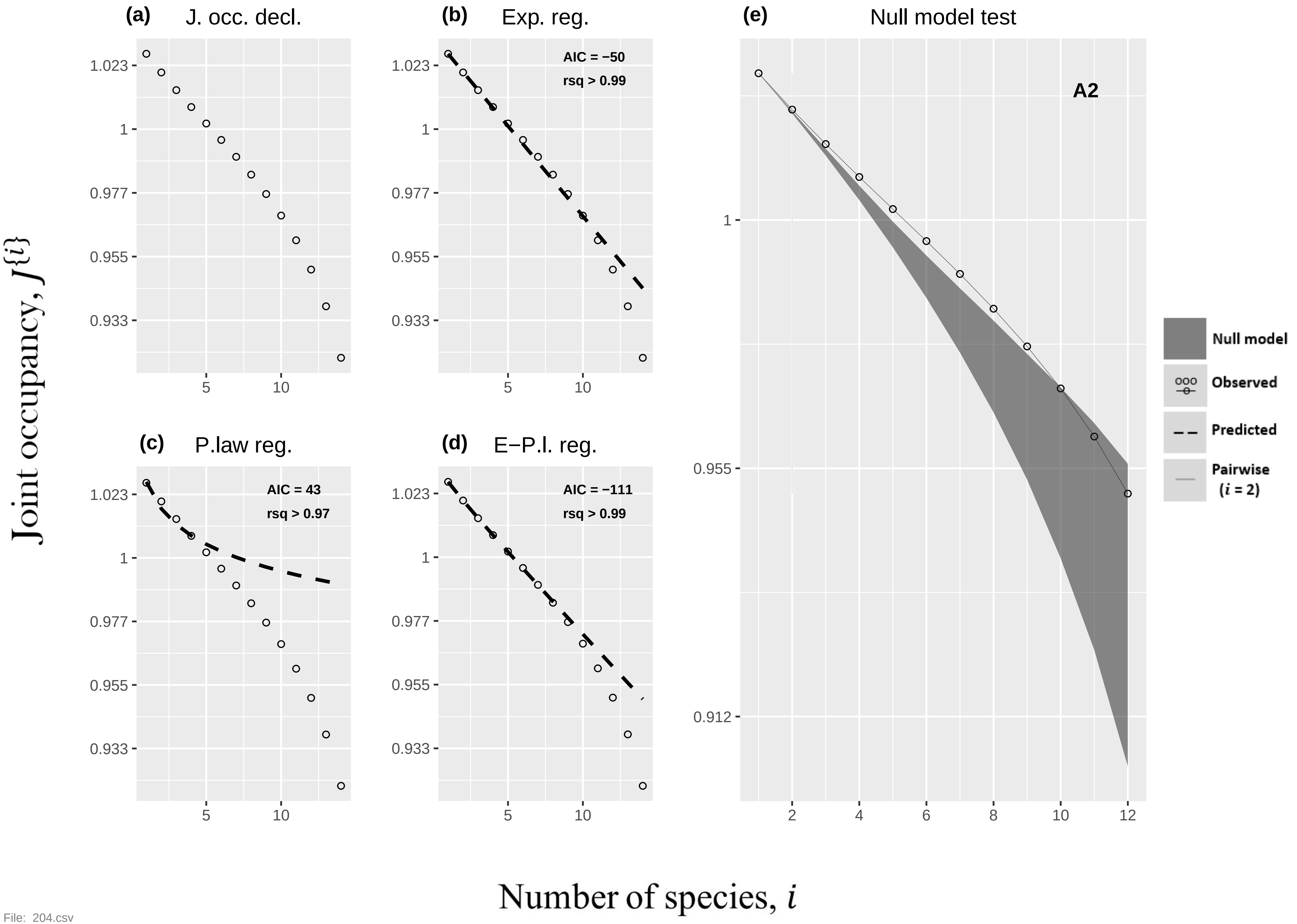

(a) J. occ. decl.
(b) Exp. reg.
AIC = −50
rsq > 0.99
(c) P.law reg.
AIC = 43
rsq > 0.97
(d) E−P.l. reg.
AIC = −111
rsq > 0.99
(e) Null model test
A2
Null model
Observed
Predicted
Pairwise (i = 2)
1.023
1
0.977
0.955
0.933
0.912
5
10
2
4
6
8
12
Joint occupancy, J{i}
Number of species, i


File: 204.csv

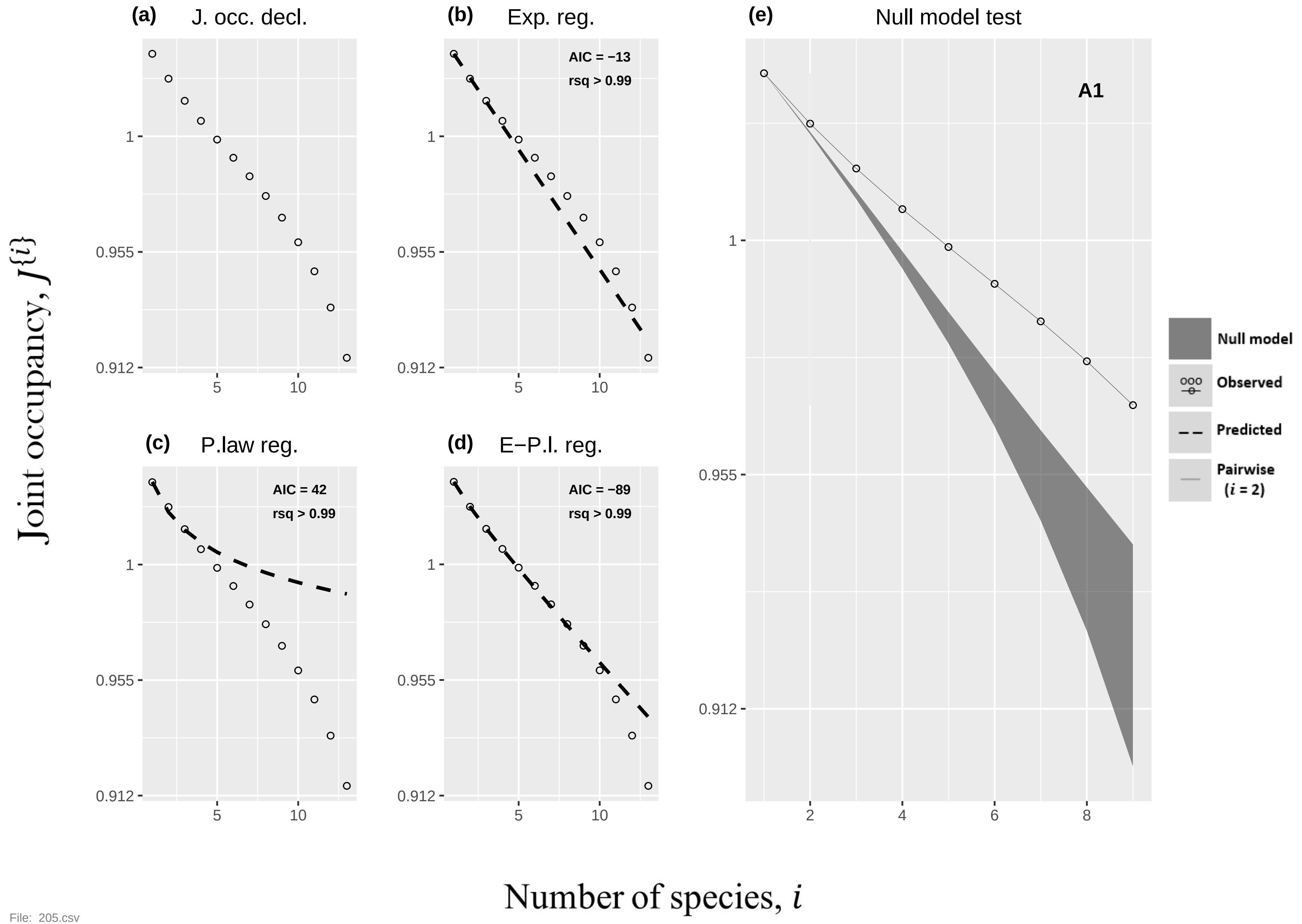

(a)
J. occ. decl.
(b)
Exp. reg.
AIC = −13
rsq > 0.99
(c)
P.law reg.
AIC = 42
rsq > 0.99
(d)
E−P.l. reg.
AIC = −89
rsq > 0.99
(e)
Null model test
A1
Null model
Observed
Predicted
Pairwise
(i = 2)
1
0.955
0.912
5
10
2
4
6
8
Joint occupancy, J{i}
Number of species, i


File: 205.csv

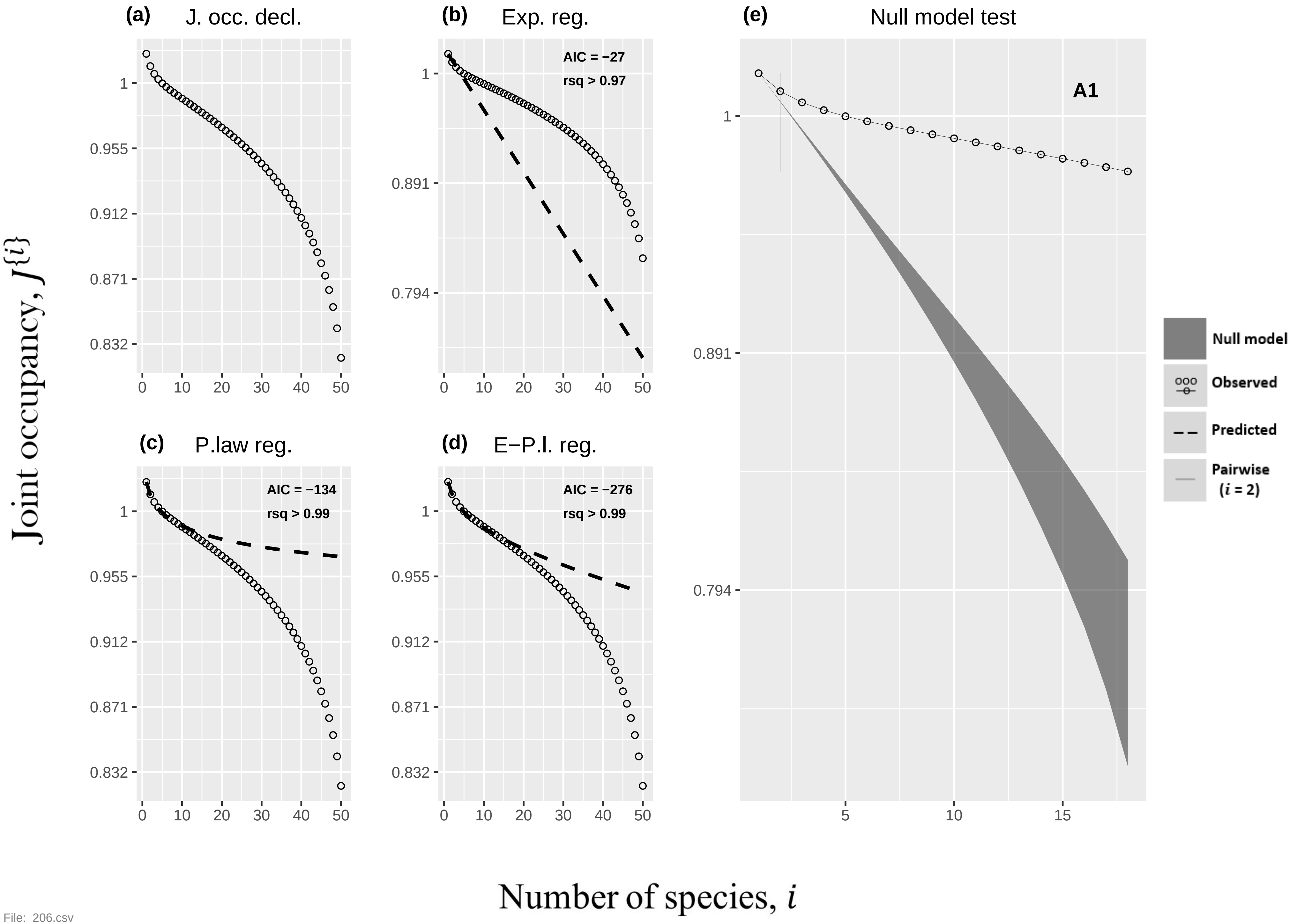
(a) J. occ. decl.
(b) Exp. reg.
AIC = −27
rsq > 0.97
(c) P.law reg.
AIC = −134
rsq > 0.99
(d) E−P.l. reg.
AIC = −276
rsq > 0.99
(e) Null model test
A1
Null model
Observed
Predicted
Pairwise ($i$ = 2)
Joint occupancy, $J\{i\}$
Number of species, $i$

File: 206.csv

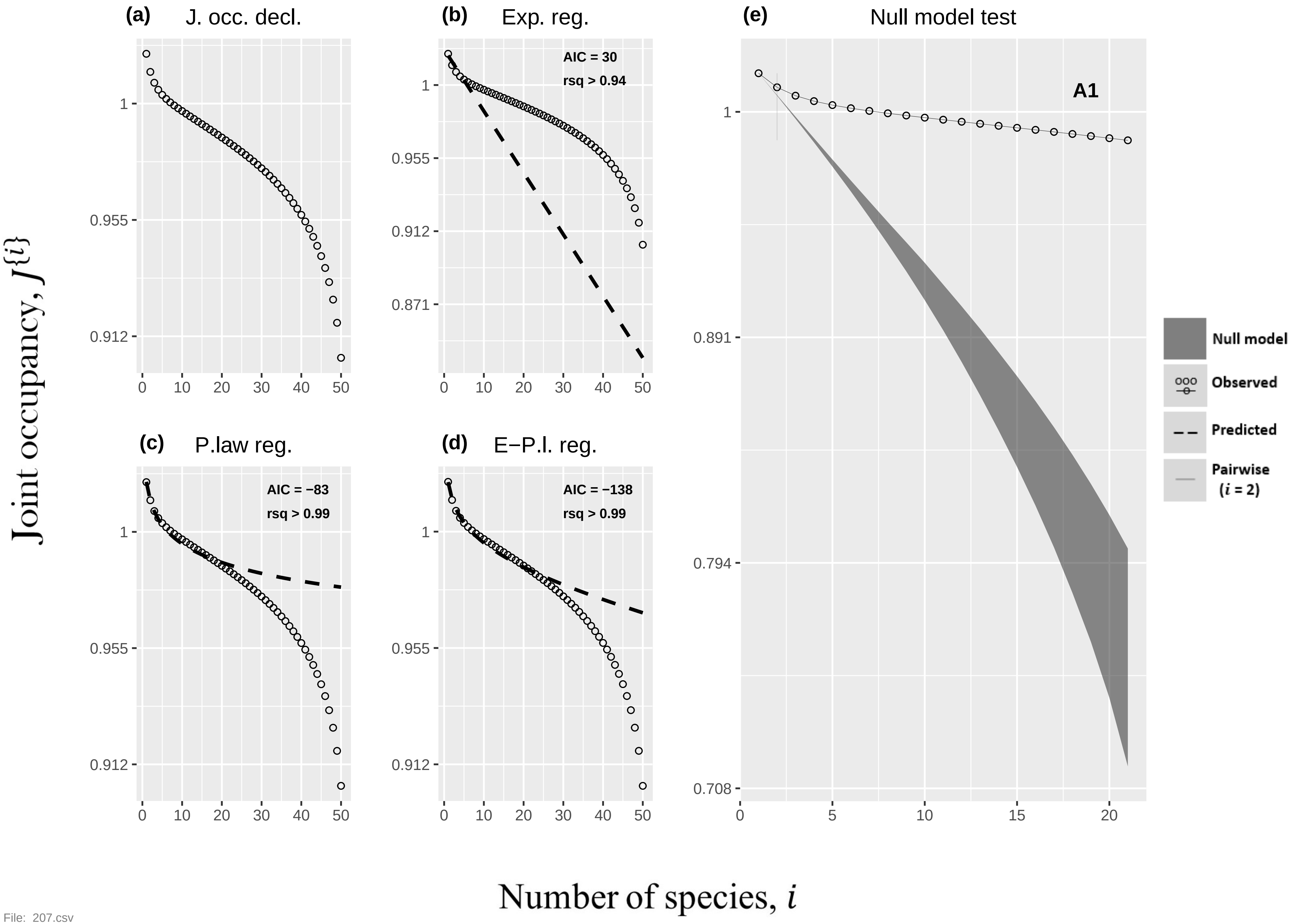

(a) J. occ. decl.
(b) Exp. reg.
AIC = 30
rsq > 0.94
(c) P.law reg.
AIC = −83
rsq > 0.99
(d) E−P.l. reg.
AIC = −138
rsq > 0.99
(e) Null model test
A1
Null model
Observed
Predicted
Pairwise (i = 2)
1
0.955
0.912
0.871
0.891
0.794
0.708
0
5
10
15
20
30
40
50
Joint occupancy, J{i}
Number of species, i


File: 207.csv

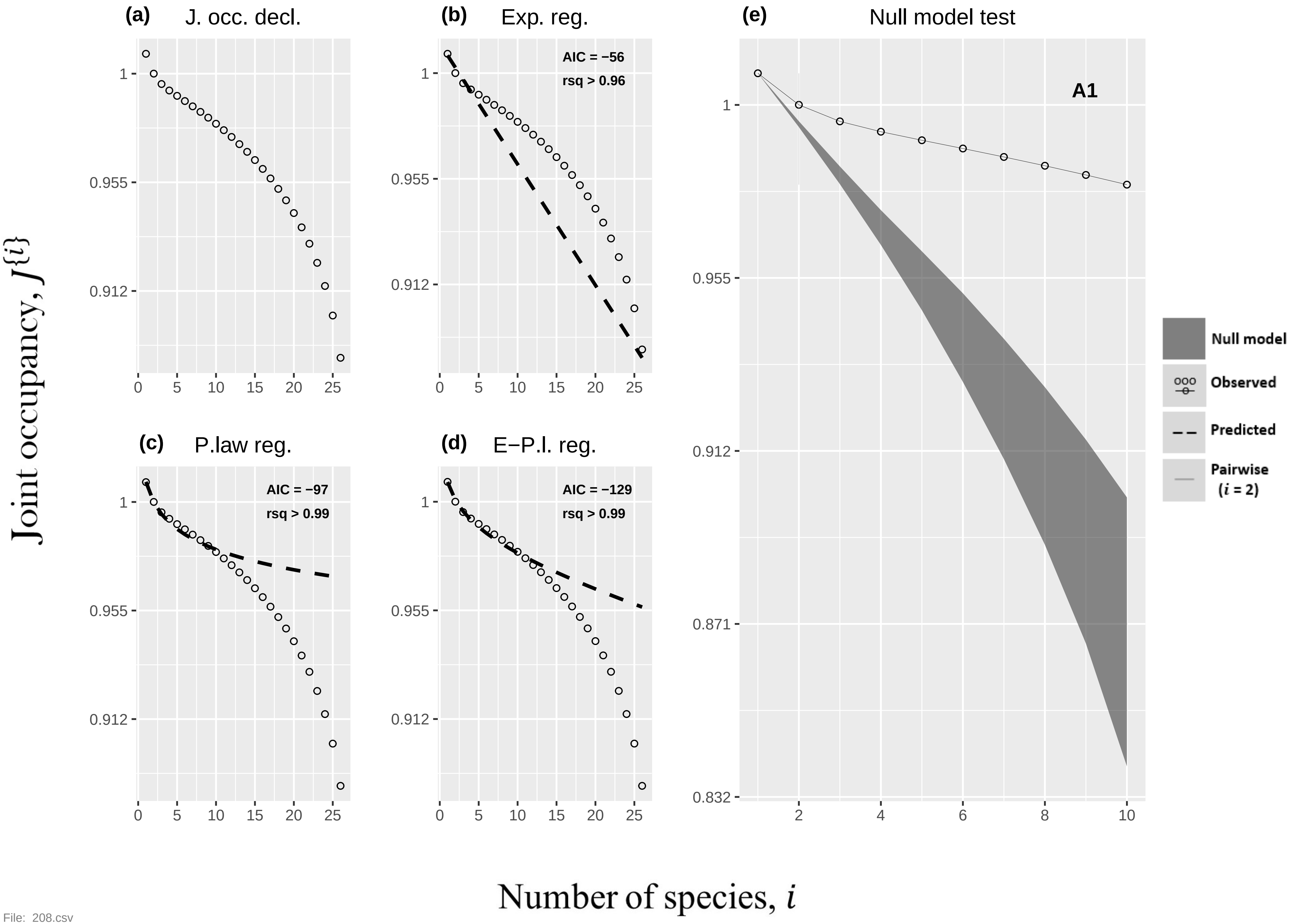

(a) J. occ. decl.
(b) Exp. reg.
AIC = −56
rsq > 0.96
(c) P.law reg.
AIC = −97
rsq > 0.99
(d) E−P.l. reg.
AIC = −129
rsq > 0.99
(e) Null model test
A1
Null model
Observed
Predicted
Pairwise (i = 2)
Joint occupancy, J{i}
Number of species, i


File: 208.csv

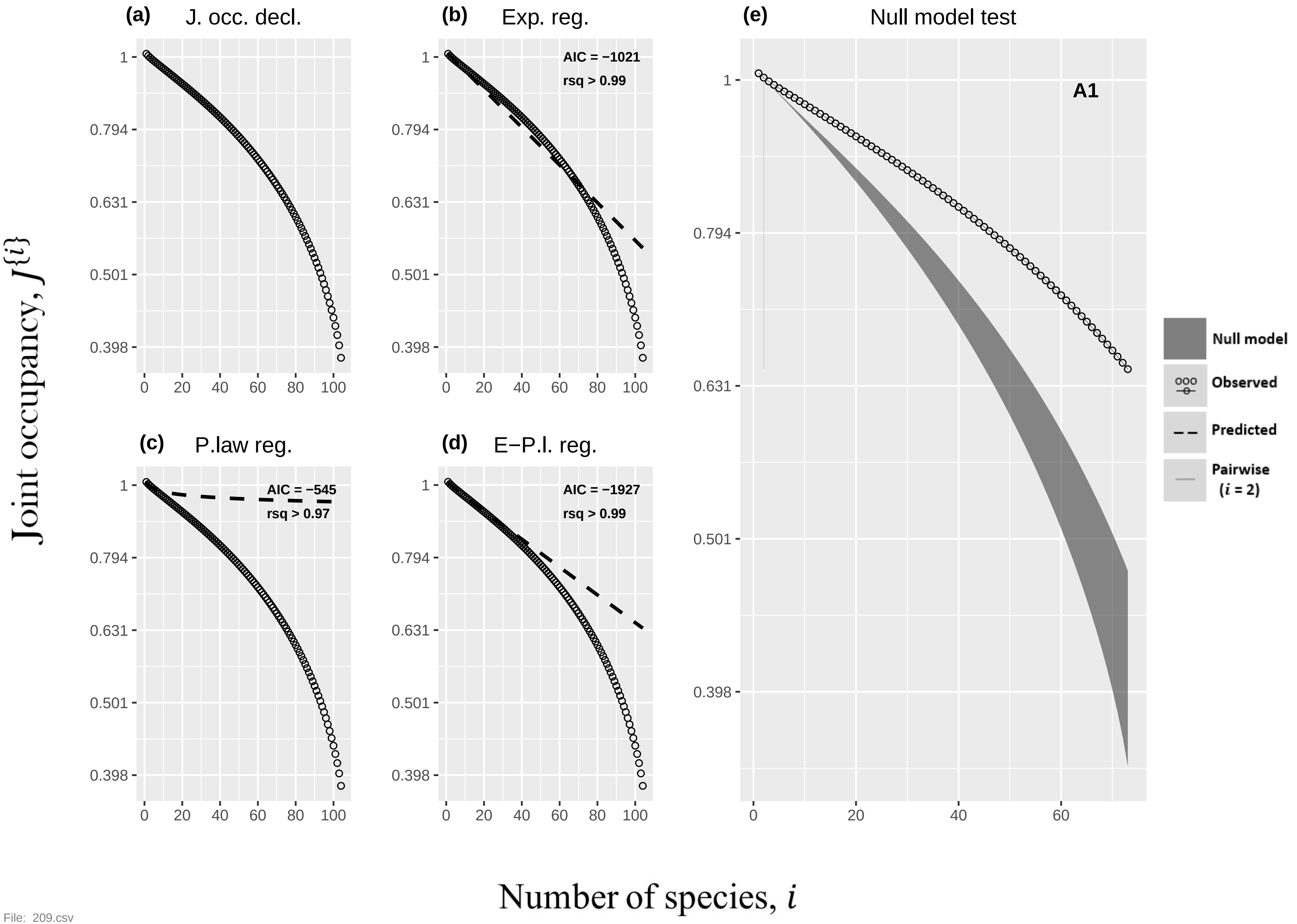

(a) J. occ. decl.
(b) Exp. reg.
AIC = −1021
rsq > 0.99
(c) P.law reg.
AIC = −545
rsq > 0.97
(d) E−P.l. reg.
AIC = −1927
rsq > 0.99
(e) Null model test
A1
Null model
Observed
Predicted
Pairwise (i = 2)
Joint occupancy, J{i}
Number of species, i
1
0.794
0.631
0.501
0.398
0
20
40
60
80
100


File: 209.csv

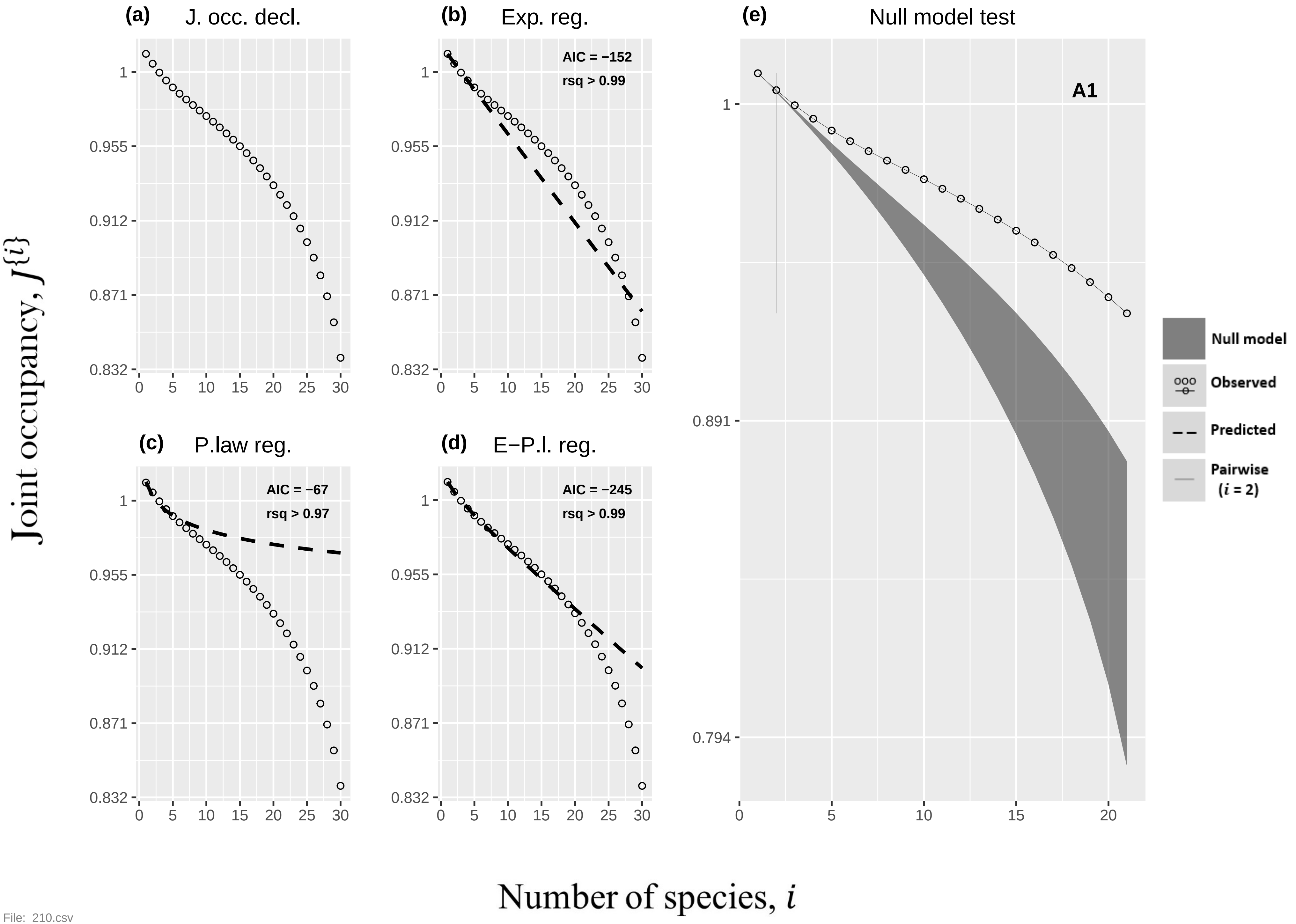

(a)
J. occ. decl.
(b)
Exp. reg.
AIC = −152
rsq > 0.99
(c)
P.law reg.
AIC = −67
rsq > 0.97
(d)
E−P.l. reg.
AIC = −245
rsq > 0.99
(e)
Null model test
A1
Null model
Observed
Predicted
Pairwise
(i = 2)
Joint occupancy, J{i}
Number of species, i
1
0.955
0.912
0.871
0.832
0.891
0.794
0
5
10
15
20
25
30


File: 210.csv

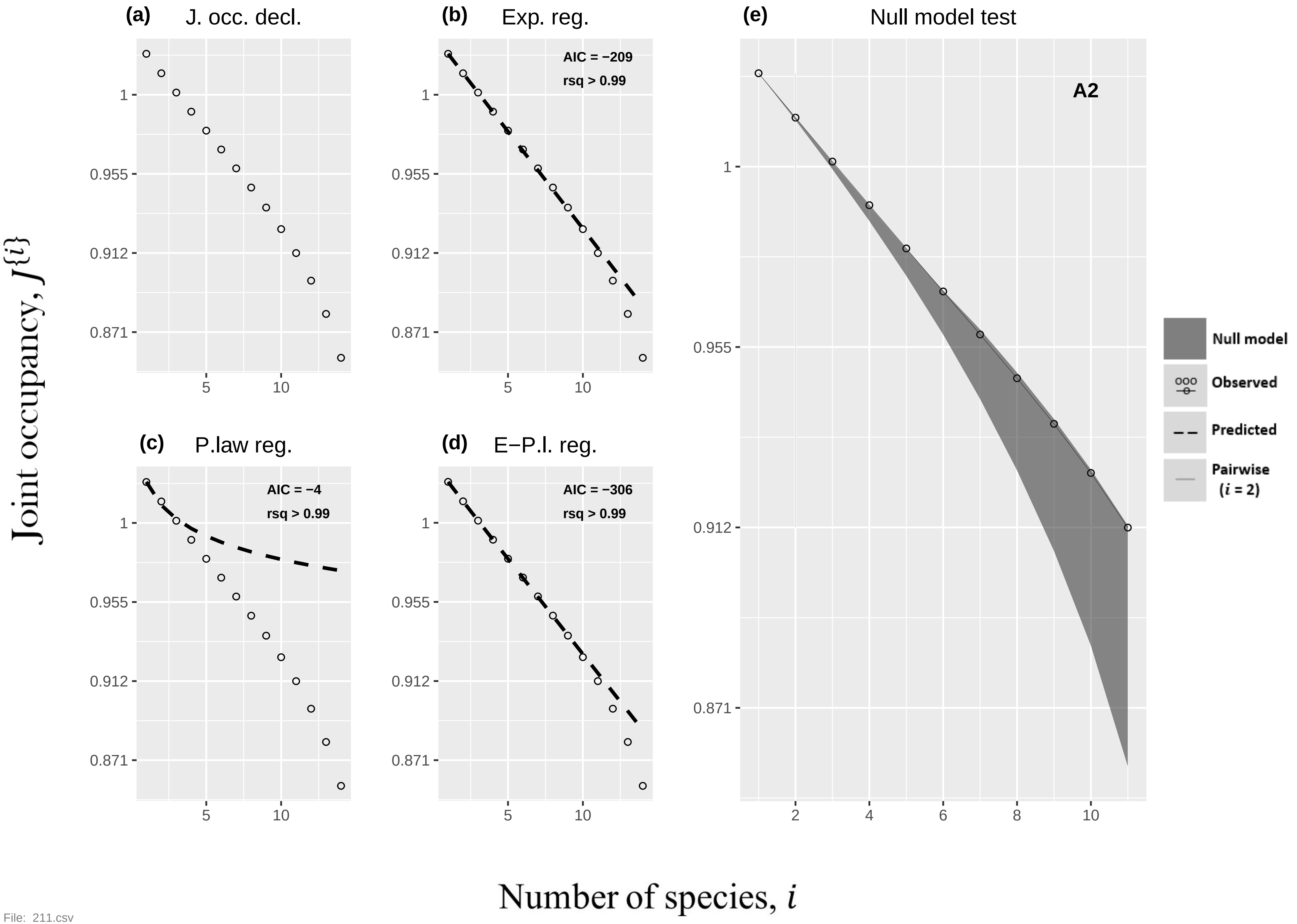

(a)
J. occ. decl.
(b)
Exp. reg.
AIC = −209
rsq > 0.99
(c)
P.law reg.
AIC = −4
rsq > 0.99
(d)
E−P.l. reg.
AIC = −306
rsq > 0.99
(e)
Null model test
A2
1
0.955
0.912
0.871
5
10
2
4
6
8
Null model
Observed
Predicted
Pairwise
(i = 2)
Joint occupancy, J{i}
Number of species, i


File: 211.csv

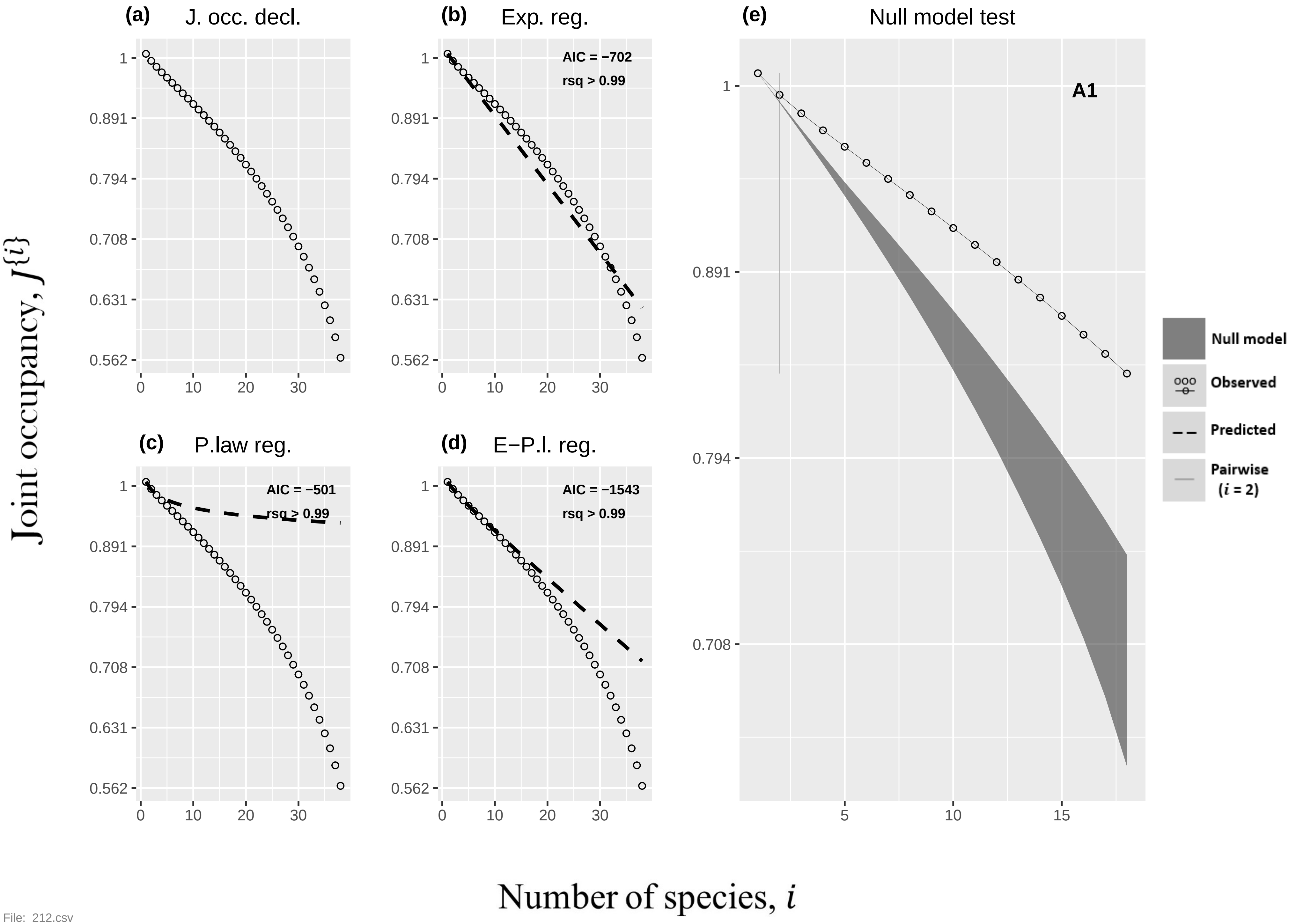
(a) J. occ. decl.
(b) Exp. reg.
AIC = −702
rsq > 0.99
(e) Null model test
A1
(c) P.law reg.
AIC = −501
rsq > 0.99
(d) E−P.l. reg.
AIC = −1543
rsq > 0.99
1
0.891
0.794
0.708
0.631
0.562
0
10
20
30
5
15
Null model
Observed
Predicted
Pairwise (i = 2)
Joint occupancy, J{i}
Number of species, i

File: 212.csv

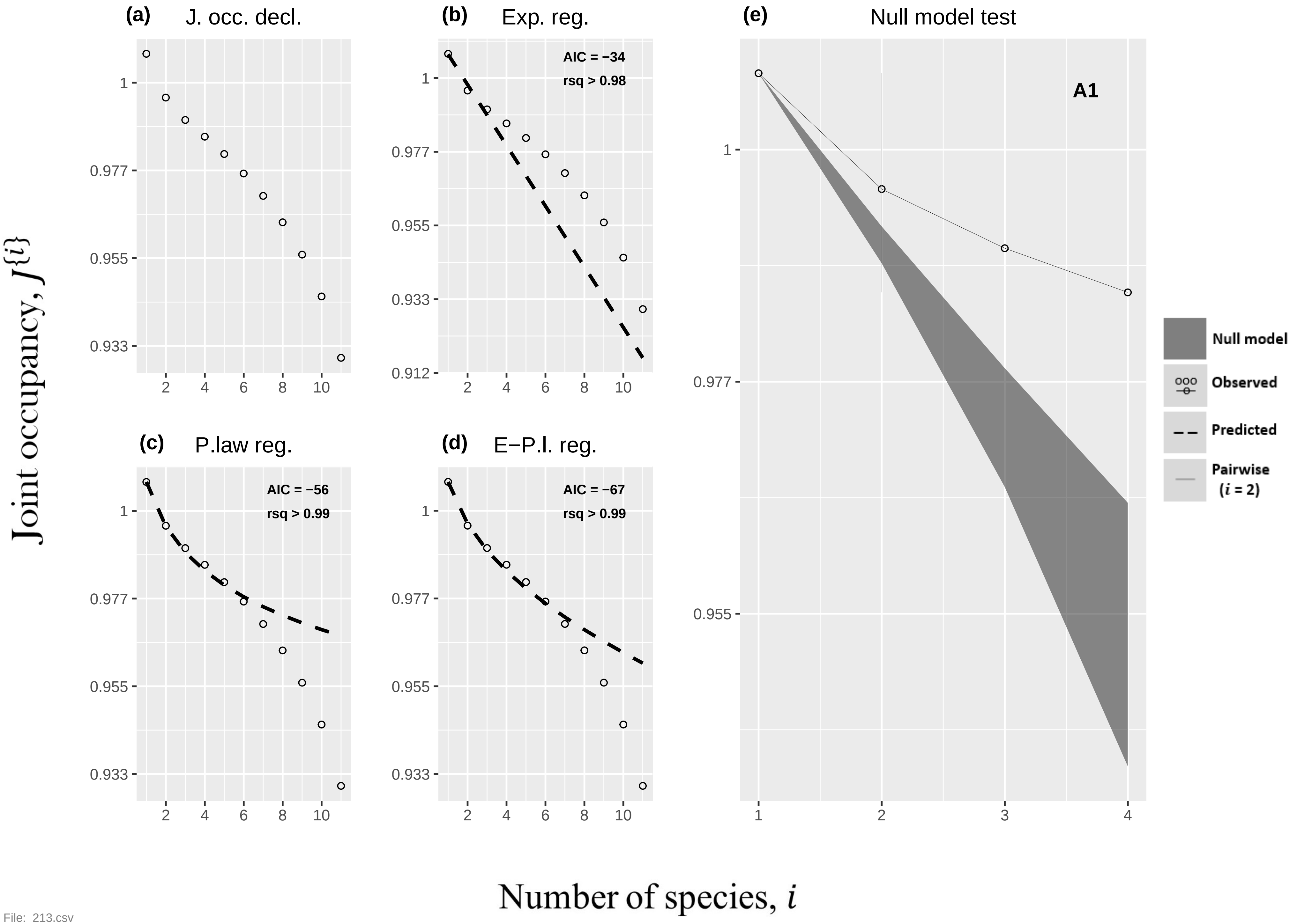
(a) J. occ. decl.
(b) Exp. reg.
AIC = −34
rsq > 0.98
(c) P.law reg.
AIC = −56
rsq > 0.99
(d) E−P.l. reg.
AIC = −67
rsq > 0.99
(e) Null model test
A1
Null model
Observed
Predicted
Pairwise (i = 2)
Joint occupancy, J{i}
Number of species, i

File: 213.csv

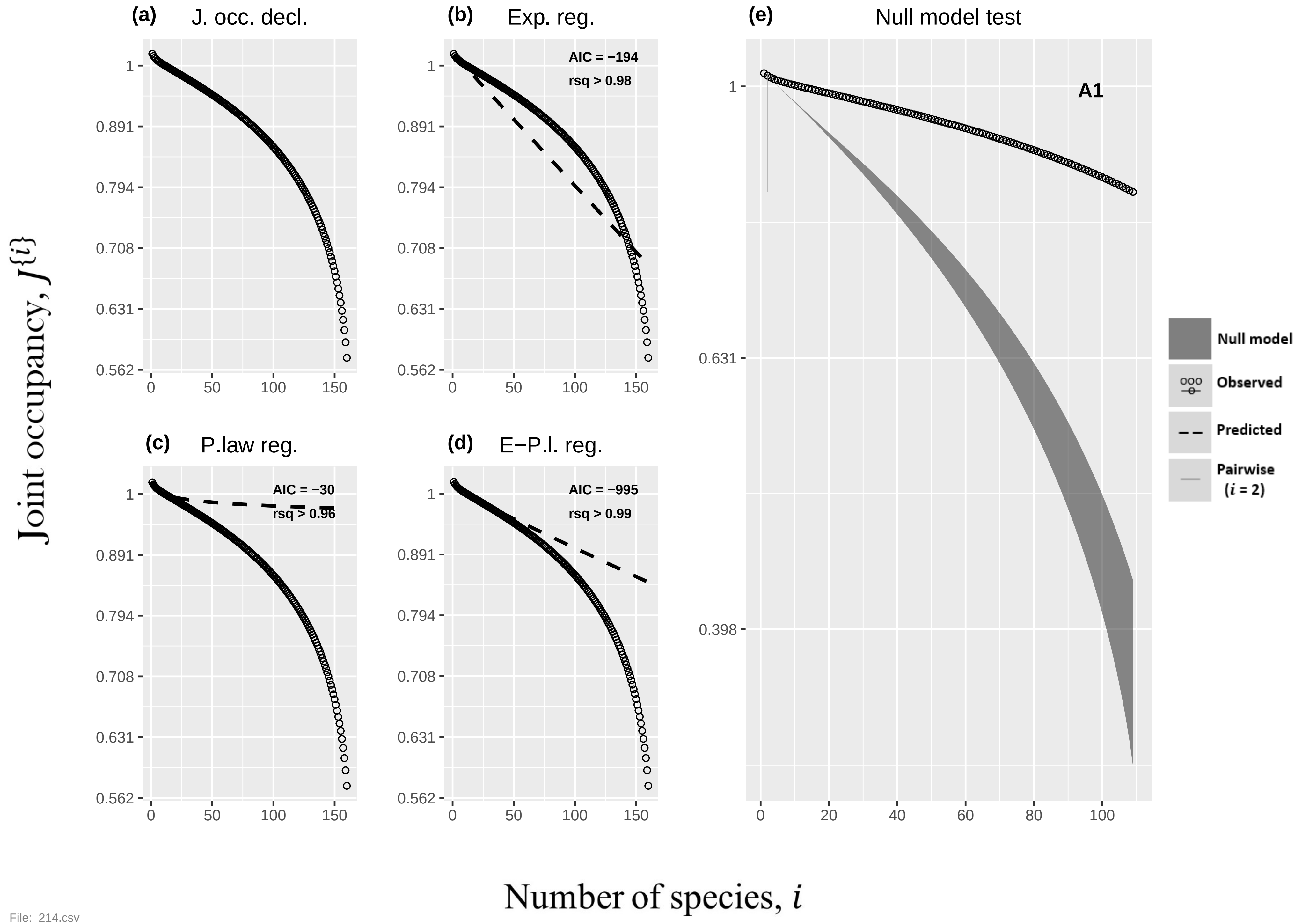

(a) J. occ. decl.
(b) Exp. reg.
AIC = −194
rsq > 0.98
(c) P.law reg.
AIC = −30
rsq > 0.96
(d) E−P.l. reg.
AIC = −995
rsq > 0.99
(e) Null model test
A1
Null model
Observed
Predicted
Pairwise (i = 2)
1
0.891
0.794
0.708
0.631
0.562
0.398
0
50
100
150
20
40
60
80
Joint occupancy, J{i}
Number of species, i


File: 214.csv

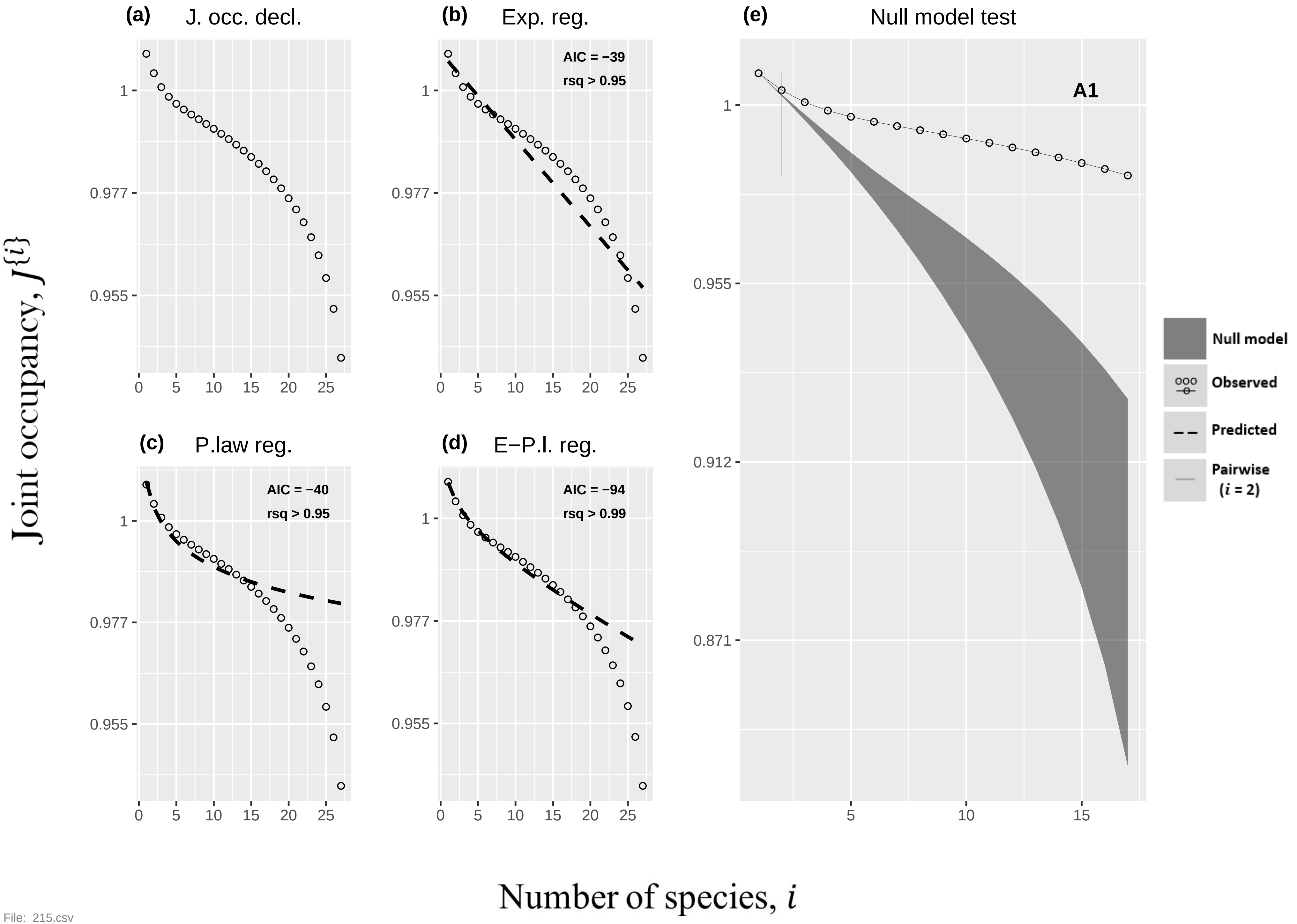
(a) J. occ. decl.
(b) Exp. reg.
AIC = −39
rsq > 0.95
(c) P.law reg.
AIC = −40
rsq > 0.95
(d) E−P.l. reg.
AIC = −94
rsq > 0.99
(e) Null model test
A1
Null model
Observed
Predicted
Pairwise
(i = 2)
Joint occupancy, J{i}
Number of species, i
1
0.977
0.955
0.912
0.871
0
5
10
15
20
25

File: 215.csv

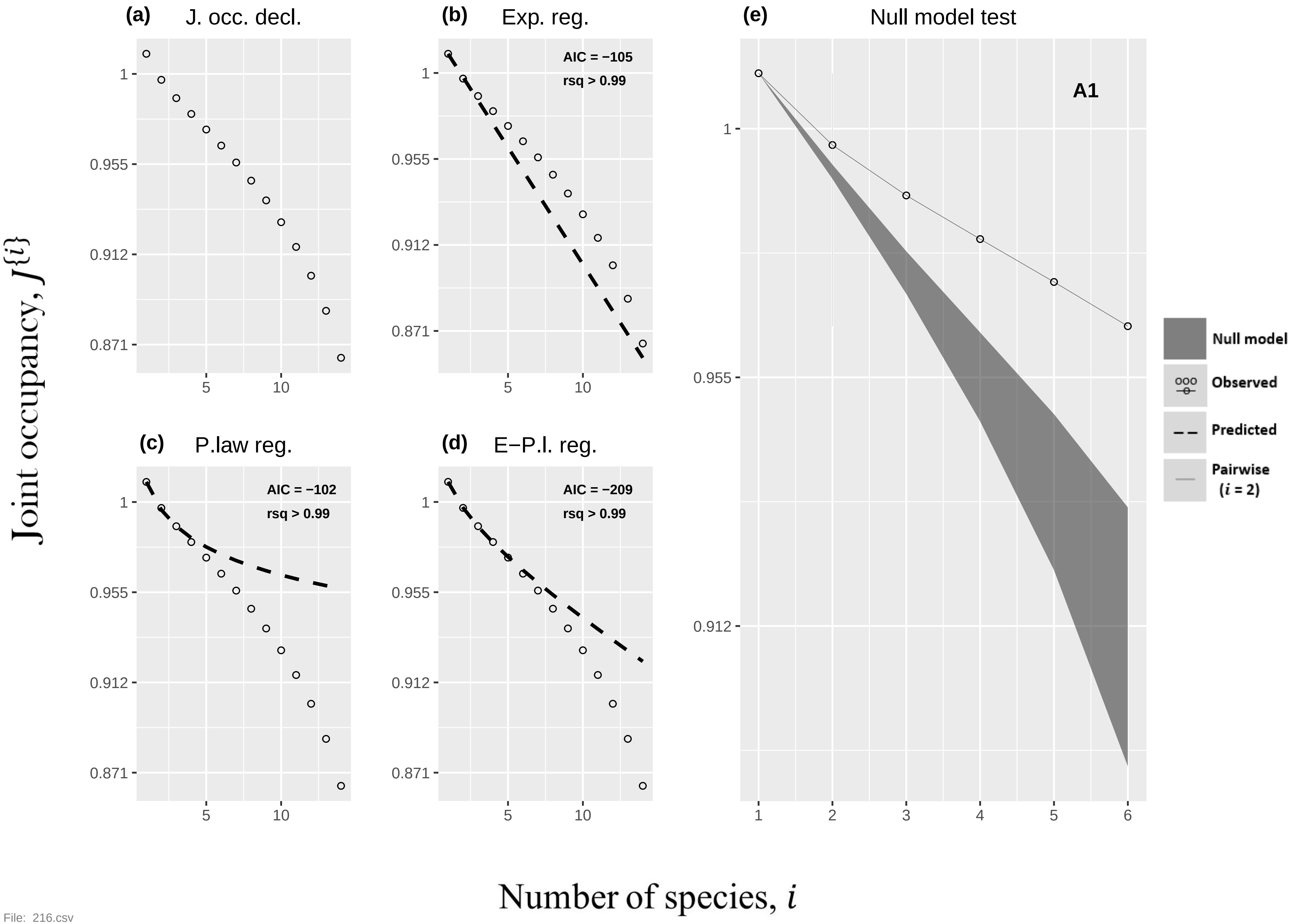

(a)
J. occ. decl.
(b)
Exp. reg.
AIC = −105
rsq > 0.99
(c)
P.law reg.
AIC = −102
rsq > 0.99
(d)
E−P.l. reg.
AIC = −209
rsq > 0.99
(e)
Null model test
A1
Null model
Observed
Predicted
Pairwise
(i = 2)
1
0.955
0.912
0.871
5
10
1
2
3
4
5
6
Joint occupancy, J{i}
Number of species, i


File: 216.csv

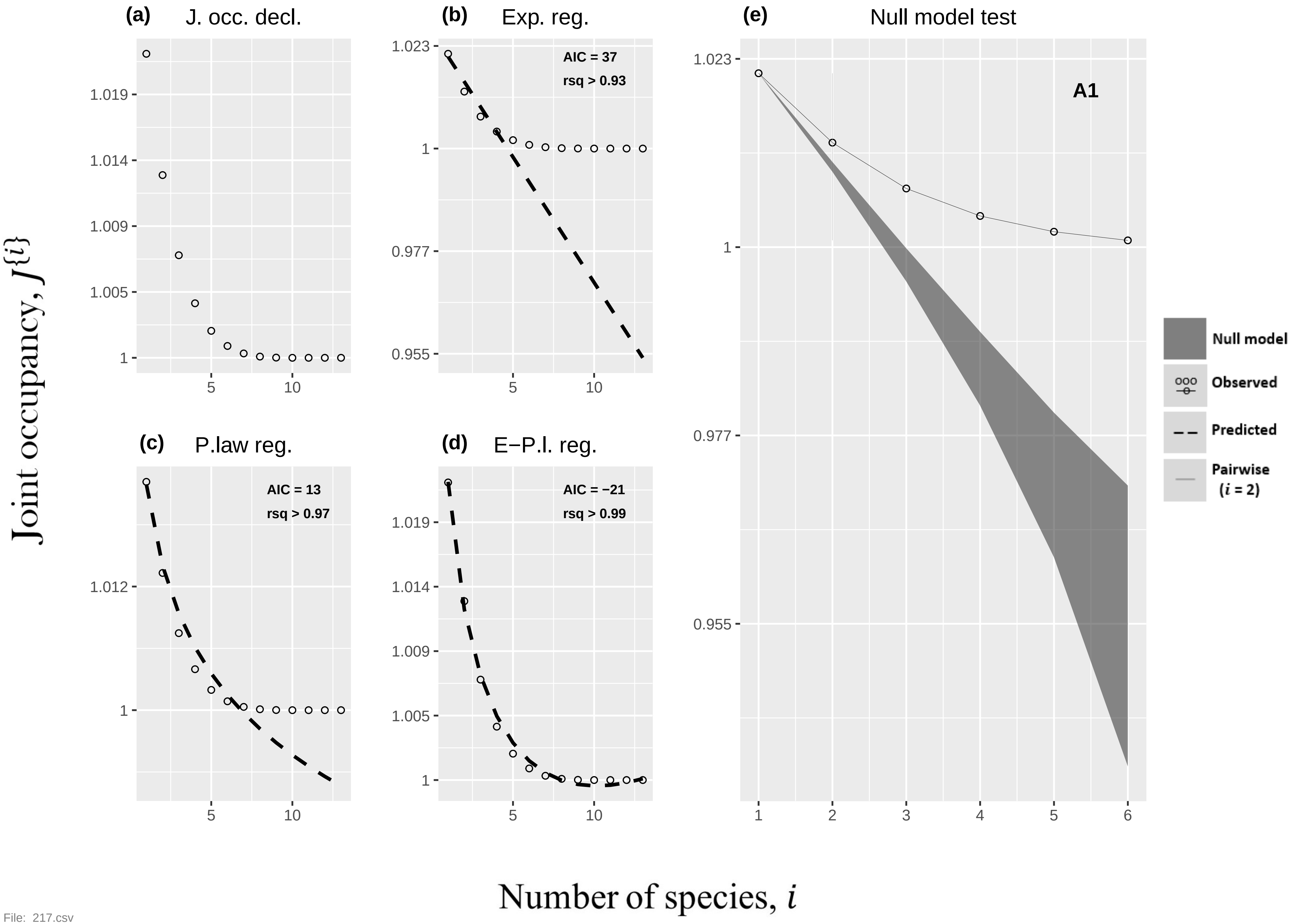
(a) J. occ. decl.
(b) Exp. reg.
AIC = 37
rsq > 0.93
(e) Null model test
A1
(c) P.law reg.
AIC = 13
rsq > 0.97
(d) E−P.l. reg.
AIC = −21
rsq > 0.99
Null model
Observed
Predicted
Pairwise (i = 2)
Joint occupancy, J{i}
Number of species, i

File: 217.csv

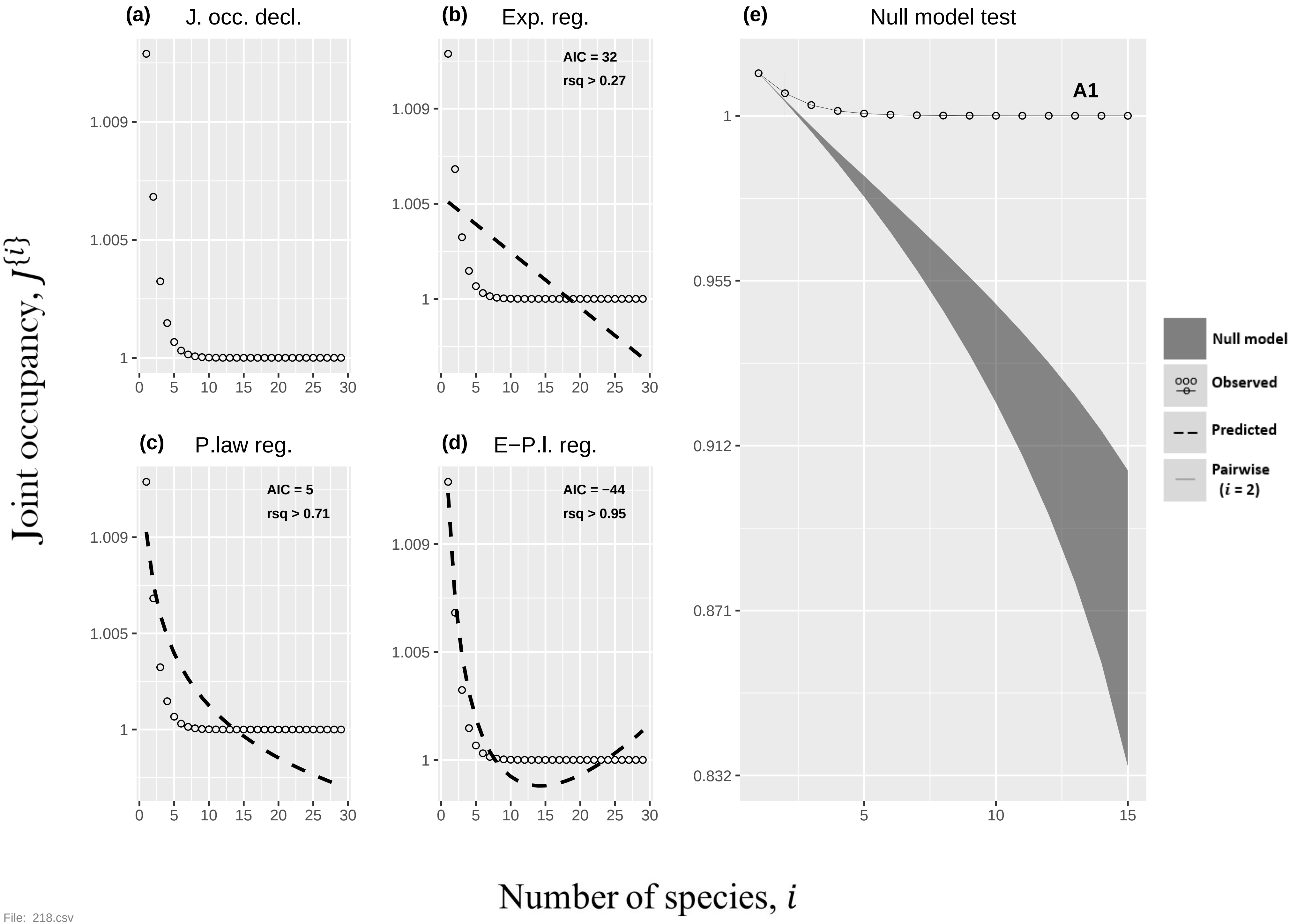
(a)
J. occ. decl.
(b)
Exp. reg.
AIC = 32
rsq > 0.27
(c)
P.law reg.
AIC = 5
rsq > 0.71
(d)
E−P.l. reg.
AIC = −44
rsq > 0.95
(e)
Null model test
A1
Null model
Observed
Predicted
Pairwise
(i = 2)
Joint occupancy, J{i}
Number of species, i
1.009
1.005
1
0.955
0.912
0.871
0.832
0
5
10
15
20
25
30

File: 218.csv

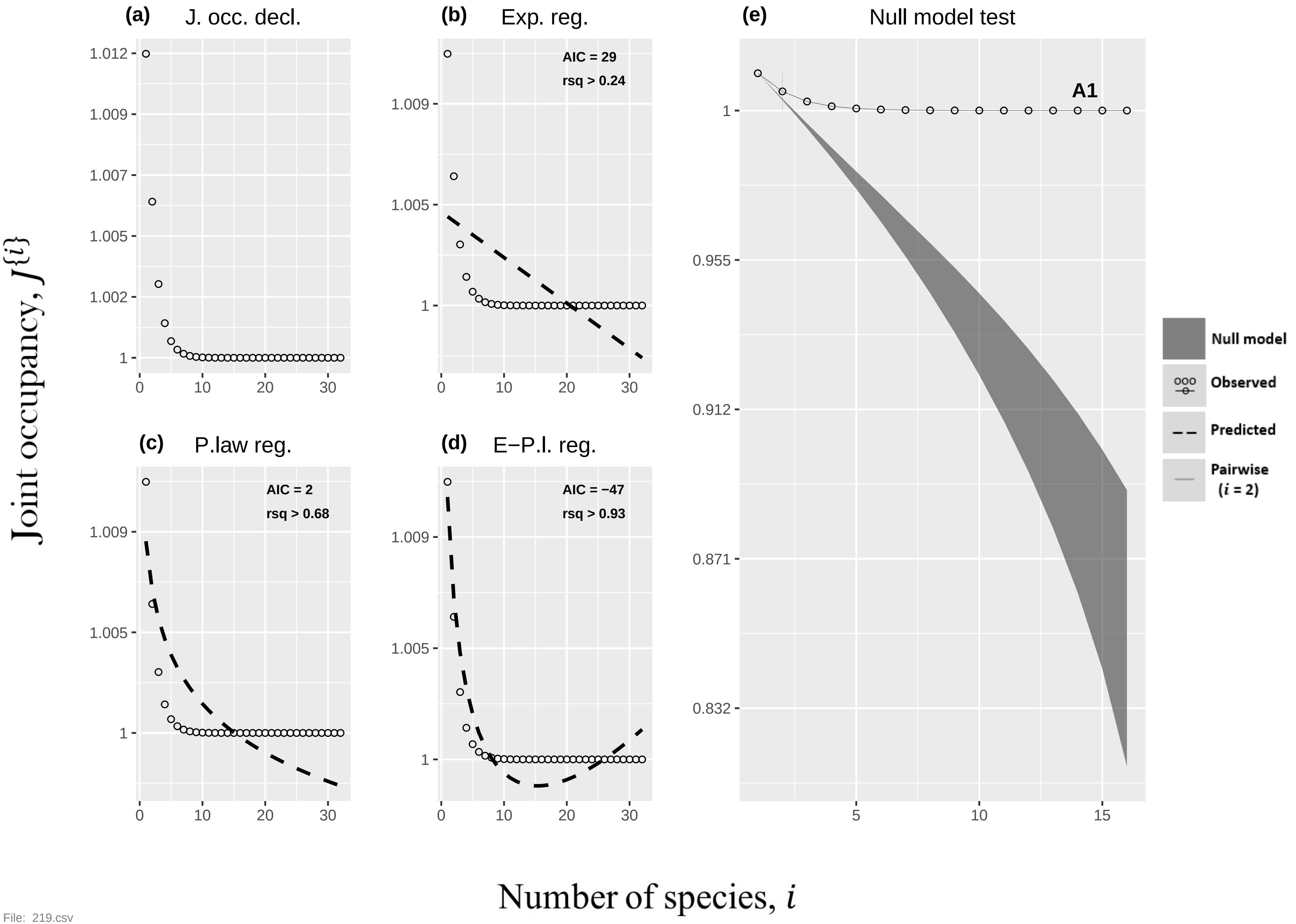

(a)
J. occ. decl.
1.012
1.009
1.007
1.005
1.002
1
0
10
20
30
(b)
Exp. reg.
AIC = 29
rsq > 0.24
1.009
1.005
1
(c)
P.law reg.
AIC = 2
rsq > 0.68
(d)
E−P.l. reg.
AIC = −47
rsq > 0.93
(e)
Null model test
A1
0.955
0.912
0.871
0.832
5
15
Null model
Observed
Predicted
Pairwise
(i = 2)
Joint occupancy, J{i}
Number of species, i

File: 219.csv

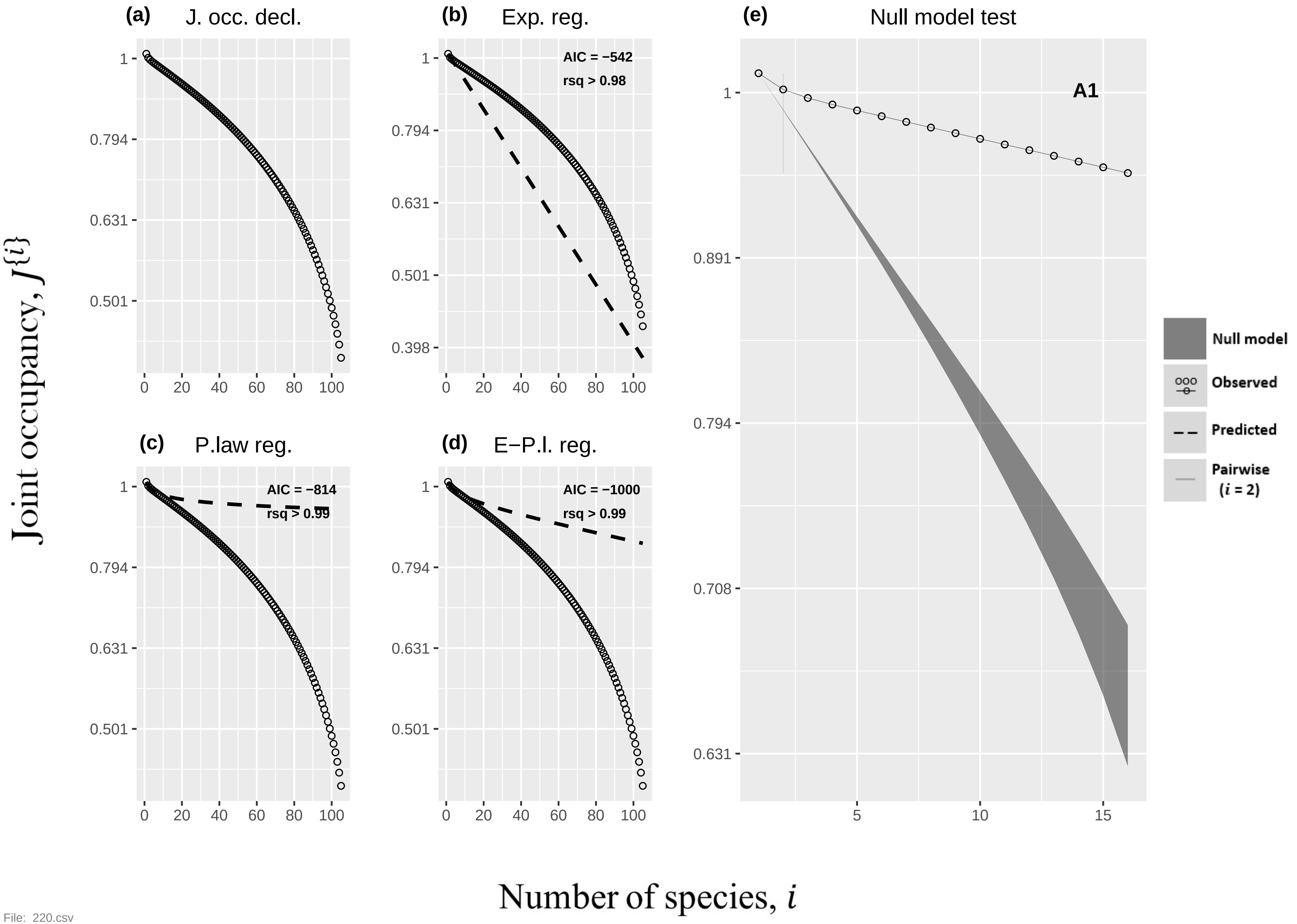

(a) J. occ. decl.
(b) Exp. reg.
AIC = −542
rsq > 0.98
(c) P.law reg.
AIC = −814
rsq > 0.99
(d) E−P.l. reg.
AIC = −1000
rsq > 0.99
(e) Null model test
A1
Null model
Observed
Predicted
Pairwise (i = 2)
Joint occupancy, J{i}
Number of species, i


File: 220.csv

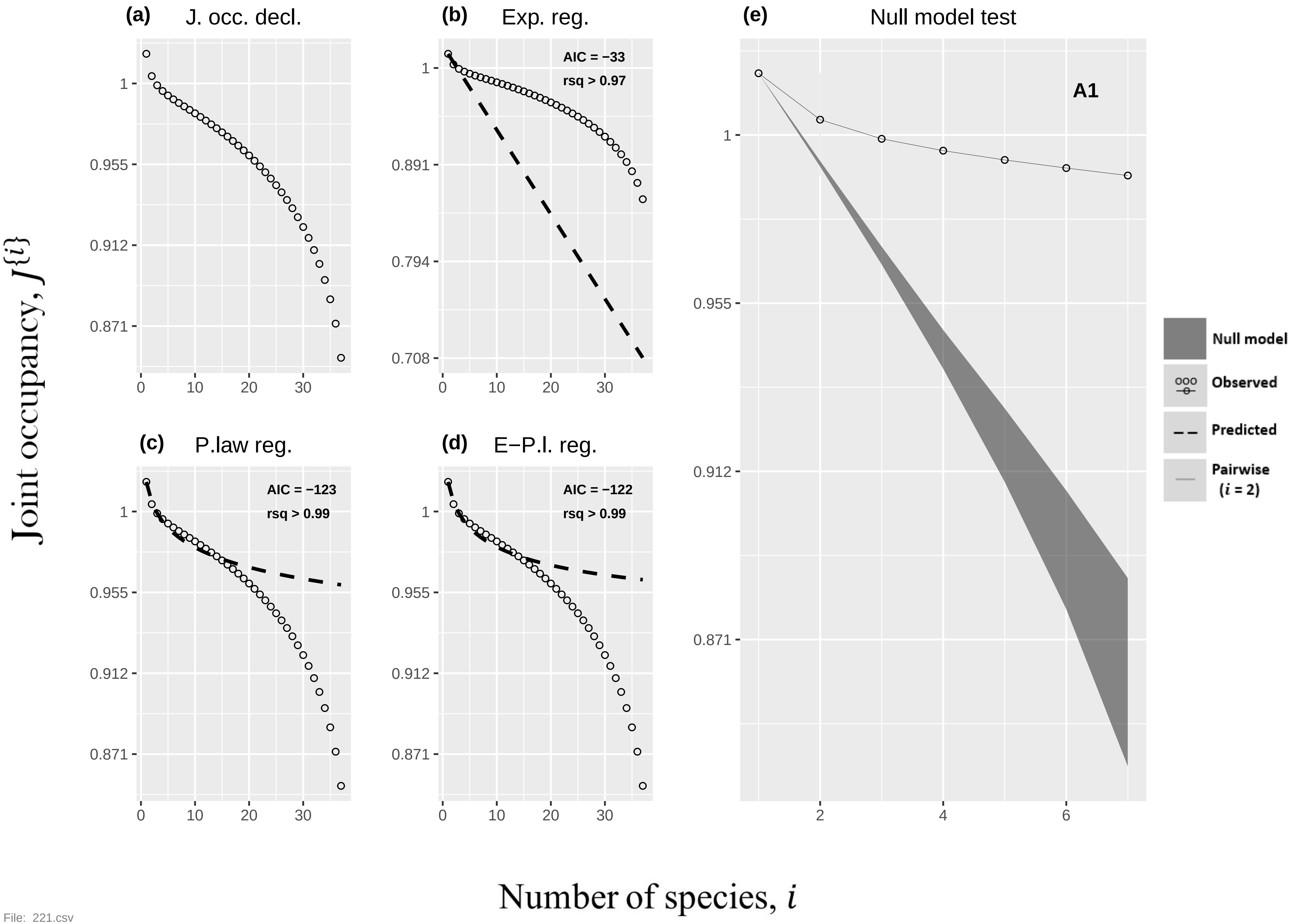

(a) J. occ. decl.
(b) Exp. reg.
AIC = −33
rsq > 0.97
(c) P.law reg.
AIC = −123
rsq > 0.99
(d) E−P.l. reg.
AIC = −122
rsq > 0.99
(e) Null model test
A1
Null model
Observed
Predicted
Pairwise (i = 2)
Joint occupancy, J{i}
Number of species, i


File: 221.csv

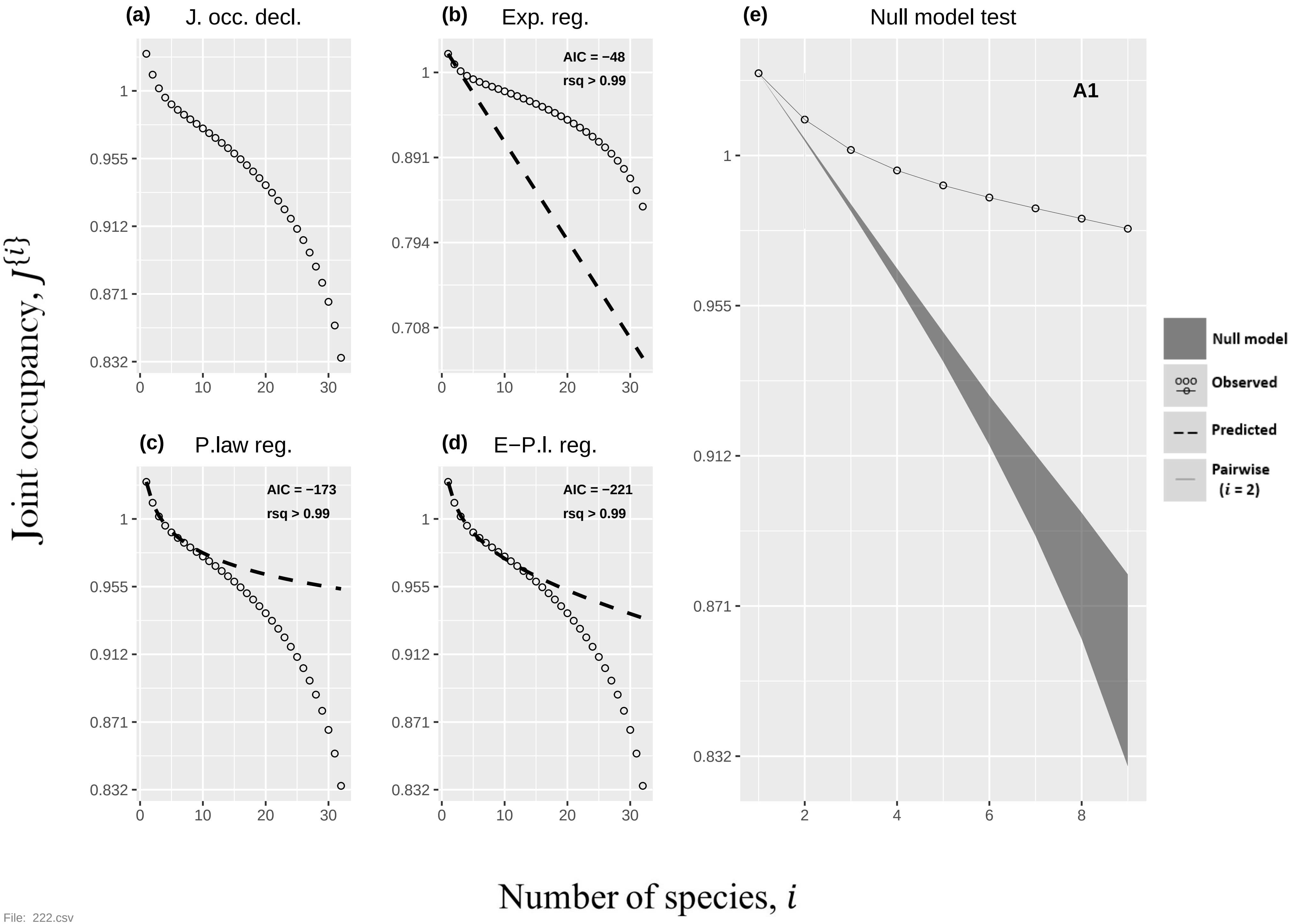

(a) J. occ. decl.
(b) Exp. reg.
AIC = −48
rsq > 0.99
(c) P.law reg.
AIC = −173
rsq > 0.99
(d) E−P.l. reg.
AIC = −221
rsq > 0.99
(e) Null model test
A1
Null model
Observed
Predicted
Pairwise
(i = 2)
Joint occupancy, J{i}
Number of species, i


File: 222.csv

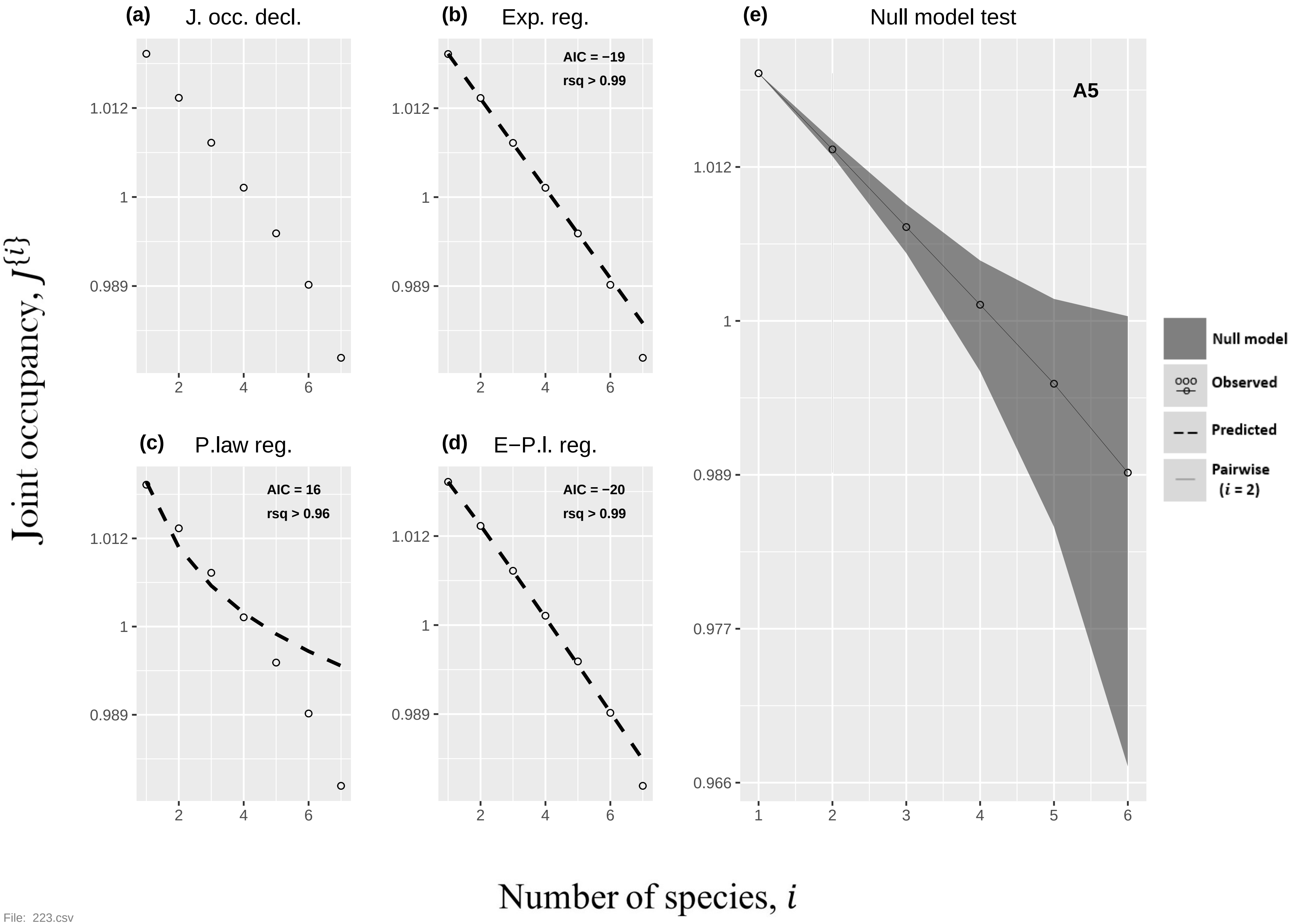

(a) J. occ. decl.
(b) Exp. reg.
AIC = −19
rsq > 0.99
(c) P.law reg.
AIC = 16
rsq > 0.96
(d) E−P.l. reg.
AIC = −20
rsq > 0.99
(e) Null model test
A5
Null model
Observed
Predicted
Pairwise (i = 2)
Joint occupancy, J{i}
Number of species, i
1.012
1
0.989
0.977
0.966


File: 223.csv

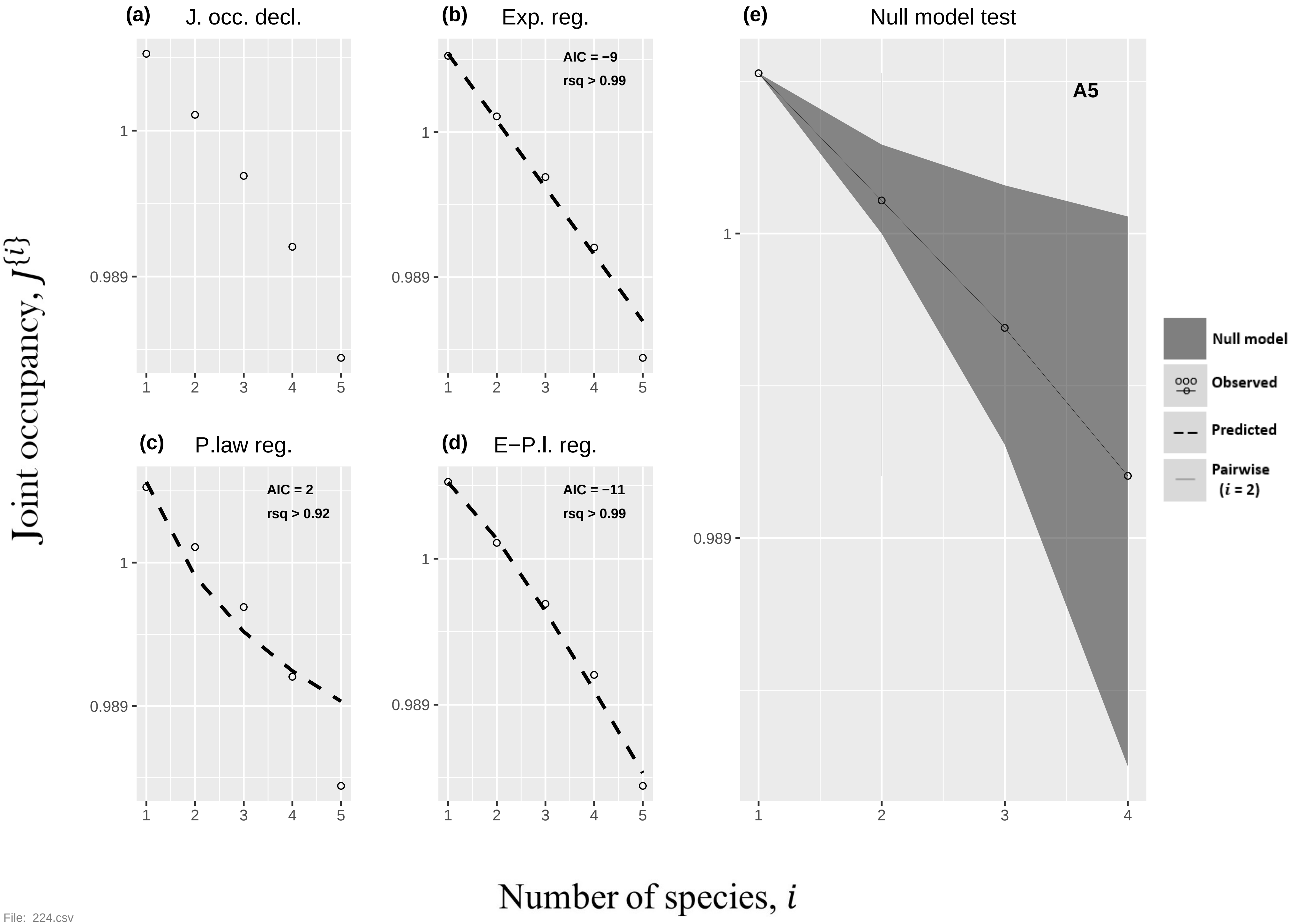

(a) J. occ. decl.
(b) Exp. reg.
AIC = −9
rsq > 0.99
(e) Null model test
A5
(c) P.law reg.
AIC = 2
rsq > 0.92
(d) E−P.l. reg.
AIC = −11
rsq > 0.99
Null model
Observed
Predicted
Pairwise ($i$ = 2)
1
0.989
1 2 3 4 5
1 2 3 4
Joint occupancy, $J^{\{i\}}$
Number of species, $i$


File: 224.csv

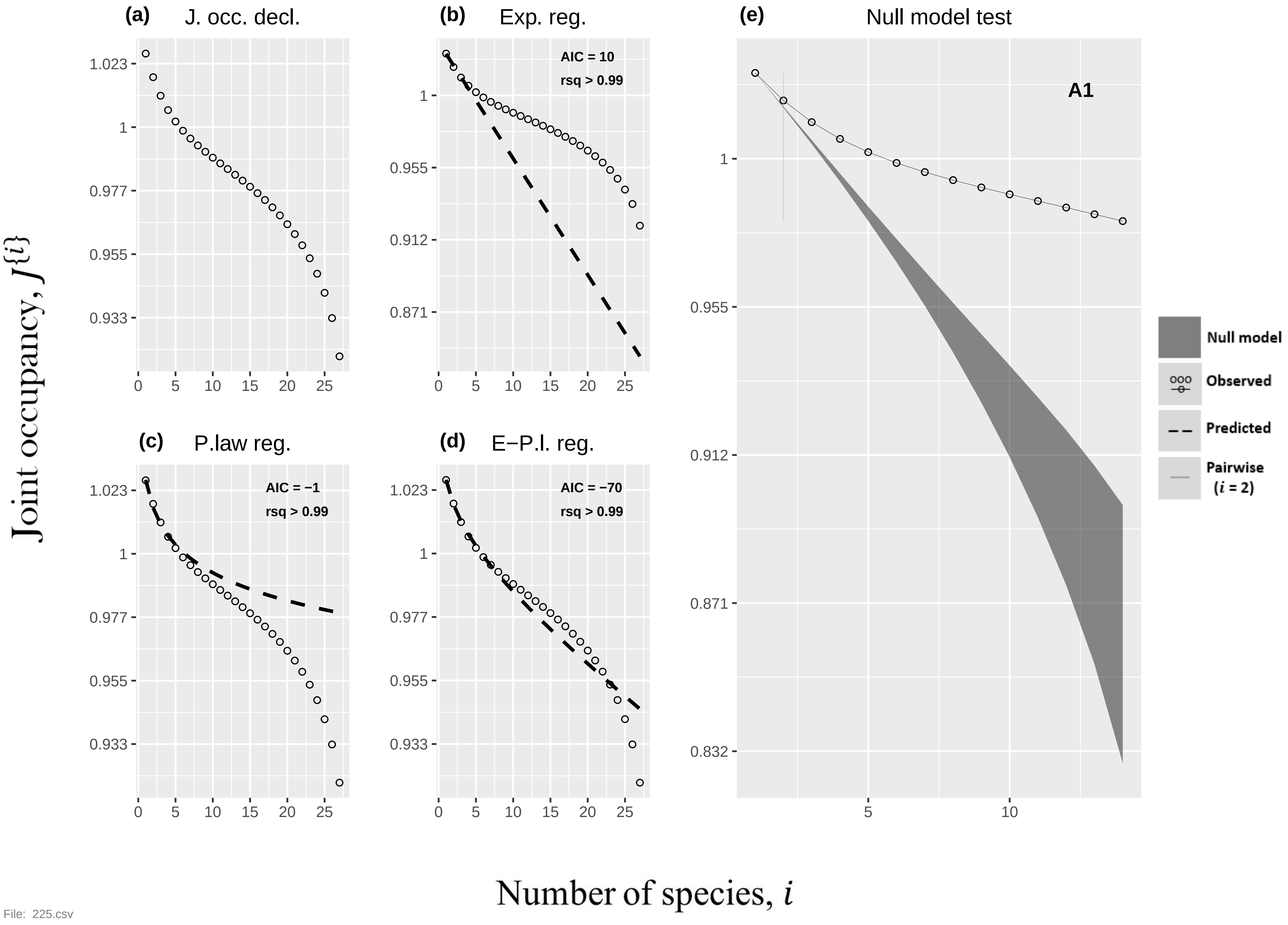
(a)
J. occ. decl.
(b)
Exp. reg.
AIC = 10
rsq > 0.99
(c)
P.law reg.
AIC = −1
rsq > 0.99
(d)
E−P.l. reg.
AIC = −70
rsq > 0.99
(e)
Null model test
A1
Null model
Observed
Predicted
Pairwise
(i = 2)
Joint occupancy, J{i}
Number of species, i
1.023
1
0.977
0.955
0.933
0.912
0.871
0.832
0
5
10
15
20
25
File: 225.csv

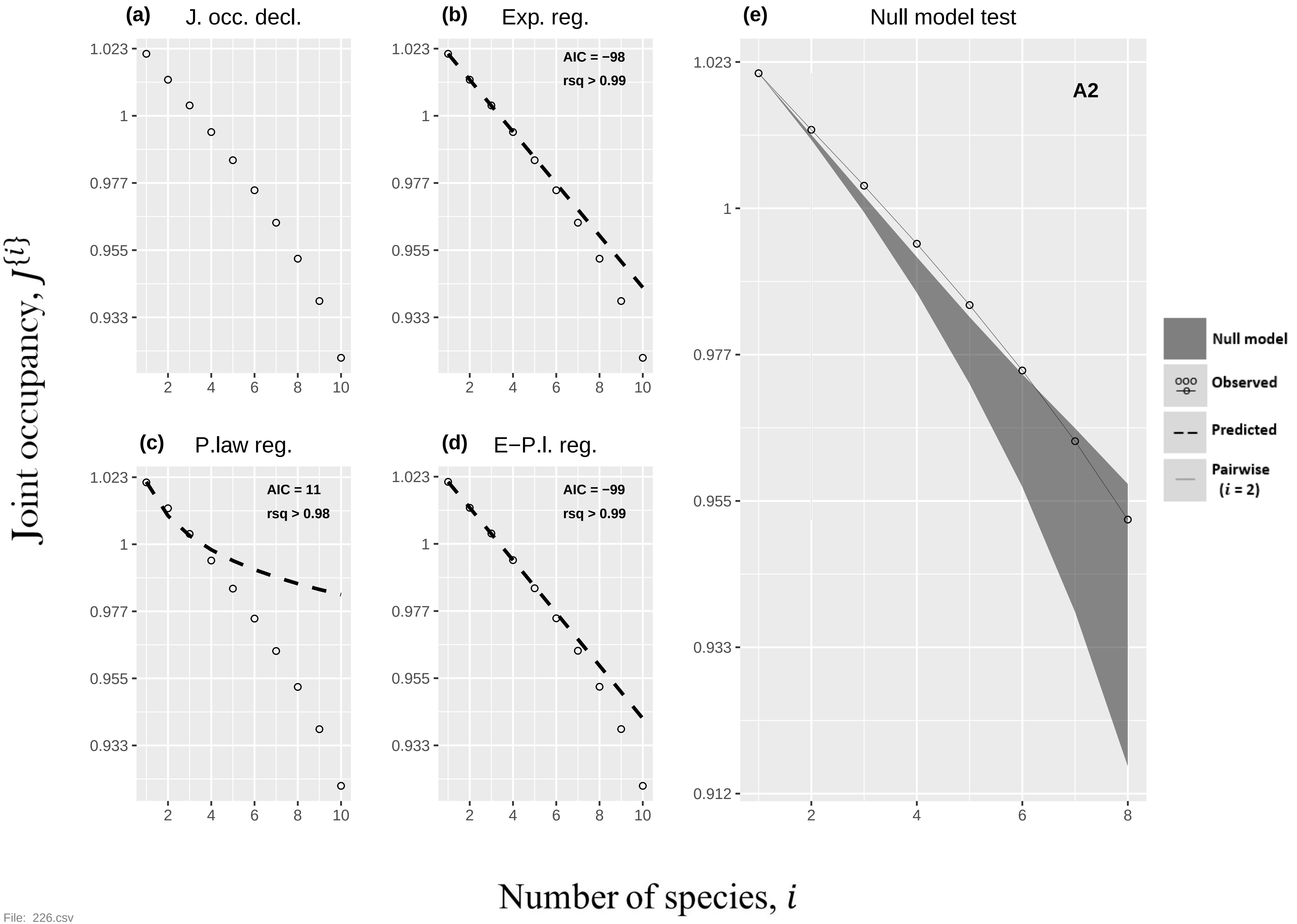
(a) J. occ. decl.
(b) Exp. reg.
AIC = −98
rsq > 0.99
(c) P.law reg.
AIC = 11
rsq > 0.98
(d) E−P.l. reg.
AIC = −99
rsq > 0.99
(e) Null model test
A2
Null model
Observed
Predicted
Pairwise (i = 2)
Joint occupancy, J{i}
Number of species, i
1.023
1
0.977
0.955
0.933
0.912
2
4
6
8
10

File: 226.csv

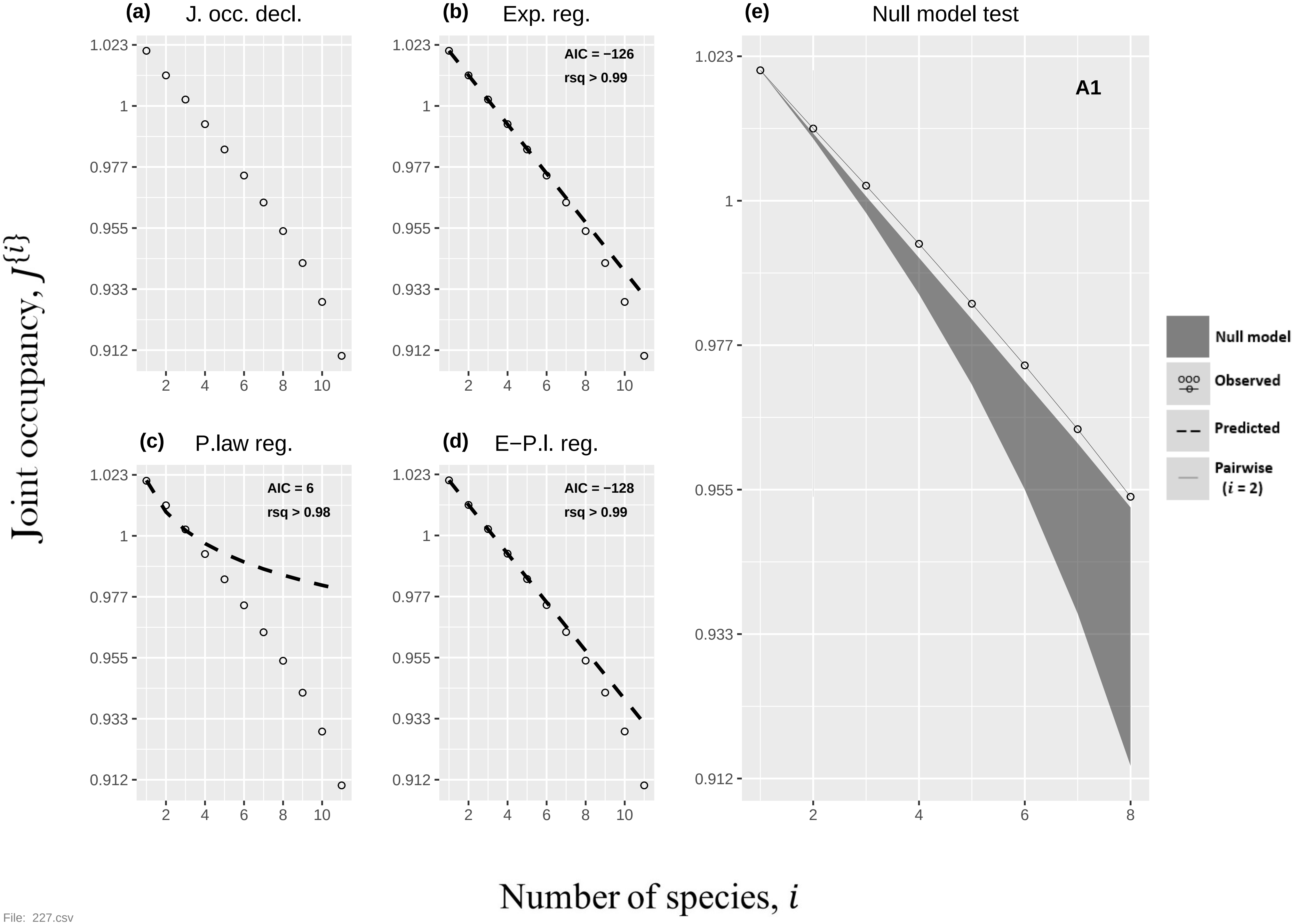

(a)
J. occ. decl.
(b)
Exp. reg.
AIC = −126
rsq > 0.99
(c)
P.law reg.
AIC = 6
rsq > 0.98
(d)
E−P.l. reg.
AIC = −128
rsq > 0.99
(e)
Null model test
A1
Null model
Observed
Predicted
Pairwise (i = 2)
1.023
1
0.977
0.955
0.933
0.912
2
4
6
8
10
Joint occupancy, J{i}
Number of species, i


File: 227.csv

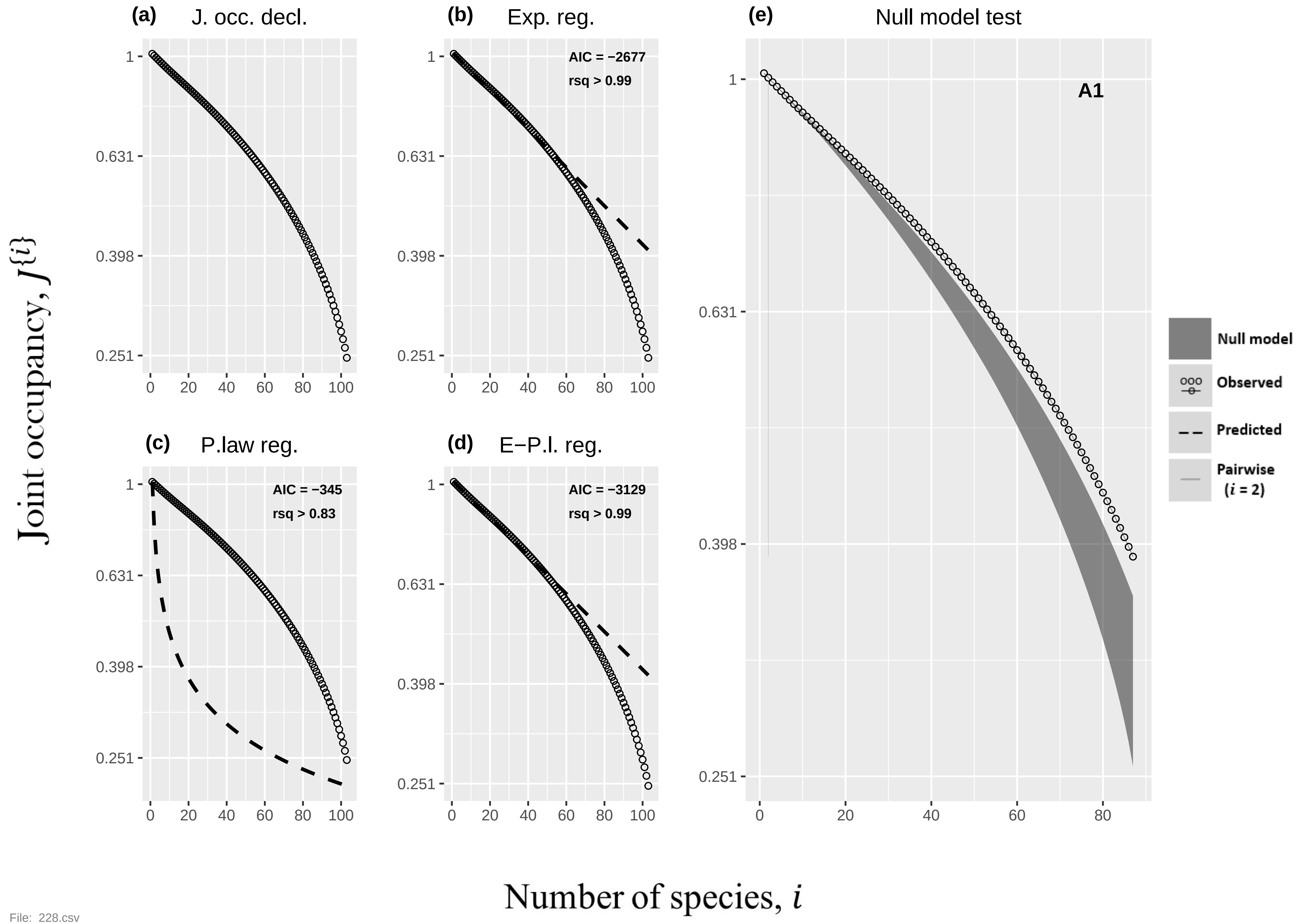

(a) J. occ. decl.
(b) Exp. reg.
AIC = −2677
rsq > 0.99
(c) P.law reg.
AIC = −345
rsq > 0.83
(d) E−P.l. reg.
AIC = −3129
rsq > 0.99
(e) Null model test
A1
Null model
Observed
Predicted
Pairwise (i = 2)
Joint occupancy, J{i}
Number of species, i
1
0.631
0.398
0.251
0
20
40
60
80
100

File: 228.csv

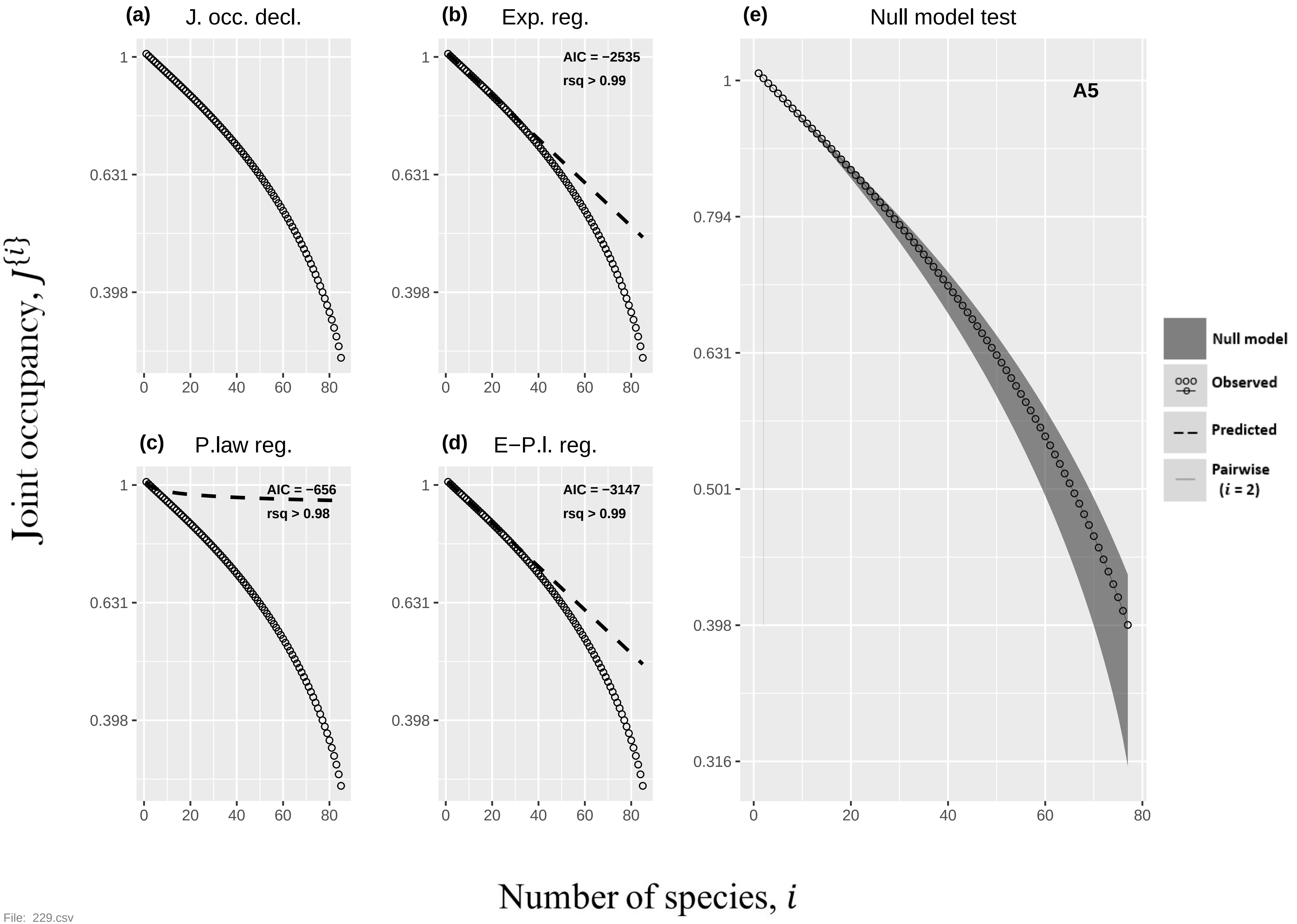
(a) J. occ. decl.
(b) Exp. reg.
AIC = −2535
rsq > 0.99
(c) P.law reg.
AIC = −656
rsq > 0.98
(d) E−P.l. reg.
AIC = −3147
rsq > 0.99
(e) Null model test
A5
Null model
Observed
Predicted
Pairwise (i = 2)
Joint occupancy, J{i}
Number of species, i

File: 229.csv

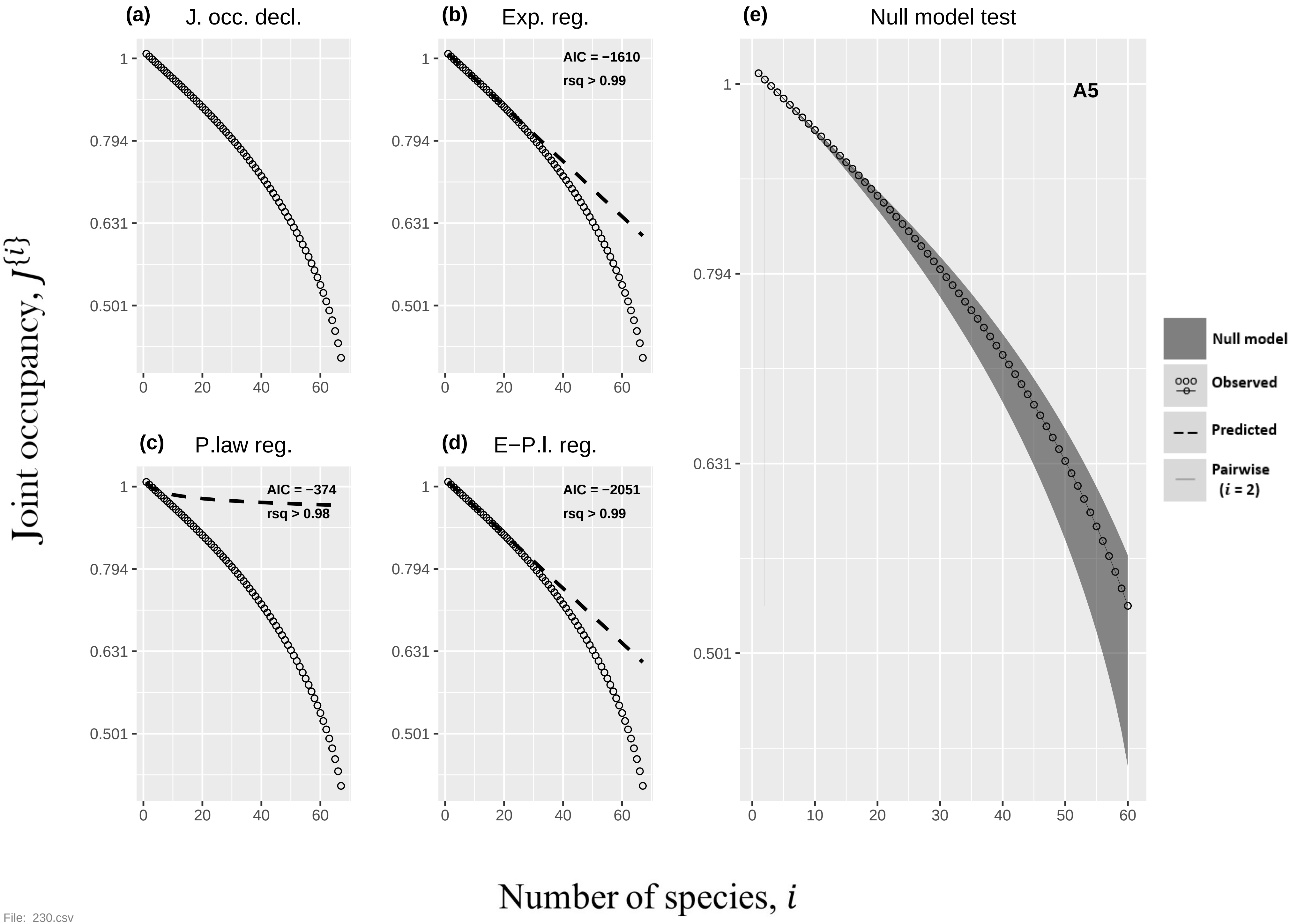
(a) J. occ. decl.
(b) Exp. reg.
AIC = −1610
rsq > 0.99
(c) P.law reg.
AIC = −374
rsq > 0.98
(d) E−P.l. reg.
AIC = −2051
rsq > 0.99
(e) Null model test
A5
Null model
Observed
Predicted
Pairwise ($i$ = 2)
Joint occupancy, $J^{\{i\}}$
Number of species, $i$
1
0.794
0.631
0.501
0
10
20
30
40
50
60

File: 230.csv

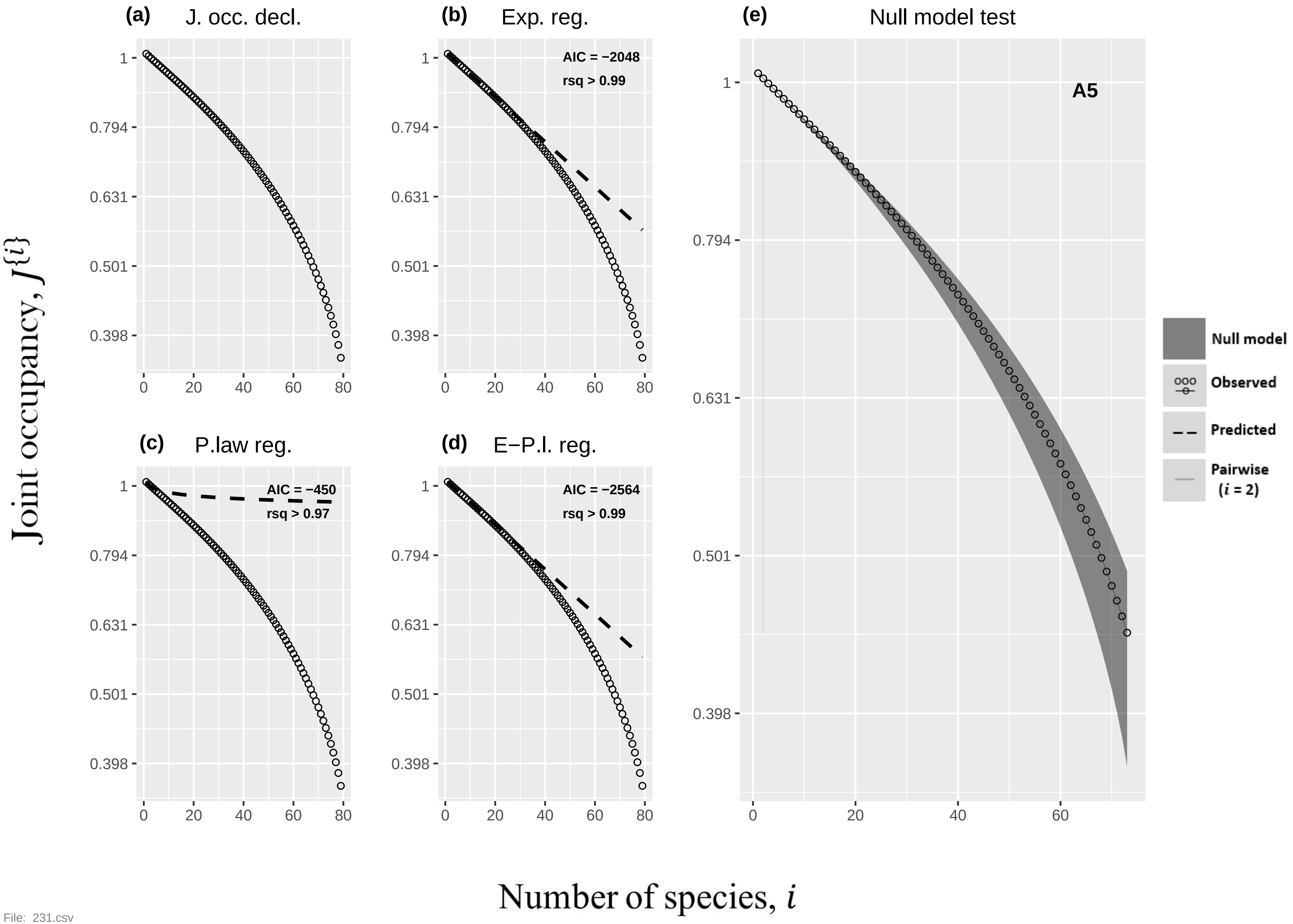

(a) J. occ. decl.
(b) Exp. reg.
AIC = −2048
rsq > 0.99
(c) P.law reg.
AIC = −450
rsq > 0.97
(d) E−P.l. reg.
AIC = −2564
rsq > 0.99
(e) Null model test
A5
Null model
Observed
Predicted
Pairwise
(i = 2)
1
0.794
0.631
0.501
0.398
0
20
40
60
80
Joint occupancy, J{i}
Number of species, i


File: 231.csv

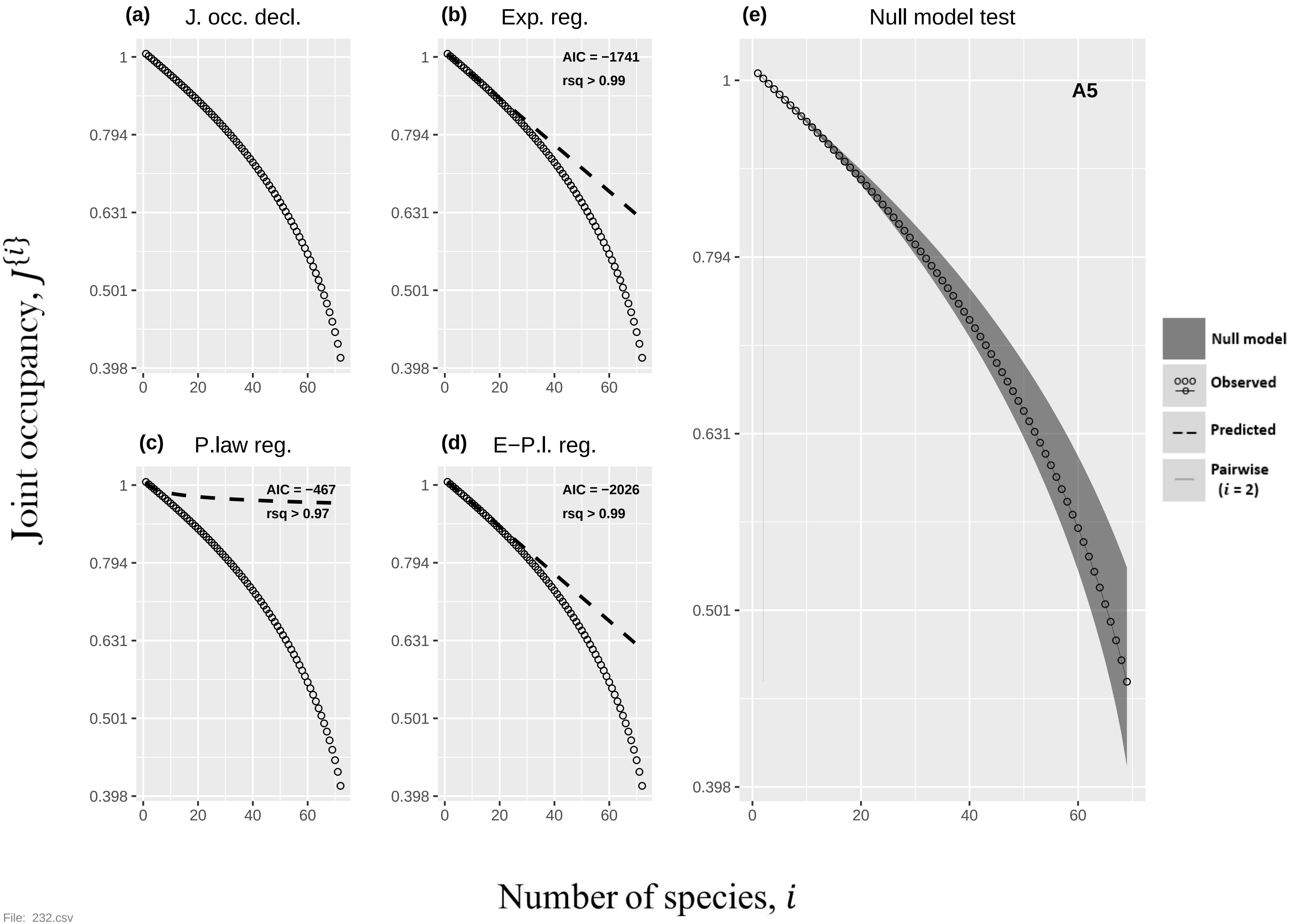

(a) J. occ. decl.
(b) Exp. reg.
AIC = −1741
rsq > 0.99
(c) P.law reg.
AIC = −467
rsq > 0.97
(d) E−P.l. reg.
AIC = −2026
rsq > 0.99
(e) Null model test
A5
Null model
Observed
Predicted
Pairwise (i = 2)
Joint occupancy, J{i}
Number of species, i
1
0.794
0.631
0.501
0.398
0
20
40
60


File: 232.csv

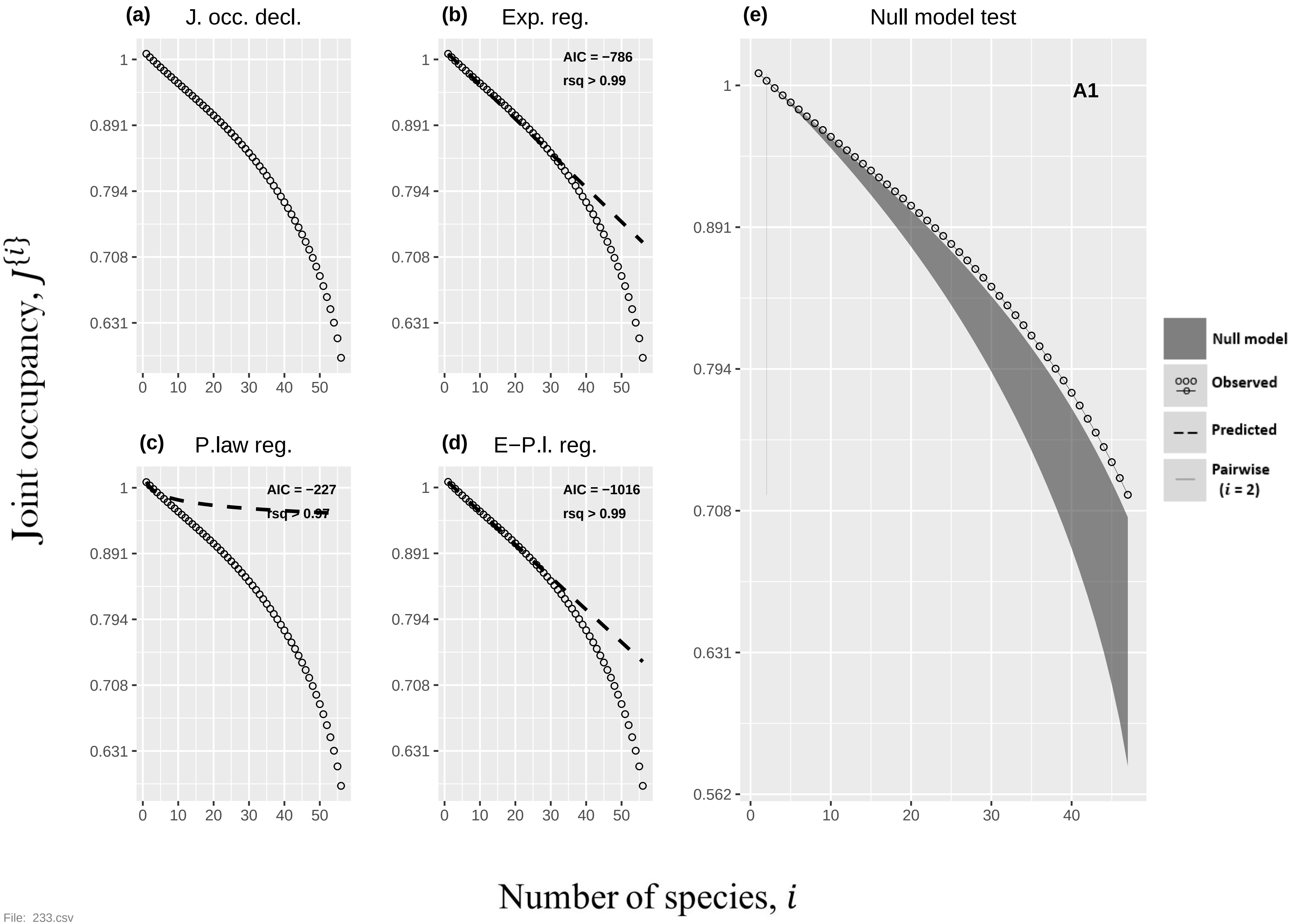

(a) J. occ. decl.
(b) Exp. reg.
AIC = −786
rsq > 0.99
(c) P.law reg.
AIC = −227
rsq > 0.97
(d) E−P.l. reg.
AIC = −1016
rsq > 0.99
(e) Null model test
A1
Null model
Observed
Predicted
Pairwise (i = 2)
Joint occupancy, J{i}
Number of species, i
1
0.891
0.794
0.708
0.631
0.562
0
10
20
30
40
50


File: 233.csv

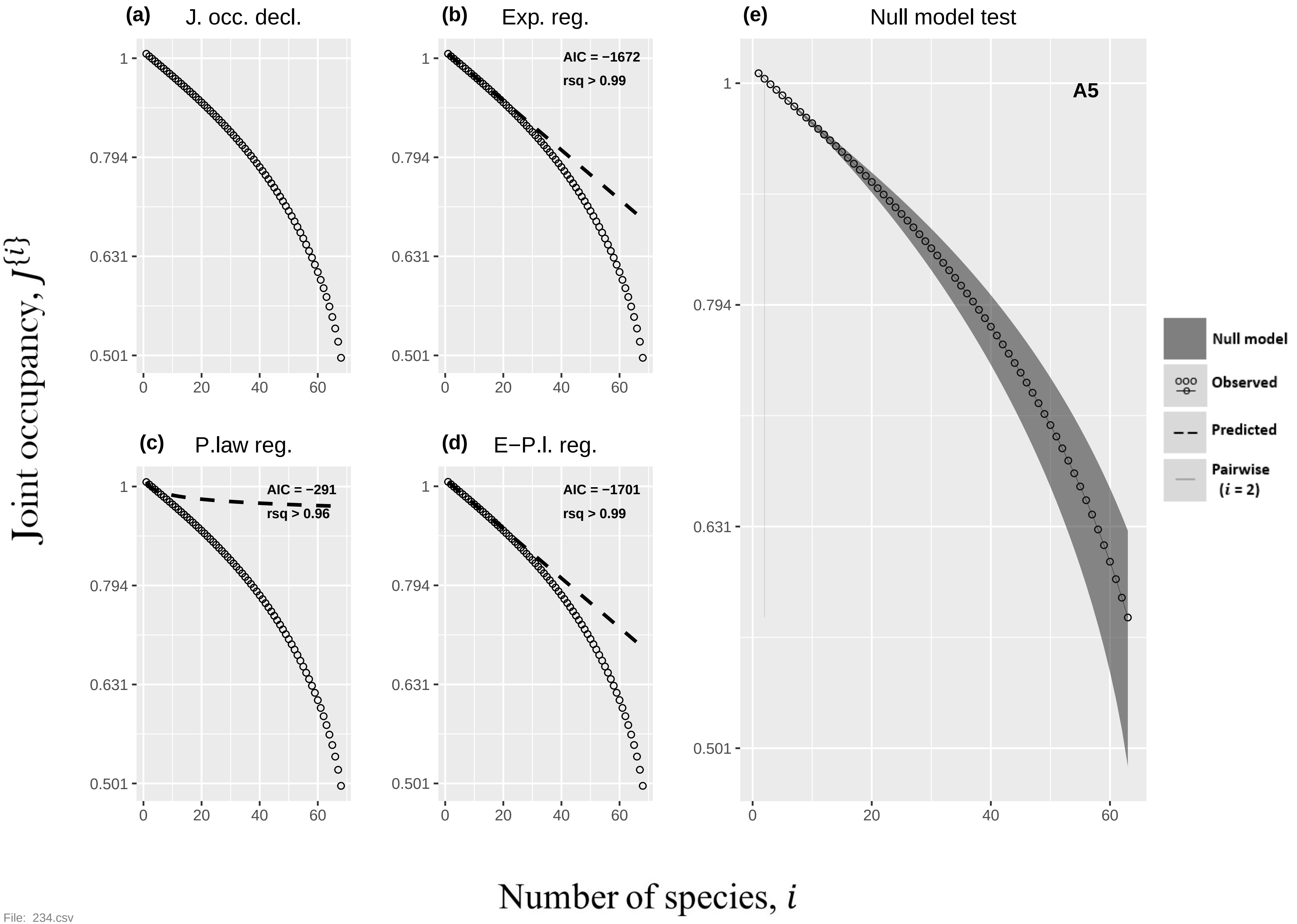

(a) J. occ. decl.
(b) Exp. reg.
AIC = −1672
rsq > 0.99
(c) P.law reg.
AIC = −291
rsq > 0.96
(d) E−P.l. reg.
AIC = −1701
rsq > 0.99
(e) Null model test
A5
Null model
Observed
Predicted
Pairwise (i = 2)
Joint occupancy, J{i}
Number of species, i
1
0.794
0.631
0.501
0
20
40
60


File: 234.csv

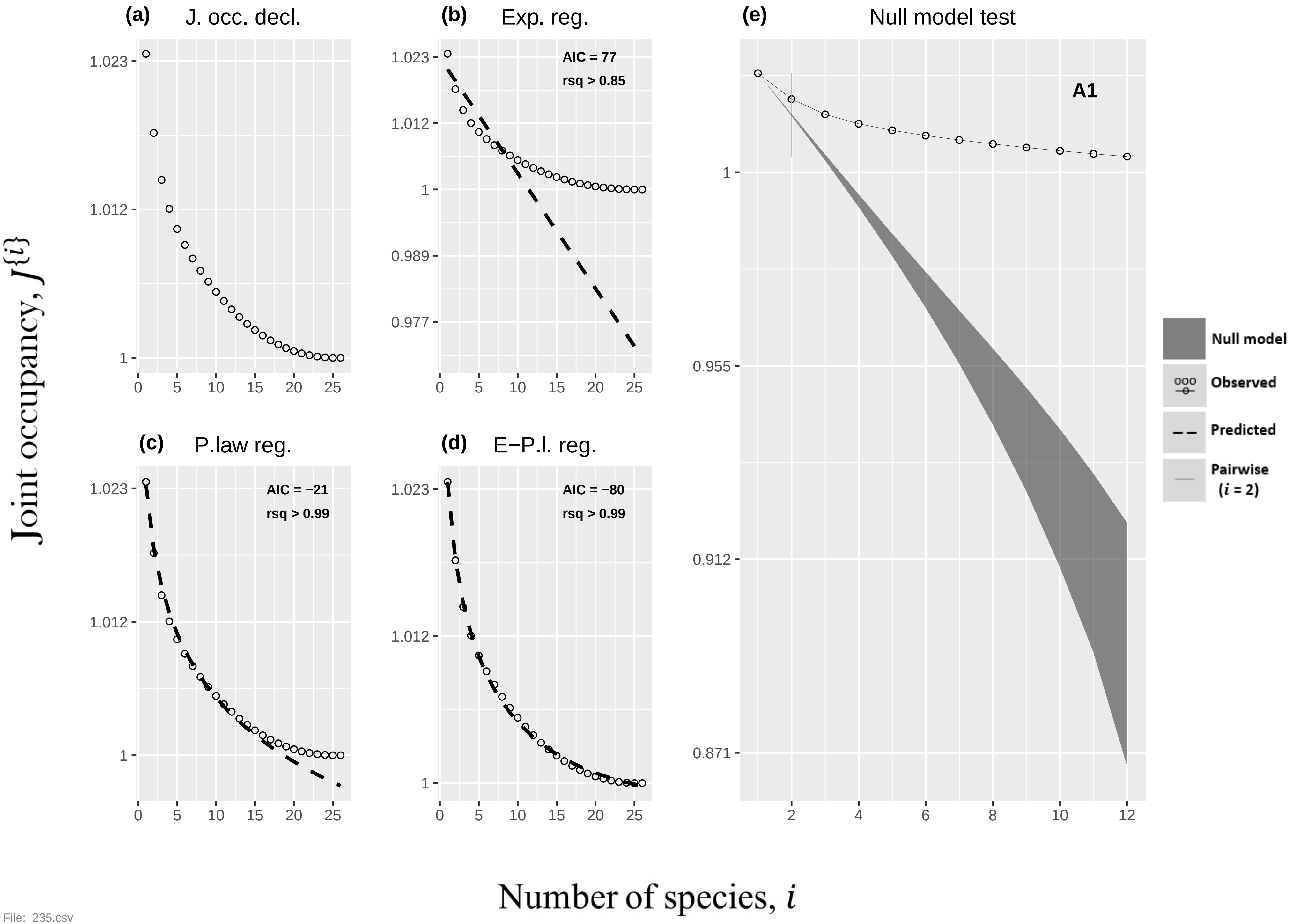

(a) J. occ. decl.
(b) Exp. reg.
AIC = 77
rsq > 0.85
(c) P.law reg.
AIC = −21
rsq > 0.99
(d) E−P.l. reg.
AIC = −80
rsq > 0.99
(e) Null model test
A1
Null model
Observed
Predicted
Pairwise (i = 2)
Joint occupancy, J{i}
Number of species, i


File: 235.csv

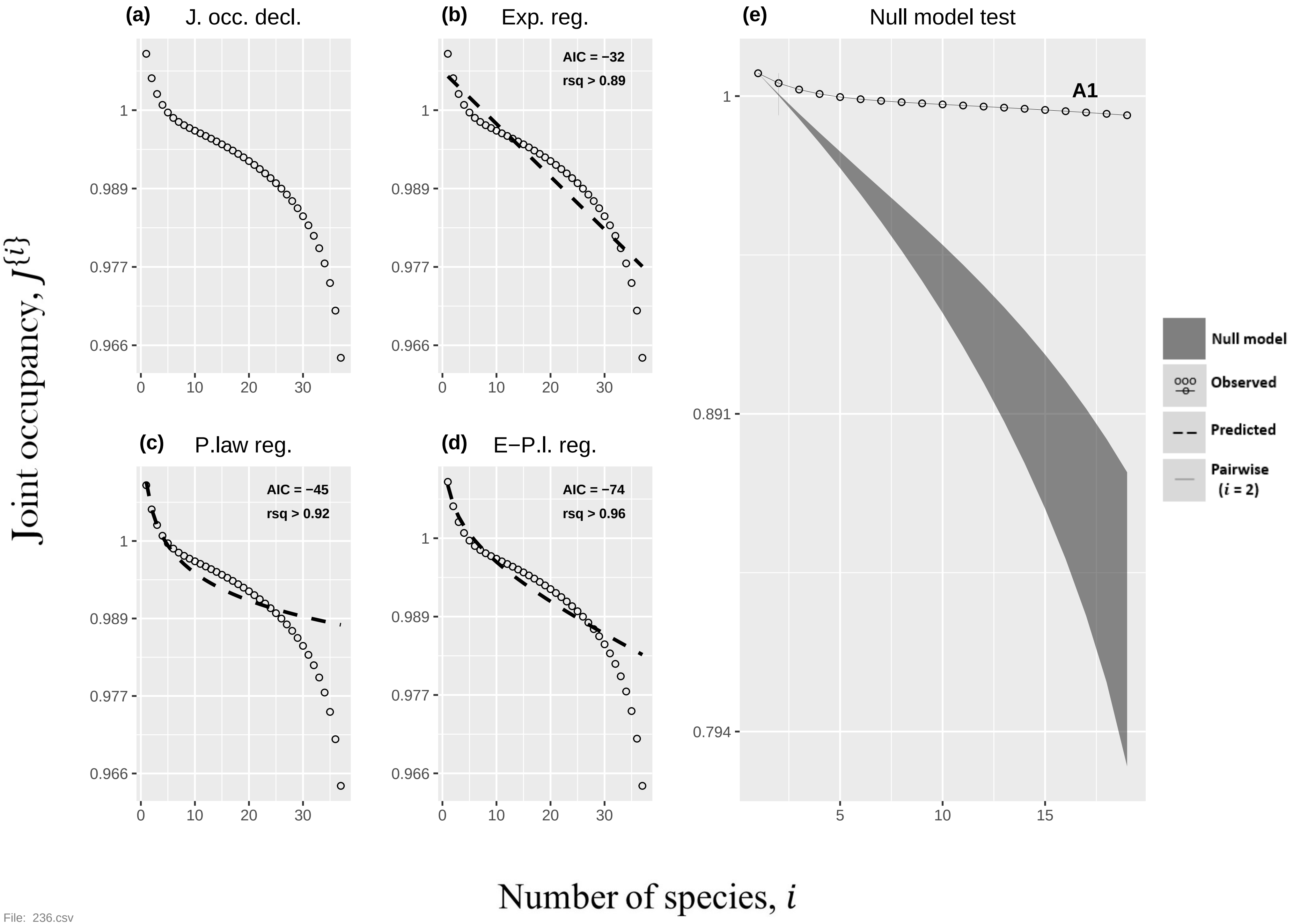

(a) J. occ. decl.
(b) Exp. reg.
AIC = −32
rsq > 0.89
(c) P.law reg.
AIC = −45
rsq > 0.92
(d) E−P.l. reg.
AIC = −74
rsq > 0.96
(e) Null model test
A1
Null model
Observed
Predicted
Pairwise (i = 2)
Joint occupancy, J{i}
Number of species, i
1
0.989
0.977
0.966
0.891
0.794
0
10
20
30
5
15


File: 236.csv

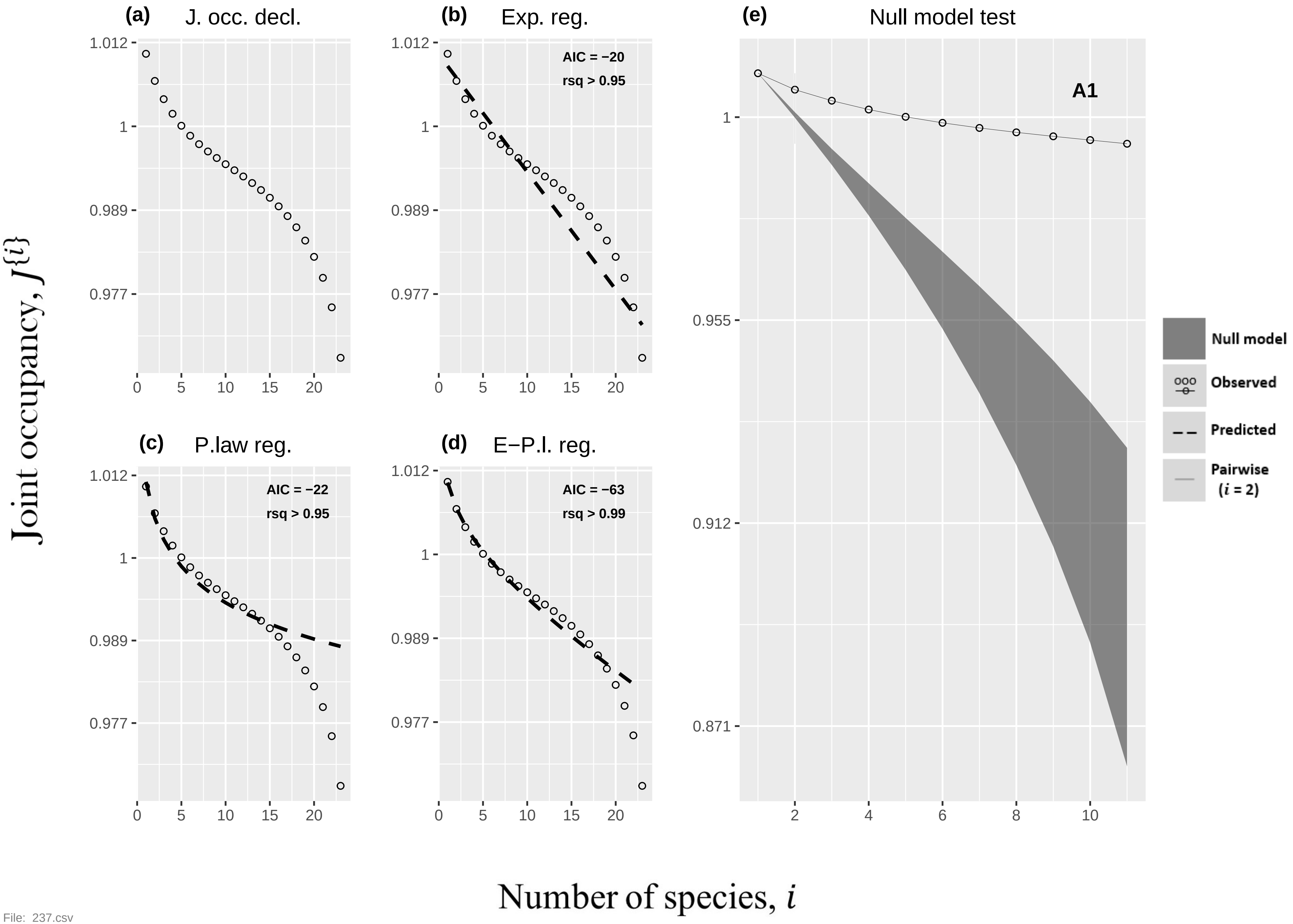

(a)
J. occ. decl.
(b)
Exp. reg.
AIC = −20
rsq > 0.95
(c)
P.law reg.
AIC = −22
rsq > 0.95
(d)
E−P.l. reg.
AIC = −63
rsq > 0.99
(e)
Null model test
A1
Null model
Observed
Predicted
Pairwise ($i$ = 2)
1.012
1
0.989
0.977
0
5
10
15
20
1
0.955
0.912
0.871
2
4
6
8
10
Joint occupancy, $J^{\{i\}}$
Number of species, $i$


File: 237.csv

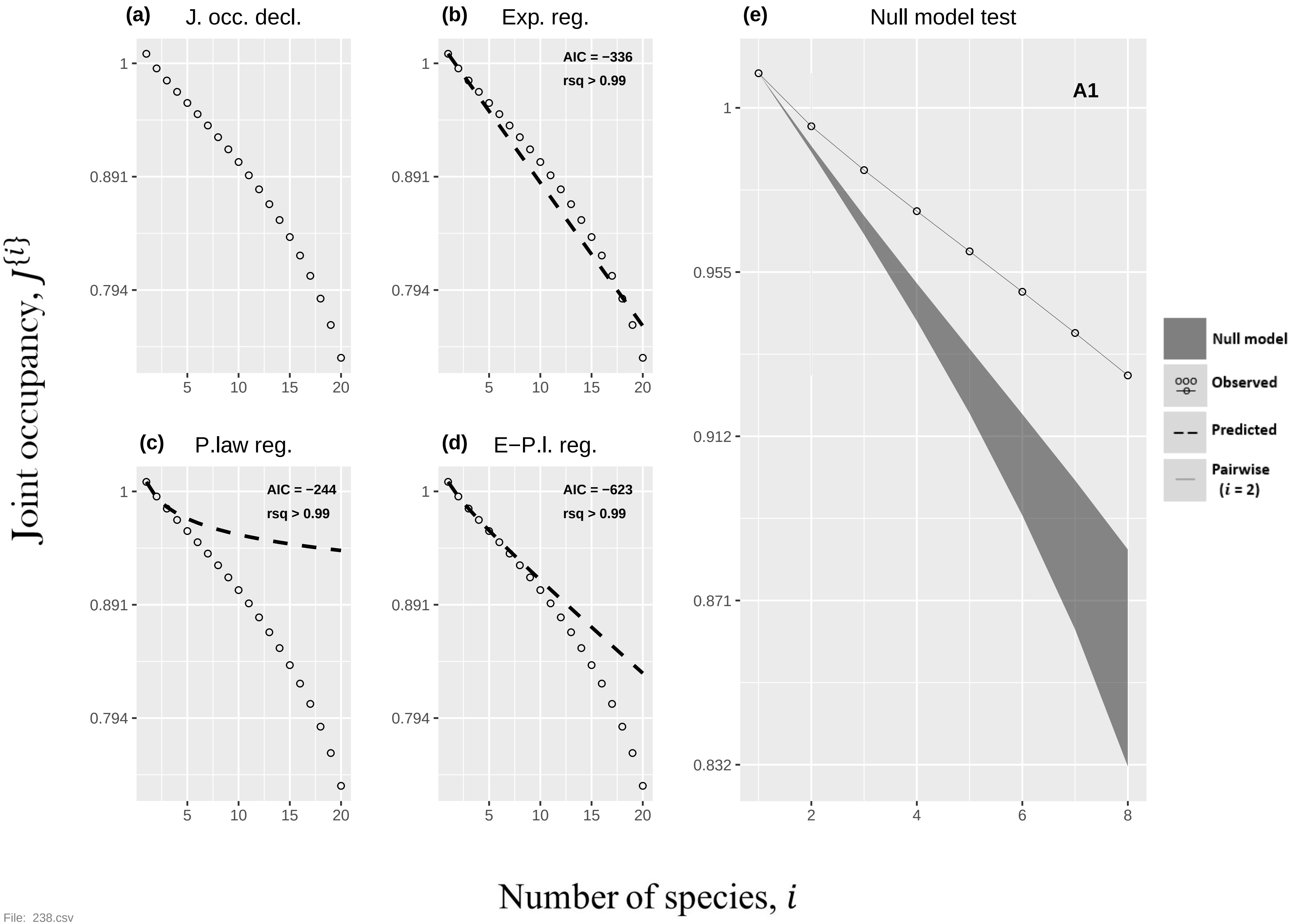

(a) J. occ. decl.
(b) Exp. reg.
AIC = −336
rsq > 0.99
(c) P.law reg.
AIC = −244
rsq > 0.99
(d) E−P.l. reg.
AIC = −623
rsq > 0.99
(e) Null model test
A1
Null model
Observed
Predicted
Pairwise (i = 2)
Joint occupancy, J{i}
Number of species, i


File: 238.csv

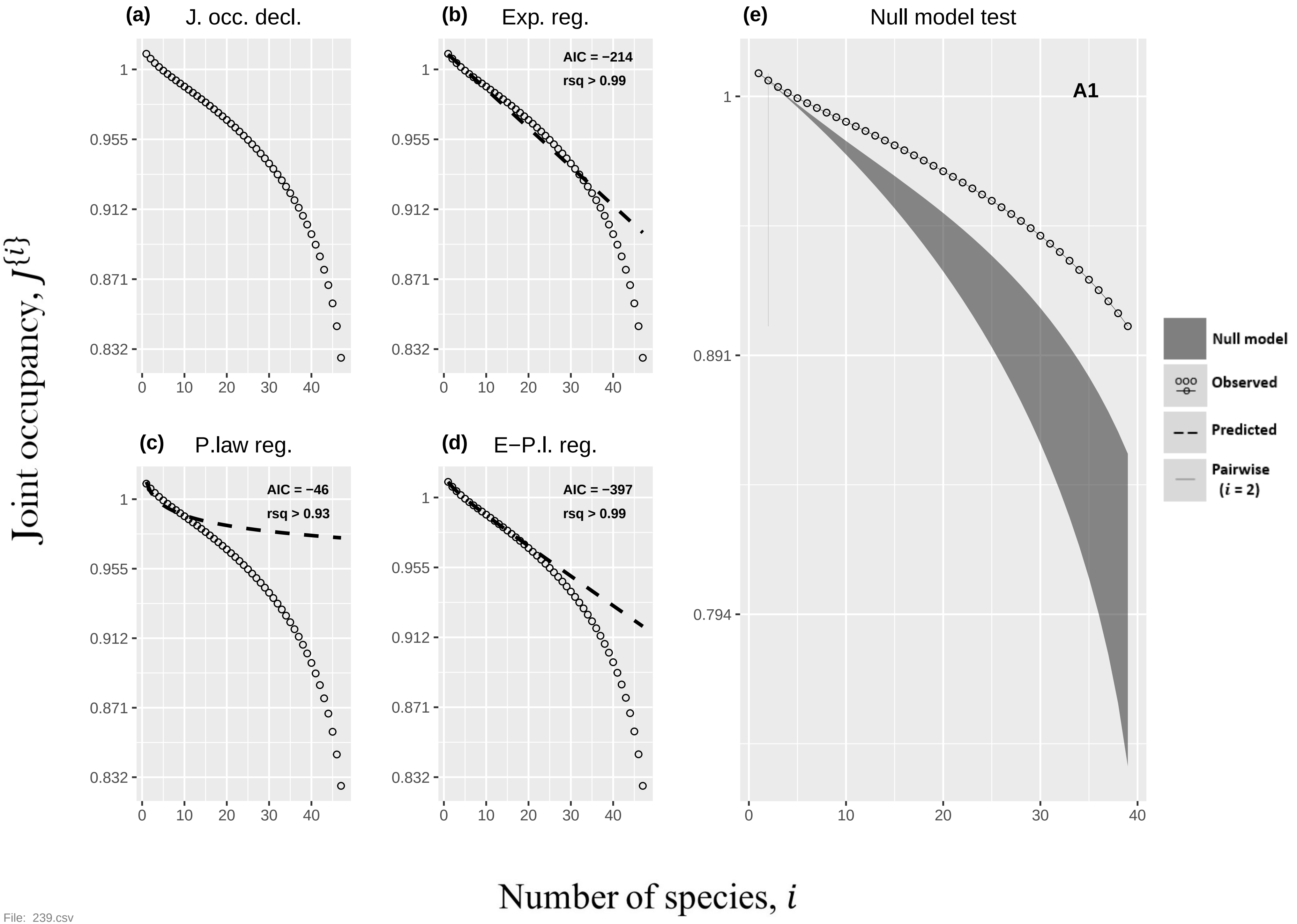

(a) J. occ. decl.
(b) Exp. reg.
AIC = −214
rsq > 0.99
(c) P.law reg.
AIC = −46
rsq > 0.93
(d) E−P.l. reg.
AIC = −397
rsq > 0.99
(e) Null model test
A1
Null model
Observed
Predicted
Pairwise
(i = 2)
Joint occupancy, J{i}
Number of species, i


File: 239.csv

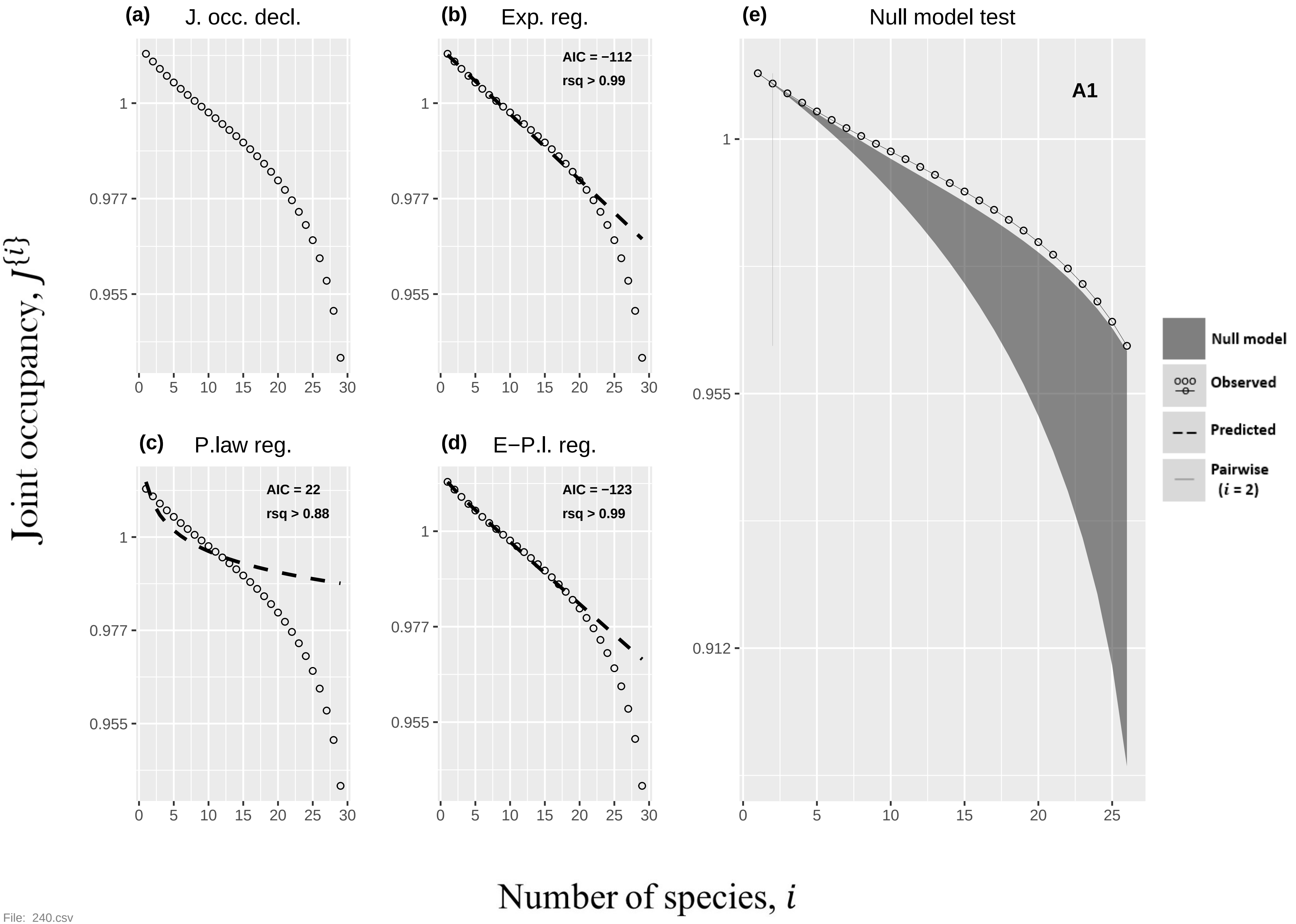

(a)
J. occ. decl.
(b)
Exp. reg.
AIC = −112
rsq > 0.99
(e)
Null model test
A1
(c)
P.law reg.
AIC = 22
rsq > 0.88
(d)
E−P.l. reg.
AIC = −123
rsq > 0.99
Null model
Observed
Predicted
Pairwise
(i = 2)
Joint occupancy, J{i}
Number of species, i


File: 240.csv

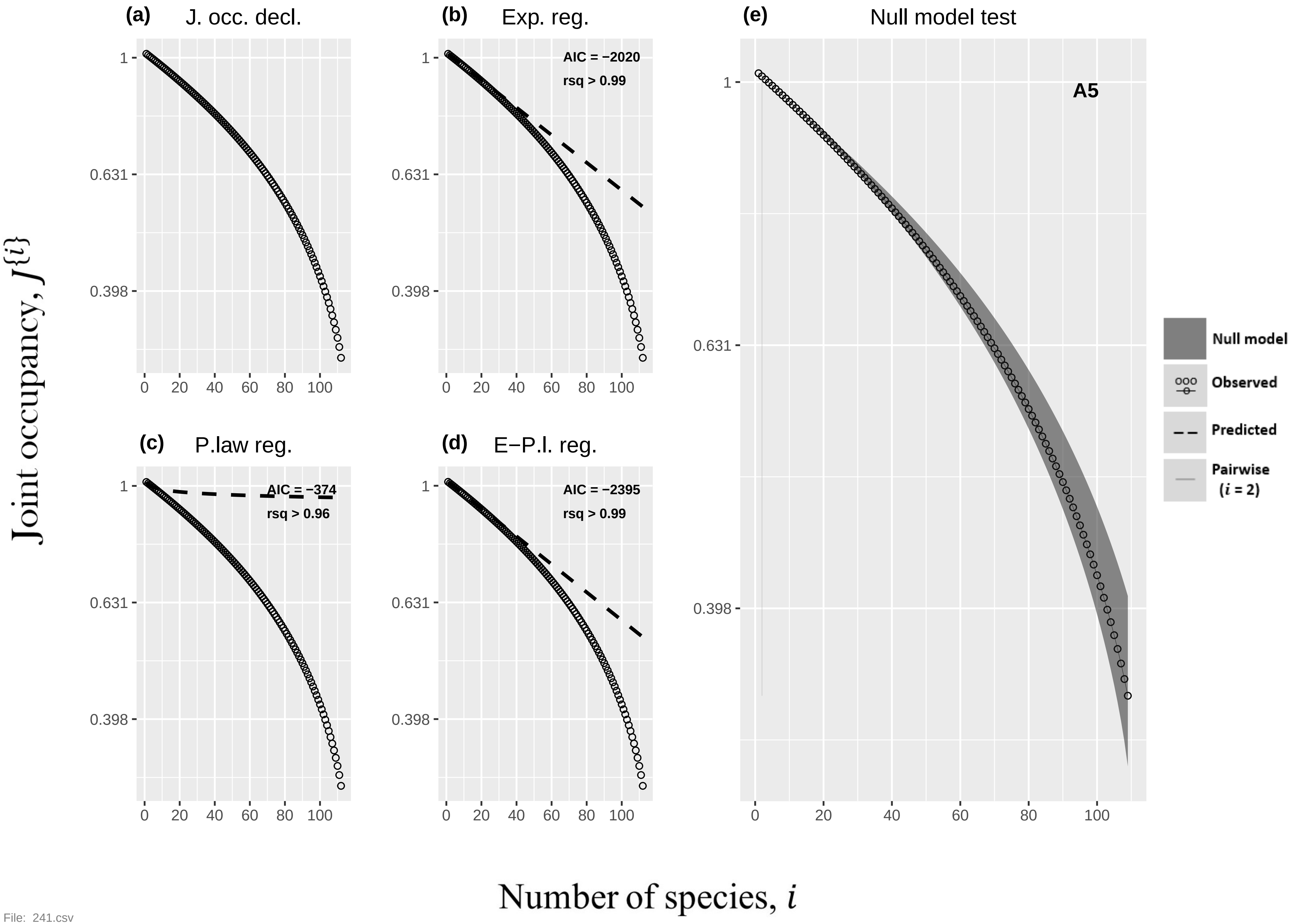

(a)
J. occ. decl.
(b)
Exp. reg.
AIC = −2020
rsq > 0.99
(c)
P.law reg.
AIC = −374
rsq > 0.96
(d)
E−P.l. reg.
AIC = −2395
rsq > 0.99
(e)
Null model test
A5
Null model
Observed
Predicted
Pairwise
(i = 2)
Joint occupancy, J{i}
Number of species, i
1
0.631
0.398
0
20
40
60
80
100

File: 241.csv

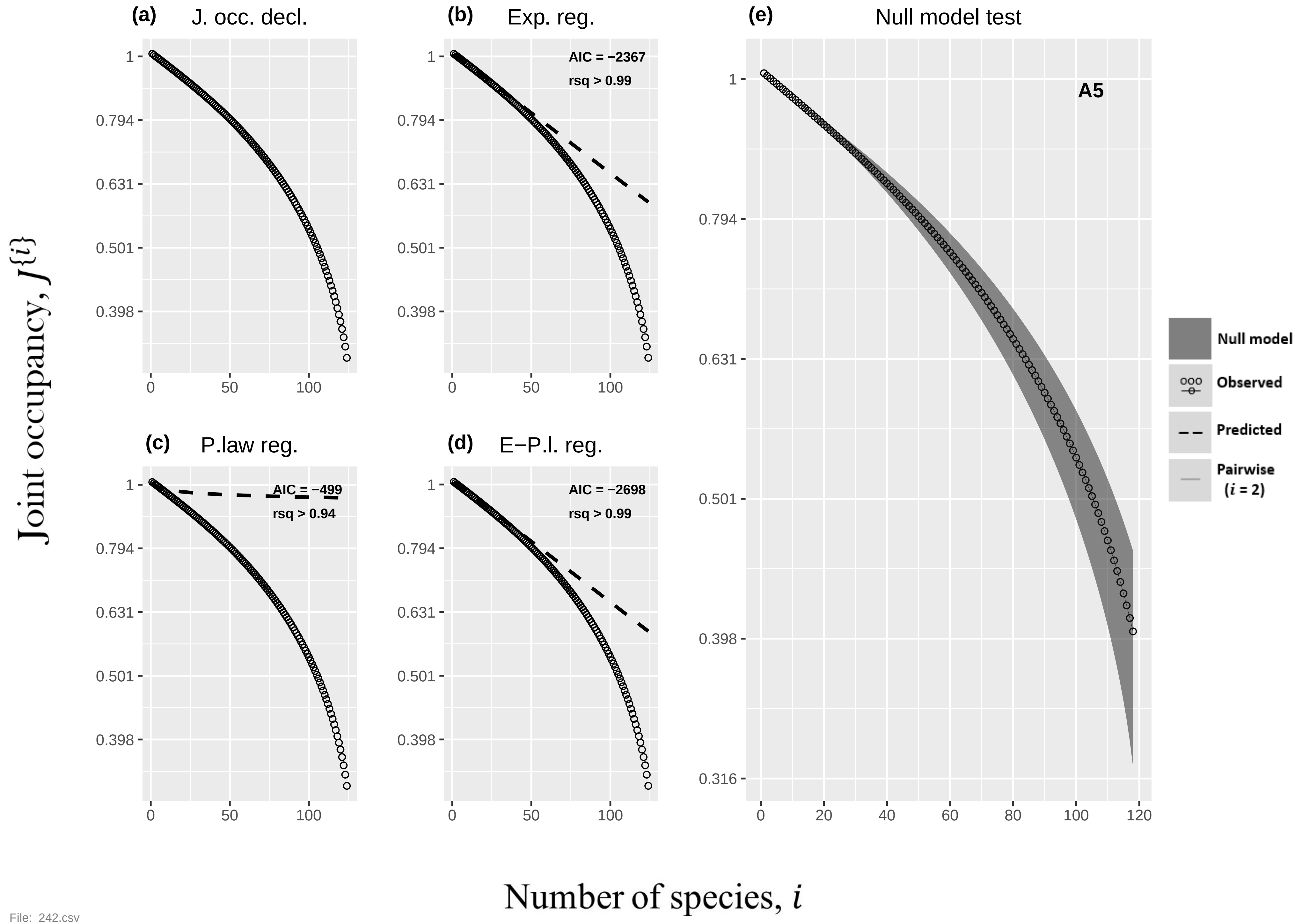

(a) J. occ. decl.
(b) Exp. reg.
AIC = −2367
rsq > 0.99
(c) P.law reg.
AIC = −499
rsq > 0.94
(d) E−P.l. reg.
AIC = −2698
rsq > 0.99
(e) Null model test
A5
Null model
Observed
Predicted
Pairwise (i = 2)
Joint occupancy, J{i}
Number of species, i
1
0.794
0.631
0.501
0.398
0.316
0
50
100
20
40
60
80
120


File: 242.csv

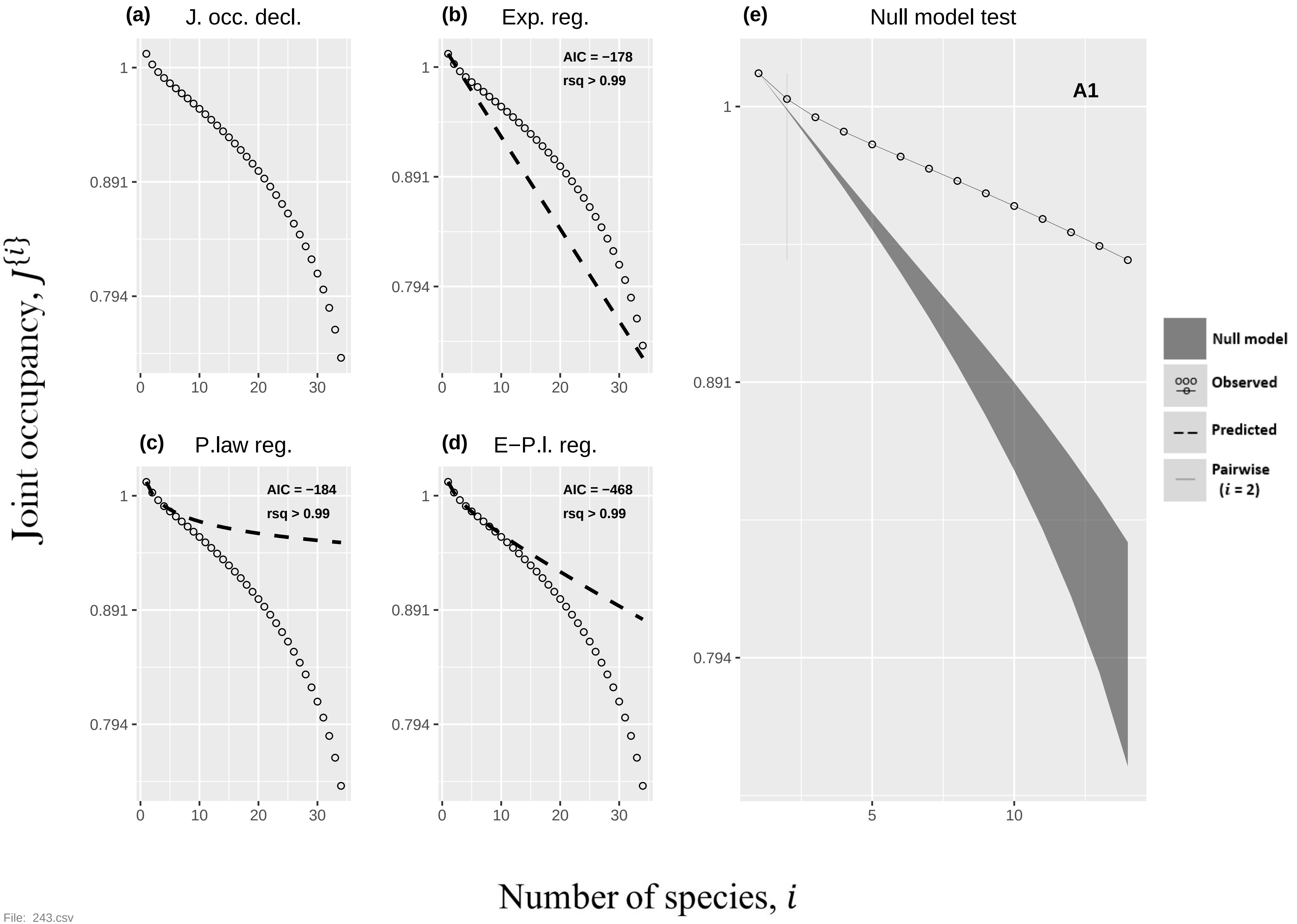

(a)
J. occ. decl.
(b)
Exp. reg.
AIC = −178
rsq > 0.99
(c)
P.law reg.
AIC = −184
rsq > 0.99
(d)
E−P.l. reg.
AIC = −468
rsq > 0.99
(e)
Null model test
A1
Null model
Observed
Predicted
Pairwise
($i$ = 2)
1
0.891
0.794
0
10
20
30
5
Joint occupancy, $J\{i\}$
Number of species, $i$


File: 243.csv

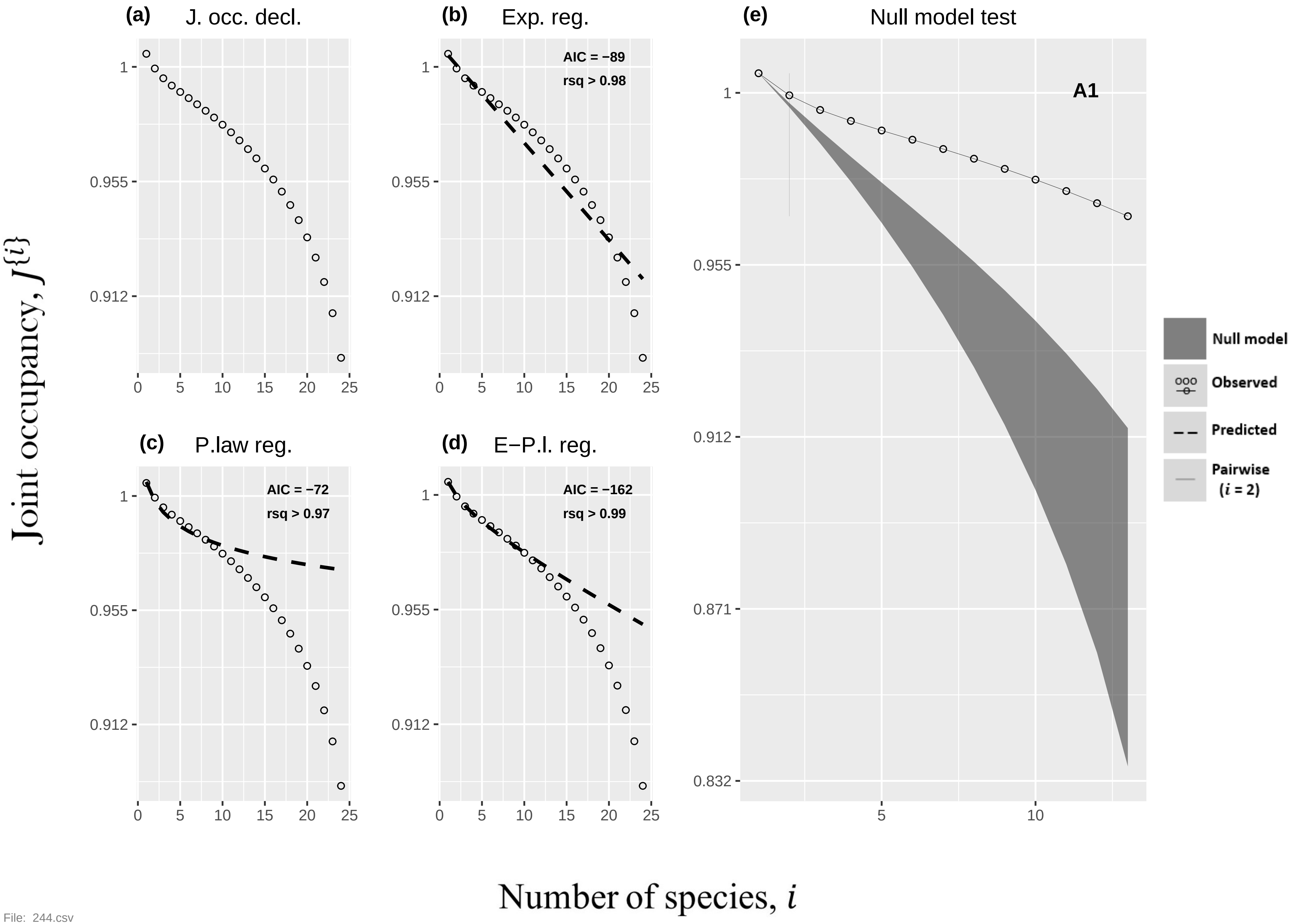

(a)
J. occ. decl.
(b)
Exp. reg.
AIC = −89
rsq > 0.98
(c)
P.law reg.
AIC = −72
rsq > 0.97
(d)
E−P.l. reg.
AIC = −162
rsq > 0.99
(e)
Null model test
A1
Null model
Observed
Predicted
Pairwise
(i = 2)
Joint occupancy, J{i}
Number of species, i
1
0.955
0.912
0.871
0.832
0
5
10
15
20
25


File: 244.csv

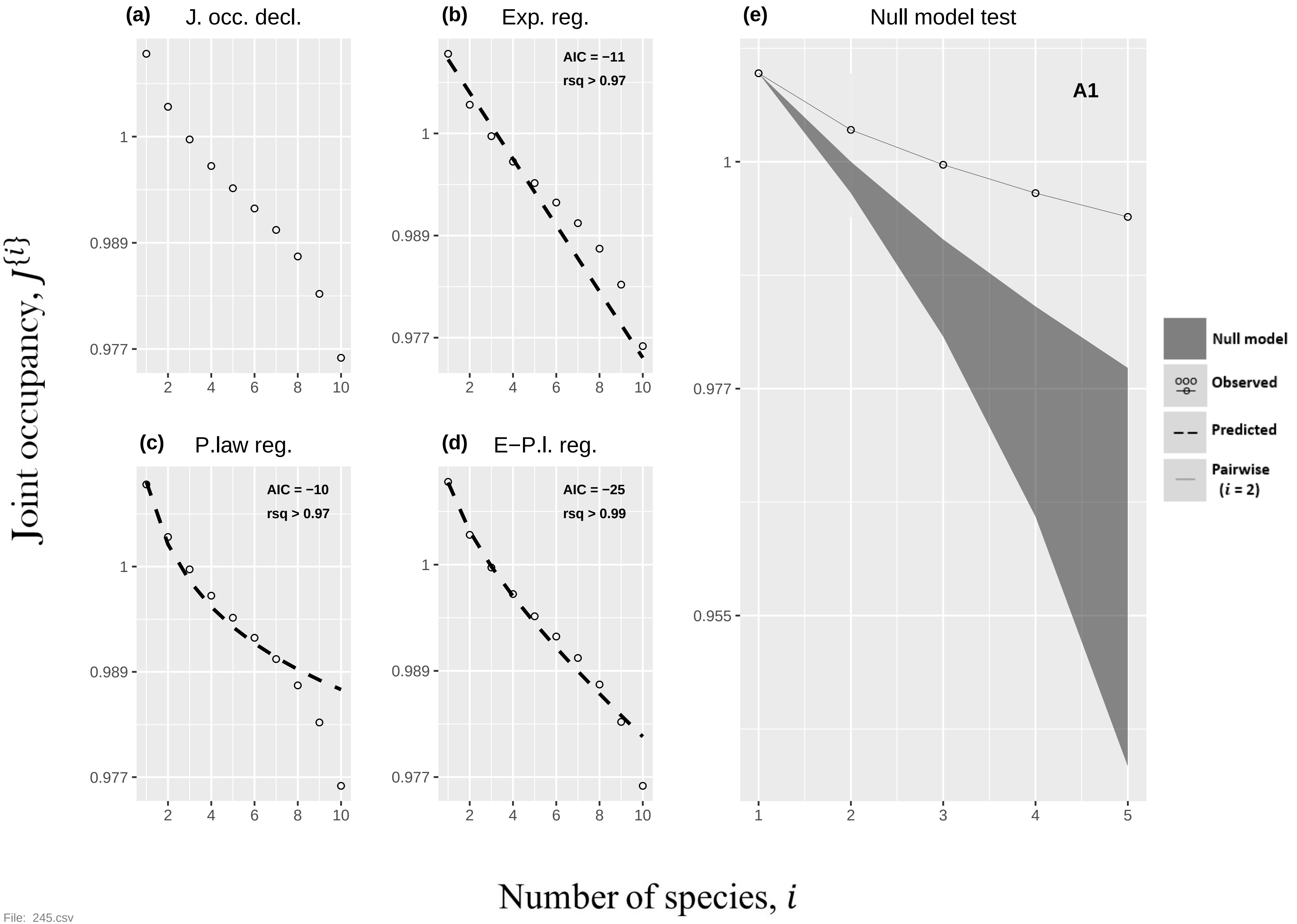

(a) J. occ. decl.
(b) Exp. reg.
AIC = −11
rsq > 0.97
(c) P.law reg.
AIC = −10
rsq > 0.97
(d) E−P.l. reg.
AIC = −25
rsq > 0.99
(e) Null model test
A1
Null model
Observed
Predicted
Pairwise (i = 2)
1
0.989
0.977
0.955
2
4
6
8
10
1
2
3
4
5
Joint occupancy, J{i}
Number of species, i


File: 245.csv

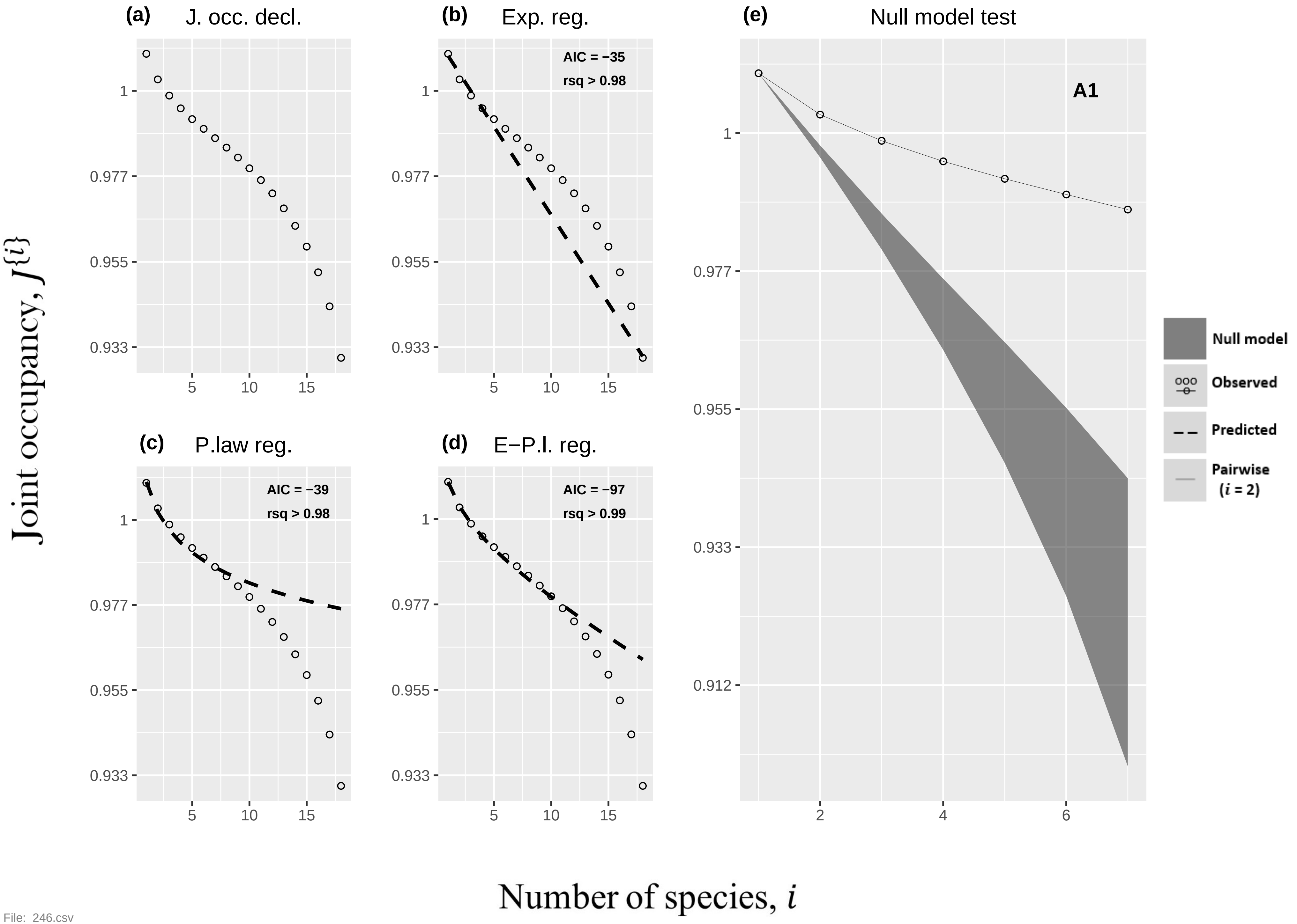

(a) J. occ. decl.
(b) Exp. reg.
AIC = −35
rsq > 0.98
(c) P.law reg.
AIC = −39
rsq > 0.98
(d) E−P.l. reg.
AIC = −97
rsq > 0.99
(e) Null model test
A1
Null model
Observed
Predicted
Pairwise (i = 2)
Joint occupancy, J{i}
Number of species, i


File: 246.csv

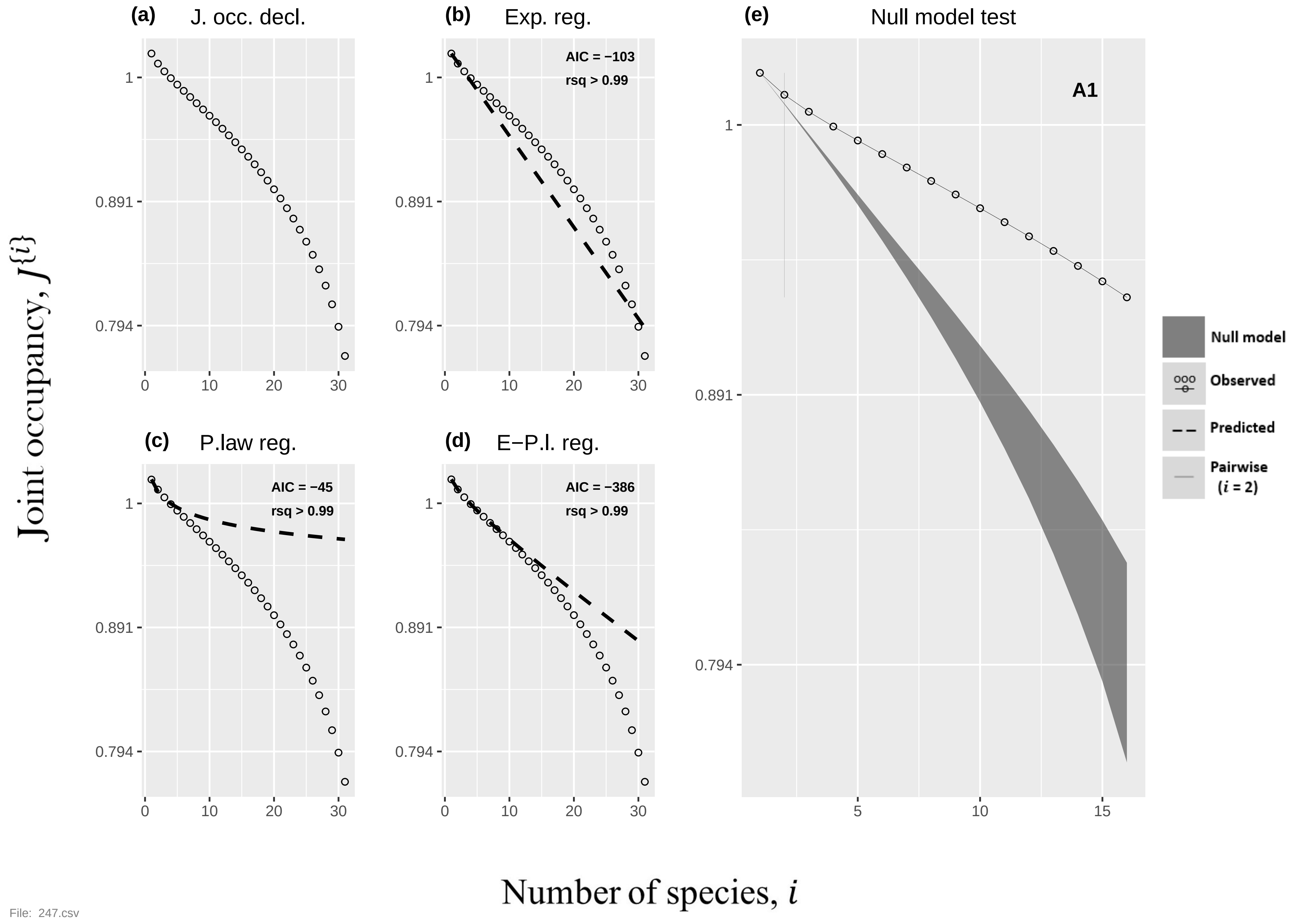
(a) J. occ. decl.
(b) Exp. reg.
AIC = −103
rsq > 0.99
(c) P.law reg.
AIC = −45
rsq > 0.99
(d) E−P.l. reg.
AIC = −386
rsq > 0.99
(e) Null model test
A1
Null model
Observed
Predicted
Pairwise (i = 2)
1
0.891
0.794
0
10
20
30
5
15
Joint occupancy, J{i}
Number of species, i
File: 247.csv

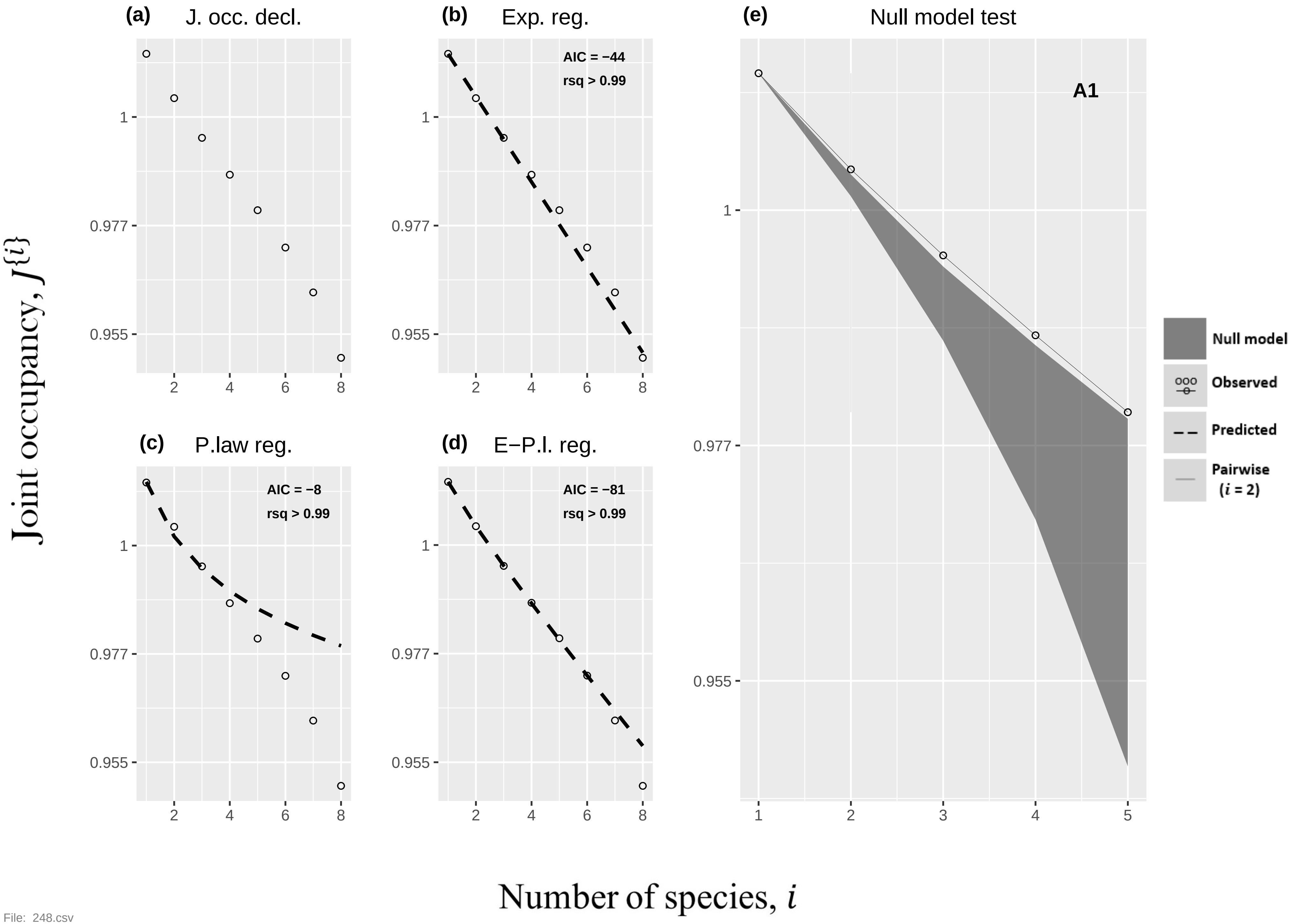

(a) J. occ. decl.
(b) Exp. reg.
AIC = −44
rsq > 0.99
(e) Null model test
A1
Null model
Observed
Predicted
Pairwise ($i$ = 2)
(c) P.law reg.
AIC = −8
rsq > 0.99
(d) E−P.l. reg.
AIC = −81
rsq > 0.99
Joint occupancy, $J^{\{i\}}$
Number of species, $i$

File: 248.csv

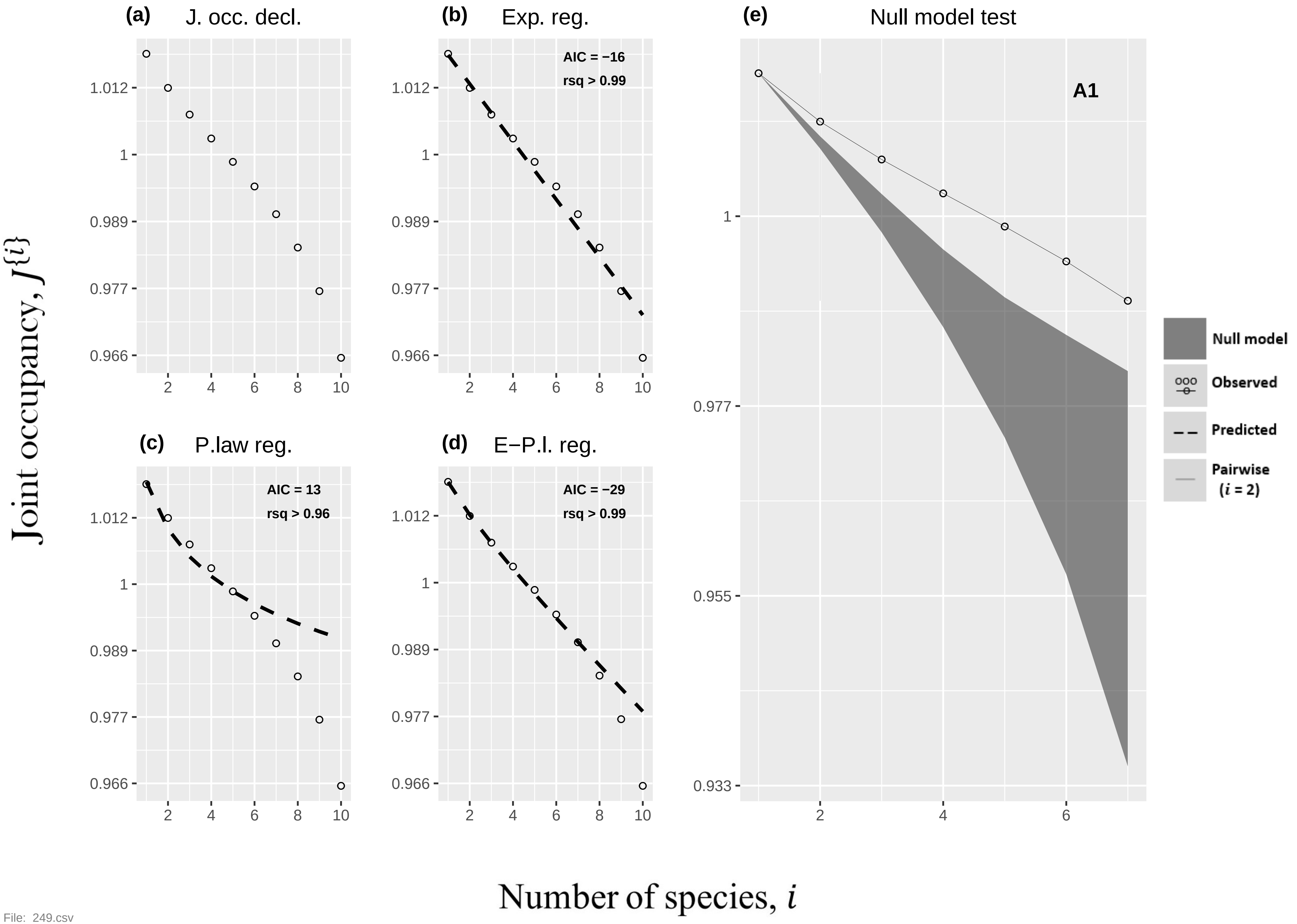

(a)
J. occ. decl.
(b)
Exp. reg.
AIC = −16
rsq > 0.99
(c)
P.law reg.
AIC = 13
rsq > 0.96
(d)
E−P.l. reg.
AIC = −29
rsq > 0.99
(e)
Null model test
A1
1.012
1
0.989
0.977
0.966
0.955
0.933
2
4
6
8
10
Null model
Observed
Predicted
Pairwise
(i = 2)
Joint occupancy, J{i}
Number of species, i


File: 249.csv

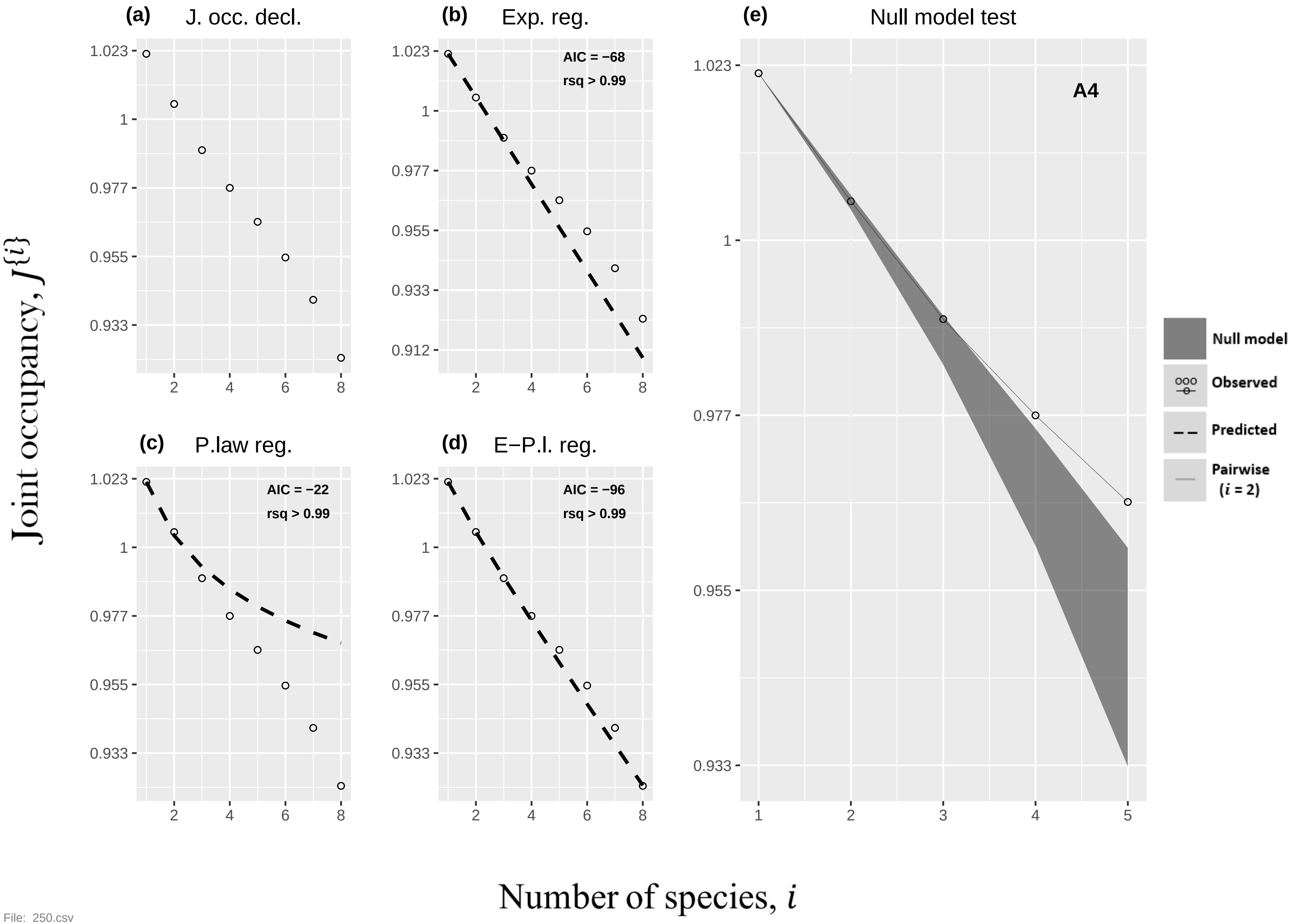

(a)
J. occ. decl.
(b)
Exp. reg.
AIC = −68
rsq > 0.99
(c)
P.law reg.
AIC = −22
rsq > 0.99
(d)
E−P.l. reg.
AIC = −96
rsq > 0.99
(e)
Null model test
A4
Null model
Observed
Predicted
Pairwise (i = 2)
Joint occupancy, J{i}
Number of species, i
1.023
1
0.977
0.955
0.933
0.912
2
4
6
8
1
3
5


File: 250.csv

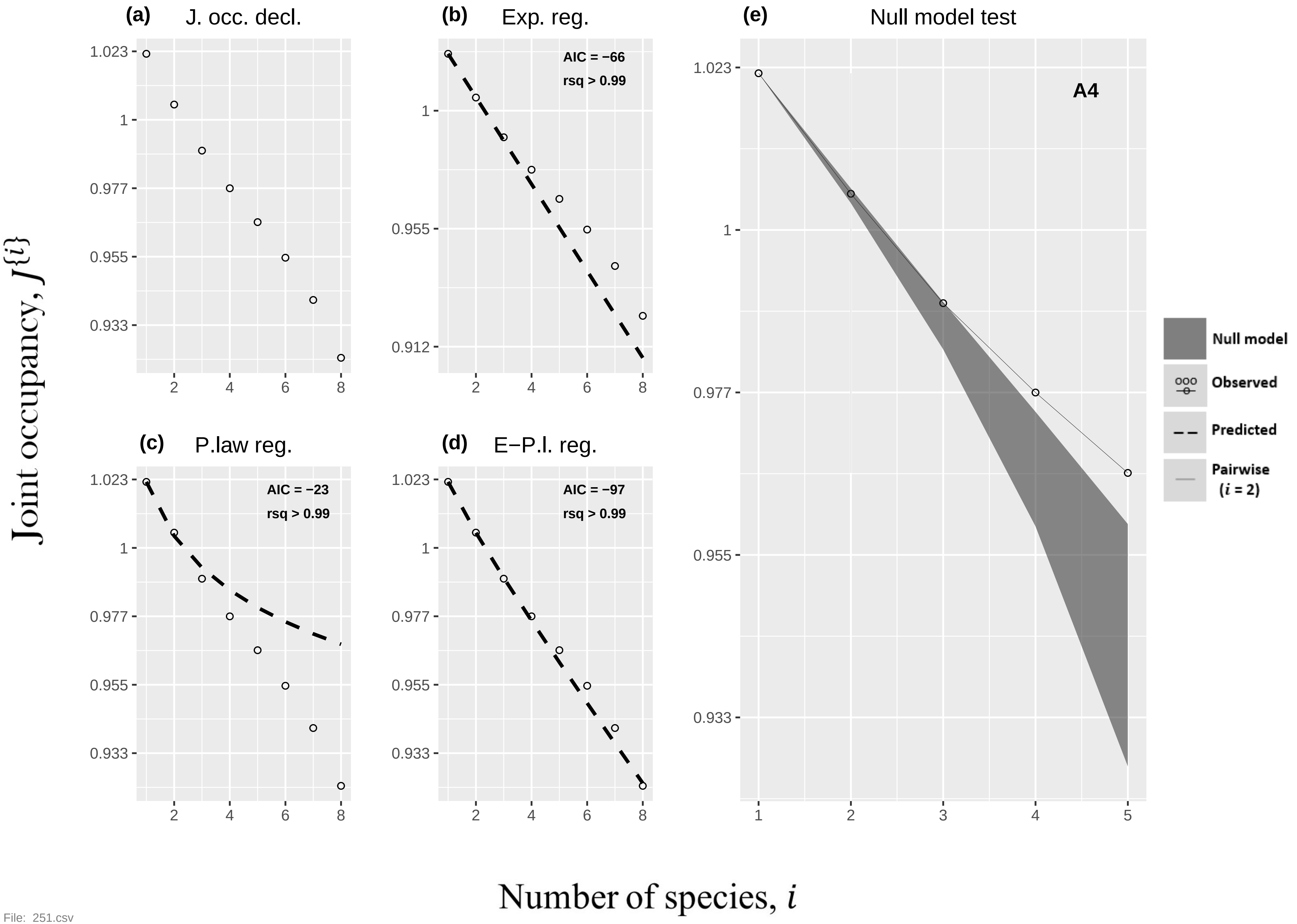
(a) J. occ. decl.
(b) Exp. reg.
AIC = −66
rsq > 0.99
(c) P.law reg.
AIC = −23
rsq > 0.99
(d) E−P.l. reg.
AIC = −97
rsq > 0.99
(e) Null model test
A4
Null model
Observed
Predicted
Pairwise ($i$ = 2)
1.023
1
0.977
0.955
0.933
0.912
1
2
3
4
5
6
8
Joint occupancy, $J\{i\}$
Number of species, $i$

File: 251.csv

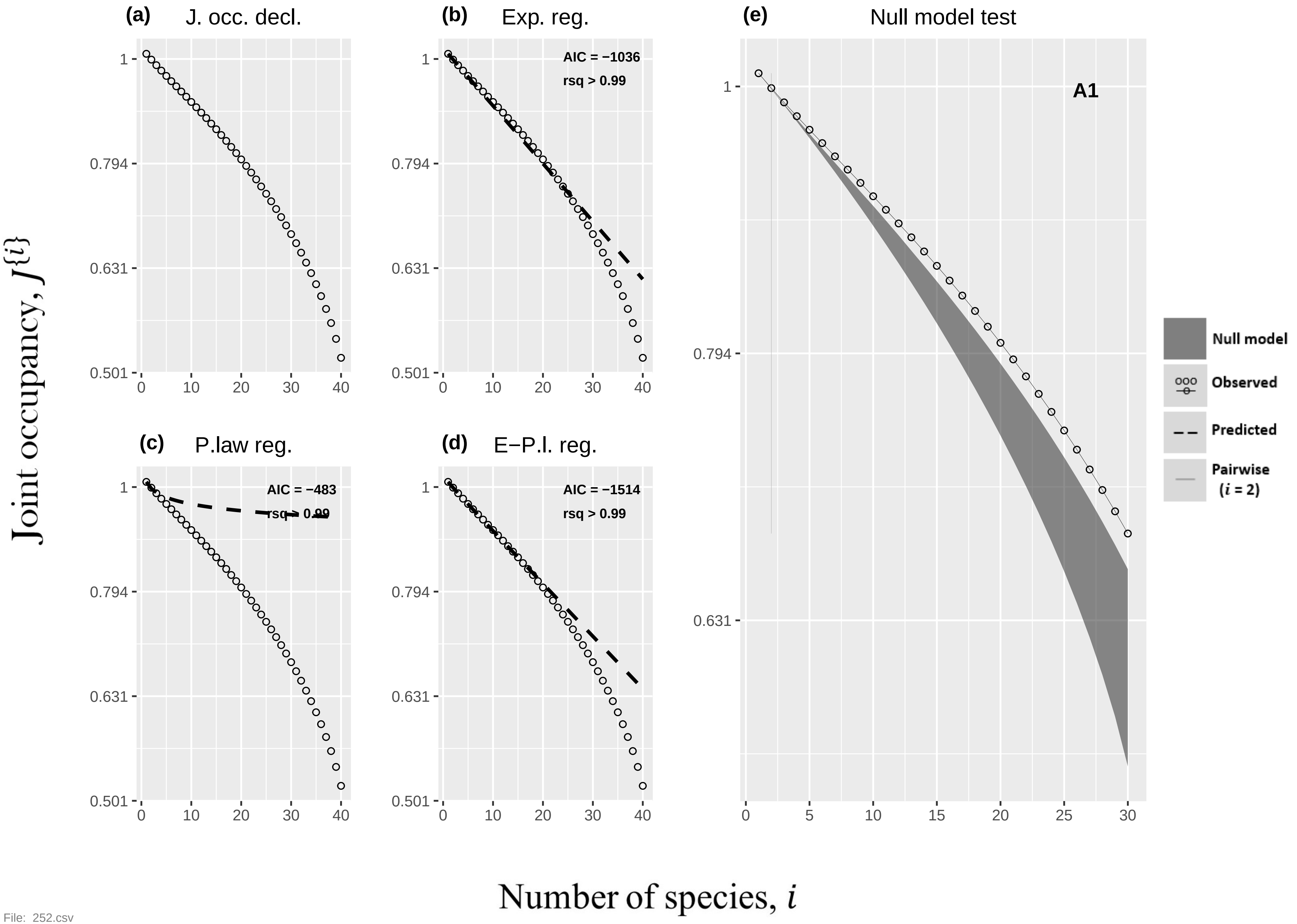

(a) J. occ. decl.
(b) Exp. reg.
AIC = −1036
rsq > 0.99
(c) P.law reg.
AIC = −483
rsq > 0.99
(d) E−P.l. reg.
AIC = −1514
rsq > 0.99
(e) Null model test
A1
Null model
Observed
Predicted
Pairwise ($i$ = 2)
Joint occupancy, $J^{\{i\}}$
Number of species, $i$


File: 252.csv

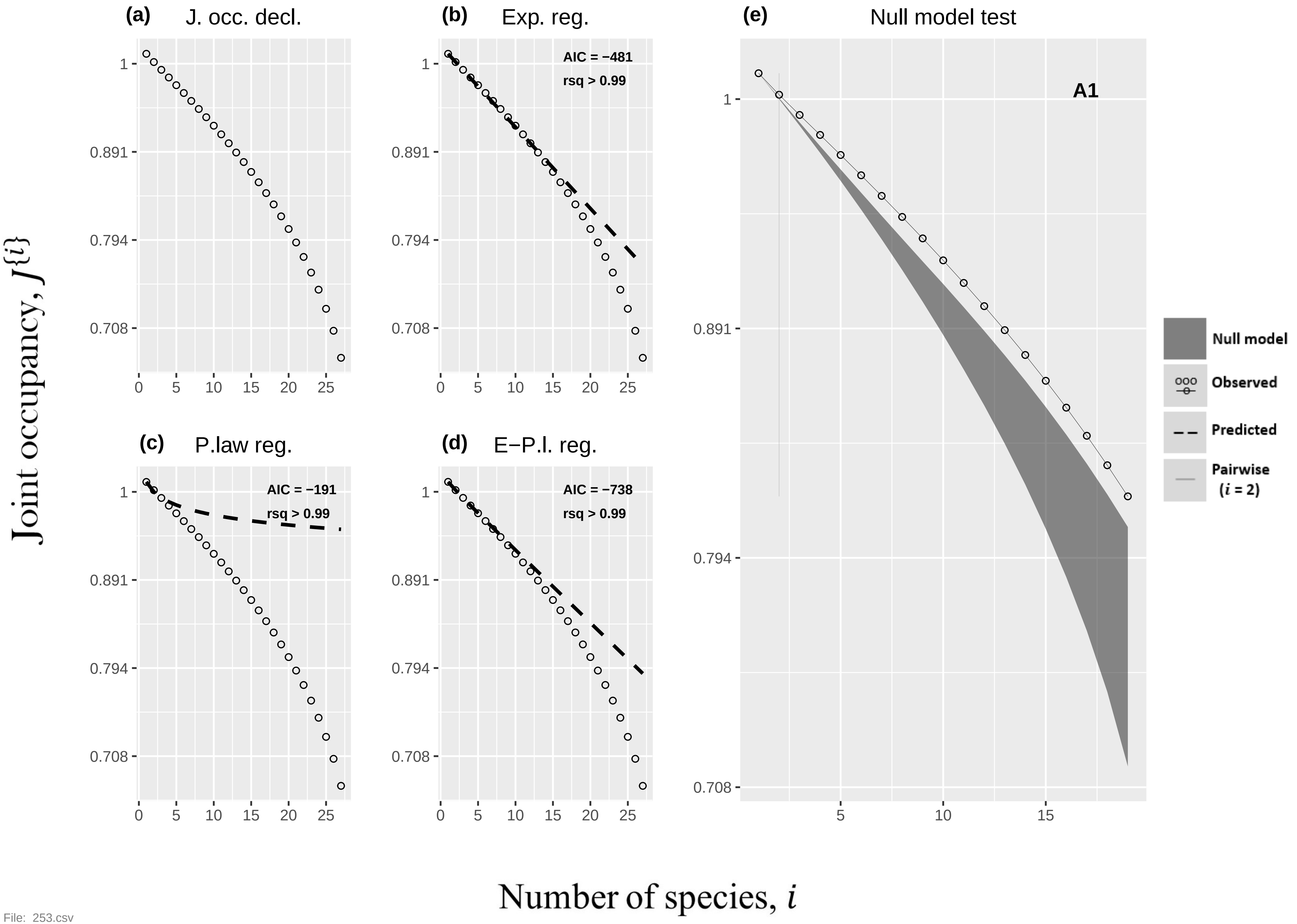
(a) J. occ. decl.
(b) Exp. reg.
AIC = −481
rsq > 0.99
(c) P.law reg.
AIC = −191
rsq > 0.99
(d) E−P.l. reg.
AIC = −738
rsq > 0.99
(e) Null model test
A1
Null model
Observed
Predicted
Pairwise
(i = 2)
1
0.891
0.794
0.708
0
5
10
15
20
25
Joint occupancy, J{i}
Number of species, i

File: 253.csv

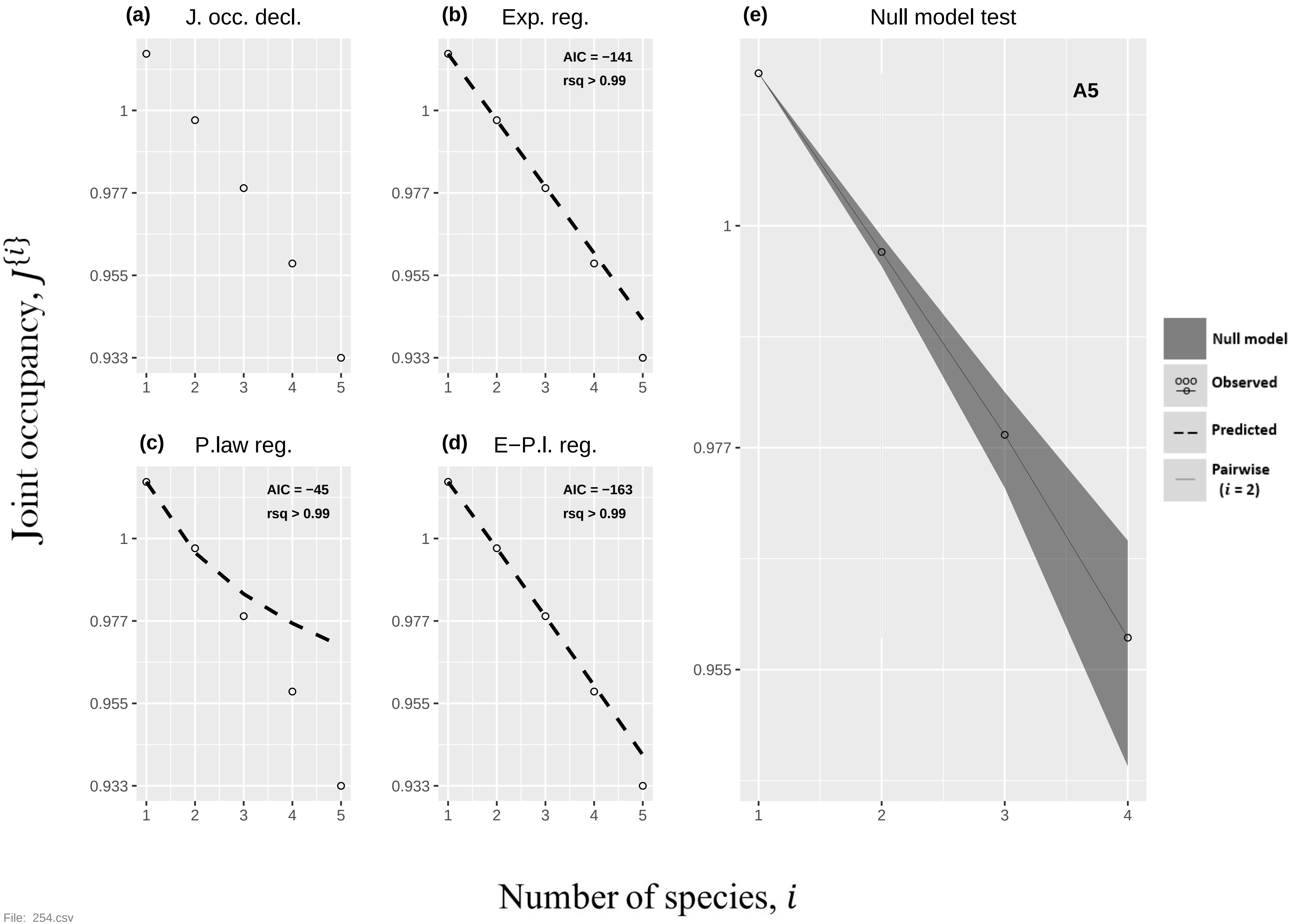

(a) J. occ. decl.
(b) Exp. reg.
AIC = −141
rsq > 0.99
(c) P.law reg.
AIC = −45
rsq > 0.99
(d) E−P.l. reg.
AIC = −163
rsq > 0.99
(e) Null model test
A5
Null model
Observed
Predicted
Pairwise ($i$ = 2)
Joint occupancy, $J^{\{i\}}$
Number of species, $i$

File: 254.csv

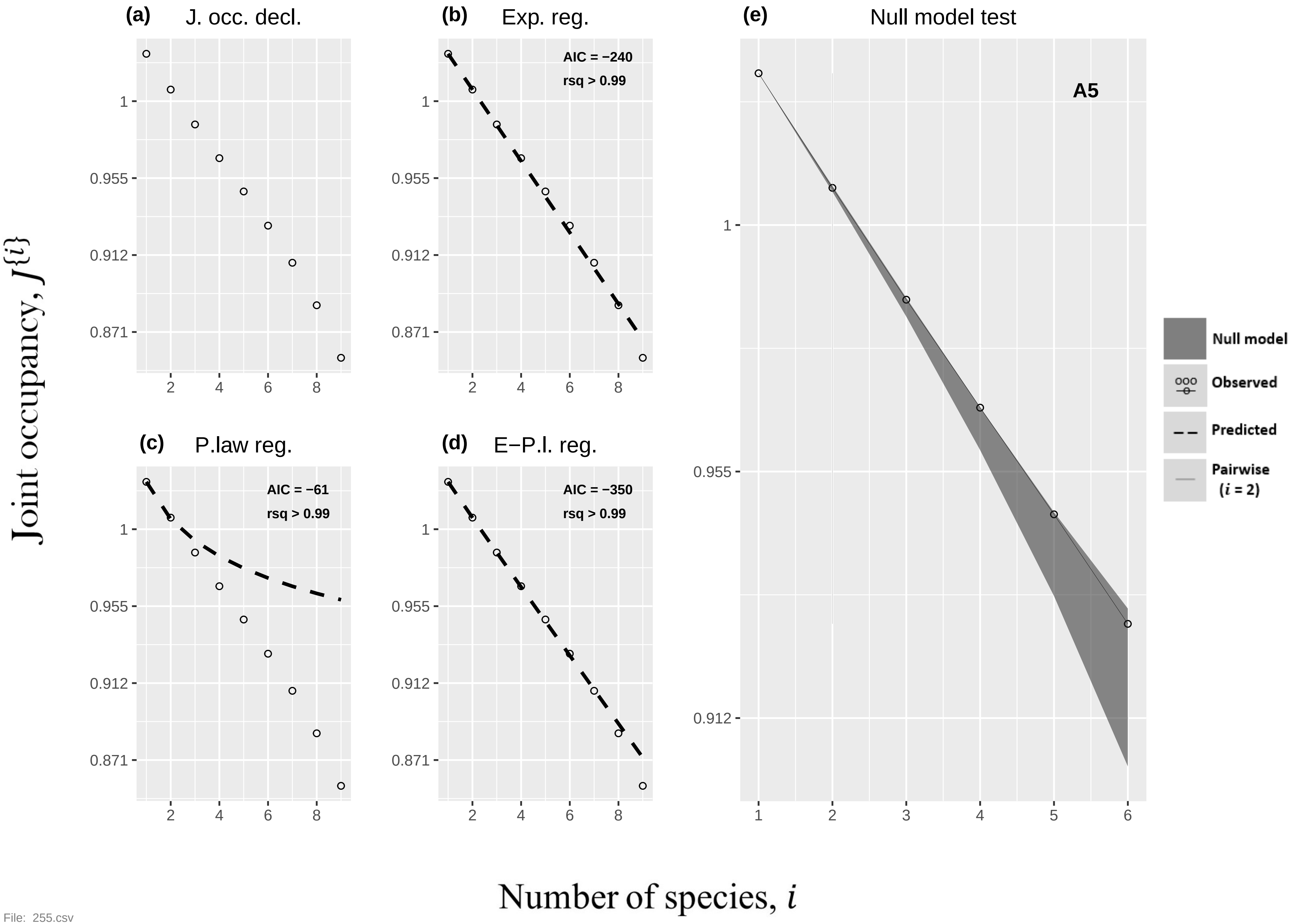

(a) J. occ. decl.
(b) Exp. reg.
AIC = −240
rsq > 0.99
(c) P.law reg.
AIC = −61
rsq > 0.99
(d) E−P.l. reg.
AIC = −350
rsq > 0.99
(e) Null model test
A5
Null model
Observed
Predicted
Pairwise
(i = 2)
1
0.955
0.912
0.871
2
4
6
8
1
2
3
4
5
6
Joint occupancy, J{i}
Number of species, i


File: 255.csv

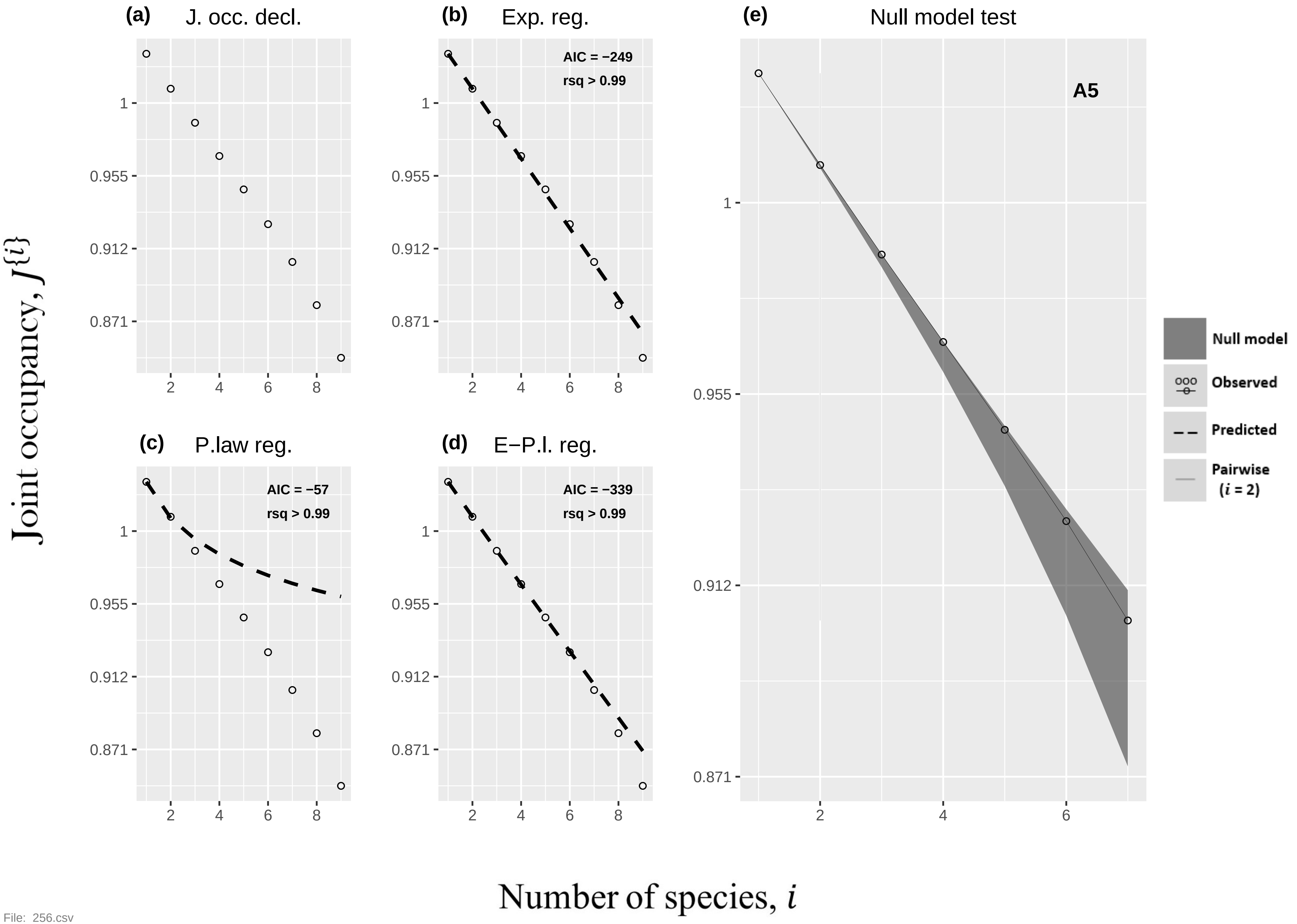
(a) J. occ. decl.
(b) Exp. reg.
AIC = −249
rsq > 0.99
(c) P.law reg.
AIC = −57
rsq > 0.99
(d) E−P.l. reg.
AIC = −339
rsq > 0.99
(e) Null model test
A5
Null model
Observed
Predicted
Pairwise (i = 2)
1
0.955
0.912
0.871
2
4
6
8
Joint occupancy, J{i}
Number of species, i

File: 256.csv

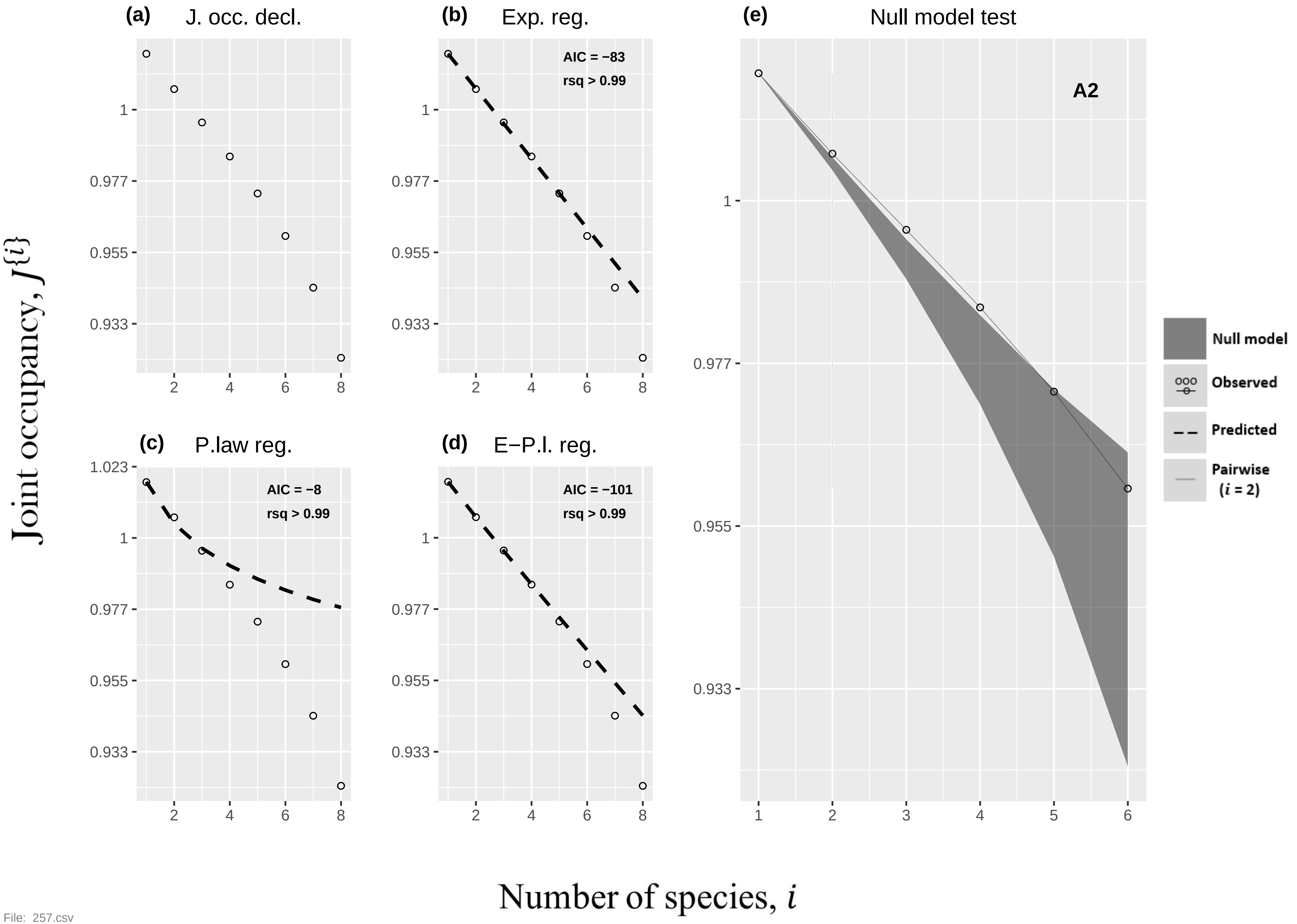

(a) J. occ. decl.
(b) Exp. reg.
AIC = −83
rsq > 0.99
(c) P.law reg.
AIC = −8
rsq > 0.99
(d) E−P.l. reg.
AIC = −101
rsq > 0.99
(e) Null model test
A2
Null model
Observed
Predicted
Pairwise
(i = 2)
1.023
1
0.977
0.955
0.933
1
2
3
4
5
6
7
8
Joint occupancy, J{i}
Number of species, i


File: 257.csv

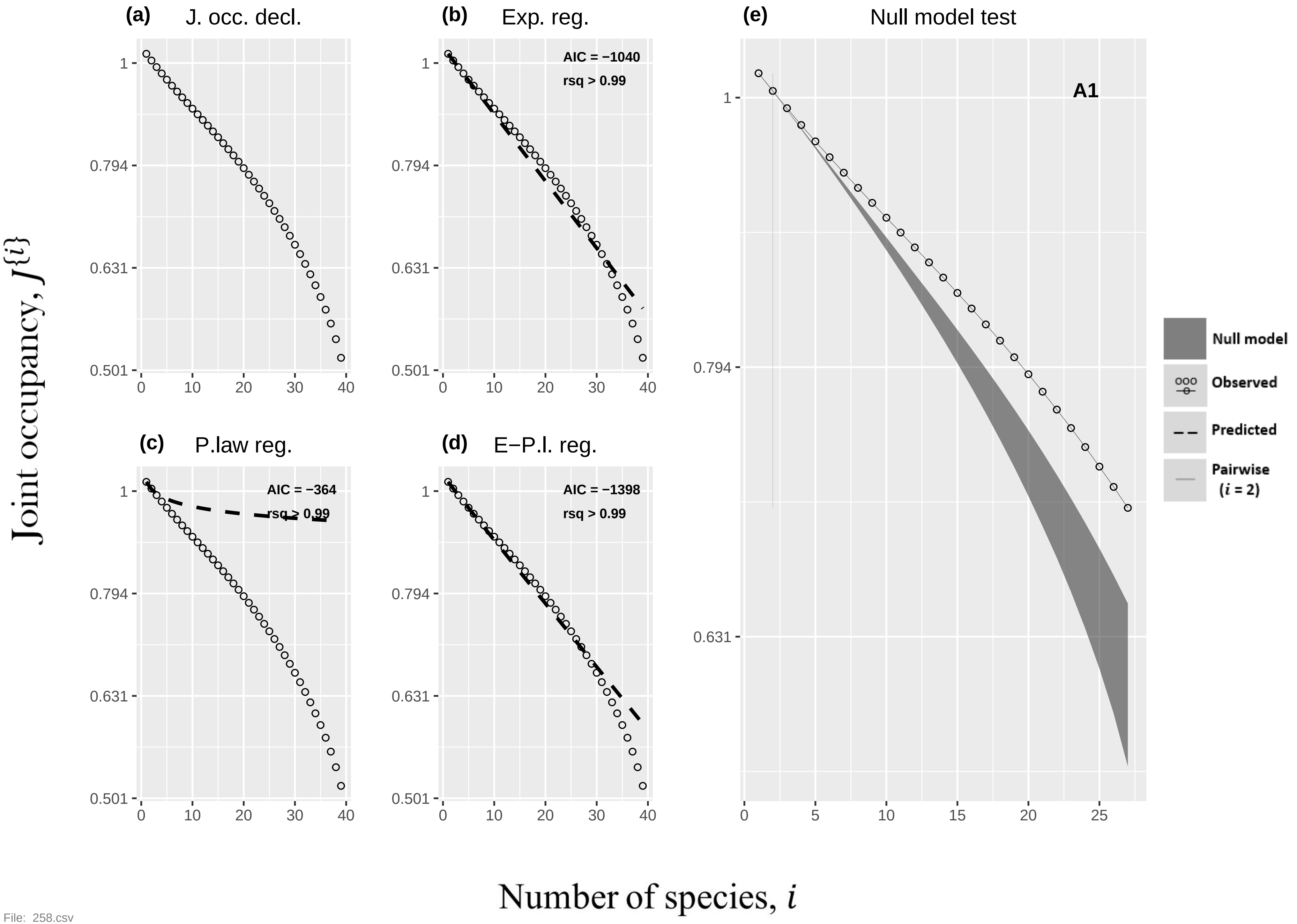

(a) J. occ. decl.
(b) Exp. reg.
AIC = −1040
rsq > 0.99
(c) P.law reg.
AIC = −364
rsq > 0.99
(d) E−P.l. reg.
AIC = −1398
rsq > 0.99
(e) Null model test
A1
Null model
Observed
Predicted
Pairwise (i = 2)
Joint occupancy, J{i}
Number of species, i


File: 258.csv

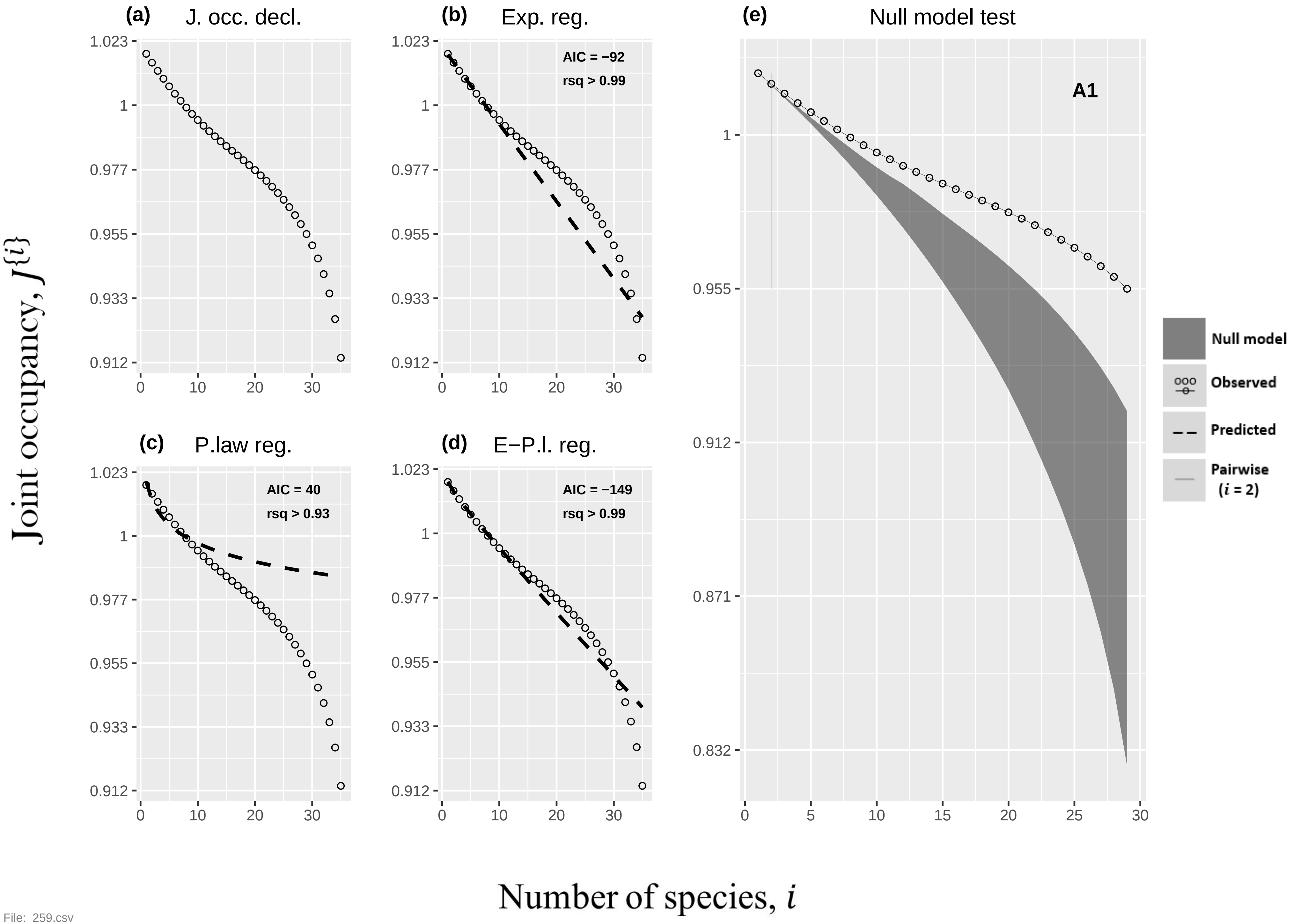

(a) J. occ. decl.
(b) Exp. reg.
AIC = −92
rsq > 0.99
(c) P.law reg.
AIC = 40
rsq > 0.93
(d) E−P.l. reg.
AIC = −149
rsq > 0.99
(e) Null model test
A1
Null model
Observed
Predicted
Pairwise (i = 2)
Joint occupancy, J{i}
Number of species, i


File: 259.csv

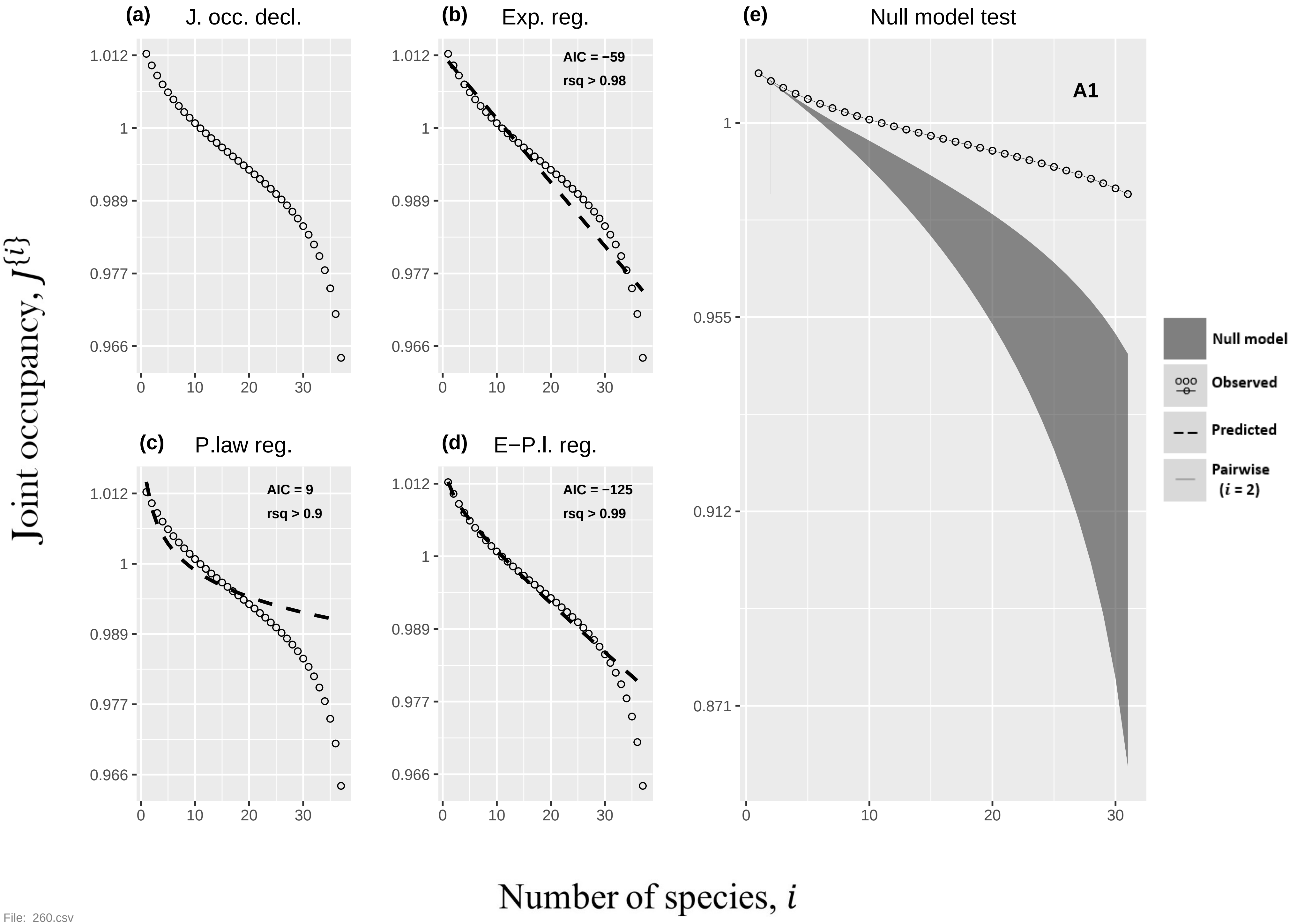

(a) J. occ. decl.
(b) Exp. reg.
AIC = −59
rsq > 0.98
(c) P.law reg.
AIC = 9
rsq > 0.9
(d) E−P.l. reg.
AIC = −125
rsq > 0.99
(e) Null model test
A1
Null model
Observed
Predicted
Pairwise (i = 2)
1.012
1
0.989
0.977
0.966
0.955
0.912
0.871
0
10
20
30
Joint occupancy, J{i}
Number of species, i


File: 260.csv

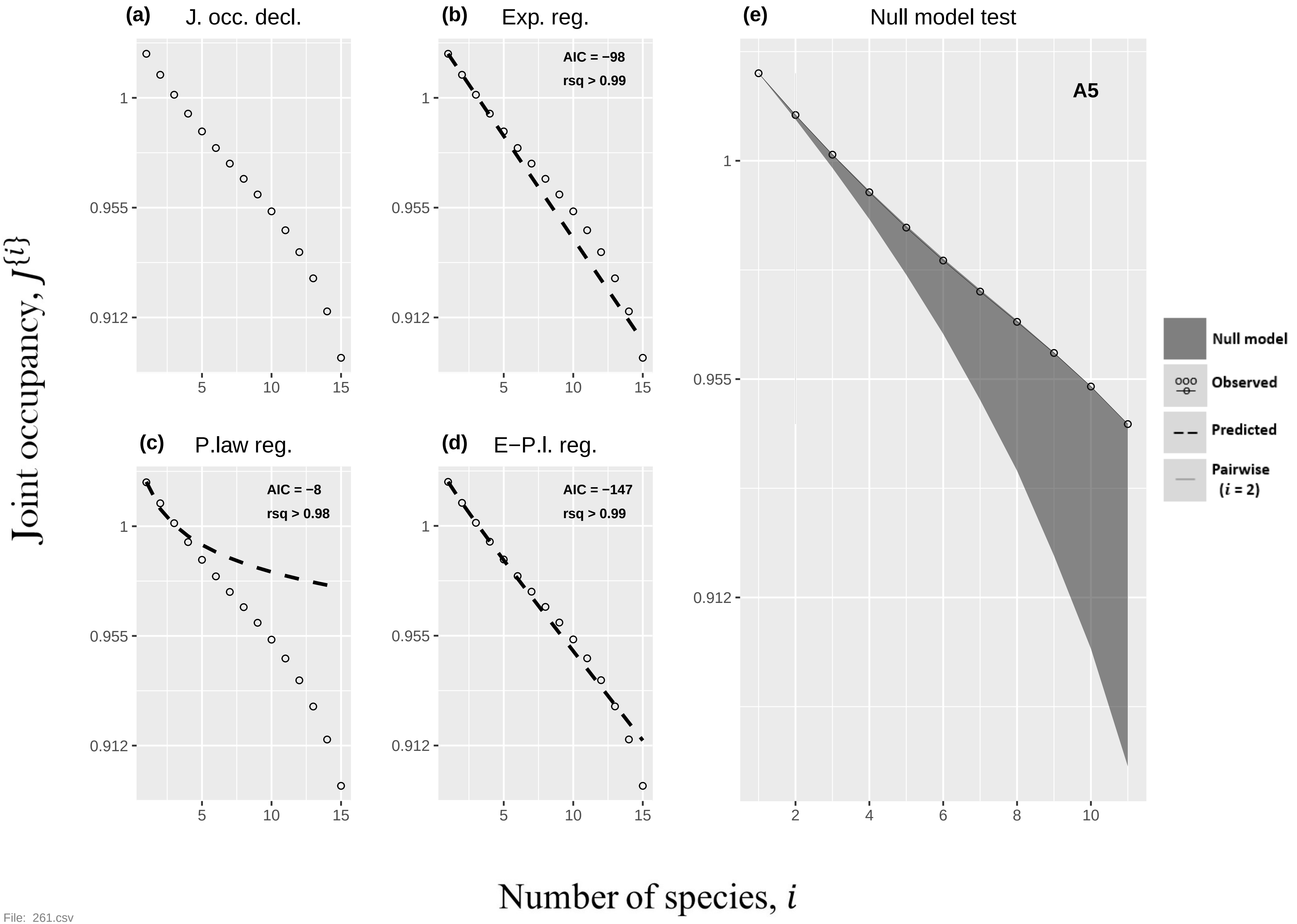
(a) J. occ. decl.
(b) Exp. reg.
AIC = −98
rsq > 0.99
(c) P.law reg.
AIC = −8
rsq > 0.98
(d) E−P.l. reg.
AIC = −147
rsq > 0.99
(e) Null model test
A5
Null model
Observed
Predicted
Pairwise
(i = 2)
Joint occupancy, J{i}
Number of species, i
1
0.955
0.912
5
10
15
2
4
6
8
10

File: 261.csv

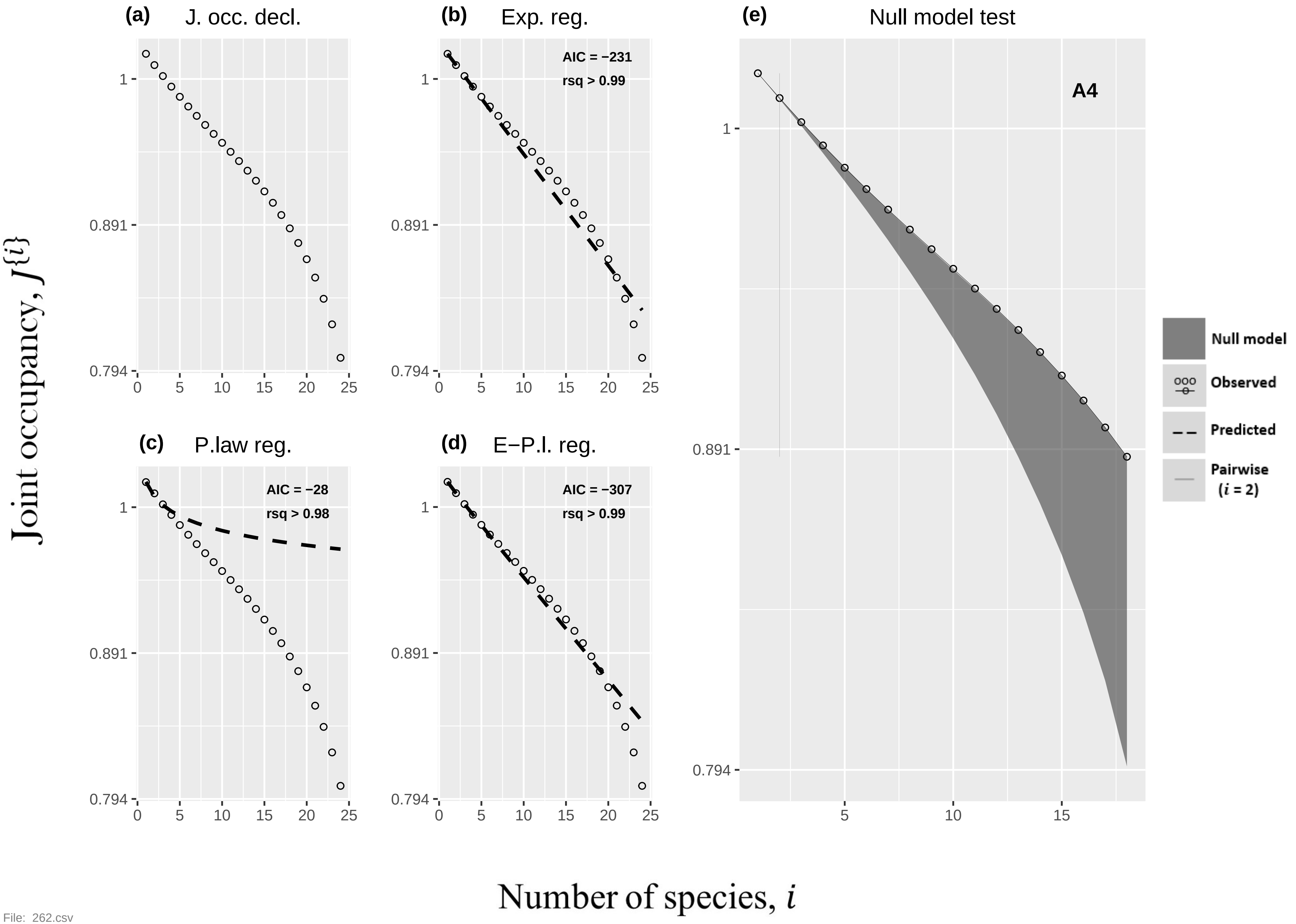

(a) J. occ. decl.
(b) Exp. reg.
AIC = −231
rsq > 0.99
(e) Null model test
A4
(c) P.law reg.
AIC = −28
rsq > 0.98
(d) E−P.l. reg.
AIC = −307
rsq > 0.99
Null model
Observed
Predicted
Pairwise (i = 2)
Joint occupancy, J{i}
Number of species, i


File: 262.csv

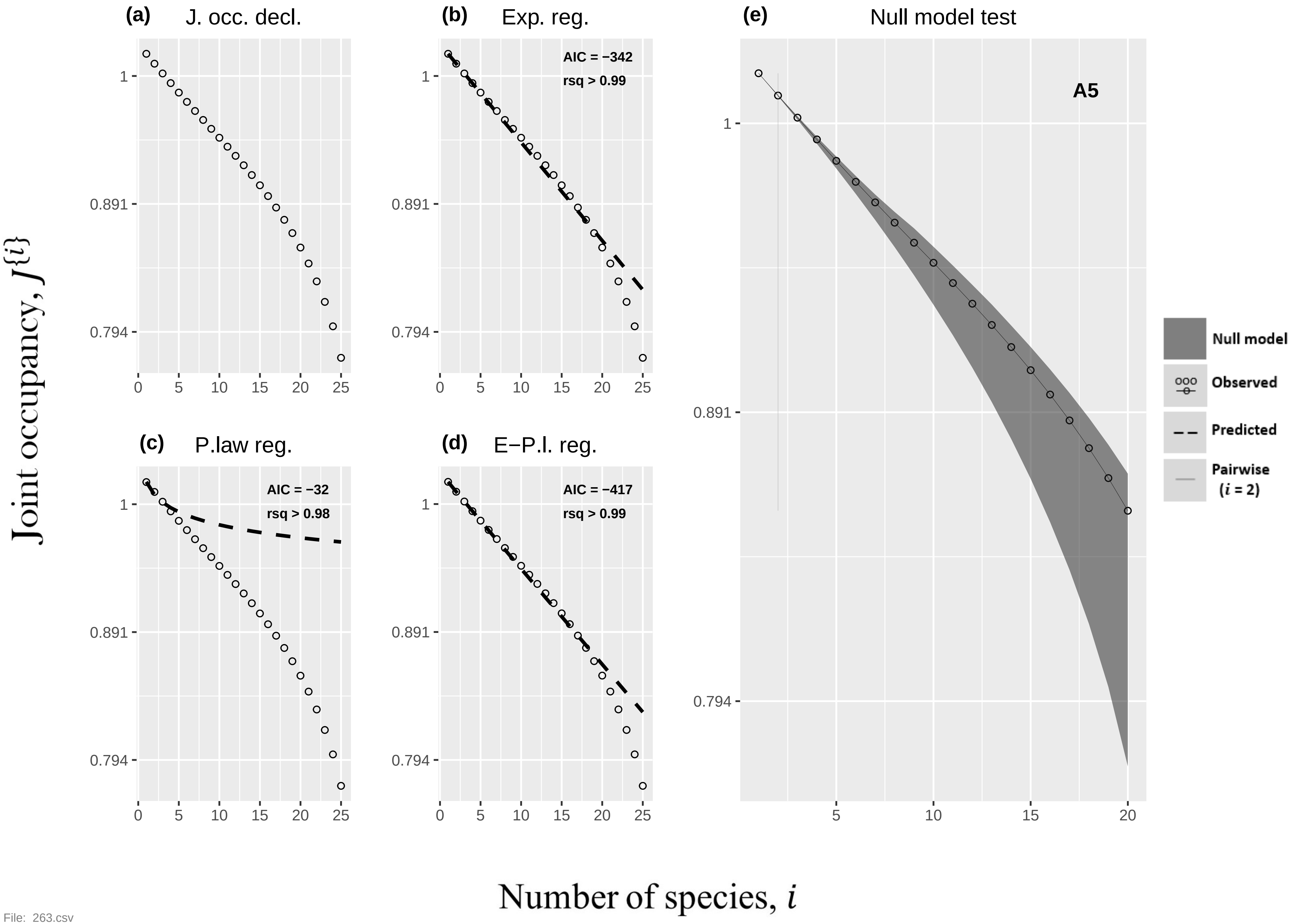

(a)
J. occ. decl.
(b)
Exp. reg.
AIC = −342
rsq > 0.99
(c)
P.law reg.
AIC = −32
rsq > 0.98
(d)
E−P.l. reg.
AIC = −417
rsq > 0.99
(e)
Null model test
A5
Null model
Observed
Predicted
Pairwise (i = 2)
1
0.891
0.794
0
5
10
15
20
25
Joint occupancy, J{i}
Number of species, i


File: 263.csv

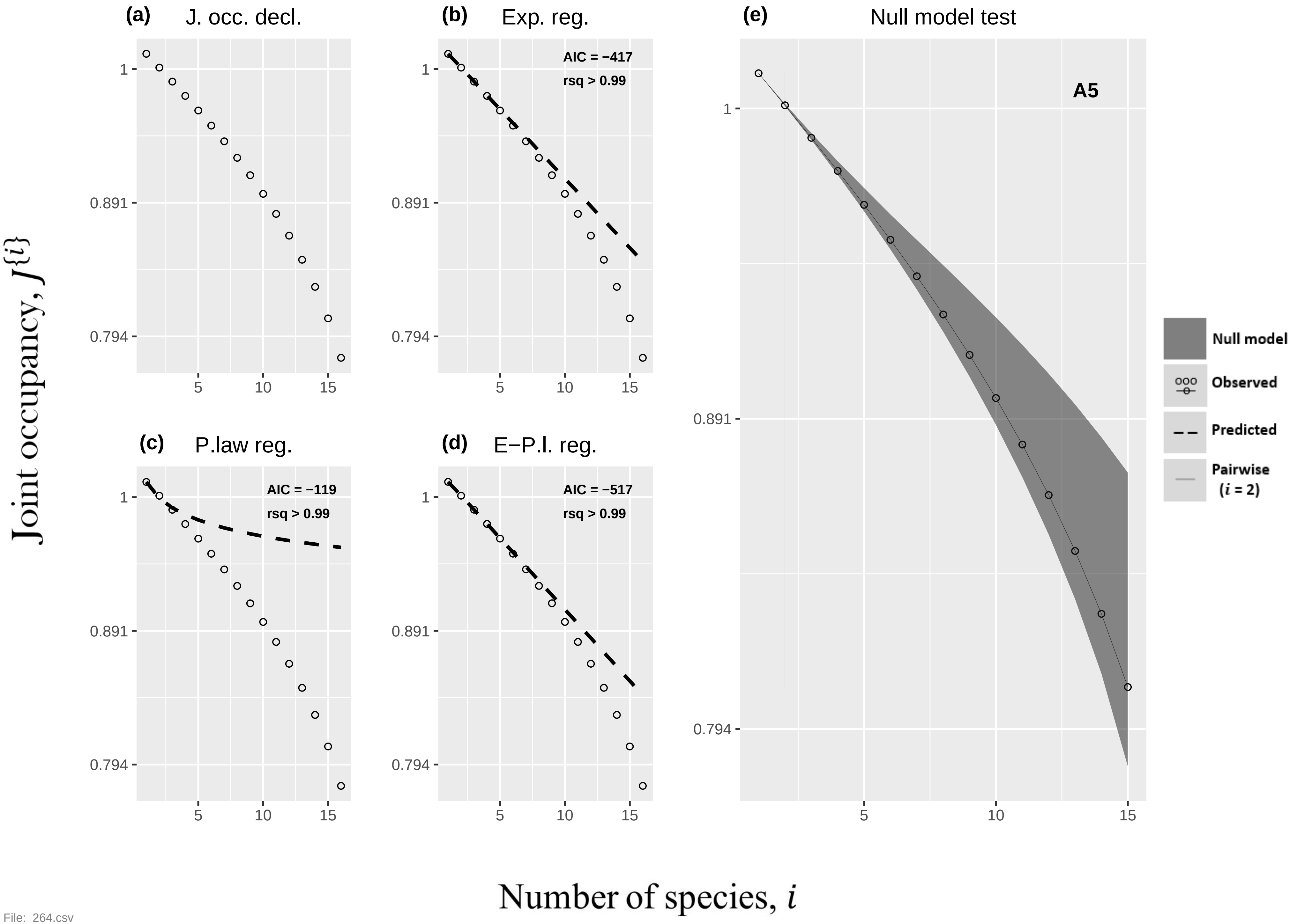

(a)
J. occ. decl.
(b)
Exp. reg.
AIC = −417
rsq > 0.99
(c)
P.law reg.
AIC = −119
rsq > 0.99
(d)
E−P.l. reg.
AIC = −517
rsq > 0.99
(e)
Null model test
A5
Null model
Observed
Predicted
Pairwise
(i = 2)
1
0.891
0.794
5
10
15
Joint occupancy, J{i}
Number of species, i

File: 264.csv

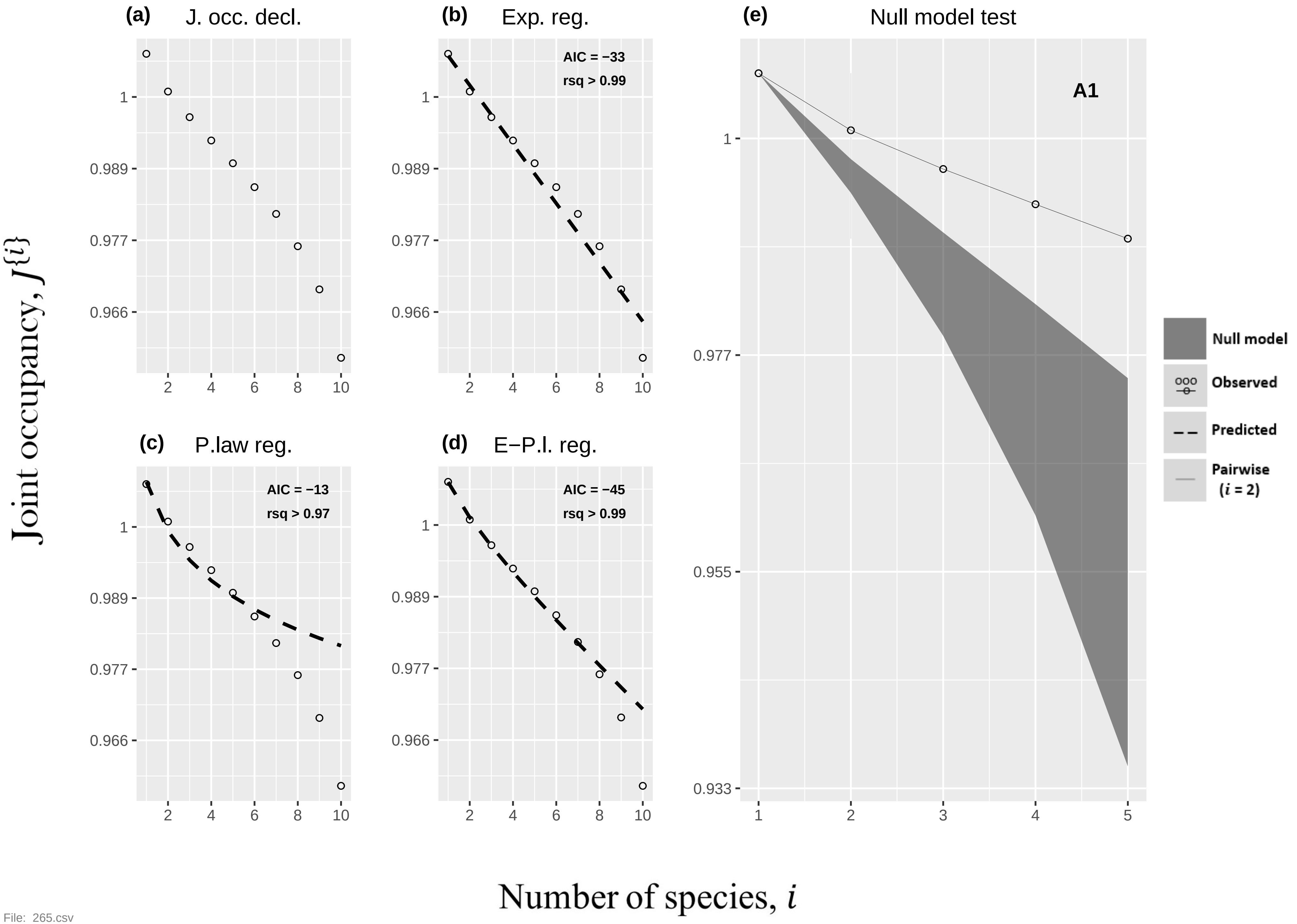

(a) J. occ. decl.
(b) Exp. reg.
AIC = −33
rsq > 0.99
(c) P.law reg.
AIC = −13
rsq > 0.97
(d) E−P.l. reg.
AIC = −45
rsq > 0.99
(e) Null model test
A1
Null model
Observed
Predicted
Pairwise (i = 2)
Joint occupancy, J{i}
Number of species, i


File: 265.csv

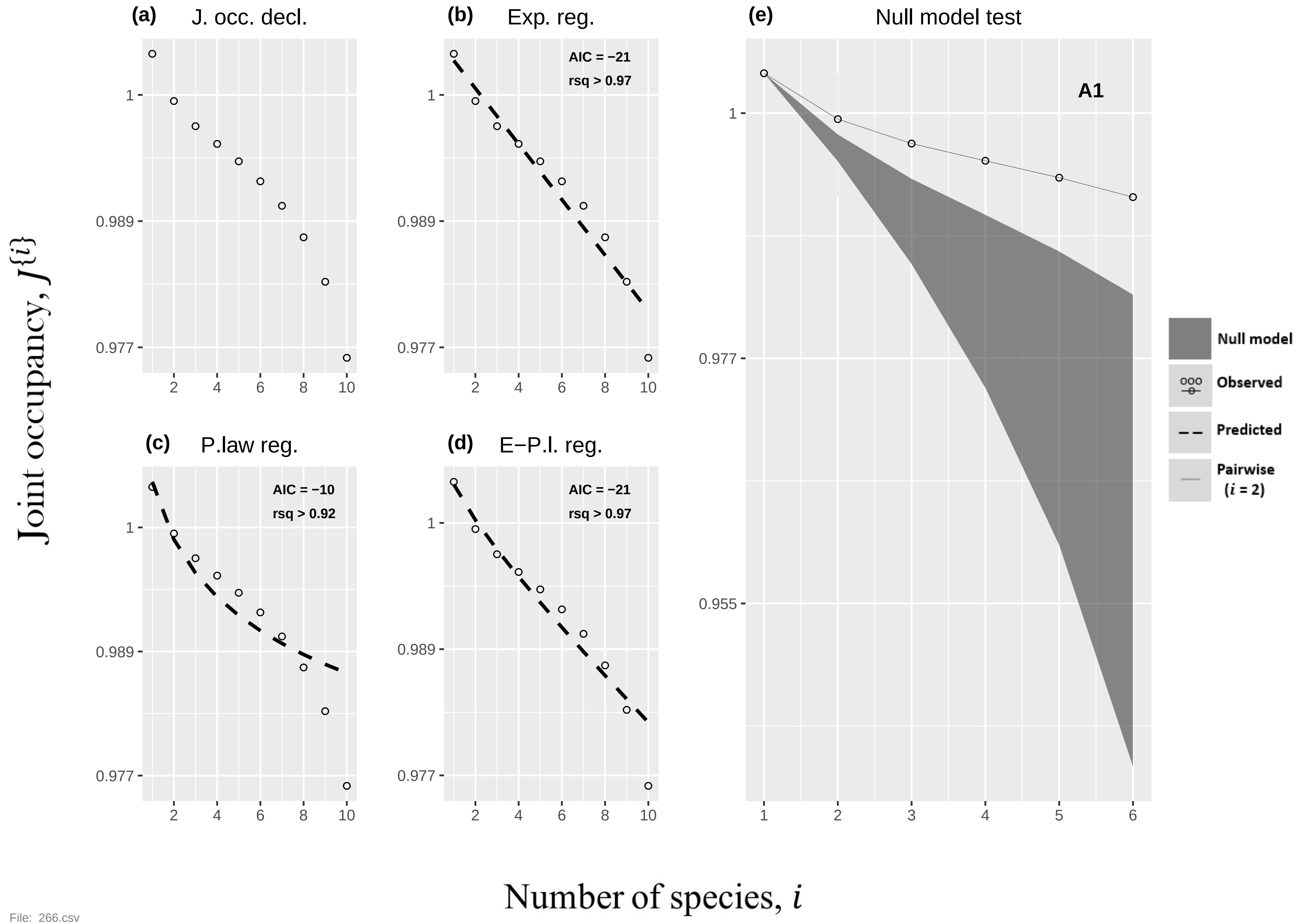
(a)
J. occ. decl.
(b)
Exp. reg.
AIC = −21
rsq > 0.97
(c)
P.law reg.
AIC = −10
rsq > 0.92
(d)
E−P.l. reg.
AIC = −21
rsq > 0.97
(e)
Null model test
A1
Null model
Observed
Predicted
Pairwise
(i = 2)
Joint occupancy, J{i}
Number of species, i
1
0.989
0.977
0.955
2
4
6
8
10
3
5
File: 266.csv

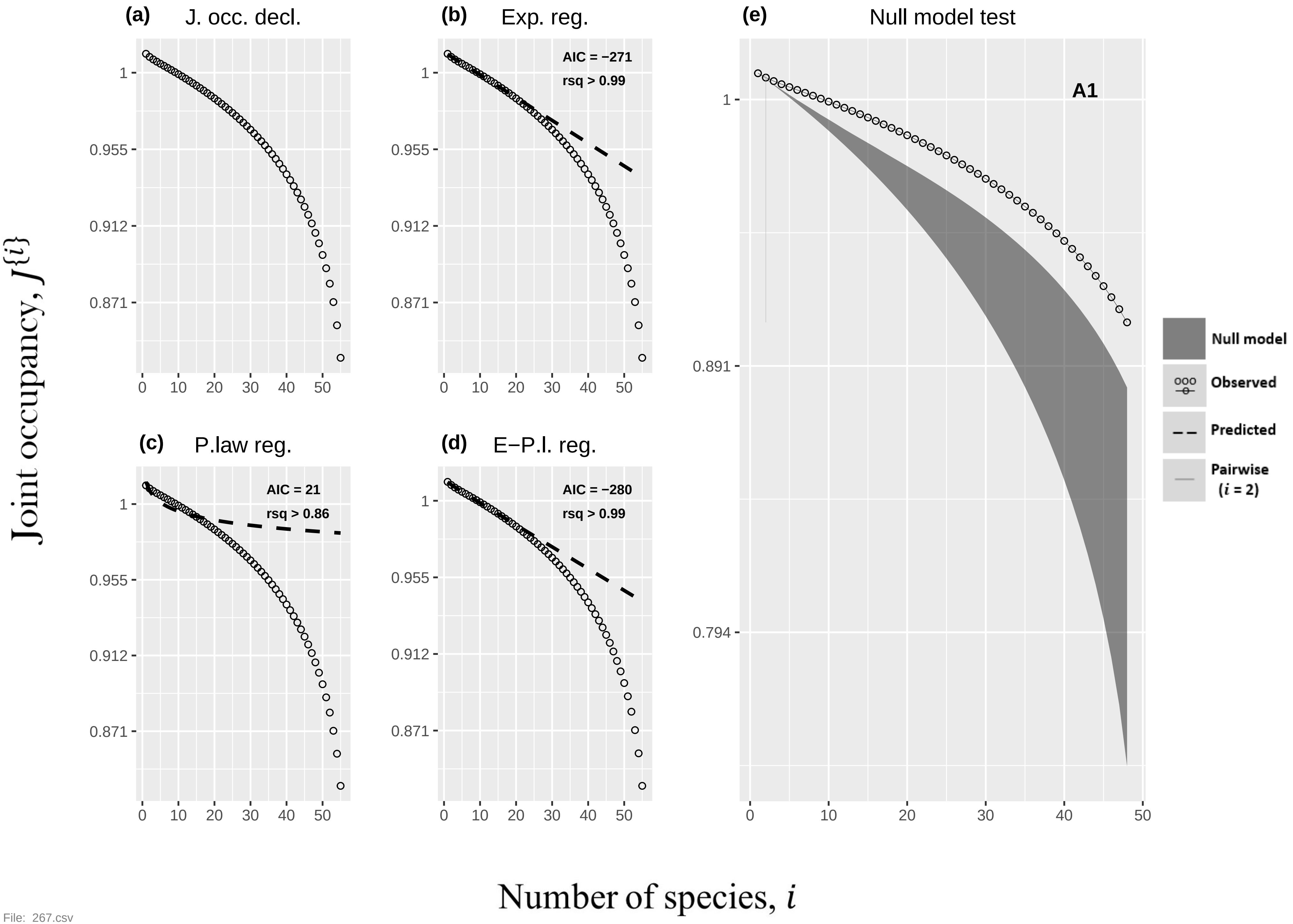
(a) J. occ. decl.
(b) Exp. reg.
AIC = −271
rsq > 0.99
(c) P.law reg.
AIC = 21
rsq > 0.86
(d) E−P.l. reg.
AIC = −280
rsq > 0.99
(e) Null model test
A1
Null model
Observed
Predicted
Pairwise
(i = 2)
Joint occupancy, J{i}
Number of species, i

File: 267.csv

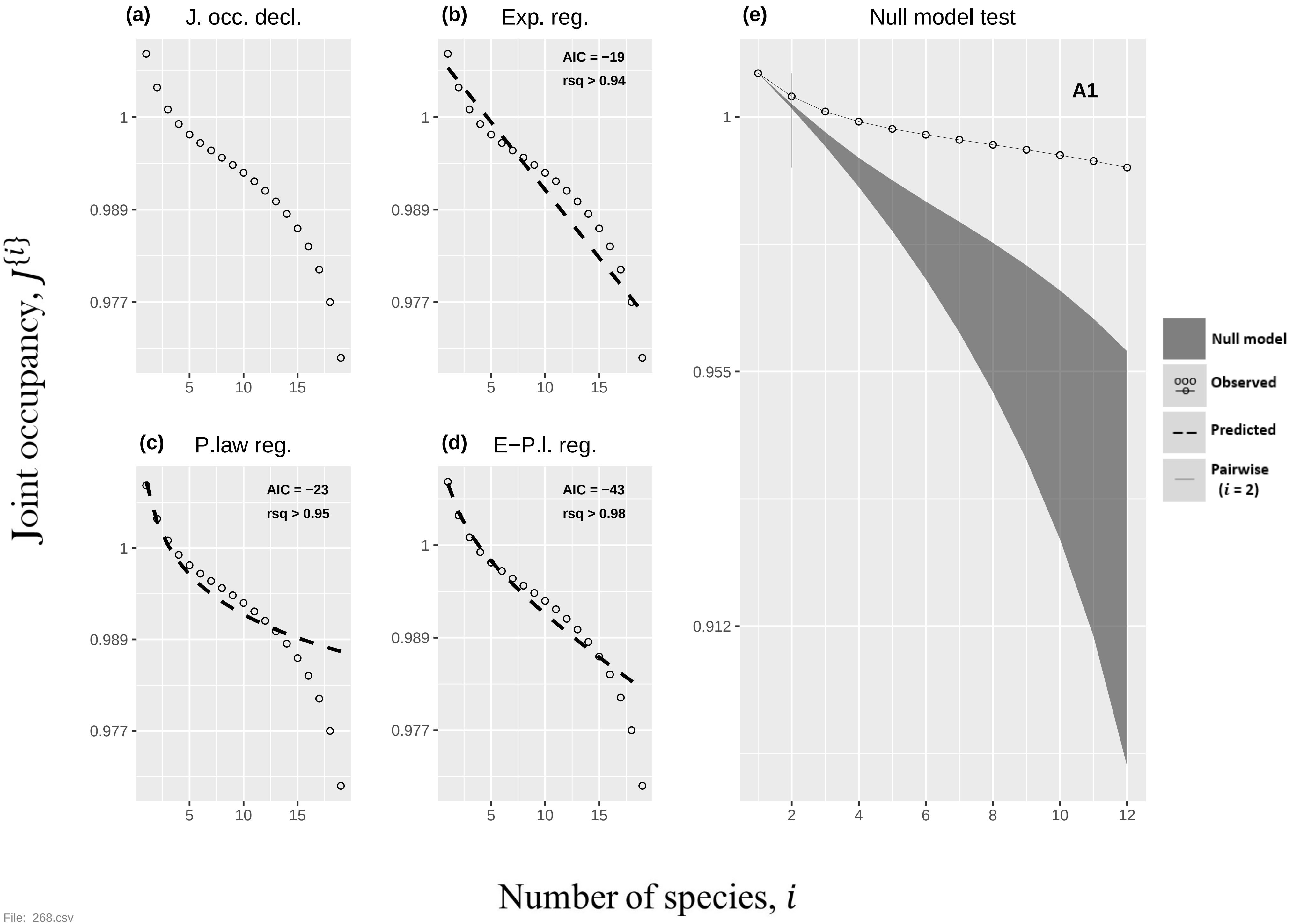
(a) J. occ. decl.
(b) Exp. reg.
AIC = −19
rsq > 0.94
(c) P.law reg.
AIC = −23
rsq > 0.95
(d) E−P.l. reg.
AIC = −43
rsq > 0.98
(e) Null model test
A1
Null model
Observed
Predicted
Pairwise (i = 2)
1
0.989
0.977
0.955
0.912
5
10
15
2
4
6
8
10
12
Joint occupancy, J{i}
Number of species, i

File: 268.csv

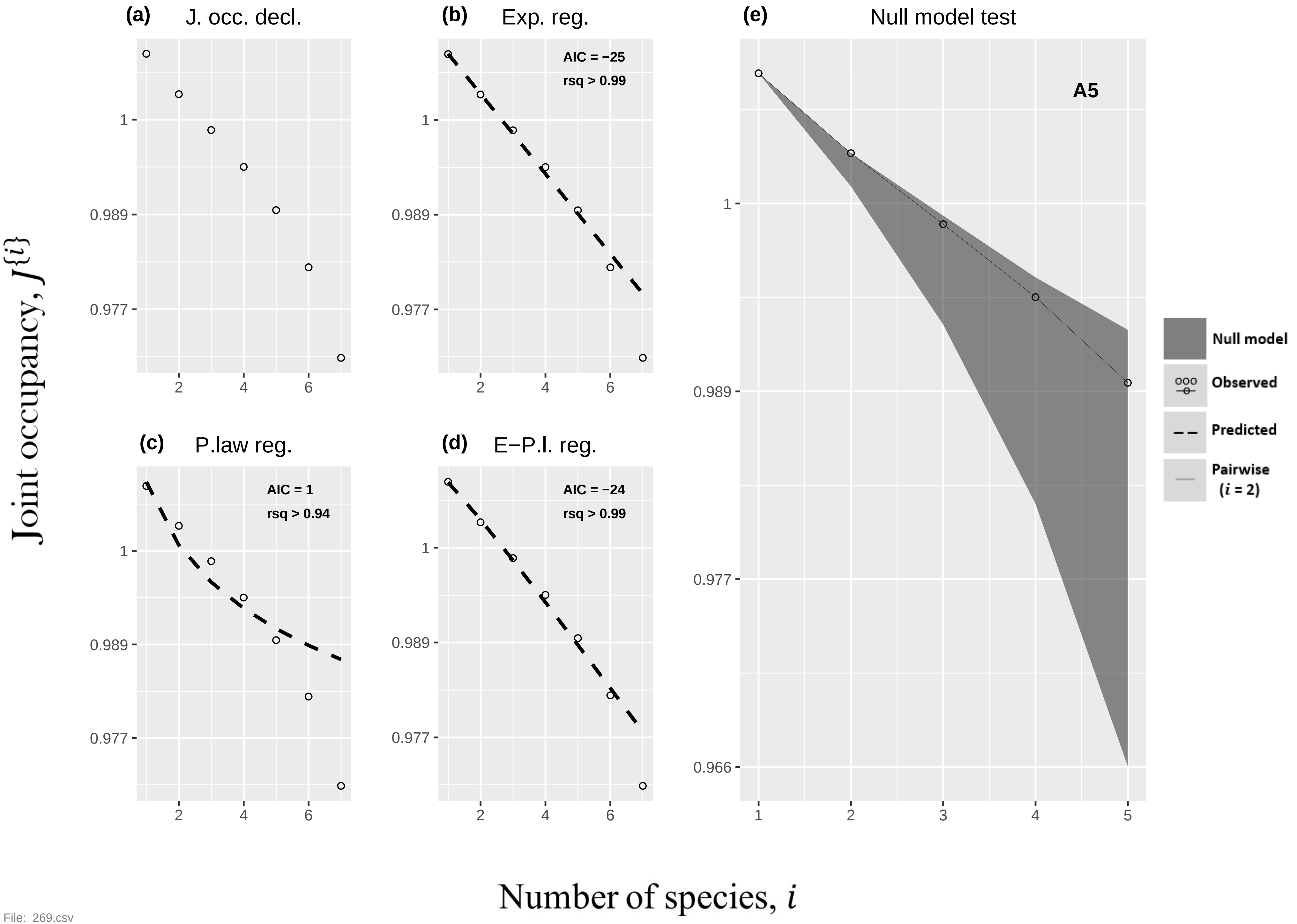

(a)
J. occ. decl.
(b)
Exp. reg.
AIC = −25
rsq > 0.99
(c)
P.law reg.
AIC = 1
rsq > 0.94
(d)
E−P.l. reg.
AIC = −24
rsq > 0.99
(e)
Null model test
A5
Null model
Observed
Predicted
Pairwise
(i = 2)
Joint occupancy, J{i}
Number of species, i
1
0.989
0.977
0.966


File: 269.csv

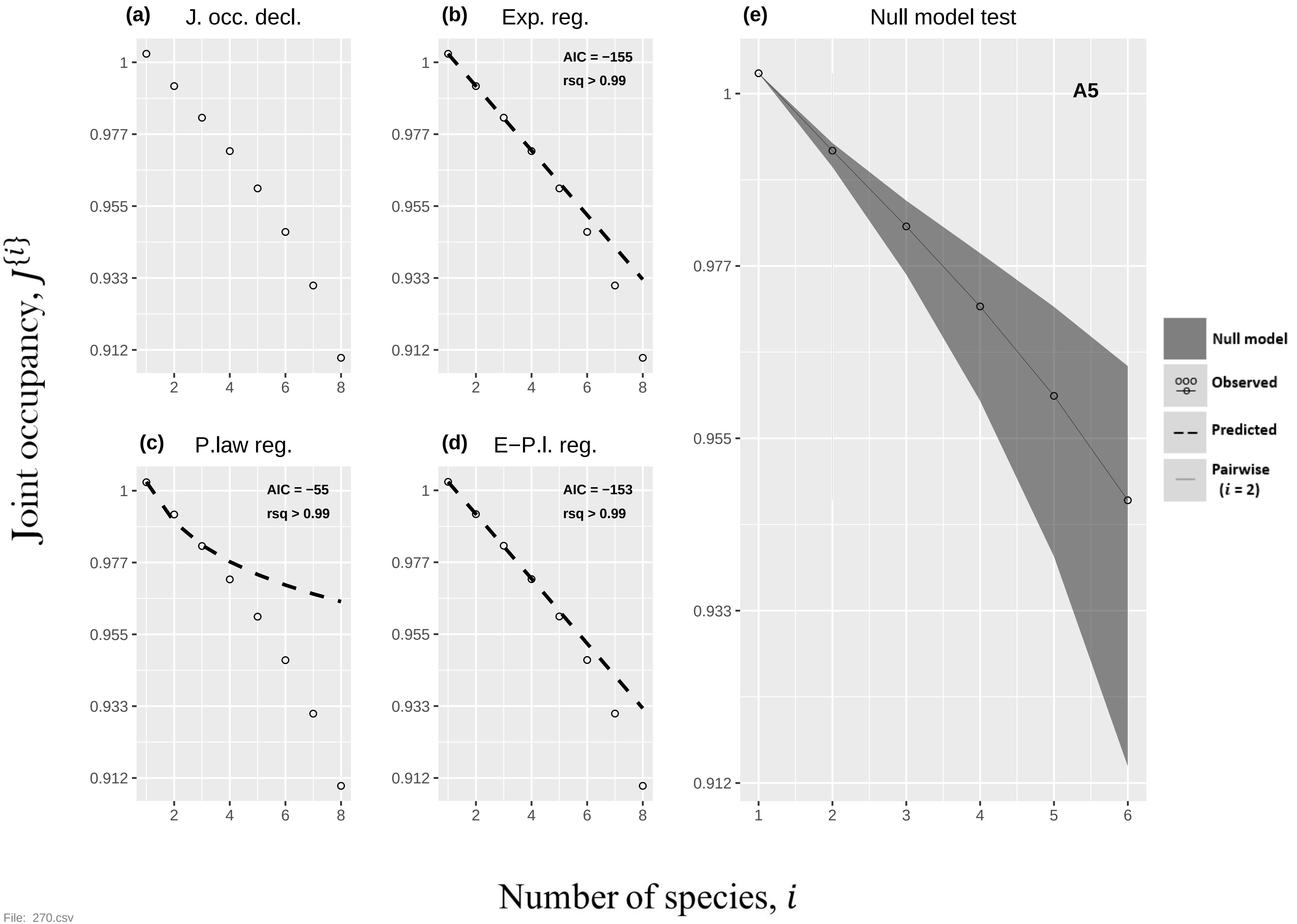

(a) J. occ. decl.
(b) Exp. reg.
AIC = −155
rsq > 0.99
(c) P.law reg.
AIC = −55
rsq > 0.99
(d) E−P.l. reg.
AIC = −153
rsq > 0.99
(e) Null model test
A5
Null model
Observed
Predicted
Pairwise
(i = 2)
1
0.977
0.955
0.933
0.912
2
4
6
8
1
2
3
4
5
6
Joint occupancy, J{i}
Number of species, i


File: 270.csv

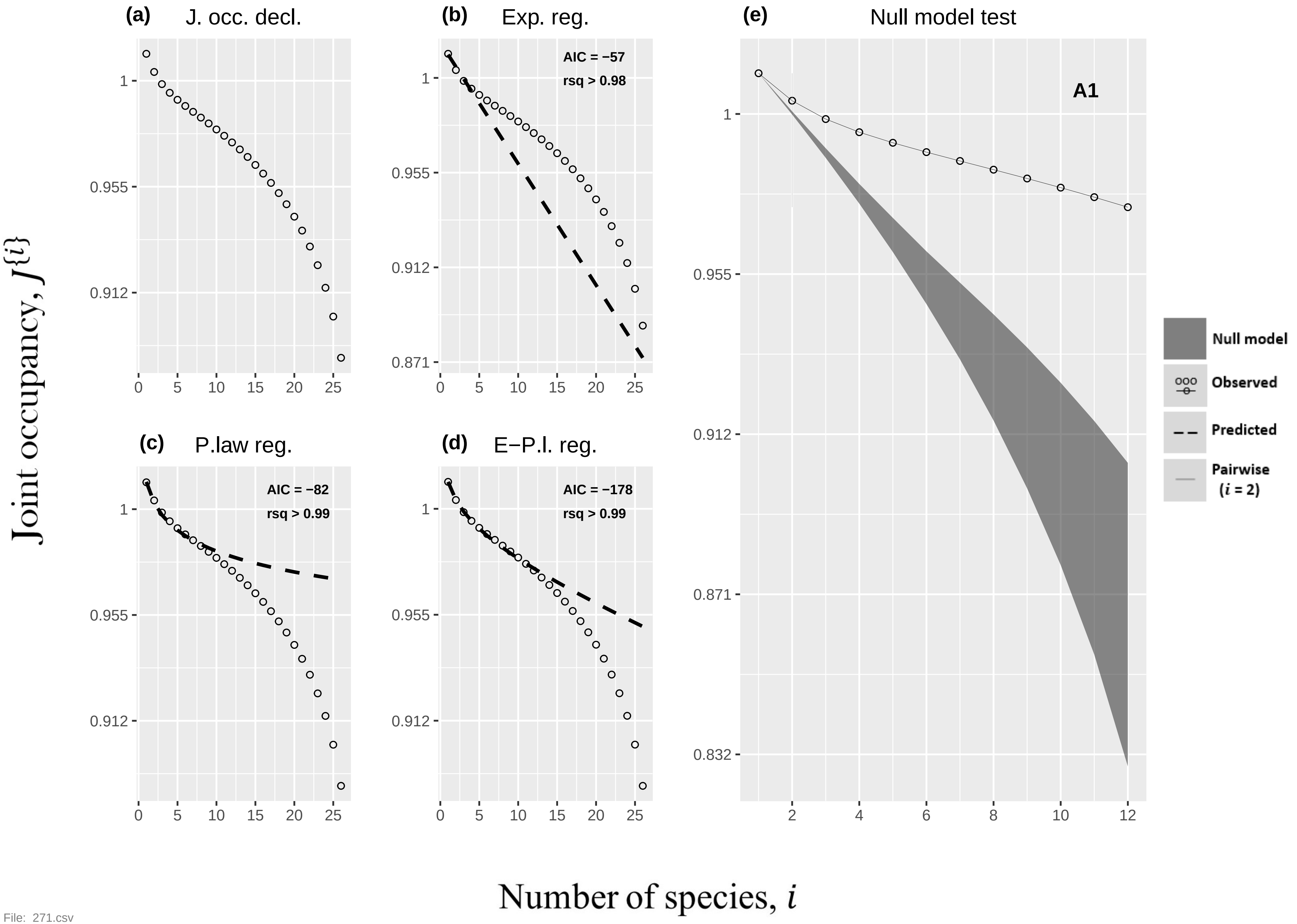

(a) J. occ. decl.
(b) Exp. reg.
AIC = −57
rsq > 0.98
(c) P.law reg.
AIC = −82
rsq > 0.99
(d) E−P.l. reg.
AIC = −178
rsq > 0.99
(e) Null model test
A1
Null model
Observed
Predicted
Pairwise
(i = 2)
Joint occupancy, J{i}
Number of species, i

File: 271.csv

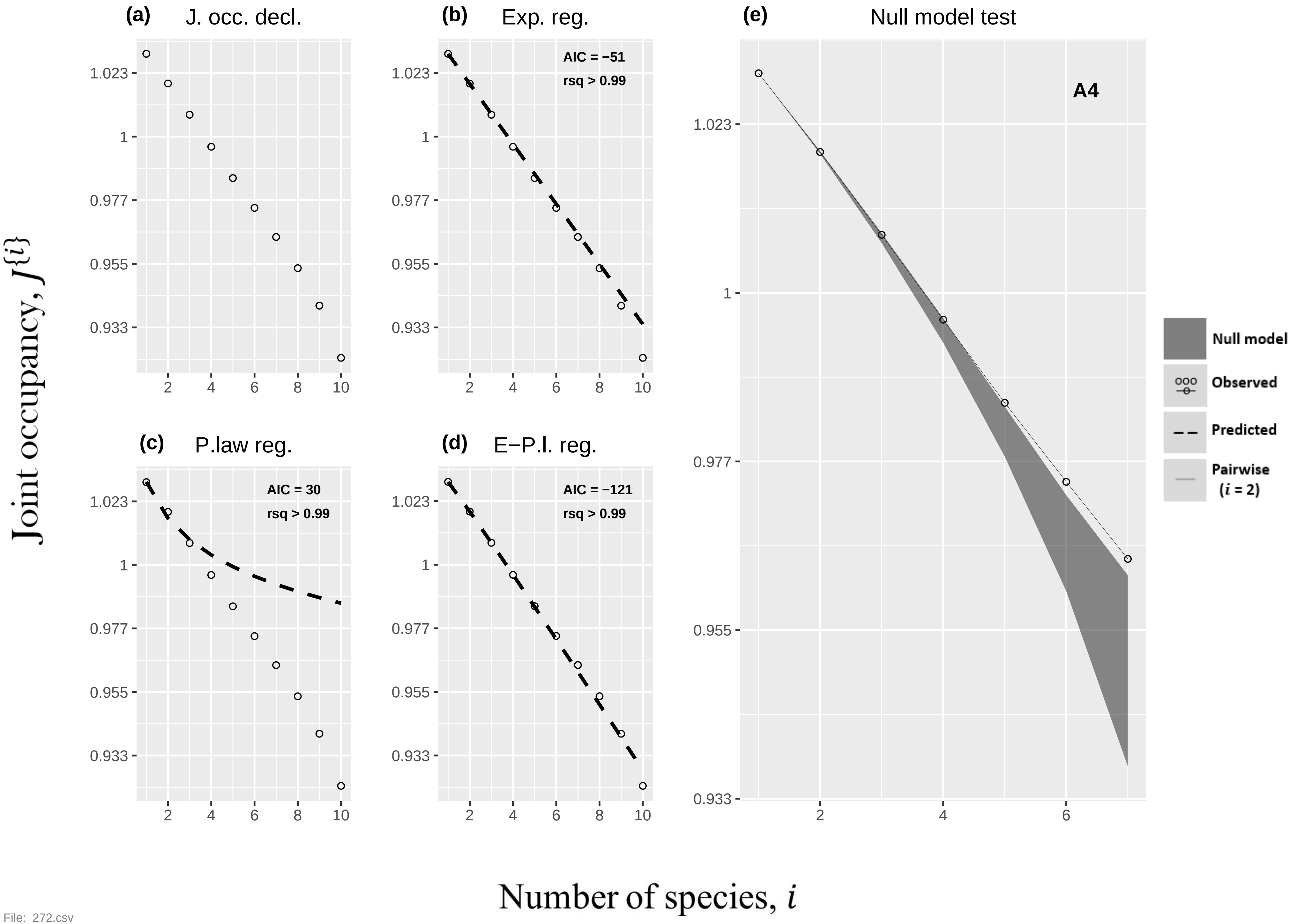

(a) J. occ. decl.
(b) Exp. reg.
AIC = −51
rsq > 0.99
(c) P.law reg.
AIC = 30
rsq > 0.99
(d) E−P.l. reg.
AIC = −121
rsq > 0.99
(e) Null model test
A4
Null model
Observed
Predicted
Pairwise (i = 2)
Joint occupancy, J{i}
Number of species, i
1.023
1
0.977
0.955
0.933
2
4
6
8
10


File: 272.csv

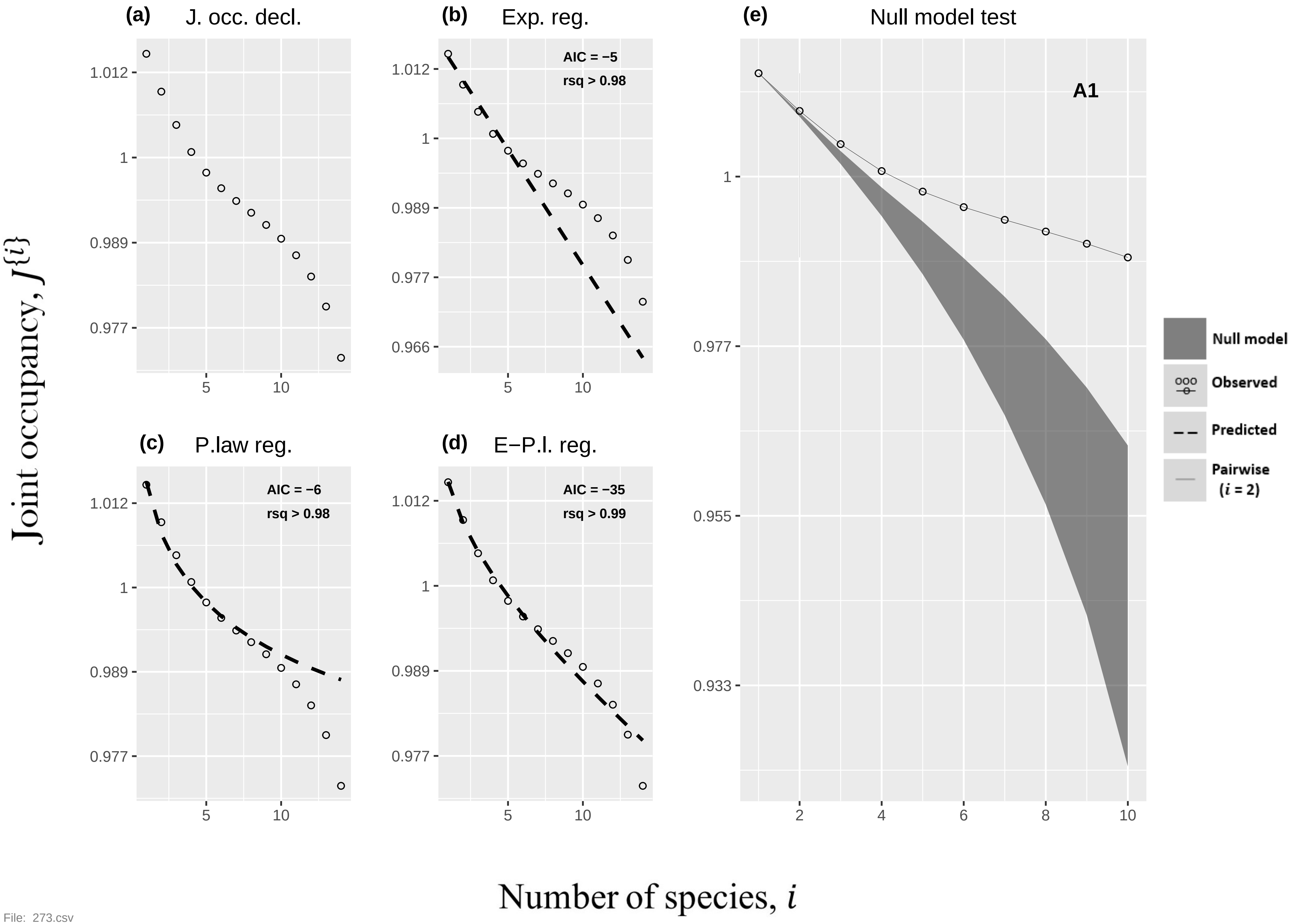
(a) J. occ. decl.
(b) Exp. reg.
AIC = −5
rsq > 0.98
(c) P.law reg.
AIC = −6
rsq > 0.98
(d) E−P.l. reg.
AIC = −35
rsq > 0.99
(e) Null model test
A1
Null model
Observed
Predicted
Pairwise (i = 2)
Joint occupancy, J{i}
Number of species, i

File: 273.csv

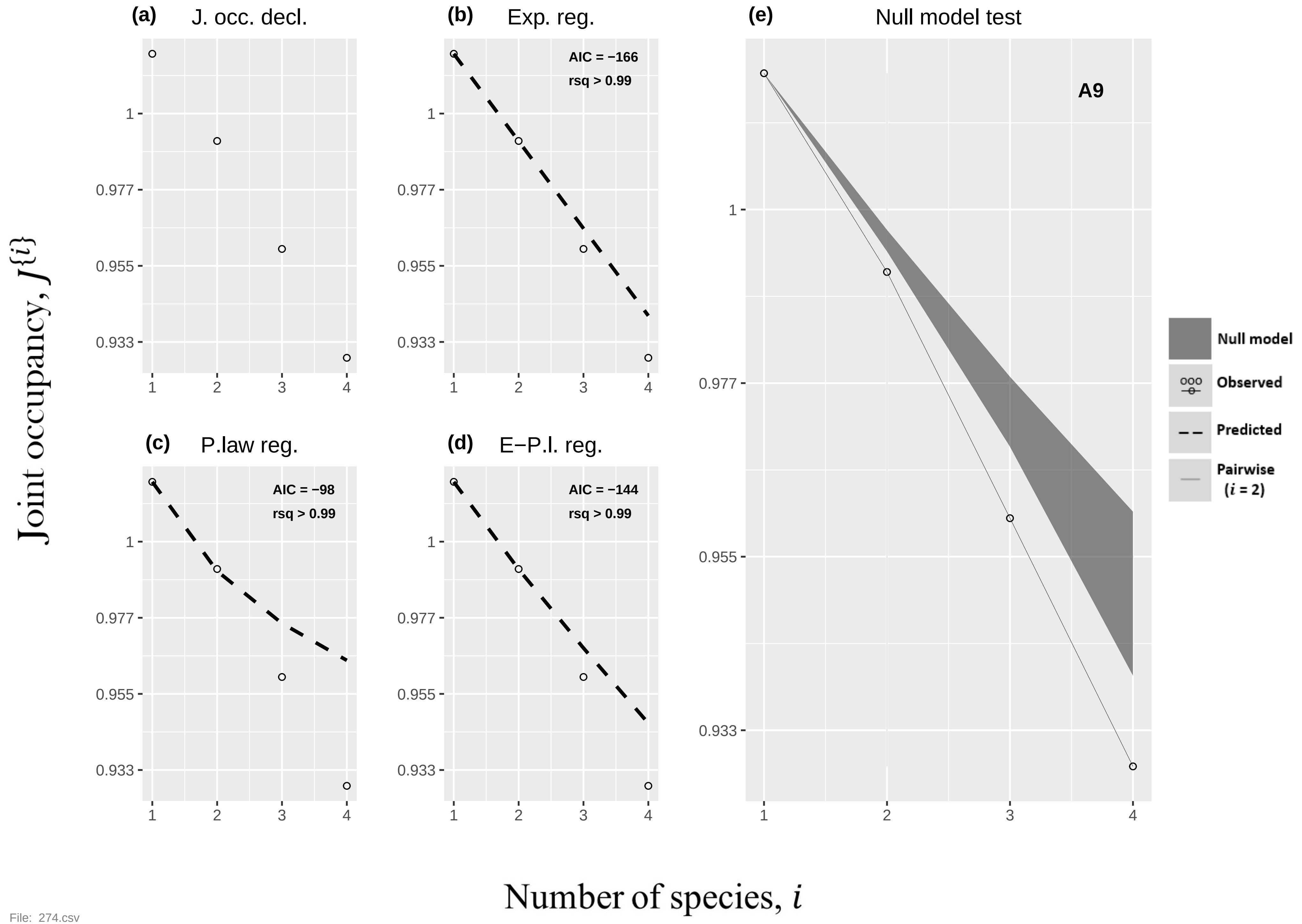

(a) J. occ. decl.
(b) Exp. reg.
AIC = −166
rsq > 0.99
(c) P.law reg.
AIC = −98
rsq > 0.99
(d) E−P.l. reg.
AIC = −144
rsq > 0.99
(e) Null model test
A9
Null model
Observed
Predicted
Pairwise (i = 2)
Joint occupancy, J{i}
Number of species, i


File: 274.csv

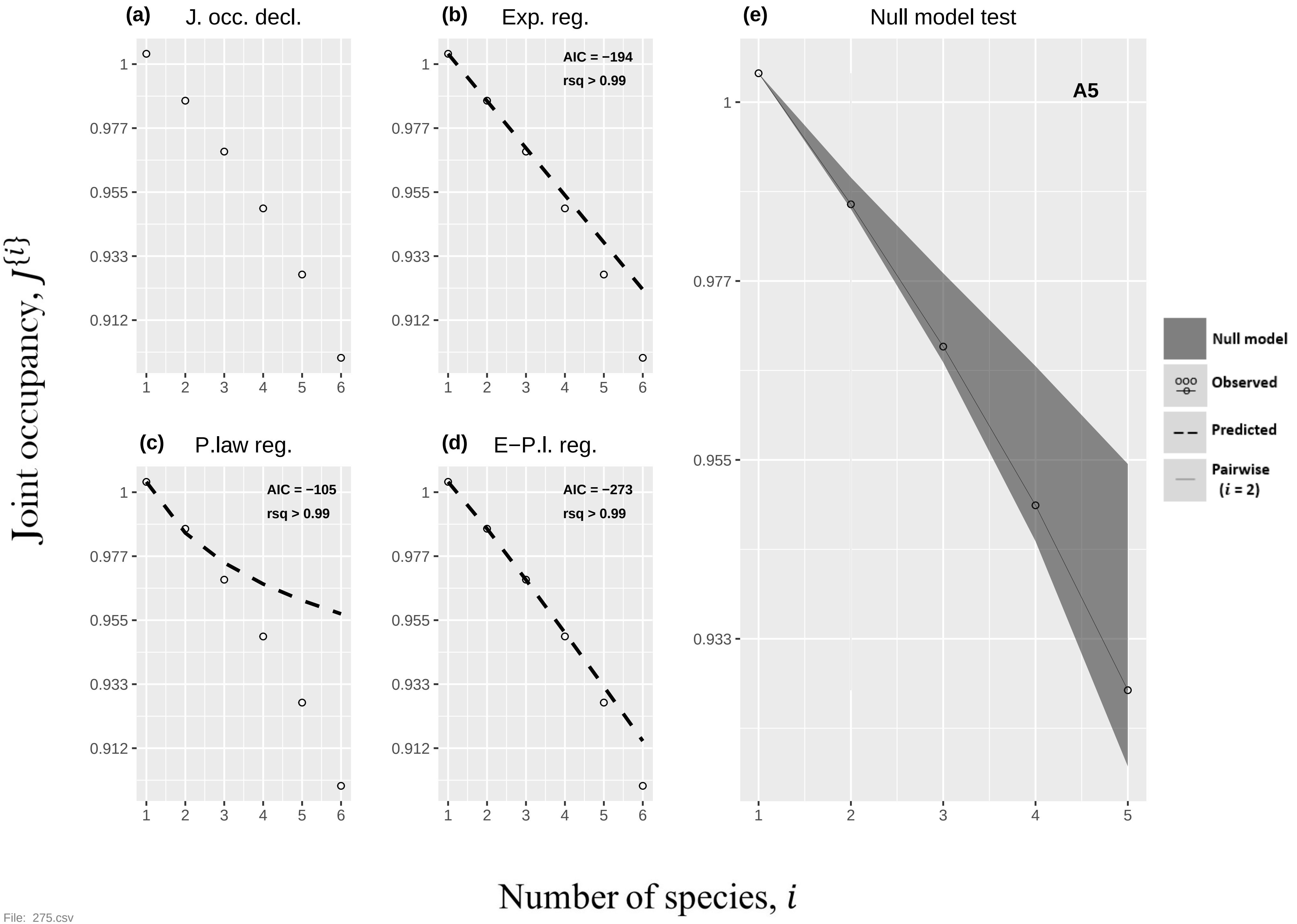
(a)
J. occ. decl.
(b)
Exp. reg.
AIC = −194
rsq > 0.99
(c)
P.law reg.
AIC = −105
rsq > 0.99
(d)
E−P.l. reg.
AIC = −273
rsq > 0.99
(e)
Null model test
A5
Null model
Observed
Predicted
Pairwise
(i = 2)
Joint occupancy, J{i}
Number of species, i
1
0.977
0.955
0.933
0.912
1 2 3 4 5 6

File: 275.csv

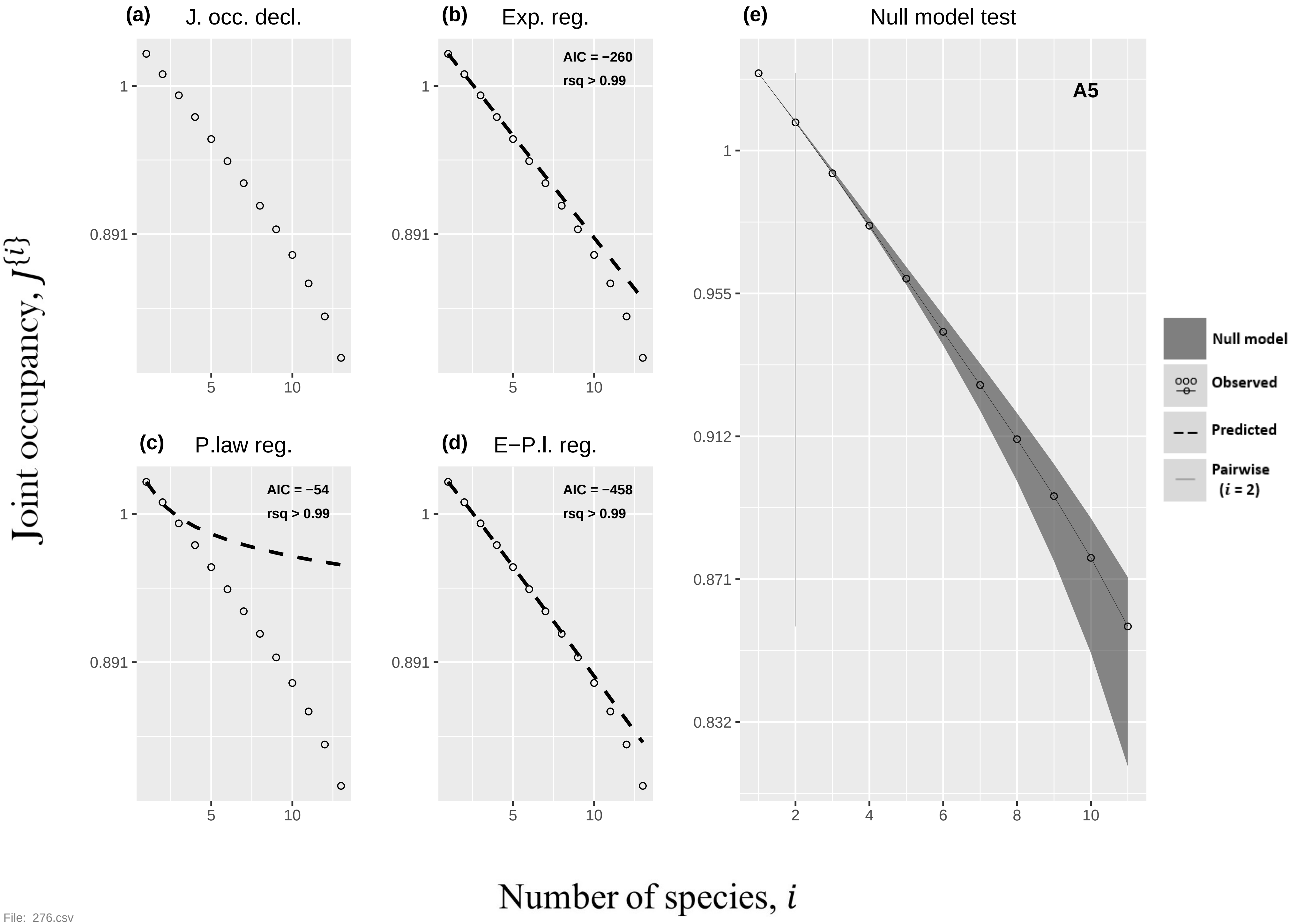

(a) J. occ. decl.
(b) Exp. reg.
AIC = −260
rsq > 0.99
(c) P.law reg.
AIC = −54
rsq > 0.99
(d) E−P.l. reg.
AIC = −458
rsq > 0.99
(e) Null model test
A5
Null model
Observed
Predicted
Pairwise
(i = 2)
Joint occupancy, J{i}
Number of species, i


File: 276.csv

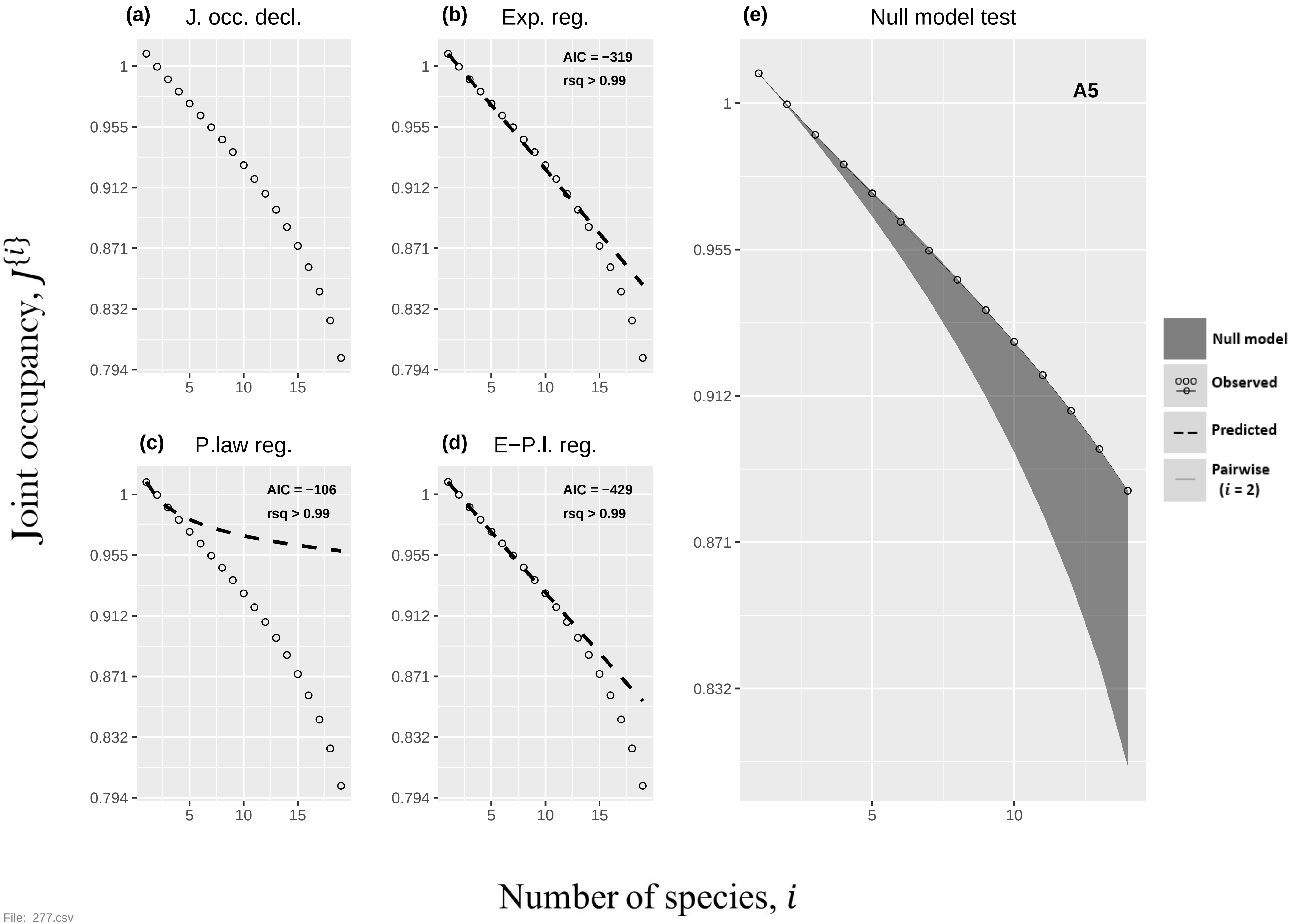
(a) J. occ. decl.
(b) Exp. reg.
AIC = −319
rsq > 0.99
(c) P.law reg.
AIC = −106
rsq > 0.99
(d) E−P.l. reg.
AIC = −429
rsq > 0.99
(e) Null model test
A5
Null model
Observed
Predicted
Pairwise (i = 2)
1
0.955
0.912
0.871
0.832
0.794
5
10
15
Joint occupancy, J{i}
Number of species, i

File: 277.csv

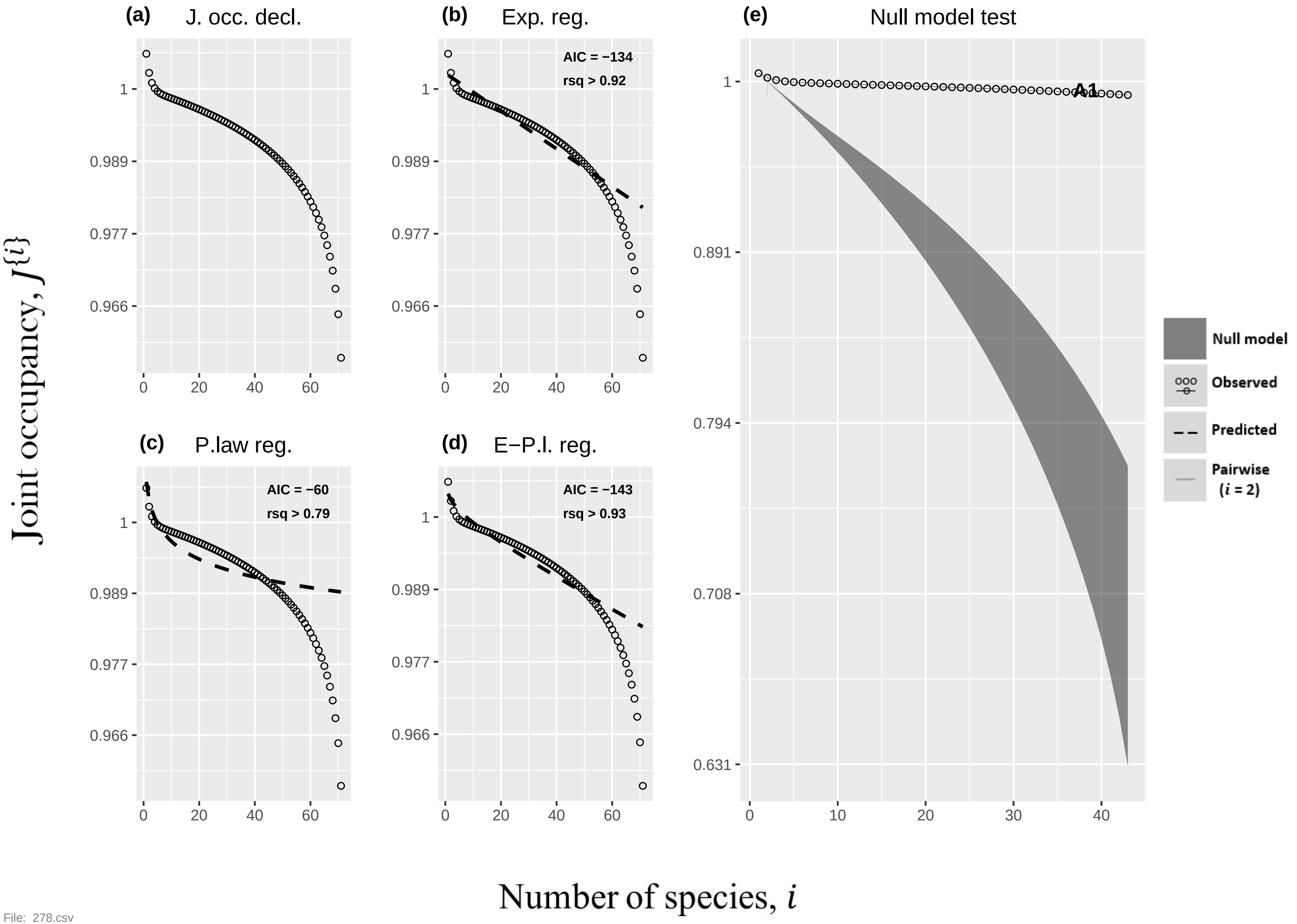
(a)
J. occ. decl.
(b)
Exp. reg.
AIC = −134
rsq > 0.92
(c)
P.law reg.
AIC = −60
rsq > 0.79
(d)
E−P.l. reg.
AIC = −143
rsq > 0.93
(e)
Null model test
A1
Null model
Observed
Predicted
Pairwise
(i = 2)
Joint occupancy, J{i}
Number of species, i
1
0.989
0.977
0.966
0.891
0.794
0.708
0.631
0
10
20
30
40
60

File: 278.csv

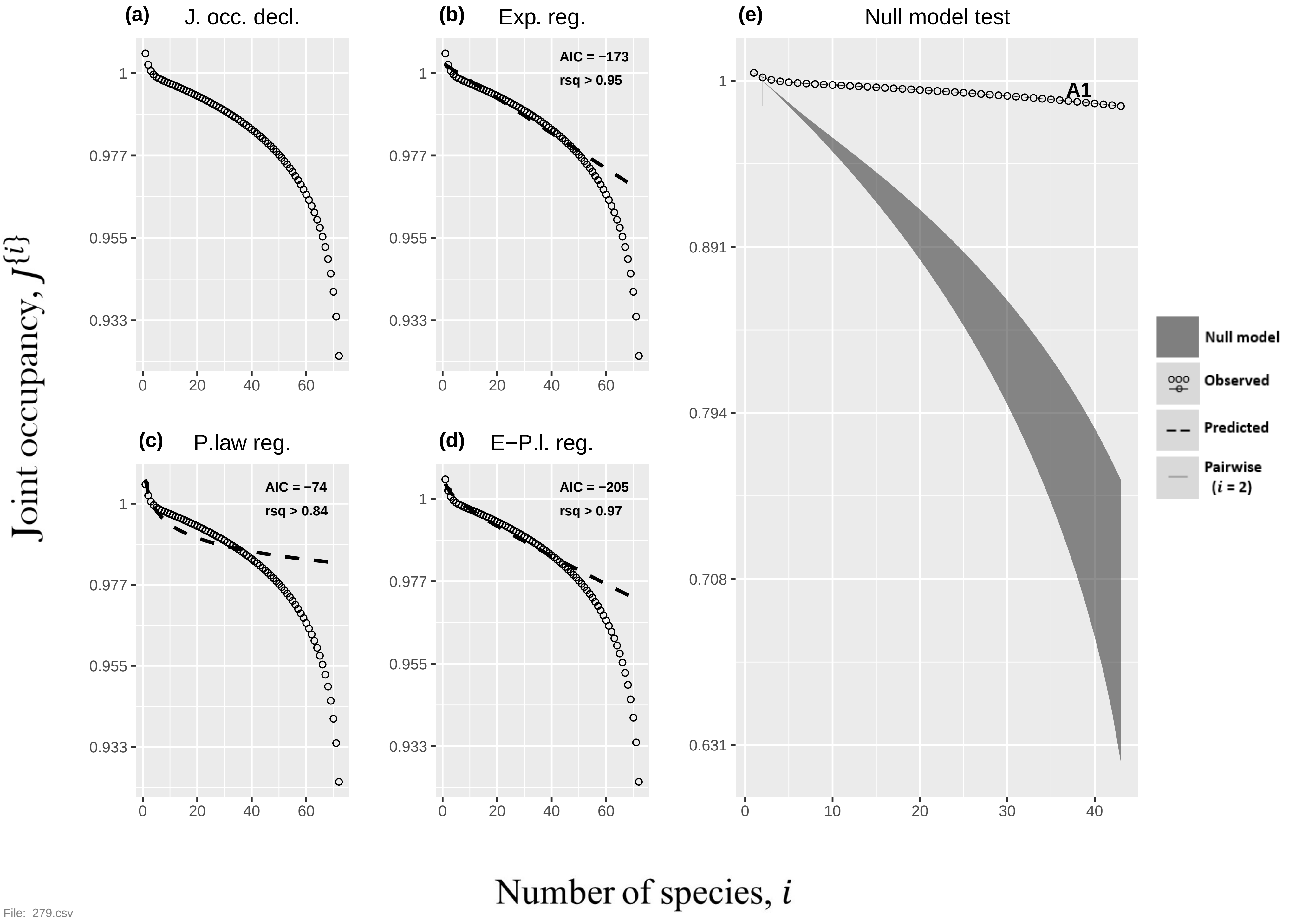
(a) J. occ. decl.
(b) Exp. reg.
AIC = −173
rsq > 0.95
(c) P.law reg.
AIC = −74
rsq > 0.84
(d) E−P.l. reg.
AIC = −205
rsq > 0.97
(e) Null model test
A1
Null model
Observed
Predicted
Pairwise
(i = 2)
Joint occupancy, J{i}
Number of species, i
File: 279.csv

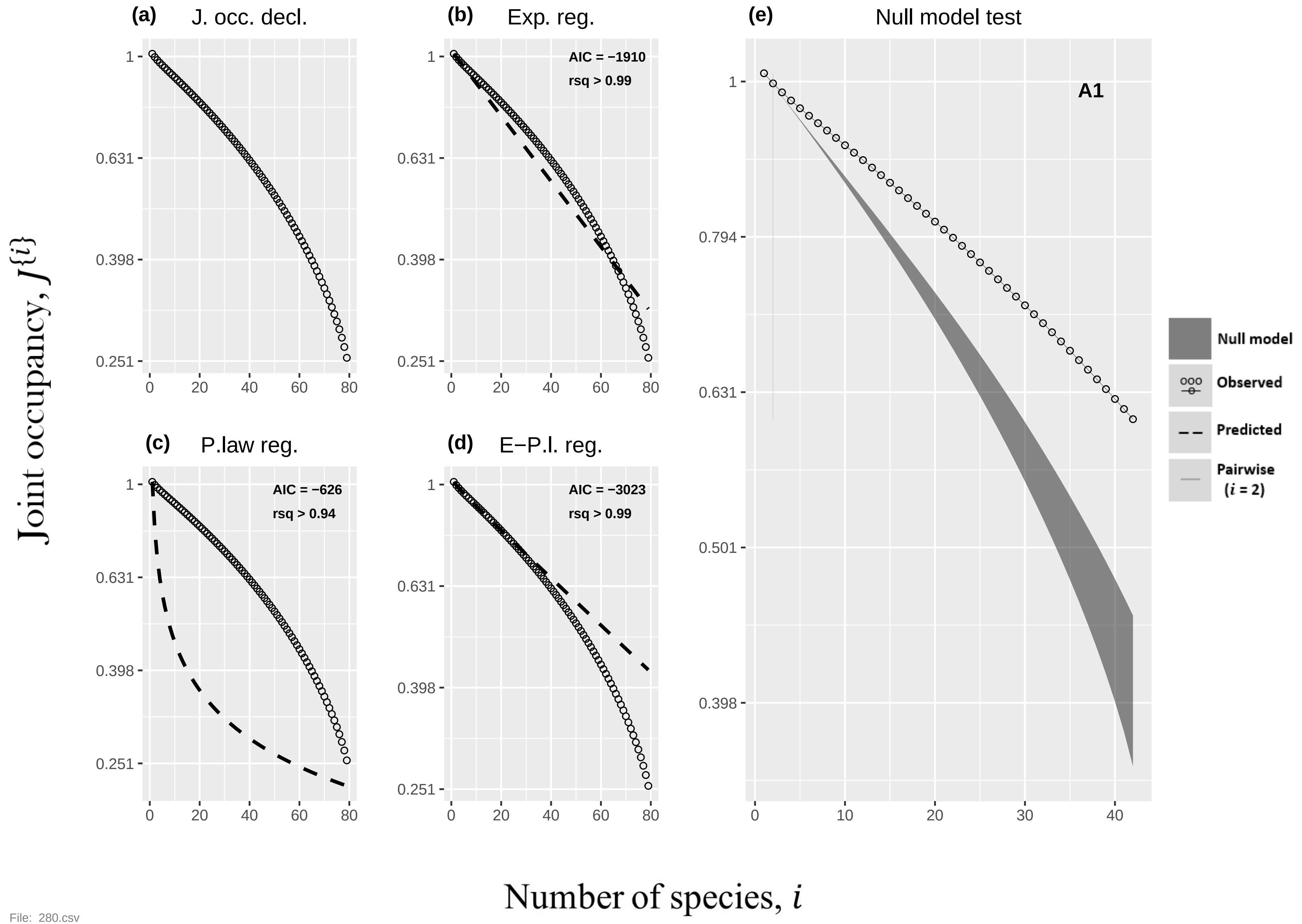
(a) J. occ. decl.
(b) Exp. reg.
AIC = −1910
rsq > 0.99
(c) P.law reg.
AIC = −626
rsq > 0.94
(d) E−P.l. reg.
AIC = −3023
rsq > 0.99
(e) Null model test
A1
Null model
Observed
Predicted
Pairwise (i = 2)
Joint occupancy, J{i}
Number of species, i
1
0.794
0.631
0.501
0.398
0.251
0
10
20
30
40
60
80

File: 280.csv

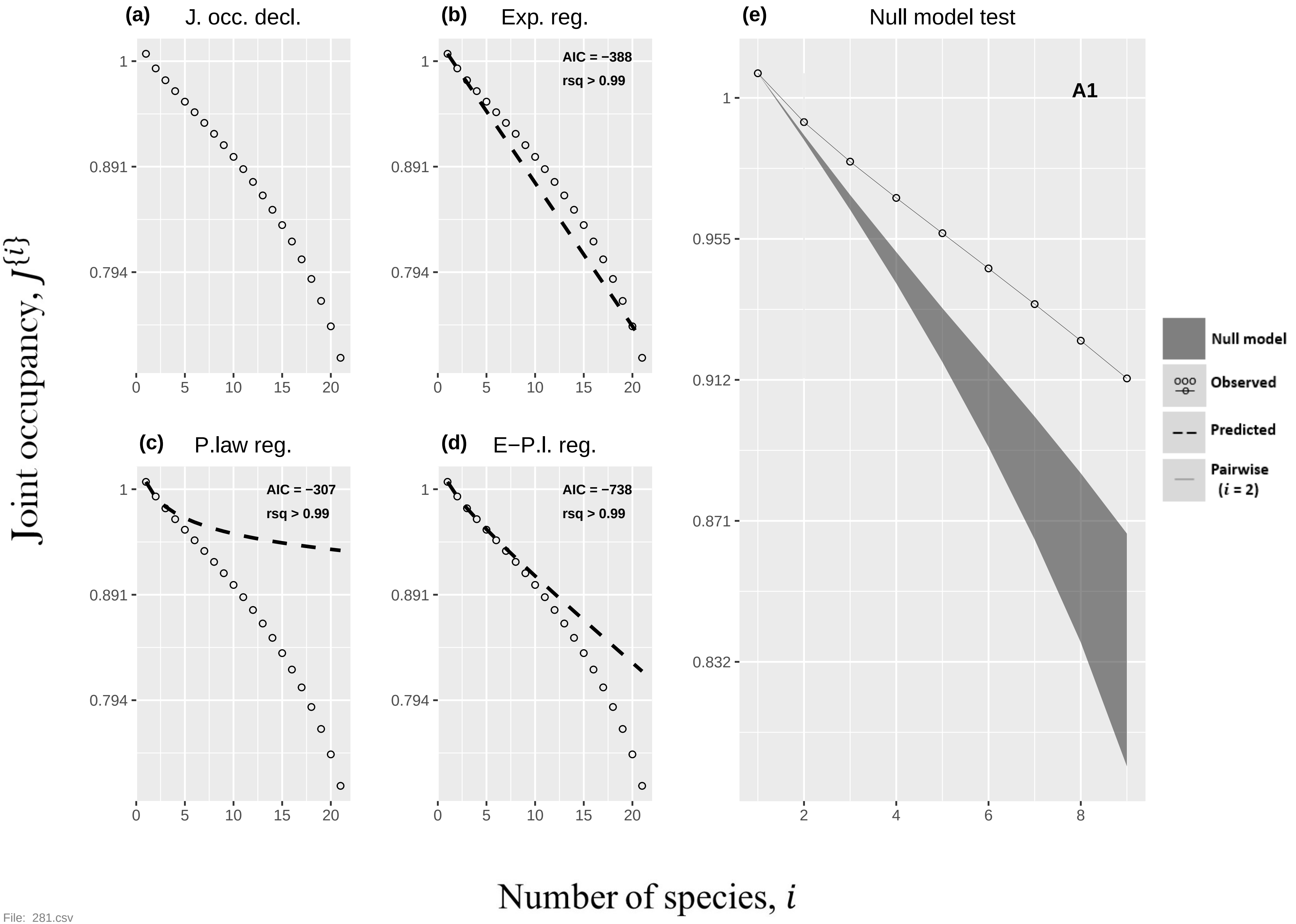

(a) J. occ. decl.
(b) Exp. reg.
AIC = −388
rsq > 0.99
(c) P.law reg.
AIC = −307
rsq > 0.99
(d) E−P.l. reg.
AIC = −738
rsq > 0.99
(e) Null model test
A1
Null model
Observed
Predicted
Pairwise ($i$ = 2)
Joint occupancy, $J^{\{i\}}$
Number of species, $i$


File: 281.csv

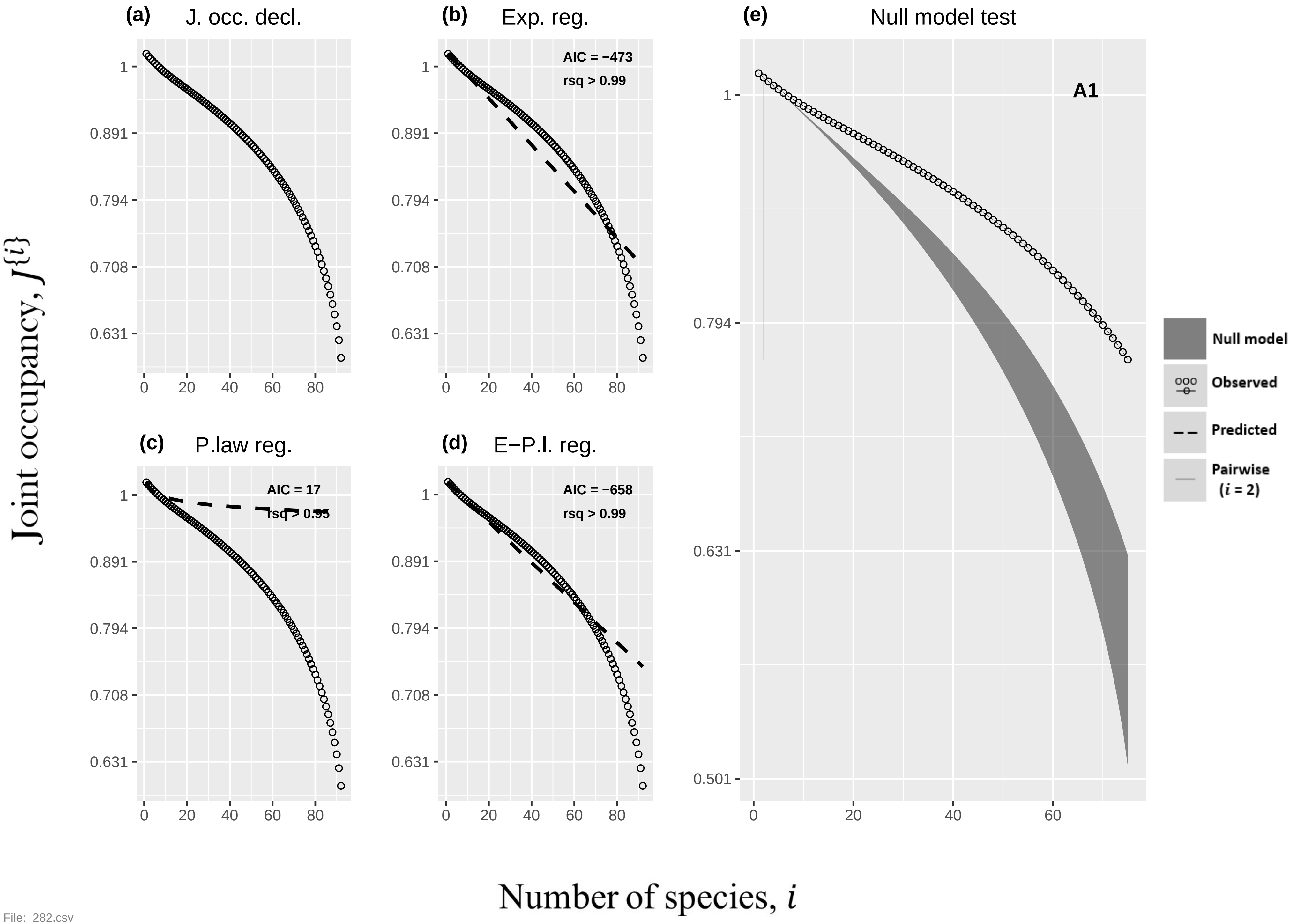

(a) J. occ. decl.
(b) Exp. reg.
AIC = −473
rsq > 0.99
(c) P.law reg.
AIC = 17
rsq > 0.95
(d) E−P.l. reg.
AIC = −658
rsq > 0.99
(e) Null model test
A1
Null model
Observed
Predicted
Pairwise
(i = 2)
1
0.891
0.794
0.708
0.631
0.501
0
20
40
60
80
Joint occupancy, J{i}
Number of species, i


File: 282.csv

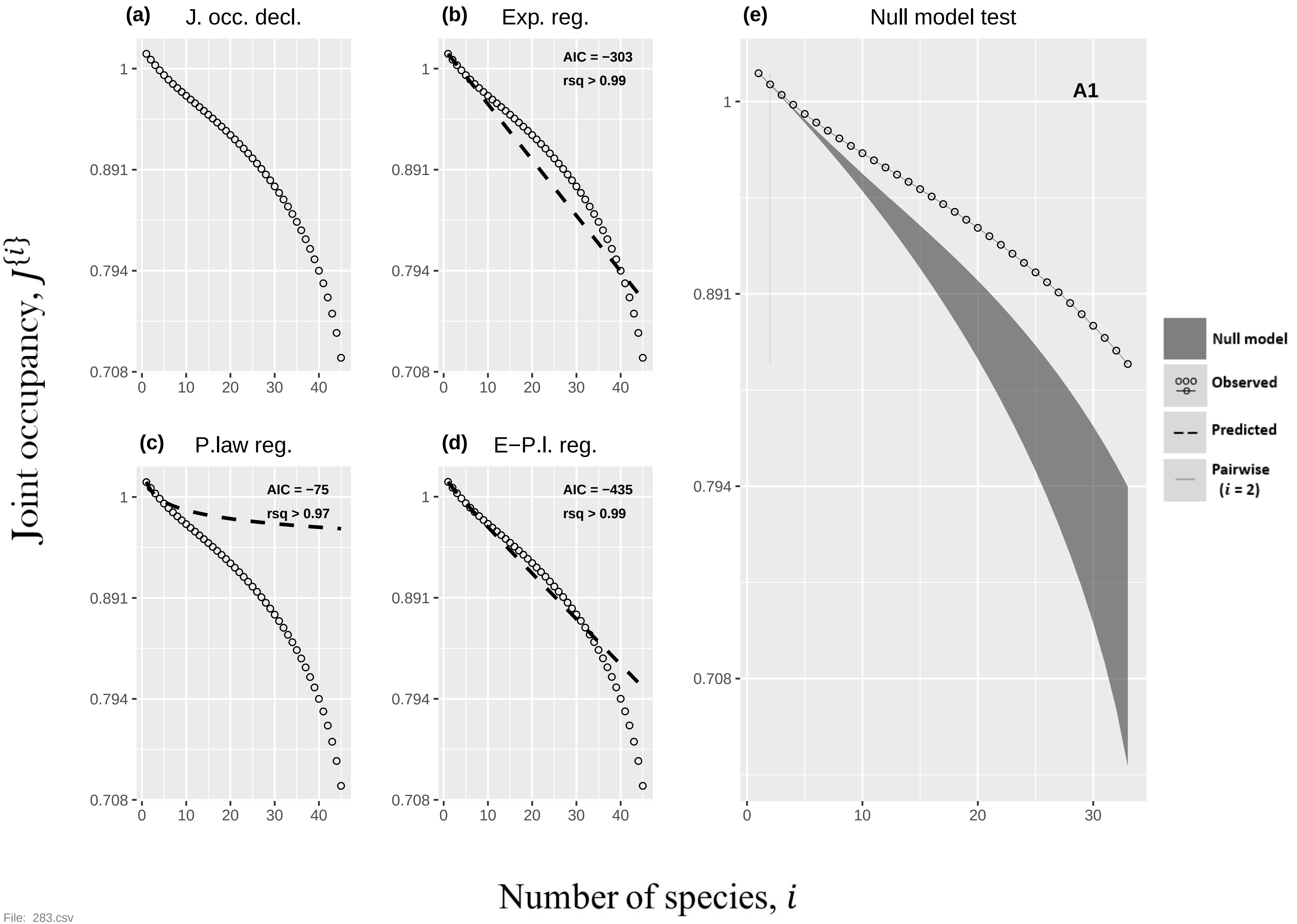

(a) J. occ. decl.
(b) Exp. reg.
AIC = −303
rsq > 0.99
(c) P.law reg.
AIC = −75
rsq > 0.97
(d) E−P.l. reg.
AIC = −435
rsq > 0.99
(e) Null model test
A1
Null model
Observed
Predicted
Pairwise (i = 2)
1
0.891
0.794
0.708
0
10
20
30
40
Joint occupancy, $J^{\{i\}}$
Number of species, $i$


File: 283.csv

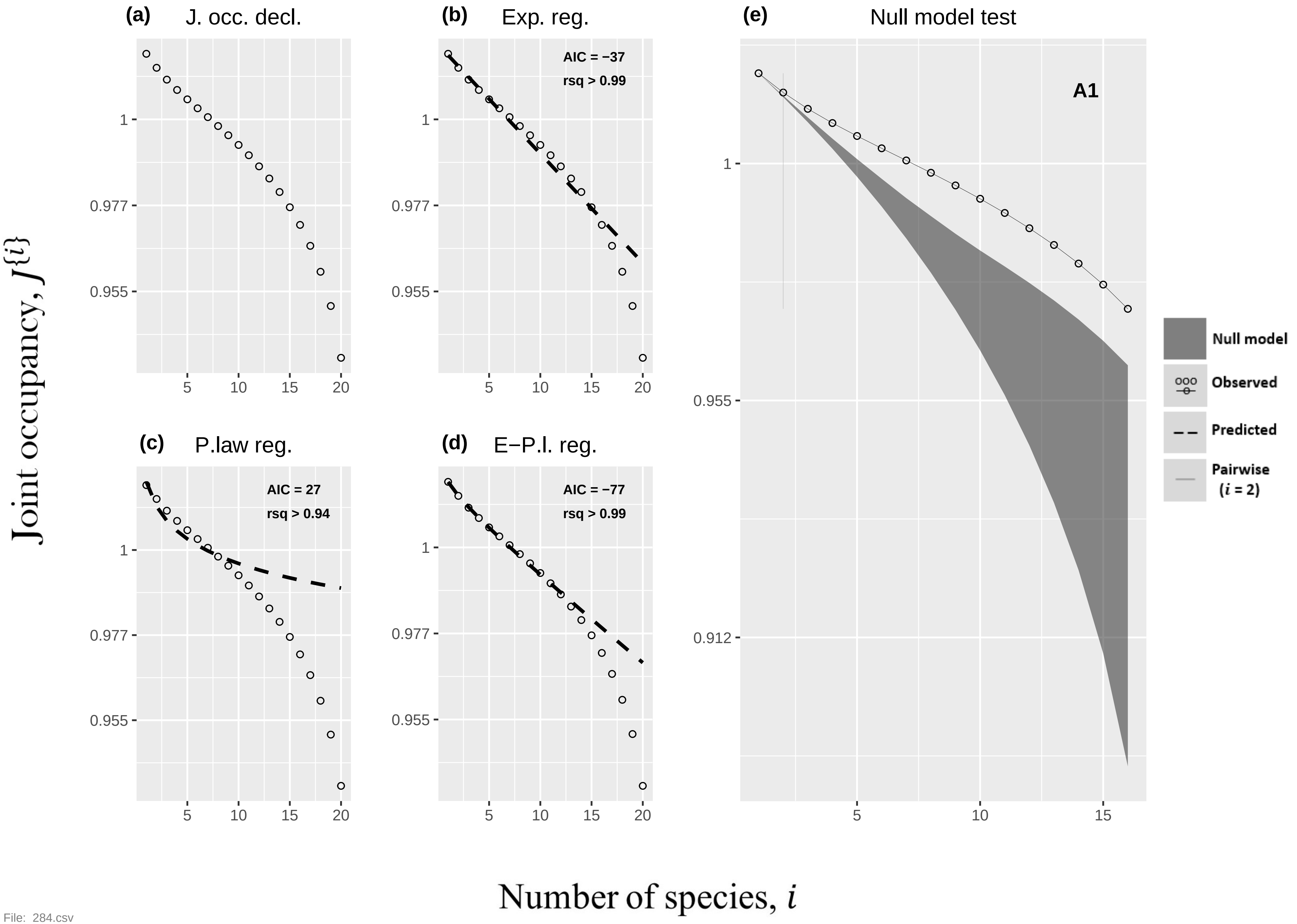

(a) J. occ. decl.
(b) Exp. reg.
AIC = −37
rsq > 0.99
(c) P.law reg.
AIC = 27
rsq > 0.94
(d) E−P.l. reg.
AIC = −77
rsq > 0.99
(e) Null model test
A1
Null model
Observed
Predicted
Pairwise (i = 2)
Joint occupancy, J{i}
Number of species, i
1
0.977
0.955
0.912
5
10
15
20


File: 284.csv

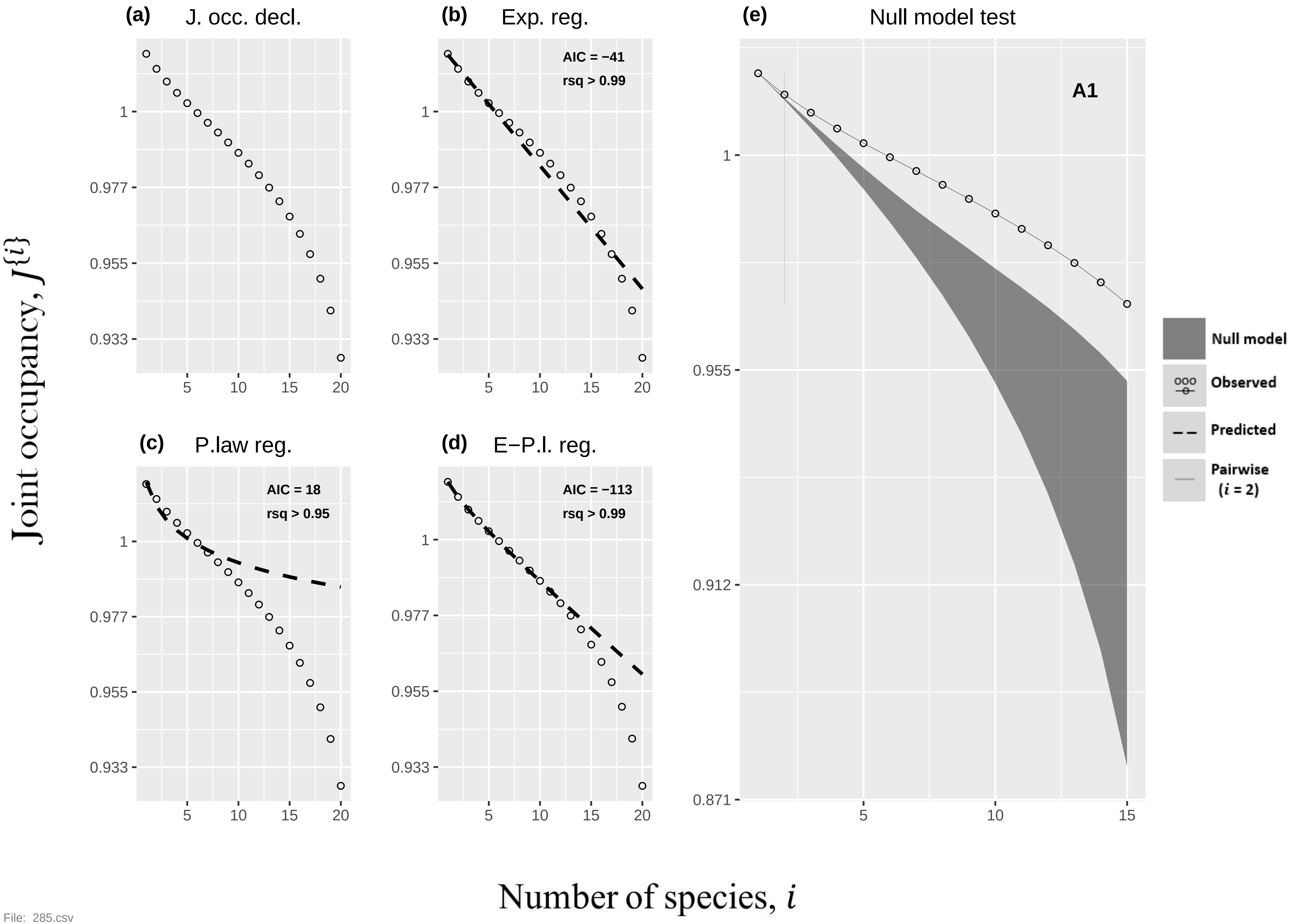

(a) J. occ. decl.
(b) Exp. reg.
AIC = −41
rsq > 0.99
(c) P.law reg.
AIC = 18
rsq > 0.95
(d) E−P.l. reg.
AIC = −113
rsq > 0.99
(e) Null model test
A1
1
0.977
0.955
0.933
0.912
0.871
5
10
15
20
Null model
Observed
Predicted
Pairwise
(i = 2)
Joint occupancy, J{i}
Number of species, i


File: 285.csv

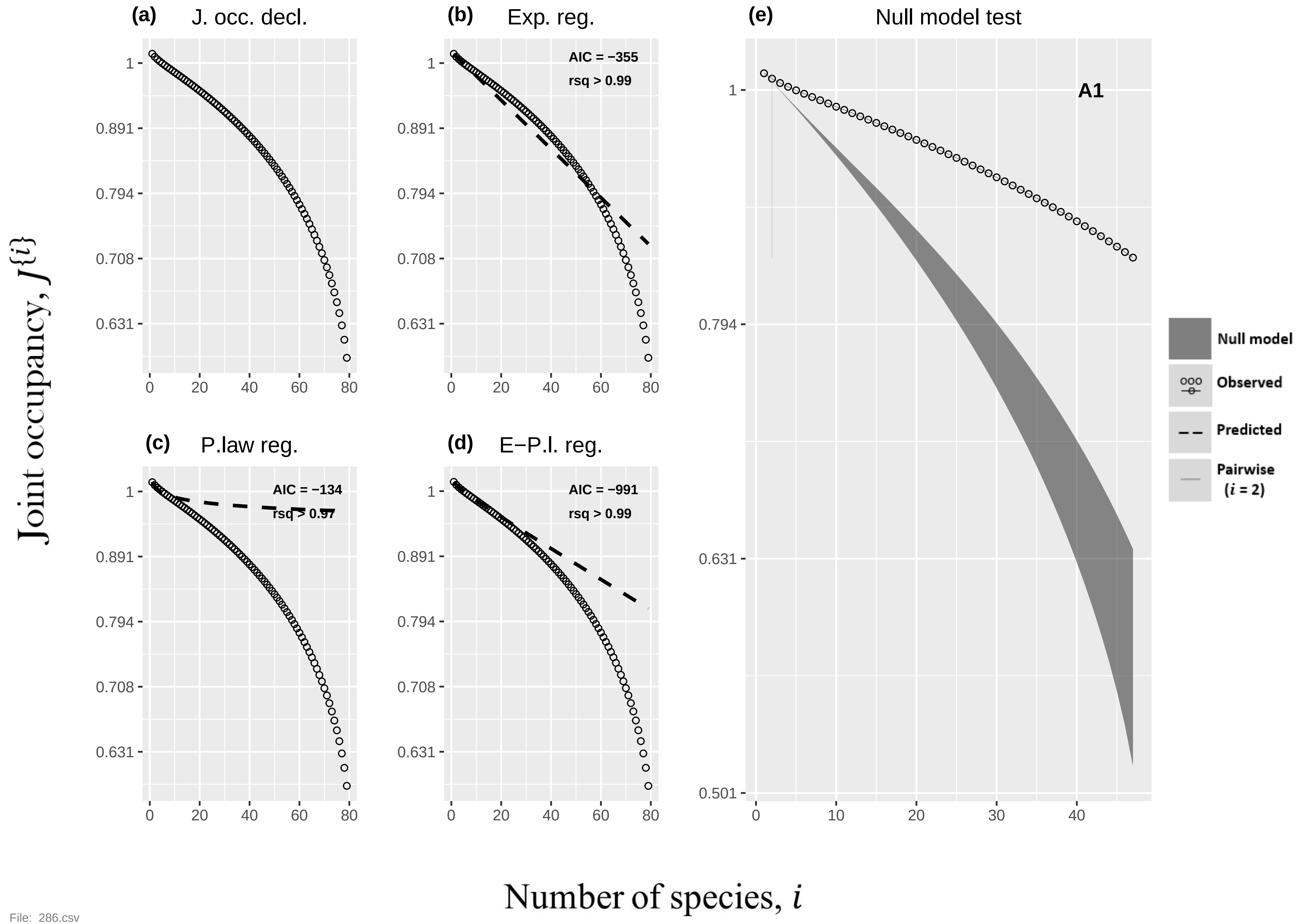

(a)
J. occ. decl.
(b)
Exp. reg.
AIC = −355
rsq > 0.99
(c)
P.law reg.
AIC = −134
rsq > 0.97
(d)
E−P.l. reg.
AIC = −991
rsq > 0.99
(e)
Null model test
A1
Null model
Observed
Predicted
Pairwise
(i = 2)
1
0.891
0.794
0.708
0.631
0.501
0
10
20
30
40
60
80
Joint occupancy, J{i}
Number of species, i


File: 286.csv

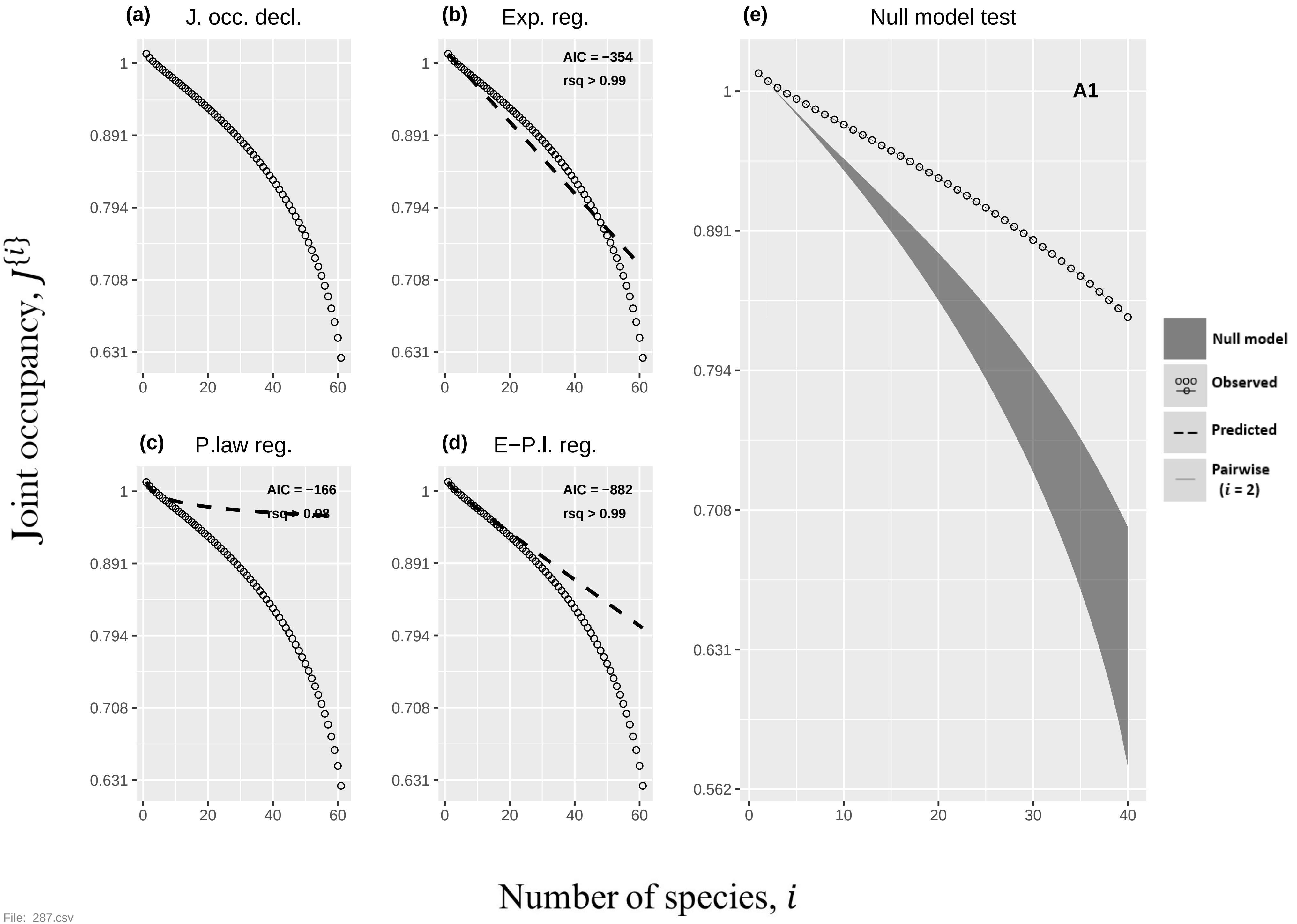

(a)
J. occ. decl.
(b)
Exp. reg.
AIC = −354
rsq > 0.99
(c)
P.law reg.
AIC = −166
rsq > 0.98
(d)
E−P.l. reg.
AIC = −882
rsq > 0.99
(e)
Null model test
A1
Null model
Observed
Predicted
Pairwise (i = 2)
Joint occupancy, J{i}
Number of species, i
1
0.891
0.794
0.708
0.631
0.562
0
10
20
30
40
60

File: 287.csv

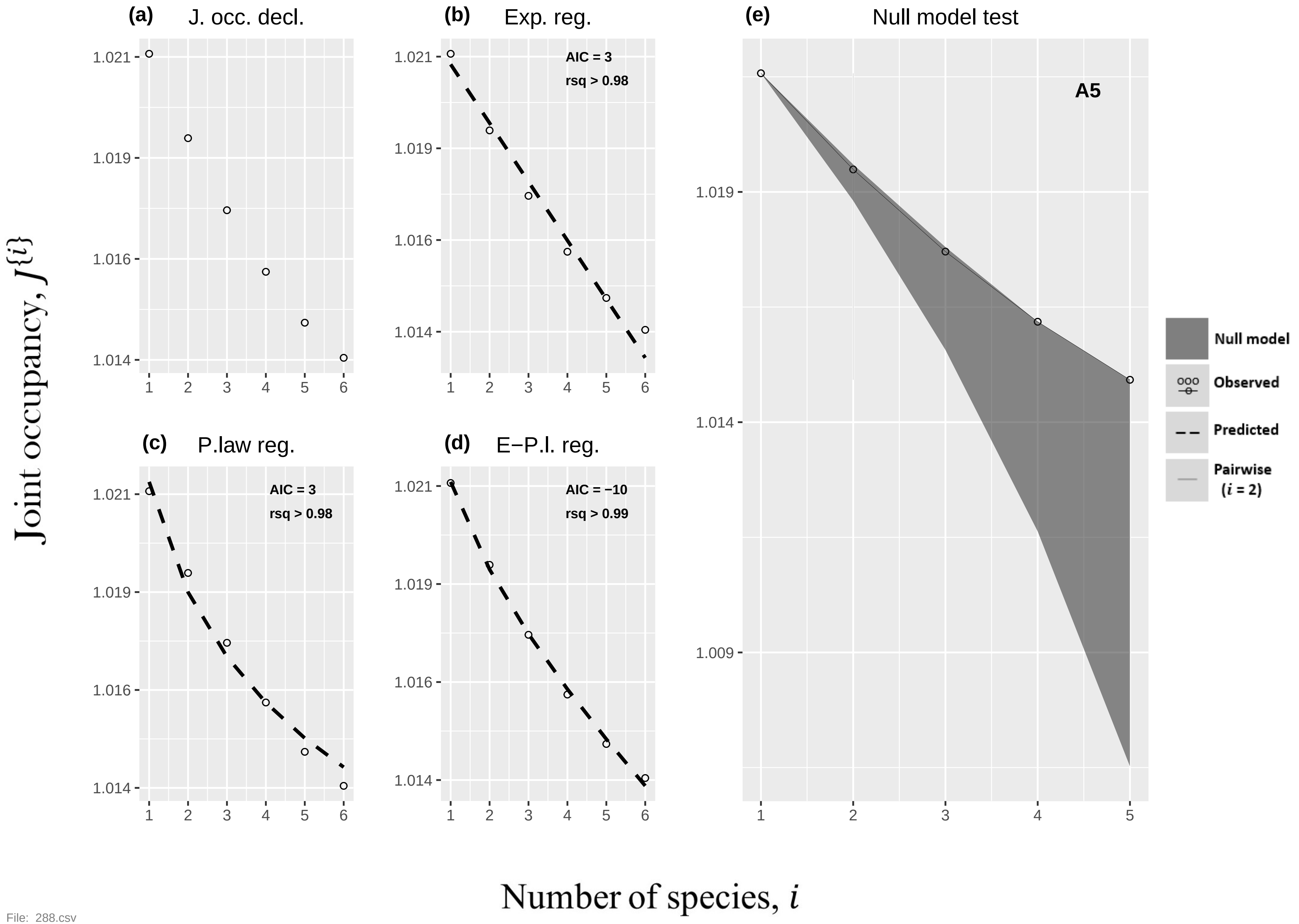

(a) J. occ. decl.
(b) Exp. reg.
AIC = 3
rsq > 0.98
(c) P.law reg.
AIC = 3
rsq > 0.98
(d) E−P.l. reg.
AIC = −10
rsq > 0.99
(e) Null model test
A5
Null model
Observed
Predicted
Pairwise (i = 2)
1.021
1.019
1.016
1.014
1.009
1 2 3 4 5 6
Joint occupancy, J{i}
Number of species, i


File: 288.csv

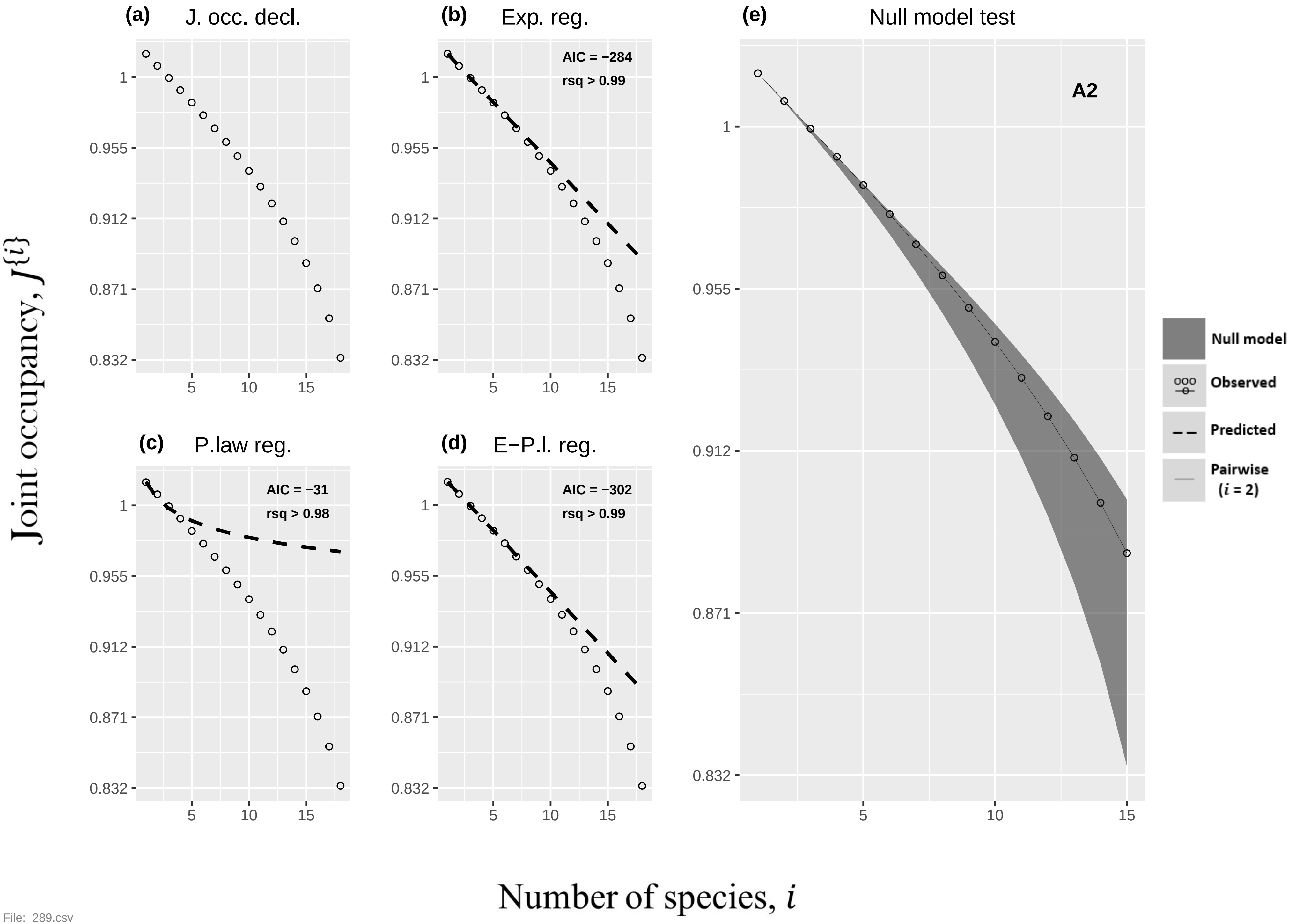
(a)
J. occ. decl.
(b)
Exp. reg.
AIC = −284
rsq > 0.99
(c)
P.law reg.
AIC = −31
rsq > 0.98
(d)
E−P.l. reg.
AIC = −302
rsq > 0.99
(e)
Null model test
A2
Null model
Observed
Predicted
Pairwise (i = 2)
1
0.955
0.912
0.871
0.832
5
10
15
Joint occupancy, J{i}
Number of species, i

File: 289.csv
